# A Pandemic-Scale Ancestral Recombination Graph for SARS-CoV-2

**DOI:** 10.1101/2023.06.08.544212

**Authors:** Shing H. Zhan, Yan Wong, Anastasia Ignatieva, Katherine Eaton, Isobel Guthrie, Benjamin Jeffery, Duncan S. Palmer, Carmen Lia Murall, Sarah P. Otto, Jerome Kelleher

**Affiliations:** Big Data Institute, Li Ka Shing Centre for Health Information and Discovery, University of Oxford, United Kingdom; Infectious Disease Epidemiology Unit (IDEU), Nuffield Department of Population Health, University of Oxford, Oxford, United Kingdom; Department of Statistics, University of Oxford, United Kingdom; National Microbiology Laboratory, Public Health Agency of Canada, Canada; The Pioneer Centre for SMARTbiomed, Big Data Institute, Li Ka Shing Centre for Health Information and Discovery, University of Oxford, United Kingdom; Department of Zoology and Biodiversity Research Centre, University of British Columbia, Vancouver BC Canada

**Keywords:** Ancestral recombination graphs, recombination, SARS-CoV-2, ARG, tskit

## Abstract

Millions of SARS-CoV-2 genome sequences were collected during the COVID-19 pandemic, forming a dataset of unprecedented richness. Estimated genealogies are fundamental to understanding this ocean of data and form the primary input to many downstream analyses. A basic assumption of methods to infer genealogies from viral genetic data is that recombination is negligible and the genealogy is a tree. However, recombinant lineages have risen to global prevalence, and simple tree representations are therefore incomplete and potentially misleading. We present sc2ts, a method to infer reticulate genealogies as an Ancestral Recombination Graph (ARG) in real time at pandemic scale. We infer an ARG for 2.48 million SARS-CoV-2 genomes, which leverages the widely used tskit software ecosystem to support further analyses and visualisation. This rich and validated resource clarifies the relationships among recombinant lineages, quantifies the rate of recombination over time, and provides a lower bound on detectable recombination.

## INTRODUCTION

Classical phylogenetics is largely concerned with inferring the evolutionary history of divergent species^1^. Recombination among lineages, however, distorts the inference of species trees^2^, biases downstream analyses^3,4^, and so requires careful consideration^5^. Modern pathogen datasets are not sparsely sampled from different species and may contain millions of genomes sampled from a single species^6–10^. In contrast to the sophisticated model-based approaches used to infer species trees^11–13^, comprehensive phylogenies of pathogen datasets are typically built with simpler distance methods or approximations^14^, primarily due to computational scaling limitations. While several methods have been proposed to incorporate recombination explicitly into model-based phylogenetic inference^15–19^, scalability is a major issue and practical application has been limited. Software support that explicitly *uses* recombination-aware inference is mostly restricted to visualisation, for example of small-scale phylogenetic networks^20–22^ or tree changes along the genome^23^.

The global surveillance data collected during the COVID-19 pandemic presents fundamental challenges to existing approaches. First, the sheer amount of data overwhelmed even approximate distance matrix-based methods. At the peak in February 2022, ∼80,000 samples per day were submitted to GISAID^6^, accumulating to a total of over 17.5 million samples as of October 2025. Regularly re-inferring trees de novo from such rapidly growing datasets is untenable. The UShER^24^ method is based on adding samples to an existing tree using maximum parsimony, with periodic rearrangements to counter the inherent greediness of the approach^25^. The benefits of this simple model and the value of a comprehensive genealogy updated in real time^26,27^ were conclusively demonstrated, as the resource quickly became an indispensable element of the pandemic response^28^. Recurrent recombination has posed an additional challenge to reconstructing the evolutionary history of SARS-CoV-2^29–34^, with multiple deep recombination events impacting both the genealogy and disease attributes of the virus^35,36^. Although several methods have been proposed to *detect* recombinant samples^37–41^, this information is not incorporated into the phylogenies used in innumerable studies^26,42^, and recombination continues to be treated in a post hoc manner, after phylogenetic reconstruction.

Ancestral recombination graphs (ARGs) describe the reticulate genealogies of sampled sequences in a recombining species^43^, and provide a natural and efficient means of systematically incorporating recombination into the study of SARS-CoV-2. While ARGs have been of theoretical interest in population genetics for several decades^44–46^, recent breakthroughs in simulation^47–50^ and inference^51–57^ have led to a surge of interest in their application in population and statistical genetics^58–60^. As the range of applications for ARGs has broadened, the term has itself evolved from a specific model of coalescence with recombination^44,46^ to a rich data structure representing recombinant ancestry^43,61^. The development of extensive software support has facilitated the broader application of ARGS. In particular, the collaboratively built tskit library^62^ underpins a rapidly growing software ecosystem of simulation^49,63–66^, visualisation^67–69^, inference^53,54,70–72^ and data analysis^73–75^ tools.

Here we present sc2ts, a new method to infer ARGs for SARS-CoV-2 that sequentially adds samples over time in a similar manner to UShER, while automatically detecting and incorporating recombination in real-time. Based on the freely-available Viridian dataset, a curated collection of SARS-CoV-2 genomes purged of various systematic errors and artifacts^76^, we infer an ARG with 2.48 million whole genome sequences. We show that the non-recombinant portions of this ARG are congruent with the UShER phylogeny built from the same dataset as well as the Pango nomenclature system^77^. The ARG also accurately captures evolutionary features such as mutational spectra^78^ and deletions^79^. We show that sc2ts automatically and accurately detects details about known recombinant lineages that were painstakingly collated through a community effort and provides novel insights into the relationships between these recombinant lineages. We compare the recombination events detected by sc2ts with the state-of-the-art combination of UShER and RIPPLES^38^, showing that sc2ts is more sensitive, detecting more of the well-characterised recombination events curated by the Pango lineage designation community. The output of sc2ts—incorporating genealogical relationships, point mutations, deletions, and recombination—provides a basis for the study of SARS-CoV-2 that systematically accounts for all of these evolutionary forces. This ARG, coupled with the rich suite of supporting software in the tskit^62^ and VCF-Zarr^80^ ecosystems, enables pandemic-scale analysis on a laptop. These resources provide a strong platform for future research, helping us to learn from the vast troves of data collected during the pandemic and to prepare for the next.

## RESULTS

### Overview of sc2ts

Sc2ts is a real-time inference method, updated incrementally with daily batches of sequence data (Figure 1). For each batch, sc2ts first infers likely “copying paths” connecting each sample to nodes in the current ARG using a simplified version of the Li and Stephens (LS) model^81^. The LS model is a Hidden Markov Model (HMM) widely used in human genomics to approximate the effects of mutation and recombination on genetic inheritance^82,83^, typically parametrised by population-scaled mutation and recombination rates varying along the genome^84^. In the interest of interpretability, we simplified this model to use a single parameter *k* that controls the number of recurrent mutations that is allowed before switching to a different parent via recombination (we use *k* = 4 here; see STAR Methods). We used the highly efficient LS HMM implementation in tsinfer^53^, with some additional optimisations enabled by our simplified model, to find exact Viterbi solutions of these copying paths for each sample given the current ARG. Note that this is a form of parsimony, with the recombination penalty *k* defining the relative parsimony of recombinant versus non-recombinant solutions to the HMM. The per-sample copying paths produced by the HMM (most of which do not involve recombination) form clusters of samples within a batch, which are then resolved using standard phylogenetic techniques. Finally, sc2ts attaches the phylogenies of the clusters to the current ARG, and applies parsimony-improving heuristics to address issues introduced by the inherent greediness of this strategy. Sc2ts is implemented in Python, building on core PyData infrastructure^85–88^ and bioinformatics packages^89–91^. See STAR Methods for details on all aspects of the method.

**Figure 1:**
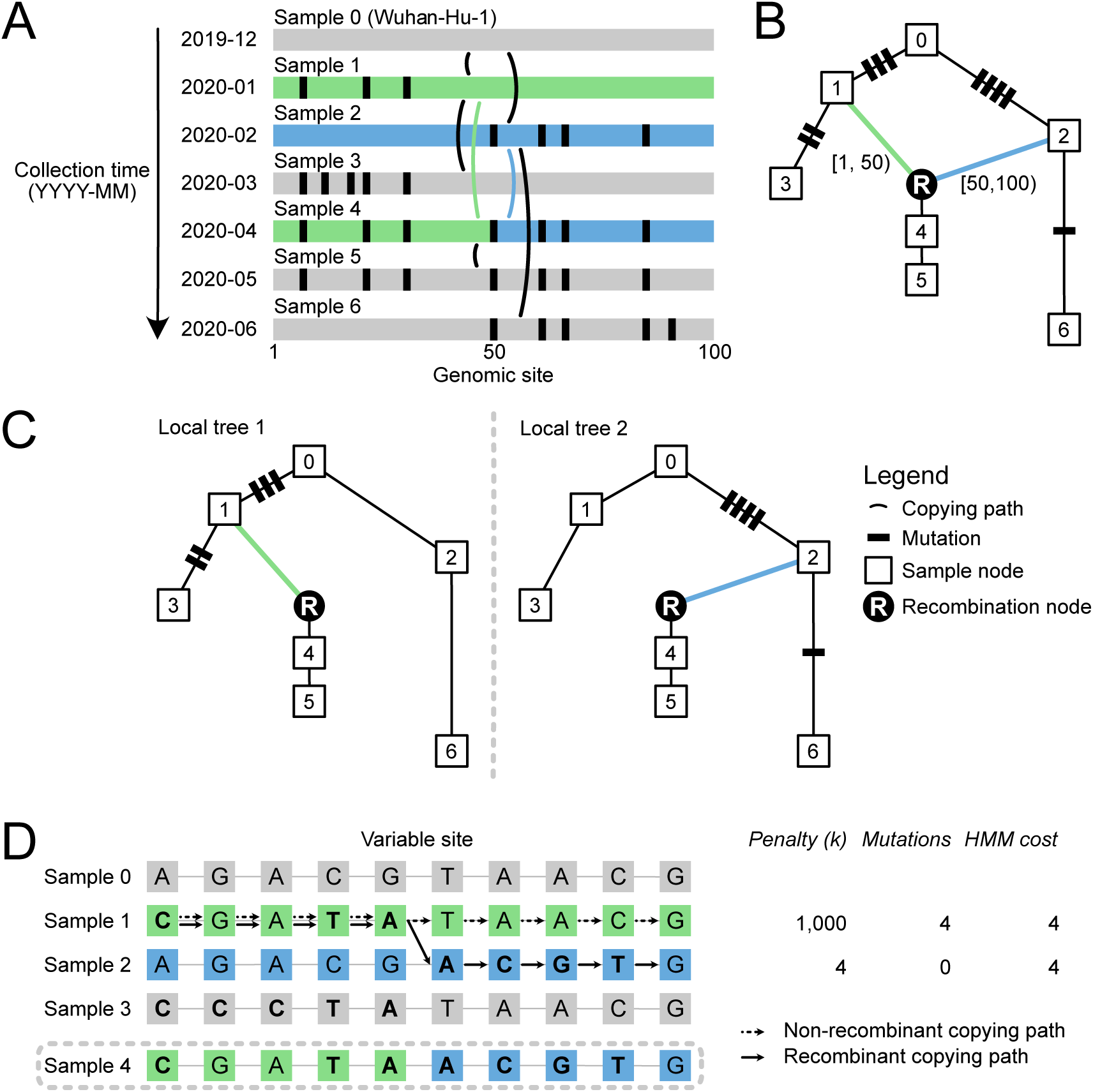
Schematic overview of sc2ts. (A) Process to build an ARG by incrementally updating it with samples sorted by collection dates. (B) The ARG resulting from the inference process. Sample 0 is used to initialize the ARG as the root. Next, Sample 1 is attached, mapping mutations onto the new branch to explain it. When Sample 4 is detected as a recombinant, a node representing the recombination event is inserted with two edges connecting to its parents, with Sample 4 itself attached to this node as its child. (C) Two local trees contained in the resulting ARG, one on each side of the inferred breakpoint which occurs at position 50. The genomic coordinates of the segments inherited from different recombinant parents are denoted by two half-open intervals. (D) Copying paths for Sample 4 inferred under the LS model while allowing recombination (*k* = 4) or disallowing it (*k* = 1000).

### A comprehensive representation of SARS-CoV-2 evolution

Using sc2ts, we built an ARG from 2,482,157 samples from the Viridian v04 dataset^76^, collected between 2020-01-01 and 2023-02-20. Inference took 21 days on a server with 64 cores (2× AMD EPYC 7502) and 512 GB of RAM (∼5 CPU years; Figures S1 S2). After post-processing, the ARG contains 2,689,054 nodes, 2,748,838 edges, and 2,285,344 mutations at 29,893 sites, and is stored in a 32 MiB file. Of the nodes, 855 represent recombination events, resulting in 316 distinct trees along the genome (some events occurred at the same breakpoints). Using the tskit Python API, the ARG can be loaded into memory in half a second, and the vectorised representation integrates tightly with powerful data science tools, such as NumPy^85^. For example, a tabular summary for the 2.75 million nodes, including sample IDs and Pango assignments (stored as metadata in the ARG), can be derived as a Pandas data frame^87^ in less than half a second. The ARG can therefore be easily accessed using familiar tools with minimal hardware requirements, enabling diverse bespoke analyses.

The ARG is a comprehensive and deeply validated record of the pandemic phase of SARS-CoV-2 evolutionary history, accurately reflecting a range of results from the literature. First, we compared the sc2ts ARG with the state-of-the-art UShER tree inferred from the same dataset by pruning both to their intersection of 2,475,418 samples and 27,507 sites (STAR Methods). Eliminating recombination nodes and their descendants (which we expected to differ), we found very similar relationships among lineages in the ARG and UShER tree based on tanglegrams for a set of representative samples from 814 well-sampled lineages (STAR Methods; Figures S3, S4, S5, S6). Overall, we also found close similarity between the ARG and UShER tree in terms of parsimony (Document S1.1) and imputation of missing and ambiguous bases (Document S1.2). We compared the mutational spectra of major Variants of Concern (VOCs) in the ARG against the UShER tree and a previous study^78^ (STAR Methods), finding close agreement (Figure S7). We also evaluated how well the ARG reflects the structure of the Pango nomenclature system by computing Pango assignments for all 2.75 million nodes in the ARG using Pangolin^92^ (STAR Methods). Only a small minority of the exported sample alignments (21,680; 0.87%) disagreed with the source consensus sequences in terms of Pango lineage assignment (comparable to the 0.27% disagreement between pangolin-data versions 1.21 and 1.29 in the source Viridian metadata). A majority of the Pango lineages (1473 of 2058) have monophyletic origins in the ARG, increasing to 1779 (86%) when we allow for multiple sibling origination nodes (Document S1.3). The ARG is time-resolved, with internal node dates estimated by a non-parametric method^93^ (STAR Methods). We show that the dates agree well with Nextstrain estimates (Figure S8) and overall accuracy is comparable to Chronumental^94^ applied to an UShER tree (Figure S9, Document S1.4). Finally, we verified that the origins of the Alpha (Figure S10, Document S1.5.1), Delta (Figure S11, Document S1.5.2) and Omicron (Figure S12, Document S1.5.3) VOCs reflect expectations from the literature^95–101^.

Deletions are an important part of SARS-CoV-2 evolution^102^ but are not incorporated into UShER trees and must be accounted for separately. Because deletions are difficult to distinguish from bioinformatic errors and their multi-base nature presents challenges to the LS HMM, we masked out all gap characters as missing data during primary inference. To approximate the effects of the most important deletions into the ARG, we remapped data using post hoc parsimony at 75 sites involved in deletions with a frequency of ≥ 1% reported by Li et al.^79^ during post-processing (STAR Methods). Although sites are treated independently in this approach, runs of mutations to the gap character at adjacent sites correctly identified many known high-frequency (Table S1) and recurrent (Table S2) deletions. For example, the three defining deletions of Alpha^95^ (11288:9, 21765:6, 21991:3) are correctly mapped above the Alpha origin in the ARG (Figure S10). The ARG captures known recurrent deletions in the N-terminal domain of the S1 unit of Spike (which is a deletion hotspot^103^), for example, the 21765:6 deletion (ΔH69/V70), which causes S-Gene Target Failure^104^ and is associated with increased efficiency in cell entry^105^, and the 21991:3 deletion which is associated with antibody escape^103^. See Document S1.6 for further analysis.

### Epidemiologically relevant recombinant lineages

Part of the success of the Pango nomenclature system^77^ can be attributed to the clarity of the rules under which new lineages are designated and their explicit phylogenetic basis. The first recombinant lineage XA was designated in 2021 based on the analysis of Jackson et al.^29^. Although the process of incorporating recombinant lineages in the Pango nomenclature system is well defined, the identification of such events is much more subjective, requiring the synthesis of multiple lines of evidence and often the manual inspection of phylogenies and sequence alignments by volunteers^106^. As of pango-designation version v1.33.1 (2025-03-31), a total of 146 distinct Pango X lineages have been designated.

By automatically detecting and incorporating recombination events into the ARG, sc2ts provides us with a systematic means of discovering and assessing evidence around these events. Table 1 summarises these results for the 44 Pango X lineages present in the ARG (see Document S1.7). For 16 Pango X lineages, the oldest node that has been assigned to the focal Pango lineage is a recombination node and that node is the ancestor of all samples assigned to that (or a descendant) Pango lineage. We consider these to be fully concordant (Type I) events, where the ARG reflects the hierarchical structure of Pango designation. Among these events, concordance with the community-consensus parent lineages and breakpoint intervals is excellent, with similar performance to the RecombinHunt^40^ and CovRecomb^41^ detection methods (STAR Methods; Table S3; see also Document S1.8 for analysis of the Jackson et al. recombinants). Of the Type I events, 13 are monophyletic for the focal Pango lineage (i.e., the descendants are entirely from that lineage); Figure 2A shows XA as an example.

**Figure 2:**
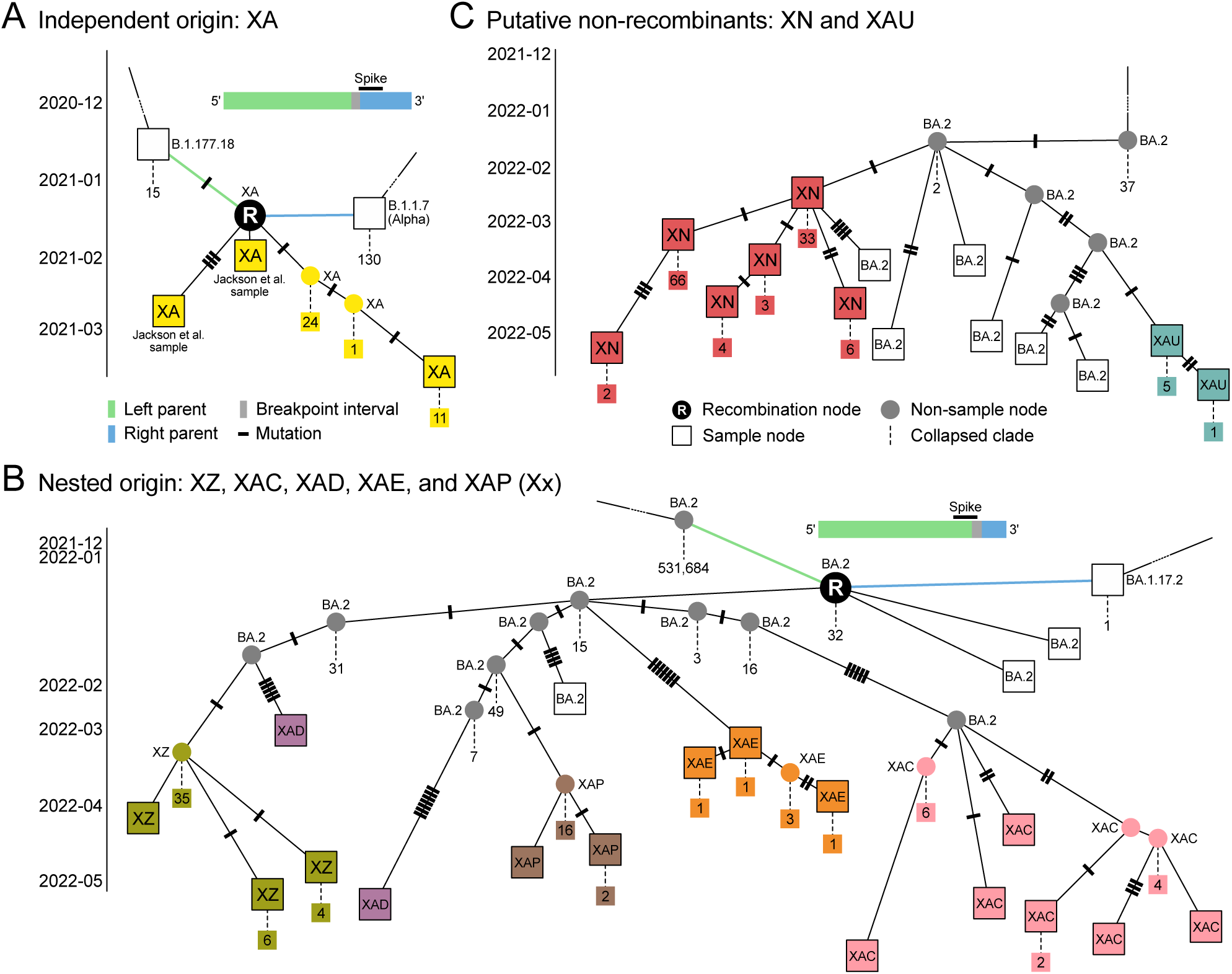
Illustrative subgraphs in the ARG. Boxes identify specific samples, coloured by their Pango X classification; smaller squares list extra samples descending from a node, hidden for brevity, with all hidden samples taking the same Pango designation as their parent. For the two recombination nodes, a genome bar above shows the location of the inferred breakpoint interval relative to the Spike gene. For simplicity, ancestral lineages above the recombinant parents or the MRCA are omitted. (A) Subgraph around the recombination node associated with XA. Two “Group A” recombinant samples reported by Jackson et al.^29^ are shown. (B) Subgraph around the recombination node associated with a group of “nested” Pango X lineages: we use the label Xx as a shorthand for this related group of recombinants; and (C) Subgraph around the MRCA of the closely related non-recombinants XN and XAU. For more detailed visualisations of these subgraphs, see Document S2.

**Table 1:**
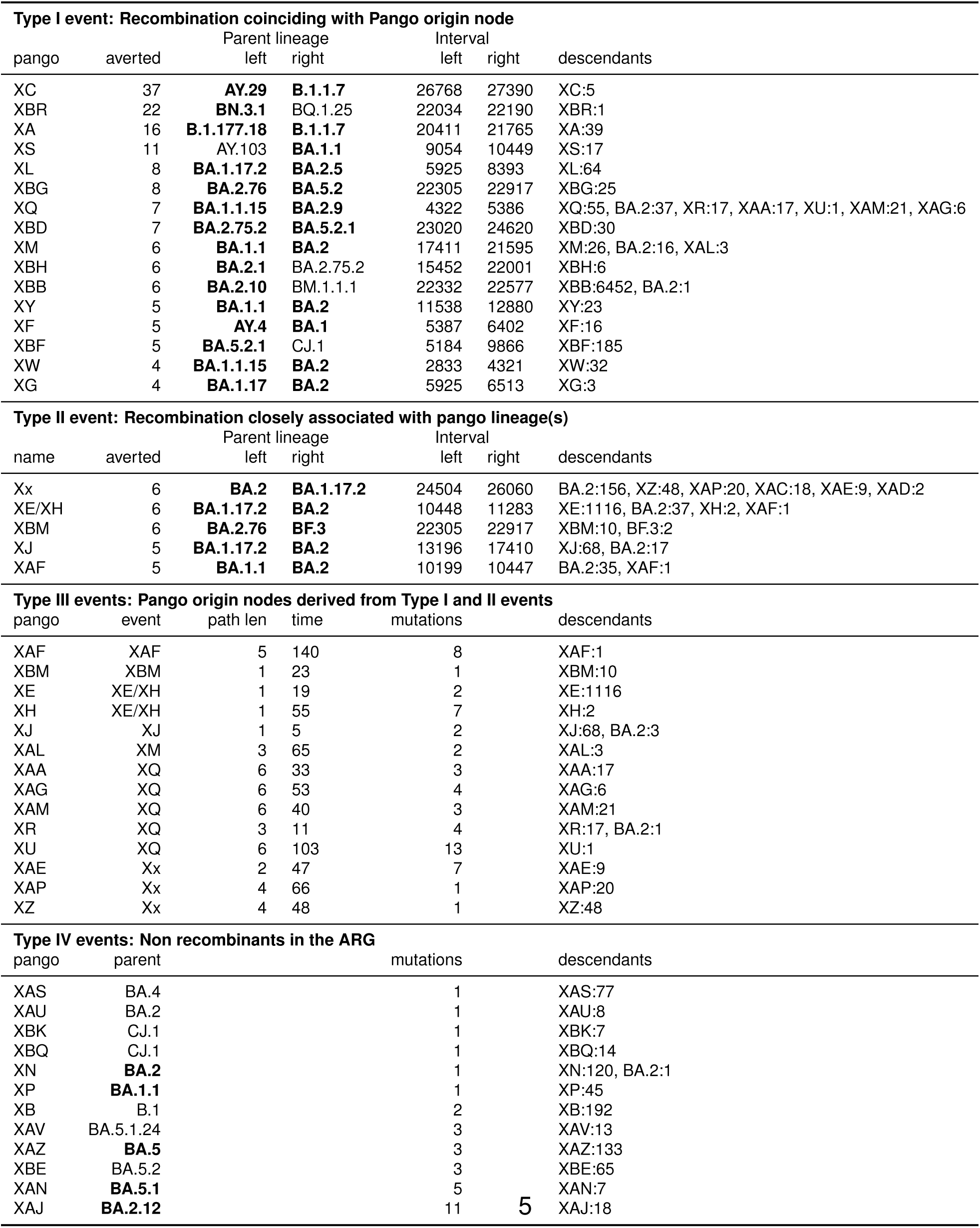
Events in the ARG associated with Pango X lineages. Each row corresponds to the origin node for a given Pango lineage (Types I, II, and IV, see text) or associated recombination event (Type II). For origin nodes that are recombinant (Type I) or near a recombination event (Type II), the “averted” column shows the number of additional mutations required without recombination, as well as the position of informative sites to the left and right of the breakpoint. Parent lineages shown in boldface are nodes exactly matched by sampled sequences. The descendants column shows the number of descendants by Pango lineage. For Pango lineages (Type III) whose origin node was a descendant of a Type I or II recombination node, the distance (path length and time in days) from the recombination event is shown, as well as the number of mutations directly ancestral to the Pango origin node. Pango X lineages not associated with a recombination node (Type IV) are shown with the parent node and number of mutations.

There are five partially concordant events (Type II) in Table 1, which are recombination events closely associated with Pango X lineages that do not meet the strict origination criteria for Type I. Type III events are Pango origin nodes in the ARG that are derived from Type I or Type II recombination events. There is significant nesting among the Pango X lineage origin nodes within the ARG, with two major clusters of particular note. Although XQ is directly associated with a recombination event in the ARG, XAA, XAG, XAM, XR and XU are all closely derived, suggesting that they may be more parsimoniously explained as sublineages of XQ than de novo recombinant lineages (Document S1.9.12). The second cluster is based around a putative undocumented recombinant that is the ancestor of lineages XZ, XAC, XAD, XAE, and XAP, which we refer to as “Xx” (Figure 2B, Document S1.9.16).

The final class of events (Type IV) in Table 1 corresponds to nodes that are not associated with a recombination event in the ARG. These vary in their plausibility of being non-recombinant. For example, XN (Figure 2C) is only one mutation distant from a BA.2 sample and has little additional evidence to suggest recombination (Document S1.9.10). XB, on the other hand, has been thoroughly characterised^32^ and is likely misclassified by sc2ts because the required parent lineages were not present (Document S1.9.2).

The remaining 10 Pango X lineages (97 samples) present in the dataset are not included in the ARG because the HMM cost exceeds the inclusion threshold and there was insufficient support for retrospective inclusion (STAR Methods). These include “complex recombinants” such as XBC and XAY^35^ which are confirmed by sc2ts to have multiple breakpoints; see Document S1.10 and Table S6 for details. See Document S1.9 for extended analyses on all the Pango X lineages in the ARG; Figure S13 for copying patterns associated with Table 1; and Document S2 for ARG visualisations of the Pango X lineages.

### Quality control and validation of recombination events

Sequencing errors in genome assemblies^107^, incorrect read alignments^76^, and multi-nucleotide substitutions^108^ can cause short regions of a sample genome to match more closely to an alternative genome rather than the true parent. Such genomic regions can be mistaken for evidence of recombination. We use a simple quality control (QC) measure that counts the number of “loci” (clusters of closely spaced sites) supporting the left and right parents of a recombinant (STAR Methods), and effectively identifies a range of issues. Figure S16 shows the net number of supporting loci for the left and right parents for all 855 recombination events. The 501 recombinants having fewer than 4 net loci supporting each parent are highly enriched for samples sequenced using the AmpliSeq V1 protocol (70% of the recombinants despite making up 2% of the Viridian dataset), and 71 (14%) are associated with 6-base deletion in the Delta lineage, known to be affected by amplicon dropout^109^. The 354 remaining recombination events that pass this QC filter contain all of the recombinants associated with Pango X lineages (Table 1) and the Jackson et al. recombinants (Document S1.8).

We validated the recombination events in the sc2ts ARG by running 3SEQ^110^ on the putative recombinant sequence trios (STAR Methods). Of the 855 recombination events identified by sc2ts, 3SEQ verified 57% of them, rising to 95% verification among the 354 recombinants that passed our QC filter. In addition to 3SEQ, we ran the CovRecomb^41^ and rebar (https://github.com/phac-nml/rebar) recombination detection methods on the putative recombinant sequences, summarised in Table S4 (see also Figures S14, and S15). CovRecomb has a substantially lower validation rate at 25% among the QC-passing recombinants. This is expected, as CovRecomb is specifically intended to detect inter-lineage recombinants and has previously been shown to be less sensitive than other methods^41^. Rebar validation is intermediate, at 57% among QC-passing recombinants.

### Assessing evidence for recombination events

The LS HMM at the heart of sc2ts provides a simple and powerful means of quantifying the plausibility of a given recombination event. To do this, we force the HMM to find a Viterbi solution for the putative recombinant sequence with no recombination by specifying a large recombination penalty *k* (Figure 1). The difference between the numbers of mutations between the two solutions—the number of mutations “averted” by recombination—is a measure of the evidence in favour of recombination (similar to the “parsimony improvement” score in RIPPLES^38^). Table 1 shows the number of averted mutations for each of the 21 recombination events associated with Pango X lineages, with support ranging from very strong in the case of XC (the most parsimonious non-recombinant placement would require 37 additional mutations) to marginal for XG and XW, which have the minimal possible number of averted mutations.

Figure 3A plots the number of mutations required in the recombinant versus non-recombinant Viterbi solutions (upper scatter plot) and the resulting distribution of the number of averted mutations (lower histogram) over all 354 recombination events that pass QC. Also shown is the 3SEQ classification status of these recombinants. The 5% of recombinants that do not pass 3SEQ validation are among the most weakly supported, with 4 or 5 averted mutations. Figures 3B,C show the distribution of recombination events as a function of the divergence between their two parents. Figures S14 and S15 plot the same information as Figure 3 broken down by CovRecomb and rebar validation status.

**Figure 3:**
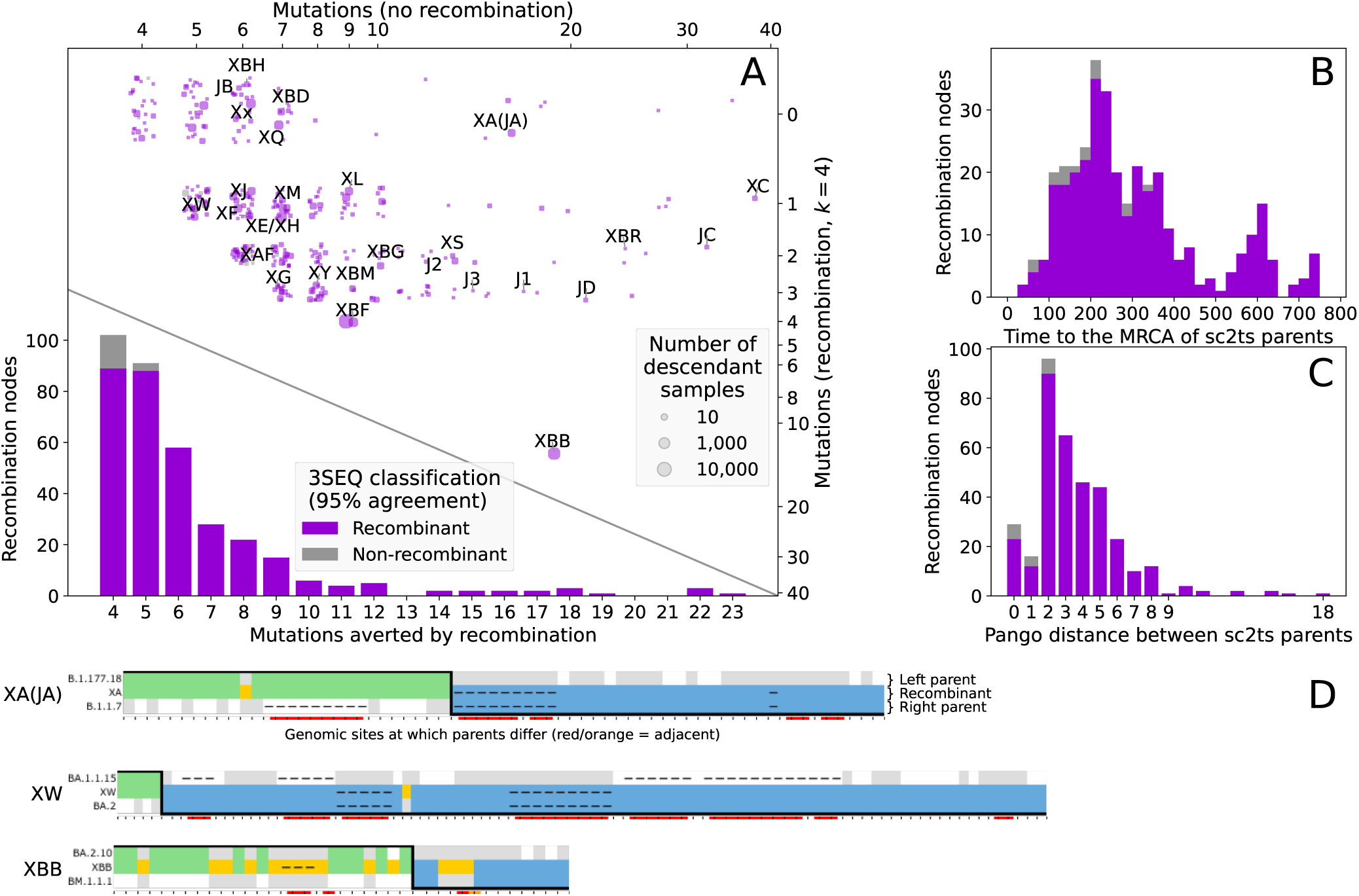
Properties of recombination events. A,B,C show summaries of the 354 sc2ts recombination nodes passing QC filters coloured by 3SEQ classification (recombinant: purple, non-recombinant: grey). (A) Upper: jittered scatterplot comparing number of mutations when recombination allowed (*y*: right axis) versus an alternative copying path disallowing recombination (*x*: top axis); recombination nodes in Table 1 are labelled starting with ‘X’ and those identified by Jackson et al.^29^ are prefixed with a ‘J’, e.g., Jackson group B (JB) and Jackson singleton recombinant 1 (J1). Point sizes reflect number of descendants. Lower: recombination nodes classified by number of mutations averted by allowing recombination (i.e., *x* − *y*). (B) Distribution of time to the MRCA of the parents of each QC-passing recombination node. (C) Distribution of the Pango lineage distance between parents. (D) Copying patterns (STAR Methods, Figure S17) for XA, XW and XBB.

Detectable recombination is rare in SARS-CoV-2, and ultimately judgments about plausibility require expert analysis. To aid such interpretation and help identify QC issues, we developed a visualisation tool inspired by SnpIt^111^ and the recombinant view of RIVET^106^, showing the recombination-informative sites between a recombinant and its parent sequences, and highlighting potentially problematic loci. Each copying pattern shows the positions where the allelic state in a recombinant (middle row) matches that of the left (upper row, coloured green) or right parent (lower row, coloured blue), or where the recombination event requires a de novo mutation (gold, with mutational change below). Figure 3D shows some example copying patterns, illustrating a spectrum of plausibility. XA is an example of a highly plausible recombinant, shown by the large number of supporting loci on the left and right. XBB is also strongly supported by many loci on the left and right, but the presence of many de novo mutations observed in the copying patterns suggests the existence of many genetically distinct, unsampled ancestors and some caution should be exercised when interpreting them (see Document S1.9.22 for more details). See Figures 3D and S13 for copying patterns of the Pango X lineages, and Figure S17 for copying patterns highlighting specific types of QC problems (and more details on the visualisations).

### Precision of recombination event times and positions

The genomic location of a recombination breakpoint in general cannot be determined exactly at single base resolution, but we can place bounds by inspecting sequences of the recombinant and its parents. The breakpoint interval of a recombinant is defined by two positions: a left position where the recombinant matches the left parent sequence but not the right, and a right position where the recombinant matches the right parent sequence but not the left. Genealogical methods such as sc2ts and UShER+RIPPLES use inferred parental sequences and are therefore in principle maximally precise. Other methods are limited in precision by the various proxies used to represent distinct parental lineages^37,39–41^.

The distribution of breakpoint intervals of the QC-passing recombination events in the sc2ts ARG shows an overall enrichment of breakpoints towards the 3’ end of the genome, particularly at the left boundary of the Spike gene (Figure 4A,B), consistent with the patterns seen in previous studies^38,41,106,112^. While this might result from a higher recombination rate in this region, it may also be due to the higher levels of polymorphism in the Spike gene, making recombination events more likely to be detected and allowing breakpoint intervals to be estimated more precisely. We found the median breakpoint interval length to be 2,207 bases (Figure 4C), indicating that the breakpoints of half of the events could be narrowed to a small region (less than ∼7% of the genome), while the uncertainty around the breakpoints of the other events is considerably larger (up to ∼54% of the genome). As expected from theory (STAR Methods), we found a negative relationship between the length of a breakpoint interval and divergence between recombinant parents (Figure 4D). Breakpoint intervals tend to be the smallest for events involving recombination between the deeply divergent Omicron BA.1 and BA.2 sublineages (the red points in Figure 4D).

**Figure 4:**
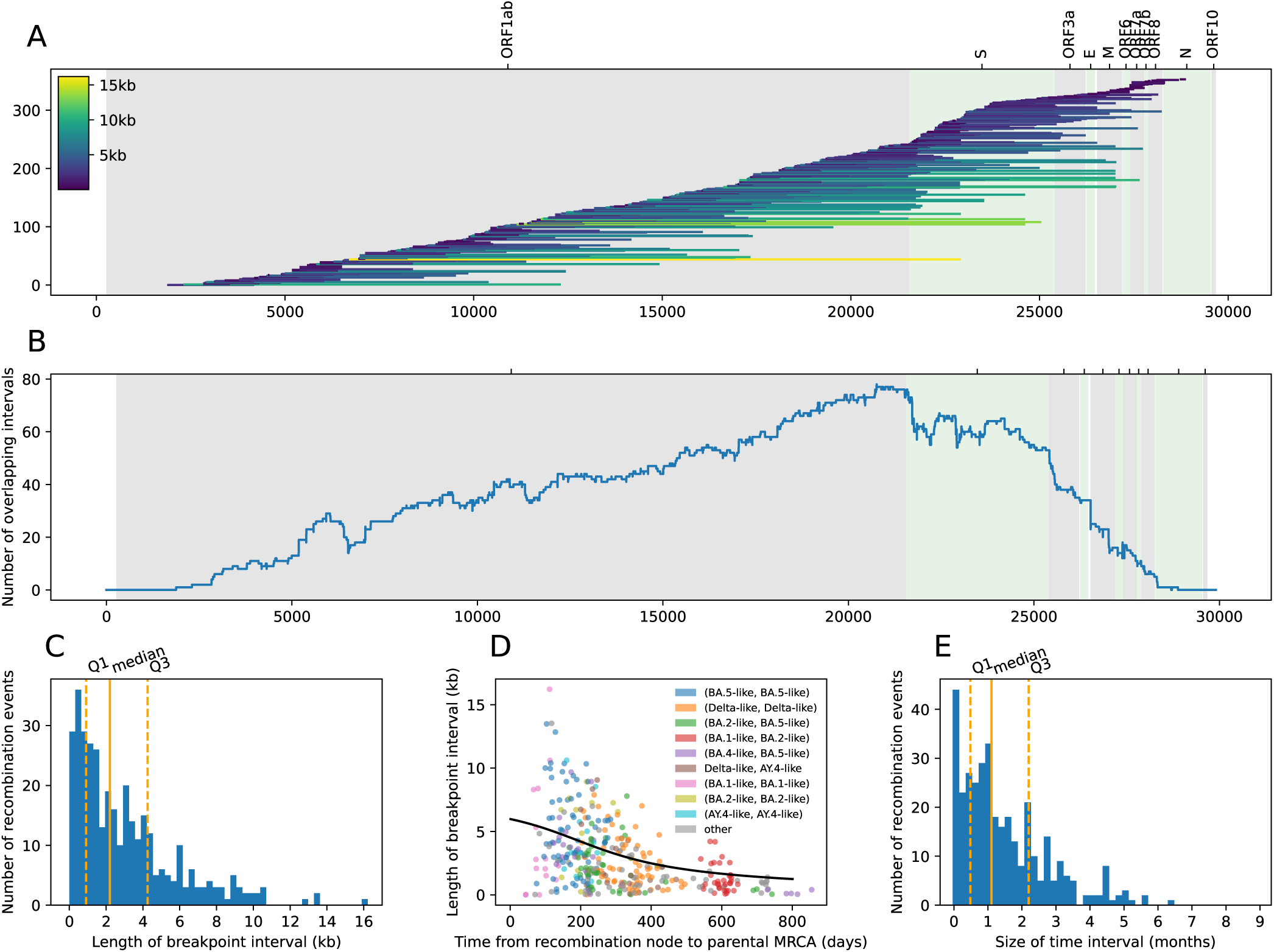
Recombination breakpoint and time intervals. (A) Distribution of the breakpoint intervals of 354 recombination events along the genome (annotated with gene labels on top). Each line represents a breakpoint interval coloured by its length. (B) Number of overlapping break-point intervals per site. (C) Length distribution of breakpoint intervals. (D) Relationship between breakpoint interval length and time between the recombination event and the parental MRCA. The black curve shows the theoretical expected relationship between these quantities, which is calculated assuming a genome-wide nucleotide substitution rate of ∼0.06 per day (STAR Methods). (E) Length distribution of time intervals around recombination events.

The time interval of a recombination node in the ARG is bounded by the age of the youngest parent and the age of the oldest child of the recombination node. We found that for ∼75% of the QC-passing recombination events, the time interval size is less than ∼2 months (Figure 4D); for example, the time intervals of the events associated with the Jackson et al. samples span 5 to 48 days. Longer time intervals imply that a parental or a recombinant lineage might have been unsampled for an extended period of time.

### Emergence of recombinants over time

We examined the frequency of emergence of the recombinants over the course of the COVID-19 pandemic (Figure 5). We explored two hypotheses for the timing of recombination. First, we would expect recombination events to occur more often when case counts are high, which increases the chance of co-infection within an individual, allowing recombination to occur. Second, we would expect recombination events to be detected more often when lineage diversity is high, as this increases the chance that two sufficiently distinct viruses co-infected an individual, increasing our power to detect recombination.

**Figure 5:**
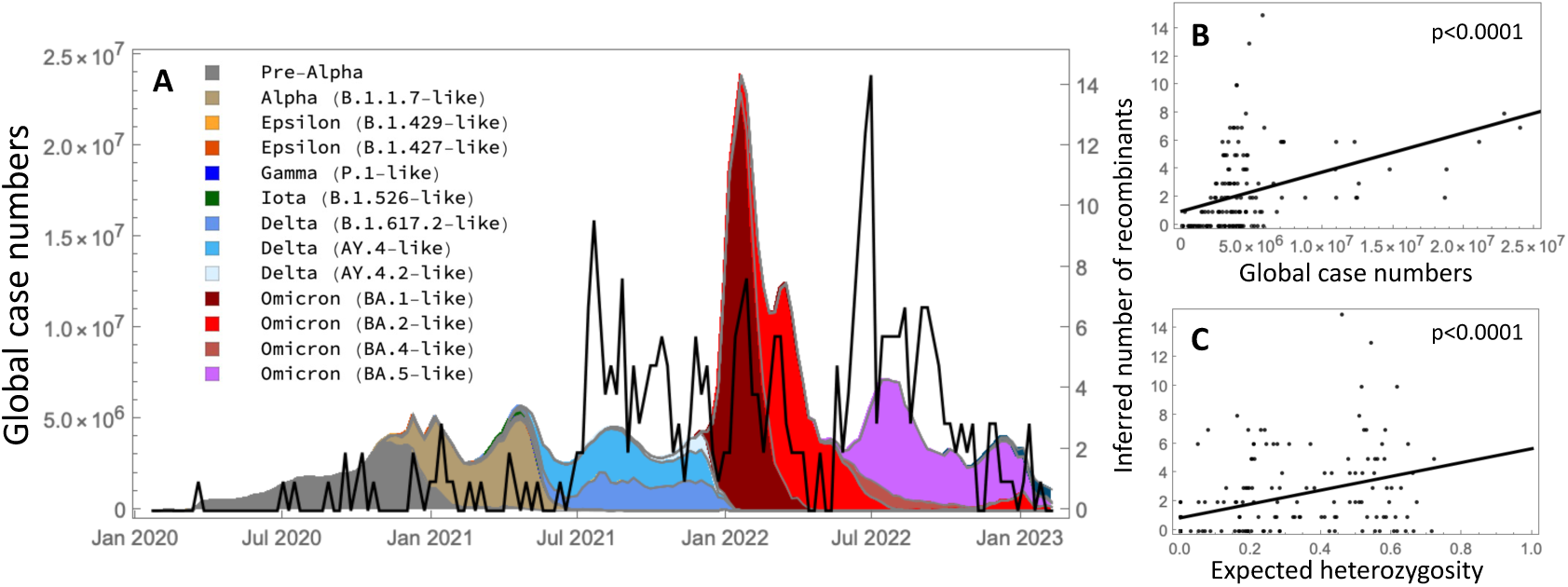
Recombination events across the pandemic. (A) Timing of the recombination events (black curve, right y-axis) relative to the global case numbers (^113^; left y-axis). Cases are colour-coded by their Scorpio labels, with the key showing those groupings with ≥ 1, 000 samples in the Viridian dataset. Recombinants more commonly occur (B) in weeks with high case numbers and (C) in weeks with extensive lineage variation, as estimated by high expected heterozygosity; solid lines show fits by linear regression, allowing for non-zero intercepts to account for unknown cases and heterozygosity not captured by the Scorpio labels.

We used the COVID-19 Data Repository at Johns Hopkins University^113^ to relate the timing of recombination events with global case numbers. We binned case numbers (*n*) by week from January, 2020 to March, 2023 and sub-divided by the proportion in each Scorpio label inferred within the ARG (shading in Figure 5). These lineages correspond loosely to a group of related lineages within a major VOC in the Viridian dataset. Alongside the case numbers, the black curve shows the number of recombinants inferred within each week along the ARG, focusing on the 354 events passing QC filters. Recombination events were inferred more often during the Delta and Omicron waves, when case counts were high. Overall, the number of recombination events rose with the global case count (Figure 5B; linear regression: 1.0 + 2.7 × 10*^−^*^7^*x*, *p* = 1.1 × 10*^−^*^7^). Recombination events also rose with the diversity of lineages in each week (Figure 5C; linear regression: 0.9 + 4.8*x*, *p* = 3.4 × 10*^−^*^7^), measured as the chance that two randomly drawn viruses have different Scorpio designations (*H*, referred to as the “expected heterozygosity”).

We then asked which single measure best explains the number of recombination events inferred each week, considering *n*, *H*, *nH*, *n*^2^, *n*^2^*H*. We included the squared number of cases each week thinking that it might better predict co-infections. The best single predictor of re-combination events involved the product of cases and expected heterozygosity (*nH*), with an adjusted *R*^2^ of 23.7%. This measure of case diversity (*nH*) explained 49% more of the variation in recombination events per week than the next best predictor (*H*, adjusted *R*^2^ of 15.9%). While consistent with the hypotheses that recombination events occur more often when infection rates are common and are detected more often when viral diversity is high, we note several sources of uncertainty. Global case numbers were underreported, particularly during the Delta and Omicron waves^114^. Furthermore, there may be substantial uncertainty in the timing of a recombination event, particularly if these occur in chronically infected individuals (as reported by^115^) with substantial delays before onward cases are detected. Finally, the filters that we applied removed all but the most strongly supported recombination events, although similar conclusions were obtained when all 855 recombinants detected by sc2ts were considered (STAR Methods).

### Comparison with UShER+RIPPLES

The most comprehensive analysis on recombination in SARS-CoV-2 to date has been performed using the combination of UShER and RIPPLES^38^. RIPPLES works by taking an UShER phylogeny and systematically breaking up sequences subtended by long branches into multiple segments that are placed independently. A recombination event at a node is flagged if the overall parsimony of these multiple placements sufficiently improves over the original single-parent origin in the phylogeny. This approach has much in common with the LS HMM used in sc2ts, but with one important distinction. While sc2ts detects and incorporates recombination in “real-time” as sequences are added to the ARG in daily batches, RIPPLES operates on a complete phylogeny, attempting to detect recombination post-hoc among millions of nodes. Like any phylogenetic reconstruction method that ignores recombination, UShER must force placement of recombinant sequences in places that minimise the discrepancy across the entire sequence, which may not be near any of the true parents. These permanent misplacements distort the phylogeny^3^, and subsequent tree reorganisations to maximise overall parsimony^25^ may break up the artefactually long branches used to detect recombination.

We compared the performance of sc2ts to detect recombination against RIPPLES run on the Viridian UShER phylogeny, subset to the 2,475,418 samples shared with the ARG (STAR Methods). To enable exact comparison, we omitted the QC filtering steps from both methods, and initially set the RIPPLES “parsimony improvement” parameter to *p* = 4 (rather than the more lenient default of 3), to match the equivalent *k* = 4 setting used in sc2ts (STAR Methods). Under this setting, RIPPLES reported 1174 recombination events of which 601 were validated by 3SEQ (51%; STAR Methods). We then compared the specific events reported by both methods by aligning them using sets of descendant samples (STAR Methods), giving 468 events shared between RIPPLES and sc2ts (Figure S18). The 706 recombination events detected by RIPPLES that do not correspond to a recombination event in the sc2ts ARG can largely be explained by a combination of the strict time ordering imposed by sc2ts and the differing sets of problematic sites masked by sc2ts and UShER (Document S1.11). The 387 events detected by sc2ts and not by UShER+RIPPLES include some of the well characterised Pango X lineages. Table S8 shows the events detected by UShER+RIPPLES associated with samples assigned to Pango X lineages, showing that at *p* = 4 only 9 of the Pango X lineages are present and with unequivocal recombinant origin for 6 (contrast with 16 Type I events in Table 1). We therefore reran RIPPLES with *p* = 3, resulting in 4111 events overall and increasing the intersection of events with sc2ts to 595 (Figure S18). At this more lenient threshold, 23 of the 44 Pango X lineages are identified as descendants of RIPPLES recombinant events, with unequivocal origins for 10 lineages, and 7 lineages associated with two competing events (Document S1.11). Thus, a more lenient threshold (*p* = 3) was needed for UShER+RIPPLES to detect a similar fraction of Pango X lineages as sct2ts with *k* = 4, resulting in more false positives. That said, XB and XAJ are among the lineages detected by UShER+RIPPLES, even at *p* = 4, strengthening evidence that these are false negatives for sc2ts (Table 1).

The choice of the *p* threshold has a large effect on the number of recombination events detected by RIPPLES, and previous studies have used the default value of 3^38,106^. For example, when applied to an UShER phylogeny of ∼1.6 million genomes sampled up to May, 2021, RIPPLES reported 589 unique recombination events with 43,104 descendant samples— corresponding to ∼2.7% of all samples of recombinant origin. Over the same time interval, the sc2ts ARG (*k* = 4) identifies 14 recombination events passing QC with 172 descendant samples among 501,235 samples (0.034%), a 40to 80-fold reduction. The sc2ts ARG identifies more of the well-characterised Pango X lineages at a more stringent threshold, and therefore provides us with a conservative lower bound on the frequency of detectable recombination events in SARS-CoV-2.

## DISCUSSION

There are two primary contributions of this work. The first is an ARG for 2.48 million SARS-CoV-2 genomes. Through multiple layers of validation against results from the literature, we have demonstrated that this ARG synthesises the evolutionary features of genetic inheritance, point mutations, deletions and recombination in a single cohesive structure. The ARG is freely available and easily accessible using the tskit software ecosystem, a rich and efficient platform for recombination-aware analyses widely used in population genomics. This combination of a deeply validated pandemic-scale ARG with a mature and highly capable software platform forms a unique resource. Previous landmark ARGs^54,116^ have helped spur the development of ARG-aware methods in human genomics^117–120^. We hope that the ARG provided here will similarly enable a range of recombination-aware analyses of SARS-CoV-2 and catalyse a wave of methodological developments integrating recombination into pathogen genomics. For instance, annotating the ARG with experimentally derived fitness estimates for mutations can help elucidate selection pressures on different recombinants; saltational variants can also be detected by analysing the rate and position of mutations along long branches (which persist after accounting for recombination). We envisage the ARG as a long-term stable resource—building on the high-quality, freely available reassemblies of the Viridian dataset^76^—to facilitate the study of the pandemic phase of SARS-CoV-2 in an open and reproducible manner.

The other primary contribution of this work is the method used to infer the ARG, sc2ts. The ability to infer an ARG over millions of whole genomes is a step change in scale over existing ARG inference methods for pathogen data^15–17,19,33^. To ensure these innovations are well-founded, we have paid significant attention to methodological detail, carefully validating the ARG (and consequently the method) against known results. Sc2ts is currently specialised to take advantage of unique features of the SARS-CoV-2 dataset, but the methods may be readily adapted to other pathogens. The LS HMM is the most important element of sc2ts, efficiently finding parsimonious placements for samples among all possible recombinant solutions. The current parametrisation—emphasising interpretability—is simplistic and could readily be extended to account for mutation rate variation along the genome^121^ or transition-to-transversion rate bias^78,122^. Our analysis of deletions shows that valuable phylogenetic signal is lost when they are ignored, and an important avenue for future work is to explore ways of incorporating them directly into the HMM. Another methodological innovation is the use of the tsdate method^54,93^ to estimate internal node dates in the ARG, illustrating the reuse of methods initially developed for population genetic applications. While results are encouraging, dating of phylogenies is a complex task^123^, and there is certainly room for improvement.

Further contributions of this work arise from the detailed study of the sc2ts ARG, providing insights into the evolution of SARS-CoV-2 and the COVID-19 pandemic. First, we clarify the relationships among recombinant Pango X lineages, finding examples of clustering of named lineages under a single shared recombination event. Second, we used the ARG to study break-point intervals for recombination events, showing that the relationship between the size of the interval in base pairs and the divergence time of the parental lineages matches theoretical expectations. Thirdly, we demonstrate a clear relationship between genetic diversity and the rate of detectable recombination, with the best predictor of the number of recombination events over time being the product of global case numbers and lineage diversity. Finally, and perhaps most fundamentally, through our conservative parameter choices and real-time handling of recombination events, we provide a stringent lower bound on the amount of recombination that occurred during the pandemic phase of SARS-CoV-2. These results underscore the utility of the ARG and provide a template for future analyses.

### Limitations of the study

We have focused on applying sc2ts to the Viridian dataset^76^ rather than the larger GISAID resource^6^ for two reasons. Firstly, the Viridian dataset is of higher quality and major efforts have been made to remove sources of error. Applying sc2ts to GISAID data is straightforward, but significant additional efforts would be required to address data quality issues. Secondly, the Viridian data is publicly available and redistributable, enabling us to share the resulting ARG without restrictions.

It is important to acknowledge that sampling is non-uniform both geographically and over time^124,125^. Sc2ts performs best in the data-dense regime, when the underlying assumption that genetic diversity is gradually accumulating within the structure holds. Large changes (i.e, saltation events) present significant challenges, and the results of the algorithm should be regarded more critically in these situations. Future methodological improvements may combine the strengths of sophisticated model-based inferences with the power of non-parametric inference^126^ when such data sparsity is detected.

## RESOURCE AVAILABILITY

### Data and code availability

- The ARG in tskit format has been deposited at Zenodo and is publicly available at https://zenodo.org/records/17558489. The UShER tree in tskit format is also available, as well as the intersection files.
- The sequences alignments in VCF Zarr format have been deposited at Zenodo and are publicly available at https://zenodo.org/records/16314739.
- The Snakemake workflow for aligning and converting the Viridian v05 dataset to VCF Zarr is publicly available at https://github.com/jeromekelleher/sc2ts-paper/blob/main/viridi dataset/.
- The complete configuration file for running sc2ts to build the ARG above is publicly available at https://github.com/jeromekelleher/sc2ts-paper/blob/main/inference/viridian_ config.toml.
- The Snakemake workflow for ARG post-processing is publicly available at https://github.com/jeromekelleher/sc2ts-paper/blob/main/arg_postprocessing/Snakefile.
- The original code of sc2ts is publicly available on GitHub at https://github.com/tskit-dev/ sc2ts.
- The Jupyter notebooks and the Mathematica script to perform the analyses in this paper are publicly available on GitHub at https://github.com/jeromekelleher/sc2ts-paper/.

## ACKNOWLEDGMENTS

We would like to thank Jotun Hein for helpful comments on the manuscript.

SHZ is supported by an NDPH Intermediate Research Fellowship (Oxford Population Health, University of Oxford). IG is supported by Wellcome (DPhil in Genomic Medicine and Statistics, 228319/Z/23/Z). DSP acknowledges funding administered by the Danish National Research Foundation in support of the Pioneer Centre for SMARTbiomed. SO acknowledges support from the Natural Sciences and Engineering Research Council of Canada (RGPIN-2022-03726). JK acknowledges support from the Oxford-Janssen Translational Genomics Fellowship, research grant R01 HG012473 from the National Institutes of Health NHGRI, research grant EP/X024881/1 from the Engineering and Physical Sciences Research Council, and the Robertson Foundation. For the purpose of Open Access, the authors have applied a CC BY public copyright licence to any Author Accepted Manuscript version arising from this submission.

Computation used the Oxford Biomedical Research Computing (BMRC) facility, a joint development between the Wellcome Centre for Human Genetics and the Big Data Institute supported by Health Data Research UK and the NIHR Oxford Biomedical Research Centre. The views expressed are those of the author(s) and not necessarily those of the NHS, the NIHR or the Department of Health.

## STAR METHODS

### Method details

#### Ancestral Recombination Graphs

The topology of an Ancestral Recombination Graph (ARG) can be defined as a set of *nodes*, representing a genome that existed at some specific time, and *edges* that describe the genetic inheritance relationships between those genomes^43^. Each edge defines the parent and child nodes and a genomic interval over which inheritance occurs. The “succinct tree sequence” ARG encoding defines these relationships in a simple and concise tabular format, sufficient to represent arbitrarily complex patterns of recombination^43^ and providing the basis for a range of highly efficient algorithms^47,62,118,119,127^. The mature and feature-rich tskit library has interfaces in Python, C and Rust, extensive documentation, and is used in numerous downstream applica_tions_^49,53,54,63–68,70–75^.

Similarly to the Mutation Annotated Tree (MAT)^26^ format (used by UShER^24^, matOptimize^25^, RIPPLES^38^, and RIVET^106^), tskit incorporates mutational information directly in the data model. This is done using the site (defining the genome position and ancestral state) and mutation (defining the site, node, and derived state) tables. In addition, tskit provides a flexible system for attaching metadata to all aspects of the data model, which can either be in JSON format for flexibility (used heavily by sc2ts for debugging information) or in a vectorisable packed binary format for efficiency (used to attach sample identifiers and Pango lineage assignments in the final ARGs). Tskit also provides a “provenance” table, which provides a mechanism for recording software version and parameter information through each step of complex processing pipelines.

#### Pandemic-scale alignment storage

To provide the efficient access to alignment data required for inference with sc2ts, we developed a new format based on the VCF Zarr^80^ specification. Briefly, Zarr is a storage format for scientific data, originally developed for the *Anopheles gambiae* 1000 Genomes Project^128^, and is now seeing widespread adoption across the sciences^129–136^. Zarr is essentially a simple mechanism for storing n-dimensional array data as a regular grid of compressed chunks, which provides high levels of data compression, efficient access across multiple dimensions, and is ideally suited to modern cloud-based deployments^80^.

We converted the Viridian v05 dataset^76^, consisting of 4,484,157 SARS-CoV-2 consensus sequences (125 GiB over 48 FASTA files) and 30 metadata fields (1.4 GiB TSV file) to VCF Zarr using a reproducible Snakemake^137^ pipeline. We aligned each sequence to the Wuhan-Hu-1 reference sequence (MN908947.3) using MAFFT v7.475^138^, with the option --keeplength to retain the reference coordinate system. We then used sc2ts import-alignments and sc2ts import-metadata to convert all alignments and metadata to VCF Zarr format, resulting in a single 401 MiB file (around 77× smaller than gzip compressed FASTA^80^).

A classical trade-off in bioinformatics is whether to store data in sample-major (FASTA) or variant-major (VCF) format, determining whether the data can be accessed efficiently either by row or column of the variant matrix. Because VCF Zarr is chunked in two dimensions, we can now access either sample alignments or the variation data for a given site across all samples efficiently. While the VCF Zarr encoding of the MAFFT-aligned Viridian sequences and meta-data can be accessed via native Zarr libraries in several popular programming languages (see Czech et al.^80^ for an overview), the sc2ts Python library provides a convenient interface. The sc2ts.Dataset class provides efficient access to alignments for samples and variant data for sites along the genome, as well as methods to export subsets of the data to FASTA if required. Document S1.2 gives an example of using this interface, where we retrieve alignment data for millions of samples in a few minutes on a standard desktop computer with minimal memory overhead. See Czech et al.^80^ for more details and benchmarks.

Both the pipeline and converted Viridian dataset are available for download; see Data and code availability for details.

#### Sample filtering and alignment pre-processing

The collection date of a sample is an important piece of information for sc2ts. We used the Viridian consensus date (“Date tree“), based on information from COG-UK, GISAID, and ENA/SRA as our source. Samples that have missing or incomplete dates (e.g., 2020-01) were omitted. Samples with a collection date of December 31, 2020 were also omitted, as the number of sample on this date (∼40,000) was a major outlier for this period of the pandemic and is likely enriched for incorrect collection dates. Manual checking (by cross-referencing the ENA and COG-UK Mutation Explorer) confirmed that some of these samples were not collected in 2020.

A set of problematic sites for SARS-CoV-2 that are enriched for systematic sequencing errors or high levels of homoplasy was compiled and updated during the pandemic^108,139^. Although these sites are recommended to be masked for UShER inference^107^, the list has not been updated to take into account the systematic improvements made by the Viridian dataset, and we therefore assembled a custom set of sites to mask. To do so, we first built a preliminary ARG from the samples collected up to June 30, 2021, without any site masking. Then, we identified the top 100 sites in terms of mutation counts in the ARG. Only 11 of these sites (635, 8835, 11074, 11083, 15521, 16887, 21304, 21305, 21575, 21987 and 28253) were present in the problematic sites file (https://github.com/W-L/ProblematicSites_SARS-CoV2), and most sites in this list were typical in terms of the number of mutations. We therefore masked our 100 most mutation-rich sites to remove sequencing errors and improve matching performance in the LS HMM.

Alignments read from the VCF Zarr file were first preprocessed to mask any non-nucleotide characters (A, C, G, or T) as missing data. Then, excluding the 100 sites identified as problematic, we filtered any samples that had missing data at more than 500 sites (this was set to 10,000 sites for the initial period up to 2020-03-01 when sampling density was low).

#### Li and Stephens model

The Li and Stephens (LS) model^81^ is an approximation of the coalescent with recombination that captures many of the key features of the joint processes of mutation and recombination. It is a Hidden Markov Model (HMM) in which a focal genome is modelled as a sequence of nucleotides that are probabilistically emitted as an imperfect mosaic of a set of reference genomes (Figure S19). The LS model is an HMM that models a focal genome as a sequence of nucleotides (the observed states), which are probabilistically emitted as an imperfect mosaic of a set of genomes in a reference panel (the hidden states). The process of switching between the genomes in the reference panel is governed by a transition matrix, and mismatches between a focal genome and a genome of the reference panel (from which it is “copied”) are permitted as emissions.

In the standard formulations of the LS model^81,84^ the probability of switching between the hidden states (i.e., genomes in a reference panel) when going from one site to the next is dependent on (1) the rate of recombination between the sites and (2) the number of genomes in the reference panel (*n*). Sc2ts uses the same formulation except that *n* is set to 1. This special case of the LS model leads to an intuitive interpretation of the ratio between the probability of switching and the probability of mismatch: When considering only two copying paths, the path having *k* mismatches and the other path having one switch but no mismatch are equally probable. We call this parameter *k* the “recombination penalty”. This parameter thus allows us to adjust the relative importance of recombination and mutation when finding the best explanation for a given sample. For simplicity, we assume that any single base mutation causing a mismatch is equally probable, ignoring transition-transversion biases and variation of mutation rate along the genome.

We use the efficient Viterbi algorithm (which finds the most likely sequence of hidden states in a HMM) implemented in tsinfer^53^ to find copying paths under this model. We updated this implementation to include a “likelihood threshold”, which allows all possible paths through the HMM with likelihood less than a given value to be considered equally improbable. This approach leads to major gains in likelihood compression, saving memory and reducing computation time. We then use this likelihood thresholding approach to iteratively process the samples for a given daily batch, first finding all exact matches, then all with 1 mismatch, etc. By setting the appropriate likelihood threshold for the HMM, we are guaranteed that any HMM solutions found at this threshold are correct, and can then run the HMM again with a lower threshold for the remaining samples. Close matches are found very quickly using this approach, and it is only samples that are more distant from the current ARG that require significant processing time and memory. This is reflected in the computational resources reported in Figures S1,S2.

#### Choice of k parameter

We explored using several values of *k*, by running sc2ts with *k* set to 3, 4, and 5 on the samples collected in 2020. We would expect that a good value for *k* should lead to parsimonious ARGs that capture plausible recombination events without incurring too many mutations, particularly reversions. A value of *k* too low could result in many artefactual recombination events, whereas a value of *k* too high could miss many genuine recombination events. We found that ARGs built with *k* set to 3 contained many dubious recombination events, but ARGs built with *k* set to 5 failed to include the well supported recombination events documented in a previous study^29^. Hence, we decided to use *k* set to 4 to infer the final ARG described in this study.

#### Tree inference from daily sample clusters

With tens of thousands of samples being added to the ARG per day, there are often clusters of hundreds of sequences with the same LS Viterbi solution. That is, large clusters of samples attach to the same node (or more generally, the same recombinant path) in the ARG. While some of these samples will require no extra mutations (as they are identical to the attachment node), in general there will be complex patterns of shared mutations among the samples reflecting their evolutionary relationships. We use a standard neighbour-joining algorithm^140^ to infer these within-cluster evolutionary relationships. We first compute the Hamming distance between all pairs of samples using SciPy^86^, and then use biotite^90,91^ to compute the neighbour-joining tree. Then, mutations are mapped back onto this daily sample cluster tree using the Fitch-Hartigan parsimony algorithm^141,142^ implemented in tskit. Finally, the resulting tree and mutations are added to the ARG at the attachment node(s) specified by the shared LS copying path.

#### Parsimony improving heuristics

Attaching trees built from the clusters of samples that copy from a particular node is an inherently greedy strategy and can produce inferences that are clearly unparsimonious. The final step in adding a daily batch of samples to the ARG is therefore to perform some local updates that target specific types of parsimony violations in the just-updated regions of the ARG. There are currently two parsimony-increasing operations applied, which we refer to as “mutation collapsing” and “reversion pushing” (Figure S20).

Given a newly attached node, mutation collapsing inspects its siblings from previous sample days to check if any of them share (a subset of) the mutations that it carries. If so, we increase the overall parsimony of the inference by creating a new node representing the ancestor that carried those shared mutations and make that new node the parent of the siblings carrying those shared mutations. The patterns of shared mutations between siblings can be complex, and the current implementation uses a simple greedy strategy for choosing the particular mutations to collapse. 119,844 nodes were added to the ARG during primary inference by mutation collapsing.

The reversion push operation inspects a newly added node to see if any of its mutations are “immediate reversions”; that is, are reversions of a mutation that occurred on the new node’s immediate parent. We increase the overall parsimony of the inference by “pushing in” a new node which descends from the original parent and carries all its mutations except those causing the reversions on the newly added node. Reversion pushing led to the addition of 52,336 nodes during primary inference.

#### Filtering time travellers

Time-travelling samples—those in which the recorded collection date differs substantially from their true collection date—pose significant problems for sc2ts and are present at a significant frequency. For example, of the 43 Scorpio^143^ designations present in the Viridian dataset studied, 32 have samples present with sampling dates in 2020, including lineages that do not arise until late 2022. Inserting these time-travellers into the ARG at the claimed sampling date causes major artefacts, and they must therefore be filtered out. We use the simple approach of filtering out samples that exceed a given “HMM cost”, i.e., requiring too many mutations or recombination breakpoints to plausibly represent the steady accumulation of diversity expected with dense sampling. The HMM cost for the Viterbi solution for a sample under the LS model is *k* times the number of recombination events plus the number of mismatches, where *k* is the recombination penalty (here, *k* = 4). After some experimentation, we set the HMM cost threshold to 7, disallowing samples that require 8 or more mutations (or, e.g., 1 recombination and 4 mutations, or 2 recombinations) from immediate inclusion (see next section) in the ARG. This approach has the benefit of filtering samples enriched for errors as well as time-travellers.

#### Inserting saltational lineages

The HMM cost threshold approach outlined in the previous section is effective at removing time-travelling samples, but it also filters out important evolutionary events such as “saltations” that characterize major VOCs. These lineages are characterised by a high number of accumulated mutations, and are hypothesised to emerge from chronic infections during which the virus is exposed to individual-specific immune selective pressures^144,145^. To distinguish time travellers from true saltational lineages, we re-evaluate the evidence for each lineage with a high HMM cost as additional samples are added. If these samples form a sufficiently plausible group over successive days (under criteria described in the next paragraph) sc2ts adds the group to the ARG as a “retrospective sample group” using same mechanisms as standard daily sample clusters.

Distinguishing a truly emerging outbreak from different forms of correlated error is non-trivial, and sc2ts has 5 parameters to fine-tune the behaviour. Firstly, we require at least 10 samples in the group to ensure there is sufficient support over a time window of seven days. Then, we build a local tree of the samples and consider some tree metrics to evaluate whether the samples within the group are closely related. If we have at least 2 mutations shared by all samples, there are at most 2 recurrent mutations within the group and at most 5 mutations unique to each sample, the retrospective group is accepted for inclusion. Although this process is highly heuristic, it successfully captured the majority of saltational lineages such as Alpha (Figure S10), and the vast majority of samples were added following the standard daily sample grouping mechanisms. In total, 882,398 daily sample groups were added to the ARG, while only 100 were added via this retrospective grouping mechanism.

The Delta, BA.1, and BA.2 lineages were particularly challenging, however, as the initial emergences of these lineages were not densely sampled. In this case, there is little information for the retrospective sampling grouping mechanism to work with and it is very difficult to distinguish short-term time travellers from true outbreak samples. For these cases, we chose specific high-quality samples with confident dates as “seeds” which are unconditionally included in the ARG on the specified date.

For Delta (first detected in December, 2020 in India^146^), we used two seeds: ERR5461562 assigned to B.1.617.1/Kappa (dated February 22, 2021 from the UK) and ERR5676810 assigned to B.1.617.2/Delta (dated March 23, 2021 from the UK). Kappa is closely related to Delta^99^, and therefore seeding it helps to reconstruct the origin of Delta. These are the earliest Kappa and Delta samples in the Viridian v04 dataset that were used for the designation of these Pango lineages. The resulting subgraph around Delta in the ARG can be seen in Figure S11; for further explanation, see Document S1.5.2.

For Omicron BA.1 and BA.2 (first detected in South Africa or Botswana in November, 2021^146^), we used SRR17041376 and SRR17461792, respectively. We chose SRR17041376 (dated November 6, 2021 from South Africa) for the BA.1 seed, as it is among the earliest African Viridian samples, and its initial placement in the ARG mirrors the initial placement of a subsequent high-quality COG-UK sample (ERR7443564). For BA.2, we chose SRR17461792 (dated November 27, 2021 from South Africa), which is the earliest BA.2 sample present in the Viridian v04 dataset. Note that this sample is dated later than SRR17041376 (the BA.1 seed), allowing time for the BA.1 lineage to become established and accumulate descendant samples in the ARG. The resulting subgraph around these Omicron lineages in the ARG can be seen in Figure S12. For further explanation, see Document S1.5.3.

With the combination of these mechanisms—gradual accumulation of diversity through daily sample groups, retrospective group inclusion and manual seeding—we successfully incorporated all major lineages in the dataset into the ARG (Figure S2).

#### Primary ARG inference

The “base ARG” we infer with sc2ts is the starting point for later analyses, and this step dominates overall processing time. To summarise, we first initialise the ARG using the Wuhan-Hu-1 reference sample (MN908947.3) as the root node, and then ran the algorithm day-by-day using the parameters described in the preceding sections and documented in the configuration file (see Data and code availability). As discussed above, computational resources are dominated by the LS model (Figures S2,S1) with all other aspects contributing negligibly.

#### Recombinant rematching

The first step in ARG postprocessing is to examine putative recombinants to determine whether a more parsimonious explanation is possible. This is done by rematching the inferred recombinant sequence against the ARG pruned back to include only samples before the date of the recombinant. Where the recent ancestry of a recombinant includes a branch containing large numbers of mutations, it is possible that a non-recombinant solution could be found by postulating an intermediate node along that branch, that contains only a subset of those mutations. In particular, for a given mutation-rich branch, artefactual recombinants may be found in which one recombinant region matches to a node like the parent whereas another region matches to a node like the child of that branch. Such recombinants can be identified by the presence of recurrent or reversion mutations in the left or right parent branches. To help remove these recombination events from the ARG, we note the genomic positions of mutations associated with the recombinant, and identify branches in their ancestry which have mutations at the same positions. Of these branches, we chose the one that contains the largest total number of mutations as a candidate for intermediate node creation. This is done by sorting the mutations on the branch such that the ones that match the allelic state in the recombinant are placed above the intermediate node, and the others are left below the node. We refer to this process as “long branch splitting”, and it creates additional nodes which are available for recombinant rematching.

Using recombinant-rematching, 58 recombinants were identified as being more parsimoniously explained by matching to existing nodes in the ARG (inserted by subsequent parsimony improving heuristics) and 16 by long branch splitting. Of these 16 recombinants, two are noteworthy as being the origins of the Delta (B.1.617.2) and Omicron-BA.2 lineages in the ARG. Both result in substantially fewer mutations (15 fewer for Delta, 6 for BA.2) than the previous recombinant HMM solution. We then updated the ARG to remove these 74 artefactual recombinants by “rewiring” the recombination node to the more parsimonious non-recombinant solution. This resulted in 16 additional nodes, 182 fewer mutations and reduced the number of trees along the genome from 348 to 317. The long branches involved in the origins of Delta and BA.2 originally joined B.1 to B.1.167 and B.1.1 to BA.1 respectively; they can be seen in Figure S11 and S12 with inserted intermediate nodes leading to the Delta and BA.2 origination nodes.

#### Postprocessing pipeline

This section described the various postprocessing steps that we performed after primary inference, which is encoded as a reproducible Snakemake workflow (see Data and code availability). The first step is to perform recombinant rematching as described in the previous section, to find any recombinants that can be more parsimoniously explained either by matching to nodes subsequently added or by targeted splitting of long branches.

For performance reasons, samples that match exactly to an existing node in the ARG are not added during primary inference, but are stored in a secondary “match database”. During postprocessing, we added 1,252,208 of these exact match samples into the ARG using the stored information.

During primary inference, gap characters (“-”) are masked out as missing data, but this excludes both important information about deletions from the ARG as well as potentially informative phylogenetic signal. We therefore performed post-hoc parsimony mapping at 163 sites to include the effects of deletions and to ensure that all Pango-informative sites are included in the ARG. Using deletions with a minimum frequency of 1% (count of sequences divided by 9,149,680 sequences analyzed) from Table S1 of Li et al.^79^, we chose 75 sites for deletion remapping (see Document S1.6 for analysis of the major deletions identified from this process). In addition, we remapped data at 88 sites that were excluded from primary inference and are defined as Pango lineage defining sites by Freyja^147^ (https://raw.githubusercontent. com/andersen-lab/Freyja/refs/heads/main/freyja/data/lineage_mutations.json). Alignment data for these 163 sites was then mapped to the ARG using the Fitch-Hartigan algorithm^141,142^ implemented in tskit, including the gap character as a state along with the four nucleotides. Following this, we iteratively applied the node parsimony heuristics in sc2ts until there were no further reductions in the numbers of mutations.

We computed Pango assignments for all 2,747,985 nodes in the ARG by first exporting their aligned sequences to a 77GiB FASTA file and then running pangolin (version 4.3.1; pangolin-data version 1.29). Note that this version of pangolin uses UShER placement for classification^148^. The resulting Pango and Scorpio designations were then stored as metadata associated with each node in the ARG.

When then estimated non-sample node dates using the “variational-gamma” algorithm in tsdate^54,93^ version 0.2.3. See Document S1.4 for more details and analysis of these dates.

Finally, we reduced the metadata stored in the ARG to a minimal set (node sample IDs, and the computed Pango and Scorpio designations), encoded to enable efficient columnar access, and compressed the file using tszip (version 0.2.5). The final file requires 32MiB of storage.

#### Quality control of recombination events

We examined the genomic positions at which the suggested parents differed in allelic state. Where such sites clustered closely (3 or fewer bases apart), we treated the entire cluster of sites as a single *locus*. For each genomic region identified by the HMM as having a single parent, we calculated the “net number of supporting loci” by counting the number of loci in the region that supported the suggested parent minus the number that did not support it. Finally, with all the recombination events in the ARG having a single breakpoint, we calculated the net number of supporting loci on the left and right sides of each breakpoint, and used this to plot all the 855 recombination events in Figure S16.

This simple QC measure effectively differentiates artefactual recombination events from those that are well supported. All the sc2ts-detected recombination events associated with the Pango X lineages (Table 1) occur in the upper right of Figure S16, with high numbers of net supporting loci on both the right and left parents. In contrast, the other regions of the plot, where one or both parents are less well supported, are dominated by recombination events associated with samples sequenced using the Ion AmpliSeq protocol. Coupled with closer inspection of copying patterns (see below), we chose a QC cutoff of 4 or more net supporting loci on both the left and right side of the breakpoint. This leads to 354 QC-passing recombination events (Figure S16, upper right quadrant), compared to 501 recombination events for which there is less support and which may be artefactual (shaded quadrants).

Inspection of the copying patterns for all the recombination events identified associations between recombination events that failed QC and known problematic loci in the SARS-CoV-2 genome. Figure S17 shows illustrative copying patterns (Q1 to Q4 correspond to the labelled quadrants in Figure S16). Q1 shows a well supported recombination event that passed QC (albeit marginally), having a net number of supporting loci of 4 on both sides of the breakpoint. Q2 shows an artefactual recombination event caused by a 6-base Delta lineage-defining deletion. There are 71 (14%) QC-failing recombination events involving this deletion, and many cases are affected by a known amplicon dropout caused by this particular deletion^109^. Q3 shows an artefactual recombination event caused by a combination of this Delta deletion and a known MNS (see below). Finally, Q4 shows an artefactual recombination event associated with the A507T-T508C-G509A MNS, which has been identified as a sequencing error due to incomplete read trimming^149^. There are 9 recombination events involving this particular MNS, of which 6 failed QC.

#### Validation of recombination events

We validated recombination events in the sc2ts ARG using 3SEQ^110^ as follows. We first exported the alignments for the recombination node and both parents to FASTA format, and then ran 3SEQ in “full mode” independently for each recombinant. Specifically, we ran 3seq -full parents.fa recombinant.fa using default thresholds. We used 3SEQ source code downloaded from https://gitlab.com/lamhm/3seq at git hash e45ee3b4.

For CovRecomb^39^, we obtained the original alignments for the “causal” samples for each recombination node, and ran CovRecomb-Local-Version with default settings. We used source code downloaded from https://github.com/wuaipinglab/CovRecomb at git hash 1fbf6b2.

We ran rebar in a similar manner, exporting the original alignments for the causal samples to FASTA and using default parameter values. We used rebar version 0.2.1 from bioconda^150^, and rebar dataset version “2025-01-28”.

These steps are included in the Snakemake postprocessing workflow (see Data and code availability).

#### Conversion of UShER tree to tskit

To facilitate comparison with the sc2ts ARG, we first converted the Viridian UShER tree from MAT^26^ to tskit format. We first downloaded the tree in JSON format (tree.all viridian.202409.jsonl) and converted the tree topology to tskit node and edge table descriptions. We then downloaded the protobuf formatted tree (tree.all viridian.202409.pb.gz), exported the mutations to nucleotide format using matUtils summary --translate (usher version 0.6.3 from bioconda), and translated these mutations into tskit site and mutation tables. The resulting file contains 5,345,019 nodes, 3,364,841 mutations and requires 29MB of storage space in tszip format (usher viridian v1.0.trees.tsz). We validated that the data encoding by comparing the variant data for the UShER tskit tree sequence against our MAFFT alignments. There was an exact match between the non-ambiguous nucleotide calls at 27,469 sites. At the remaining 39 sites for which the UShER tree had mutations there were a significant number of gap characters (deletion calls), suggesting that the mismatches could be due to differences in the alignments used as input to UShER. See the notebook validate usher tree.ipynb for details.

We then computed intersection ARGs for sc2ts (sc2ts viridian inter v1.2.trees.tsz) and UShE (usher viridian inter v1.2.trees.tsz) by simplifying^43,48^ both to the 2,475,418 samples and 27,507 sites that are shared (the UShER tree has mutations at 27,508 sites and sc2ts ARG has 29,893). As there was no documented means of deriving node dates from the UShER tree, and we wish to visually compare the phylogenetic backbones, we also dated the intersection tree by fixing the sample dates using the “Date tree” metadata field, and using tsdate (version 0.2.3) to date internal nodes.

All conversions were performed as part of the reproducible pipeline described in the Postprocessing pipeline section. The full UShER tree and the intersection files are available for download in tszip format (see Data and code availability).

#### Comparison of phylogenetic backbones

We compare the non-recombinant parts of the ARG with the UShER topology by simplifying the phylogenies to a smaller number of Pango representative samples. We focus on 1286 Pango lineages comprising those which are defined by perfect “origination events” (Document S1.3). but which are not acknowledged recombinant lineages (i.e. Pango X lineages and their descendants). To ensure that each sample is a tip in the simplified topologies, a representative sample is then chosen as the oldest sample descending from the origination node whose own descendants (if any) are entirely of the focal Pango type. We then further remove 4 lineages which are singleton descendants of a putative ARG recombination event, This results in a simplified ARG of 1282 samples, each sample from a different Pango lineage.

The simplified ARG contains only two recombination nodes, which are the two likely artefactual events discussed in Document S1.12, both of which have breakpoints near the 5’ end of genome. We therefore compare the UShER topology against the first tree in the ARG (covering positions 1-26858; the majority of the genome). To reduce plot density, we consider only those Pango lineages with greater than 10 samples. Figure S3 shows the resulting tree compared with the equivalent dated UShER tree as a tanglegram. We also show subtrees for Delta (Figure S4), BA.2 (Figure S5), and BA.5 (Figure S6).

The order of tips in each plot was chosen to reduce tangling, as calculated using the NeighborNet algorithm from Dendroscope^20,151^, with tanglegrams plotted using the SVG outputting facility built into tskit. See supp pdf-cophylogeny-tanglegrams.ipynb for details.

#### Comparison of mutational spectra

We calculated the mutational spectra of Alpha, Delta, Omicron BA.1, BA.2, BA.4, and BA.5 in the sc2ts ARG, considering mutations (ignoring indels) above the sample nodes and internal nodes which have Scorpio labels associated with the VOCs (e.g., for Alpha, B.1.1.7-like and B.1.1.7-like+E484K). Note that ∼98% of all the mutations in the ARG occur above the nodes that have a Scorpio label associated with these VOCs (excluding the nodes with unassigned Scorpio labels).

We calculated the mutational spectra of the same VOCs in the intersection UShER tree in tskit format using a similar approach. Here, we used the Scorpio sample assignments provided in the Viridian metadata, and therefore restricted to mutations above sample nodes only.

Next, we calculated the mutational spectra of the same VOCs using the mutational counts from an UShER phylogeny analysed by Bloom et al.^78^, which was built using a Nextclade pipeline and has samples collected worldwide up to November, 2022. To obtain the WHO label corresponding to a Nextclade-defined clade, we used the following mapping: 20I, Alpha; 21I and 21J, Delta; 21K, BA.1; 21L, BA.2; 22A, BA.4; and 22B, BA.5. The mutation counts were downloaded from https://github.com/jbloomlab/SARS2-mut-spectrum/.

See the fig mutational spectra.ipynb notebook for details.

#### Concordance of recombination events

We compare concordance rates of sc2ts in identifying the Pango lineages of recombinant parents and the location of breakpoint intervals with RecombinHunt^40^ and CovRecomb^39^. We use these two methods because detection results on the Pango X lineages were reported and made available in the original publications. We do not compare here with UShER+RIPPLES+RIVET because curated data on the X lineages is not available for this pipeline (see Document S1.11 and Table S8, however). We use the data from the Pango designation community (https://github.com/cov-lineages/pango-designation/) as the “truth” when comparing performance in identifying parent Pango lineages and breakpoints between the methods. We focus on the 16 Type I recombination events (Table 1) identified by sc2ts for simplicity. While this is certainly biased in favour of sc2ts (i.e., by ignoring likely sc2ts false negatives such as XB), it helps to understand the probable accuracy of sc2ts on the true positives. It is important to emphasise that the “true” truth set among the Pango X lineages is currently unknown.

For RecombinHunt, we obtained two sets of detection results of the Pango X lineages from the supplementary materials of Alfonso et al.^40^. One set of results is based on mutational profiles calculated from a global GISAID dataset, and the other set from a global Nextstrain dataset. The breakpoint intervals identified using RecombinHunt are defined as the left and right recombination-informative sites.

For CovRecomb, we summarized the detection results for the Pango X lineages as follows. First, we obtained the “independent” recombination events identified by Li et al.^41^ (see Table S4 of the study) which correspond to the Pango X lineages (if available). When multiple events corresponded to a Pango X lineage, we took the results of the event which had the highest number of epidemic recombinants. For other Pango X lineages, we downloaded the full list of putative recombinants identified in the study (https://github.com/wuaipinglab/CovRecomb/blob/main/ CovRecomb-Global-Version/putative_recombinants/putative%20recombinants.csv). For eac of these Pango X lineages, we took the most commonly observed combination of the left parent, the right parent, and the breakpoint interval. The breakpoint intervals are defined as the left and right recombination-informative sites.

The parents of a Pango X inferred using a method are deemed concordant if they have Pango lineage assignments that are at least as specific as those proposed by the Pango designation community. The breakpoint interval of a Pango X inferred using a method is deemed concordant if it overlaps with that proposed by the community by at least one base.

The results are reported in Table S3. See the tab methods concordance.ipynb notebook for details.

#### Calculation of Pango lineage distance using pangonet

We define a “pseudo-phylogenetic” distance to quantify the divergence between two Pango lineages. The idea is to count the number of edges separating two given Pango lineages in a pseudo-phylogeny which relates all the designated Pango lineages. To construct this pseudo-phylogeny, we used pangonet (https://github.com/phac-nml/pangonet) which mines the information on the Pango lineages in the “alias key.json” and “lineage notes.txt” files defining the designations (https://github.com/cov-lineages/pango-designation). The distance between two Pango lineages is then the number of edges separating the corresponding nodes in this graph.

#### Expected lengths of recombination breakpoint intervals

Recombination breakpoint intervals are imprecise when the recombining parents are identical in sequence at sites spanning the breakpoint. Here, we estimate the interval lengths expected around a breakpoint, given a genome-wide nucleotide substitution rate (*µ*), total time between two samples (*T*), and sufficient support to detect the recombination event. If a breakpoint occurs a fraction *P* along a genome of length *L* and *k* mutations have occurred to the left of the breakpoint, then we expect the nearest mutation to occur *PL/*(*k* + 1) bases to the left. If we require *n* mutations on both sides of the breakpoint, we can calculate the expected value of the interval to the left by summing across a Poisson-distributed number of mutations, conditioned on having at least *n* mutations, using the following equation (where Γ(*x*) is the gamma function, and Γ(*x, y*) is the upper incomplete gamma function):

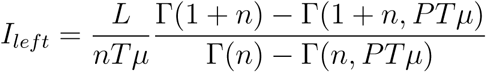

A similar result holds for the right interval (*I_right_*), but with *P* replaced with 1 − *P*. Summing the two intervals (*I_left_* + *I_right_*) gives the expected interval length. As detectable recombination events tend to be closer to the center of the genome, we use *P* = 0.5 in Figure 4D.

In Figure 4D, we calculated the expected lengths of breakpoint intervals by setting *µ* to ∼0.0598 per day. This is based on published estimates of the per-site nucleotide substitution rate of ∼ 2 x 10*^−^*^6^ per site per day^145,152,153^.

#### Comparison with UShER+RIPPLES

We began with the Viridian UShER tree downloaded from Figshare. To simplify the comparison with sc2ts, we subset this file down to the 2,475,418 samples shared with the sc2ts ARG by running matUtils extract. We then ran ripples-fast^106^ on this file, using -n1 so that all detected recombinants were reported (equivalent to sc2ts). We ran RIPPLES with the parsimony improvement parameter *p* equal to 3 (the default) and 4.

The output of RIPPLES is two text files, “recombination.tsv” and “descendants.tsv“. We use the “descendants.tsv” file, which lists the descendant samples of all recombination events keyed by the recombinant node ID, to match events with sc2ts. Specifically, for each RIPPLES event identified a list of descendant samples, we find the MRCA of that set of samples in the first tree of the sc2ts ARG. The clades are then said to agree if the number of additional samples descending from the sc2ts node is *<* 2 (Figure S18).

We ran 3SEQ on the RIPPLES recombinants by extracting the alignments for the child and parent sequences for each recombination event. We first obtained the parent node IDs from the “recombination.tsv” file by choosing the most parsimonious event associated with each recombinant node, arbitrarily choosing among the possibilities in the case of ties. We then used this list of child and parent node IDs as input to matUtils extract -v to extract their sequences in VCF format. Following this, we used vcflib^154^ to convert to alignments in FASTA format, as recommended by the UShER documentation (vcf2fasta -f NC 045512v2.fa). We then ran 3SEQ in “full mode” independently for each trio, in the same manner as described when validating the sc2ts recombinants.

The UShER tree is not dated, and so to obtain dates for parent nodes we used sample metadata (Date Tree from the Viridian metadata). For the minority of parent sequences that are not samples, we used the oldest sample descending from a given internal node as a proxy for its date. We used the BTE Python library (https://jmcbroome.github.io/BTE/build/html/ index.html) to perform the required traversals on the UShER tree.

The full workflow is encoded in the Snakemake postprocessing pipeline. See Document S1.11 for detailed analysis of the RIPPLES events, and the analysis ripples.ipynb notebook for full methodological details of the analysis.

### Quantification and statistical analysis

#### Analysing global case counts

To explore the timing of emergence of recombination events, global case count data were downloaded from the COVID-19 Data Repository by the Center for Systems Science and Engineering (CSSE) at Johns Hopkins University^113^ and binned by week from January, 2020 to March, 2023. The proportion of each lineage *i* in each week (*p_i_*) was then inferred from the Scorpio designation at each node within the ARG for that week. Genetic variation was measured as the “expected heterozygosity” across lineages, which is the chance that two lineages chosen at random from that week had different Scorpio designations 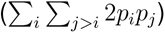. The linear regression models reported in Figure 5 allowed for a non-zero intercept, because there may be unreported cases even without any reported cases. The slopes were highly significant, and the p-values were confirmed by randomly permuting the predictor variable 10,000 times. ANOVAs were conducted to determine which single predictor (*n*, *H*, *n H*, *n*^2^, *n*^2^*H*) explained the most variation in recombination events by week. These statistical tests and graphics were conducted in *Mathematica* 14.2 (Wolfram 2024) and available at https://github.com/jeromekelleher/ sc2ts-paper/blob/main/notebooks/TemporalTrends_RecEvent25August2025.nb.

In addition to the analysis of the 354 recombination events that pass QC, we analyzed the relationship between the full set of 855 recombinants and case numbers. These additional recombinants occurred disproportionately during the Delta and first Omicron wave (July, 2021 to February, 2022). Again, the number of recombinant events rose with the global case count (linear regression: 3.1 + 5.3 10*^−^*^7^*x*, *p* = 0.0006) and with the diversity of lineages in each week (linear regression: 0.42 + 18.3*x*, *p* = 2 10*^−^*^12^). The single best predictor of recombination events among the predictors tested (*n*, *H*, *n H*, *n*^2^, *n*^2^*H*) was again the product of cases and diversity (*n H*), with an adjusted *R*^2^ of 26.9%, although this was now followed closely by expected heterozygosity on its own (*H*), with an adjusted *R*^2^ of 26.5%. Thus, similar conclusions are reached with either dataset, although filtering the recombination events to the 354 with better support revealed a stronger relationship with case numbers.

## Supplementary Material

### S1.1 Parsimony comparison of sc2ts and UShER

We performed an in-depth analysis of the sc2ts and UShER ARGs (we refer to the UShER tree as an ARG for simplicity here, as a single tree is also an ARG) in terms of parsimony. A direct comparison is difficult because of the different alignments used as input, the different sets of sites used for inference and the fact that we are including deletions for a subset of sites in the sc2ts ARG.

After site remapping, and including all mutations involving deletion characters, sc2ts has 143,690 mutations more (2,078,370 vs 1,934,680). If we exclude mutations involving deletions in sc2ts, this is reduced to 46,371 more (2.3%) than UShER. If we restrict to the 26,192 sites that were not remapped, sc2ts has 4,785 (0.26%) fewer mutations than UShER, and has around 2.5 times more mutations than UShER on the 163 sites that were post-hoc remapped.

To provide a more objective measure of the overall parsimony of the inferred ARGs with respect to the Viridian alignments, we remapped all 29903 sites in both ARGs using Fitch-Hartigan parsimony. Here the nucleotides ACGT and “-” were treated as distinct characters, with all other values regarded as missing data. In this case, the sc2ts ARG has 629,864 (5.5%) fewer mutations (10,840,687 vs 11,470,551). Some of this difference may be due to the post-hoc incorporation of phylogenetic signal from deletions by sc2ts, but more work is required to fully understand the differences between the two approaches.

See notebook analysis usher sc2ts.ipynb for details.

### S1.2 Imputation comparison of sc2ts and UShER

Both sc2ts and UShER support missing data in input alignments, and effectively “hard call” these values when the samples are integrated into the tree/ARG. In doing so, we impute the missing and ambiguous data for these samples. We evaluate the imputation performance of sc2ts and UShER by comparing their outputs on the Viridian alignments.

The total aligned dataset for 2,475,418 samples and 27,507 sites in the sc2ts and UShER intersection ARGs represents 63.41 GiB of nucleotide calls. Of the 162,838,461 missing data calls (Ns) in the alignments, sc2ts and UShER disagreed in their imputed values for only 203,579 (0.125%). Additionally, 956,859 calls made use of the IUPAC uncertainty codes. Of these sc2ts imputed 435 (0.05%) incorrectly (i.e., with a base that is not compatible with the ambiguity code). UShER imputed only 7 calls (0.007%) from this set incorrectly, likely due to special handling of ambiguous bases in the parsimony calculations.

This computation is a good example of type of integrated large-scale data analysis that can be easily performed using the combination of tskit and VCF Zarr. The entire analysis is performed in a short Jupyter notebook^155^, with the core calculation requiring about 30 lines of Python code. Retrieving data from the base alignments and ARGs for 2.48 million samples at the same time over all 27,507 sites took a little under 5 minutes, with a peak memory usage of about 4.5GiB of RAM on a standard laptop.

See notebook example data processing.ipynb for details.

### S1.3 Pango lineage “origination” events

To analyse how well the ARG reflects the phylogenetic structure implicit in the Pango lineage naming system, we considered the Pango assignments generated by Pangolin on the alignments for each node in the ARG (STAR Methods). If the ARG perfectly reflected the Pango lineage structure, each lineage would form a clade descending from a single originating node with that label. We identified putative origination nodes among those labelled with a given lineage as the node with the maximum number of descendant samples and the earliest time (if there is a tie).

Of the 2058 distinct Pango lineages in the ARG, 1473 of these (comprising 717798 samples) match perfectly, with unique origination events in the ARG where all samples assigned a given lineage descend from the first node assigned that lineage. A further 245 lineages (589197 samples) match perfectly when we count the descendants of the parent of the first node (accounting for polytomies in which multiple originating nodes for a given lineage are siblings). We then have 306 lineages (755840 samples) where the difference in the number descendants of the first node’s parent is *<* 100. The remaining 35 lineages (420047 samples) are dominated by a few large lineages such as BA.1.1 (147271 samples) and AY.4.2 (54607 samples) which have multiple non-sibling origins within the ARG.

See notebook analysis pango events in arg.ipynb for details.

### S1.4 Evaluating inferred node dates

To validate the accuracy of times for non-sample internal nodes inserted into the ARG (inferred using tsdate), we compared the dates of first emergence of Nextstrain clades against the Nextstrain SARS-CoV-2 tree (downloaded on 2023-01-21). Figure S8 shows relatively close agreement for most clades, with only three disagreeing by more than 28 days: 20C (an early lineage from 2020), 20E (Pango B.1.177 also known as EU1, emerging mid-2020), and 21B (Pango B.1.617.1 also known as Kappa, emerging in late 2020 to early 2021).

Then, to gauge internal consistency of the time estimation procedure, we selected a subset of sample nodes, matching the distribution of node times and average number of children of the non-sample internal nodes. These selected sample nodes were then re-dated, and the inferred dates compared against their recorded collection dates. The results in Figure S9 demonstrate that although the inferred and true times are generally very close, there is a clear bias with re-estimated times being consistently older (by an average of 37 days). This is in line with the findings of other recent work^94^, and is likely to be a consequence of edges carrying more mutations than would be expected under a neutral molecular clock prior (either due to errors in tree-building or genuinely varying mutation rates), as well as inaccuracies in the recording of sample dates.

### S1.5 Origins of major VOCs

#### S1.5.1 Alpha

The B.1.1.7 (Alpha) variant emerged in the UK as the first major VOC. Its origin is well reconstructed in the ARG, with Figure S10 showing it to be a saltational lineage characterized by a long branch preceding its origin. Most of the characteristic Alpha mutations^95^ (20 out of 23) in the ARG occur immediately above the Alpha root. However, a characteristic three-base correlated mutation is instead associated with the parent node (labelled B.1.1) of the Alpha root, suggesting that this mutation was acquired prior to the emergence of Alpha. This parent node has an additional 5 children (leading to 12 samples), of which we display one (ERR4413600) with no associated mutations, meaning that it has an identical sequence to the inferred ancestor of Alpha. Below the root of Alpha, a single sample (ERR5178290) appears without the characteristic mutation C3267T (or any additional mutations). However, this is only a single sample and given that it also appears two months after the first Alpha samples we treat its placement with caution. Note that the list of mutations in ref^95^ includes a T26801C in the M gene which should no longer be considered characteristic of Alpha, as the Wuhan-Hu-1 reference at 26,801 has since been updated to C. In the ARG, a one-base non-coding deletion (position 28,271) is also associated with the Alpha root; although this is not in the original list of characteristic Alpha mutations, it has been suggested elsewhere as an Alpha-defining mutation^79^.

#### S1.5.2 Delta

The B.1.617.2 lineage (Delta) was first detected in India^96^. The subgraph in Figure S11 shows a mutation-rich branch leading to the Delta root, which contains 11 single nucleotide mutations and a 6-base deletion. The placement of this branch in the ARG is a result of long branch splitting of the lineage above B.1.617.

Delta belongs to the “B.1.617” clade, which also includes Kappa (B.1.617.1) and B.1.617.3^96^. Previous phylogenetic analyses suggest that Delta and B.1.617.3 are more closely related to each other than either is to Kappa^97,98^. In the ARG, however, Delta and Kappa appear as sibling lineages (see the subgraph in Figure S11). This reflects the closest approximation to the relationships suggested in the previous studies that could be inferred, given that no B.1.617.3 samples passed our QC filters and no seed B.1.617.3 samples were specified for ARG inference.

In the ARG, all six of the shared Delta-Kappa mutations (red) occur above the B.1-labelled common ancestor of Kappa (left branch) and Delta (right branch). All nine of the Kappa-specific mutations (blue) occur above the B.1.617-labelled node leading to Kappa samples. Seven out of nine Delta-specific mutations (purple) occur above the Delta root (the other two are above the MRCA of Kappa and Delta). We further examined the mutations associated with two major branches of Delta. Using a phylogenetic analysis, Stern et al.^99^ suggest that the Delta sublineages can be classified into distinct clades and identified the characteristic mutations of these clades. In the Nextstrain phylogeny shown in Stern et al.^99^, the first major split separates clades A to D from clade E; this topology is reflected in the ARG. All three of the clade A-D mutations (G15451A, C16466T, and a 6-base deletion at 28,248) occur on the uppermost branch of Delta (which corresponds to clades A to D). There are ten clade E mutations: five occur on the right branch of Delta (which corresponds to clade E) and three (C1191T, C5184T, and C28253T) occur above the Delta root. The placement of these mutations in the ARG generally reflects what is proposed in Stern et al.^99^.

There are four positions with multiple mutations (orange at position 1267, blue at 210, green at 20396, and red at 25276). Of these, the only clear topological change that could make the graph more parsimonious would be to attach the node above SRR12316669 one position higher up the tree, which would allow the placement of two recurrent red mutations (a C25276T above SRR12316669 and another above ERR4561562 / B.1.617) rather than one C25276T mutation followed by two reversions. The other recurrent mutations can only be made more parsimonious by topological rearrangements that would cause recurrence at other sites. We therefore believe that the inferred Delta origins in the sc2ts ARG are largely correct.

#### S1.5.3 Omicron BA.1 and BA.2

The B.1.1.529 lineage (Omicron) is believed to have emerged in southern Africa, and it is a saltational lineage with two highly successful sublineages, BA.1 and BA.2^100^. BA.1 and BA.2 are genetically distinct from each other and from their parental B.1.1.529 lineage^101^. In the subgraph shown in Figure S12, BA.1 and BA.2 differ from the non-Omicron B.1.1 ancestor (above the B.1.1.529-labelled MRCA of BA.1 and BA.2) by 27 mutations (including two deletions), reflecting its saltational origin. The ARG captures the split between BA.1 and BA.2 shown in previous phylogenetic analysis^100^. The 5-base deletion highlighted in green above the BA.2 root is an artefact of our naive deletion remapping procedure, which splits two 9-base deletions in BA.1 and BA.2 into three deletions (one in green at 11288, one in blue at 11283, and one in orange 11292; see Document S1.6). Also, there is uncertainty in the sequence alignments leading to ambiguity in the start position of these deletions (if these deletions occur at 11283, there would be a single deletion event before the BA.1/BA.2 split), as noted in the Pango designation issue^101^. We selected a BA.1 seed sample that led to the fewest mutations (see https://github.com/ jeromekelleher/sc2ts-paper/issues/264). This strategy resulted in five recurrent mutations highlighted in blue/orange/green/red/purple which are absent in the chosen seed but present in many of its inferred descendants. Inspection of the subgraph reveals that an alternative choice for a seed would be the later sample SRR17051902. Seeding with this sample would be likely to generate a topology in which those five recurrent mutations are shared between BA.1 and BA.2 and subsequently reverted in a number of the deep branching BA.1 lineages such as those to ERR7671333 and ERR7858953.

### S1.6 Analysis of deletion events

Although uncommon relative to single nucleotide mutations, deletions are an important component of SARS-CoV-2 evolution^102^, contributing to the emergence of VOCs by enhancing, e.g., efficiency for cell entry^156^ and potential for antibody escape^103^. Using the ARG, we identified major deletion events (which are propagated to many descendants) as well as recurrent deletions (which occur on at least two branches) in the ARG. In Table S1, we list the major deletion events in the ARG which are each inherited by at least 10,000 descendant samples; and in Table S2, we list the recurrent deletions in the ARG which are each observed in at least 10,000 (0.4%) samples. A deletion is said to be observed in a sample here if the gap character is present at the specified positions, and the immediately adjacent positions are not the gap character.

Most of the deletions in these tables occur in coding regions (except the 1-base deletions at 28,271) and do not cause frame shifts, involving 3, 6, or 9 sites (except the 4-base deletion and the two 5-base deletions, which are likely artefacts; see below). All the deletions in these tables (besides the artefactual cases) have been linked to the emergence of major VOCs (Alpha^95^, Delta^99^, Omicron B.1.1.529^101^, BA.1^101^, BA.2^101^, and BA.4^157^) and/or reported to be prevalent in previous studies of deletions^79^. All these major deletion events (again, besides the artefactual cases) are associated with the origins of major VOCs in the ARG (see the Alpha subgraph in Figure S10; the Delta subgraph in Figure S11; and the Omicron subgraph in Figure S12), except the 1-base deletion at 28,271 on node 1436808 and the 9-base deletion at 686 on the BA.4-labeled node 2698748 (Table S1). Most of these deletions have been identified as characteristic mutations of the major VOCs in the studies or analysis cited in Table S1. Notably, eight of the major deletion events occur in the N-terminal domain of the S1 unit of Spike (a known hotspot of recurrent deletions^103^ that plays a substantial role in antigenicity^158^), which are all highly recurrent in the ARG (Table S2). These include known recurrent deletions: the 6-base deletion at 21,765 and the 3-base deletion at 21,991, which are associated with increased efficiency in cell entry^105^ and antibody escape^103^, respectively. In addition, we found recurrent deletions in ORF8 and NSP3, which have been suggested to be indel hotspots^159^. Together, these findings demonstrate that the ARG captures genuine deletion events in SARS-CoV-2 evolution. A comprehensive evolutionary analysis of the deletion events in the ARG is beyond the scope of the current study; however, we believe that the ARG, as a resource, will enable and facilitate future studies to better understand the role of deletions in SARS-CoV-2 evolution (alongside recombination and single nucleotide mutations), which to date has been understudied in great part due to a lack of suitable data models, methods, and tools.

Detection of deletions and inference of deletion events remain challenging, especially when the dataset is enormous. Here we took a naive approach to map deletions onto the ARG, treating each site involved in a deletion as independent of the other sites. Consequently, some deletion events might be split up and appear as multiple smaller deletion events, which are likely artefactual. The three contiguous out-of-frame deletions in NSP6 (5-base at 11,283, 4-base at 11,288, and 5-base at 11,292) are examples of such artefacts (Table S1). In the sequence alignments, there are two 9-base deletions, one at 11,283 and the other at 11,288, which are found in 339,810 BA.1 samples and 843,291 BA.2 samples in the Viridian v04 dataset, respectively. Assuming that the alignments are correct, there should be two deletion events, but they were mapped as three deletion events as they had four overlapping sites from 11,288 to 11,292, which were inferred by parsimony as the 4-base deletion event at 11,288. Future improvements in deletion remapping should remove such artefactual deletions.

**Table S1:** Major deletion events in the ARG. The deletions caused by these events are inherited by at least 10,000 samples. The deletions are characterized by a site position in the genome (start) and the number of sites involved (length); they are annotated by the gene in which they occur (non-coding otherwise). The IDs of the nodes in the ARG above which the deletions occur and the Pango lineages assigned to these nodes are shown. Frequency is the percentage of samples inheriting a deletion. If a deletion has been identified as a characteristic mutation of a VOC by a study (or an analysis), a reference to the study (or the analysis) is provided. Abbreviations: NSP, non-structural protein; ORF, open reading frame; S, Spike; S1-NTD, N-terminal domain in Spike S1 subunit.

| Start | Length | Region | Node | Pango lineage | Frequency (%) | Ref |
| --- | --- | --- | --- | --- | --- | --- |
| 28271 | 1 | non-coding | 1436808 | B.1 | 45.06 |  |
| 28248 | 6 | ORF8 | 220186 | B.1.617.2 (Delta) | 44.91 | 99 |
| 22029 | 6 | S / S1-NTD | 200039 | B.1.617.2 (Delta) | 44.82 | 99 |
| 11288 | 4 | ORF1ab / NSP6 | 1436802 | B.1.1.529 (Omicron) | 35.12 |  |
| 28362 | 9 | N | 1436802 | B.1.1.529 (Omicron) | 35.12 | 101 |
| 21633 | 9 | S / S1-NTD | 822854 | BA.2 (Omicron) | 21.44 | 101 |
| 11292 | 5 | ORF1ab / NSP6 | 822854 | BA.2 (Omicron) | 21.43 |  |
| 6513 | 3 | ORF1ab / NSP3 | 851246 | BA.1 (Omicron) | 13.73 | 101 |
| 11283 | 5 | ORF1ab / NSP6 | 851246 | BA.1 (Omicron) | 13.69 |  |
| 21765 | 6 | S / S1-NTD | 851246 | BA.1 (Omicron) | 13.68 | 101 |
| 21987 | 9 | S / S1-NTD | 851246 | BA.1 (Omicron) | 13.68 | 101 |
| 22194 | 3 | S / S1-NTD | 851246 | BA.1 (Omicron) | 12.60 | 101 |
| 28271 | 1 | non-coding | 86456 | B.1.1.7 (Alpha) | 11.70 |  |
| 11288 | 9 | ORF1ab / NSP6 | 86456 | B.1.1.7 (Alpha) | 11.70 | 95 |
| 21991 | 3 | S / S1-NTD | 86456 | B.1.1.7 (Alpha) | 11.64 | 95 |
| 21765 | 6 | S / S1-NTD | 86456 | B.1.1.7 (Alpha) | 11.70 | 95 |
| 21765 | 6 | S / S1-NTD | 1265302 | BA.4 (Omicron) | 7.67 | 157 |
| 686 | 9 | ORF1ab / NSP1 | 2698748 | BA.4 (Omicron) | 0.94 |  |

### S1.7 Pango X lineage events

Most of the Pango X lineages had straightforward origination events as defined in Section S1.3 and are represented in Table 1. Three lineages were treated specially, corresponding to 23 omitted samples in total.

XM has 4 separate origins, 3 singletons and one dominant node with 26 descendants. We include the dominant node in Table 1 and omit the singletons. See Document S1.9.9 for further details. XAC has 4 separate origins, and has been omitted from Table 1. However, as Figure 2B shows, these are all descending from a single BA.2 classified node descending from the putative Xx event. Similarly, XAD has two independent origins descending from the Xx event, and we omit it for simplicity. See Document S1.9.16 for more details on the Xx event.

**Table S2:** Recurrent deletions in the ARG. These deletions occur on at least two branches in the ARG and are found in the alignments of at least 10,000 (0.4%) samples in the Viridian v04 dataset. For each deletion, the number of occurrences in the ARG and the number of samples where it is observed are shown. For further explanation, see the caption of Table S1. Abbreviations: NSP, non-structural protein; ORF, open reading frame; S, Spike; S1-NTD, N-terminal domain in Spike S1 subunit.

| Start | Length | Region | Occurrences | Samples |
| --- | --- | --- | --- | --- |
| 686 | 9 | ORF1ab / NSP1 | 6436 | 43354 |
| 21991 | 3 | S / S1-NTD | 2088 | 309629 |
| 21765 | 6 | S / S1-NTD | 482 | 820194 |
| 22029 | 6 | S / S1-NTD | 275 | 1107941 |
| 22194 | 3 | S / S1-NTD | 200 | 310349 |
| 6513 | 3 | ORF1ab / NSP3 | 181 | 341380 |
| 21987 | 9 | S / S1-NTD | 70 | 338633 |
| 28248 | 6 | ORF8 | 28 | 1113392 |
| 28271 | 1 | non-coding | 24 | 1331409 |
| 11288 | 9 | ORF1ab / NSP6 | 19 | 843291 |
| 11283 | 9 | ORF1ab / NSP6 | 13 | 339810 |
| 28362 | 9 | N | 7 | 850483 |
| 21633 | 9 | S / S1-NTD | 7 | 531973 |

**Table S3:** Concordance among methods in characterizing Pango X lineages. These comparisons focus on the Pango X lineages associated with the Type 1 recombination events (Table 1) For each method, we calculated the number of Pango X lineages which have inferred parent Pango lineages and breakpoint intervals concordant with those proposed by the community. The denominators indicate the number of Pango X lineages for which detection results are available from the original studies^40,41^. For the Pango X lineages with non-overlapping breakpoint intervals, the distances between the inferred breakpoint intervals and those proposed by the community are shown (median and range).

| Method | Parent lineages | Breakpoint intervals |  |
| --- | --- | --- | --- |
|  | Concordant (%) | Concordant (%) | Distance (bases) |
| Sc2ts | 14 / 16 (87.5) | 14 / 16 (87.5) | 540.5 (315, 766) |
| RecombinHunt-GISAID | 13 / 15 (86.7) | 10 / 15 (66.7) | 764.0 (4, 1202) |
| RecombinHunt-Nextstrain | 12 / 15 (80.0) | 11 / 15 (73.3) | 1160.0 (559, 2785) |
| CovRecomb | 7 / 8 (87.5) | 4 / 8 (50.0) | 1562.5 (217, 13208) |

**Table S4:** Validation of sc2ts recombinants by 3SEQ, CovRecomb and rebar over all 855 recombinants and the 354 passing and 501 failing QC.

| Method | Total | QC Pass | QC Fail |
| --- | --- | --- | --- |
| 3SEQ | 484 | 338 | 146 |
| CovRecomb | 92 | 90 | 2 |
| rebar | 213 | 203 | 10 |

In cases where nested samples not assigned to the focal Pango lineage are either from different Pango X lineages or given a generic BA.2 assignment. Of these 59 generic assignments 6 had conflicts in their UShER placements and could have been assigned an X lineage (XJ: 3, XU: 2, XN: 1), and 25 were cases in which Scorpio^143^ failed to find a specific lineage (accounting for 7% of the 357 “Omicron (Unassigned)“ Scorpio calls across 2,747,985 alignments).

See notebook tab pango x events.ipynb for details.

### S1.8 Analysis of the Jackson et al. recombinants

Jackson et al.^29^ conducted a detailed analysis of early recombinants involving B.1.1.7 (Alpha), including the first designated Pango X lineage, XA. The authors first performed a targeted search for candidate recombinants by scanning sampled sequences for sequence motifs associated with Alpha, followed by an analysis to reduce the list of plausible recombinants displaying signs of onward transmission. Here we compared the sc2ts detection results of these recombinants with the detection results reported by Jackson et al.

Jackson et al. described four groups of recombinants (named A to D) and four singleton recombinants. Of the 16 samples analysed by Jackson et al., 12 were included in the ARG; the remaining 4 were excluded because the sequences did not pass the QC filtering criteria or because of a high HMM cost. Groups A-D are summarized in Table S5 and show excellent concordance in terms of the parental Pango lineages and breakpoint intervals. All four groups are directly associated with strongly supported recombination events in the ARG in terms of the number of averted mutations (A: 16, B: 6, C: 30, D: 18). In the ARG, the singleton CAMC-CBA018 has the parents B.1.177+B.1.1.7 and the breakpoint interval 16,177–20,133, and is strongly supported by 14 averted mutations. In the Jackson et al. results, this singleton is almost identical with the same parent lineages and a breakpoint interval of 11,395—21,993. In the ARG, the singleton MILK-103C712 has the parents B.1.177.16+B.1.1.7 and a breakpoint interval of 24915–27972, again strongly supported by 11 averted mutations. In the Jackson et al. results, this lineage has the same parents, with the breakpoint interval 26,800–27,974. In the ARG, the singleton QEUH-1067DEF has the parents B.1.1.7+B.1.177.9 and a breakpoint interval of 7,729–10,870, supported by 11 averted mutations. In the Jackson et al. results, this lineage has the same parents, with the breakpoint interval 6,953–10,872.

See notebook analysis jackson recombinants.ipynb for details.

### S1.9 Additional analyses of the Pango X lineages

In this section we provide some additional analysis on Pango X lineages in the ARG. These analyses are intended to be read in conjunction with the subgraph illustrations in Document S2.

#### S1.9.1 XA

XA traces to a well supported recombination node, with no reversions. Note that the “causal” sample ERR5308556 (which triggers the initial recombination event) is identical to the recombination node, and therefore lacks C8090T, which is one of the XA consensus mutations (identified as those shared by over 90% of XA samples). As mentioned elsewhere, the sc2ts detection results in terms of the Pango lineages of the parents and the breakpoint intervals agree with the results reported by Jackson et al. (Table S5). Two of the Jackson et al. XA samples are present in the ARG (ERR5308556 and ERR5414941); these samples are specifically marked in Figure 2, which is a summary of this subgraph.

**Table S5:** Comparison of recombination breakpoint intervals and parent lineages for Groups A-D reported by Jackson et al. with the corresponding recombination events in the sc2ts ARG. The second column gives the number of sequences in the group. See the text for details of the groups and the sequences included. limited to the samples considered by Jackson et al. The breakpoint coordinates in Table 1 of Jackson et al. have been altered as follows: we subtract one to the left coordinates and add one to the right coordinates to correspond to the tskit definition of inheritance on either side of a breakpoint, and add one to the right coordinate to make the intervals right-exclusive.

| Group | Sequences | Method | Breakpoint interval | Parent lineages |
| --- | --- | --- | --- | --- |
| A (XA) | 2 | Jackson | 21,254–21,767 | B.1.177+B.1.1.7 |
|  |  | sc2ts | 20,411–21,765 | B.1.177.18+B.1.1.7 |
| B | 2 | Jackson | 6,527–6,956 | B.1.36.28+B.1.1.7 |
|  |  | sc2ts | 6,529–6,954 | B.1.36.28+B.1.1.7 |
| C | 3 | Jackson | 24,913–28,653 | B.1.1.7+B.1.221 |
|  |  | sc2ts | 25,997–27,972 | B.1.1.7+B.1.221.1 |
| D | 3 | Jackson | 21,574–23,065 | B.1.36.17+B.1.1.7 |
|  |  | sc2ts | 22,445–23,063 | B.1.36.39+B.1.1.7 |

#### S1.9.2 XB

The displayed subgraph shows a small selection of the 192 XB samples. XB was originally defined as a recombinant between the closely related lineages B.1.634 (left parent) and B.1.631 (right parent) from North and Central America^32^. However, these parent lineages were not added to the ARG due to high HMM costs. Together with other lineages such as B.1.627 (sister to B.1.631^32^), they are likely to attach somewhere along the long branch in the subgraph visualisation (with *>* 30 mutations). If samples from these parental lineages were available for attachment, then XB would likely be detected as a recombinant by sc2ts.

See github:sc2ts-paper/discussions/780 for further details.

#### S1.9.3 XC

The five XC samples descend from a well supported recombination node. The two parents that are both identical to actual samples (ERR4908740 on the left and DRR321274 on the right). There is one reversion (G21987A) on the left side of the breakpoint, but the site 21987 was masked during ARG inference because it was identified to be experiencing frequent mutation in a preliminary ARG.

#### S1.9.4 XE/XH

The XE samples descend from a well supported recombination node, with one mutation and one reversion (T22792C). This reversion was mapped onto the ARG as a part of postprocessing, and could be placed more parsimoniously by a single C22792T that groups together XH and the lineage leading to the BA.2 recombination nodes displayed on the lower left. Nevertheless, the exact arrangement of the mutations at position 22792 should not affect the proposed breakpoint, which is very distant, at position 11283. This breakpoint coincides with a repeated homopolymeric sequence involving 5-base deletions, so that alignment errors could have a small effect on the breakpoint position. Note that the two lower recombination nodes are both well supported, the youngest being postulated as the origin of XAF (see below).

See github:sc2ts-paper/discussions/633 for further details.

#### S1.9.5 XF

XF is a well-supported recombinant between Delta and Omicron BA.1, with the breakpoint on the far left of the genome. All XF samples are the unique descendants of a well supported recombination node, with both parent nodes (ERR6193241: AY.4 and ERR8042040: BA.1) and the single child node (ERR8076129) being samples.

#### S1.9.6 XG

The three XG samples descend from a reconstructed internal node, which in turn descends from a well supported recombination node. The left parent is a sample (SRR20179863), and the right parent is a reconstructed internal node assigned to Omicron BA.2, which is supported by a sample (ERR9124067) that is identical to it.

#### S1.9.7 XJ

All the XJ samples descend from a well supported recombination node. There are 14 samples (including e.g. ERR8683190) which sc2ts identified as descendants of the same recombination node but are labelled BA.2 by pangolin (displayed on the left), and which do not possess the XJ consensus mutations C22792T and C27945T. These BA.2 samples could be worthy of further investigation.

#### S1.9.8 XL

All the XL samples descend from a well supported recombination node, with three child samples, which are identical to it. The left parent is a sample (ERR7989424), and the right parent is a reconstructed internal node assigned to BA.2.5 with a large number of descendant samples.

#### S1.9.9 XM/XAL

XM is a recombinant of BA.1 (left parent) and BA.2 (right parent). Sc2ts identifies multiple recombinant origins of XM. One of these origins leads to a dominant clade (node 1003220 at position 21,595), which also contains the only three XAL samples.

Independent evidence for multiple XM origins comes from consideration of a 9-base deletion in the Spike gene at position 21633, which is characteristic of BA.2 but not BA.1 https://github.com/cov-lineages/pango-designation/issues/361. Most of the XM-labeled samples in ARG have this deletion, but three (SRR19024311, ERR9447529, and SRR18775874) do not, suggesting that XM samples trace to at least two recombination events, one to the right of the deletion (which leads to the majority of the XM samples), and one to the right. In the subgraph, it can be seen that one of the samples, SRR18775874, has been (presumably wrongly) placed within the main XM deletion-containing clade. As sc2ts does not use deletions for inference, deletion remapping (STAR Methods) has reverted the deletion in this case, creating many insertions on the branch above the sample (magenta). A better use of deletions in sc2ts would likely result in placing this sample within one of the two XM clades which do not have the deletion.

See github:sc2ts-paper/discussions/656 for further details.

#### S1.9.10 XN/XAU

In the sc2ts ARG, the initial XN sample (ERR8986821) is well explained as a non-recombinant, copying from an inferred internal node assigned to BA.2 with only one mutation. This internal node is directly supported by the sample ERR9502469. All the other XN samples descend from ERR8986821. A small clade containing XAU is the sister to XN, and there are a series of samples (ERR9453882, ERR9502469, and ERR9823912) which support the gradual emergence of both groups from a BA.2 ancestral lineage via accumulation of single mutations.

The evidence supporting the original designation of XN and XAU as recombinants may come from inaccurate or problematic data sources. In the case of XN, support for a recombinant BA.1+BA.2 origin is from three mutations that are claimed to be BA-1-specific: C241T, A2832G, and C3037T (https://github.com/cov-lineages/pango-designation/issues/480). However, the BA.1/BA.2 split (https://github.com/cov-lineages/pango-designation/issues/361) does not list C241T and C3037T as BA.1-specific, meaning that these two mutations are not supportive of recombination. Moreover, although the remaining mutation claimed to support a recombinant origin, A2832G, is typical of BA.1 and not BA.2, it is seen in several BA.2 samples (e.g. ERR9453882), supporting the idea that it has independently arisen within BA.2. Hence, none of the sites used to designate XN as a recombinant reliably do so.

In the case of XAU, a recombinant origin was proposed on the basis of the same three sites, plus C2470T (https://github.com/cov-lineages/pango-designation/issues/894). However, C2470T is not BA.1-specific (https://github.com/cov-lineages/pango-designation/issues/ 361). Moreover, some BA.2-designated samples such as ERR9823912 also have C2470T, so we do not believe this to be a recombinant-supporting mutation either.

See github:sc2ts-paper/discussions/781 for further details on XN. See github:sc2ts-paper/discussions/928 for further details on XAU.

#### S1.9.11 XP

This is a case where the Pango X lineage is designated on the basis of two loci on the far right of the genome: a single nucleotide mutation (A29510C) and a rare, large deletion at position 29734 in the S2M step loop (https://github.com/cov-lineages/pango-designation/issues/ 481), which is not common enough to map in sc2ts during postprocessing and hence not shown here. Therefore, unless this deletion were given more weight than a single locus, sc2ts would not identify XP as a recombinant, even if deletions were used during ARG inference. If this deletion were recurrent, the evidence from sc2ts suggests that XP might not be a recombinant, because it is only one mutation different from the sample ERR8628084.

See github:sc2ts-paper/discussions/775 for further details.

#### S1.9.12 XQ/XR/XU/XAA/XAG/XAM

In the ARG, XR, XU, XAA, XAG, and XAM are nested under the XQ recombinant lineage. These lineages were designated around the same time, approximately within a five-month period (Mar. to Jul., 2022). The breakpoint intervals of these lineages, as suggested in the Pango designation issues, all occur at the 5’ end of the genome, within a ∼5 kb segment (∼4,300 to ∼9,400): XQ (4321-5387), XAG (6515-8394), XU (6517-9345), XAM (8087-9193), and XAA (8392-9345).

Upon closer inspection, the right-hand branch above the recombination node appears problematic, as it displays three reversion mutations, which revert mutations at positions 22792, 24023, and 25624 in the ancestral lineage just above. These sites were excluded during ARG inference, and remapped onto the ARG by parsimony during ARG postprocessing. If these three sites were used during primary ARG inference, the right parent of the recombination node may have been chosen to be the ancestral BA.2 sample on the right-hand path with 18 child nodes. A more parsimonious reconstruction would then have the XQ clade as a sibling to a group of (monophyletic) XAA, XAM, XAG, and XR lineages. This would remove between 4 and 5 reversion mutations.

See github:sc2ts-paper/discussions/655 for further details.

#### S1.9.13 XS

In the ARG, the XS samples appear to have originated from two sequential recombination events (separated by a single branch). However, the later event is likely an artifact, because the XS sample (SRR18343220) associated with this event failed detection-level QC (i.e., the minimum net number of supporting loci; see the lower copying pattern). In addition, this sample was sequenced using the Ion AmpliSeq V1 kit, which was also used to sequence eight other samples descending from the same event (clustered due to a shared sequencing error due to AmpliSeq). These eight samples would likely attach to the nodes descending from the first recombination event. We excluded this likely artefactual event from further analysis and consider XS to have emerged from a single recombinant origin.

See github:sc2ts-paper/discussions/782 for further details.

#### S1.9.14 XW

All the XW samples descend from a well supported recombination node. Its left parent is a BA.1.1.15-labelled sample (ERR8069658), and its right parent is a BA.2-labelled sample (ERR9086669).

#### S1.9.15 XY

All the XY samples descend from a well supported recombination node, through a shared reconstructed internal node. Its left parent is a reconstructed internal node assigned to BA.1.1, which is supported by a sample (ERR8627048) identical to it. Its right parent is a BA.2-labelled sample (ERR9399983). There are a pair of (orange) reversion mutations within the XY clade at position 23948: these were not used for inference but remapped using parsimony during postprocessing, and indicate that the topology within the XY clade could potentially be improved by alternative phylogenetic reconstruction methods.

#### S1.9.16 Xx: XZ/XAC/XAD/XAE/XAP

This nested set of Pango X lineages descends from a single recombination node in the ARG, which we tentatively refer to as “Xx”. Note that all were designated around the same time, with their samples collected roughly within a three-month period (Mar., 2022 to Jul., 2022). Several samples labelled as BA.2 by Pangolin also lie within this recombinant clade, including two that are identical to the proposed originating recombinant (ERR8146303 and ERR8163061).

Exact relationships within the Xx clade are somewhat tentative. For example, there are three highly mutable sites (22792, 28877, and 28878) which were not used during inference, but were remapped onto the ARG during postprocessing. The inferred topology places these sites as recurrent (coloured orange, grey, and faded yellow), but a more parsimonious arrangement would, for example, group the two XAD samples as siblings (removing the need to postulate mutations A1250G, A28877T, and G28878C to be recurrent). It is possible that a more parsimonious arrangement would also reduce the recurrent mutations and reversions seen at sites 22,792 (four orange mutations) and 26,060 (two parallel brown mutations), but we do not show such a reconstruction, which is complicated by the presence of additional (non-displayed) samples descending from internal nodes in the plotted subgraph.

For XAC, in particular, the Pango designation issue (https://github.com/cov-lineages/ pango-designation/issues/590) suggests it to be a recombinant with two breakpoints: BA.2 on the left, BA.1 in the middle, and a switch back to a very small region of BA.2 on the right. This contrasts with the one-breakpoint BA.2+BA.1 origin inferred in the ARG. However, only two loci support the right hand BA.2 assignment: A29510C and a large deletion (deletions are treated as missing data during primary ARG inference). Sc2ts does not count this as enough evidence to insert an additional breakpoint.

See github:sc2ts-paper/discussions/631 for further details.

#### S1.9.17 XAF

Of the 21 XAF samples in the Viridian v04 dataset, only one XAF sample passed our QC filters and was added to the ARG. However, we find a small recombinant BA1.1/BA.2 clade of 37 samples, associated with the BA.2-labelled sample ERR9755541, within which the single XAF sample is nested. The inferred parents and breakpoint at 10,447 aligns with that proposed for XAF (https://github.com/cov-lineages/pango-designation/issues/676). Inspection of the omitted XAF samples with high HMM cost shows that they would attach as close sisters to the included XAF sample, albeit on branches with many new mutations.

The mutation-rich branch in the subgraph with a 5-base deletion (11283-11287) and insertion (11292-11296) may be due to a simple deletion alignment error.

See github:sc2ts-paper/discussions/638 for further details.

#### S1.9.18 XAJ

In the Pango designation issue (https://github.com/cov-lineages/pango-designation/issue 826), XAJ is suggested to be a recombinant between BA.2.12.1 (left parent) and BA.4 (right parent). However, XAJ is not inferred to be a recombinant lineage in the ARG. From the subgraph, which shows the 18 XAJ samples plus a small number of relevant BA.4 samples, it can be seen that 9 sites leading to XAJ indeed recur in the lineage to the closest BA.4 sample (SRR18848050), which strongly indicates that this is a recombinant that is missed by sc2ts. The reason that is is not a recombinant in the ARG is that 6 sites represent a 6bp deletion and so are not used for inference. The remaining three sites (T22917G, T23018G, and G23040A) are not enough to exceed our k=4 threshold for recombinant designation. Including deletions in inference would certainly lead to XAJ being detected as a recombinant, and with a lower threshold of k=3 the distribution of mutations on the ARG topology suggests that it is is even possible that XAJ is a 2-breakpoint recombinant with the region from 21765-23535 coming from the BA.4 labelled sample SRR18848050.

See github:sc2ts-paper/discussions/634 for further details.

#### S1.9.19 XAN/XAV

XAN and XAV are shown in this subgraph, as they share some mutations. In the Pango designation issues (https://github.com/cov-lineages/pango-designation/issues/771; https://github.com/cov-lineages/pango-designation/issues/911), XAN and XAV are suggested to be a recombinant between BA.2 (left parent) and BA.5 (right parent). Both these Pango lineages were detected as non-recombinants by sc2ts, because the number of supporting loci fell below our threshold of detection. In the ARG, we found only one mutation (C9866T) supporting the suggested BA.2 parent for both XAN and XAV.

See github:sc2ts-paper/discussions/783 for further details on XAN. See github:sc2ts-paper/discussions/785 for further details on XAV.

#### S1.9.20 XAS

XAS was proposed as a recombination between BA.5 (left parent) and BA.2 (right parent). However, we find that this is only two mutations different from e.g. ERR9929846, which is a BA.4 sample. In fact, XAS only differs from the BA.4 root by a single mutation, C27945T, which was identified in the original Pango designation issue (https://github.com/cov-lineages/ pango-designation/issues/882) as an important ORF truncation mutation which is also present in a few BA.2.65 samples. It is possible that C27945T was acquired by recombination, but this is clearly below our threshold of detection.

See github:sc2ts-paper/discussions/784 for further details.

#### S1.9.21 XAZ

In the Pango designation issue (https://github.com/cov-lineages/pango-designation/issue 797), XAZ is suggested as a two-breakpoint recombinant between BA.2.5 and BA.5 (the middle parent), with three sites supporting the BA.2.5 parent on the left end (C2232T, C3317T, and T3358C). In the ARG, this is very close to being a recombinant of BA.2.5 (ERR9615610) on the left side of the breakpoint, as not only are the three mutations present in this sample, but there is another site which supports recombination C1912T, which is not reported in the Pango designation issue). XAZ has C44T which is lacking in all BA.2.5 samples, leaving only three net supporting loci for a putative recombinant. However, position 44 is often clipped, so this is not strong evidence against a recombinant, and therefore XAZ is a marginal case of recombination. We found no evidence for the third segment inherited from a BA2.5 parent on the right end. In the ARG, all the XAZ samples form a clade. Here we display only the first 20 XAZ samples.

See github:sc2ts-paper/discussions/786 for further details.

#### S1.9.22 XBB

Most of the XBB samples descend from a well supported recombination node. In the Pango designation issue (https://github.com/cov-lineages/pango-designation/issues/1058), the Pango lineages of the suggested left and right parents are BJ.1 and BM.1.1.1, respectively. In the ARG, the Pango lineage of the right parent is identical, but that of the left parent is a related ancestral Pango lineage (BA.2.10 instead of BA.2.10.1.1 = BJ.1). This is likely due to the absence of BJ.1 in the ARG: only four BJ.1 samples exist in the Viridian v04 dataset, a single one passed sequence-level QC, and that was then rejected due to high HMM cost. Note that BJ.1 differs from BA.2.10 by over 20 mutations (https://github.com/cov-lineages/ pango-designation/issues/915), explaining the large number of (gold) de novo mutations in the XBB copying pattern. Had BJ.1 samples been present in the ARG, the left branch of the

XBB-associated recombination node may have attached to a BJ.1-labelled node, and the origin of XBB would required fewer mutations.

See github:sc2ts-paper/discussions/929 for further details.

#### S1.9.23 XBD

All the XBD samples descend from a well supported recombination node. Its left and right parents are reconstructed internal nodes, both of which are directly supported by samples (SRR21382561 and ERR9825609, respectively) that are identical to them.

#### S1.9.24 XBE

In the Pango designation issue (https://github.com/cov-lineages/pango-designation/issue 1246), XBE was suggested as a recombinant between BA.5.2 (left parent; most of the genome) and BE.4 (right parent; somewhere from 23,609 onwards). However, sc2ts treats the 65 XBE samples as non-recombinant descendants of BA.5.2 because the number of supporting sites fell below our threshold of detection. We have included the BE.4 lineage on the subgraph, revealing two mutations (C27800T, and G28681T) which occur in parallel above the root of the XBE clade and along the path ascending from the displayed BE.4 sample, suggesting that if we used k=2 as the recombinant threshold, sc2ts would indeed detect XBE as a recombinant.

See github:sc2ts-paper/discussions/930 for further details.

#### S1.9.25 XBF

All the XBF samples descend from a well supported recombination node, with three mutations on the left of the breakpoint. Its left parent is a sample assigned to BA.5.2.1 (SRR20305553). Its right parent is a reconstructed internal node that is supported by a CJ.1-labelled sample (SRR21948068), which differs by three mutations.

#### S1.9.26 XBG

All the XBG samples descend from a well supported recombination node, with two mutations, one on each side of the breakpoint. Both its left and right parents are samples (SRR19920084 and ERR9852351, respectively).

#### S1.9.27 XBH

All the XBH samples descend from a well supported recombination node, despite two reversions (T222C and T12789C, respectively), which occur at excluded sites and were remapped by parsimony during postprocessing. If these two sites had been used during ARG inference, the branch attached to BA.2.1 would have been attached to the BA.2.5 shown on the top left instead, leading to a more parsimonious local topology with two mutations on the left branch. This would also remove the two blue lines on the left and side of the copying pattern.

#### S1.9.28 XBQ/XBK/XBK.1

XBK and XBQ are siblings, and neither appear to be recombinants in the ARG, as they are closely related to the non-recombinant node that originates CJ.1.3. At the time of designating XBK a hypothesis was made that a missing common ancestor could exist (https://github.com/cov-lineages/pango-designation/issues/1440#issuecomment-1355264973), making XBK non recombinant. Exactly such a missing ancestor is found as the upper internal node labelled CJ.1 in the ARG, constructed by sc2ts as part of its tree building steps.

**Table S6:** Sc2ts detection results for Pango X lineages not added to the ARG. The number of samples in Viridian v04 dataset assigned to these Pango X lineages, the number of samples that passed sequence-level quality control, and the number of recombination breakpoints in the Viterbi solutions of the samples are shown.

| Pango | Samples in Viridian | Samples passing QC | Breakpoints |
| --- | --- | --- | --- |
| XBJ | 2 | 1 | 1 |
| XBP | 8 | 2 | 1 |
| XBS | 19 | 15 | 1 |
| XBW | 1 | 1 | 1 |
| XCA | 11 | 4 | 1 |
| XAK | 2 | 2 | 1 |
| XAY | 15 | 11 | 6 |
| XBC | 36 | 21 | 3 |
| XBL | 2 | 2 | 0 |
| XBT | 1 | 1 | 2 |

See github:sc2ts-paper/discussions/931 for further details on XBQ. See github:sc2ts-paper/discussions/932 for further details on XBK.

#### S1.9.29 XBM

All the XBM samples descend from a well supported recombination node. Its left parent is a reconstructed internal node that is directly supported by a BA.2.76-labelled sample identical to it. Its right parent is a BF.3-labelled sample.

#### S1.9.30 XBR

This one XBR sample descends from a well supported recombination node. Its left parent is a BN.3.1-labeled sample, and its right parent is a reconstructed internal node that is directly supported by a BQ.1.25-labeled sample identical to it.

### S1.10 Pango X lineages not added to the ARG

Ten Pango X lineages which are present in the Viridian v04 dataset were not integrated into the ARG due to high HMM costs and/or insufficient sample support, but their samples were matched against the ARG during inference. Table S6 summarizes the HMM matching results of these Pango X lineages. All were identified as recombinants by sc2ts, except XBL. (For XBL, the evidence for recombination proposed in the Pango designation issue^160^ involves only three informative sites supporting the suggested rightmost parent, and therefore this likely fell below our threshold of detection.) Importantly, these results demonstrate that sc2ts can detect recombinants with multiple breakpoints (XAY, XBC, and XBT). XBC and XBT are relatively clean examples of multiple-breakpoint recombinants, with the Pango lineages of the parents and the breakpoint intervals concordant with those suggested in the Pango designation issues, but XAY is more complex, possibly involving up to six breakpoints as per sc2ts.

**Table S7:** Distribution of numbers of descendants for UShER+RIPPLES events and sc2ts. The first column shows the number of descendants (binned for larger values) and the others show the corresponding number of events.

| Count | RIPPLES p3 | RIPPLES p4 | sc2ts |
| --- | --- | --- | --- |
| 1 | 2616 | 757 | 525 |
| 2 | 575 | 178 | 124 |
| 3 | 232 | 65 | 47 |
| 4 | 124 | 38 | 32 |
| 5 | 74 | 14 | 17 |
| 5-100 | 437 | 108 | 96 |
| 100-1000 | 32 | 6 | 8 |
| 1000-10k | 15 | 6 | 4 |
| 10k-100k | 2 | 0 | 0 |
| 100k-1m | 4 | 2 | 2 |

### S1.11 Detailed RIPPLES analysis

We ran RIPPLES and aligned events with the sc2ts ARG as described in STAR Methods. We begin by describing the overall composition of events output by RIPPLES at the two parameter values used.

At *p* = 4, RIPPLES returned a total of 1174 events, and 610 pass 3SEQ validation (51%). 1052 events (89.6%) were in close clade agreement with the sc2ts ARG, and 468 (40%) corresponded to recombination events in the ARG (Figure S18A). We investigated the 701 RIPPLES events that could be aligned with the ARG but were not close to a recombination node to determine why they were not found by sc2ts. For 408 of these RIPPLES identified recombination events, one or more parents were *younger* than the recombinant sequence based on metadata, and therefore impossible under the strict time ordering enforced by sc2ts. Secondly, of the 179 remaining events with matching clades and in which the sc2ts node has more than 4 mutations, of the total 1,167 mutations, 452 of these occurred at sites that were remapped during post-hoc parsimony (and thus the mutations were not observed by the LS HMM during primary inference). The remaining “missing” events are likely attributable to the finer details of the differences in tree building approaches between UShER and sc2ts.

At *p* = 3, RIPPLES returned a total of 4111 events, and 2718 pass 3SEQ validation (66%). 3755 events (91.3%) were in close clade agreement with the sc2ts ARG, and 595 (14%) corresponded to recombination events in the ARG (Figure S18B). We did not investigate the *p* = 3 events any further as the additional events detected at the lower threshold are not expected to be detected by sc2ts at the parameter value used (*k* = 4).

Next we considered the overall distribution of the numbers of descendants of the events returned by sc2ts and RIPPLES (Table S7). Overall, the distributions were similar, with the majority having very few descendants and being dominated by singletons. The events subtending *>* 100,000 samples in all three cases are noteworthy (see Document S1.12 for analysis of the two sc2ts events).

We next examined the RIPPLES events referencing Pango X lineages. Table S8 shows a summary of all recombination events associated with Pango X lineages detected by RIPPLES at *p* = 3 and their relationship with the sc2ts ARG. Of the 16 Class I events (X lineages unequivocally associated with a recombination node in the sc2ts ARG) in Table 1, 8 have corresponding recombination events (4, at *p* = 4). Of the 5 Type II events (recombination nodes with more complex assocation with X lineages), two have close matches. XBM matches exactly (including 2 descending BF.3 samples), and XE/XH is similar (but without including the 2 XH samples in the clade). UShER and sc2ts agree on creating two separate XAM clades, but while sc2ts makes these both descendants of the Xx event, RIPPLES creates two independent recombinations. Similarly, XAC and XAE have exactly matching clades, but RIPPLES infers two independent recombination events while sc2ts groups them both under the Xx event. XB is notable as a likely sc2ts false negative that is identified by RIPPLES at both *p* = 3 and *p* = 4. The remaining events associated with X lineages output by RIPPLES cannot be easily mapped to the sc2ts ARG as one or more descendants are distantly placed. XAJ is an outlier for sc2ts, being the Pango X lineage identified as a non-recombinant with the largest number of mutations (Table 1), and a likely false negative. The remaining 4 events are mixtures of multiple lineages that are difficult to reconcile with either the other events in the table or the sc2ts ARG.

**Table S8:** RIPPLES events associated with Pango X lineages. Each row is a recombination event identified by RIPPLEs (at *p* = 3) in which Pango X lineages appear, along with the corresponding sc2ts event from Table 1, where applicable, grouped by class. Events also present at *p* = 4 are indicated. Clades are matched by finding the most recent common ancestor of the samples associated with each UShER+RIPPLES event in the sc2ts ARG. The clade differences indicates the closeness of the match.

| sc2ts event | in $p = 4$ | UShER desc | sc2ts class | clade diff | descendants |
| --- | --- | --- | --- | --- | --- |
| Equivalent Sc2ts and RIPPLES recombination events |  |  |  |  |  |
| XC | ✓ | 5 | I | 0 | XC:5 |
| XBR | ✓ | 1 | I | 0 | XBR:1 |
| XS | ✓ | 17 | I | 0 | XS:17 |
| XA | ✓ | 39 | I | 0 | XA:39 |
| XY |  | 23 | I | 0 | XY:23 |
| XW |  | 32 | I | 0 | XW:32 |
| XL |  | 64 | I | 0 | XL:64 |
| XBB |  | 6452 | I | 1 | XBB:6452 |
| RIPPLES recombination event close to sc2ts recombination event |  |  |  |  |  |
| XBM | ✓ | 12 | II | 0 | XBM:10, BF.3:2 |
| XM | ✓ | 8 | II | 0 | XM:4, BA.2:4 |
| Xx | ✓ | 18 | II | 0 | XAC:18 |
| Xx |  | 9 | II | 0 | XAE:9 |
| XQ |  | 1 | II | 0 | XAM:1 |
| XQ |  | 21 | II | 0 | XAM:21 |
| XE/XH |  | 1113 | II | 3 | XE:1113 |
| RIPPLES recombination event equivalent to sc2ts non-recombination event |  |  |  |  |  |
| XB | ✓ | 411 | IV | 0 | XB:192, B.1:219 |
| RIPPLES recombination event not comparable to sc2ts |  |  |  |  |  |
| NA | ✓ | 19 | NA | 45190 | XAJ:18, BA.4:1 |
| NA | ✓ | 35 | NA | 7147 | XBD:30, BA.2.75:5 |
| NA | ✓ | 36 | NA | 7146 | XBR:1, XBD:30, BA.2.75:5 |
| NA |  | 20 | NA | 1061775 | XS:17, BA.1:1, BA.1.1:1, BA.1.15:1 |
| NA |  | 1321 | NA | 340501 | XE:1113, XJ:68, BA.2:82, XM:29, XY:23, XAL:3, XAF:1, XH:2 |

See notebook analysis ripples.ipynb for details.

#### S1.12 Likely spurious recombinants with many descendants

In the sc2ts ARG, there are two recombination events which have a large number of descendants. The first of these corresponds to recombination event within BA.2 whose breakpoint is at position 27382 on the far right of the genome. Although supported by 5 sites, three of them are adjacent so that they only represent 3 loci, meaning that this event does not pass our QC filter.. The second of these involves a single breakpoint at position 26858, and is the origin of the BA.5 lineage in the ARG. Although supported by 5 loci on the right, this recombinant also requires 3 de-novo mutations, some of which are adjacent to the supporting loci. Furthermore, when the first BA.5 sample is added to the ARG, it is possible to construct a more parsimonious non-recombinant origin by long branch splitting, although subsequent tree building does not make this a parsimonious way to rewire the final ARG. It is therefore likely to be spurious.

## Supplementary Figures

**Figure S1:**
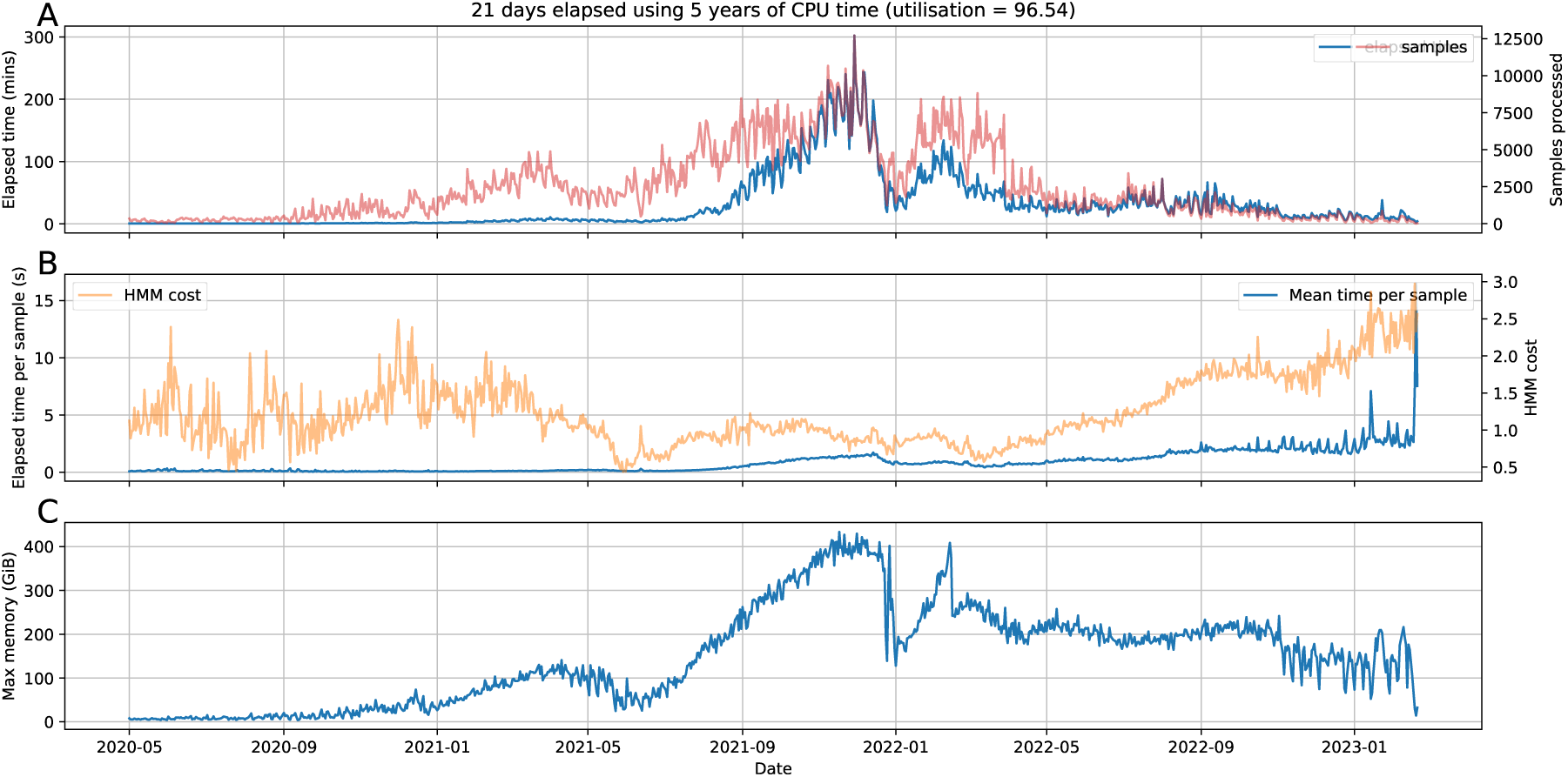
Supplementary Figures Computational resources used during the ARG inference process. (A) The amount of time to process a daily batch of samples and the number of samples processed per day. (B) Mean time to process a sample is less than 5s except when the number of samples is small. (C) RAM usage is the most difficult resource to manage, and depends on the sample composition. The sharp drop in 2022-01 corresponds to the sweep in which Omicron BA.1 replaced Delta and there was very little sample diversity. The drop in RAM usage around 2022-02 corresponds to a change in configuration where we reduced the number of HMM threads from 128 to 80 to better fit into available memory.

**Figure S2:**
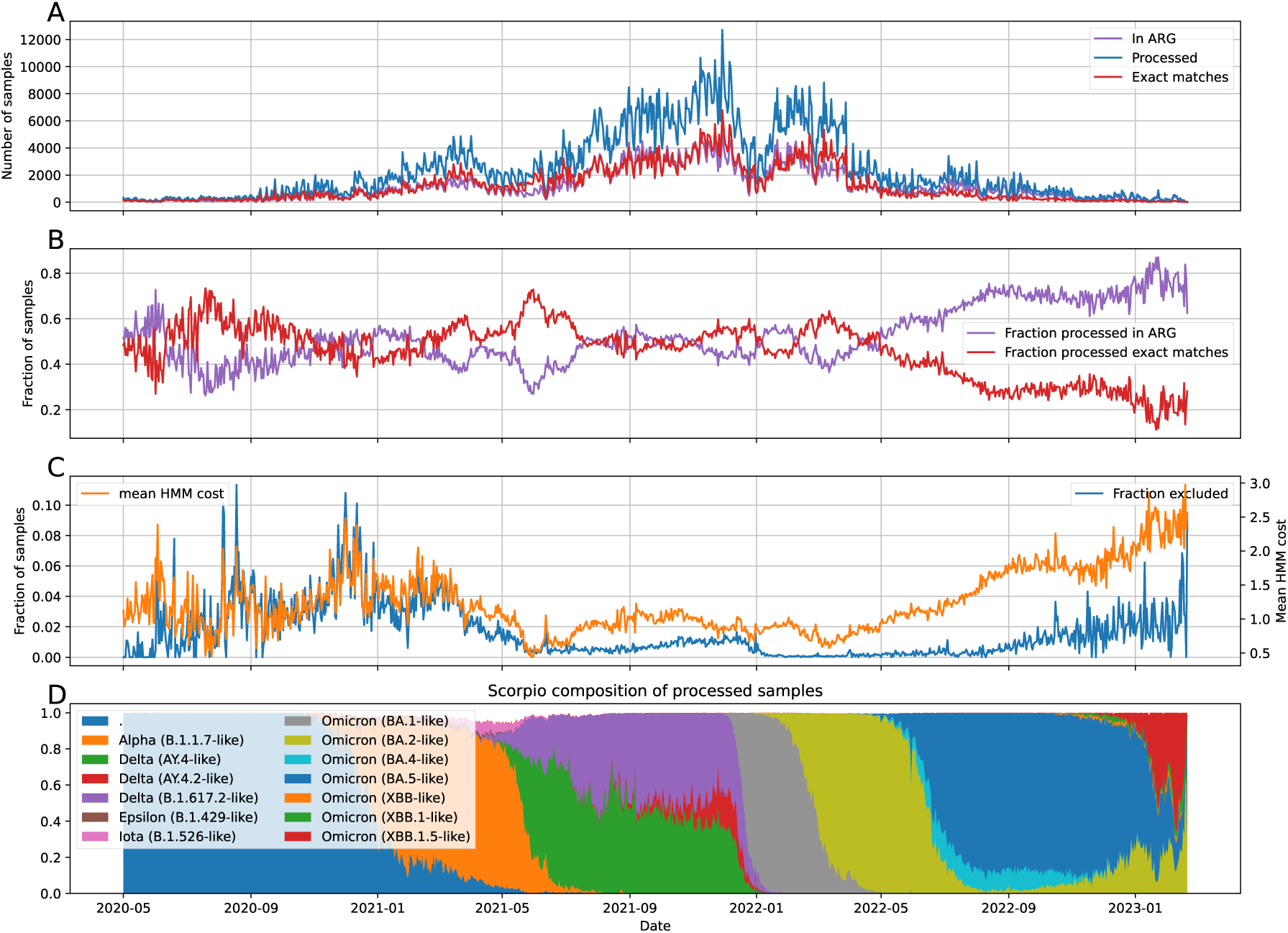
Daily statistics of samples analysed during the ARG inference process. (A) The number of samples processed from the input sequence alignments, exactly matched to a node in the current ARG, and added to the ARG per day. (B) The fraction of exactly matched samples and the fraction of samples added to the ARG mirror each other. The fraction of exact matches is a function of both sampling depth and composition of the global population. (C) The mean HMM cost per sample is a useful metric. (D) Proportion of samples classified by (simplified) Scorpio designation.

**Figure S3:**
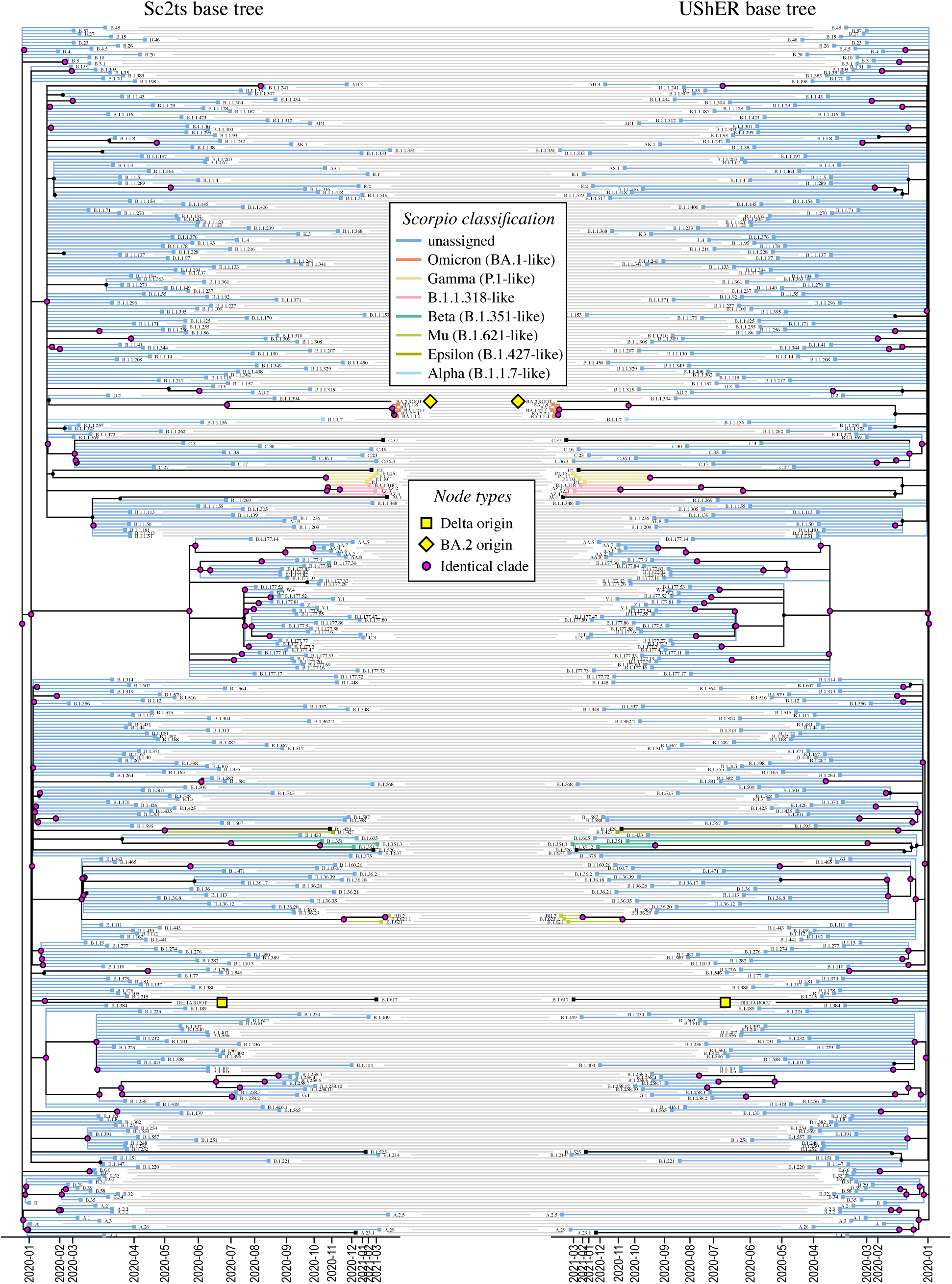
Tanglegram comparing the basic phylogenetic backbones of sc2ts and UShER.

**Figure S4:**
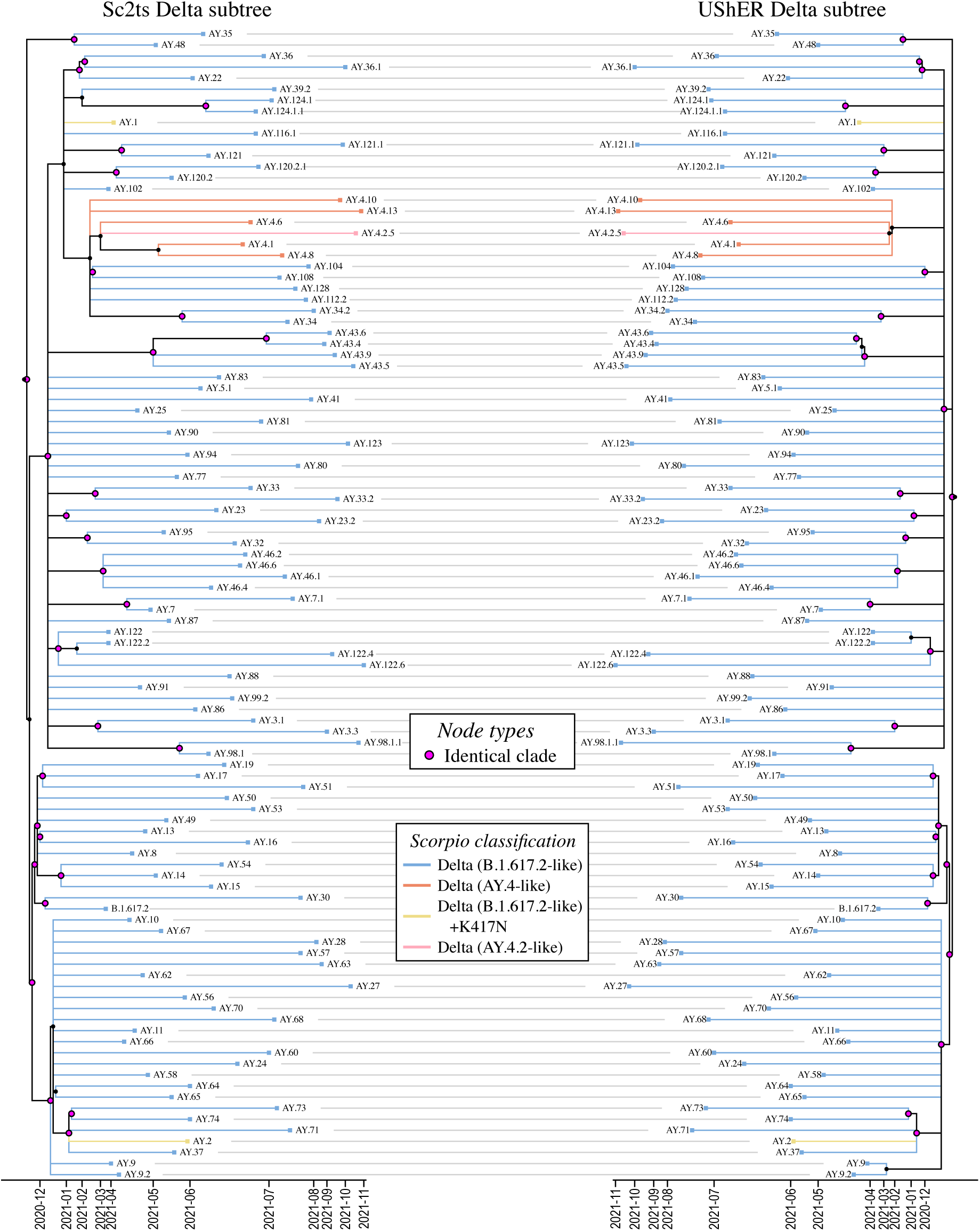
Tanglegram comparing sc2ts and UShER on the Delta subtree.

**Figure S5:**
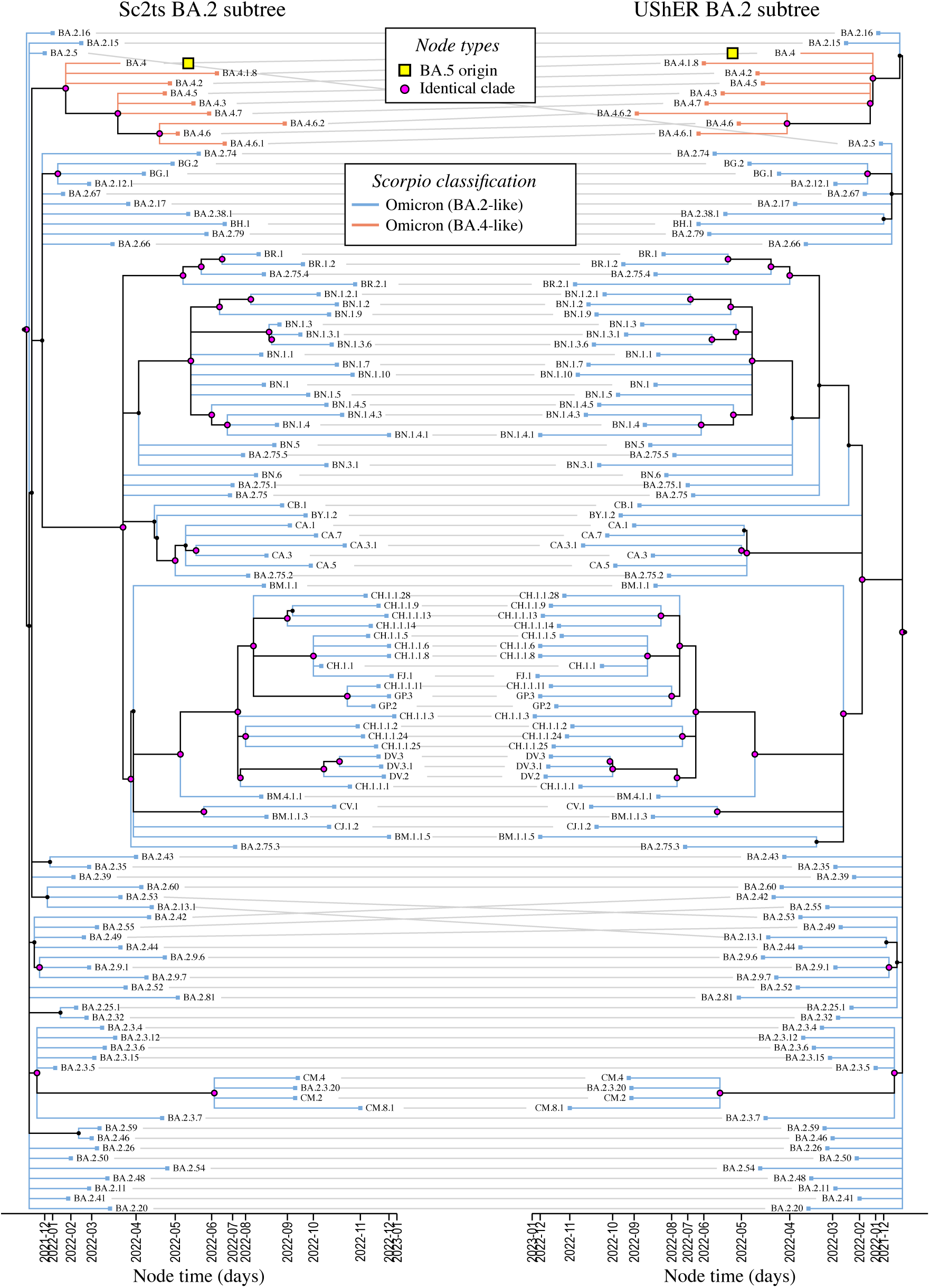
Tanglegram comparing sc2ts and UShER on the BA.2 subtree.

**Figure S6:**
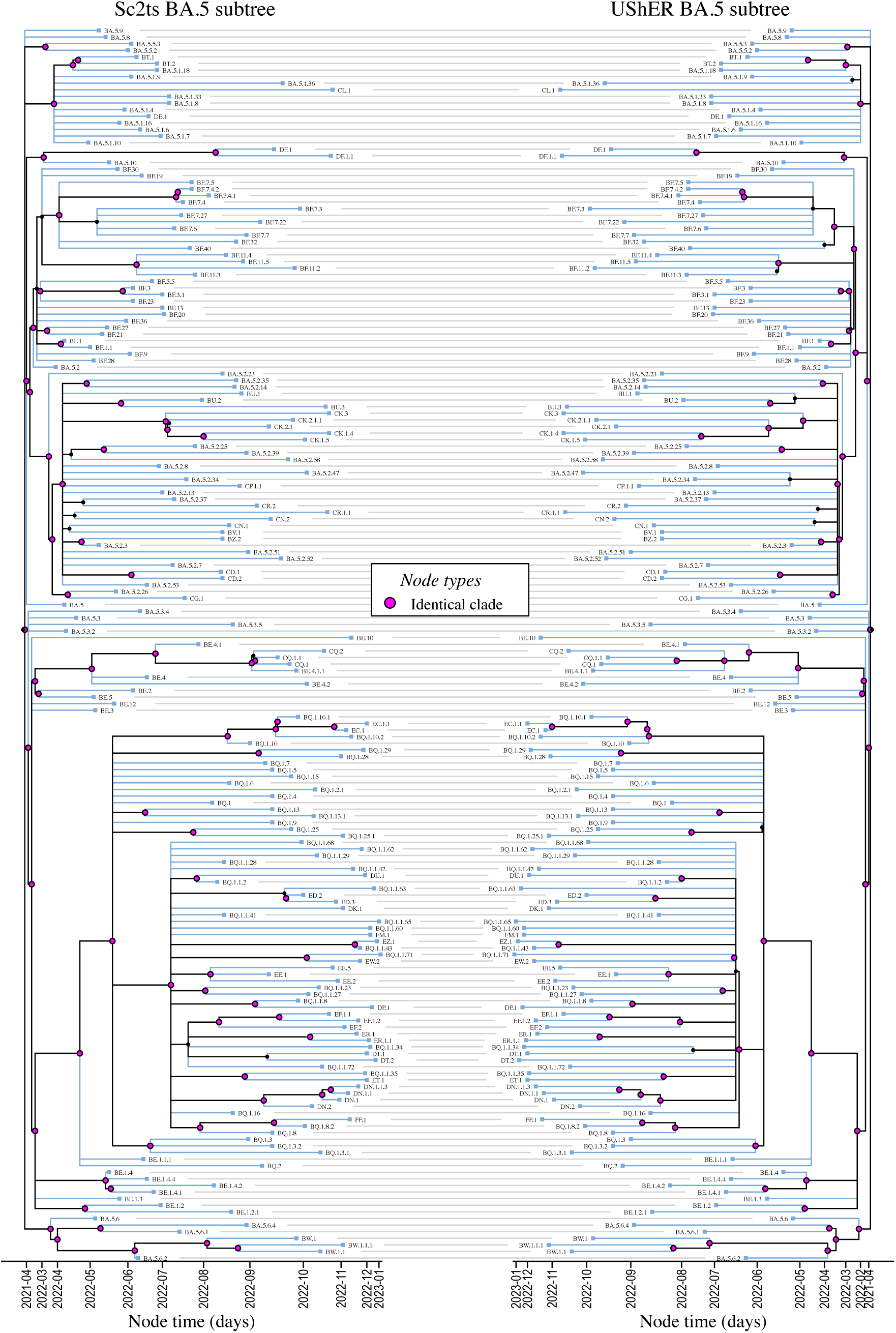
Tanglegram comparing sc2ts and UShER on the BA.5 subtree.

**Figure S7:**
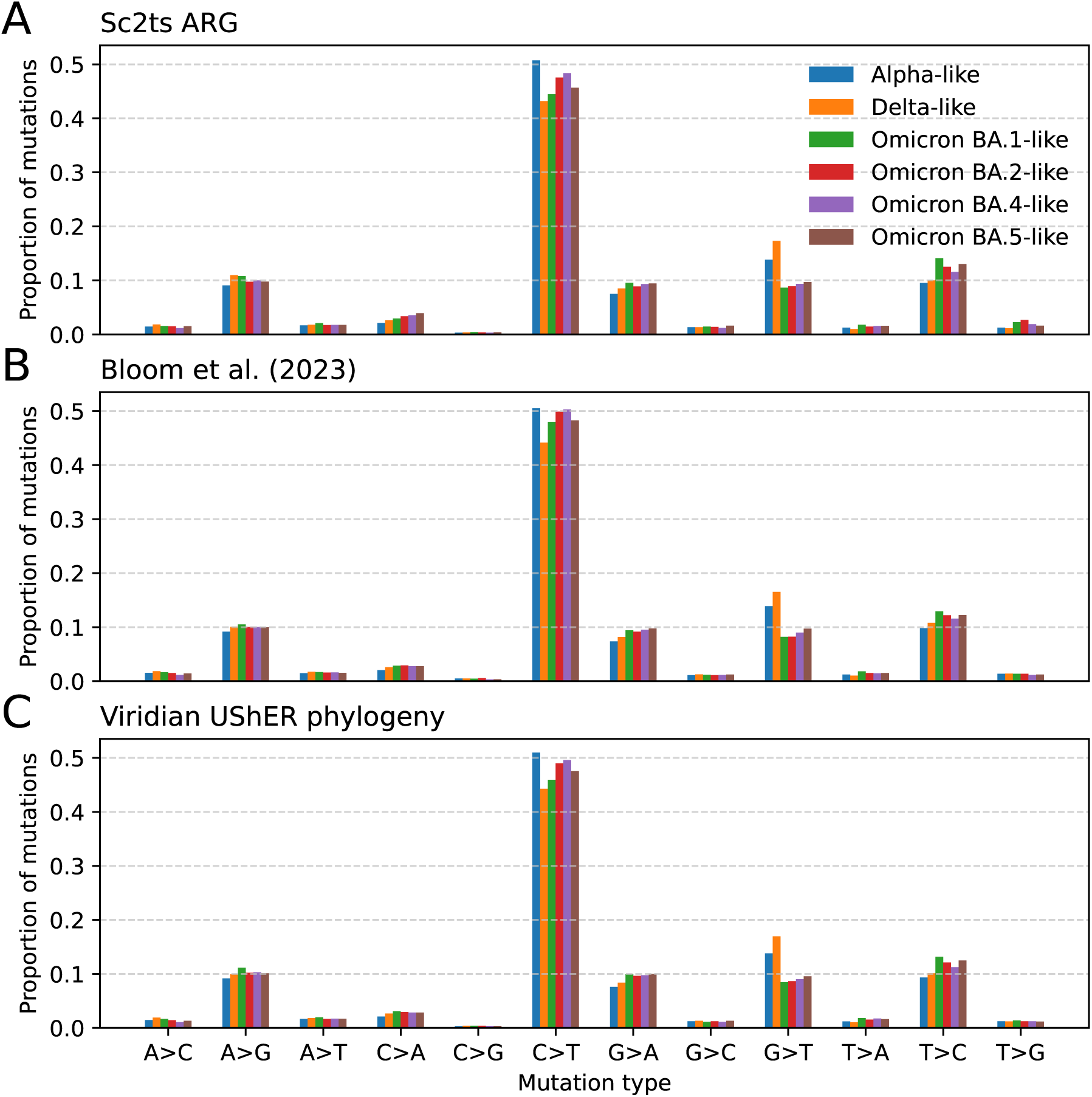
All-site mutational spectra of major VOCs calculated from the sc2ts ARG (A), the mutation count data from Bloom et al. (2023)^78^ (B), and the Viridian UShER phylogeny from Hunt et al. (2024)^76^.

**Figure S8:**
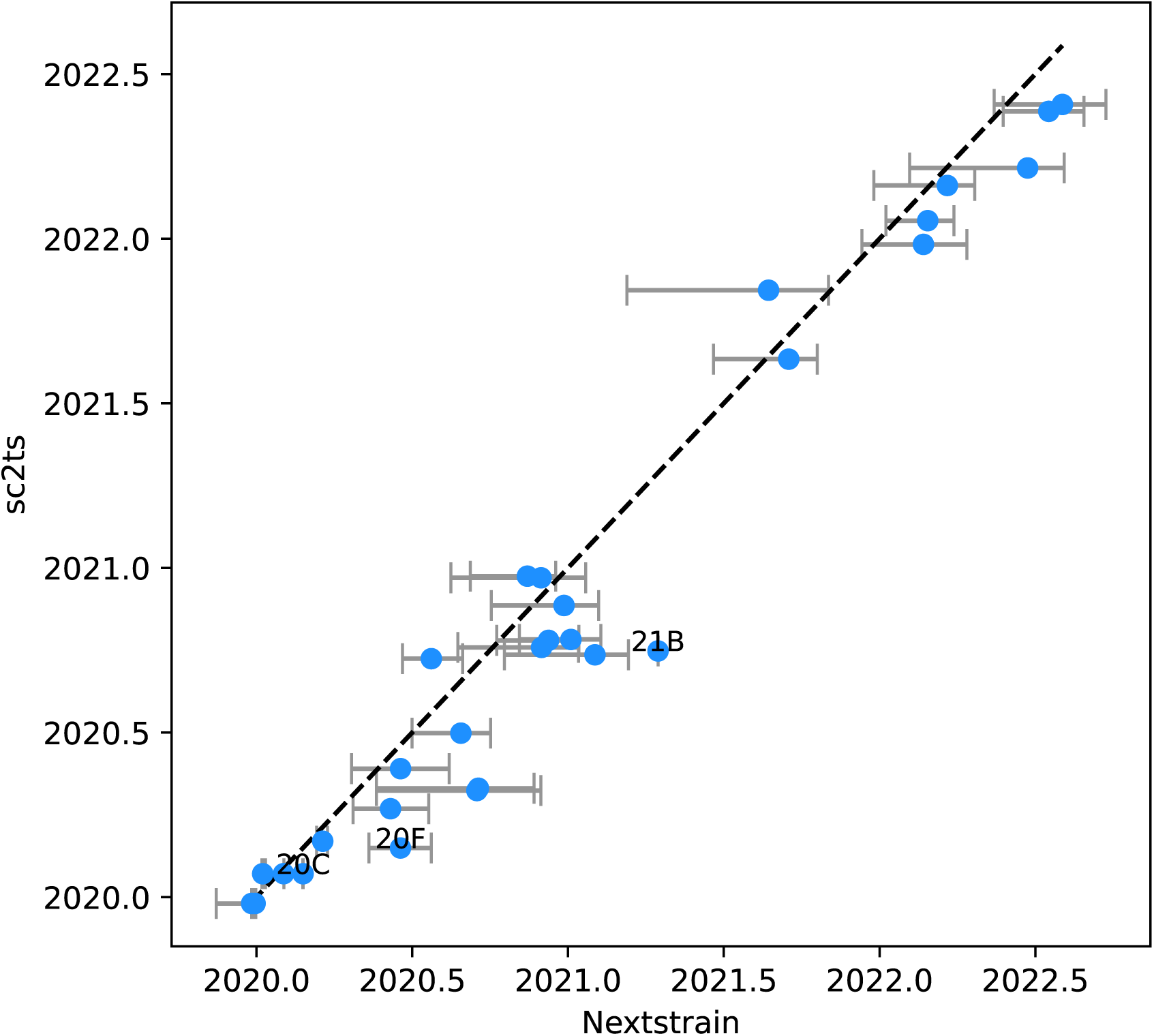
Inferred times of internal nodes corresponding to origins of Nextclade clades (*y*-axis) against those estimated in the Nextclade tree (*x*-axis). Confidence intervals shown as horizontal bars. Clades are labelled where dates differ by more than 28 days.

**Figure S9:**
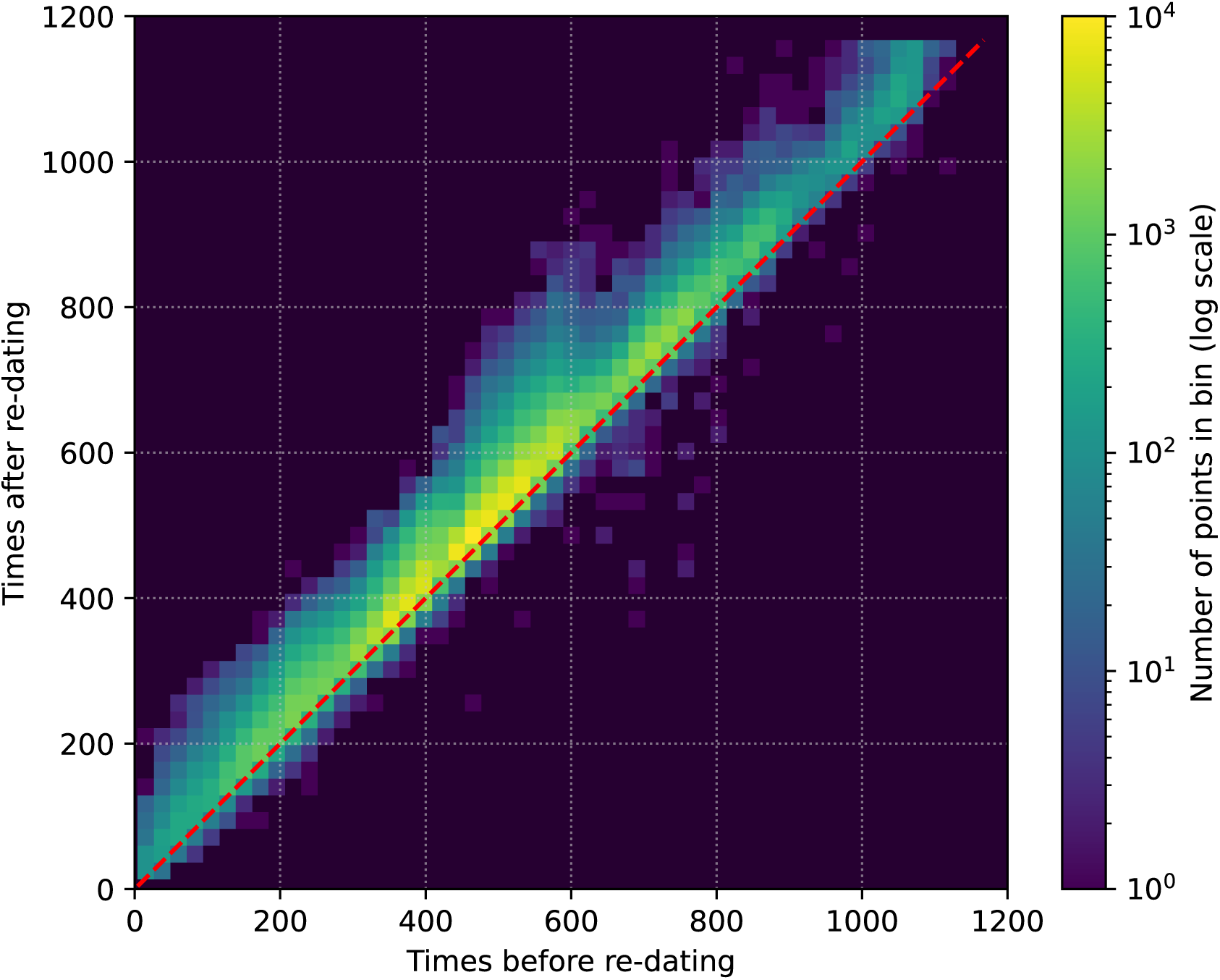
Heat map showing differences (in days) between actual and re-estimated sample dates for a subset of internal sample nodes.

**Figure S10:**
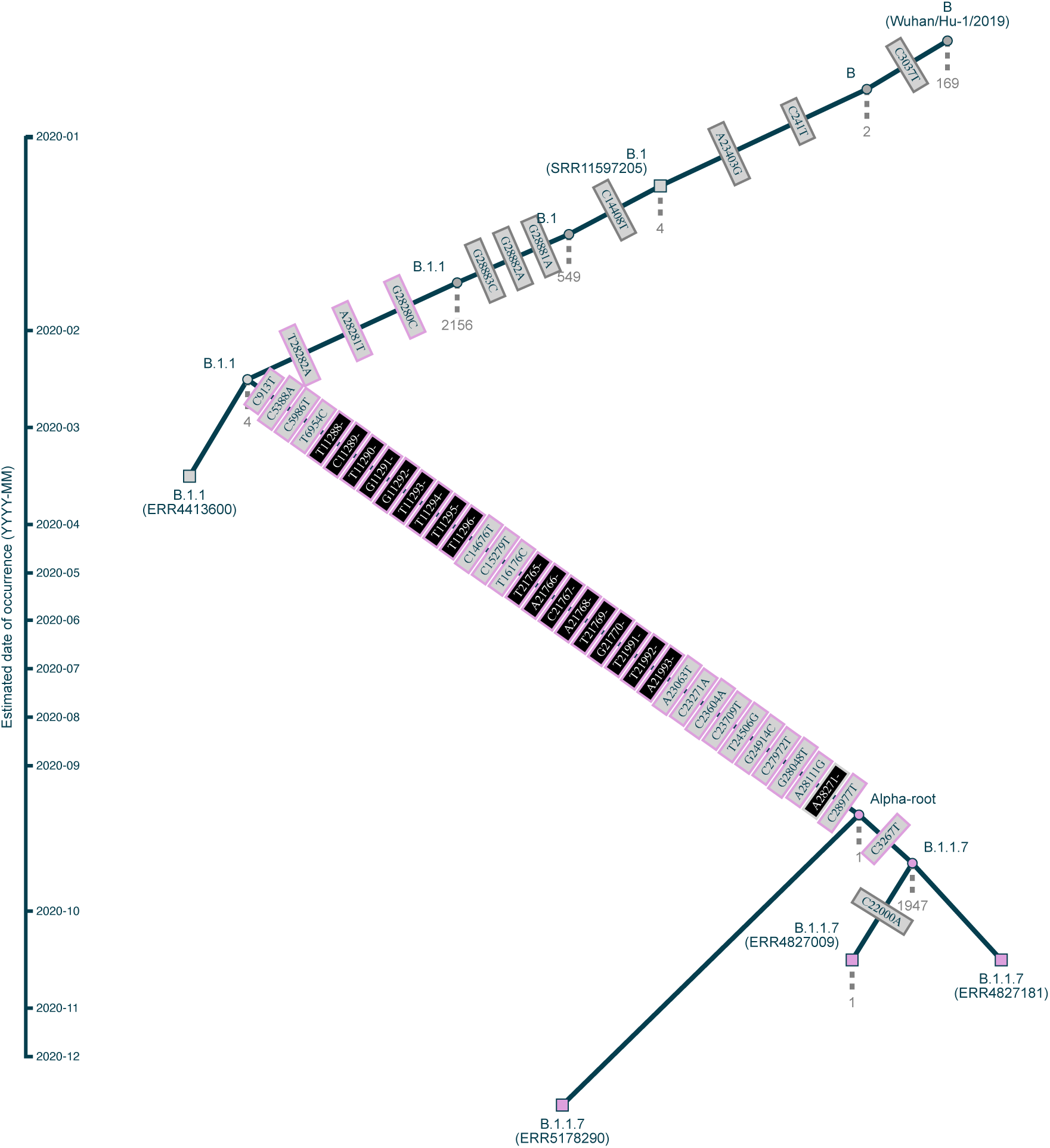
Subgraph illustrating the saltational origin of B.1.1.7 (Alpha). Squares represent sample nodes, circles represent reconstructed internal nodes; nodes classified as B.1.1.7 by Pangolin v4.3.1 are in pink. Mutations are shown as rectangles along branches, ordered by position; deletions are black. Characteristic mutations associated with the emergence of Alpha are highlighted with a pink outline.

**Figure S11:**
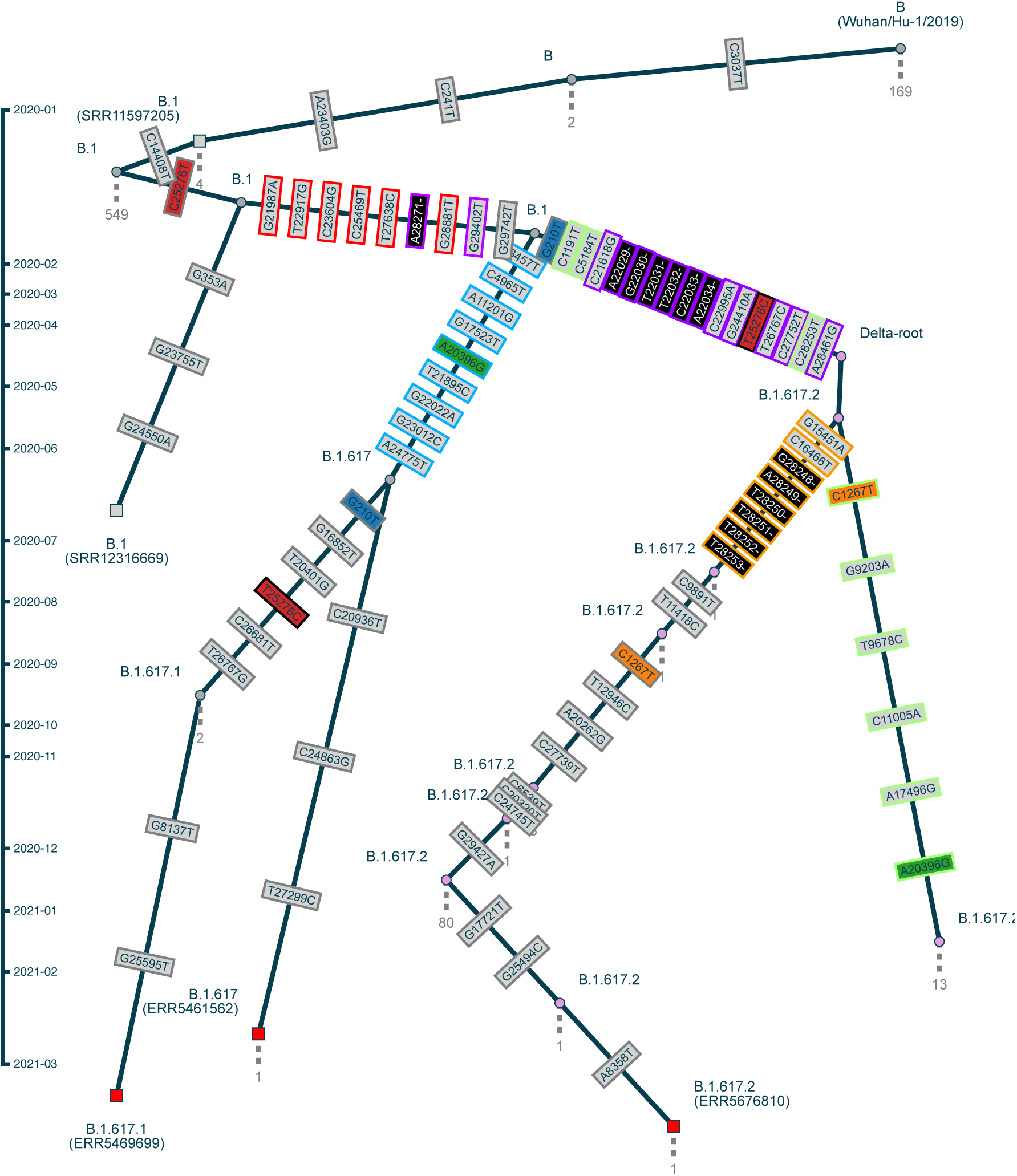
Subgraph illustrating the origin of B.1.617.2 (Delta). Symbols are described in S10, except that we highlight the characteristic mutations of Kappa and Delta: those with a blue outline are listed in the Pango designation issue^161^ or the Kappa constellation^162^, but absent in the Delta constellation; those with a purple outline are listed in the Delta constellation^163^ or are Delta-specific deletions reported in Stern et al.^99^, but absent in the Kappa constellation; and those with a red outline are listed in both the Kappa and Delta constellations. We also outline in orange the mutations associated with the lineage containing clades A to D and in green those associated with lineage containing clade E. Sites associated with multiple mutations in the subgraph have their mutations assigned a unique fill colour (orange, blue, green, red, etc); reversions are further highlighted with a black outline. Sc2ts seed samples (see STAR methods) for B.1.617, B.1.617.1, and B.1.617.2 are plotted as red squares.

**Figure S12:**
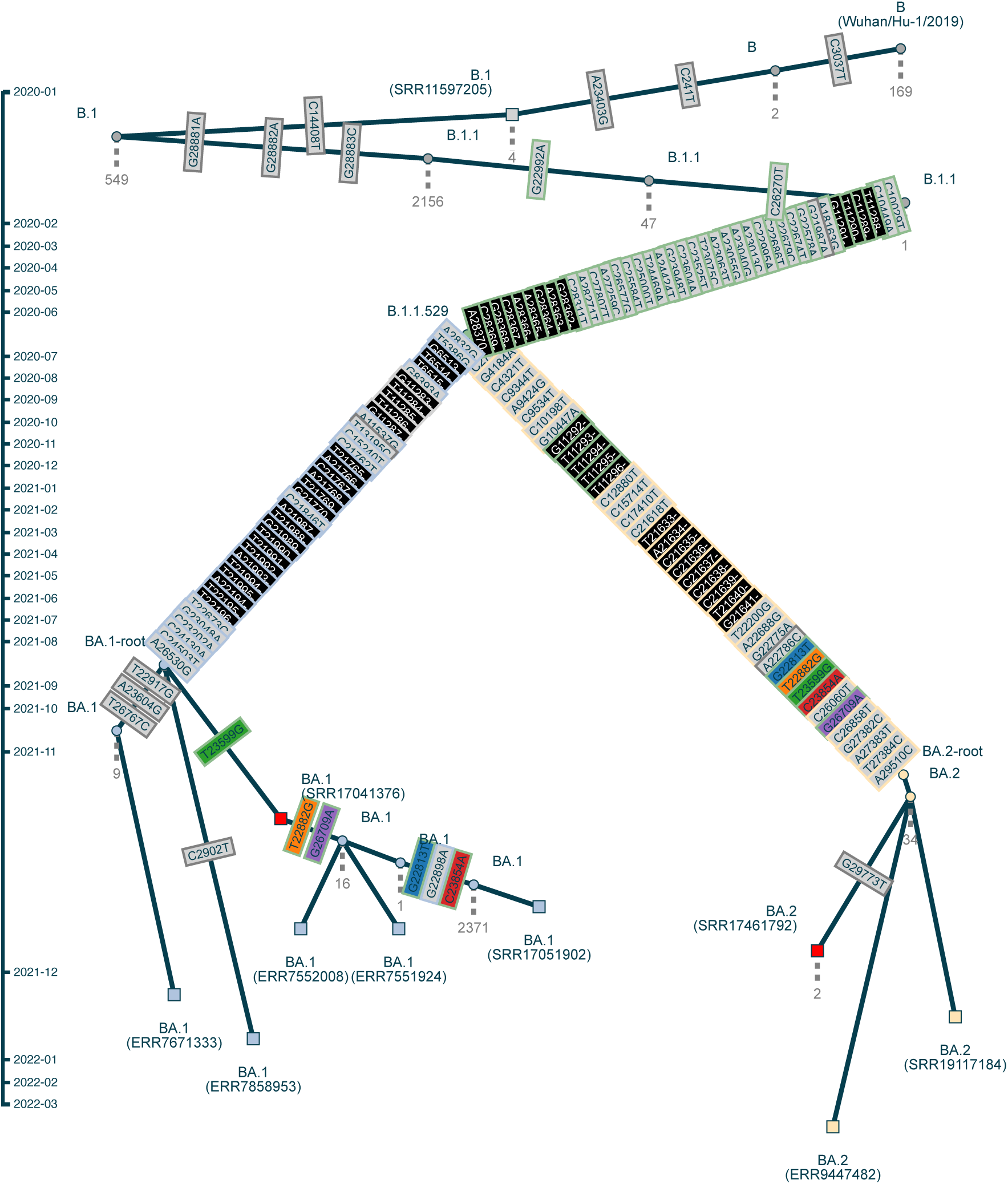
Subgraph illustrating the saltational origin of the major Omicron lineages BA.1 and BA.2. Symbols are described in Figure S11, except that BA.1 samples are filled in light blue (also used to outline BA.1 characteristic mutations), BA.2 samples in light yellow (also used to outline BA.2 characteristic mutations), and B.1.1.529 mutations which are neither BA.1 or BA.2 characteristic mutations are outlined in green. These colours correspond to those used in the BA.1/BA.2 Pango designation issue^101^.

**Figure S13:**
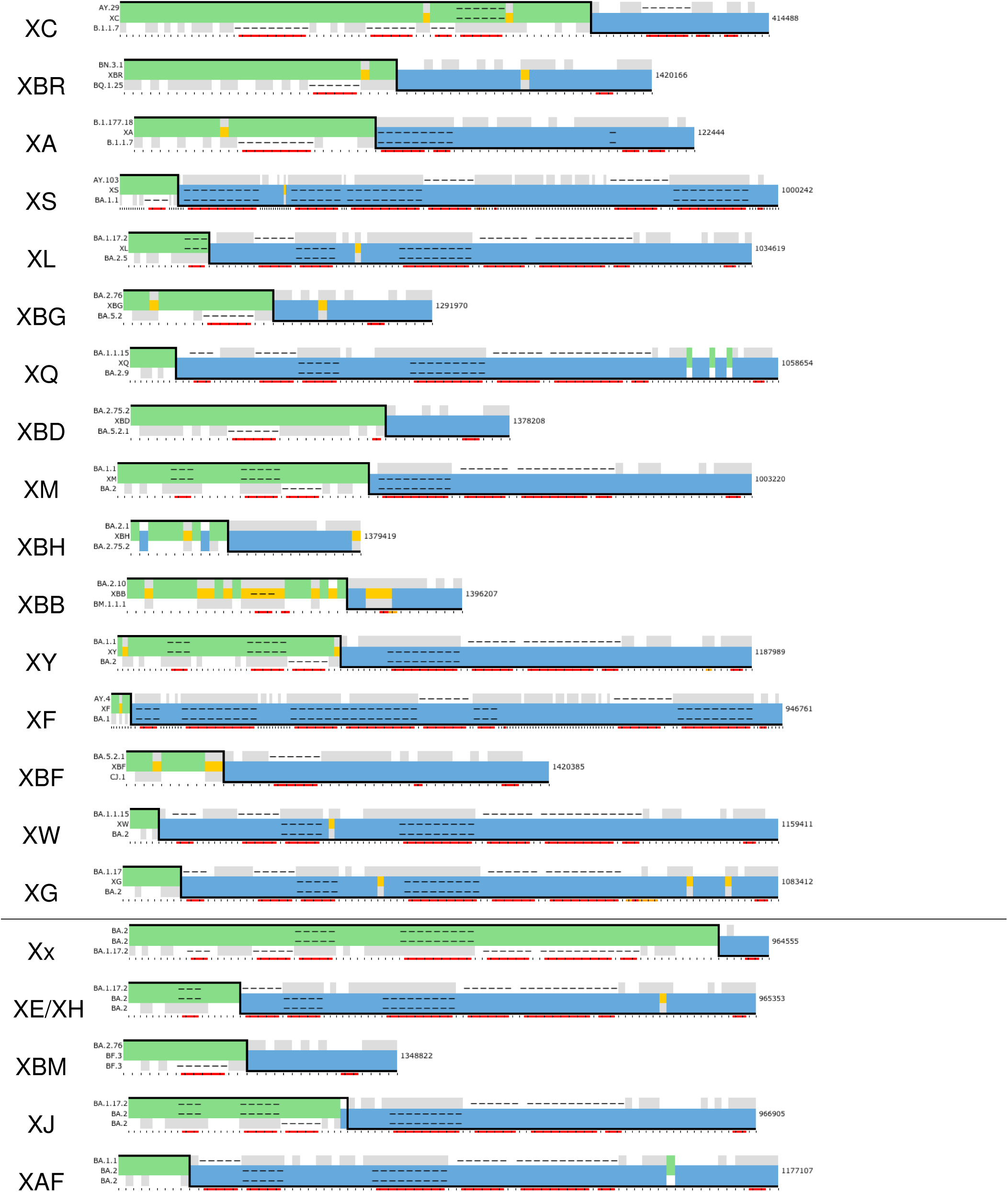
Copying patterns for the 21 recombination events associated with Pango X lineages listed in Table 1 (in the same order). The horizontal line separates Type I and Type II events. See also Document S3 for exact positions and nucleotide bases.

**Figure S14:**
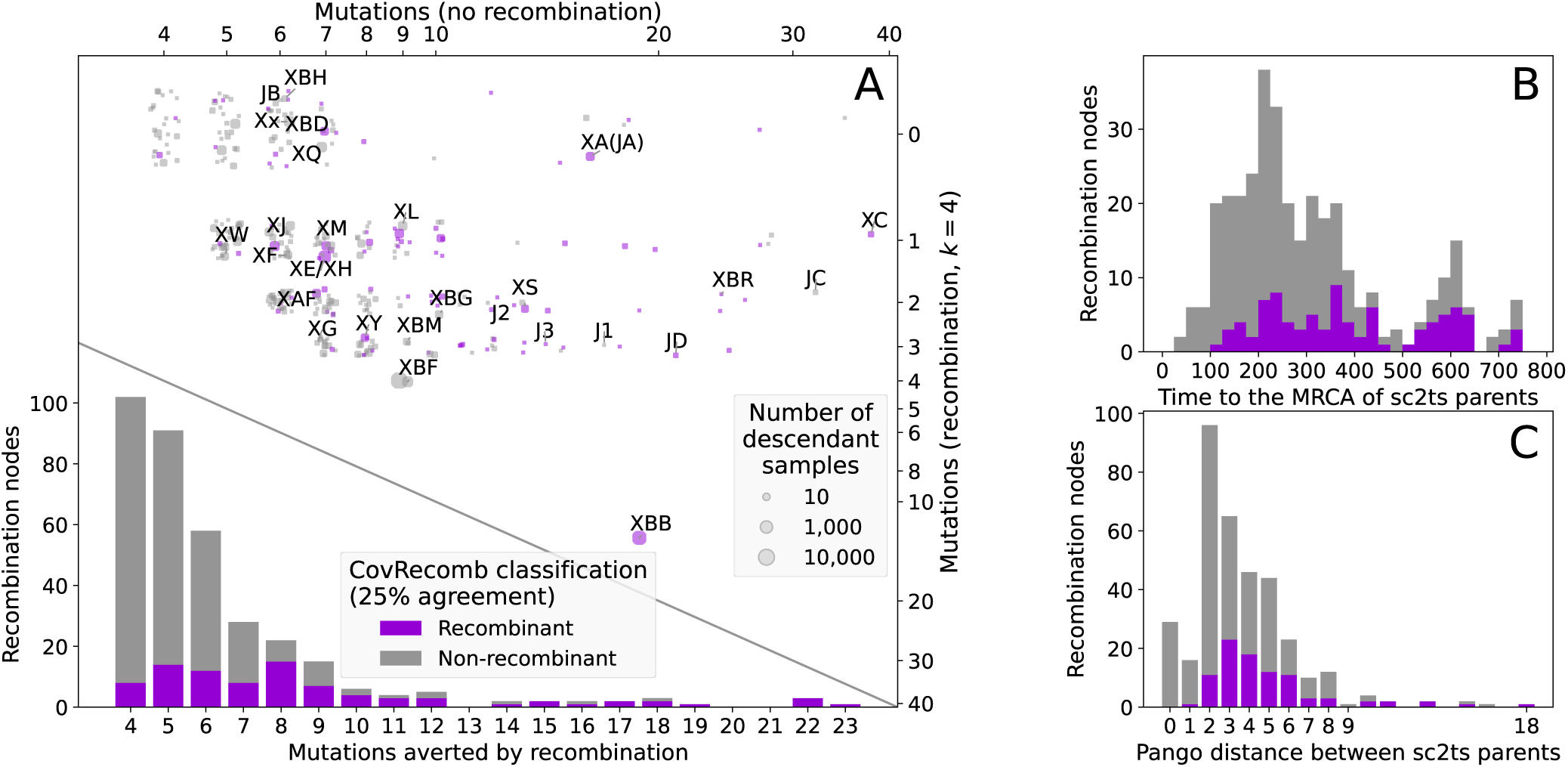
Properties of sc2ts recombination events, coloured by CovRecomb classification. All other details as per Figure 3.

**Figure S15:**
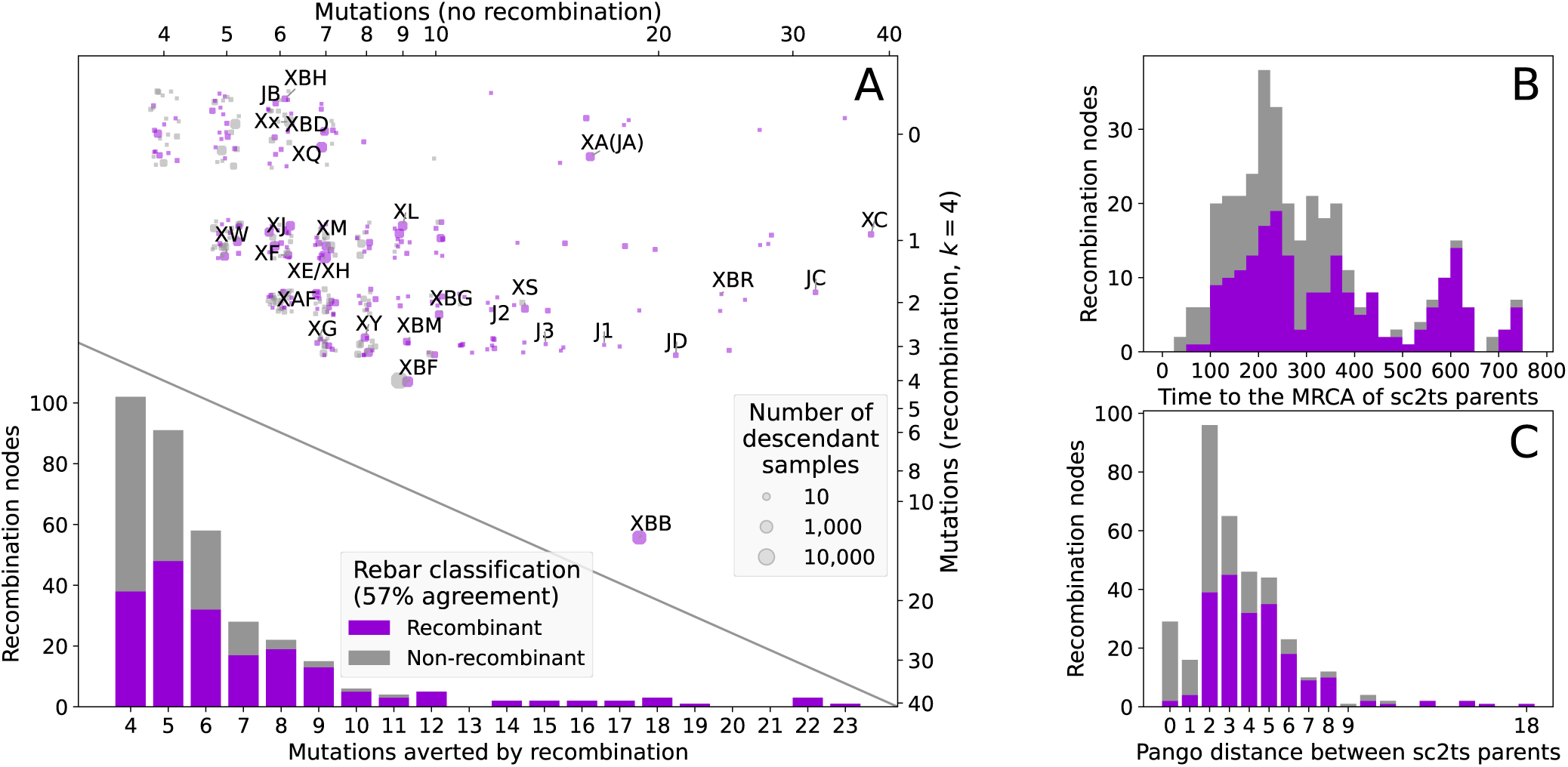
Properties of sc2ts recombination events, coloured by rebar classification. All other details as per Figure 3.

**Figure S16:**
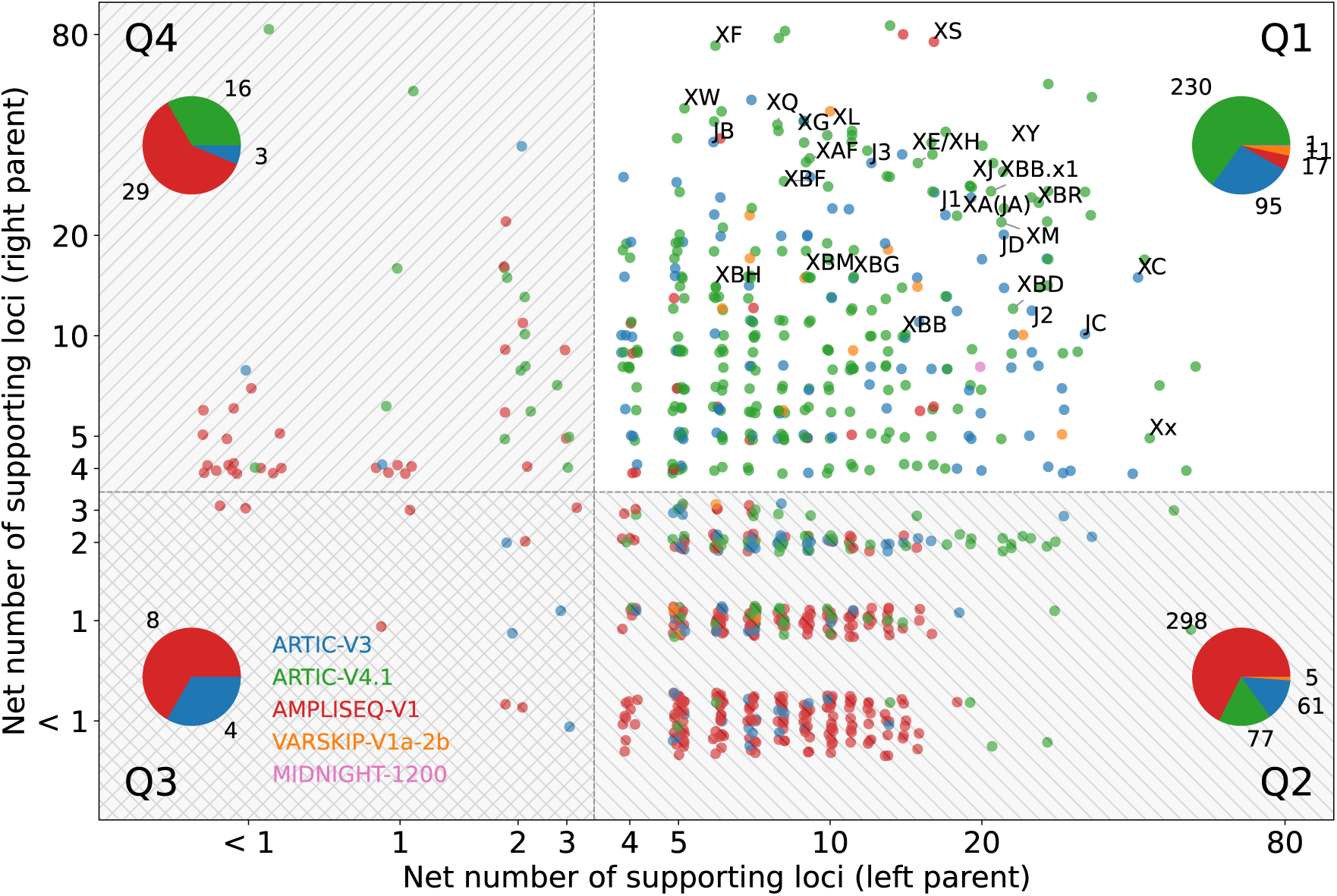
Quality control of recombination events. Scatterplot of recombination events by the net number of supporting loci on the left parent (x-axis) and the right parent (y-axis). The shaded regions highlight potentially artefactual recombination events, which have fewer than 4 net supporting loci on one side or both sides of the suggested breakpoint. colours indicate the primer scheme used for sequencing. The pie charts show breakdowns of the recombination events by primer scheme per quadrant (labeled Q1 to Q4). Recombination events associated with the origins of Pango X and Jackson^29^ lineages are labeled.

**Figure S17:**
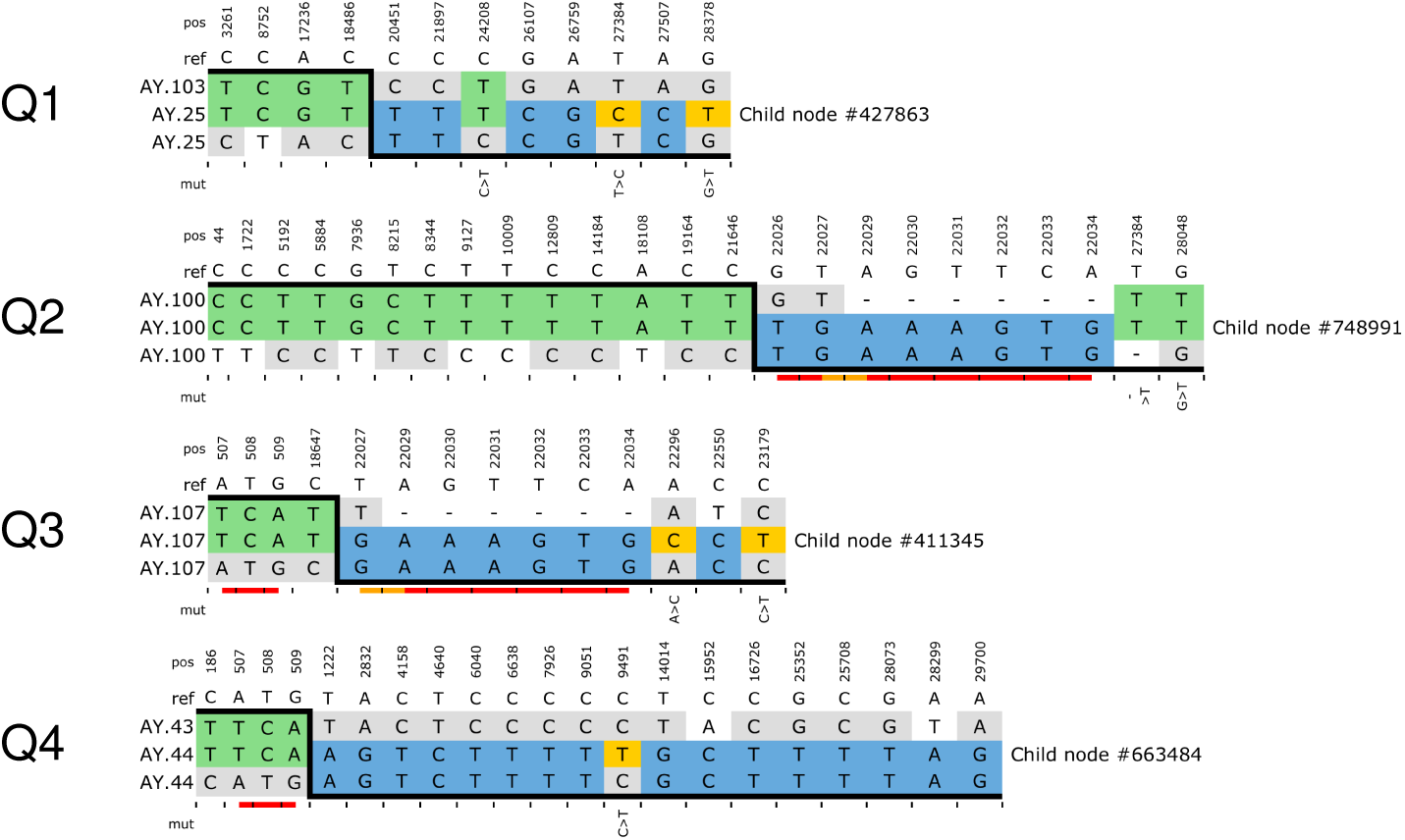
Illustrative copying patterns drawn from the quadrants labelled in Figure S16. Each copying pattern shows the the positions where the allelic state in a recombinant (middle row, labelled by node ID) matches that of the left (P0: upper row, coloured green) or right parent (P1: lower row, coloured blue), or where the recombination event requires a de-novo mutation (gold, with mutational change below). Parental states that correspond to the reference but are not inherited by the recombinant are shown with a gray background. Genome position (“pos”) and reference allele (“ref”) are shown for each column. Underneath the copying pattern, adjacent genomic positions are underlined in red, and near-adjacent sites (within 3 bases) in orange.

**Figure S18:**
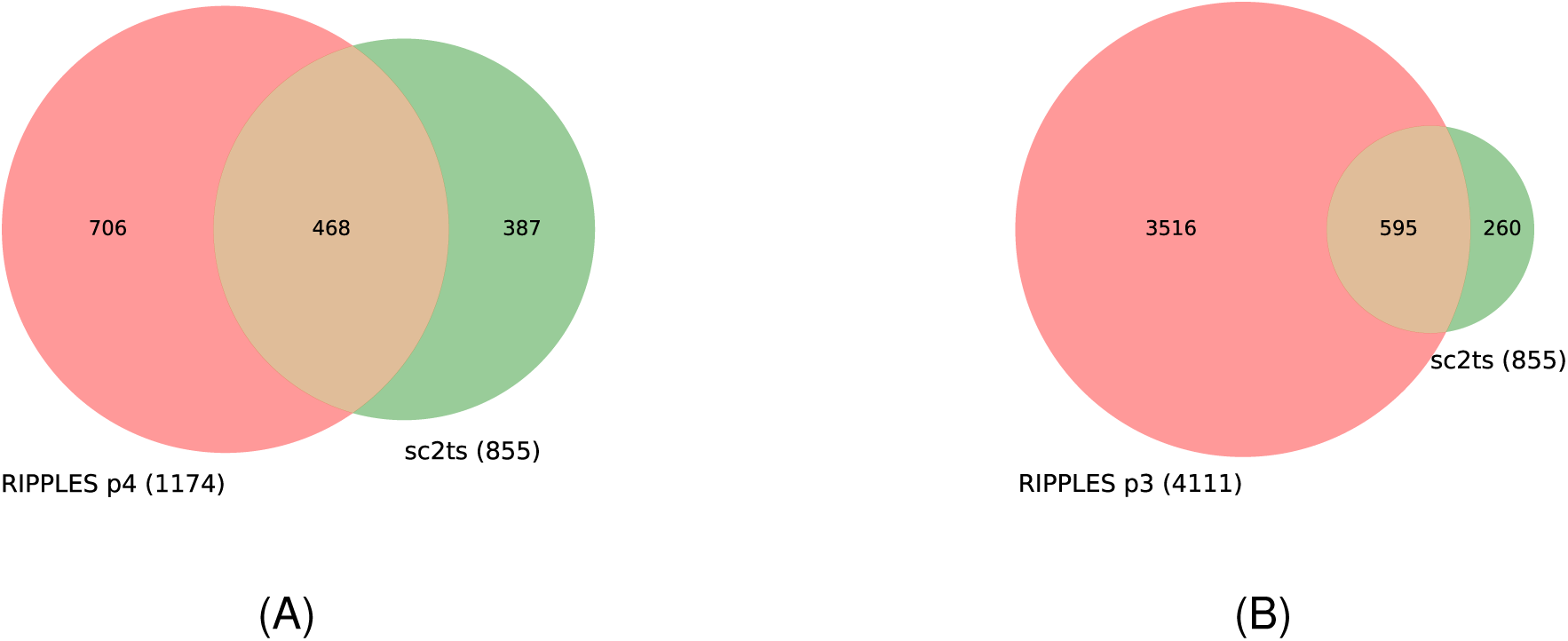
Intersection of the RIPPLES and sc2ts recombination events at different values of the RIPPLES parsimony parameter, p.

**Figure S19:**
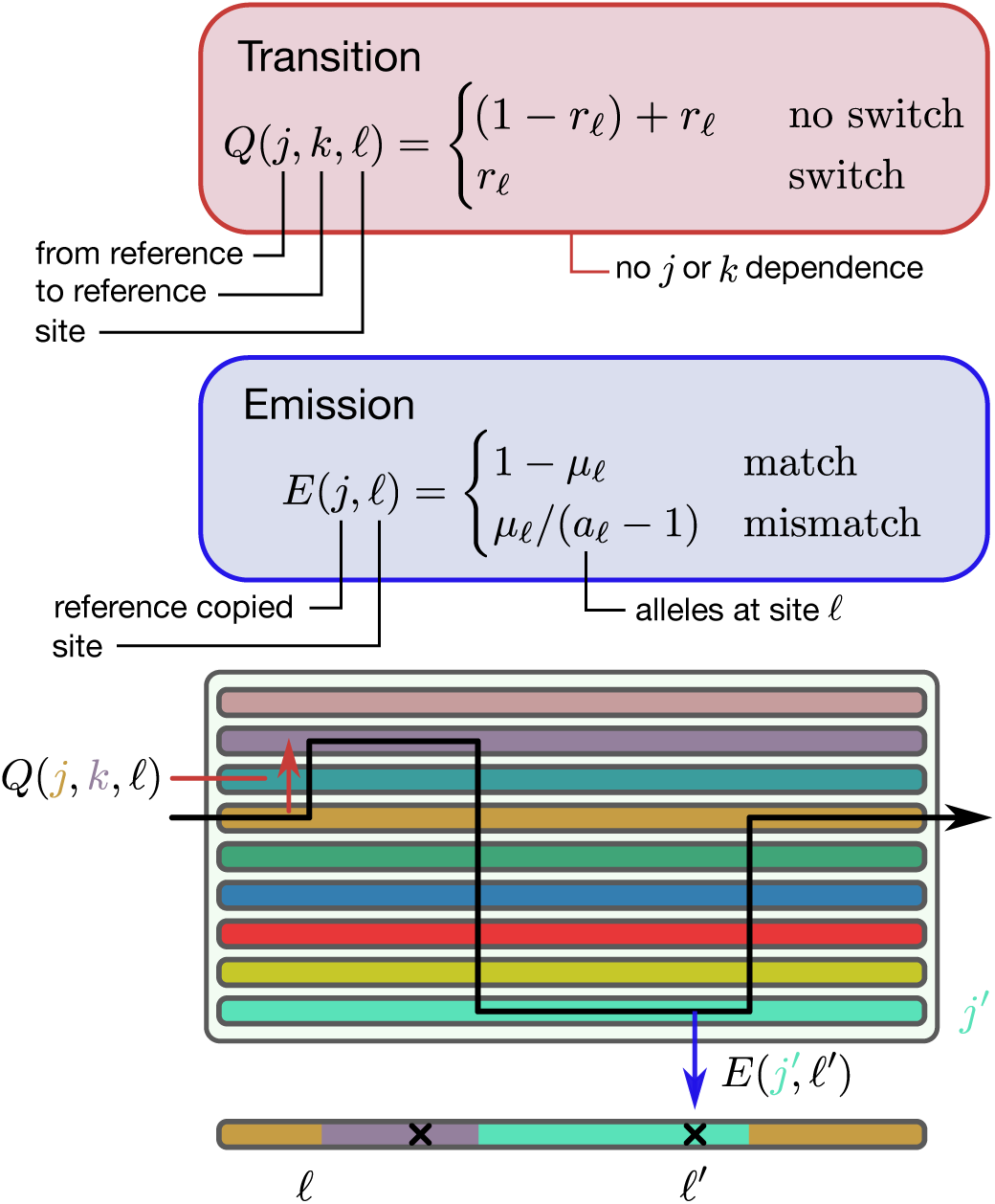
A schematic of the Li and Stephens (LS) model, in which a focal sequence (bottom) is described as an imperfect mosaic of the sequences in a reference panel. Black crosses along the focal sequence show sequencing errors or mutations. In the standard formulation, at site *ℓ*, the recombination probability is *r_ℓ_*, the mutation probability is *µ_ℓ_* and *n* denotes the size of the reference panel. The Viterbi algorithm can be used to find a “copying path” through the reference panel for a given focal sequence that maximises the likelihood under these parameters. Unseen states in the reference panel are shown as coloured lines enclosed by the grey box. The black arrow describes the true path through the data which leads to the emitted focal sequence below. Examples of transition and emission probabilities along this trajectory are shown by the red and blue arrows, respectively.

**Figure S20:**
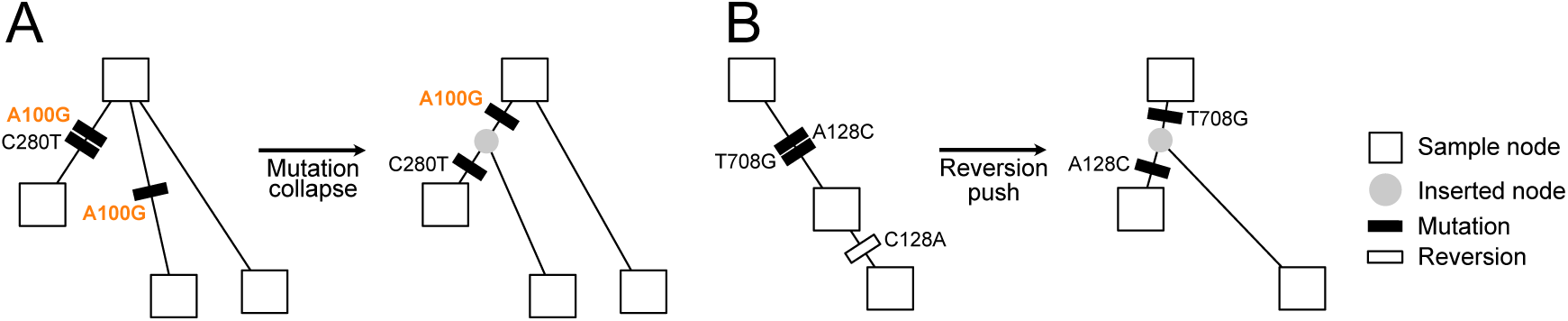
Parsimony improving heuristics. (A) Mutation collapsing. Mutation A100G is shared by two siblings, and we create a new node to represent the ancestor on which this mutation occurred. (B) Reversion pushing. Mutation A128C is immediately reverted by C128A, and we create a new node to represent the ancestor that did not carry A128C.

