## Supplementary material for "A Pandemic-Scale Ancestral Recombination Graph for SARS-CoV-2": Document S2

```
Using a 434.8 megabyte ARG up to 2023-02-21, with 2482157 sampled SARS-CoV2 sequences
(317 trees, 2484587 mutations over 29904.0bp with 855 recomb. events)
```

| 44 main pango-X lineages | 135 sub pango-X lineages |
|---|---|
| XA, XAA, XAC, XAD, XAE, XAF, XAG, XAJ, XAL, XAM, XAN, XAP, XAS, XAU, XAV, XAZ, XB, XBB, XBD, XBE, XBF, XBG, XBH, XBK, XBM, XBQ, XBR, XC, XE, XF, XG, XH, XJ, XL, XM, XN, XP, XQ, XR, XS, XU, XW, XY, XZ | XBB.1, XBB.1.1, XBB.1.11, XBB.1.13, XBB.1.14, XBB.1.15, XBB.1.18, XBB.1.18.1, XBB.1.22.1, XBB.1.28, XBB.1.29, XBB.1.3, XBB.1.30, XBB.1.32, XBB.1.34, XBB.1.35, XBB.1.37, XBB.1.4, XBB.1.4.1, XBB.1.43, XBB.1.43.1, XBB.1.45, XBB.1.45.1, XBB.1.46, XBB.1.5, XBB.1.5.1, XBB.1.5.10, XBB.1.5.100, XBB.1.5.102, XBB.1.5.104, XBB.1.5.107, XBB.1.5.11, XBB.1.5.12, XBB.1.5.13, XBB.1.5.14, XBB.1.5.15, XBB.1.5.16, XBB.1.5.17, XBB.1.5.18, XBB.1.5.19, XBB.1.5.2, XBB.1.5.20, XBB.1.5.21, XBB.1.5.23, XBB.1.5.24, XBB.1.5.25, XBB.1.5.26, XBB.1.5.28, XBB.1.5.3, XBB.1.5.30, XBB.1.5.31, XBB.1.5.32, XBB.1.5.33, XBB.1.5.34, XBB.1.5.35, XBB.1.5.36, XBB.1.5.37, XBB.1.5.38, XBB.1.5.39, XBB.1.5.4, XBB.1.5.40, XBB.1.5.41, XBB.1.5.43, XBB.1.5.46, XBB.1.5.47, XBB.1.5.48, XBB.1.5.49, XBB.1.5.5, XBB.1.5.51, XBB.1.5.52, XBB.1.5.55, XBB.1.5.56, XBB.1.5.57, XBB.1.5.59, XBB.1.5.6, XBB.1.5.60, XBB.1.5.61, XBB.1.5.62, XBB.1.5.63, XBB.1.5.64, XBB.1.5.65, XBB.1.5.66, XBB.1.5.67, XBB.1.5.69, XBB.1.5.7, XBB.1.5.73, XBB.1.5.75, XBB.1.5.77, XBB.1.5.78, XBB.1.5.79, XBB.1.5.8, XBB.1.5.80, XBB.1.5.86, XBB.1.5.88, XBB.1.5.9, XBB.1.5.90, XBB.1.5.91, XBB.1.5.93, XBB.1.5.95, XBB.1.5.96, XBB.1.5.97, XBB.1.7, XBB.1.9, XBB.1.9.1, XBB.1.9.2, XBB.1.9.4, XBB.2, XBB.2.1, XBB.2.10, XBB.2.11.1, XBB.2.2, XBB.2.4, XBB.2.5, XBB.2.6, XBB.2.7, XBB.2.8, XBB.3, XBB.3.1, XBB.3.2, XBB.3.5, XBB.4, XBB.6, XBB.6.1, XBB.8, XBB.9, XBF.1, XBF.10, XBF.2, XBF.3, XBF.4, XBF.5, XBF.6, XBF.7, XBF.9, XBK.1 |

```
Consensus mutations for each lineage taken from https://covidcg.org
```

Bold = main pango

| *RE node* | *pango* | *parents* | *break@* | *# descendants* | *Most common* |
|---|---|---|---|---|---|
| **122444** | **XA** | B.1.177.18/B.1.1.7 | 21765 | 39 of which 39 XA | XA: 39 |
| | ~~XB~~ | | | 0 of which 0 XB | |
| **414488** | **XC** | AY.29/B.1.1.7 | 27390 | 5 of which 5 XC | XC: 5 |
| | *XD* | not in dataset | | | |
| **965353** | **XE** | BA.1.17.2/BA.2 | 11283 | 1156 of which 1116 XE | XE: 1116, BA.2: 37, XH: 2 |
| **946761** | **XF** | AY.4/BA.1 | 6402 | 16 of which 16 XF | XF: 16 |
| **1083412** | **XG** | BA.1.17/BA.2 | 6513 | 3 of which 3 XG | XG: 3 |
| 965353 | XH | BA.1.17.2/BA.2 | 11283 | 1156 of which 2 XH | XE: 1116, BA.2: 37, XH: 2 |
| **966905** | **XJ** | BA.1.17.2/BA.2 | 17410 | 85 of which 68 XJ | XJ: 68, BA.2: 17 |
| | *XK* | not in dataset | | | |
| **1034619** | **XL** | BA.1.17.2/BA.2 | 8393 | 64 of which 64 XL | XL: 64 |
| | ~~XM~~ | | | 0 of which 0 XM | |
| | ~~XN~~ | | | 0 of which 0 XN | |
| | ~~XP~~ | | | 0 of which 0 XP | |
| **1058654** | **XQ** | BA.1.1.15/BA.2.9 | 5386 | 154 of which 55 XQ | XQ: 55, BA.2: 37, XAM: 21 |
| 1058654 | XR | BA.1.1.15/BA.2.9 | 5386 | 154 of which 17 XR | XQ: 55, BA.2: 37, XAM: 21 |
| **1000242** | **XS** | AY.103/BA.1.1 | 10449 | 17 of which 17 XS | XS: 17 |
| | *XT* | not in dataset | | | |
| 1058654 | XU | BA.1.1.15/BA.2.9 | 5386 | 154 of which 1 XU | XQ: 55, BA.2: 37, XAM: 21 |
| | *XV* | not in dataset | | | |
| **1159411** | **XW** | BA.1.1.15/BA.2 | 4321 | 32 of which 32 XW | XW: 32 |
| **1187989** | **XY** | BA.1.1/BA.2 | 12880 | 23 of which 23 XY | XY: 23 |
| **964555** | **XZ** | BA.2/BA.1.17.2 | 26060 | 253 of which 48 XZ | BA.2: 156, XZ: 48, XAP: 20 |
| 1058654 | XAA | BA.1.1.15/BA.2.9 | 5386 | 154 of which 17 XAA | XQ: 55, BA.2: 37, XAM: 21 |
| | *XAB* | not in dataset | | | |
| 964555 | XAC | BA.2/BA.1.17.2 | 26060 | 253 of which 18 XAC | BA.2: 156, XZ: 48, XAP: 20 |
| 964555 | XAD | BA.2/BA.1.17.2 | 26060 | 253 of which 2 XAD | BA.2: 156, XZ: 48, XAP: 20 |
| 964555 | XAE | BA.2/BA.1.17.2 | 26060 | 253 of which 9 XAE | BA.2: 156, XZ: 48, XAP: 20 |
| **1177107** | **XAF** | BA.1.1/BA.2 | 10447 | 36 of which 1 XAF | BA.2: 35, XAF: 1 |
| 1058654 | XAG | BA.1.1.15/BA.2.9 | 5386 | 154 of which 6 XAG | XQ: 55, BA.2: 37, XAM: 21 |
| | *XAH* | not in dataset | | | |
| | ~~XAJ~~ | | | 0 of which 0 XAJ | |
| | *XAK* | not in dataset | | | |
| 1003220 | XAL | BA.1.1/BA.2 | 21595 | 45 of which 3 XAL | XM: 26, BA.2: 16, XAL: 3 |
| 1058654 | XAM | BA.1.1.15/BA.2.9 | 5386 | 154 of which 21 XAM | XQ: 55, BA.2: 37, XAM: 21 |
| | ~~XAN~~ | | | 167127 of which 7 XAN | BA.5.2.1: 30582, BA.5.1: 18649, BA.5.2: 15522 |
| 964555 | XAP | BA.2/BA.1.17.2 | 26060 | 253 of which 20 XAP | BA.2: 156, XZ: 48, XAP: 20 |
| | *XAQ* | not in dataset | | | |
| | *XAR* | not in dataset | | | |
| | ~~XAS~~ | | | 0 of which 0 XAS | |
| | *XAT* | not in dataset | | | |
| | ~~XAU~~ | | | 0 of which 0 XAU | |
| | ~~XAV~~ | | | 167127 of which 13 XAV | BA.5.2.1: 30582, BA.5.1: 18649, BA.5.2: 15522 |
| | *XAW* | not in dataset | | | |
| | *XAY* | not in dataset | | | |
| | ~~XAZ~~ | | | 167127 of which 133 XAZ | BA.5.2.1: 30582, BA.5.1: 18649, BA.5.2: 15522 |
| | *XBA* | not in dataset | | | |
| 1396207 | XBB | BA.2.10/BM.1.1.1 | 22577 | 6455 of which 71 XBB | XBB.1.5: 3338, XBB.1: 524, XBB.1.5.7: 224 |
| | *XBC* | not in dataset | | | |
| **1378208** | **XBD** | BA.2.75.2/BA.5.2.1 | 24620 | 30 of which 30 XBD | XBD: 30 |
| | ~~XBE~~ | | | 167127 of which 65 XBE | BA.5.2.1: 30582, BA.5.1: 18649, BA.5.2: 15522 |
| **1420385** | **XBF** | BA.5.2.1/CJ.1 | 9866 | 185 of which 124 XBF | XBF: 124, XBF.4: 22, XBF.3: 22 |
| **1291970** | **XBG** | BA.2.76/BA.5.2 | 22917 | 25 of which 25 XBG | XBG: 25 |
| **1379419** | **XBH** | BA.2.1/BA.2.75.2 | 22001 | 6 of which 2 XBH | BA.2: 4, XBH: 2 |
| | ~~XBK~~ | | | 0 of which 0 XBK | |
| **1348822** | **XBM** | BA.2.76/BF.3 | 22917 | 12 of which 10 XBM | XBM: 10, BF.3: 2 |
| | ~~XBQ~~ | | | 0 of which 0 XBQ | |
| **1420166** | **XBR** | BN.3.1/BQ.1.25 | 22190 | 1 of which 1 XBR | XBR: 1 |

```
21 total pango X recombinant origins of which 19 include all descendants of the dominant gro
up (exceptions: XM and XBB)
```

### Pango-X Subgraphs

Below we display subgraphs for all the main PangoX lineages that have samples present in the *sc2ts* ARG. Pango designations for both samples and internal nodes were assigned using XX TODO: fill out details XXX. For nodes with large numbers of descendants, only a selected sample of (say) 20-50 Pango X samples are shown. Extra descendants of a node are shown with dotted lines indicating additional immediate children of a node. In some cases, additional descendant nodes of different Pango designations (e.g. BA.2) are shown for context.

Recombination nodes are presented as larger circular nodes, with a Pango designation followed by the breakpoint position(s) surrounded by slashes, e.g. a breakpoint at position 1234 bp is indicated as **/1234/** (but note that PangoX lineages that are not of recombinant origin in sc2ts will not have a clear recombination node). Mutations within each subgraph (tickmarks along edges) are coloured pink if they are flagged as consensus mutations for those lineages: often such mutations occur in lineages above the PangoX origination node. Alternatively, if there are multiple mutations at the same site within a subgraph (indicating reversions or recurrent mutations) they plotted in a unique colour. For example, two green mutation tickmarks will represent mutations at the same site. If one is a reversion of a previous mutation (often indicating an unparsimonious reconstruction of topology), then the mutation is emphasised with a solid black outline. Deletion mutations are filled in black, and reversions of deletions (expected not to happen spontaneously) are magenta with a black outline.

In the PDF version of this document, hovering over node names will reveal the sample_id of a node, and hovering over a mutation will reveal the position of the mutation and the inherited vs derived state. E.g. a mouseover label of `mut:A1234T` denotes a mutation from an A to a T at position 1234 in the genome. Technially this is implemented by faking a URL (this leads to the slightly annoying behaviour that actually clicking on the hover-over text will attempt to open a non-existent URL).

Subgraph of pango XA: (39 samples, 39 shown)

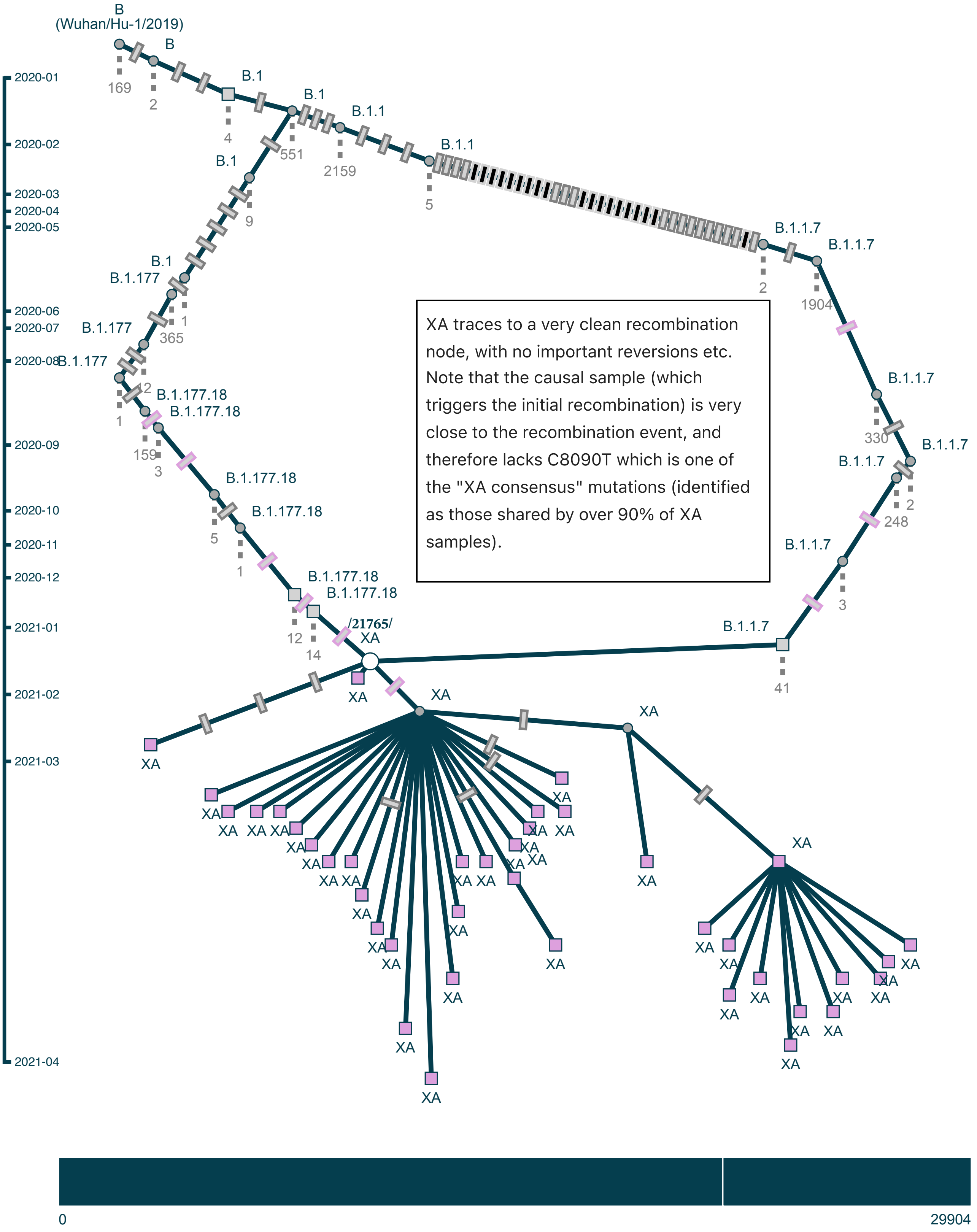

Subgraph of pango XB: (192 samples, 22 shown)

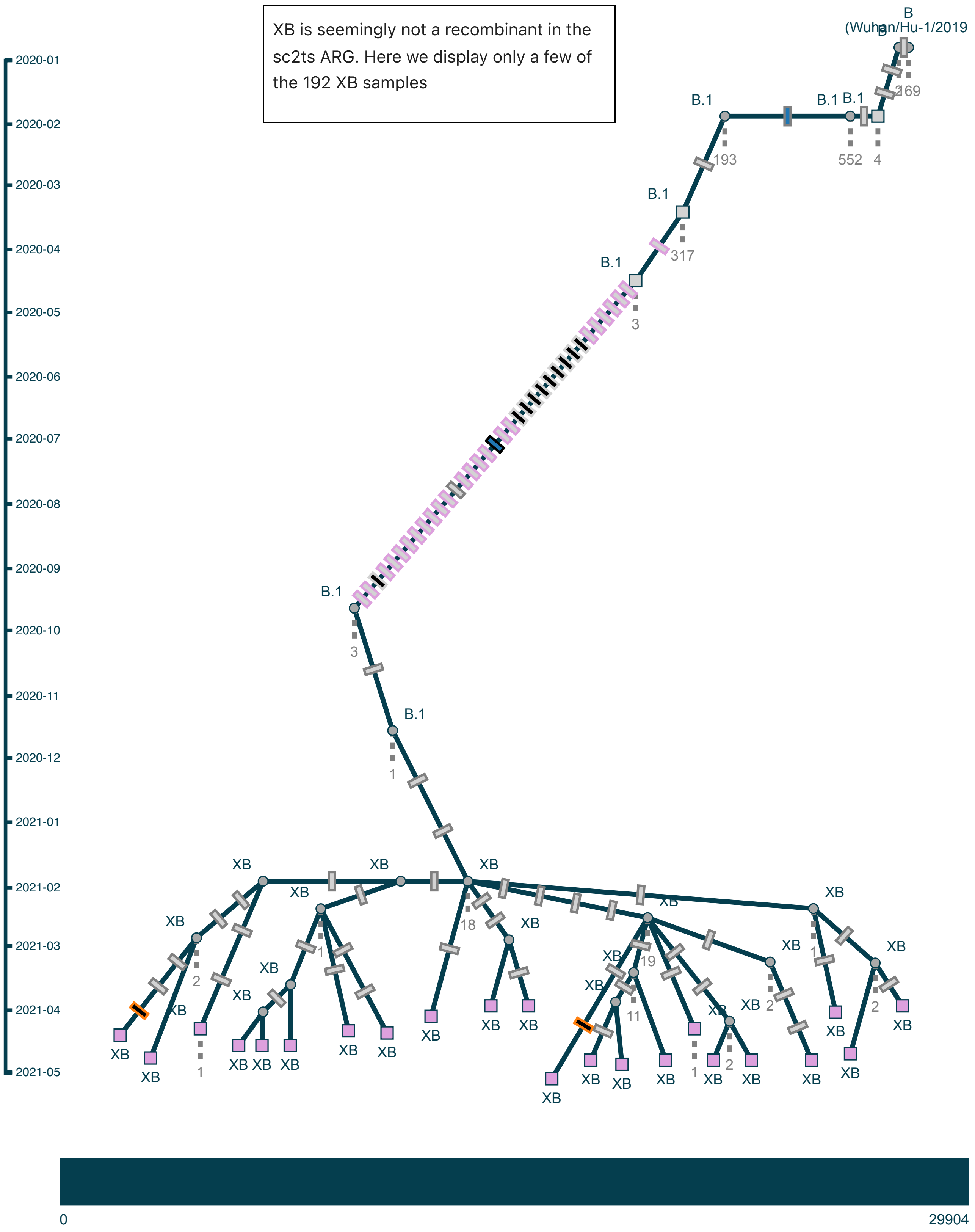

Subgraph of pango XC: (5 samples, 5 shown)

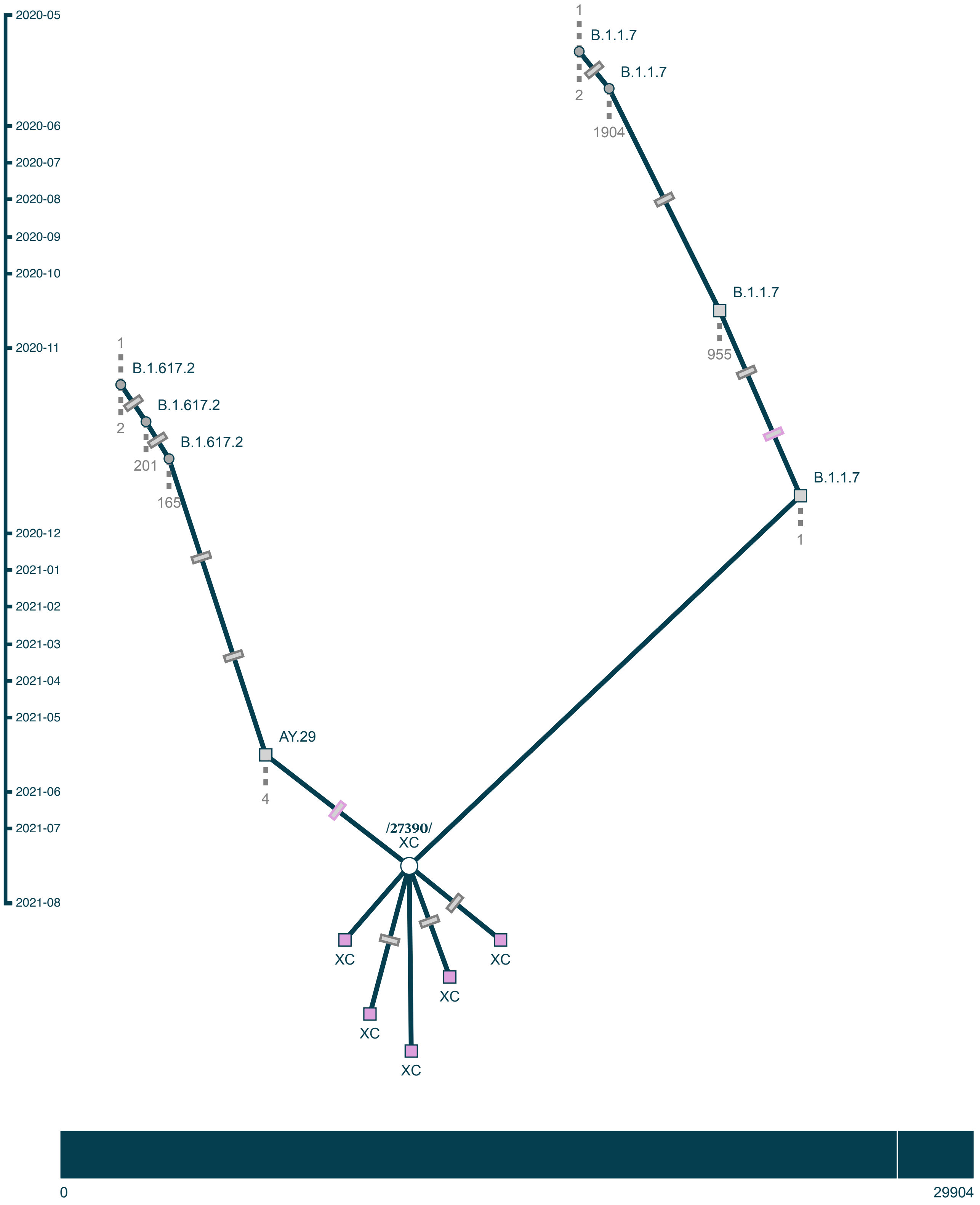

### Subgraph of pango XE/XH: (1118 samples, 22 shown)

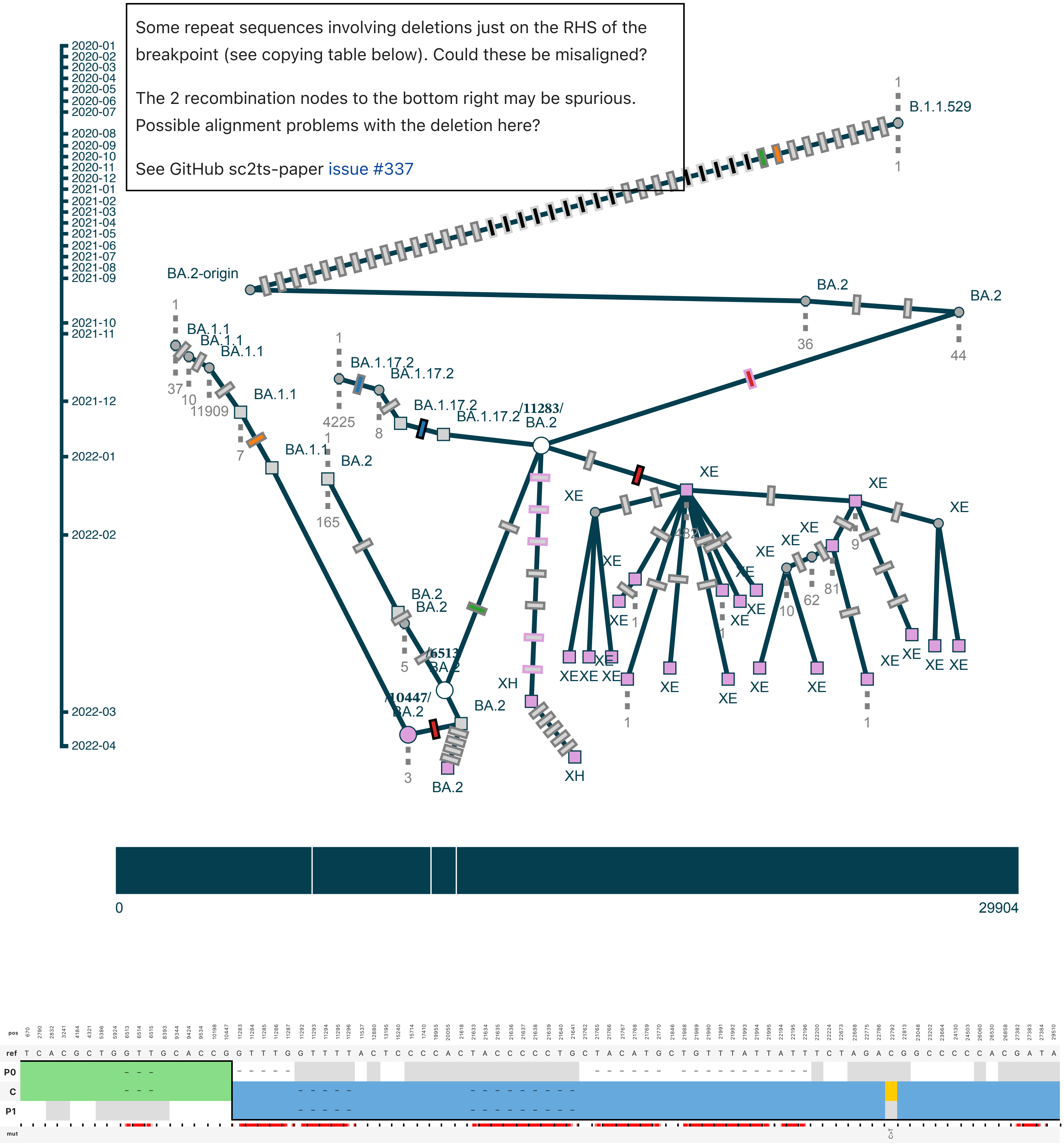

Subgraph of pango XF: (16 samples, 16 shown)

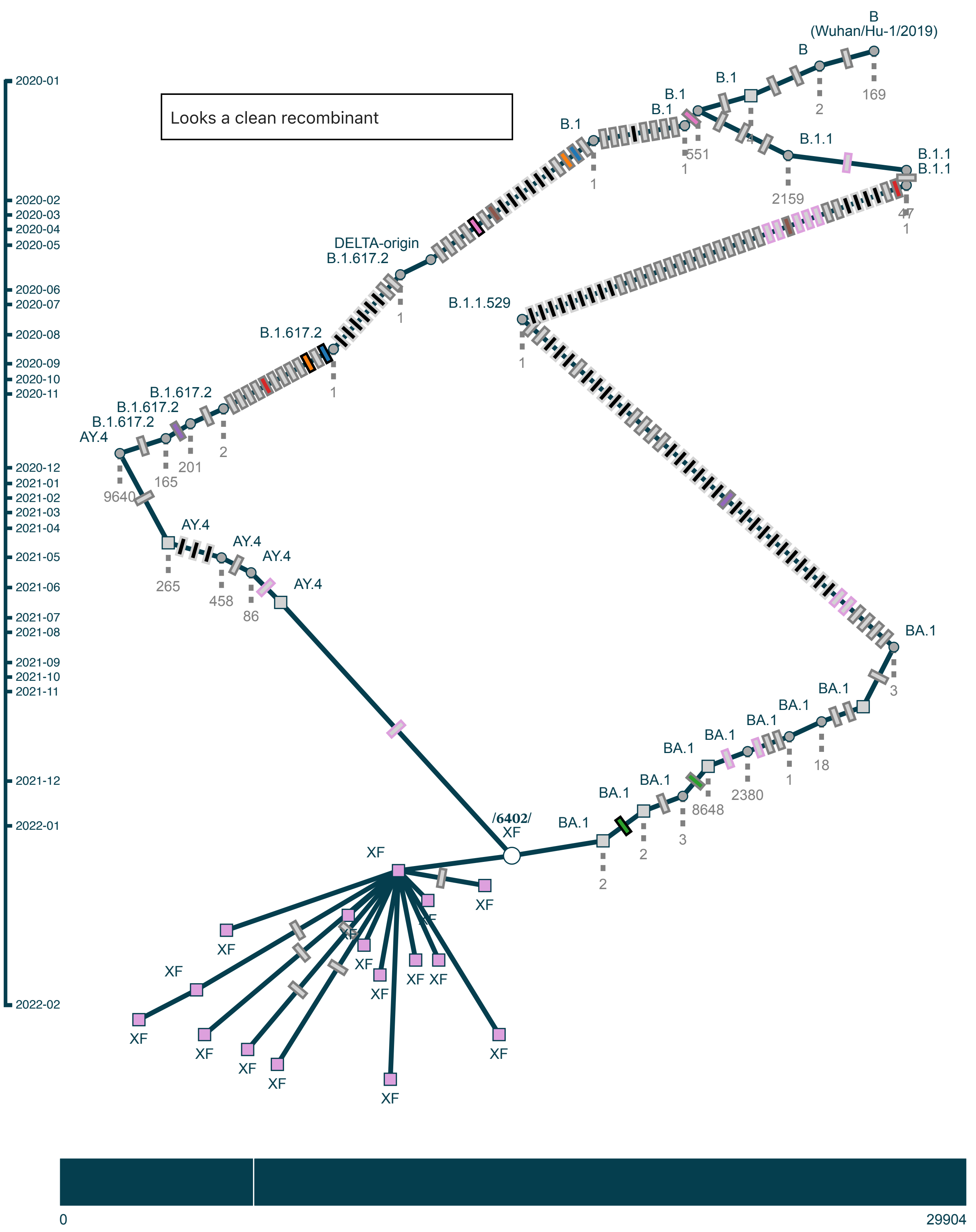

Subgraph of pango XG: (3 samples, 3 shown)

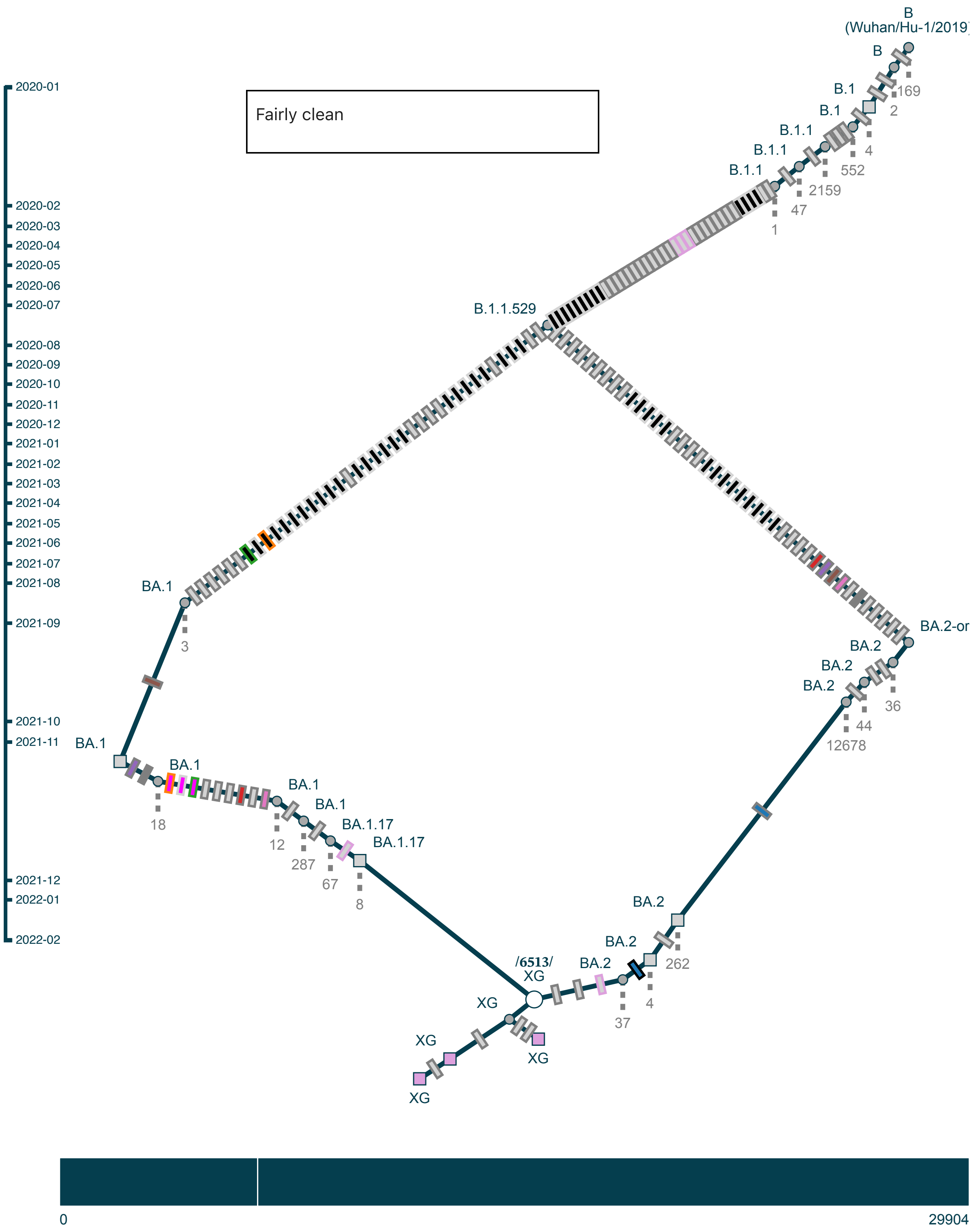

Subgraph of pango XJ: (68 samples, 68 shown)

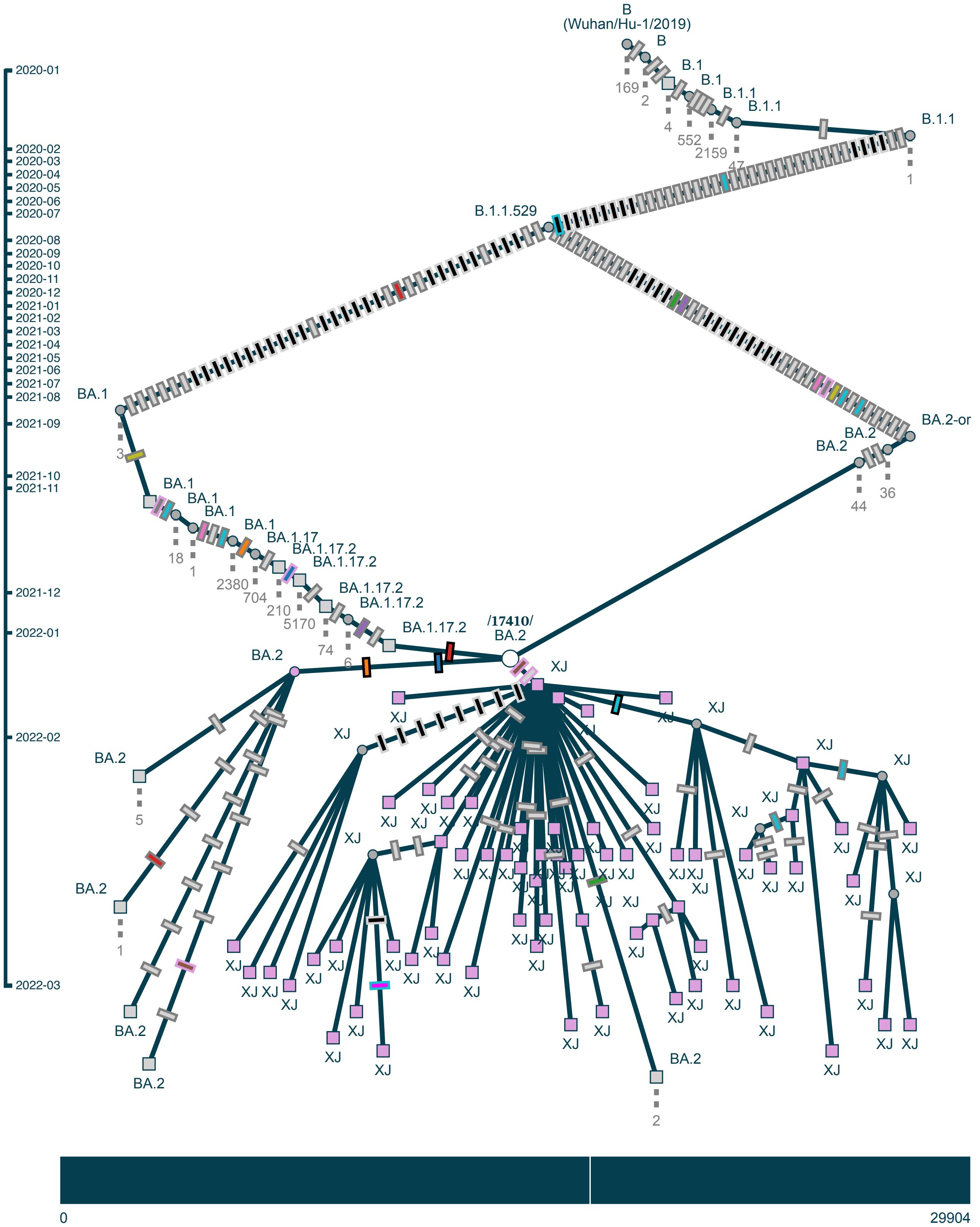

Subgraph of pango XL: (64 samples, 64 shown)

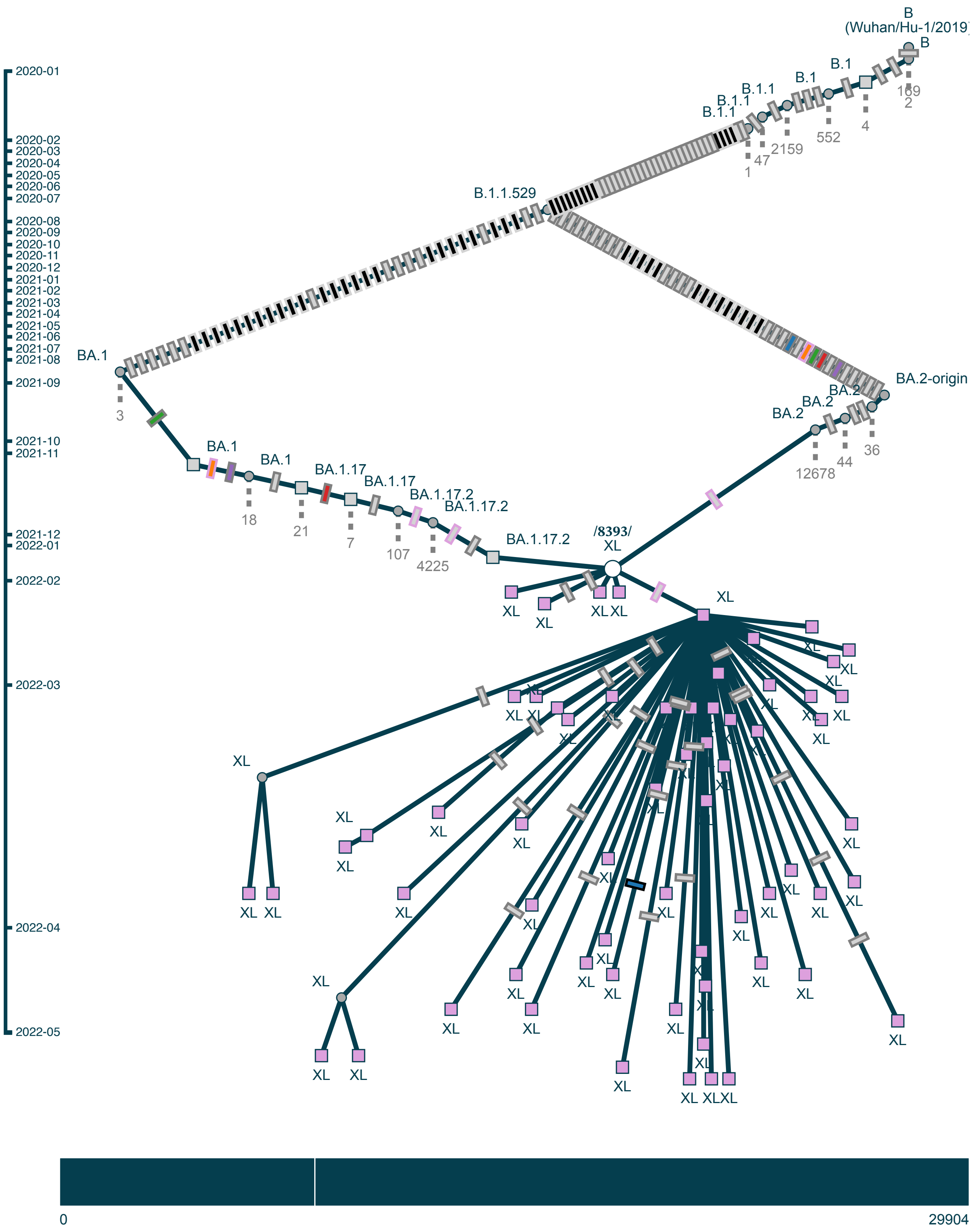

Subgraph of pango XM/XAL: (32 samples, 32 shown)

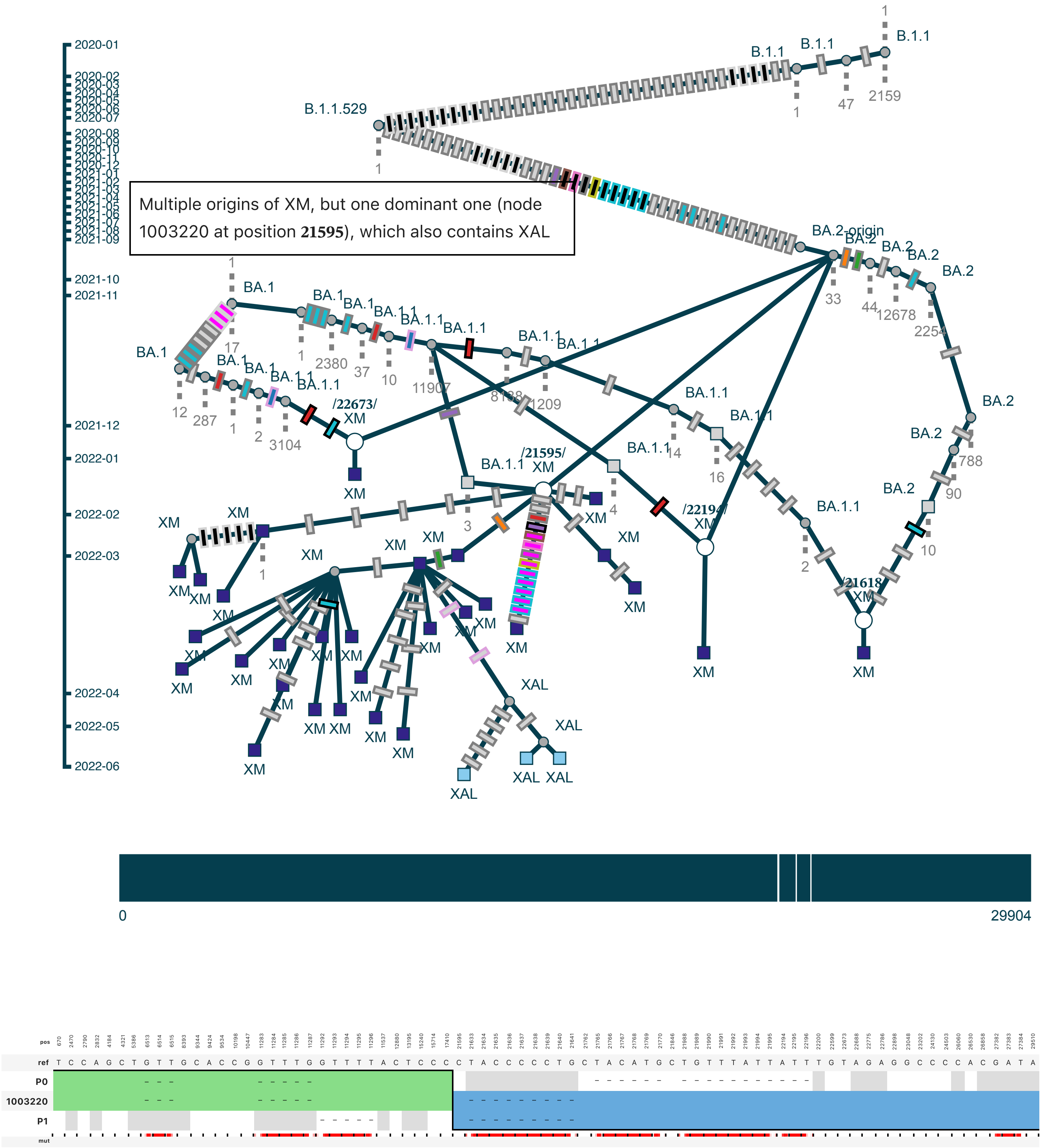

Subgraph of pango XN/XAU: (128 samples, 128 shown)

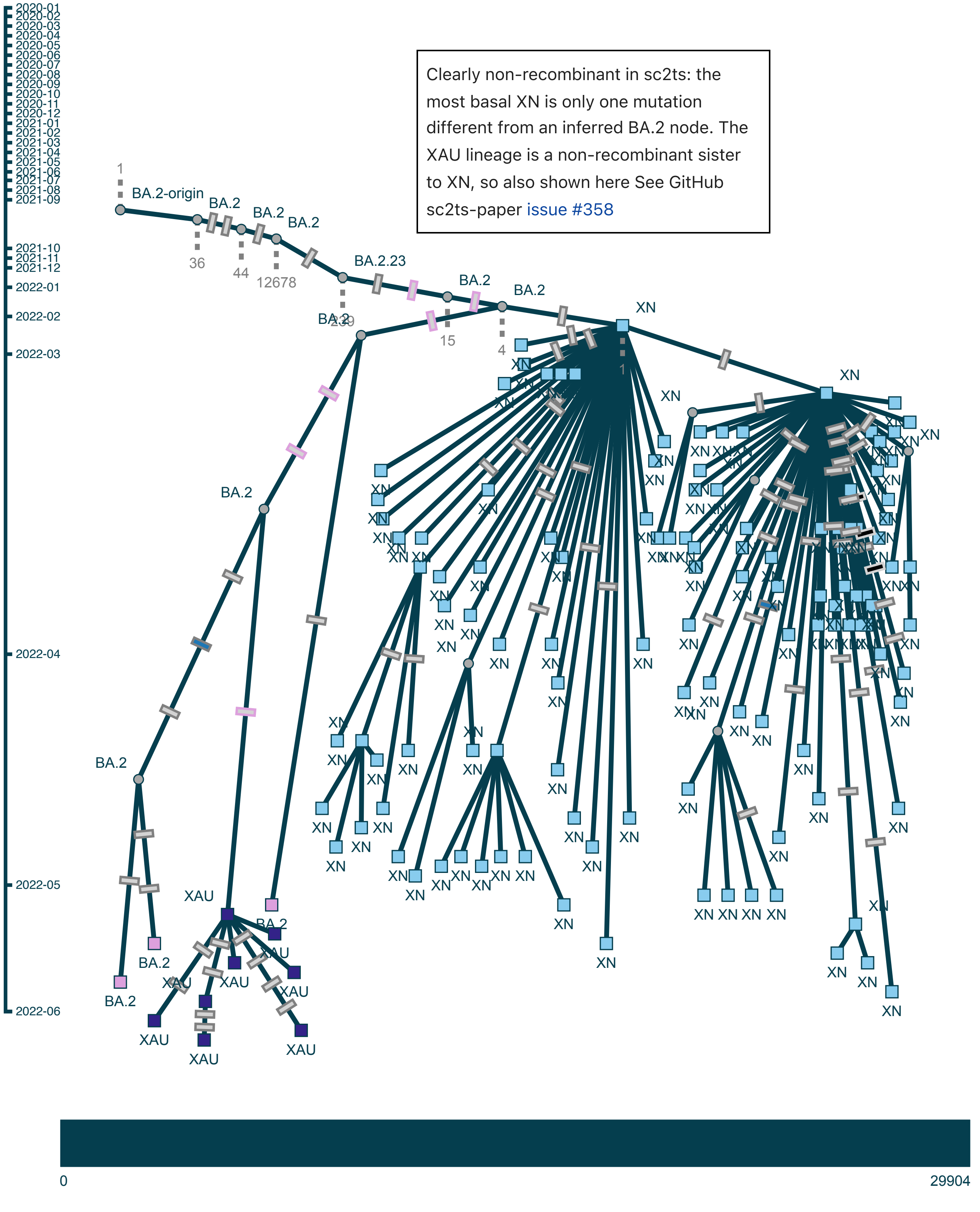

Subgraph of pango XP: (45 samples, 45 shown)

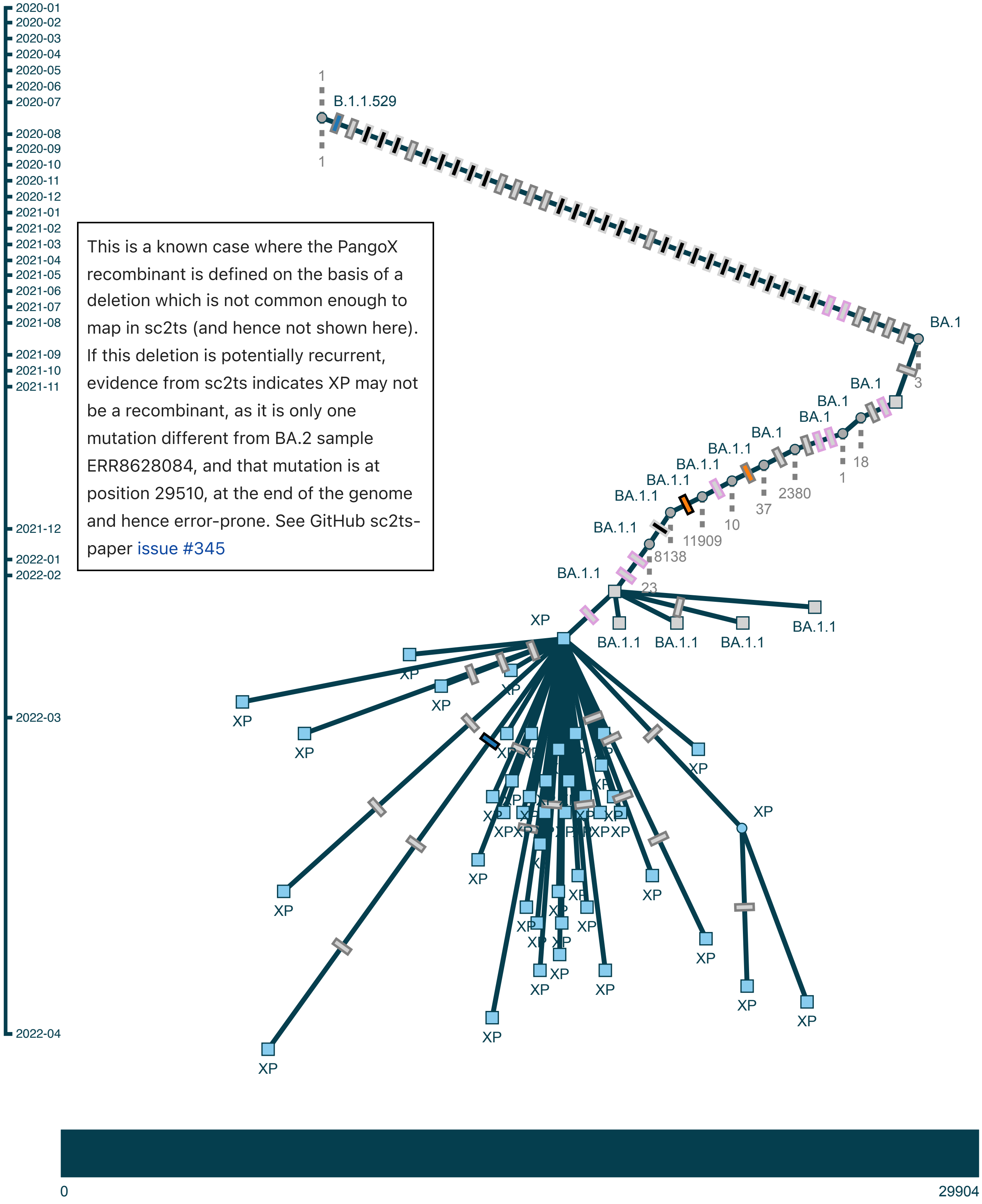

Subgraph of pango XQ/XR/XU/XAA/XAG/XAM: (117 samples, 49 shown)

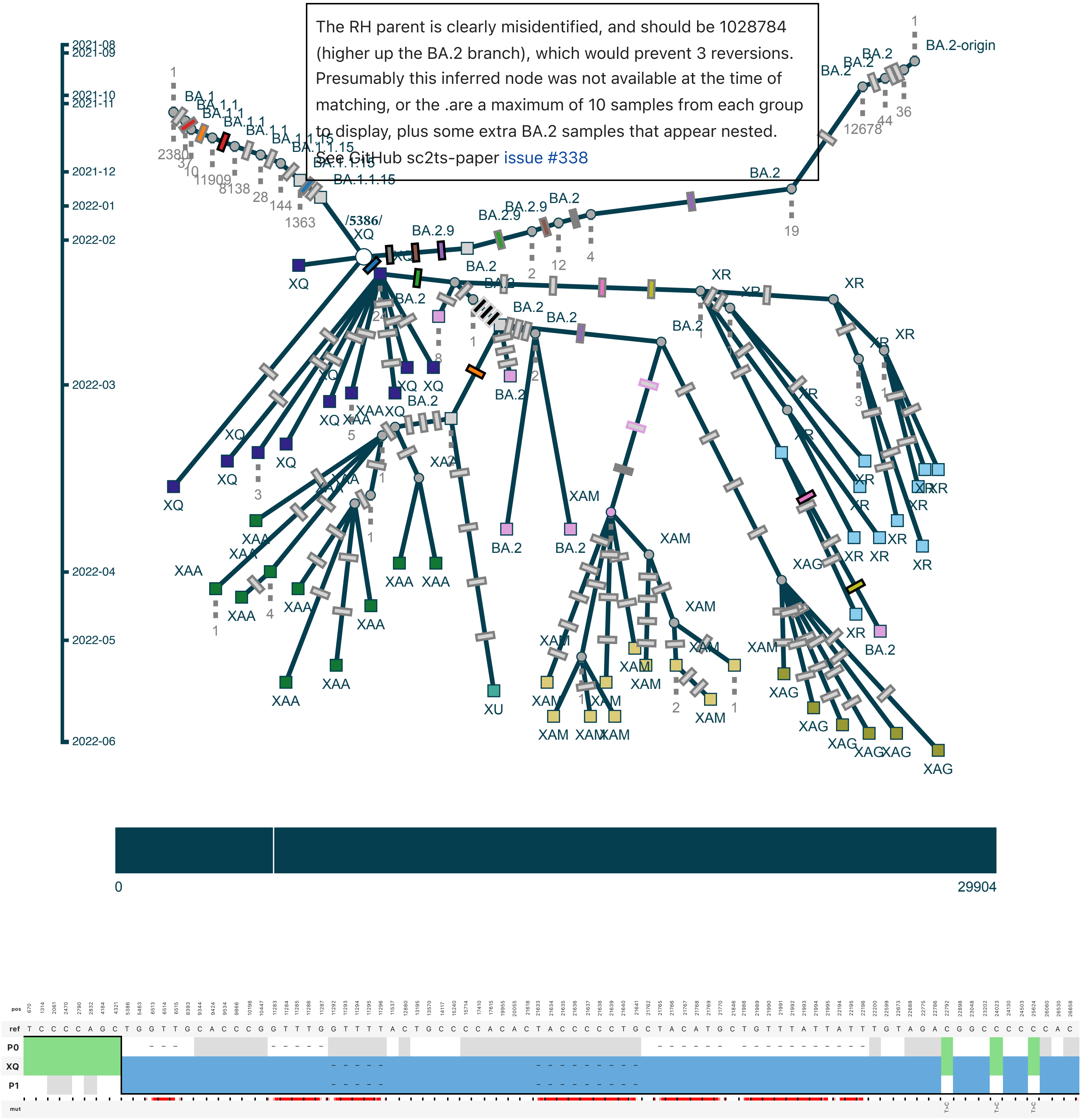

#### Subgraph of pango XS: (17 samples, 17 shown)

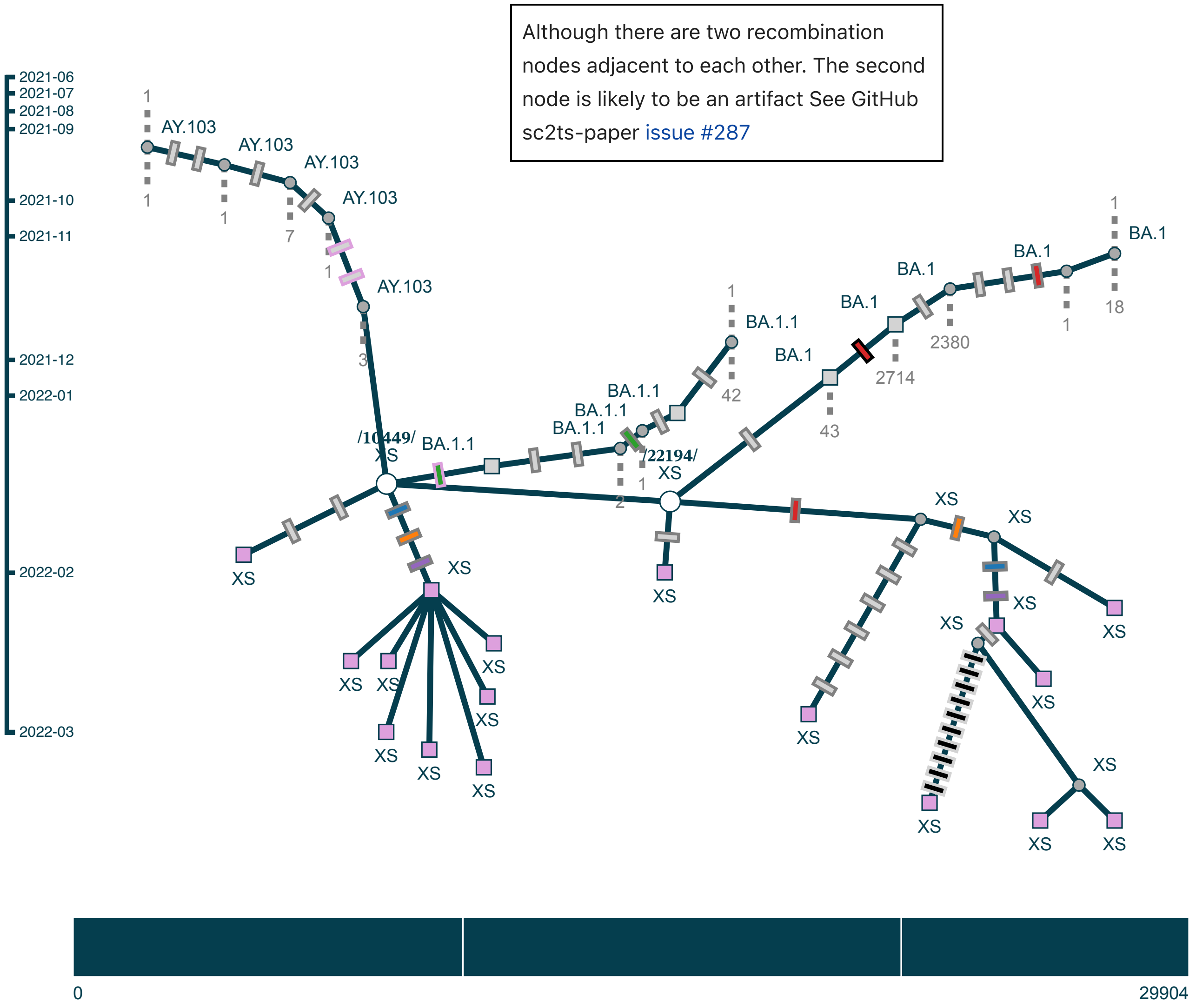

Subgraph of pango XW: (32 samples, 32 shown)

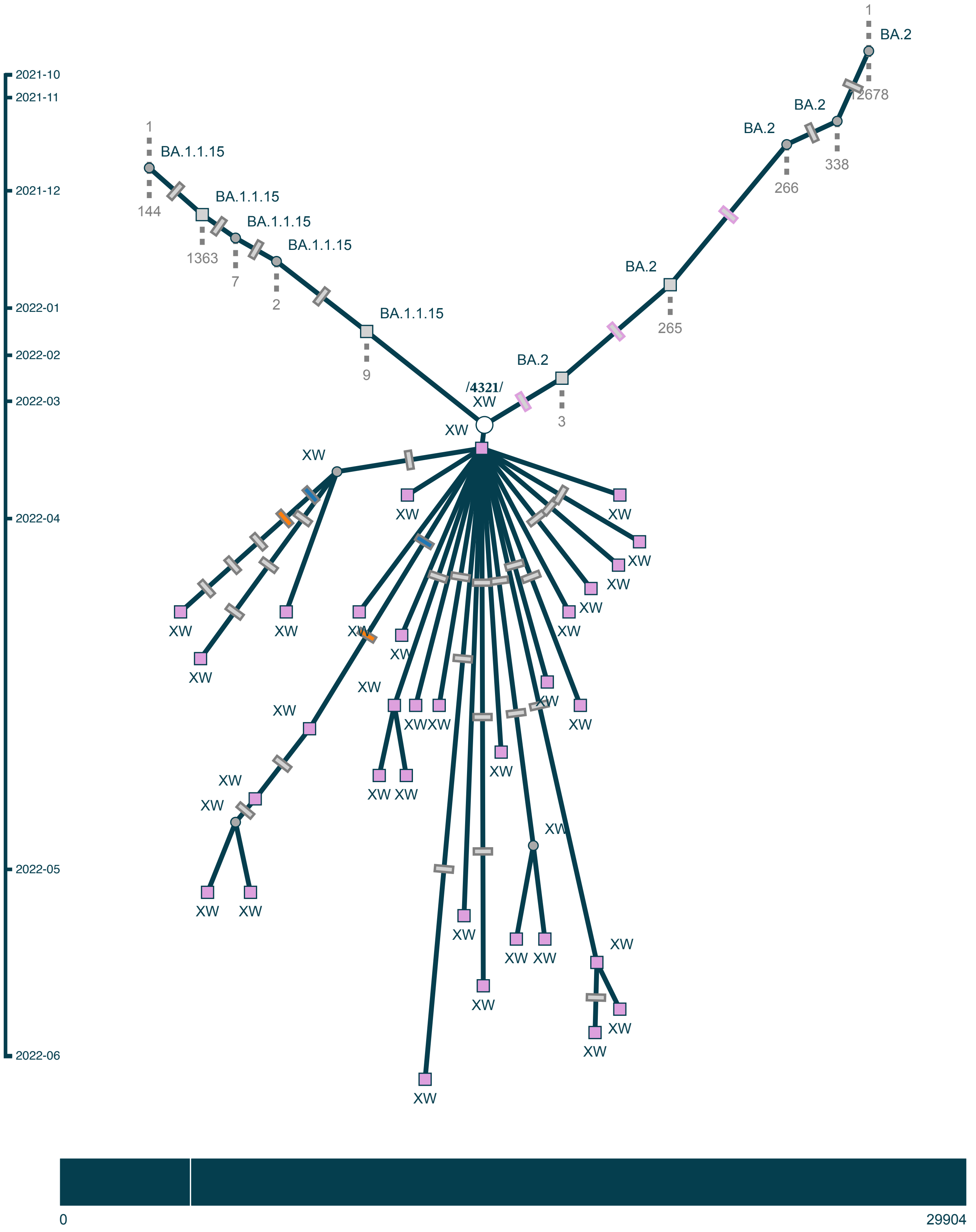

Subgraph of pango XY: (23 samples, 23 shown)

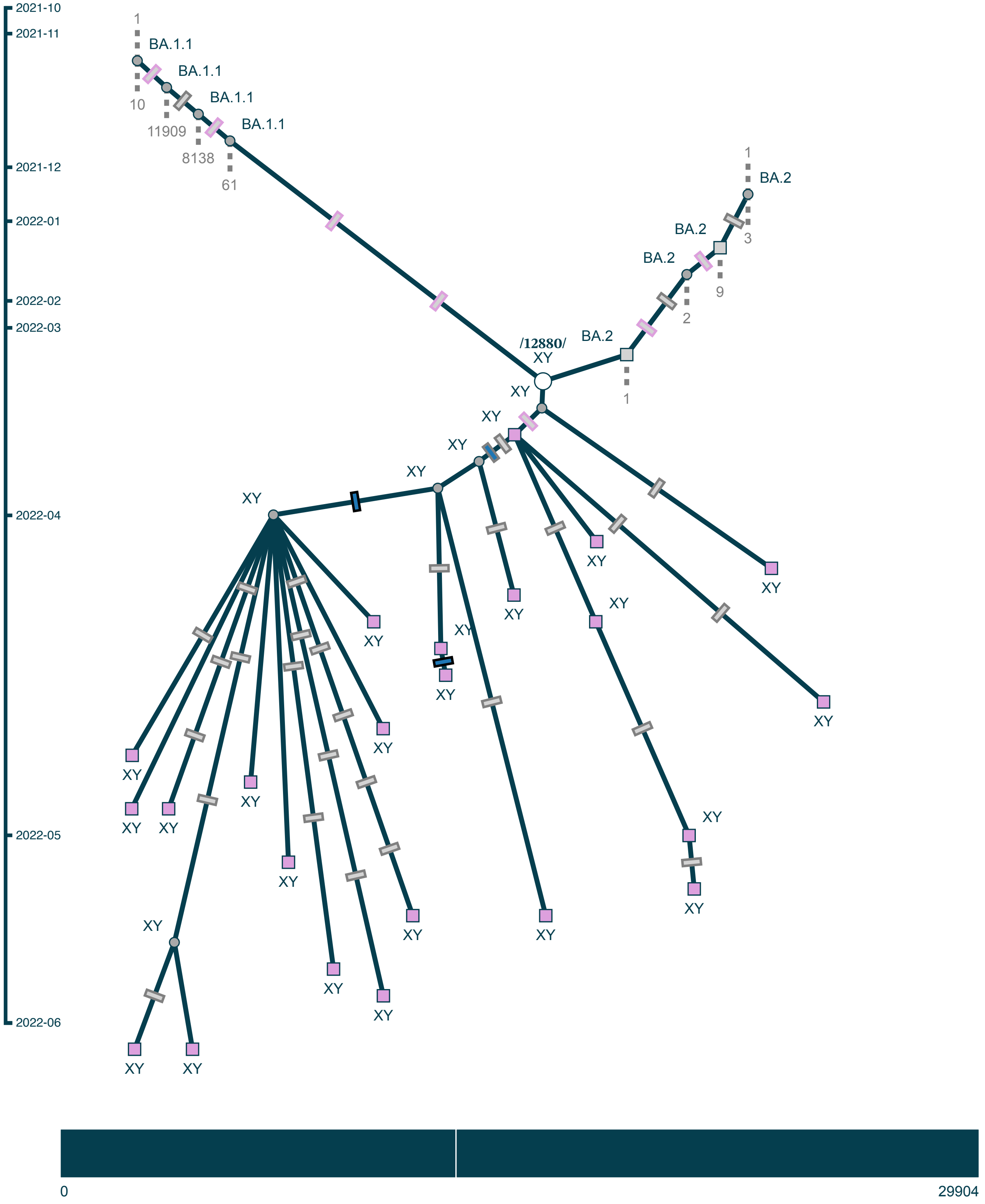

Subgraph of pango XZ/XAC/XAD/XAE/XAP: (97 samples, 97 shown)

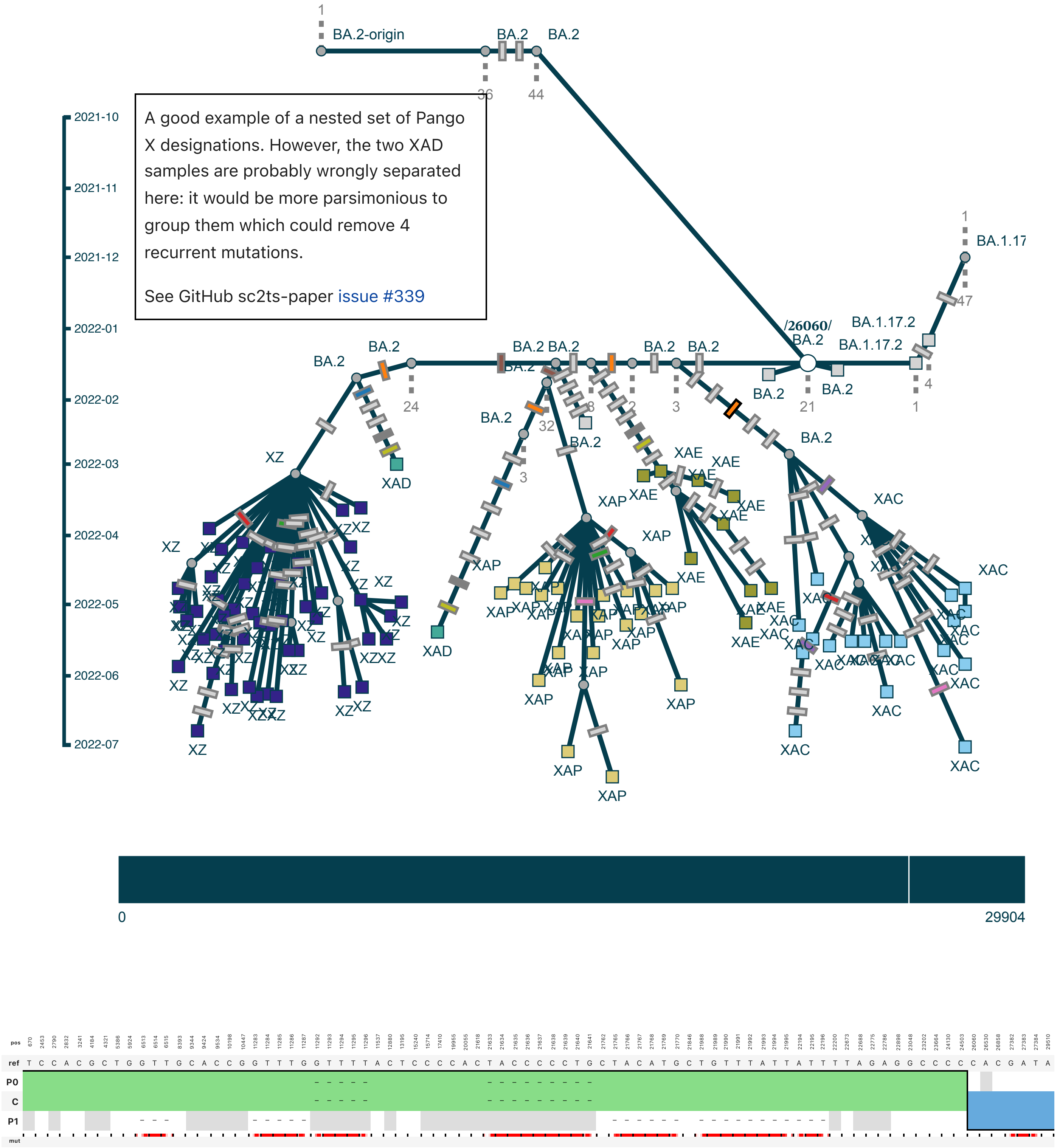

Subgraph of pango XAF: (1 sample, 1 shown)

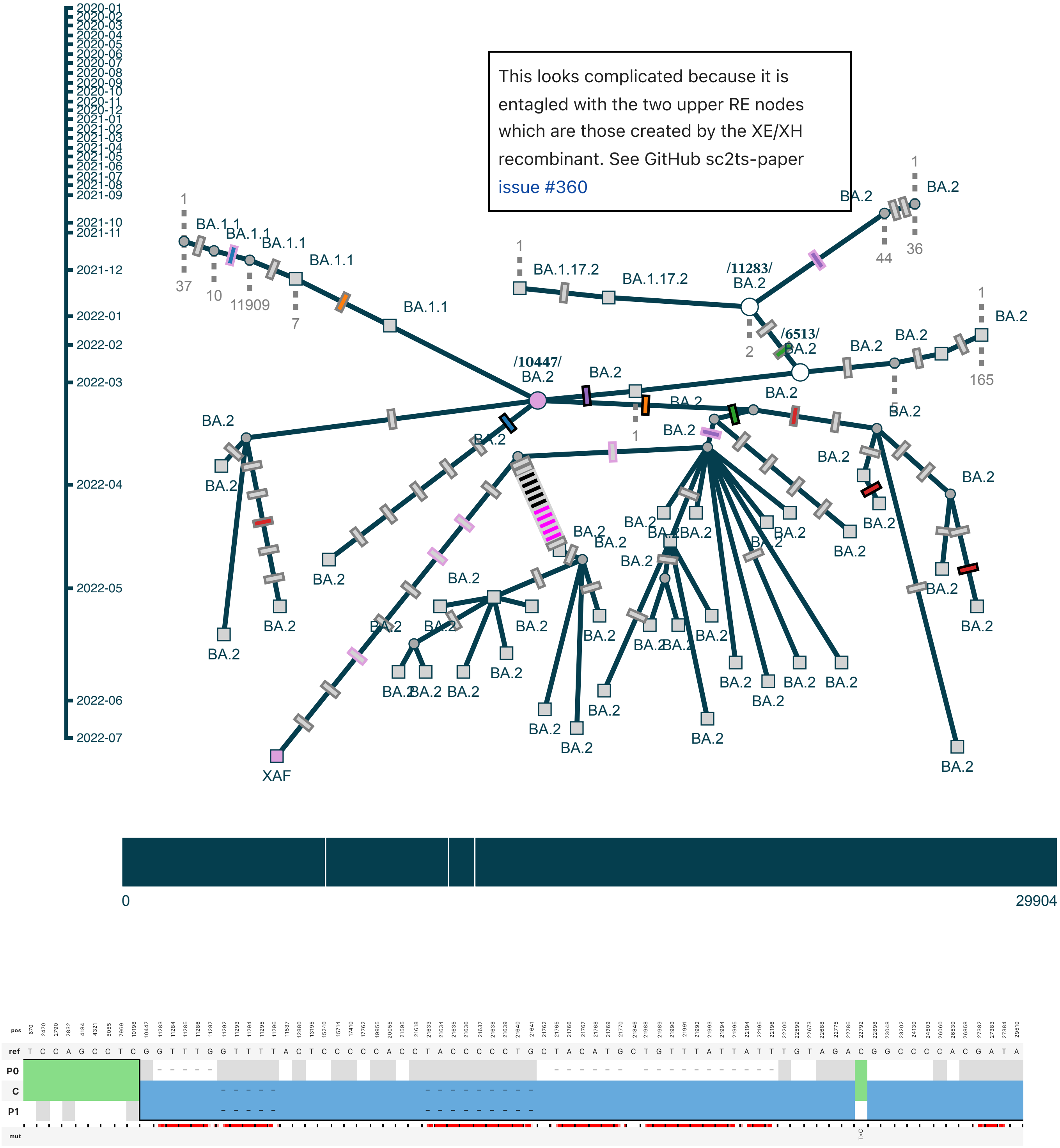

Subgraph of pango XAJ: (18 samples, 18 shown)

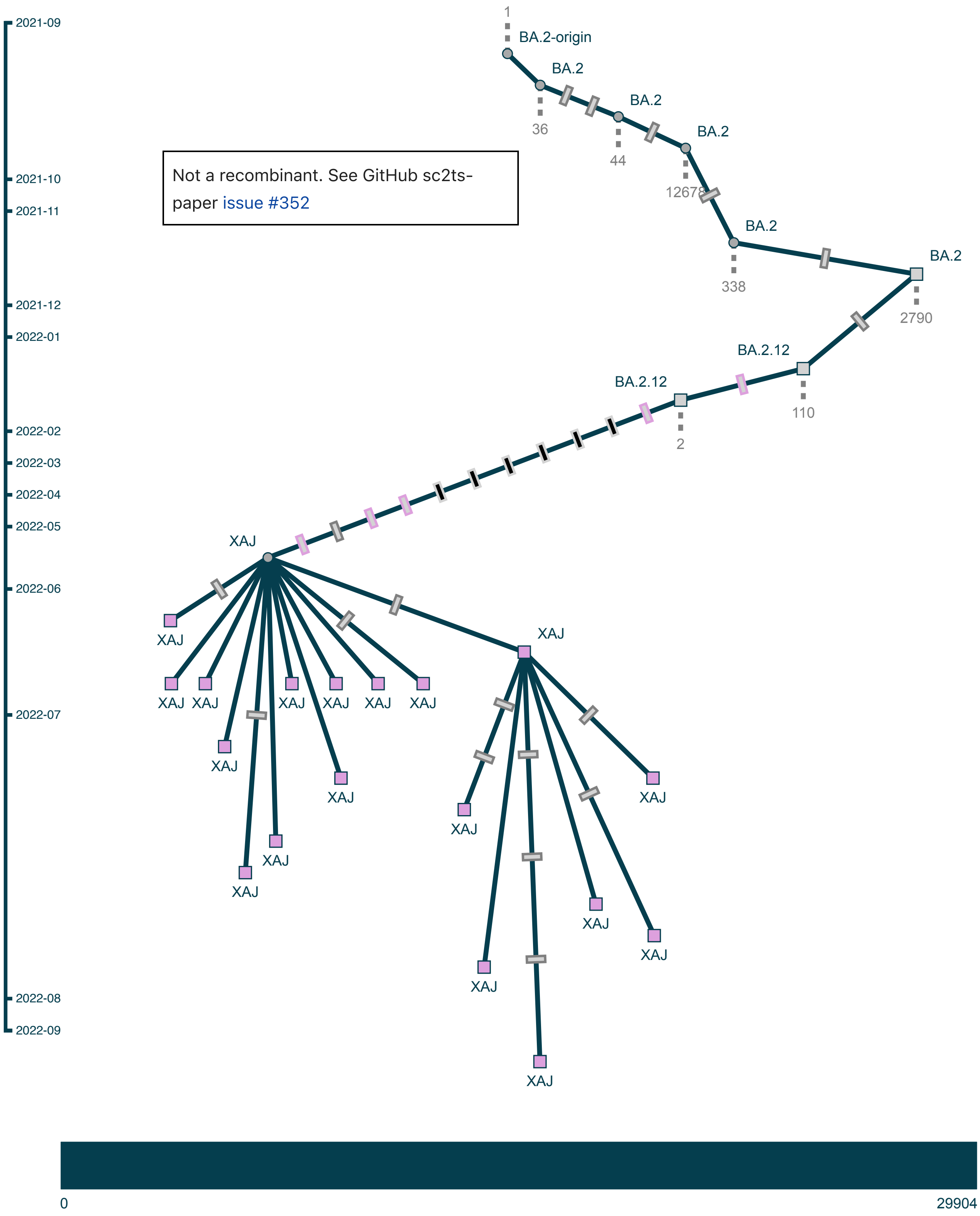

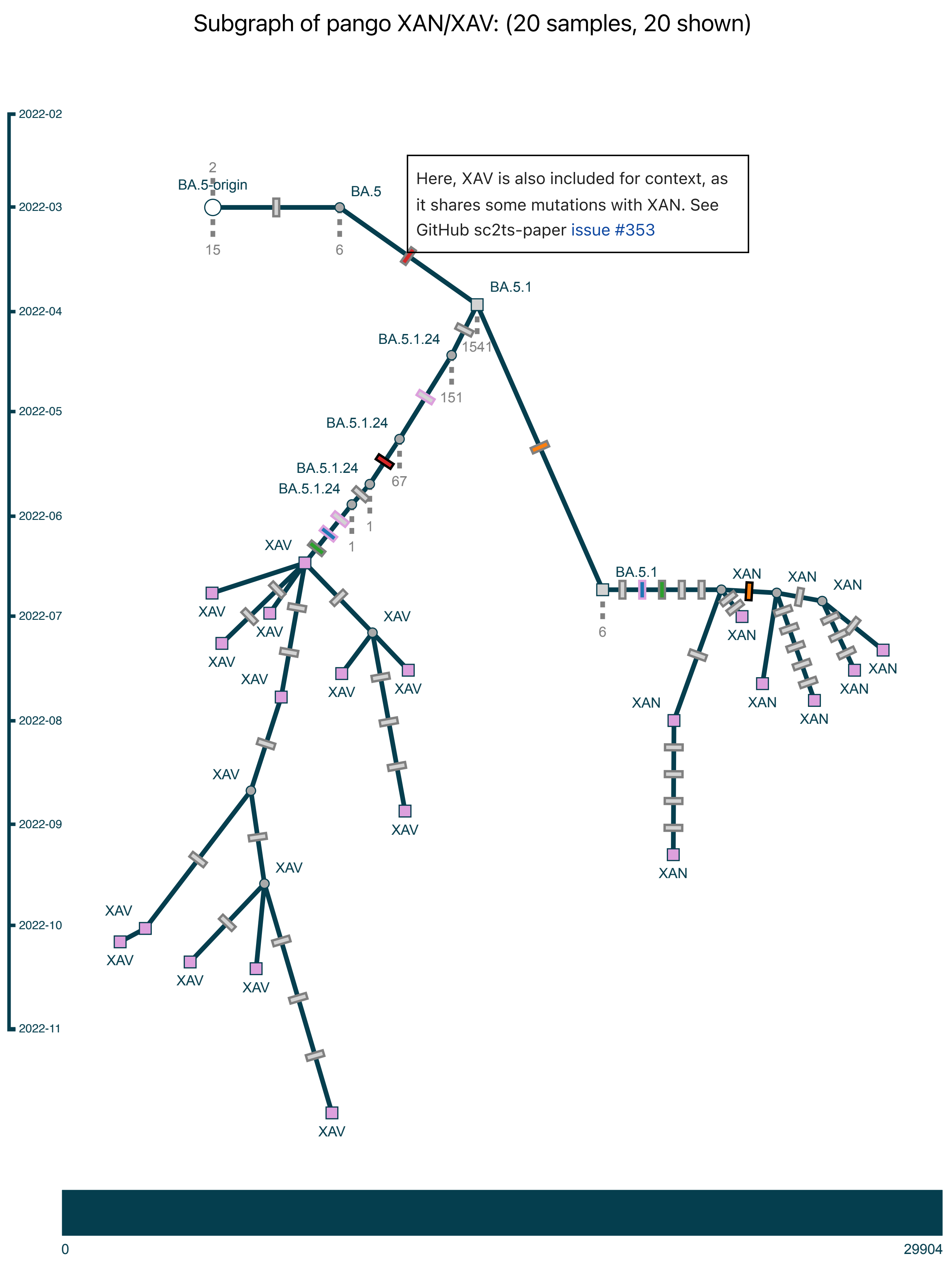

Subgraph of pango XAN/XAV: (20 samples, 20 shown)
2022-02
2022-03
2022-04
2022-05
2022-06
2022-07
2022-08
2022-09
2022-10
2022-11
BA.5-origin
BA.5
BA.5.1
BA.5.1.24
BA.5.1.24
BA.5.1.24
BA.5.1.24
2
15
6
1541
151
67
1
1
6
Here, XAV is also included for context, as it shares some mutations with XAN. See GitHub sc2ts-paper issue #353
XAV
BA.5.1
XAN
0
29904

Subgraph of pango XAS: (77 samples, 77 shown)

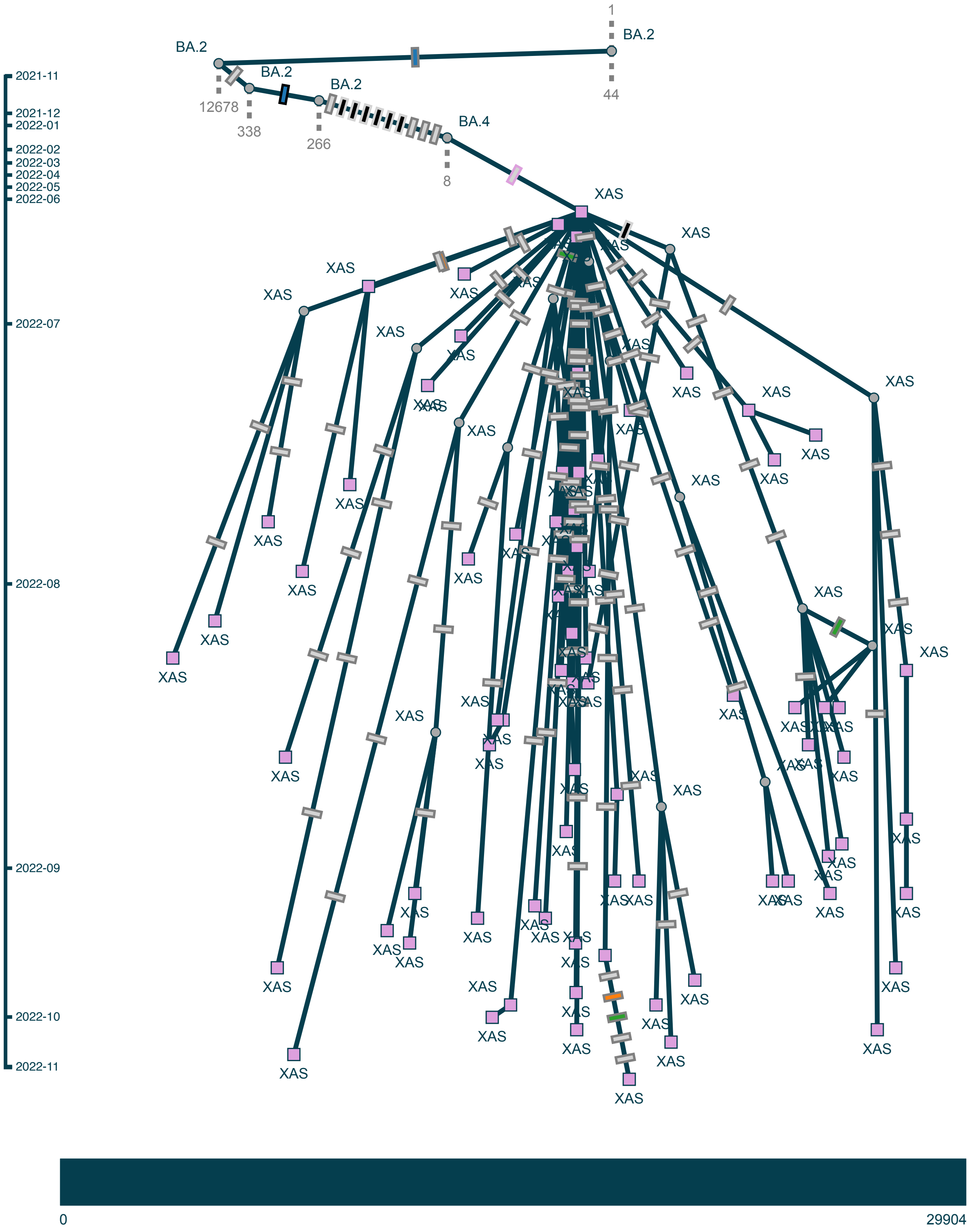

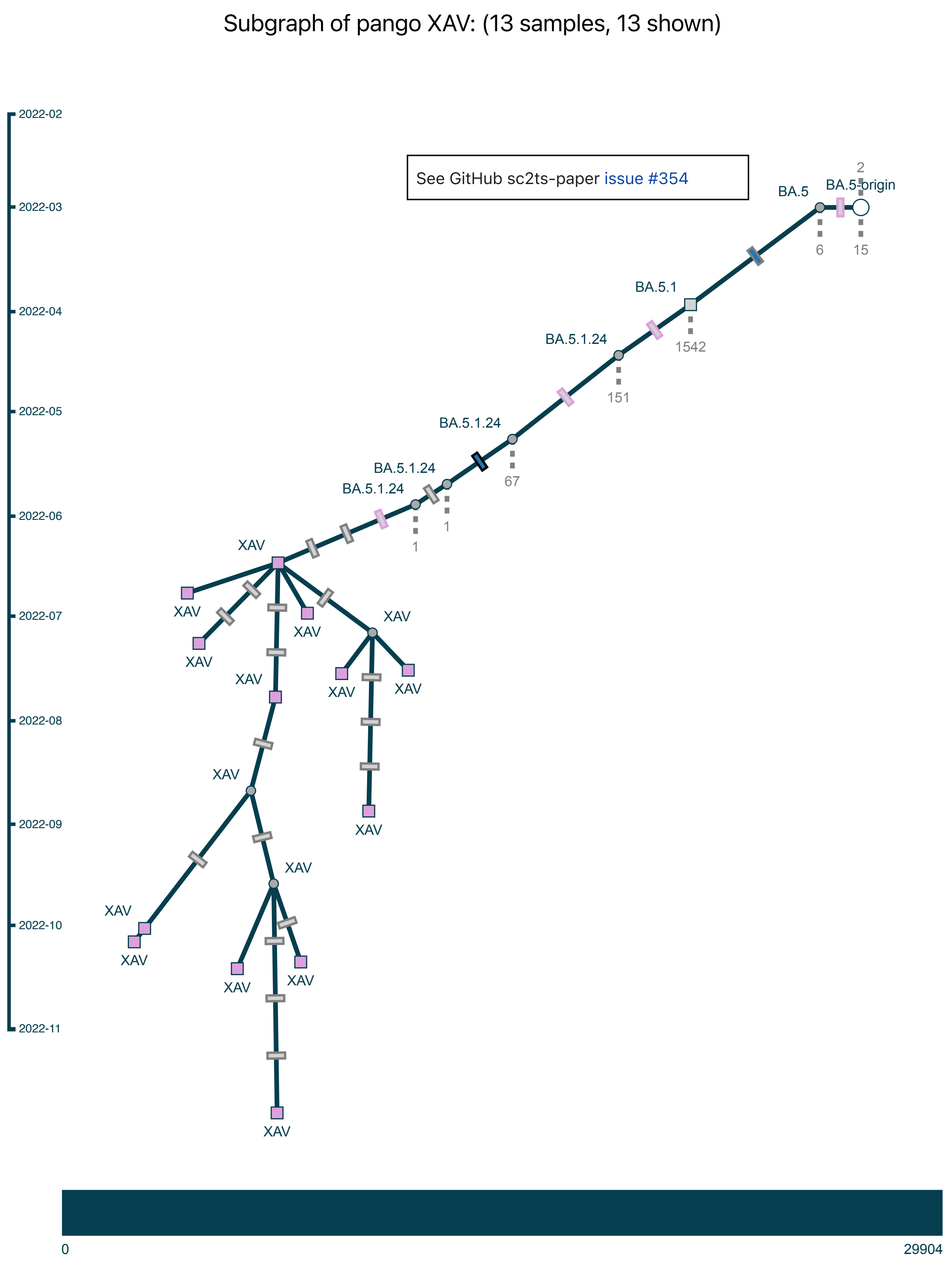
Subgraph of pango XAV: (13 samples, 13 shown)
See GitHub sc2ts-paper issue #354
2022-02
2022-03
2022-04
2022-05
2022-06
2022-07
2022-08
2022-09
2022-10
2022-11
BA.5
BA.5-origin
2
6
15
BA.5.1
1542
BA.5.1.24
151
BA.5.1.24
67
BA.5.1.24
1
BA.5.1.24
1
XAV
0
29904

Subgraph of pango XAZ: (133 samples, 20 shown)

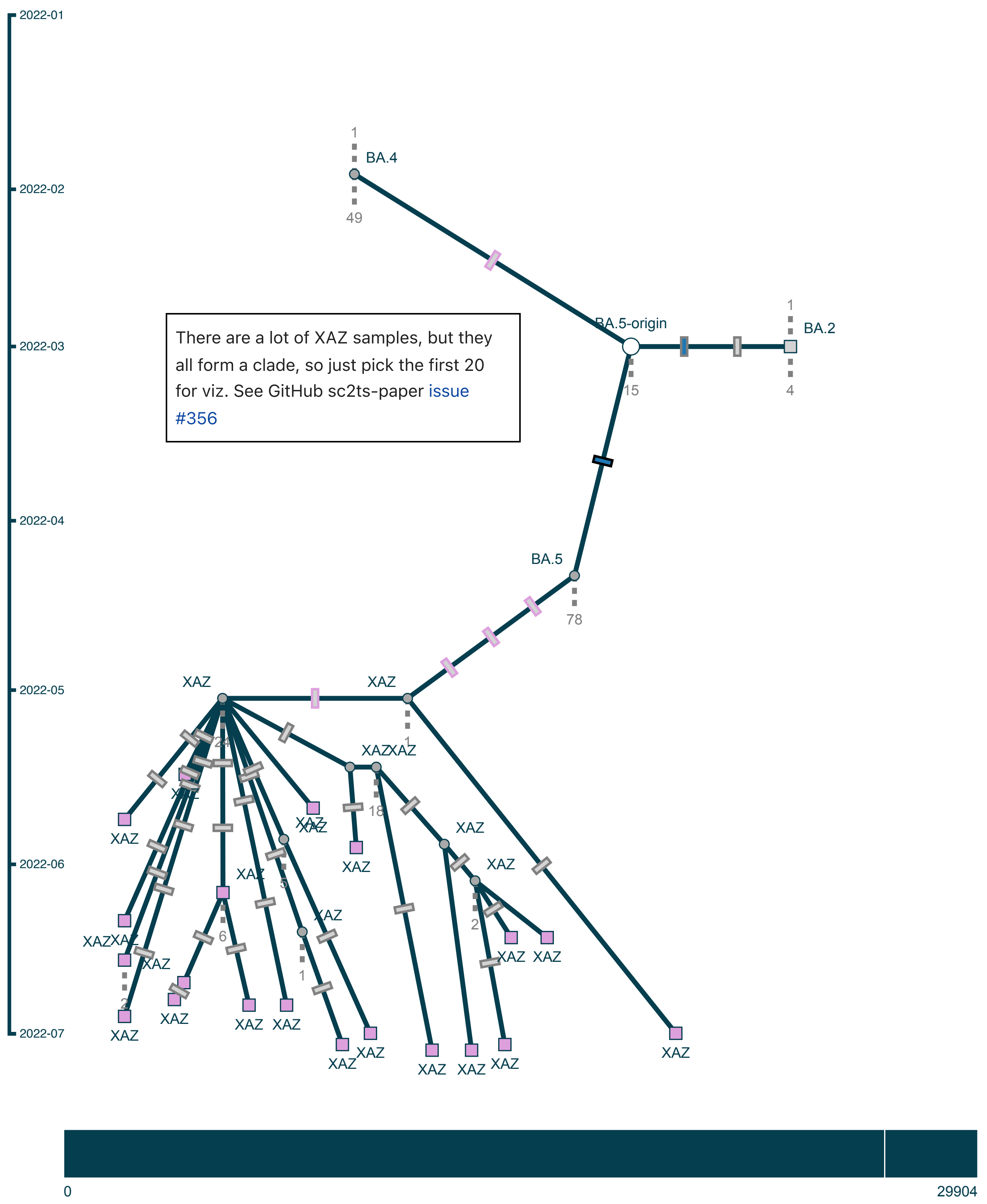

Subgraph of pango XBB: (71 samples, 20 shown)

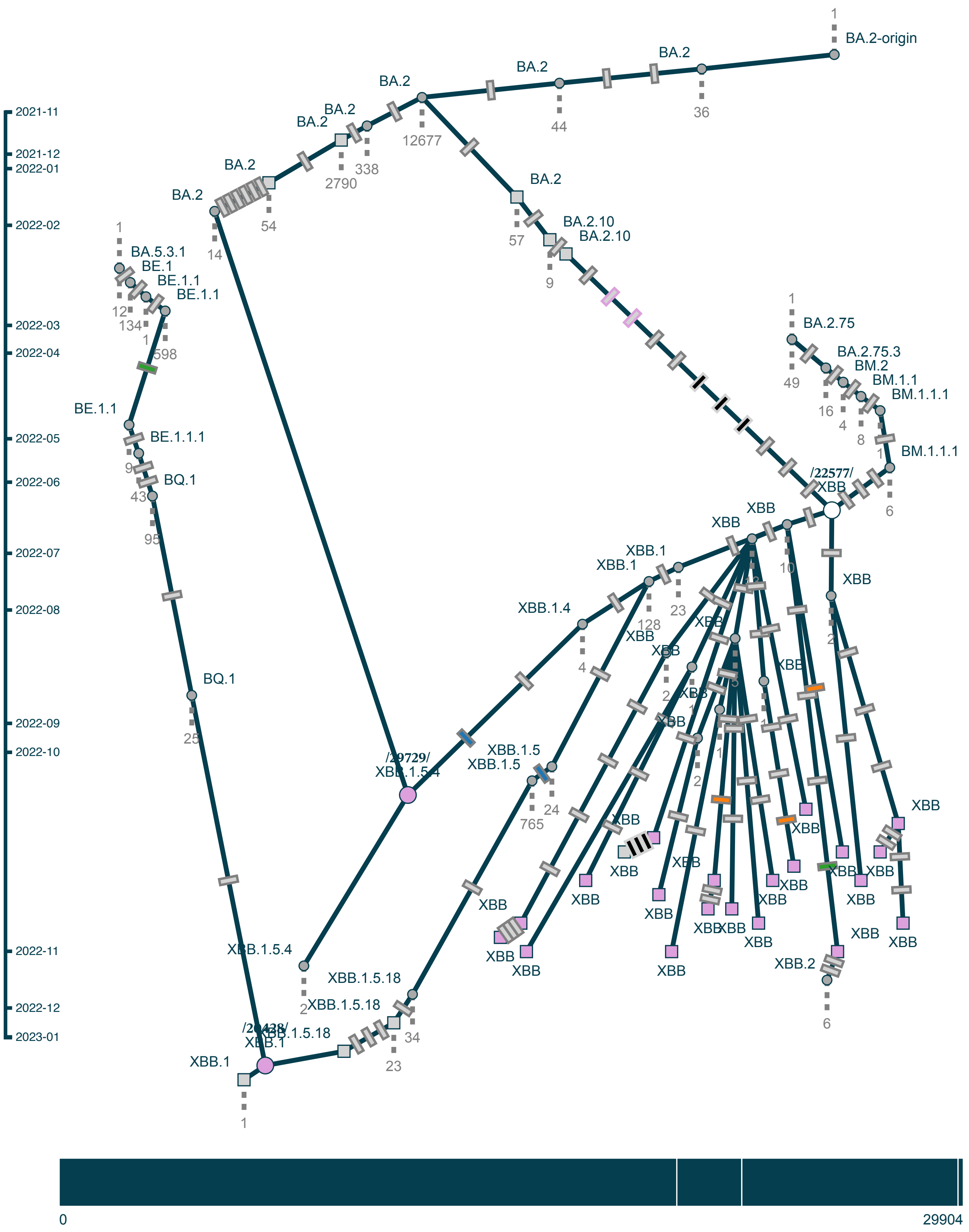

Subgraph of pango XBD: (30 samples, 30 shown)

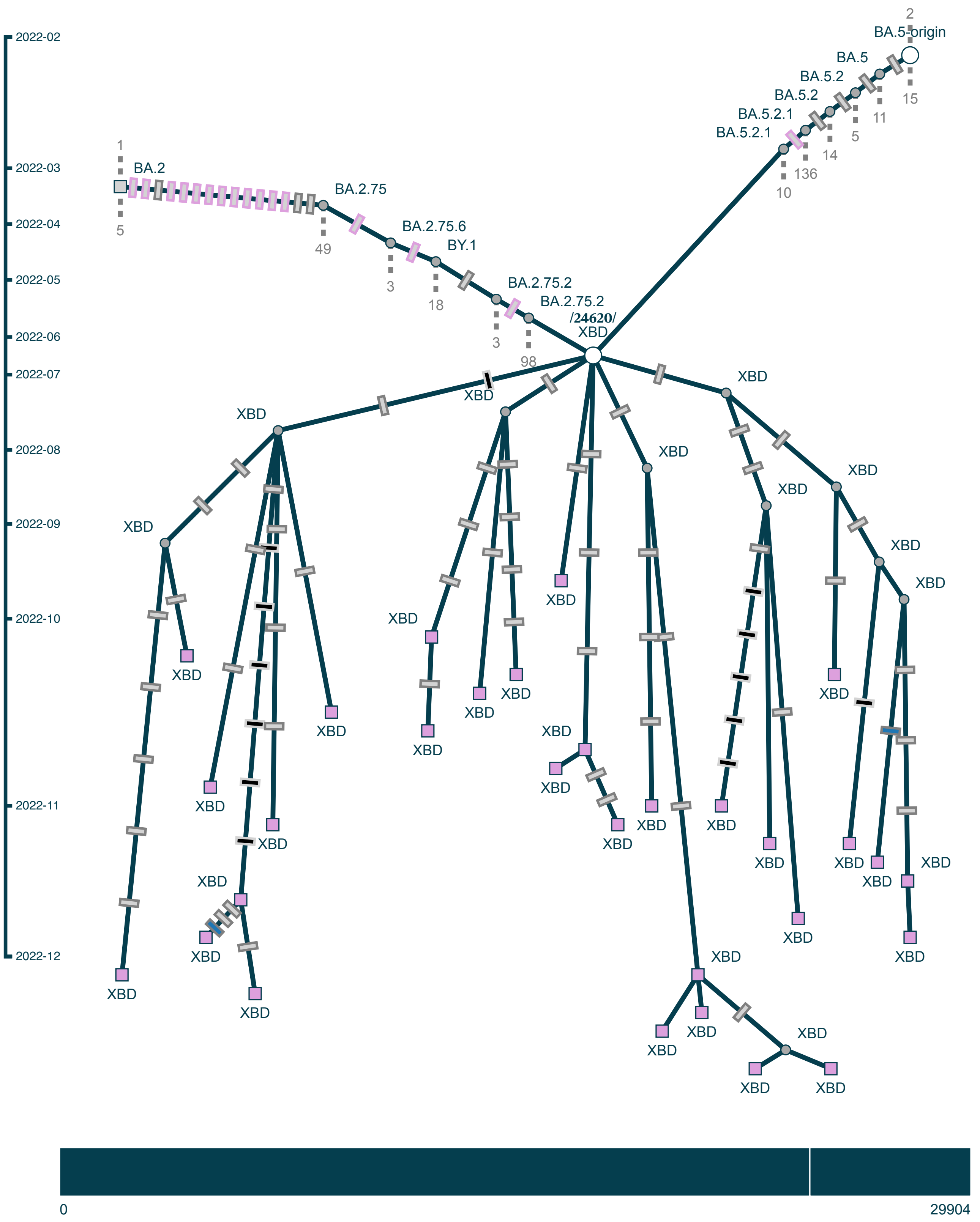

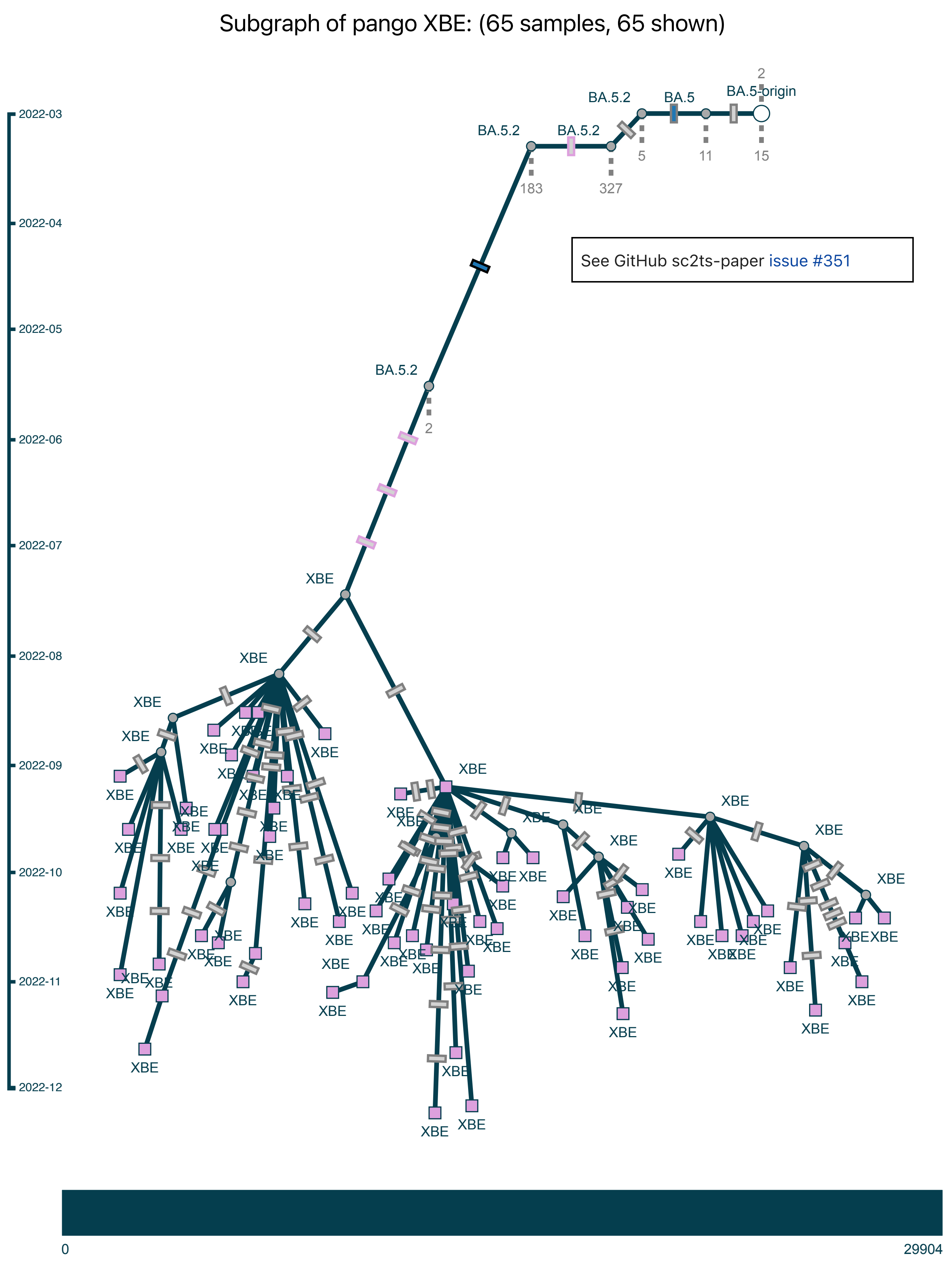

Subgraph of pango XBE: (65 samples, 65 shown)
See GitHub sc2ts-paper issue #351
BA.5-origin
BA.5
BA.5.2
XBE
2022-03
2022-04
2022-05
2022-06
2022-07
2022-08
2022-09
2022-10
2022-11
2022-12
0
29904

Subgraph of pango XBF: (124 samples, 124 shown)

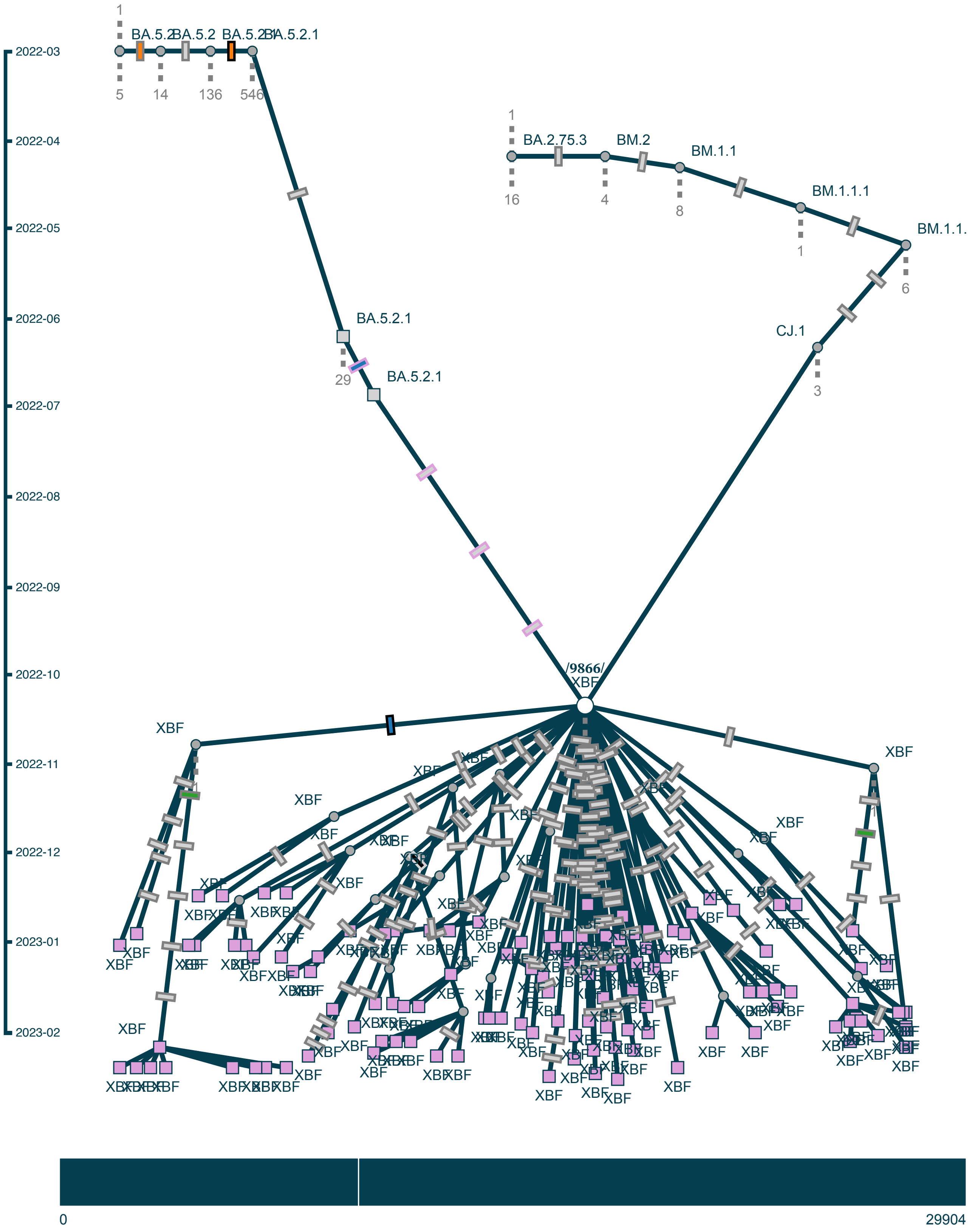

Subgraph of pango XBG: (25 samples, 25 shown)

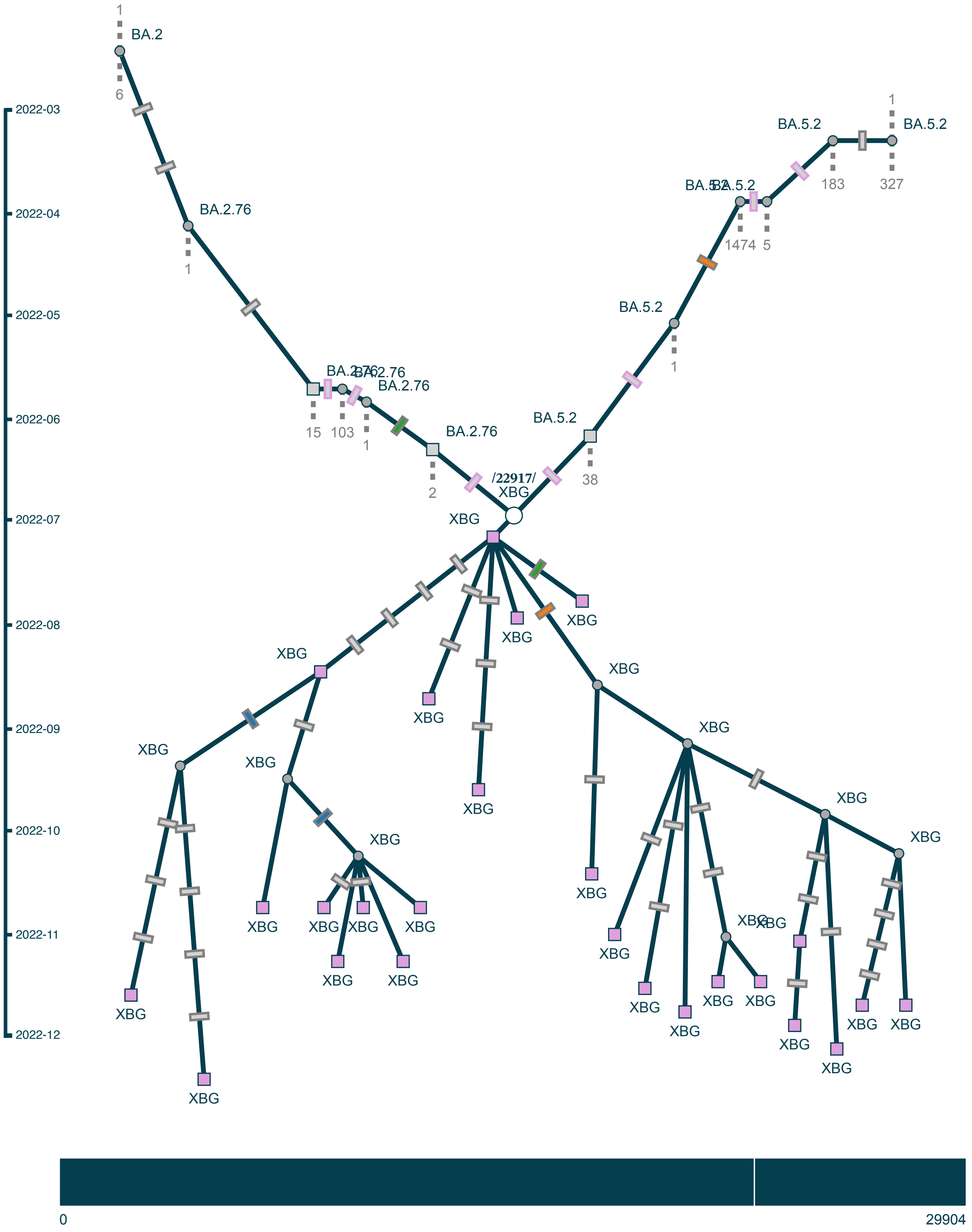

Subgraph of pango XBH: (2 samples, 2 shown)

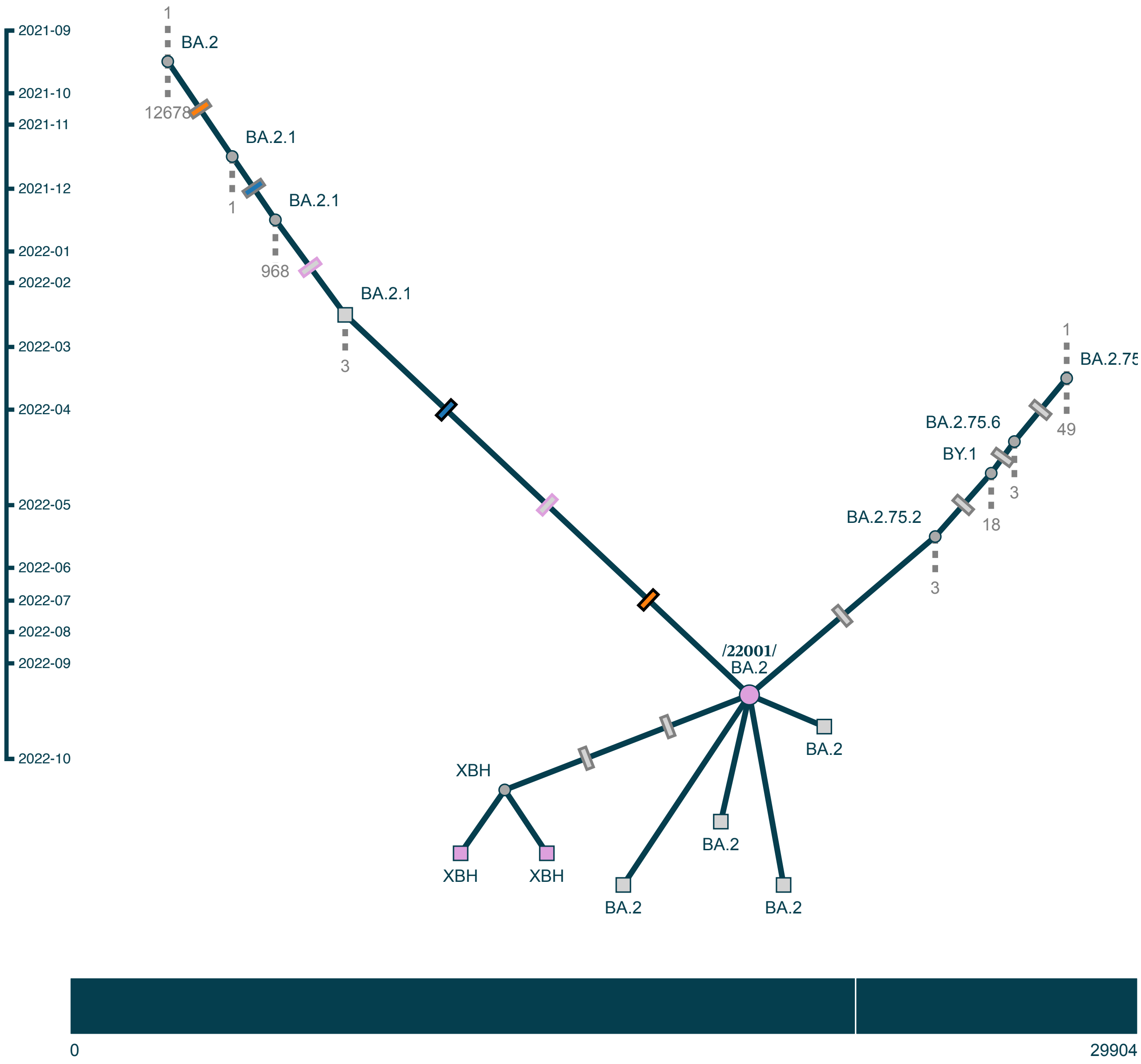

Subgraph of pango XBK/XBK.1/XBQ/CJ.1.3: (27 samples, 27 shown)

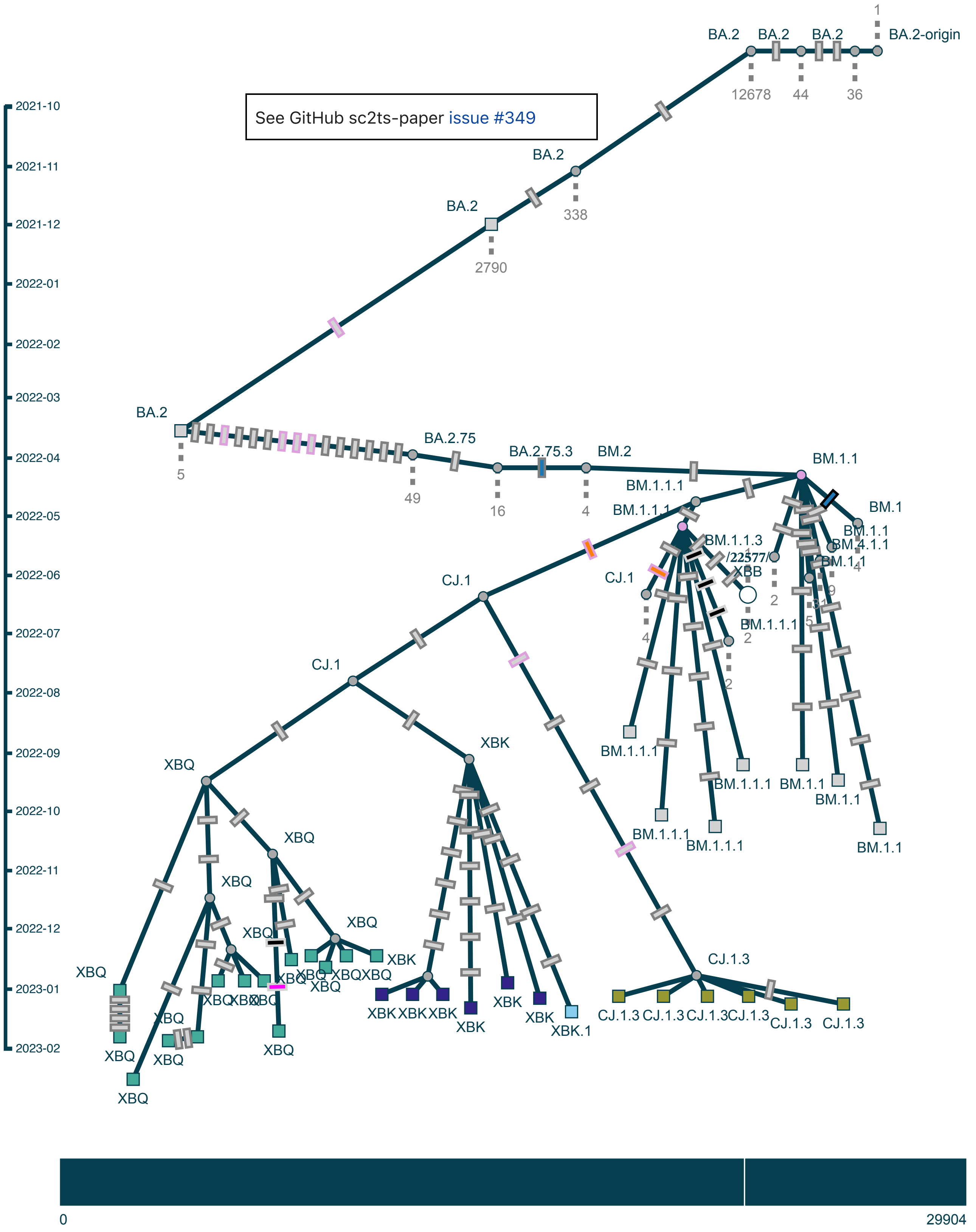

Subgraph of pango XBM: (10 samples, 10 shown)

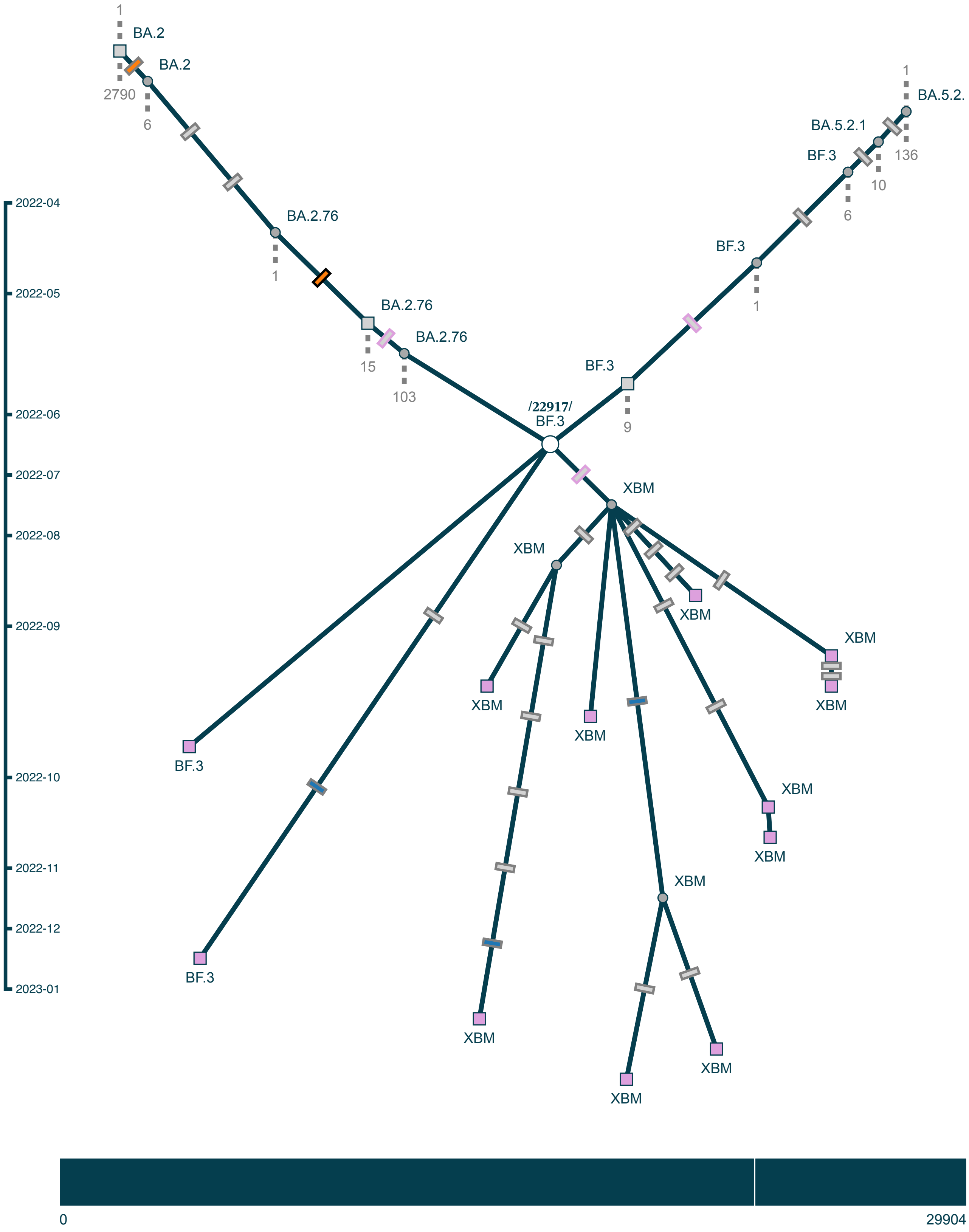

#### Subgraph of pango XBR: (1 sample, 1 shown)

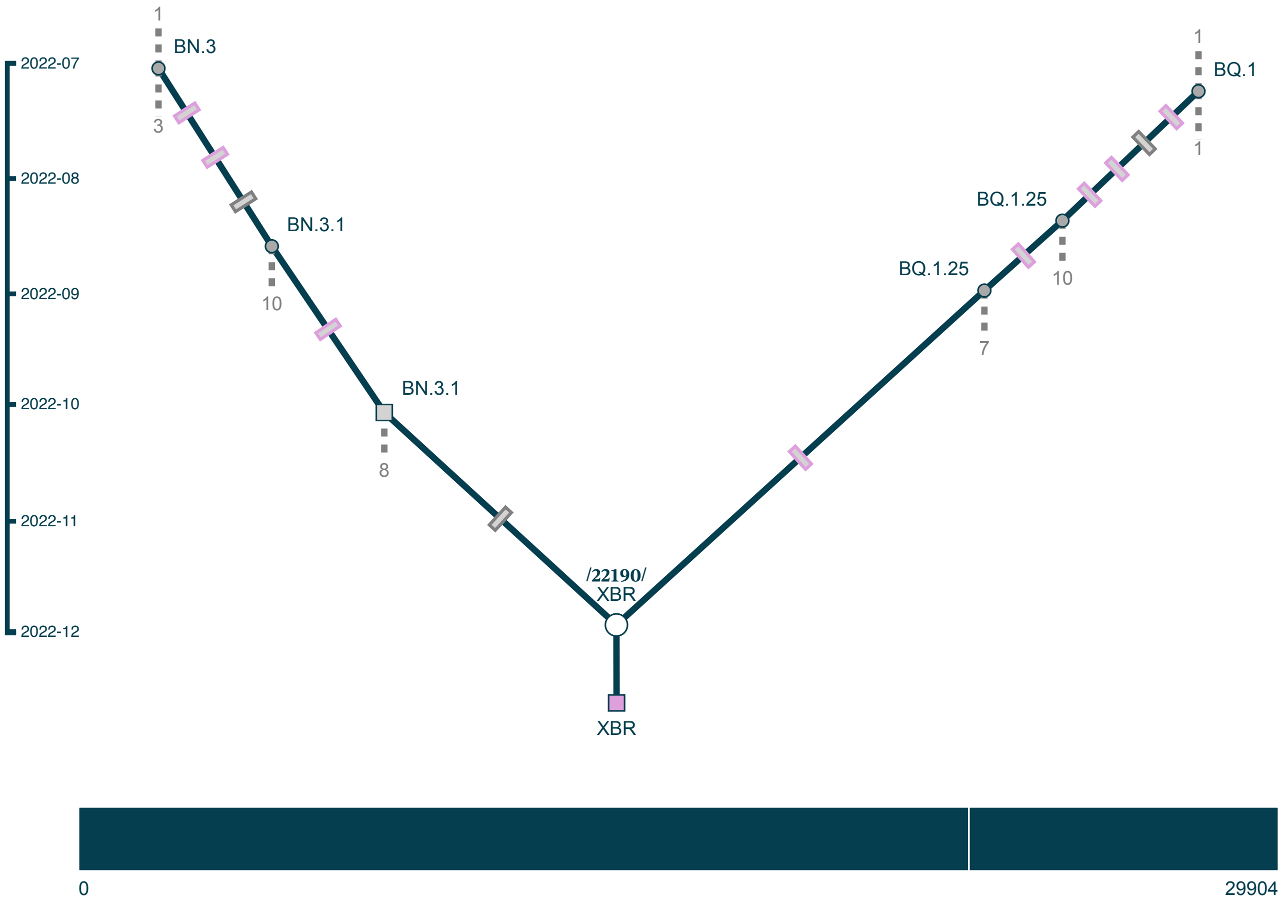
