## Supplementary material for "A Pandemic-Scale Ancestral Recombination Graph for SARS-CoV-2": Document S3: S3-copy_patterns.html

 

|  |  |  |  |  |  |  |  |  |  |  |  |  |  |  |
| --- | --- | --- | --- | --- | --- | --- | --- | --- | --- | --- | --- | --- | --- | --- |
| pos | 241 | 850 | 3037 | 6449 | 8782 | 13617 | 14408 | 17126 | 20268 | 23403 | 24197 | 26088 | 28144 |  |
| ref | C | C | C | C | C | G | C | T | A | A | G | C | T |  |
| B | C | T | C | T | T | T | C | T | A | A | G | T | C |  |
| B.1.157 | C | T | C | T | T | G | T | C | G | G | A | C | C | Copying pattern for recombinant child, node #1530 |
| B.1.157 | T | C | T | C | C | G | T | C | G | G | G | C | T |  |
| mut |  |  |  |  |  |  |  |  |  |  | G>A |  | T>C |  |

|  |  |  |  |  |  |  |  |  |  |  |  |  |  |  |  |  |  |  |
| --- | --- | --- | --- | --- | --- | --- | --- | --- | --- | --- | --- | --- | --- | --- | --- | --- | --- | --- |
| pos | 222 | 445 | 3602 | 6286 | 6941 | 15324 | 21255 | 21855 | 22227 | 25505 | 25906 | 25996 | 26801 | 28651 | 28869 | 28932 | 29645 |  |
| ref | C | T | C | C | C | C | G | C | C | A | G | G | C | C | C | C | G |  |
| B.1.177 | C | C | C | T | C | C | C | C | T | A | G | G | G | C | C | T | T |  |
| B.1.221 | T | C | C | T | C | C | G | T | C | G | C | T | C | T | T | C | G | #26465 |
| B.1.221 | C | T | T | C | T | T | G | T | C | G | C | T | C | T | T | C | G |  |
| mut | C>T |  |  |  |  |  | C>G |  |  |  |  |  |  |  |  |  |  |  |

|  |  |  |  |  |  |  |  |  |  |  |  |  |  |  |  |  |  |  |  |  |  |  |  |  |  |  |  |  |
| --- | --- | --- | --- | --- | --- | --- | --- | --- | --- | --- | --- | --- | --- | --- | --- | --- | --- | --- | --- | --- | --- | --- | --- | --- | --- | --- | --- | --- |
| pos | 4543 | 5629 | 9526 | 10265 | 11497 | 13993 | 15720 | 15766 | 15972 | 16889 | 17019 | 18877 | 19480 | 19718 | 22992 | 25563 | 25710 | 26700 | 26735 | 26876 | 28759 | 28881 | 28882 | 28883 | 28975 | 29348 | 29399 |  |
| ref | C | G | G | G | C | G | C | G | A | A | G | C | G | C | G | G | C | G | C | T | T | G | G | G | G | G | G |  |
| B.1.160 | T | T | T | G | T | T | C | T | A | G | T | T | G | C | A | T | T | G | T | C | T | G | G | G | C | G | A |  |
| B.1.160 | T | T | T | G | T | T | C | T | A | G | T | T | T | C | A | G | C | T | C | T | C | A | A | C | G | T | G | #27003 |
| B.1.1.70 | C | G | G | A | C | G | T | G | G | A | G | C | G | T | G | G | C | T | C | T | C | A | A | C | G | T | G |  |
| mut |  |  |  |  |  |  |  |  |  |  |  |  | G>T |  |  |  |  |  |  |  |  |  |  |  |  |  |  |  |

|  |  |  |  |  |  |  |  |  |  |  |  |  |  |  |  |  |  |  |  |  |  |  |  |  |  |
| --- | --- | --- | --- | --- | --- | --- | --- | --- | --- | --- | --- | --- | --- | --- | --- | --- | --- | --- | --- | --- | --- | --- | --- | --- | --- |
| pos | 833 | 2003 | 2473 | 5512 | 6541 | 9515 | 10943 | 11379 | 16260 | 17125 | 17550 | 19175 | 19839 | 21221 | 22320 | 25177 | 25523 | 26230 | 27434 | 27916 | 27964 | 28821 | 29466 | 29685 |  |
| ref | T | C | A | C | C | C | G | C | C | C | C | A | T | C | A | G | G | G | C | G | C | C | C | T |  |
| B.1.369 | T | T | A | C | C | C | A | C | T | C | T | A | T | C | A | T | G | C | C | T | C | A | T | T |  |
| B.1.426 | T | T | A | C | C | T | G | T | C | T | C | C | C | T | G | G | T | G | T | G | T | C | C | G | #28379 |
| B.1.426 | C | C | G | T | T | T | G | T | C | T | C | C | C | T | G | G | T | G | T | G | T | C | C | G |  |
| mut |  |  |  |  |  |  |  |  |  |  |  |  |  |  |  |  |  |  |  |  |  |  |  |  |  |

|  |  |  |  |  |  |  |  |  |  |  |  |  |
| --- | --- | --- | --- | --- | --- | --- | --- | --- | --- | --- | --- | --- |
| pos | 204 | 7926 | 9745 | 11522 | 20404 | 25049 | 25062 | 27944 | 28657 | 29366 | 29401 |  |
| ref | G | C | C | T | C | G | G | C | C | C | T |  |
| B.1.177.56 | T | T | T | G | C | G | G | T | C | C | T |  |
| B.1.177 | T | T | T | G | T | T | T | C | T | T | C | #34811 |
| B.1.177.11 | G | C | C | T | C | T | T | C | T | T | T |  |
| mut |  |  |  |  | C>T |  |  |  |  |  | T>C |  |

|  |  |  |  |  |  |  |  |  |  |  |  |  |  |  |  |  |  |  |  |  |  |  |
| --- | --- | --- | --- | --- | --- | --- | --- | --- | --- | --- | --- | --- | --- | --- | --- | --- | --- | --- | --- | --- | --- | --- |
| pos | 213 | 1150 | 1210 | 1344 | 4291 | 8683 | 9802 | 12067 | 13667 | 15406 | 15957 | 16260 | 19017 | 19862 | 19960 | 21637 | 27865 | 27866 | 27999 | 28169 | 29785 |  |
| ref | G | C | G | T | A | C | G | G | C | G | G | C | C | C | A | C | A | T | C | A | T |  |
| B.1.1.37 | A | C | G | C | T | T | G | G | C | T | G | T | C | T | G | T | A | T | C | G | T |  |
| B.1.1.307 | A | T | G | C | T | C | T | T | T | G | T | C | T | C | A | C | T | A | T | A | A | #37970 |
| B.1.1.307 | G | C | T | T | A | C | T | T | T | G | T | C | T | C | A | C | T | A | T | A | A |  |
| mut |  | C>T |  |  |  |  |  |  |  |  |  |  |  |  |  |  |  |  |  |  |  |  |

|  |  |  |  |  |  |  |  |  |  |  |  |  |  |  |  |  |  |  |  |  |  |  |  |  |  |  |  |  |  |
| --- | --- | --- | --- | --- | --- | --- | --- | --- | --- | --- | --- | --- | --- | --- | --- | --- | --- | --- | --- | --- | --- | --- | --- | --- | --- | --- | --- | --- | --- |
| pos | 66 | 445 | 1987 | 3602 | 6286 | 6941 | 10870 | 11781 | 15324 | 17334 | 21255 | 21855 | 21991 | 21992 | 21993 | 22227 | 25505 | 25614 | 25906 | 25996 | 26801 | 27870 | 27944 | 28651 | 28869 | 28932 | 29645 | 29769 |  |
| ref | C | T | A | C | C | C | G | A | C | G | G | C | T | T | A | C | A | C | G | G | C | G | C | C | C | C | G | C |  |
| B.1.177.4 | T | C | G | C | T | C | T | G | C | T | C | C | T | T | A | T | A | T | G | G | G | T | T | C | C | T | T | C |  |
| B.1.177.4 | T | C | G | C | T | C | T | G | C | T | C | C | - | - | - | T | A | T | G | G | G | T | C | T | T | C | G | T | #39337 |
| B.1.221 | C | T | A | T | C | T | G | A | T | G | G | T | T | T | A | C | G | C | C | T | C | G | C | T | T | C | G | C |  |
| mut |  |  |  |  |  |  |  |  |  |  |  |  | T>- | T>- | A>- |  |  |  |  |  |  |  |  |  |  |  |  | C>T |  |

|  |  |  |  |  |  |  |  |  |  |  |  |  |  |
| --- | --- | --- | --- | --- | --- | --- | --- | --- | --- | --- | --- | --- | --- |
| pos | 204 | 222 | 5000 | 6070 | 20661 | 21614 | 24334 | 26951 | 27769 | 27944 | 29366 | 29386 |  |
| ref | G | C | C | C | T | C | C | G | C | C | C | C |  |
| B.1.177 | G | T | T | T | C | C | C | G | C | C | T | T |  |
| B.1.177 | G | T | T | T | C | C | T | T | T | T | C | C | #42494 |
| B.1.177 | T | C | C | C | T | T | T | T | T | T | C | C |  |
| mut |  |  |  |  |  |  |  |  |  |  |  |  |  |

|  |  |  |  |  |  |  |  |  |  |  |  |  |  |  |  |  |  |  |  |  |  |  |  |  |  |  |  |  |  |  |  |
| --- | --- | --- | --- | --- | --- | --- | --- | --- | --- | --- | --- | --- | --- | --- | --- | --- | --- | --- | --- | --- | --- | --- | --- | --- | --- | --- | --- | --- | --- | --- | --- |
| pos | 3602 | 4442 | 4543 | 5629 | 6525 | 6941 | 9526 | 10377 | 11497 | 13993 | 15324 | 15766 | 16044 | 16889 | 17019 | 18877 | 21855 | 22992 | 25505 | 25563 | 25710 | 25731 | 25906 | 25996 | 26735 | 26876 | 28651 | 28869 | 28975 | 29399 |  |
| ref | C | G | C | G | C | C | G | C | C | G | C | G | A | A | G | C | C | G | A | G | C | C | G | G | C | T | C | C | G | G |  |
| B.1.221.1 | T | T | C | G | T | T | G | T | C | G | T | G | T | A | G | C | T | G | G | G | C | T | C | T | C | T | T | T | G | G |  |
| B.1 | T | T | C | G | T | T | G | T | C | G | T | G | T | A | G | C | T | G | A | T | T | C | G | G | T | C | C | C | C | A | #42873 |
| B.1.160 | C | G | T | T | C | C | T | C | T | T | C | T | A | G | T | T | C | A | A | T | T | C | G | G | T | C | C | C | C | A |  |
| mut |  |  |  |  |  |  |  |  |  |  |  |  |  |  |  |  |  |  |  |  |  |  |  |  |  |  |  |  |  |  |  |

|  |  |  |  |  |  |  |  |  |  |  |  |  |  |  |  |
| --- | --- | --- | --- | --- | --- | --- | --- | --- | --- | --- | --- | --- | --- | --- | --- |
| pos | 1590 | 1820 | 2232 | 14877 | 18176 | 19185 | 24034 | 25617 | 27874 | 27875 | 27881 | 27882 | 27883 | 28382 |  |
| ref | G | G | C | C | C | C | C | G | C | T | C | G | C | T |  |
| B.1.1.308 | T | G | C | C | C | C | C | T | C | T | C | G | C | T |  |
| B.1.1.308 | T | A | C | C | T | C | C | T | T | C | T | C | T | T | #44468 |
| B.1.1.29 | G | G | T | T | C | T | T | G | T | C | T | C | T | C |  |
| mut |  | G>A |  |  | C>T |  |  |  |  |  |  |  |  | C>T |  |

|  |  |  |  |  |  |  |  |  |  |  |  |  |  |  |  |  |  |
| --- | --- | --- | --- | --- | --- | --- | --- | --- | --- | --- | --- | --- | --- | --- | --- | --- | --- |
| pos | 204 | 445 | 4002 | 6286 | 11396 | 21255 | 21614 | 22227 | 22329 | 26801 | 27944 | 28881 | 28882 | 28883 | 28932 | 29645 |  |
| ref | G | T | C | C | C | G | C | C | C | C | C | G | G | G | C | G |  |
| B.1.1 | G | T | T | C | C | G | C | C | C | C | C | A | A | C | C | G |  |
| B.1.177 | G | T | T | C | T | C | T | T | T | G | T | G | G | G | T | T | #45340 |
| B.1.177 | T | C | C | T | T | C | T | T | T | G | T | G | G | G | T | T |  |
| mut |  |  |  |  |  |  |  |  |  |  |  |  |  |  |  |  |  |

|  |  |  |  |  |  |  |  |  |  |  |  |  |  |  |  |  |  |  |  |  |  |  |  |  |  |  |  |  |  |  |  |  |  |  |  |  |  |  |  |  |  |  |  |  |  |  |  |  |  |  |  |  |  |  |  |  |  |  |  |  |  |  |  |  |
| --- | --- | --- | --- | --- | --- | --- | --- | --- | --- | --- | --- | --- | --- | --- | --- | --- | --- | --- | --- | --- | --- | --- | --- | --- | --- | --- | --- | --- | --- | --- | --- | --- | --- | --- | --- | --- | --- | --- | --- | --- | --- | --- | --- | --- | --- | --- | --- | --- | --- | --- | --- | --- | --- | --- | --- | --- | --- | --- | --- | --- | --- | --- | --- | --- |
| pos | 204 | 445 | 913 | 3267 | 3291 | 4136 | 5388 | 5986 | 6286 | 6310 | 6954 | 7252 | 10833 | 11288 | 11289 | 11290 | 11291 | 11292 | 11293 | 11294 | 11295 | 11296 | 11396 | 14676 | 15017 | 15279 | 16176 | 20133 | 21255 | 21614 | 21765 | 21766 | 21767 | 21768 | 21769 | 21770 | 21991 | 21992 | 21993 | 22227 | 23063 | 23271 | 23604 | 23709 | 24506 | 24914 | 25006 | 26645 | 26801 | 27944 | 27972 | 28048 | 28111 | 28271 | 28280 | 28281 | 28282 | 28881 | 28882 | 28883 | 28932 | 28977 | 29645 |  |
| ref | G | T | C | C | A | G | C | C | C | C | T | G | C | T | C | T | G | G | T | T | T | T | C | C | C | C | T | C | G | C | T | A | C | A | T | G | T | T | A | C | A | C | C | C | T | G | C | C | C | C | C | G | A | A | G | A | T | G | G | G | C | C | G |  |
| B.1.177 | T | C | C | C | A | T | C | C | T | T | T | G | C | T | C | T | G | G | T | T | T | T | T | C | C | C | T | C | C | T | T | A | C | A | T | G | T | T | A | T | A | C | C | C | T | G | C | T | G | T | C | G | A | A | G | A | T | G | G | G | T | C | T |  |
| B.1.1 | T | C | C | C | G | T | C | C | T | T | T | G | C | T | C | T | G | G | T | T | T | T | T | C | T | C | T | T | G | C | - | - | - | - | - | - | - | - | - | C | T | A | A | T | G | C | C | C | C | C | T | T | G | - | C | T | A | A | A | C | C | T | G | #79763 |
| B.1.1.7 | G | T | T | T | A | G | A | T | C | C | C | A | T | - | - | - | - | - | - | - | - | - | C | T | C | T | C | T | G | C | - | - | - | - | - | - | - | - | - | C | T | A | A | T | G | C | T | C | C | C | T | T | G | - | C | T | A | A | A | C | C | T | G |  |
| mut |  |  |  |  | A>G |  |  |  |  |  |  |  |  |  |  |  |  |  |  |  |  |  |  |  | C>T |  |  |  |  |  |  |  |  |  |  |  |  |  |  |  |  |  |  |  |  |  | T>C |  |  |  |  |  |  |  |  |  |  |  |  |  |  |  |  |  |

|  |  |  |  |  |  |  |  |  |  |  |  |  |  |  |  |  |  |  |  |  |  |  |  |  |  |  |  |  |  |  |  |  |  |  |  |  |  |  |  |  |  |  |  |  |  |  |  |  |  |  |  |  |  |  |  |  |  |  |  |  |  |  |  |  |
| --- | --- | --- | --- | --- | --- | --- | --- | --- | --- | --- | --- | --- | --- | --- | --- | --- | --- | --- | --- | --- | --- | --- | --- | --- | --- | --- | --- | --- | --- | --- | --- | --- | --- | --- | --- | --- | --- | --- | --- | --- | --- | --- | --- | --- | --- | --- | --- | --- | --- | --- | --- | --- | --- | --- | --- | --- | --- | --- | --- | --- | --- | --- | --- | --- |
| pos | 913 | 1947 | 3267 | 5388 | 5986 | 6528 | 6954 | 7292 | 8410 | 9092 | 9165 | 10833 | 11288 | 11289 | 11290 | 11291 | 11292 | 11293 | 11294 | 11295 | 11296 | 11339 | 11572 | 12262 | 14676 | 15279 | 16176 | 16293 | 17615 | 18877 | 21765 | 21766 | 21767 | 21768 | 21769 | 21770 | 21991 | 21992 | 21993 | 22444 | 22484 | 23063 | 23271 | 23604 | 23709 | 23986 | 24506 | 24914 | 25563 | 26735 | 27972 | 28048 | 28111 | 28271 | 28280 | 28281 | 28282 | 28854 | 28881 | 28882 | 28883 | 28977 | 29645 |  |
| ref | C | T | C | C | C | A | T | T | T | G | C | C | T | C | T | G | G | T | T | T | T | C | C | T | C | C | T | C | A | C | T | A | C | A | T | G | T | T | A | C | G | A | C | C | C | T | T | G | G | C | C | G | A | A | G | A | T | C | G | G | G | C | G |  |
| B.1.36.28 | C | C | C | C | C | G | T | C | C | A | C | T | T | C | T | G | G | T | T | T | T | C | T | T | C | C | T | T | A | T | T | A | C | A | T | G | T | T | A | T | G | A | C | C | C | C | T | G | T | T | C | G | A | A | G | A | T | T | G | G | G | C | G |  |
| B.1.1.7 | C | C | C | C | C | G | C | T | T | G | T | C | - | - | - | - | - | - | - | - | - | T | C | C | T | T | C | C | G | C | - | - | - | - | - | - | - | - | - | C | T | T | A | A | T | T | G | C | G | C | T | T | G | - | C | T | A | C | A | A | C | T | T | #82697 |
| B.1.1.7 | T | T | T | A | T | A | C | T | T | G | T | C | - | - | - | - | - | - | - | - | - | T | C | C | T | T | C | C | G | C | - | - | - | - | - | - | - | - | - | C | T | T | A | A | T | T | G | C | G | C | T | T | G | - | C | T | A | C | A | A | C | T | T |  |
| mut |  |  |  |  |  |  |  |  |  |  |  |  |  |  |  |  |  |  |  |  |  |  |  |  |  |  |  |  |  |  |  |  |  |  |  |  |  |  |  |  |  |  |  |  |  |  |  |  |  |  |  |  |  |  |  |  |  |  |  |  |  |  |  |  |

|  |  |  |  |  |  |  |  |  |  |  |  |  |  |  |  |  |  |  |  |  |  |  |  |  |  |  |  |  |  |  |  |  |  |  |  |  |  |  |  |  |  |  |  |  |  |  |  |  |  |  |  |  |  |  |  |  |  |  |  |
| --- | --- | --- | --- | --- | --- | --- | --- | --- | --- | --- | --- | --- | --- | --- | --- | --- | --- | --- | --- | --- | --- | --- | --- | --- | --- | --- | --- | --- | --- | --- | --- | --- | --- | --- | --- | --- | --- | --- | --- | --- | --- | --- | --- | --- | --- | --- | --- | --- | --- | --- | --- | --- | --- | --- | --- | --- | --- | --- | --- |
| pos | 445 | 913 | 5388 | 5986 | 6286 | 6954 | 7348 | 10523 | 11288 | 11289 | 11290 | 11291 | 11292 | 11293 | 11294 | 11295 | 11296 | 14676 | 15279 | 16176 | 17215 | 21255 | 21306 | 21765 | 21766 | 21767 | 21768 | 21769 | 21770 | 21991 | 21992 | 21993 | 22227 | 23063 | 23271 | 23604 | 23709 | 24106 | 24506 | 24914 | 25970 | 26801 | 27972 | 28048 | 28111 | 28271 | 28280 | 28281 | 28282 | 28292 | 28310 | 28881 | 28882 | 28883 | 28932 | 28977 | 29645 | 29738 |  |
| ref | T | C | C | C | C | T | T | G | T | C | T | G | G | T | T | T | T | C | C | T | A | G | C | T | A | C | A | T | G | T | T | A | C | A | C | C | C | C | T | G | G | C | C | G | A | A | G | A | T | C | C | G | G | G | C | C | G | C |  |
| B.1.177.16 | C | C | C | C | T | T | T | A | T | C | T | G | G | T | T | T | T | C | C | T | G | C | T | T | A | C | A | T | G | T | T | A | T | A | C | C | C | T | T | G | G | G | C | G | A | A | G | A | T | A | A | G | G | G | T | C | T | T |  |
| B.1.1 | C | C | C | C | T | T | C | A | T | C | T | G | G | T | T | T | T | C | C | T | G | C | T | T | A | C | A | T | G | T | T | A | T | A | C | C | C | T | T | G | T | G | T | T | G | - | C | T | A | C | C | A | A | C | C | T | G | C | #99000 |
| B.1.1.7 | T | T | A | T | C | C | T | G | - | - | - | - | - | - | - | - | - | T | T | C | A | G | C | - | - | - | - | - | - | - | - | - | C | T | A | A | T | C | G | C | G | C | T | T | G | - | C | T | A | C | C | A | A | C | C | T | G | C |  |
| mut |  |  |  |  |  |  | T>C |  |  |  |  |  |  |  |  |  |  |  |  |  |  |  |  |  |  |  |  |  |  |  |  |  |  |  |  |  |  |  |  |  | G>T |  |  |  |  |  |  |  |  |  |  |  |  |  |  |  |  |  |  |

|  |  |  |  |  |  |  |  |  |  |  |  |  |  |  |  |  |  |  |  |  |  |  |  |  |  |  |  |  |  |  |  |  |  |  |  |  |  |  |  |  |  |  |  |  |  |  |  |  |  |  |  |  |  |  |  |  |  |  |  |  |  |  |  |  |
| --- | --- | --- | --- | --- | --- | --- | --- | --- | --- | --- | --- | --- | --- | --- | --- | --- | --- | --- | --- | --- | --- | --- | --- | --- | --- | --- | --- | --- | --- | --- | --- | --- | --- | --- | --- | --- | --- | --- | --- | --- | --- | --- | --- | --- | --- | --- | --- | --- | --- | --- | --- | --- | --- | --- | --- | --- | --- | --- | --- | --- | --- | --- | --- | --- |
| pos | 204 | 445 | 913 | 3267 | 5388 | 5986 | 6286 | 6954 | 9165 | 10974 | 11288 | 11289 | 11290 | 11291 | 11292 | 11293 | 11294 | 11295 | 11296 | 11396 | 12262 | 14676 | 15279 | 16176 | 17615 | 21255 | 21614 | 21765 | 21766 | 21767 | 21768 | 21769 | 21770 | 21991 | 21992 | 21993 | 22205 | 22227 | 23063 | 23271 | 23604 | 23709 | 24506 | 24914 | 26801 | 27301 | 27881 | 27944 | 27972 | 28048 | 28111 | 28271 | 28280 | 28281 | 28282 | 28697 | 28717 | 28881 | 28882 | 28883 | 28932 | 28956 | 28977 |  |
| ref | G | T | C | C | C | C | C | T | C | G | T | C | T | G | G | T | T | T | T | C | T | C | C | T | A | G | C | T | A | C | A | T | G | T | T | A | G | C | A | C | C | C | T | G | C | A | C | C | C | G | A | A | G | A | T | C | C | G | G | G | C | A | C |  |
| B.1.177 | T | C | C | C | C | C | T | T | C | A | T | C | T | G | G | T | T | T | T | T | T | C | C | T | A | C | T | T | A | C | A | T | G | T | T | A | C | T | A | C | C | C | T | G | G | G | A | T | C | G | A | A | G | A | T | T | T | G | G | G | T | C | C |  |
| B.1.177 | T | C | C | C | C | C | T | T | C | A | T | C | T | G | G | T | T | T | T | T | T | C | C | T | A | C | T | T | A | C | A | T | G | T | T | A | C | T | A | C | C | C | T | G | G | G | A | T | C | G | A | A | G | A | T | T | T | A | A | C | C | A | T | #103390 |
| B.1.1.7 | G | T | T | T | A | T | C | C | T | G | - | - | - | - | - | - | - | - | - | C | C | T | T | C | G | G | C | - | - | - | - | - | - | - | - | - | G | C | T | A | A | T | G | C | C | A | C | C | T | T | G | - | C | T | A | C | C | A | A | C | C | A | T |  |
| mut |  |  |  |  |  |  |  |  |  |  |  |  |  |  |  |  |  |  |  |  |  |  |  |  |  |  |  |  |  |  |  |  |  |  |  |  |  |  |  |  |  |  |  |  |  |  |  |  |  |  |  |  |  |  |  |  |  |  |  |  |  |  |  |  |

|  |  |  |  |  |  |  |  |  |  |  |  |  |  |  |  |  |  |  |  |  |  |  |  |  |  |  |  |  |  |  |  |  |  |  |  |  |  |  |  |  |  |  |  |  |  |  |  |  |  |  |  |  |  |  |  |  |  |  |  |  |  |  |  |  |  |  |  |  |  |  |
| --- | --- | --- | --- | --- | --- | --- | --- | --- | --- | --- | --- | --- | --- | --- | --- | --- | --- | --- | --- | --- | --- | --- | --- | --- | --- | --- | --- | --- | --- | --- | --- | --- | --- | --- | --- | --- | --- | --- | --- | --- | --- | --- | --- | --- | --- | --- | --- | --- | --- | --- | --- | --- | --- | --- | --- | --- | --- | --- | --- | --- | --- | --- | --- | --- | --- | --- | --- | --- | --- | --- |
| pos | 445 | 913 | 1643 | 3267 | 5388 | 5986 | 6286 | 6500 | 6954 | 7037 | 7534 | 7728 | 10870 | 11288 | 11289 | 11290 | 11291 | 11292 | 11293 | 11294 | 11295 | 11296 | 13176 | 14676 | 15279 | 16176 | 16305 | 19274 | 20223 | 21255 | 21765 | 21766 | 21767 | 21768 | 21769 | 21770 | 21800 | 21991 | 21992 | 21993 | 22053 | 22227 | 22346 | 23063 | 23271 | 23604 | 23709 | 24506 | 24914 | 26013 | 26801 | 27972 | 28048 | 28095 | 28111 | 28251 | 28253 | 28254 | 28271 | 28280 | 28281 | 28282 | 28881 | 28882 | 28883 | 28932 | 28977 | 29218 | 29645 |  |
| ref | T | C | A | C | C | C | C | C | T | G | T | C | G | T | C | T | G | G | T | T | T | T | C | C | C | T | A | C | A | G | T | A | C | A | T | G | G | T | T | A | A | C | G | A | C | C | C | T | G | C | C | C | G | A | A | T | C | A | A | G | A | T | G | G | G | C | C | C | G |  |
| B.1.1.7 | T | T | T | T | A | T | C | C | C | G | T | T | G | - | - | - | - | - | - | - | - | - | T | T | T | C | A | C | A | G | - | - | - | - | - | - | G | - | - | - | A | C | G | T | A | A | T | G | C | C | C | T | T | T | G | C | T | C | - | C | T | A | A | A | C | C | T | C | G |  |
| B.1.177.9 | T | T | T | T | A | T | C | C | C | G | T | T | T | T | C | T | G | G | T | T | T | T | C | C | C | T | G | T | G | C | T | A | C | A | T | G | T | T | T | A | C | T | T | A | C | C | C | T | G | T | G | C | G | A | A | T | C | A | A | G | A | T | G | G | G | T | C | T | T | #104754 |
| B.1.177.9 | C | C | A | C | C | C | T | T | T | T | C | C | T | T | C | T | G | G | T | T | T | T | C | C | C | T | A | T | A | C | T | A | C | A | T | G | T | T | T | A | C | T | T | A | C | C | C | T | G | T | G | C | G | A | A | T | C | A | A | G | A | T | G | G | G | T | C | C | T |  |
| mut |  |  |  |  |  |  |  |  |  |  |  |  |  |  |  |  |  |  |  |  |  |  |  |  |  |  | A>G |  | A>G |  |  |  |  |  |  |  |  |  |  |  |  |  |  |  |  |  |  |  |  |  |  |  |  |  |  |  |  |  |  |  |  |  |  |  |  |  |  | C>T |  |  |

|  |  |  |  |  |  |  |  |  |  |  |  |  |  |  |  |  |  |  |  |  |  |  |  |  |  |  |  |  |  |  |  |  |  |  |  |  |  |  |  |  |  |  |  |  |  |  |  |  |  |  |  |  |  |  |  |  |  |  |  |  |  |  |  |
| --- | --- | --- | --- | --- | --- | --- | --- | --- | --- | --- | --- | --- | --- | --- | --- | --- | --- | --- | --- | --- | --- | --- | --- | --- | --- | --- | --- | --- | --- | --- | --- | --- | --- | --- | --- | --- | --- | --- | --- | --- | --- | --- | --- | --- | --- | --- | --- | --- | --- | --- | --- | --- | --- | --- | --- | --- | --- | --- | --- | --- | --- | --- | --- |
| pos | 913 | 920 | 3267 | 3602 | 4442 | 5388 | 5986 | 6525 | 6941 | 6954 | 9857 | 10193 | 10195 | 11288 | 11289 | 11290 | 11291 | 11292 | 11293 | 11294 | 11295 | 11296 | 14676 | 15279 | 15324 | 16044 | 16176 | 21765 | 21766 | 21767 | 21768 | 21769 | 21770 | 21855 | 21991 | 21992 | 21993 | 23063 | 23271 | 23604 | 23709 | 24106 | 24506 | 24914 | 25505 | 25731 | 25906 | 25996 | 27972 | 28048 | 28111 | 28271 | 28280 | 28281 | 28282 | 28651 | 28869 | 28881 | 28882 | 28883 | 28899 | 28977 |  |
| ref | C | C | C | C | G | C | C | C | C | T | C | G | A | T | C | T | G | G | T | T | T | T | C | C | C | A | T | T | A | C | A | T | G | C | T | T | A | A | C | C | C | C | T | G | A | C | G | G | C | G | A | A | G | A | T | C | C | G | G | G | G | C |  |
| B.1.1.7 | T | T | T | C | G | A | T | C | C | C | C | A | T | - | - | - | - | - | - | - | - | - | T | T | C | A | C | - | - | - | - | - | - | C | - | - | - | T | A | A | T | C | G | C | A | C | G | G | T | T | G | - | C | T | A | C | C | A | A | C | G | T |  |
| B.1.1.7 | T | T | T | C | G | A | T | C | C | C | C | A | T | - | - | - | - | - | - | - | - | - | T | T | C | A | C | - | - | - | - | - | - | C | - | - | - | T | A | A | T | C | G | C | A | C | G | G | C | G | A | A | G | A | T | T | T | G | G | G | A | C | #105931 |
| B.1.221.1 | C | C | C | T | T | C | C | T | T | T | T | G | A | T | C | T | G | G | T | T | T | T | C | C | T | T | T | T | A | C | A | T | G | T | T | T | A | A | C | C | C | T | T | G | G | T | C | T | C | G | A | A | G | A | T | T | T | G | G | G | A | C |  |
| mut |  |  |  |  |  |  |  |  |  |  |  |  |  |  |  |  |  |  |  |  |  |  |  |  |  |  |  |  |  |  |  |  |  |  |  |  |  |  |  |  |  |  |  |  |  |  |  |  |  |  |  |  |  |  |  |  |  |  |  |  |  |  |  |

|  |  |  |  |  |  |  |  |  |  |  |  |  |  |  |  |  |  |  |  |  |  |  |  |  |  |  |  |  |  |  |  |  |  |  |  |  |  |  |  |  |  |  |  |  |  |  |  |  |  |  |  |  |  |  |  |  |  |  |  |  |  |  |  |  |  |  |
| --- | --- | --- | --- | --- | --- | --- | --- | --- | --- | --- | --- | --- | --- | --- | --- | --- | --- | --- | --- | --- | --- | --- | --- | --- | --- | --- | --- | --- | --- | --- | --- | --- | --- | --- | --- | --- | --- | --- | --- | --- | --- | --- | --- | --- | --- | --- | --- | --- | --- | --- | --- | --- | --- | --- | --- | --- | --- | --- | --- | --- | --- | --- | --- | --- | --- | --- |
| pos | 445 | 913 | 2019 | 2110 | 3267 | 4999 | 5388 | 5986 | 6286 | 6954 | 9430 | 10323 | 11288 | 11289 | 11290 | 11291 | 11292 | 11293 | 11294 | 11295 | 11296 | 13945 | 14120 | 14676 | 15279 | 16176 | 20410 | 21255 | 21765 | 21766 | 21767 | 21768 | 21769 | 21770 | 21991 | 21992 | 21993 | 22227 | 23063 | 23208 | 23271 | 23311 | 23604 | 23709 | 24506 | 24914 | 25440 | 25614 | 25855 | 26801 | 27752 | 27972 | 28048 | 28095 | 28111 | 28271 | 28280 | 28281 | 28282 | 28881 | 28882 | 28883 | 28932 | 28977 | 29645 |  |
| ref | T | C | T | C | C | C | C | C | C | T | C | A | T | C | T | G | G | T | T | T | T | C | C | C | C | T | G | G | T | A | C | A | T | G | T | T | A | C | A | C | C | G | C | C | T | G | G | C | G | C | C | C | G | A | A | A | G | A | T | G | G | G | C | C | G |  |
| B.1.177.18 | C | C | C | C | C | T | C | C | T | T | C | G | T | C | T | G | G | T | T | T | T | T | C | C | C | T | A | C | T | A | C | A | T | G | T | T | A | T | A | C | C | C | C | C | T | G | T | T | G | G | T | C | G | A | A | A | G | A | T | G | G | G | T | C | T |  |
| XA | C | C | C | C | C | T | C | C | T | T | T | G | T | C | T | G | G | T | T | T | T | T | C | C | C | T | A | C | - | - | - | - | - | - | - | - | - | C | T | T | A | G | A | T | G | C | G | C | T | C | C | T | T | T | G | - | C | T | A | A | A | C | C | T | G | #122444 |
| B.1.1.7 | T | T | T | T | T | C | A | T | C | C | C | A | - | - | - | - | - | - | - | - | - | C | T | T | T | C | G | G | - | - | - | - | - | - | - | - | - | C | T | T | A | G | A | T | G | C | G | C | T | C | C | T | T | T | G | - | C | T | A | A | A | C | C | T | G |  |
| mut |  |  |  |  |  |  |  |  |  |  | C>T |  |  |  |  |  |  |  |  |  |  |  |  |  |  |  |  |  |  |  |  |  |  |  |  |  |  |  |  |  |  |  |  |  |  |  |  |  |  |  |  |  |  |  |  |  |  |  |  |  |  |  |  |  |  |  |

|  |  |  |  |  |  |  |  |  |  |  |  |  |  |  |  |  |  |  |  |  |  |  |  |  |  |  |  |  |  |  |  |  |  |  |  |  |  |  |  |  |  |  |  |  |  |  |  |  |  |  |  |  |  |  |  |  |  |  |  |  |  |  |  |  |  |
| --- | --- | --- | --- | --- | --- | --- | --- | --- | --- | --- | --- | --- | --- | --- | --- | --- | --- | --- | --- | --- | --- | --- | --- | --- | --- | --- | --- | --- | --- | --- | --- | --- | --- | --- | --- | --- | --- | --- | --- | --- | --- | --- | --- | --- | --- | --- | --- | --- | --- | --- | --- | --- | --- | --- | --- | --- | --- | --- | --- | --- | --- | --- | --- | --- | --- |
| pos | 186 | 913 | 1947 | 2319 | 2453 | 3267 | 5388 | 5986 | 6258 | 6573 | 6954 | 7528 | 9142 | 9936 | 11288 | 11289 | 11290 | 11291 | 11292 | 11293 | 11294 | 11295 | 11296 | 12454 | 14676 | 15279 | 15981 | 16176 | 18877 | 21123 | 21765 | 21766 | 21767 | 21768 | 21769 | 21770 | 21991 | 21992 | 21993 | 22444 | 23063 | 23271 | 23604 | 23709 | 24305 | 24506 | 24914 | 25563 | 25904 | 26735 | 27972 | 28048 | 28111 | 28271 | 28280 | 28281 | 28282 | 28854 | 28881 | 28882 | 28883 | 28977 | 29555 | 29742 |  |
| ref | C | C | T | T | C | C | C | C | C | C | T | C | C | C | T | C | T | G | G | T | T | T | T | T | C | C | C | T | C | G | T | A | C | A | T | G | T | T | A | C | A | C | C | C | G | T | G | G | C | C | C | G | A | A | G | A | T | C | G | G | G | C | C | G |  |
| B.1.36.39 | C | C | C | C | C | C | C | C | C | T | T | T | T | A | T | C | T | G | G | T | T | T | T | T | C | C | T | T | T | G | T | A | C | A | T | G | T | T | A | T | A | C | C | C | A | T | G | T | T | T | C | G | A | A | G | A | T | T | G | G | G | C | T | G |  |
| B.1.1 | C | C | C | C | C | C | C | C | T | T | T | T | T | A | T | C | T | G | G | T | T | T | T | G | C | C | T | T | T | T | T | A | C | A | T | G | T | T | A | T | T | A | A | T | G | G | C | G | C | C | T | T | G | - | C | T | A | C | A | A | C | T | C | A | #125722 |
| B.1.1.7 | T | T | T | T | T | T | A | T | C | C | C | C | C | C | - | - | - | - | - | - | - | - | - | T | T | T | C | C | C | G | - | - | - | - | - | - | - | - | - | C | T | A | A | T | G | G | C | G | C | C | T | T | G | - | C | T | A | C | A | A | C | T | C | A |  |
| mut |  |  |  |  |  |  |  |  | C>T |  |  |  |  |  |  |  |  |  |  |  |  |  |  | T>G |  |  |  |  |  | G>T |  |  |  |  |  |  |  |  |  |  |  |  |  |  |  |  |  |  |  |  |  |  |  |  |  |  |  |  |  |  |  |  |  |  |  |

|  |  |  |  |  |  |  |  |  |  |  |  |  |  |  |  |  |  |  |  |  |  |  |  |  |  |  |  |  |  |  |  |  |  |  |
| --- | --- | --- | --- | --- | --- | --- | --- | --- | --- | --- | --- | --- | --- | --- | --- | --- | --- | --- | --- | --- | --- | --- | --- | --- | --- | --- | --- | --- | --- | --- | --- | --- | --- | --- |
| pos | 204 | 445 | 583 | 832 | 2272 | 3004 | 4002 | 4890 | 6286 | 6558 | 8353 | 9289 | 10097 | 13536 | 16776 | 21255 | 21614 | 21746 | 21850 | 22227 | 22326 | 23731 | 24334 | 24781 | 26801 | 27513 | 27944 | 28881 | 28882 | 28883 | 28932 | 29645 | 29734 |  |
| ref | G | T | C | C | G | G | C | C | C | C | G | T | G | C | T | G | C | G | G | C | C | C | C | G | C | C | C | G | G | G | C | G | G |  |
| C.30 | G | T | T | T | G | T | T | T | C | C | A | C | A | T | T | G | C | G | G | C | T | T | C | T | C | T | C | A | A | C | C | G | C |  |
| B.1.177 | G | T | T | T | G | T | T | T | T | T | G | T | G | C | C | C | T | C | A | T | C | C | T | G | G | C | T | G | G | G | T | T | G | #130133 |
| B.1.177 | T | C | C | C | T | G | C | C | T | C | G | T | G | C | C | C | T | G | G | T | C | C | T | G | G | C | T | G | G | G | T | T | G |  |
| mut |  |  |  |  |  |  |  |  |  | C>T |  |  |  |  |  |  |  | G>C | G>A |  |  |  |  |  |  |  |  |  |  |  |  |  |  |  |

|  |  |  |  |  |  |  |  |  |  |  |  |  |  |  |  |  |  |
| --- | --- | --- | --- | --- | --- | --- | --- | --- | --- | --- | --- | --- | --- | --- | --- | --- | --- |
| pos | 752 | 2485 | 3811 | 7168 | 10369 | 15096 | 16575 | 16609 | 18998 | 21646 | 26485 | 26488 | 26491 | 26492 | 26497 | 29781 |  |
| ref | A | C | C | T | C | T | C | A | C | C | T | T | T | A | T | G |  |
| B.1.1.7 | G | T | T | T | C | C | C | A | C | C | T | T | T | A | T | G |  |
| B.1.1.7 | G | T | T | T | C | C | C | A | C | C | G | G | C | T | C | G | #143250 |
| B.1.1.7 | A | C | C | C | T | T | T | G | T | T | T | G | C | T | C | T |  |
| mut |  |  |  |  |  |  |  |  |  |  | T>G |  |  |  |  | T>G |  |

|  |  |  |  |  |  |  |  |  |  |  |  |  |  |  |  |  |  |  |  |  |  |  |  |  |  |  |  |  |  |  |  |
| --- | --- | --- | --- | --- | --- | --- | --- | --- | --- | --- | --- | --- | --- | --- | --- | --- | --- | --- | --- | --- | --- | --- | --- | --- | --- | --- | --- | --- | --- | --- | --- |
| pos | 1171 | 1346 | 5747 | 6479 | 9535 | 10319 | 10789 | 13335 | 17259 | 18424 | 18563 | 21304 | 21431 | 21765 | 21766 | 21767 | 21768 | 21769 | 21770 | 24007 | 25844 | 25907 | 26664 | 27879 | 27964 | 28472 | 28690 | 28869 | 28887 | 29705 |  |
| ref | T | C | A | G | T | C | C | C | G | A | A | C | C | T | A | C | A | T | G | G | C | G | A | C | C | C | G | C | C | G |  |
| B.1.375 | C | A | G | G | T | C | T | T | T | A | G | C | T | - | - | - | - | - | - | A | T | G | G | C | C | C | T | C | T | T |  |
| B.1.375 | C | A | G | G | C | C | T | T | T | A | G | C | T | - | - | - | - | - | - | A | T | G | A | T | T | T | G | T | C | G | #148227 |
| B.1.2 | T | C | A | A | T | T | C | C | G | G | A | T | C | T | A | C | A | T | G | G | C | T | A | T | T | T | G | T | C | G |  |
| mut |  |  |  |  | T>C |  |  |  |  |  |  |  |  |  |  |  |  |  |  |  |  |  |  |  |  |  |  |  |  |  |  |

|  |  |  |  |  |  |  |  |  |  |  |  |  |  |  |
| --- | --- | --- | --- | --- | --- | --- | --- | --- | --- | --- | --- | --- | --- | --- |
| pos | 337 | 3177 | 3327 | 8251 | 17799 | 18647 | 21306 | 21855 | 21975 | 26714 | 26721 | 26722 | 26723 |  |
| ref | C | C | A | C | A | C | C | C | A | T | C | T | T |  |
| B.1.1.7 | T | C | G | T | A | C | T | T | G | T | C | T | T |  |
| B.1.1.7 | C | C | G | T | A | T | T | T | G | G | T | A | C | #153040 |
| B.1.1.7 | C | T | A | C | G | C | C | C | A | G | T | A | C |  |
| mut | T>C |  |  |  |  | C>T |  |  |  |  |  |  |  |  |

|  |  |  |  |  |  |  |  |  |  |  |  |  |  |  |  |  |  |  |  |  |  |  |  |  |  |  |  |  |  |  |  |
| --- | --- | --- | --- | --- | --- | --- | --- | --- | --- | --- | --- | --- | --- | --- | --- | --- | --- | --- | --- | --- | --- | --- | --- | --- | --- | --- | --- | --- | --- | --- | --- |
| pos | 203 | 1829 | 2607 | 3484 | 3627 | 5541 | 5730 | 6394 | 7399 | 8083 | 10319 | 10874 | 11651 | 13138 | 14634 | 14805 | 17436 | 18131 | 18424 | 18555 | 21304 | 25677 | 25907 | 27964 | 28376 | 28472 | 28849 | 28869 | 28892 | 29518 |  |
| ref | C | A | C | A | A | A | C | T | A | G | C | A | C | C | A | C | C | C | A | C | C | G | G | C | G | C | C | C | C | C |  |
| B.1.349 | T | G | T | G | A | A | C | C | G | G | C | A | C | C | G | C | C | C | A | T | C | G | G | C | G | C | C | C | T | T |  |
| B.1.2 | T | G | T | G | C | A | C | C | G | G | T | G | T | T | A | T | T | T | G | C | T | T | T | T | T | T | T | T | C | C | #165974 |
| B.1.2 | C | A | C | A | A | C | T | T | A | A | T | G | T | T | A | T | T | T | G | C | T | T | T | T | T | T | T | T | C | C |  |
| mut |  |  |  |  | A>C |  |  |  |  |  |  |  |  |  |  |  |  |  |  |  |  |  |  |  |  |  |  |  |  |  |  |

|  |  |  |  |  |  |  |  |  |  |  |  |  |  |  |
| --- | --- | --- | --- | --- | --- | --- | --- | --- | --- | --- | --- | --- | --- | --- |
| pos | 2110 | 7984 | 14120 | 18687 | 19390 | 21462 | 22264 | 27880 | 27881 | 27882 | 27883 | 28087 | 28737 |  |
| ref | C | T | C | C | C | T | C | A | C | G | C | C | C |  |
| B.1.1.7 | T | C | T | C | T | T | C | A | C | G | C | C | C |  |
| B.1.1.7 | T | C | T | C | T | T | C | T | T | C | T | C | T | #179752 |
| B.1.1.7 | C | T | C | T | C | C | T | T | T | C | T | T | C |  |
| mut |  |  |  |  |  |  |  |  |  |  |  | T>C | C>T |  |

|  |  |  |  |  |  |  |  |  |  |  |  |
| --- | --- | --- | --- | --- | --- | --- | --- | --- | --- | --- | --- |
| pos | 695 | 2306 | 4582 | 23506 | 26416 | 28233 | 28238 | 28239 | 28240 | 28242 |  |
| ref | G | C | C | C | G | G | T | G | T | G |  |
| B.1.1.7 | G | T | C | T | T | G | T | G | T | G |  |
| B.1.1.7 | G | T | C | T | T | T | A | T | C | C | #191383 |
| B.1.1.7 | T | C | T | C | G | T | A | T | C | C |  |
| mut |  |  |  |  |  |  |  |  |  |  |  |

|  |  |  |  |  |  |  |  |  |  |  |  |  |  |  |  |  |  |  |  |  |  |  |  |  |  |  |  |  |  |  |  |  |  |  |  |  |  |  |  |  |  |  |  |  |  |
| --- | --- | --- | --- | --- | --- | --- | --- | --- | --- | --- | --- | --- | --- | --- | --- | --- | --- | --- | --- | --- | --- | --- | --- | --- | --- | --- | --- | --- | --- | --- | --- | --- | --- | --- | --- | --- | --- | --- | --- | --- | --- | --- | --- | --- | --- |
| pos | 626 | 3411 | 5512 | 6026 | 6101 | 6394 | 7201 | 7869 | 8683 | 8809 | 9867 | 10319 | 11288 | 11289 | 11290 | 11291 | 11292 | 11293 | 11294 | 11295 | 11296 | 13742 | 16017 | 16500 | 17336 | 20188 | 20262 | 21846 | 22320 | 22992 | 24432 | 25517 | 25968 | 27630 | 27739 | 27925 | 27929 | 27964 | 28271 | 28311 | 28328 | 28879 | 28887 | 29719 |  |
| ref | G | C | C | C | G | T | A | C | C | C | T | C | T | C | T | G | G | T | T | T | T | T | C | A | C | C | A | C | A | G | A | C | A | C | C | C | T | C | A | C | G | T | C | C |  |
| B.1.526 | G | C | C | C | A | T | G | C | C | T | C | C | - | - | - | - | - | - | - | - | - | G | C | C | C | C | G | T | G | A | G | T | G | T | T | T | T | C | - | T | G | G | C | C |  |
| B.1.526 | G | C | C | C | A | T | G | C | C | T | C | C | - | - | - | - | - | - | - | - | - | G | C | C | T | C | G | T | G | A | G | T | G | T | T | T | T | C | - | C | A | T | T | T | #200331 |
| B.1.595 | A | T | T | T | G | C | A | T | T | C | T | T | T | C | T | G | G | T | T | T | T | T | T | A | C | T | A | C | A | G | A | C | A | C | C | C | A | T | A | C | G | T | T | T |  |
| mut |  |  |  |  |  |  |  |  |  |  |  |  |  |  |  |  |  |  |  |  |  |  |  |  | C>T |  |  |  |  |  |  |  |  |  |  |  |  |  |  |  | G>A |  |  |  |  |

|  |  |  |  |  |  |  |  |  |  |  |  |  |  |  |  |
| --- | --- | --- | --- | --- | --- | --- | --- | --- | --- | --- | --- | --- | --- | --- | --- |
| pos | 2110 | 7042 | 12015 | 14120 | 18687 | 21462 | 22264 | 24049 | 27880 | 27881 | 27882 | 27883 | 28087 | 28847 |  |
| ref | C | G | T | C | C | T | C | A | A | C | G | C | C | A |  |
| B.1.1.7 | T | T | G | T | C | T | C | T | A | C | G | C | C | A |  |
| B.1.1.7 | T | T | G | T | C | T | C | T | T | T | C | T | C | G | #213224 |
| B.1.1.7 | C | G | T | C | T | C | T | A | T | T | C | T | T | A |  |
| mut |  |  |  |  |  |  |  |  |  |  |  |  | T>C | A>G |  |

|  |  |  |  |  |  |  |  |  |  |  |  |  |  |  |  |  |  |
| --- | --- | --- | --- | --- | --- | --- | --- | --- | --- | --- | --- | --- | --- | --- | --- | --- | --- |
| pos | 695 | 1387 | 4582 | 7765 | 23426 | 26834 | 26894 | 26895 | 27469 | 28233 | 28238 | 28239 | 28240 | 28314 | 28877 | 28878 |  |
| ref | G | C | C | C | G | G | C | C | G | G | T | G | T | G | A | G |  |
| B.1.1.7 | G | T | C | T | T | T | A | A | A | G | T | G | T | T | T | C |  |
| B.1.1.7 | G | T | C | T | T | T | A | A | A | T | A | T | C | G | T | C | #214402 |
| B.1.1.7 | T | C | T | C | G | G | C | C | G | T | A | T | C | G | A | G |  |
| mut |  |  |  |  |  |  |  |  |  |  |  |  |  |  | A>T | G>C |  |

|  |  |  |  |  |  |  |  |  |  |  |  |  |
| --- | --- | --- | --- | --- | --- | --- | --- | --- | --- | --- | --- | --- |
| pos | 3446 | 20961 | 23426 | 25066 | 25690 | 28243 | 28245 | 28246 | 28248 | 28249 | 28251 |  |
| ref | G | T | G | C | G | T | T | T | G | A | T |  |
| B.1.1.7 | A | C | T | T | T | T | G | T | G | A | C |  |
| B.1.1.7 | A | C | T | T | T | G | A | G | T | C | G | #229637 |
| B.1.1.7 | G | T | G | C | G | T | A | G | T | C | G |  |
| mut |  |  |  |  |  | T>G |  |  |  |  |  |  |

|  |  |  |  |  |  |  |  |  |  |  |  |  |  |  |  |  |  |  |  |  |  |  |  |  |  |  |  |  |  |  |  |  |  |  |  |  |  |  |  |  |  |  |  |  |  |  |  |  |  |  |  |  |  |  |
| --- | --- | --- | --- | --- | --- | --- | --- | --- | --- | --- | --- | --- | --- | --- | --- | --- | --- | --- | --- | --- | --- | --- | --- | --- | --- | --- | --- | --- | --- | --- | --- | --- | --- | --- | --- | --- | --- | --- | --- | --- | --- | --- | --- | --- | --- | --- | --- | --- | --- | --- | --- | --- | --- | --- |
| pos | 241 | 913 | 2110 | 3037 | 3267 | 5388 | 5986 | 6539 | 6954 | 11288 | 11289 | 11290 | 11291 | 11292 | 11293 | 11294 | 11295 | 11296 | 14120 | 14408 | 14676 | 15279 | 16176 | 21765 | 21766 | 21767 | 21768 | 21769 | 21770 | 21991 | 21992 | 21993 | 23063 | 23271 | 23403 | 23604 | 23709 | 24506 | 24914 | 27972 | 28006 | 28048 | 28095 | 28111 | 28271 | 28280 | 28281 | 28282 | 28881 | 28882 | 28883 | 28887 | 28977 |  |
| ref | C | C | C | C | C | C | C | C | T | T | C | T | G | G | T | T | T | T | C | C | C | C | T | T | A | C | A | T | G | T | T | A | A | C | A | C | C | T | G | C | C | G | A | A | A | G | A | T | G | G | G | C | C |  |
| B.1.1.7 | T | T | T | T | T | A | T | C | C | - | - | - | - | - | - | - | - | - | T | T | T | T | C | - | - | - | - | - | - | - | - | - | T | A | G | A | T | G | C | T | C | T | T | G | - | C | T | A | A | A | C | C | T |  |
| B.1.1.7 | T | T | T | T | T | A | T | T | C | - | - | - | - | - | - | - | - | - | T | T | T | T | C | - | - | - | - | - | - | - | - | - | T | A | G | A | T | G | C | T | T | T | T | G | - | C | T | A | G | G | G | T | C | #237642 |
| B | C | C | C | C | C | C | C | C | T | T | C | T | G | G | T | T | T | T | C | C | C | C | T | T | A | C | A | T | G | T | T | A | A | C | A | C | C | T | G | C | C | G | A | A | A | G | A | T | G | G | G | T | C |  |
| mut |  |  |  |  |  |  |  | C>T |  |  |  |  |  |  |  |  |  |  |  |  |  |  |  |  |  |  |  |  |  |  |  |  |  |  |  |  |  |  |  |  | C>T |  |  |  |  |  |  |  |  |  |  |  |  |  |

|  |  |  |  |  |  |  |  |  |  |  |  |  |  |  |  |  |  |  |  |  |  |  |  |  |
| --- | --- | --- | --- | --- | --- | --- | --- | --- | --- | --- | --- | --- | --- | --- | --- | --- | --- | --- | --- | --- | --- | --- | --- | --- |
| pos | 2110 | 3391 | 3602 | 7042 | 10870 | 11833 | 14120 | 18465 | 19366 | 21161 | 21991 | 21992 | 21993 | 21994 | 21995 | 23429 | 23593 | 27879 | 27880 | 27881 | 27882 | 27883 | 28095 |  |
| ref | C | T | C | G | G | C | C | T | C | C | T | T | A | T | T | G | G | C | A | C | G | C | A |  |
| B.1.1.7 | T | C | C | T | G | T | T | C | T | C | T | T | A | C | C | G | G | C | A | C | G | C | T |  |
| B.1.1.7 | T | C | C | T | G | T | T | C | T | C | T | T | A | C | C | G | G | A | G | T | C | T | A | #238621 |
| B.1.1.7 | C | T | T | G | T | C | C | T | C | T | - | - | - | T | T | T | T | A | G | T | C | T | A |  |
| mut |  |  |  |  |  |  |  |  |  |  |  |  |  |  |  |  |  |  |  |  |  |  |  |  |

|  |  |  |  |  |  |  |  |  |  |  |  |  |  |  |  |  |  |  |  |  |  |  |  |  |  |  |  |  |  |  |  |  |  |  |
| --- | --- | --- | --- | --- | --- | --- | --- | --- | --- | --- | --- | --- | --- | --- | --- | --- | --- | --- | --- | --- | --- | --- | --- | --- | --- | --- | --- | --- | --- | --- | --- | --- | --- | --- |
| pos | 686 | 687 | 688 | 689 | 690 | 691 | 692 | 693 | 694 | 1191 | 1267 | 4181 | 5184 | 6402 | 7124 | 8986 | 9053 | 9891 | 10029 | 11201 | 11332 | 11418 | 12946 | 18176 | 19220 | 20262 | 21846 | 25562 | 26607 | 27476 | 27739 | 27874 | 28916 |  |
| ref | A | A | G | T | C | A | T | T | T | C | C | G | C | C | C | C | G | C | C | A | A | T | T | C | C | A | C | A | C | C | C | C | G |  |
| B.1.617.2 | A | A | G | T | C | A | T | T | T | C | C | T | C | T | T | T | T | C | T | G | G | T | T | C | T | A | T | A | C | C | C | T | T |  |
| B.1.617.2 | - | - | - | - | - | - | - | - | - | C | C | T | C | T | T | T | T | C | T | G | G | T | T | C | T | A | T | G | T | T | T | C | G | #248276 |
| B.1.617.2 | A | A | G | T | C | A | T | T | T | T | T | G | T | C | C | C | G | T | C | A | A | C | C | T | C | G | C | G | C | C | T | C | G |  |
| mut | A>- | A>- | G>- | T>- | C>- | A>- | T>- | T>- | T>- |  |  |  |  |  |  |  |  |  |  |  |  |  |  |  |  |  |  |  | C>T | C>T |  |  |  |  |

|  |  |  |  |  |  |  |  |  |  |  |  |  |  |  |  |  |
| --- | --- | --- | --- | --- | --- | --- | --- | --- | --- | --- | --- | --- | --- | --- | --- | --- |
| pos | 1002 | 8603 | 11112 | 12308 | 17615 | 19164 | 23683 | 25437 | 26787 | 27874 | 27875 | 27881 | 27882 | 27883 | 29466 |  |
| ref | A | T | T | C | A | C | C | G | G | C | T | C | G | C | C |  |
| B.1.1.7 | A | C | C | A | G | T | C | T | A | C | T | C | G | C | T |  |
| B.1.1.7 | G | C | C | A | G | T | T | T | A | T | C | T | C | T | T | #251428 |
| B.1.1.7 | A | T | T | C | A | C | C | G | G | T | C | T | C | T | C |  |
| mut | A>G |  |  |  |  |  | C>T |  |  |  |  |  |  |  | C>T |  |

|  |  |  |  |  |  |  |  |  |  |  |  |  |  |  |  |  |  |  |  |  |  |  |  |  |  |  |  |  |  |  |  |  |  |  |  |  |  |  |  |  |  |  |  |  |  |  |  |  |  |  |  |  |  |  |  |  |  |  |  |  |  |  |  |
| --- | --- | --- | --- | --- | --- | --- | --- | --- | --- | --- | --- | --- | --- | --- | --- | --- | --- | --- | --- | --- | --- | --- | --- | --- | --- | --- | --- | --- | --- | --- | --- | --- | --- | --- | --- | --- | --- | --- | --- | --- | --- | --- | --- | --- | --- | --- | --- | --- | --- | --- | --- | --- | --- | --- | --- | --- | --- | --- | --- | --- | --- | --- | --- |
| pos | 143 | 174 | 388 | 398 | 913 | 1059 | 2110 | 2782 | 3267 | 4113 | 4456 | 5388 | 5986 | 6954 | 9867 | 11312 | 14120 | 14676 | 15279 | 15720 | 16176 | 16500 | 16733 | 19390 | 20262 | 21765 | 21766 | 21767 | 21768 | 21769 | 21770 | 21846 | 21898 | 21991 | 21992 | 21993 | 22320 | 23012 | 23063 | 23271 | 23604 | 23664 | 23709 | 24506 | 24904 | 24914 | 25517 | 25563 | 27925 | 27972 | 28048 | 28095 | 28111 | 28280 | 28281 | 28282 | 28869 | 28881 | 28882 | 28883 | 28975 | 28977 |  |
| ref | A | G | G | C | C | C | C | G | C | C | C | C | C | T | T | G | C | C | C | C | T | A | C | C | A | T | A | C | A | T | G | C | G | T | T | A | A | G | A | C | C | C | C | T | C | G | C | G | C | C | G | A | A | G | A | T | C | G | G | G | G | C |  |
| B.1.1.7 | A | G | G | C | T | C | T | T | T | T | T | A | T | C | T | A | T | T | T | C | C | A | T | T | A | - | - | - | - | - | - | C | G | - | - | - | A | G | T | A | A | C | T | G | T | C | C | G | C | T | T | T | G | C | T | A | C | A | A | C | G | T |  |
| B.1.526 | A | G | A | C | T | C | T | T | T | T | T | A | T | C | T | G | C | C | C | T | T | C | C | C | G | T | A | C | A | T | G | T | T | T | T | A | G | A | A | C | C | T | C | T | C | G | T | T | T | C | G | A | A | G | A | T | T | G | G | G | A | C | #254203 |
| B.1.526 | G | T | G | T | C | T | C | G | C | C | C | C | C | T | C | G | C | C | C | T | T | C | C | C | G | T | A | C | A | T | G | T | T | T | T | A | G | A | A | C | C | T | C | T | C | G | T | T | T | C | G | A | A | G | A | T | T | G | G | G | A | C |  |
| mut |  |  | G>A |  |  |  |  |  |  |  |  |  |  |  |  |  |  |  |  |  |  |  |  |  |  |  |  |  |  |  |  |  |  |  |  |  |  |  |  |  |  |  |  |  |  |  |  |  |  |  |  |  |  |  |  |  |  |  |  |  |  |  |  |

|  |  |  |  |  |  |  |  |  |  |  |  |  |  |  |  |  |  |  |  |  |  |  |  |  |  |  |  |  |  |
| --- | --- | --- | --- | --- | --- | --- | --- | --- | --- | --- | --- | --- | --- | --- | --- | --- | --- | --- | --- | --- | --- | --- | --- | --- | --- | --- | --- | --- | --- |
| pos | 143 | 1588 | 4106 | 5577 | 6070 | 6101 | 6813 | 7201 | 8809 | 9435 | 10029 | 11573 | 12880 | 15720 | 21614 | 21658 | 21707 | 22992 | 23012 | 23664 | 24432 | 25968 | 27739 | 27758 | 28311 | 28869 | 28879 | 28975 |  |
| ref | A | A | A | T | C | G | C | A | C | T | C | T | C | C | C | C | C | G | G | C | A | A | C | G | C | C | T | G |  |
| B.1.526 | A | A | C | C | T | A | C | G | T | C | C | G | T | C | T | T | C | A | G | C | G | G | T | G | T | C | G | G |  |
| B.1.526 | A | G | C | C | T | A | C | G | T | C | C | G | T | C | T | T | T | G | A | T | A | A | C | T | C | T | T | A | #260179 |
| B.1.526 | G | A | A | T | C | G | T | A | C | T | T | T | C | T | C | C | T | G | A | T | A | A | C | G | C | T | T | A |  |
| mut |  | A>G |  |  |  |  |  |  |  |  |  |  |  |  |  |  |  |  |  |  |  |  |  | G>T |  |  |  |  |  |

|  |  |  |  |  |  |  |  |  |  |  |  |  |  |  |  |
| --- | --- | --- | --- | --- | --- | --- | --- | --- | --- | --- | --- | --- | --- | --- | --- |
| pos | 7279 | 9086 | 15957 | 17122 | 22841 | 25563 | 25572 | 28254 | 28255 | 28256 | 28257 | 28258 | 28262 | 28708 |  |
| ref | C | G | G | G | G | G | C | A | T | C | T | A | G | C |  |
| P.1 | C | A | A | T | G | T | C | A | T | C | T | A | G | T |  |
| P.1 | C | A | A | T | G | T | C | T | C | T | C | G | A | T | #260769 |
| P.1 | T | G | G | G | A | G | T | T | C | T | C | G | A | C |  |
| mut |  |  |  |  |  |  |  | A>T |  |  |  |  |  | C>T |  |

|  |  |  |  |  |  |  |  |  |  |  |  |  |  |  |  |  |  |  |  |  |  |  |  |  |  |  |  |  |  |  |  |  |  |  |  |  |  |  |  |  |  |  |  |  |  |  |  |  |  |  |  |  |  |  |
| --- | --- | --- | --- | --- | --- | --- | --- | --- | --- | --- | --- | --- | --- | --- | --- | --- | --- | --- | --- | --- | --- | --- | --- | --- | --- | --- | --- | --- | --- | --- | --- | --- | --- | --- | --- | --- | --- | --- | --- | --- | --- | --- | --- | --- | --- | --- | --- | --- | --- | --- | --- | --- | --- | --- |
| pos | 913 | 1059 | 2323 | 3267 | 5388 | 5986 | 6954 | 9127 | 9867 | 12162 | 12357 | 14403 | 14676 | 15243 | 15279 | 16176 | 16500 | 17615 | 20262 | 21765 | 21766 | 21767 | 21768 | 21769 | 21770 | 21846 | 21991 | 21992 | 21993 | 22320 | 23012 | 23063 | 23271 | 23604 | 23664 | 23709 | 24506 | 24914 | 25517 | 25563 | 27925 | 27972 | 28048 | 28111 | 28280 | 28281 | 28282 | 28869 | 28881 | 28882 | 28883 | 28975 | 28977 |  |
| ref | C | C | G | C | C | C | T | T | T | A | C | C | C | G | C | T | A | A | A | T | A | C | A | T | G | C | T | T | A | A | G | A | C | C | C | C | T | G | C | G | C | C | G | A | G | A | T | C | G | G | G | G | C |  |
| B.1.1.7 | T | C | G | T | A | T | C | T | T | G | C | C | T | G | T | C | A | G | A | - | - | - | - | - | - | C | - | - | - | A | G | T | A | A | C | T | G | C | C | G | C | T | T | G | C | T | A | C | A | A | C | G | T |  |
| B.1.526 | T | C | G | T | A | T | C | C | T | G | T | T | C | T | C | T | C | A | G | T | A | C | A | T | G | T | T | T | A | G | A | A | C | C | T | C | T | G | T | T | T | C | G | A | G | A | T | T | G | G | G | A | C | #263782 |
| B.1.526 | C | T | A | C | C | C | T | T | C | A | C | T | C | T | C | T | C | A | G | T | A | C | A | T | G | T | T | T | A | G | A | A | C | C | T | C | T | G | T | T | T | C | G | A | G | A | T | T | G | G | G | A | C |  |
| mut |  |  |  |  |  |  |  | T>C |  |  | C>T |  |  |  |  |  |  |  |  |  |  |  |  |  |  |  |  |  |  |  |  |  |  |  |  |  |  |  |  |  |  |  |  |  |  |  |  |  |  |  |  |  |  |  |

|  |  |  |  |  |  |  |  |  |  |  |  |  |  |  |  |
| --- | --- | --- | --- | --- | --- | --- | --- | --- | --- | --- | --- | --- | --- | --- | --- |
| pos | 583 | 1437 | 8668 | 10190 | 23757 | 27371 | 27903 | 28273 | 28274 | 28276 | 28280 | 28281 | 28282 | 29358 |  |
| ref | C | C | C | T | C | C | C | A | A | G | G | A | T | C |  |
| B.1.214 | C | C | C | T | C | T | C | T | G | C | A | A | T | C |  |
| B.1.214 | C | C | C | T | C | T | C | A | A | G | C | T | A | C | #264915 |
| B.1.214 | T | T | T | G | T | C | T | A | A | G | C | T | A | T |  |
| mut |  |  |  |  |  |  |  |  |  |  |  |  |  | T>C |  |

|  |  |  |  |  |  |  |  |  |  |  |  |  |  |  |  |  |  |  |  |
| --- | --- | --- | --- | --- | --- | --- | --- | --- | --- | --- | --- | --- | --- | --- | --- | --- | --- | --- | --- |
| pos | 1738 | 2110 | 4900 | 7042 | 12162 | 13887 | 14053 | 14120 | 15720 | 17615 | 17762 | 21137 | 21722 | 22624 | 22675 | 25135 | 28095 | 28967 |  |
| ref | G | C | C | G | A | C | C | C | C | A | C | A | T | C | C | G | A | A |  |
| B.1.1.7 | T | C | T | G | G | T | T | C | T | G | C | G | C | C | C | T | A | G |  |
| B.1.1.7 | T | C | T | G | G | C | C | T | C | A | T | A | T | T | T | G | T | A | #277217 |
| B.1.1.7 | G | T | C | T | A | C | C | T | C | A | T | A | T | T | T | G | T | A |  |
| mut |  |  |  |  |  |  |  |  |  |  |  |  |  |  |  |  |  |  |  |

|  |  |  |  |  |  |  |  |  |  |  |  |  |  |  |  |  |  |  |  |  |  |  |  |  |  |  |  |  |  |  |  |  |  |  |  |  |  |  |  |  |  |  |  |  |  |  |  |  |  |  |  |  |  |  |  |  |
| --- | --- | --- | --- | --- | --- | --- | --- | --- | --- | --- | --- | --- | --- | --- | --- | --- | --- | --- | --- | --- | --- | --- | --- | --- | --- | --- | --- | --- | --- | --- | --- | --- | --- | --- | --- | --- | --- | --- | --- | --- | --- | --- | --- | --- | --- | --- | --- | --- | --- | --- | --- | --- | --- | --- | --- | --- |
| pos | 913 | 3177 | 3267 | 5388 | 5986 | 6954 | 9559 | 11288 | 11289 | 11290 | 11291 | 11292 | 11293 | 11294 | 11295 | 11296 | 14676 | 15096 | 15279 | 15324 | 16176 | 21765 | 21766 | 21767 | 21768 | 21769 | 21770 | 21991 | 21992 | 21993 | 23063 | 23271 | 23604 | 23709 | 24506 | 24914 | 25516 | 25785 | 27532 | 27637 | 27661 | 27972 | 28048 | 28095 | 28111 | 28271 | 28280 | 28281 | 28282 | 28697 | 28881 | 28882 | 28883 | 28916 | 28977 |  |
| ref | C | C | C | C | C | T | C | T | C | T | G | G | T | T | T | T | C | T | C | C | T | T | A | C | A | T | G | T | T | A | A | C | C | C | T | G | C | G | C | G | C | C | G | A | A | A | G | A | T | C | G | G | G | G | C |  |
| B.1.1.7 | T | T | T | A | T | C | T | - | - | - | - | - | - | - | - | - | T | C | T | C | C | - | - | - | - | - | - | - | - | - | T | A | A | T | G | C | T | T | - | A | T | T | T | T | G | - | C | T | A | T | A | A | C | G | T |  |
| B.1.1.7 | T | T | T | A | T | C | T | - | - | - | - | - | - | - | - | - | T | C | T | C | C | - | - | - | - | - | - | - | - | - | T | A | A | T | G | C | T | T | - | A | T | T | T | T | G | - | C | T | A | T | T | G | G | T | C | #286975 |
| B.1 | C | C | C | C | C | T | C | T | C | T | G | G | T | T | T | T | C | T | C | T | T | T | A | C | A | T | G | T | T | A | A | C | C | C | T | G | C | G | C | G | C | C | G | A | A | A | G | A | T | C | T | G | G | G | C |  |
| mut |  |  |  |  |  |  |  |  |  |  |  |  |  |  |  |  |  |  |  |  |  |  |  |  |  |  |  |  |  |  |  |  |  |  |  |  |  |  |  |  |  |  |  |  |  |  |  |  |  |  |  |  |  | G>T |  |  |

|  |  |  |  |  |  |  |  |  |  |  |  |  |  |  |  |  |  |
| --- | --- | --- | --- | --- | --- | --- | --- | --- | --- | --- | --- | --- | --- | --- | --- | --- | --- |
| pos | 826 | 884 | 4321 | 4720 | 10271 | 16914 | 19677 | 22027 | 22029 | 22030 | 22031 | 22032 | 22033 | 22034 | 24325 | 27338 |  |
| ref | T | C | C | G | G | G | G | T | A | G | T | T | C | A | A | A |  |
| AY.114 | T | T | C | T | G | T | T | T | - | - | - | - | - | - | T | A |  |
| AY.114 | T | T | C | T | G | T | T | G | A | A | A | G | T | G | A | A | #293131 |
| AY.118 | C | C | T | G | A | G | G | G | A | A | A | G | T | G | A | T |  |
| mut |  |  |  |  |  |  |  |  |  |  |  |  |  |  |  | T>A |  |

|  |  |  |  |  |  |  |  |  |  |  |  |  |  |  |  |
| --- | --- | --- | --- | --- | --- | --- | --- | --- | --- | --- | --- | --- | --- | --- | --- |
| pos | 884 | 4720 | 7851 | 14829 | 16914 | 17193 | 19677 | 22027 | 22029 | 22030 | 22031 | 22032 | 22033 | 22034 |  |
| ref | C | G | C | G | G | G | G | T | A | G | T | T | C | A |  |
| AY.4 | C | G | T | T | G | T | G | T | - | - | - | - | - | - |  |
| AY.4 | C | G | T | T | G | T | G | G | A | A | A | G | T | G | #295320 |
| AY.114 | T | T | C | G | T | G | T | G | A | A | A | G | T | G |  |
| mut |  |  |  |  |  |  |  |  |  |  |  |  |  |  |  |

|  |  |  |  |  |  |  |  |  |  |  |  |  |  |  |  |  |
| --- | --- | --- | --- | --- | --- | --- | --- | --- | --- | --- | --- | --- | --- | --- | --- | --- |
| pos | 805 | 884 | 4237 | 4720 | 7851 | 16914 | 17040 | 19677 | 22027 | 22029 | 22030 | 22031 | 22032 | 22033 | 22034 |  |
| ref | G | C | T | G | C | G | T | G | T | A | G | T | T | C | A |  |
| AY.4 | T | C | C | G | T | G | C | G | T | - | - | - | - | - | - |  |
| AY.4 | T | C | C | G | T | G | C | G | G | A | A | A | G | T | G | #296729 |
| AY.114 | G | T | T | T | C | T | T | T | G | A | A | A | G | T | G |  |
| mut |  |  |  |  |  |  |  |  |  |  |  |  |  |  |  |  |

|  |  |  |  |  |  |  |  |  |  |  |  |
| --- | --- | --- | --- | --- | --- | --- | --- | --- | --- | --- | --- |
| pos | 526 | 7851 | 19524 | 21846 | 23856 | 25959 | 27875 | 27881 | 27882 | 27883 |  |
| ref | G | C | C | C | G | T | T | C | G | C |  |
| AY.4 | G | T | C | T | T | C | T | C | G | C |  |
| AY.4 | G | T | C | T | T | T | C | T | C | T | #296997 |
| AY.98 | T | C | T | C | G | T | C | T | C | T |  |
| mut |  |  |  |  |  |  |  |  |  |  |  |

|  |  |  |  |  |  |  |  |  |  |  |  |  |  |  |  |
| --- | --- | --- | --- | --- | --- | --- | --- | --- | --- | --- | --- | --- | --- | --- | --- |
| pos | 1684 | 6285 | 8772 | 11563 | 21641 | 22026 | 22027 | 22029 | 22030 | 22031 | 22032 | 22033 | 22034 | 26042 |  |
| ref | C | C | C | C | G | G | T | A | G | T | T | C | A | C |  |
| AY.39 | T | T | T | T | T | G | T | - | - | - | - | - | - | T |  |
| AY.39 | T | T | T | T | T | G | G | A | A | A | G | T | G | T | #300908 |
| AY.39 | C | C | C | C | G | T | G | A | A | A | G | T | G | C |  |
| mut |  |  |  |  |  |  |  |  |  |  |  |  |  | C>T |  |

|  |  |  |  |  |  |  |  |  |  |  |  |  |  |  |  |  |  |  |
| --- | --- | --- | --- | --- | --- | --- | --- | --- | --- | --- | --- | --- | --- | --- | --- | --- | --- | --- |
| pos | 884 | 2047 | 4720 | 7851 | 16914 | 19292 | 19677 | 21048 | 21485 | 22027 | 22029 | 22030 | 22031 | 22032 | 22033 | 22034 | 27513 |  |
| ref | C | T | G | C | G | G | G | T | G | T | A | G | T | T | C | A | C |  |
| AY.4 | C | C | G | T | G | T | G | C | T | T | - | - | - | - | - | - | T |  |
| AY.4 | C | C | G | T | G | T | G | C | T | G | A | A | A | G | T | G | T | #313917 |
| AY.114 | T | T | T | C | T | G | T | T | G | G | A | A | A | G | T | G | C |  |
| mut |  |  |  |  |  |  |  |  |  |  |  |  |  |  |  |  | C>T |  |

|  |  |  |  |  |  |  |  |  |  |  |  |  |  |  |  |  |
| --- | --- | --- | --- | --- | --- | --- | --- | --- | --- | --- | --- | --- | --- | --- | --- | --- |
| pos | 3087 | 4543 | 6896 | 9712 | 9714 | 14184 | 20571 | 21614 | 25614 | 26013 | 27881 | 28093 | 28254 | 28310 | 29762 |  |
| ref | T | C | C | C | C | C | T | C | C | C | C | C | A | C | C |  |
| B.1.351 | C | T | T | T | T | T | T | C | T | C | C | C | C | C | C |  |
| B.1.351 | C | T | T | T | T | C | C | T | C | T | T | T | A | T | T | #320592 |
| B.1.351 | T | C | C | C | C | C | T | T | C | T | C | C | A | T | C |  |
| mut |  |  |  |  |  |  | T>C |  |  |  | C>T | C>T |  |  | C>T |  |

|  |  |  |  |  |  |  |  |  |  |  |  |  |  |  |  |  |  |  |  |  |  |  |  |  |  |
| --- | --- | --- | --- | --- | --- | --- | --- | --- | --- | --- | --- | --- | --- | --- | --- | --- | --- | --- | --- | --- | --- | --- | --- | --- | --- |
| pos | 1181 | 1191 | 1267 | 4181 | 5184 | 6402 | 7124 | 8266 | 8986 | 9053 | 9891 | 10029 | 11201 | 11332 | 11418 | 12946 | 16049 | 19220 | 20262 | 24700 | 24928 | 27739 | 27874 | 28916 |  |
| ref | G | C | C | G | C | C | C | C | C | G | C | C | A | A | T | T | C | C | A | C | G | C | C | G |  |
| AY.5 | T | C | C | T | C | T | T | T | T | T | C | T | G | G | T | T | T | T | A | T | T | C | T | T |  |
| AY.5 | T | C | C | T | C | T | T | T | T | T | C | T | G | G | T | T | T | T | A | C | G | T | C | G | #325811 |
| B.1.617.2 | G | T | T | G | T | C | C | C | C | G | T | C | A | A | C | C | C | C | G | C | G | T | C | G |  |
| mut |  |  |  |  |  |  |  |  |  |  |  |  |  |  |  |  |  |  |  |  |  |  |  |  |  |

|  |  |  |  |  |  |  |  |  |  |  |  |  |  |  |
| --- | --- | --- | --- | --- | --- | --- | --- | --- | --- | --- | --- | --- | --- | --- |
| pos | 204 | 526 | 7851 | 18197 | 18348 | 19524 | 21846 | 22051 | 27384 | 27875 | 27881 | 27882 | 27883 |  |
| ref | G | G | C | G | C | C | C | G | T | T | C | G | C |  |
| AY.4 | T | G | T | A | T | C | T | C | C | T | C | G | C |  |
| AY.4 | T | G | T | A | T | C | T | C | C | C | T | C | T | #326037 |
| AY.98 | G | T | C | G | C | T | C | G | T | C | T | C | T |  |
| mut |  |  |  |  |  |  |  |  |  |  |  |  |  |  |

|  |  |  |  |  |  |  |  |  |  |  |  |  |  |  |  |  |  |  |  |  |  |  |  |  |  |  |  |  |  |
| --- | --- | --- | --- | --- | --- | --- | --- | --- | --- | --- | --- | --- | --- | --- | --- | --- | --- | --- | --- | --- | --- | --- | --- | --- | --- | --- | --- | --- | --- |
| pos | 1190 | 3177 | 5023 | 6568 | 9529 | 9559 | 10396 | 12473 | 15096 | 17615 | 20962 | 22093 | 22094 | 22227 | 25516 | 25785 | 26730 | 27468 | 27532 | 27637 | 27661 | 28095 | 28697 | 28881 | 28882 | 28883 | 28916 | 28977 |  |
| ref | C | C | G | C | A | C | G | C | T | A | G | G | G | C | C | G | G | A | C | G | C | A | C | G | G | G | G | C |  |
| Q.3 | T | C | A | T | G | C | T | T | T | G | A | C | C | T | C | G | C | A | C | G | C | A | C | A | A | C | G | T |  |
| Q.3 | T | C | A | T | G | C | T | T | T | G | A | C | C | T | C | G | C | G | C | G | C | A | T | T | G | G | T | C | #330618 |
| B.1.1.7 | C | T | G | C | A | T | G | C | C | A | G | G | G | C | T | T | G | A | - | A | T | T | T | T | G | G | T | C |  |
| mut |  |  |  |  |  |  |  |  |  |  |  |  |  |  |  |  |  | A>G |  |  |  |  |  |  |  |  |  |  |  |

|  |  |  |  |  |  |  |  |  |  |  |  |  |  |  |  |  |  |  |  |  |  |  |  |
| --- | --- | --- | --- | --- | --- | --- | --- | --- | --- | --- | --- | --- | --- | --- | --- | --- | --- | --- | --- | --- | --- | --- | --- |
| pos | 1191 | 1267 | 1387 | 1877 | 2872 | 4890 | 5584 | 5644 | 7417 | 10323 | 11514 | 12946 | 13019 | 18176 | 19160 | 19519 | 20262 | 20718 | 22227 | 22346 | 27368 | 27739 |  |
| ref | C | C | C | T | T | C | A | T | C | A | C | T | C | C | C | T | A | G | C | G | A | C |  |
| AY.14 | T | T | C | G | T | C | A | T | C | A | C | C | C | T | T | T | G | T | C | G | A | T |  |
| AY.67 | T | T | C | G | T | T | G | C | T | G | T | T | T | C | C | C | A | G | T | T | G | C | #334514 |
| AY.67 | C | C | T | T | A | T | G | C | T | A | T | T | T | C | C | C | A | G | T | T | G | C |  |
| mut |  |  |  |  |  |  |  |  |  | A>G |  |  |  |  |  |  |  |  |  |  |  |  |  |

|  |  |  |  |  |  |  |  |  |  |  |  |  |  |  |  |  |  |
| --- | --- | --- | --- | --- | --- | --- | --- | --- | --- | --- | --- | --- | --- | --- | --- | --- | --- |
| pos | 884 | 2258 | 4720 | 9508 | 12073 | 15237 | 16914 | 19677 | 22027 | 22029 | 22030 | 22031 | 22032 | 22033 | 22034 | 25872 |  |
| ref | C | G | G | T | C | C | G | G | T | A | G | T | T | C | A | C |  |
| AY.42 | C | A | G | C | T | T | G | G | T | - | - | - | - | - | - | T |  |
| AY.42 | C | A | G | C | T | T | G | G | G | A | A | A | G | T | G | T | #336012 |
| AY.114 | T | G | T | T | C | C | T | T | G | A | A | A | G | T | G | C |  |
| mut |  |  |  |  |  |  |  |  |  |  |  |  |  |  |  | C>T |  |

|  |  |  |  |  |  |  |  |  |  |  |  |  |  |  |  |
| --- | --- | --- | --- | --- | --- | --- | --- | --- | --- | --- | --- | --- | --- | --- | --- |
| pos | 507 | 508 | 509 | 6968 | 11115 | 11456 | 15576 | 17236 | 17795 | 24208 | 26270 | 28858 | 29050 | 29509 |  |
| ref | A | T | G | C | G | A | T | A | C | C | C | A | G | C |  |
| AY.103 | T | C | A | C | G | A | T | G | T | T | C | A | G | C |  |
| AY.3 | T | C | A | C | G | G | C | A | C | C | T | C | A | T | #336516 |
| AY.3 | A | T | G | T | T | G | C | A | C | C | T | C | A | T |  |
| mut |  |  |  |  |  |  |  |  |  |  |  |  |  |  |  |

|  |  |  |  |  |  |  |  |  |  |  |  |  |  |  |  |  |  |
| --- | --- | --- | --- | --- | --- | --- | --- | --- | --- | --- | --- | --- | --- | --- | --- | --- | --- |
| pos | 884 | 3884 | 4720 | 7173 | 7756 | 7851 | 9170 | 16914 | 19677 | 22027 | 22029 | 22030 | 22031 | 22032 | 22033 | 22034 |  |
| ref | C | C | G | C | G | C | C | G | G | T | A | G | T | T | C | A |  |
| AY.4 | C | T | G | T | T | T | T | G | G | T | - | - | - | - | - | - |  |
| AY.4 | C | T | G | T | T | T | T | G | G | G | A | A | A | G | T | G | #346285 |
| AY.114 | T | C | T | C | G | C | C | T | T | G | A | A | A | G | T | G |  |
| mut |  |  |  |  |  |  |  |  |  |  |  |  |  |  |  |  |  |

|  |  |  |  |  |  |  |  |  |  |  |  |  |  |  |  |  |  |
| --- | --- | --- | --- | --- | --- | --- | --- | --- | --- | --- | --- | --- | --- | --- | --- | --- | --- |
| pos | 884 | 4720 | 10691 | 12071 | 16914 | 19160 | 19677 | 20968 | 21793 | 22027 | 22029 | 22030 | 22031 | 22032 | 22033 | 22034 |  |
| ref | C | G | A | G | G | C | G | G | G | T | A | G | T | T | C | A |  |
| AY.116.1 | C | G | G | A | G | T | G | A | A | T | - | - | - | - | - | - |  |
| AY.116.1 | C | G | G | A | G | T | G | A | A | G | A | A | A | G | T | G | #349841 |
| AY.114 | T | T | A | G | T | C | T | G | G | G | A | A | A | G | T | G |  |
| mut |  |  |  |  |  |  |  |  |  |  |  |  |  |  |  |  |  |

|  |  |  |  |  |  |  |  |  |  |  |  |  |  |  |  |  |  |  |  |  |  |
| --- | --- | --- | --- | --- | --- | --- | --- | --- | --- | --- | --- | --- | --- | --- | --- | --- | --- | --- | --- | --- | --- |
| pos | 208 | 884 | 4720 | 6040 | 7926 | 14014 | 14396 | 16914 | 19677 | 20832 | 21846 | 22027 | 22029 | 22030 | 22031 | 22032 | 22033 | 22034 | 26873 | 29700 |  |
| ref | G | C | G | C | C | T | C | G | G | A | C | T | A | G | T | T | C | A | C | A |  |
| AY.44 | A | C | G | T | T | G | C | G | G | A | C | T | - | - | - | - | - | - | C | G |  |
| AY.44 | A | C | G | T | T | G | T | G | G | G | C | T | - | - | A | G | T | G | T | G | #355137 |
| AY.114 | G | T | T | C | C | T | C | T | T | A | T | G | A | A | A | G | T | G | C | A |  |
| mut |  |  |  |  |  |  | C>T |  |  | A>G |  |  |  |  |  |  |  |  | C>T | A>G |  |

|  |  |  |  |  |  |  |  |  |  |  |  |  |
| --- | --- | --- | --- | --- | --- | --- | --- | --- | --- | --- | --- | --- |
| pos | 526 | 5991 | 7851 | 18651 | 19524 | 21846 | 23625 | 27875 | 27881 | 27882 | 27883 |  |
| ref | G | A | C | G | C | C | C | T | C | G | C |  |
| AY.4 | G | C | T | T | C | T | T | T | C | G | C |  |
| AY.4 | G | C | T | T | C | T | T | C | T | C | T | #356439 |
| AY.98 | T | A | C | G | T | C | C | C | T | C | T |  |
| mut |  |  |  |  |  |  |  |  |  |  |  |  |

|  |  |  |  |  |  |  |  |  |  |  |  |  |  |  |  |  |  |  |  |  |  |  |  |  |  |  |  |  |  |  |  |  |  |  |  |  |  |  |  |  |  |  |  |  |  |  |  |  |  |  |  |  |  |
| --- | --- | --- | --- | --- | --- | --- | --- | --- | --- | --- | --- | --- | --- | --- | --- | --- | --- | --- | --- | --- | --- | --- | --- | --- | --- | --- | --- | --- | --- | --- | --- | --- | --- | --- | --- | --- | --- | --- | --- | --- | --- | --- | --- | --- | --- | --- | --- | --- | --- | --- | --- | --- | --- |
| pos | 210 | 745 | 4181 | 6402 | 7124 | 8986 | 9053 | 9165 | 10029 | 10323 | 11201 | 11332 | 12772 | 15240 | 15451 | 16466 | 19220 | 21255 | 21618 | 21846 | 21987 | 22029 | 22030 | 22031 | 22032 | 22033 | 22034 | 22917 | 22995 | 23604 | 24410 | 25469 | 26767 | 27638 | 27752 | 28248 | 28249 | 28250 | 28251 | 28252 | 28253 | 28271 | 28461 | 28877 | 28878 | 28881 | 28882 | 28883 | 28916 | 29402 | 29742 | 29834 |  |
| ref | G | C | G | C | C | C | G | C | C | A | A | A | A | C | G | C | C | G | C | C | G | A | G | T | T | C | A | T | C | C | G | C | T | T | C | G | A | T | T | T | C | A | A | A | G | G | G | G | G | G | G | T |  |
| AY.113 | T | C | T | T | T | T | T | T | T | G | G | G | G | T | A | T | T | G | G | T | A | - | - | - | - | - | - | G | A | G | A | T | C | C | T | - | - | - | - | - | - | - | G | A | G | T | G | G | T | T | T | T |  |
| B.1.617.2 | T | C | T | T | T | T | T | T | T | G | G | G | G | T | A | T | T | G | G | T | A | - | - | - | - | - | - | G | A | G | A | T | T | T | C | - | - | - | - | - | - | - | G | T | C | A | A | C | G | G | G | A | #357756 |
| B.1.1 | G | T | G | C | C | C | G | C | C | A | A | A | A | C | G | C | C | T | C | C | G | A | G | T | T | C | A | T | C | C | G | C | T | T | C | G | A | T | T | T | C | A | A | A | G | A | A | C | G | G | G | T |  |
| mut |  |  |  |  |  |  |  |  |  |  |  |  |  |  |  |  |  |  |  |  |  |  |  |  |  |  |  |  |  |  |  |  |  |  |  | G>- | A>- | T>- | T>- | T>- | C>- | A>- | A>G | A>T | G>C |  |  |  |  |  |  | T>A |  |

|  |  |  |  |  |  |  |  |  |  |  |  |  |  |  |  |  |  |
| --- | --- | --- | --- | --- | --- | --- | --- | --- | --- | --- | --- | --- | --- | --- | --- | --- | --- |
| pos | 1594 | 1779 | 7869 | 13026 | 22026 | 22027 | 22029 | 22030 | 22031 | 22032 | 22033 | 22034 | 25401 | 27527 | 28077 | 29362 |  |
| ref | C | C | C | C | G | T | A | G | T | T | C | A | G | C | G | C |  |
| AY.39 | T | T | T | T | G | T | - | - | - | - | - | - | T | T | T | T |  |
| AY.39 | T | T | T | T | T | G | A | A | A | G | T | G | T | T | T | T | #360755 |
| AY.39 | C | C | C | C | T | G | A | A | A | G | T | G | G | C | G | C |  |
| mut |  |  |  |  |  |  |  |  |  |  |  |  | G>T | C>T | G>T | C>T |  |

|  |  |  |  |  |  |  |  |  |  |  |  |  |  |  |  |  |  |
| --- | --- | --- | --- | --- | --- | --- | --- | --- | --- | --- | --- | --- | --- | --- | --- | --- | --- |
| pos | 884 | 2485 | 4720 | 7851 | 11824 | 16474 | 16914 | 18529 | 19677 | 22027 | 22029 | 22030 | 22031 | 22032 | 22033 | 22034 |  |
| ref | C | C | G | C | C | A | G | A | G | T | A | G | T | T | C | A |  |
| AY.4 | C | T | G | T | T | G | G | G | G | T | - | - | - | - | - | - |  |
| AY.4 | C | T | G | T | T | G | G | G | G | G | A | A | A | G | T | G | #362625 |
| AY.114 | T | C | T | C | C | A | T | A | T | G | A | A | A | G | T | G |  |
| mut |  |  |  |  |  |  |  |  |  |  |  |  |  |  |  |  |  |

|  |  |  |  |  |  |  |  |  |  |  |  |
| --- | --- | --- | --- | --- | --- | --- | --- | --- | --- | --- | --- |
| pos | 1181 | 2447 | 6408 | 8266 | 16049 | 16074 | 21846 | 24700 | 24928 | 25003 |  |
| ref | G | G | C | C | C | T | C | C | G | A |  |
| AY.5 | T | T | C | T | T | C | C | T | T | A |  |
| AY.5 | T | T | C | T | T | T | T | C | G | G | #364956 |
| AY.120 | G | G | T | C | C | T | T | C | G | G |  |
| mut |  |  |  |  |  |  |  |  |  |  |  |

|  |  |  |  |  |  |  |  |  |  |  |  |  |  |  |  |  |  |  |  |  |  |  |  |  |
| --- | --- | --- | --- | --- | --- | --- | --- | --- | --- | --- | --- | --- | --- | --- | --- | --- | --- | --- | --- | --- | --- | --- | --- | --- |
| pos | 1191 | 1267 | 1387 | 2872 | 4079 | 4890 | 5584 | 5644 | 5649 | 7417 | 10323 | 11514 | 12946 | 13019 | 18176 | 19160 | 19519 | 20262 | 20718 | 22227 | 22346 | 27368 | 27739 |  |
| ref | C | C | C | T | A | C | A | T | A | C | A | C | T | C | C | C | T | A | G | C | G | A | C |  |
| AY.67 | C | C | T | A | G | T | G | C | A | T | G | T | T | T | C | C | C | A | G | T | T | G | C |  |
| AY.67 | C | C | T | A | G | T | G | C | G | T | G | T | T | T | C | T | T | G | T | C | G | A | T | #366278 |
| AY.14 | T | T | C | T | A | C | A | T | A | C | A | C | C | C | T | T | T | G | T | C | G | A | T |  |
| mut |  |  |  |  |  |  |  |  | A>G |  |  |  |  |  |  |  |  |  |  |  |  |  |  |  |

|  |  |  |  |  |  |  |  |  |  |  |  |  |  |  |  |  |  |
| --- | --- | --- | --- | --- | --- | --- | --- | --- | --- | --- | --- | --- | --- | --- | --- | --- | --- |
| pos | 6040 | 6638 | 6661 | 7926 | 14014 | 16726 | 17236 | 24208 | 25323 | 27291 | 28073 | 28460 | 28585 | 29593 | 29700 | 29781 |  |
| ref | C | C | T | C | T | C | A | C | G | T | G | G | C | G | A | G |  |
| AY.103 | C | C | C | C | T | C | G | T | G | C | G | G | C | G | A | - |  |
| AY.44 | C | C | C | C | G | T | A | C | T | T | T | A | T | T | G | G | #367440 |
| AY.44 | T | T | T | T | G | T | A | C | G | T | T | A | C | T | G | G |  |
| mut |  |  |  |  |  |  |  |  | G>T |  |  |  | C>T |  |  |  |  |

|  |  |  |  |  |  |  |  |  |  |  |  |  |  |  |  |  |  |  |  |  |  |  |  |  |  |  |  |  |  |  |  |  |  |  |  |  |
| --- | --- | --- | --- | --- | --- | --- | --- | --- | --- | --- | --- | --- | --- | --- | --- | --- | --- | --- | --- | --- | --- | --- | --- | --- | --- | --- | --- | --- | --- | --- | --- | --- | --- | --- | --- | --- |
| pos | 512 | 526 | 1191 | 1267 | 4181 | 5184 | 6402 | 7124 | 8986 | 9053 | 9891 | 10029 | 11201 | 11332 | 11418 | 11455 | 12946 | 14170 | 15738 | 18176 | 19220 | 19524 | 20262 | 22224 | 26065 | 26936 | 27077 | 27739 | 27874 | 27875 | 27881 | 27882 | 27883 | 28916 | 29744 |  |
| ref | C | G | C | C | G | C | C | C | C | G | C | C | A | A | T | C | T | A | C | C | C | C | A | C | G | C | G | C | C | T | C | G | C | G | G |  |
| AY.98 | T | T | C | C | T | C | T | T | T | T | C | T | G | G | T | C | T | A | C | C | T | T | A | C | T | C | G | C | T | C | T | C | T | T | G |  |
| AY.8 | T | T | C | C | T | C | T | T | T | T | C | T | G | A | C | T | C | G | T | T | C | C | G | T | G | T | A | T | C | T | C | G | C | G | A | #368048 |
| AY.8 | C | G | T | T | G | T | C | C | C | G | T | C | A | A | C | T | C | G | T | T | C | C | G | T | G | T | A | T | C | T | C | G | C | G | A |  |
| mut |  |  |  |  |  |  |  |  |  |  |  |  |  |  |  |  |  |  |  |  |  |  |  |  |  |  |  |  |  |  |  |  |  |  |  |  |

|  |  |  |  |  |  |  |  |  |  |  |
| --- | --- | --- | --- | --- | --- | --- | --- | --- | --- | --- |
| pos | 4621 | 7851 | 8660 | 15351 | 16075 | 21648 | 21846 | 24928 | 27610 |  |
| ref | T | C | C | T | G | C | C | G | C |  |
| AY.5 | C | C | C | C | G | C | C | T | C |  |
| B.1.617.2 | C | C | C | C | G | T | T | G | T | #368521 |
| AY.4 | T | T | T | T | A | T | T | G | T |  |
| mut |  |  |  |  |  |  |  |  |  |  |

|  |  |  |  |  |  |  |  |  |  |  |  |  |  |  |  |  |  |  |  |  |  |  |  |  |  |  |  |  |  |  |  |  |  |  |
| --- | --- | --- | --- | --- | --- | --- | --- | --- | --- | --- | --- | --- | --- | --- | --- | --- | --- | --- | --- | --- | --- | --- | --- | --- | --- | --- | --- | --- | --- | --- | --- | --- | --- | --- |
| pos | 1729 | 2164 | 4181 | 4321 | 5184 | 5584 | 6402 | 7124 | 7401 | 7851 | 8986 | 9053 | 9891 | 10029 | 11201 | 11332 | 11418 | 11514 | 13019 | 14030 | 14829 | 17193 | 18086 | 19220 | 21846 | 21987 | 22227 | 22792 | 26416 | 27621 | 27874 | 28291 | 28916 |  |
| ref | G | G | G | C | C | A | C | C | C | C | C | G | C | C | A | A | T | C | C | G | G | G | C | C | C | G | C | C | G | G | C | C | G |  |
| AY.4.7 | G | T | T | C | C | A | T | T | C | T | T | T | C | T | G | G | T | C | C | G | T | T | C | T | T | G | C | C | T | G | T | T | T |  |
| AY.9 | G | T | T | C | C | A | C | C | T | C | C | G | T | C | A | A | C | T | T | A | G | G | T | C | C | A | T | T | G | A | C | C | G | #369130 |
| AY.9.2 | A | G | G | T | T | G | C | C | C | C | C | G | T | C | A | A | C | T | T | A | G | G | T | C | C | A | T | T | G | A | C | C | G |  |
| mut |  |  |  |  |  |  |  |  | C>T |  |  |  |  |  |  |  |  |  |  |  |  |  |  |  |  |  |  |  |  |  |  |  |  |  |

|  |  |  |  |  |  |  |  |  |  |  |  |  |  |  |  |  |  |  |  |
| --- | --- | --- | --- | --- | --- | --- | --- | --- | --- | --- | --- | --- | --- | --- | --- | --- | --- | --- | --- |
| pos | 2258 | 3688 | 4081 | 5526 | 12008 | 12073 | 15237 | 21846 | 21987 | 23263 | 23718 | 24928 | 25654 | 25872 | 26067 | 29253 | 29262 | 29711 |  |
| ref | G | C | T | C | C | C | C | C | G | A | C | G | G | C | T | C | G | G |  |
| AY.42 | A | T | A | C | C | T | T | T | G | C | T | G | G | T | C | C | G | G |  |
| AY.5 | A | T | A | C | C | T | T | C | A | A | C | T | T | C | T | T | T | T | #371112 |
| AY.5 | G | C | T | A | T | C | C | C | A | A | C | T | T | C | T | T | T | T |  |
| mut |  |  |  |  |  |  |  |  |  |  |  |  |  |  |  |  |  |  |  |

|  |  |  |  |  |  |  |  |  |  |  |  |  |  |  |  |  |  |  |  |  |  |  |  |  |  |  |  |  |  |  |  |  |
| --- | --- | --- | --- | --- | --- | --- | --- | --- | --- | --- | --- | --- | --- | --- | --- | --- | --- | --- | --- | --- | --- | --- | --- | --- | --- | --- | --- | --- | --- | --- | --- | --- |
| pos | 2911 | 3882 | 4181 | 5184 | 5584 | 5842 | 6402 | 6796 | 7124 | 8986 | 9053 | 9891 | 10029 | 11201 | 11332 | 11418 | 11456 | 11514 | 13019 | 15342 | 19220 | 20629 | 22227 | 23202 | 25352 | 27345 | 27874 | 28916 | 29050 | 29509 | 29781 |  |
| ref | T | T | G | C | A | C | C | A | C | C | G | C | C | A | A | T | A | C | C | C | C | C | C | C | G | A | C | G | G | C | G |  |
| AY.3 | T | T | T | C | A | C | T | A | T | T | T | C | T | G | G | T | G | C | C | C | T | C | C | C | G | A | T | T | A | T | G |  |
| AY.3 | T | T | T | C | A | C | T | A | T | T | T | C | T | G | G | T | G | C | C | C | C | T | T | C | T | T | C | G | G | C | - | #371354 |
| AY.26 | G | C | G | T | G | T | C | G | C | C | G | T | C | A | A | C | A | T | T | T | C | T | T | T | T | T | C | G | G | C | G |  |
| mut |  |  |  |  |  |  |  |  |  |  |  |  |  |  |  |  |  |  |  |  |  |  |  | T>C |  |  |  |  |  |  | G>- |  |

|  |  |  |  |  |  |  |  |  |  |  |  |  |  |  |  |  |  |  |  |  |
| --- | --- | --- | --- | --- | --- | --- | --- | --- | --- | --- | --- | --- | --- | --- | --- | --- | --- | --- | --- | --- |
| pos | 884 | 4720 | 6472 | 12385 | 16914 | 19404 | 19677 | 19764 | 21387 | 22027 | 22029 | 22030 | 22031 | 22032 | 22033 | 22034 | 25831 | 28568 | 29095 |  |
| ref | C | G | C | G | G | T | G | A | T | T | A | G | T | T | C | A | C | G | C |  |
| B.1.617.2 | C | G | A | A | G | C | G | G | G | T | - | - | - | - | - | - | T | A | T |  |
| B.1.617.2 | C | G | A | A | G | C | G | G | G | G | A | A | A | G | T | G | T | A | T | #373052 |
| AY.114 | T | T | C | G | T | T | T | A | T | G | A | A | A | G | T | G | C | G | C |  |
| mut |  |  |  |  |  |  |  |  |  |  |  |  |  |  |  |  | C>T | G>A | C>T |  |

|  |  |  |  |  |  |  |  |  |  |  |  |  |  |  |  |  |  |  |  |  |  |
| --- | --- | --- | --- | --- | --- | --- | --- | --- | --- | --- | --- | --- | --- | --- | --- | --- | --- | --- | --- | --- | --- |
| pos | 745 | 884 | 4720 | 6730 | 13027 | 13944 | 15906 | 16914 | 18905 | 19677 | 21893 | 22027 | 22029 | 22030 | 22031 | 22032 | 22033 | 22034 | 25471 | 26786 |  |
| ref | C | C | G | C | T | C | G | G | G | G | G | T | A | G | T | T | C | A | G | A |  |
| AY.125 | T | C | G | T | C | T | T | G | A | G | C | T | - | - | - | - | - | - | T | G |  |
| AY.125 | T | C | G | T | C | T | T | G | A | G | G | G | A | A | A | G | T | G | T | G | #376350 |
| AY.114 | C | T | T | C | T | C | G | T | G | T | G | G | A | A | A | G | T | G | G | A |  |
| mut |  |  |  |  |  |  |  |  |  |  |  |  |  |  |  |  |  |  | G>T | A>G |  |

|  |  |  |  |  |  |  |  |  |  |  |  |  |  |  |  |  |  |  |  |  |  |  |  |  |  |  |  |  |  |  |  |  |  |  |  |  |  |  |  |  |  |  |  |
| --- | --- | --- | --- | --- | --- | --- | --- | --- | --- | --- | --- | --- | --- | --- | --- | --- | --- | --- | --- | --- | --- | --- | --- | --- | --- | --- | --- | --- | --- | --- | --- | --- | --- | --- | --- | --- | --- | --- | --- | --- | --- | --- | --- |
| pos | 1191 | 1267 | 4181 | 4321 | 5184 | 6402 | 7124 | 8986 | 9053 | 9203 | 9678 | 10029 | 10399 | 10868 | 11005 | 11201 | 11332 | 15451 | 16466 | 17236 | 17496 | 18360 | 19220 | 20396 | 21792 | 21974 | 23202 | 24208 | 25572 | 25702 | 26501 | 27874 | 27948 | 28248 | 28249 | 28250 | 28251 | 28252 | 28253 | 28895 | 28916 | 29669 |  |
| ref | C | C | G | C | C | C | C | C | G | G | T | C | A | C | C | A | A | G | C | A | A | A | C | A | A | G | C | C | C | C | G | C | G | G | A | T | T | T | C | G | G | A |  |
| AY.48 | T | T | G | T | T | C | C | C | G | A | C | C | G | T | A | A | A | G | C | A | G | A | C | G | C | G | T | C | T | C | G | C | T | G | A | T | T | T | T | G | G | G |  |
| AY.103 | T | T | G | T | T | C | C | C | G | A | C | T | A | C | C | G | G | A | T | G | A | G | T | A | A | T | C | T | C | T | T | T | G | - | - | - | - | - | - | T | T | A | #376921 |
| AY.103 | C | C | T | C | C | T | T | T | T | G | T | T | A | C | C | G | G | A | T | G | A | G | T | A | A | G | C | T | C | T | G | T | G | - | - | - | - | - | - | T | T | A |  |
| mut |  |  |  |  |  |  |  |  |  |  |  |  |  |  |  |  |  |  |  |  |  |  |  |  |  | G>T |  |  |  |  | G>T |  |  |  |  |  |  |  |  |  |  |  |  |

|  |  |  |  |  |  |  |  |  |  |  |
| --- | --- | --- | --- | --- | --- | --- | --- | --- | --- | --- |
| pos | 355 | 4237 | 4891 | 7948 | 10642 | 12685 | 17040 | 24138 | 26822 |  |
| ref | C | T | C | C | G | G | T | C | C |  |
| AY.4 | T | C | C | T | T | G | C | C | C |  |
| AY.4 | T | C | C | T | T | T | T | T | T | #378870 |
| AY.4 | C | T | T | C | G | T | T | T | T |  |
| mut |  |  |  |  |  |  |  |  |  |  |

|  |  |  |  |  |  |  |  |  |  |  |  |  |  |  |  |  |  |  |  |  |  |  |  |  |  |  |  |  |  |  |  |  |  |  |  |  |  |  |  |  |  |  |  |  |  |  |  |  |  |  |  |  |  |  |  |  |  |  |  |  |  |  |  |  |  |  |  |  |  |  |  |  |  |  |  |  |  |  |  |  |  |  |  |  |  |  |  |  |  |  |  |  |  |  |  |  |  |
| --- | --- | --- | --- | --- | --- | --- | --- | --- | --- | --- | --- | --- | --- | --- | --- | --- | --- | --- | --- | --- | --- | --- | --- | --- | --- | --- | --- | --- | --- | --- | --- | --- | --- | --- | --- | --- | --- | --- | --- | --- | --- | --- | --- | --- | --- | --- | --- | --- | --- | --- | --- | --- | --- | --- | --- | --- | --- | --- | --- | --- | --- | --- | --- | --- | --- | --- | --- | --- | --- | --- | --- | --- | --- | --- | --- | --- | --- | --- | --- | --- | --- | --- | --- | --- | --- | --- | --- | --- | --- | --- | --- | --- | --- | --- | --- | --- | --- |
| pos | 210 | 337 | 733 | 2749 | 3644 | 3828 | 4181 | 5648 | 5812 | 6319 | 6402 | 6613 | 7124 | 8986 | 9053 | 10029 | 11201 | 11288 | 11289 | 11290 | 11291 | 11292 | 11293 | 11294 | 11295 | 11296 | 11332 | 11456 | 12778 | 13860 | 15194 | 15451 | 16466 | 17259 | 18115 | 19220 | 20199 | 21302 | 21304 | 21305 | 21614 | 21618 | 21621 | 21638 | 21987 | 22029 | 22030 | 22031 | 22032 | 22033 | 22034 | 22132 | 22812 | 22917 | 22995 | 23012 | 23063 | 23284 | 23525 | 23604 | 24410 | 24642 | 25088 | 25339 | 25469 | 26149 | 26767 | 27638 | 27752 | 27874 | 28167 | 28248 | 28249 | 28250 | 28251 | 28252 | 28253 | 28271 | 28461 | 28512 | 28877 | 28878 | 28881 | 28882 | 28883 | 28916 | 29050 | 29402 | 29509 | 29700 | 29742 | 29762 | 29769 | 29774 | 29779 | 29781 |  |
| ref | G | C | T | C | G | C | G | A | C | A | C | A | C | C | G | C | A | T | C | T | G | G | T | T | T | T | A | A | C | C | C | G | C | G | C | C | C | C | C | G | C | C | C | C | G | A | G | T | T | C | A | G | A | T | C | G | A | T | C | C | G | C | G | C | C | T | T | T | C | C | G | G | A | T | T | T | C | A | A | C | A | G | G | G | G | G | G | G | C | A | G | C | C | C | G | G |  |
| P.1 | G | C | C | T | G | T | G | C | C | G | C | G | C | C | G | C | A | - | - | - | - | - | - | - | - | - | A | A | T | T | C | G | C | T | C | C | C | C | C | G | T | C | A | T | G | A | G | T | T | C | A | T | C | T | C | A | T | T | T | C | G | T | T | C | C | C | T | T | C | C | A | G | A | T | T | T | C | A | A | G | T | C | A | A | C | G | G | G | C | A | G | C | C | C | G | G |  |
| AY.3.1 | G | T | C | T | G | T | G | C | T | A | T | A | T | T | T | T | G | T | C | T | G | G | T | T | T | T | G | G | C | C | T | A | T | G | T | T | T | T | A | A | C | G | C | C | A | - | - | - | - | - | - | G | A | G | A | G | A | C | C | G | A | C | G | T | T | T | C | C | T | T | G | - | - | - | - | - | - | - | G | C | A | G | T | G | G | T | A | T | T | A | T | C | C | C | G | G | #380339 |
| AY.3.1 | T | C | T | C | T | C | T | A | C | A | T | A | T | T | T | T | G | T | C | T | G | G | T | T | T | T | G | G | C | C | T | A | T | G | T | T | T | T | A | A | C | G | C | C | A | - | - | - | - | - | - | G | A | G | A | G | A | C | C | G | A | C | G | T | T | T | C | C | T | T | G | - | - | - | - | - | - | - | G | C | A | G | T | G | G | T | A | T | T | - | T | - | - | - | - | - |  |
| mut |  | C>T |  |  |  |  |  |  | C>T |  |  |  |  |  |  |  |  |  |  |  |  |  |  |  |  |  |  |  |  |  |  |  |  |  |  |  |  |  |  |  |  |  |  |  |  |  |  |  |  |  |  |  |  |  |  |  |  |  |  |  |  |  |  |  |  |  |  |  |  |  |  |  |  |  |  |  |  |  |  |  |  |  |  |  |  |  |  |  |  | ->A |  | ->C | ->C | ->C | ->G | ->G |  |

|  |  |  |  |  |  |  |  |  |  |  |  |  |  |  |  |  |  |  |  |  |  |  |  |  |  |  |  |  |  |  |  |  |  |  |  |  |  |
| --- | --- | --- | --- | --- | --- | --- | --- | --- | --- | --- | --- | --- | --- | --- | --- | --- | --- | --- | --- | --- | --- | --- | --- | --- | --- | --- | --- | --- | --- | --- | --- | --- | --- | --- | --- | --- | --- |
| pos | 1699 | 3948 | 4068 | 4181 | 5184 | 5584 | 6402 | 7124 | 7393 | 8986 | 9053 | 9891 | 10029 | 11201 | 11332 | 11418 | 11514 | 11562 | 13019 | 13812 | 15786 | 16731 | 19220 | 21622 | 22227 | 23575 | 25693 | 26062 | 26107 | 27112 | 27507 | 27516 | 27874 | 28789 | 28916 | 29738 |  |
| ref | A | A | C | G | C | A | C | C | G | C | G | C | C | A | A | T | C | G | C | G | T | T | C | C | C | C | C | G | G | G | A | G | C | C | G | C |  |
| AY.75 | A | A | C | G | T | G | C | C | T | C | G | T | C | A | A | C | T | G | T | G | T | T | C | T | T | T | C | T | G | A | A | A | C | C | G | A |  |
| AY.75 | A | A | C | G | T | G | C | C | T | C | G | T | C | A | A | C | T | G | T | T | T | T | C | T | T | T | C | T | G | A | C | G | T | T | T | C | #383230 |
| AY.25.1 | G | G | T | T | C | A | T | T | G | T | T | C | T | G | G | T | C | T | C | G | C | C | T | C | C | C | A | G | C | G | C | G | T | T | T | C |  |
| mut |  |  |  |  |  |  |  |  |  |  |  |  |  |  |  |  |  |  |  | G>T |  |  |  |  |  |  |  |  |  |  |  |  |  |  |  |  |  |

|  |  |  |  |  |  |  |  |  |  |  |  |  |  |  |  |  |
| --- | --- | --- | --- | --- | --- | --- | --- | --- | --- | --- | --- | --- | --- | --- | --- | --- |
| pos | 2336 | 5629 | 6145 | 7851 | 7875 | 11963 | 13210 | 17040 | 17331 | 18115 | 18167 | 20269 | 21846 | 24630 | 29420 |  |
| ref | G | G | C | C | C | T | T | T | G | C | C | G | C | A | C |  |
| AY.6 | A | T | T | C | T | C | C | T | T | C | T | T | C | A | T |  |
| AY.6 | A | T | T | C | T | C | C | T | T | T | C | G | T | C | C | #389317 |
| AY.4 | G | G | C | T | C | T | T | C | G | T | C | G | T | A | C |  |
| mut |  |  |  |  |  |  |  |  |  |  |  |  |  | A>C |  |  |

|  |  |  |  |  |  |  |  |  |  |  |  |  |  |  |  |  |  |  |  |  |
| --- | --- | --- | --- | --- | --- | --- | --- | --- | --- | --- | --- | --- | --- | --- | --- | --- | --- | --- | --- | --- |
| pos | 1911 | 5221 | 7851 | 10262 | 12083 | 13424 | 14919 | 19006 | 20814 | 21987 | 22026 | 22027 | 22029 | 22030 | 22031 | 22032 | 22033 | 22034 | 25183 |  |
| ref | C | C | C | G | A | C | C | G | A | G | G | T | A | G | T | T | C | A | G |  |
| AY.119 | T | T | C | T | G | C | C | T | G | G | G | T | - | - | - | - | - | - | T |  |
| AY.119 | T | T | C | T | A | C | C | T | G | G | T | G | A | A | A | G | T | G | T | #391756 |
| AY.4 | C | C | T | G | A | T | T | G | A | A | T | G | A | A | A | G | T | G | G |  |
| mut |  |  |  |  | G>A |  |  |  |  |  |  |  |  |  |  |  |  |  | G>T |  |

|  |  |  |  |  |  |  |  |  |  |  |  |  |  |  |  |  |  |  |  |  |
| --- | --- | --- | --- | --- | --- | --- | --- | --- | --- | --- | --- | --- | --- | --- | --- | --- | --- | --- | --- | --- |
| pos | 884 | 4720 | 7851 | 8179 | 11599 | 13638 | 16914 | 17302 | 19677 | 21638 | 22027 | 22029 | 22030 | 22031 | 22032 | 22033 | 22034 | 28077 | 29366 |  |
| ref | C | G | C | G | T | T | G | G | G | C | T | A | G | T | T | C | A | G | C |  |
| AY.4.3 | C | G | T | T | C | C | G | T | G | T | T | - | - | - | - | - | - | C | T |  |
| AY.4.3 | C | G | T | T | C | C | G | T | G | T | G | A | A | A | G | T | G | C | T | #392422 |
| AY.114 | T | T | C | G | T | T | T | G | T | C | G | A | A | A | G | T | G | G | C |  |
| mut |  |  |  |  |  |  |  |  |  |  |  |  |  |  |  |  |  | G>C | C>T |  |

|  |  |  |  |  |  |  |  |  |  |  |  |  |  |  |  |  |  |  |  |  |  |  |  |  |  |  |  |  |  |  |
| --- | --- | --- | --- | --- | --- | --- | --- | --- | --- | --- | --- | --- | --- | --- | --- | --- | --- | --- | --- | --- | --- | --- | --- | --- | --- | --- | --- | --- | --- | --- |
| pos | 1045 | 1820 | 4181 | 5184 | 5584 | 6402 | 7124 | 8986 | 9053 | 9891 | 10029 | 11201 | 11332 | 11418 | 11514 | 11743 | 13019 | 19220 | 20085 | 21372 | 21846 | 21987 | 22227 | 27604 | 27874 | 28093 | 28916 | 29440 | 29751 |  |
| ref | A | G | G | C | A | C | C | C | G | C | C | A | A | T | C | G | C | C | A | G | C | G | C | G | C | C | G | G | G |  |
| AY.39 | A | G | T | C | A | T | T | T | T | C | T | G | G | T | C | G | C | T | A | T | T | G | C | A | T | C | T | G | G |  |
| B.1.617.2 | A | G | T | C | A | T | T | T | T | C | T | G | G | T | C | T | C | T | A | T | T | A | C | G | C | T | G | A | C | #394632 |
| AY.62 | T | A | G | T | G | C | C | C | G | T | C | A | A | C | T | G | T | C | G | G | C | A | T | G | C | T | G | A | C |  |
| mut |  |  |  |  |  |  |  |  |  |  |  |  |  |  |  | G>T |  |  |  |  |  | G>A |  |  |  |  |  |  |  |  |

|  |  |  |  |  |  |  |  |  |  |  |  |  |  |  |  |  |  |  |  |
| --- | --- | --- | --- | --- | --- | --- | --- | --- | --- | --- | --- | --- | --- | --- | --- | --- | --- | --- | --- |
| pos | 884 | 2706 | 3047 | 4720 | 7851 | 13665 | 16914 | 17040 | 19677 | 21303 | 21516 | 22027 | 22029 | 22030 | 22031 | 22032 | 22033 | 22034 |  |
| ref | C | C | G | G | C | C | G | T | G | A | C | T | A | G | T | T | C | A |  |
| AY.4 | C | T | T | G | T | T | G | C | G | G | T | T | - | - | - | - | - | - |  |
| AY.4 | C | T | T | G | T | T | G | C | G | G | T | G | A | A | A | G | T | G | #395805 |
| AY.114 | T | C | G | T | C | C | T | T | T | A | C | G | A | A | A | G | T | G |  |
| mut |  |  |  |  |  |  |  |  |  |  |  |  |  |  |  |  |  |  |  |

|  |  |  |  |  |  |  |  |  |  |  |  |  |  |  |  |  |  |  |  |  |  |  |  |  |  |  |  |  |  |  |  |  |  |
| --- | --- | --- | --- | --- | --- | --- | --- | --- | --- | --- | --- | --- | --- | --- | --- | --- | --- | --- | --- | --- | --- | --- | --- | --- | --- | --- | --- | --- | --- | --- | --- | --- | --- |
| pos | 1729 | 4181 | 5184 | 5584 | 6402 | 6943 | 7124 | 7851 | 8986 | 9053 | 9891 | 10029 | 10318 | 11201 | 11332 | 11418 | 11514 | 12400 | 13019 | 14030 | 15368 | 17574 | 18086 | 18340 | 19220 | 21846 | 22227 | 22792 | 26019 | 27874 | 28706 | 28916 |  |
| ref | G | G | C | A | C | G | C | C | C | G | C | C | G | A | A | T | C | C | C | G | C | T | C | G | C | C | C | C | A | C | C | G |  |
| AY.4 | G | T | C | A | T | T | T | T | T | T | C | T | T | G | G | T | C | C | C | G | T | T | C | A | T | T | C | C | A | T | C | T |  |
| AY.9.2 | G | T | C | A | T | T | C | C | C | G | T | C | G | A | A | C | T | T | T | A | C | C | T | G | C | C | T | T | G | C | T | G | #396951 |
| AY.9.2 | A | G | T | G | C | G | C | C | C | G | T | C | G | A | A | C | T | C | T | A | C | C | T | G | C | C | T | T | A | C | C | G |  |
| mut |  |  |  |  |  |  |  |  |  |  |  |  |  |  |  |  |  | C>T |  |  |  |  |  |  |  |  |  |  | A>G |  | C>T |  |  |

|  |  |  |  |  |  |  |  |  |  |  |  |  |  |  |  |  |  |  |
| --- | --- | --- | --- | --- | --- | --- | --- | --- | --- | --- | --- | --- | --- | --- | --- | --- | --- | --- |
| pos | 8139 | 13482 | 14246 | 17615 | 18457 | 19725 | 20762 | 21987 | 22026 | 22027 | 22029 | 22030 | 22031 | 22032 | 22033 | 22034 | 27670 |  |
| ref | C | A | A | A | C | T | C | G | G | T | A | G | T | T | C | A | G |  |
| AY.39 | C | A | G | A | T | A | C | G | G | T | - | - | - | - | - | - | T |  |
| AY.39 | C | A | G | A | T | A | C | G | T | G | A | A | A | G | T | G | T | #398296 |
| AY.39.1 | T | G | A | G | C | T | T | A | T | G | A | A | A | G | T | G | G |  |
| mut |  |  |  |  |  |  |  |  |  |  |  |  |  |  |  |  | G>T |  |

|  |  |  |  |  |  |  |  |  |  |  |  |  |  |  |  |  |  |  |
| --- | --- | --- | --- | --- | --- | --- | --- | --- | --- | --- | --- | --- | --- | --- | --- | --- | --- | --- |
| pos | 484 | 884 | 4158 | 4720 | 5907 | 7851 | 9969 | 16914 | 19677 | 21987 | 22027 | 22029 | 22030 | 22031 | 22032 | 22033 | 22034 |  |
| ref | T | C | C | G | C | C | C | G | G | G | T | A | G | T | T | C | A |  |
| AY.4 | C | C | T | G | T | T | T | G | G | G | T | - | - | - | - | - | - |  |
| AY.4 | C | C | T | G | T | T | T | G | G | G | G | A | A | A | G | T | G | #398986 |
| AY.114 | T | T | C | T | C | C | C | T | T | A | G | A | A | A | G | T | G |  |
| mut |  |  |  |  |  |  |  |  |  |  |  |  |  |  |  |  |  |  |

|  |  |  |  |  |  |  |  |  |  |  |  |  |  |  |
| --- | --- | --- | --- | --- | --- | --- | --- | --- | --- | --- | --- | --- | --- | --- |
| pos | 526 | 7851 | 15138 | 15277 | 17040 | 17427 | 19524 | 21846 | 27875 | 27881 | 27882 | 27883 | 28639 |  |
| ref | G | C | G | C | T | G | C | C | T | C | G | C | C |  |
| AY.4 | G | T | G | T | C | T | C | T | T | C | G | C | C |  |
| AY.4 | G | T | T | T | C | T | C | T | C | T | C | T | T | #400258 |
| AY.98 | T | C | G | C | T | G | T | C | C | T | C | T | C |  |
| mut |  |  | G>T |  |  |  |  |  |  |  |  |  | C>T |  |

|  |  |  |  |  |  |  |  |  |  |  |  |  |  |  |  |  |  |  |  |  |  |  |  |  |  |  |  |  |  |  |  |
| --- | --- | --- | --- | --- | --- | --- | --- | --- | --- | --- | --- | --- | --- | --- | --- | --- | --- | --- | --- | --- | --- | --- | --- | --- | --- | --- | --- | --- | --- | --- | --- |
| pos | 1191 | 1267 | 1877 | 2564 | 3948 | 4181 | 5184 | 6402 | 7124 | 8986 | 9053 | 9267 | 9891 | 10029 | 11201 | 11332 | 11418 | 11562 | 12501 | 12946 | 18176 | 19160 | 19220 | 20262 | 20718 | 26107 | 27507 | 27739 | 27874 | 28916 |  |
| ref | C | C | T | G | A | G | C | C | C | C | G | G | C | C | A | A | T | G | C | T | C | C | C | A | G | G | A | C | C | G |  |
| AY.14 | T | T | G | G | A | G | T | C | C | C | G | G | T | C | A | A | C | G | C | C | T | T | C | G | T | G | A | T | C | G |  |
| AY.25 | T | T | G | G | A | G | T | C | C | T | T | T | C | T | G | G | T | T | T | T | C | C | T | A | G | C | C | C | T | T | #405949 |
| AY.25.1 | C | C | T | A | G | T | C | T | T | T | T | G | C | T | G | G | T | T | T | T | C | C | T | A | G | C | C | C | T | T |  |
| mut |  |  |  |  |  |  |  |  |  |  |  | G>T |  |  |  |  |  |  |  |  |  |  |  |  |  |  |  |  |  |  |  |

|  |  |  |  |  |  |  |  |  |  |  |  |  |  |  |  |  |  |  |
| --- | --- | --- | --- | --- | --- | --- | --- | --- | --- | --- | --- | --- | --- | --- | --- | --- | --- | --- |
| pos | 3948 | 6408 | 7600 | 11095 | 11562 | 11565 | 12964 | 21846 | 21987 | 25003 | 25452 | 26107 | 27507 | 27572 | 27573 | 27575 | 27576 |  |
| ref | A | C | C | C | G | C | A | C | G | A | C | G | A | G | C | C | T |  |
| AY.25.1 | G | C | C | C | T | T | A | C | G | A | C | C | C | G | C | C | T |  |
| AY.25.1 | G | C | C | C | T | T | G | C | G | A | C | C | C | A | T | A | C | #408173 |
| AY.120.1 | A | T | T | T | G | C | A | T | A | G | T | G | A | A | T | A | C |  |
| mut |  |  |  |  |  |  | A>G |  |  |  |  |  |  |  |  |  |  |  |

|  |  |  |  |  |  |  |  |  |  |  |  |  |  |
| --- | --- | --- | --- | --- | --- | --- | --- | --- | --- | --- | --- | --- | --- |
| pos | 1559 | 1741 | 3896 | 6408 | 7851 | 12223 | 18264 | 25003 | 26151 | 26188 | 27147 | 28378 |  |
| ref | G | A | G | C | C | G | C | A | C | G | G | G |  |
| AY.120 | A | A | G | T | C | G | T | G | T | G | T | T |  |
| AY.120 | A | A | G | T | C | G | T | A | C | A | G | G | #410200 |
| AY.4 | G | G | A | C | T | A | C | A | C | A | G | G |  |
| mut |  |  |  |  |  |  |  |  |  |  |  |  |  |

|  |  |  |  |  |  |  |  |  |  |  |  |  |  |  |  |  |  |  |  |  |  |  |  |  |  |  |  |  |
| --- | --- | --- | --- | --- | --- | --- | --- | --- | --- | --- | --- | --- | --- | --- | --- | --- | --- | --- | --- | --- | --- | --- | --- | --- | --- | --- | --- | --- |
| pos | 686 | 687 | 688 | 689 | 690 | 691 | 692 | 693 | 694 | 884 | 4720 | 7851 | 9967 | 15277 | 16914 | 17040 | 17427 | 19677 | 20677 | 21057 | 22027 | 22029 | 22030 | 22031 | 22032 | 22033 | 22034 |  |
| ref | A | A | G | T | C | A | T | T | T | C | G | C | C | C | G | T | G | G | C | C | T | A | G | T | T | C | A |  |
| AY.4 | - | - | - | - | - | - | - | - | - | C | G | T | T | T | G | C | T | G | T | T | T | - | - | - | - | - | - |  |
| AY.4 | - | - | - | - | - | - | - | - | - | C | G | T | T | T | G | C | T | G | T | T | G | A | A | A | G | T | G | #410707 |
| AY.114 | A | A | G | T | C | A | T | T | T | T | T | C | C | C | T | T | G | T | C | C | G | A | A | A | G | T | G |  |
| mut |  |  |  |  |  |  |  |  |  |  |  |  |  |  |  |  |  |  |  |  |  |  |  |  |  |  |  |  |

|  |  |  |  |  |  |  |  |  |  |  |  |  |  |  |  |  |
| --- | --- | --- | --- | --- | --- | --- | --- | --- | --- | --- | --- | --- | --- | --- | --- | --- |
| pos | 745 | 2258 | 4012 | 6730 | 10525 | 12073 | 15237 | 15598 | 15906 | 18905 | 21161 | 23718 | 25471 | 25872 | 26786 |  |
| ref | C | G | C | C | C | C | C | G | G | G | C | C | G | C | A |  |
| AY.42 | C | A | C | C | T | T | T | G | G | G | C | T | G | T | A |  |
| AY.42 | C | A | T | C | T | T | T | T | G | A | T | C | T | C | G | #410724 |
| AY.125 | T | G | C | T | C | C | C | G | T | A | C | C | T | C | G |  |
| mut |  |  | C>T |  |  |  |  | G>T |  |  | C>T |  |  |  |  |  |

|  |  |  |  |  |  |  |  |  |  |  |  |  |  |  |  |  |  |  |  |  |  |
| --- | --- | --- | --- | --- | --- | --- | --- | --- | --- | --- | --- | --- | --- | --- | --- | --- | --- | --- | --- | --- | --- |
| pos | 884 | 1738 | 4720 | 4794 | 10691 | 13372 | 13712 | 14925 | 16914 | 19677 | 19983 | 21987 | 22027 | 22029 | 22030 | 22031 | 22032 | 22033 | 22034 | 27298 |  |
| ref | C | G | G | C | A | C | A | C | G | G | C | G | T | A | G | T | T | C | A | A |  |
| AY.116 | C | T | G | T | G | T | G | T | G | G | T | G | T | - | - | - | - | - | - | G |  |
| AY.116 | C | T | G | T | G | T | G | T | G | G | T | G | G | A | A | A | G | T | G | G | #411301 |
| AY.114 | T | G | T | C | A | C | A | C | T | T | C | A | G | A | A | A | G | T | G | A |  |
| mut |  |  |  |  |  |  |  |  |  |  |  |  |  |  |  |  |  |  |  | A>G |  |

|  |  |  |  |  |  |  |  |  |  |  |  |  |  |  |  |
| --- | --- | --- | --- | --- | --- | --- | --- | --- | --- | --- | --- | --- | --- | --- | --- |
| pos | 507 | 508 | 509 | 18647 | 22027 | 22029 | 22030 | 22031 | 22032 | 22033 | 22034 | 22296 | 22550 | 23179 |  |
| ref | A | T | G | C | T | A | G | T | T | C | A | A | C | C |  |
| AY.107 | T | C | A | T | T | - | - | - | - | - | - | A | T | C |  |
| AY.107 | T | C | A | T | G | A | A | A | G | T | G | C | C | T | #411345 |
| AY.107 | A | T | G | C | G | A | A | A | G | T | G | A | C | C |  |
| mut |  |  |  |  |  |  |  |  |  |  |  | A>C |  | C>T |  |

|  |  |  |  |  |  |  |  |  |  |  |  |  |  |  |  |  |  |
| --- | --- | --- | --- | --- | --- | --- | --- | --- | --- | --- | --- | --- | --- | --- | --- | --- | --- |
| pos | 1048 | 1441 | 4002 | 4455 | 5583 | 7851 | 13501 | 21846 | 22051 | 23425 | 25494 | 26065 | 26885 | 27526 | 27527 | 27737 |  |
| ref | G | C | C | C | C | C | C | C | G | T | G | G | C | C | C | C |  |
| AY.122 | T | T | C | T | T | C | T | C | G | T | T | T | C | T | T | C |  |
| AY.122 | T | T | C | T | T | C | T | T | C | C | G | G | T | C | C | T | #413359 |
| AY.4 | G | C | T | C | C | T | C | T | C | C | G | G | T | C | C | T |  |
| mut |  |  |  |  |  |  |  |  |  |  |  |  |  |  |  |  |  |

|  |  |  |  |  |  |  |  |  |  |  |  |  |
| --- | --- | --- | --- | --- | --- | --- | --- | --- | --- | --- | --- | --- |
| pos | 3523 | 5668 | 6419 | 17913 | 22027 | 22029 | 22030 | 22031 | 22032 | 22033 | 22034 |  |
| ref | A | G | G | C | T | A | G | T | T | C | A |  |
| AY.117 | C | A | A | T | T | - | - | - | - | - | - |  |
| AY.117 | C | A | A | T | T | A | A | A | G | T | G | #413874 |
| AY.117 | A | G | G | C | G | A | A | A | G | T | G |  |
| mut |  |  |  |  |  |  |  |  |  |  |  |  |

|  |  |  |  |  |  |  |  |  |  |  |  |  |  |  |  |  |  |  |  |  |  |  |  |  |  |  |  |  |  |  |  |  |  |  |  |  |  |  |  |  |  |  |  |  |  |  |  |  |  |  |  |  |  |  |  |  |  |  |  |  |  |  |  |  |  |  |  |  |  |  |  |  |  |  |  |  |  |  |  |  |  |  |  |  |  |  |  |  |
| --- | --- | --- | --- | --- | --- | --- | --- | --- | --- | --- | --- | --- | --- | --- | --- | --- | --- | --- | --- | --- | --- | --- | --- | --- | --- | --- | --- | --- | --- | --- | --- | --- | --- | --- | --- | --- | --- | --- | --- | --- | --- | --- | --- | --- | --- | --- | --- | --- | --- | --- | --- | --- | --- | --- | --- | --- | --- | --- | --- | --- | --- | --- | --- | --- | --- | --- | --- | --- | --- | --- | --- | --- | --- | --- | --- | --- | --- | --- | --- | --- | --- | --- | --- | --- | --- | --- | --- | --- |
| pos | 210 | 913 | 3177 | 3267 | 4181 | 5239 | 5388 | 5514 | 5986 | 6402 | 6954 | 7124 | 8986 | 9053 | 9211 | 10029 | 11201 | 11288 | 11289 | 11290 | 11291 | 11292 | 11293 | 11294 | 11295 | 11296 | 11332 | 14676 | 15279 | 15451 | 16176 | 16466 | 19220 | 21618 | 21765 | 21766 | 21767 | 21768 | 21769 | 21770 | 21846 | 21987 | 21991 | 21992 | 21993 | 22029 | 22030 | 22031 | 22032 | 22033 | 22034 | 22081 | 22917 | 22995 | 23063 | 23271 | 23604 | 23709 | 24410 | 24506 | 24914 | 25469 | 26767 | 27390 | 27638 | 27752 | 27874 | 27972 | 28048 | 28111 | 28248 | 28249 | 28250 | 28251 | 28252 | 28253 | 28280 | 28281 | 28282 | 28461 | 28881 | 28882 | 28883 | 28916 | 28977 | 29402 | 29742 |  |
| ref | G | C | C | C | G | C | C | T | C | C | T | C | C | G | G | C | A | T | C | T | G | G | T | T | T | T | A | C | C | G | T | C | C | C | T | A | C | A | T | G | C | G | T | T | A | A | G | T | T | C | A | G | T | C | A | C | C | C | G | T | G | C | T | G | T | C | C | C | G | A | G | A | T | T | T | C | G | A | T | A | G | G | G | G | C | G | G |  |
| AY.29 | T | C | C | C | T | T | C | C | C | T | T | T | T | T | G | T | G | T | C | T | G | G | T | T | T | T | G | C | C | A | T | T | T | G | T | A | C | A | T | G | T | G | T | T | A | - | - | - | - | - | - | G | G | A | A | C | G | C | A | T | G | T | C | G | C | T | T | C | G | A | - | - | - | - | - | - | G | A | T | G | T | G | G | T | C | T | T |  |
| XC | T | C | C | C | T | T | C | C | C | T | T | T | T | T | G | T | G | T | C | T | G | G | T | T | T | T | G | C | C | A | T | T | T | G | T | A | C | A | T | G | T | A | T | T | A | - | - | - | - | - | - | T | G | A | A | C | G | C | A | T | G | T | C | T | T | C | C | T | T | G | G | A | T | T | T | C | C | T | A | A | A | A | C | G | T | G | G | #414488 |
| B.1.1.7 | G | T | T | T | G | C | A | T | T | C | C | C | C | G | A | C | A | - | - | - | - | - | - | - | - | - | A | T | T | G | C | C | C | C | - | - | - | - | - | - | C | G | - | - | - | A | G | T | T | C | A | G | T | C | T | A | A | T | G | G | C | C | T | T | T | C | C | T | T | G | G | A | T | T | T | C | C | T | A | A | A | A | C | G | T | G | G |  |
| mut |  |  |  |  |  |  |  |  |  |  |  |  |  |  |  |  |  |  |  |  |  |  |  |  |  |  |  |  |  |  |  |  |  |  |  |  |  |  |  |  |  | G>A |  |  |  |  |  |  |  |  |  | G>T |  |  |  |  |  |  |  |  |  |  |  |  |  |  |  |  |  |  |  |  |  |  |  |  |  |  |  |  |  |  |  |  |  |  |  |  |

|  |  |  |  |  |  |  |  |  |  |  |  |  |  |  |  |  |  |  |  |  |  |  |  |  |  |  |  |  |  |  |  |  |  |  |  |
| --- | --- | --- | --- | --- | --- | --- | --- | --- | --- | --- | --- | --- | --- | --- | --- | --- | --- | --- | --- | --- | --- | --- | --- | --- | --- | --- | --- | --- | --- | --- | --- | --- | --- | --- | --- |
| pos | 2061 | 2651 | 2974 | 4181 | 5184 | 5584 | 5706 | 6402 | 7124 | 8986 | 9053 | 9891 | 10029 | 10525 | 11201 | 11332 | 11418 | 11514 | 13019 | 14487 | 15009 | 19220 | 19846 | 21846 | 21987 | 22227 | 23758 | 23896 | 24872 | 25352 | 26079 | 27345 | 27874 | 28916 |  |
| ref | C | A | T | G | C | A | A | C | C | C | G | C | C | C | A | A | T | C | C | A | T | C | G | C | G | C | C | C | G | G | C | A | C | G |  |
| AY.26 | C | A | T | G | T | G | A | C | C | C | G | T | C | C | A | A | C | T | T | G | C | C | A | C | A | T | C | C | G | T | T | T | C | G |  |
| B.1.617.2 | C | A | T | G | T | G | A | C | C | C | G | T | C | C | A | A | C | T | T | G | C | C | A | C | A | T | T | T | T | G | C | A | T | T | #415855 |
| AY.20 | T | G | C | T | C | A | G | T | T | T | T | C | T | T | G | G | T | C | C | A | T | T | G | T | G | C | T | T | T | G | C | A | T | T |  |
| mut |  |  |  |  |  |  |  |  |  |  |  |  |  |  |  |  |  |  |  |  |  |  |  |  |  |  |  |  |  |  |  |  |  |  |  |

|  |  |  |  |  |  |  |  |  |  |  |  |  |  |  |  |  |  |  |  |  |  |  |  |  |  |  |  |  |  |  |  |  |  |  |  |  |
| --- | --- | --- | --- | --- | --- | --- | --- | --- | --- | --- | --- | --- | --- | --- | --- | --- | --- | --- | --- | --- | --- | --- | --- | --- | --- | --- | --- | --- | --- | --- | --- | --- | --- | --- | --- | --- |
| pos | 376 | 1191 | 1267 | 1473 | 1877 | 4181 | 5184 | 5866 | 6402 | 7124 | 8986 | 9053 | 9559 | 9733 | 9891 | 10029 | 10870 | 11201 | 11332 | 11418 | 12946 | 15240 | 17122 | 17790 | 18176 | 19160 | 19220 | 20262 | 20718 | 21846 | 24130 | 27739 | 27874 | 28007 | 28916 |  |
| ref | G | C | C | C | T | G | C | C | C | C | C | G | C | C | C | C | G | A | A | T | T | C | G | G | C | C | C | A | G | C | C | C | C | T | G |  |
| AY.14 | T | T | T | C | G | G | T | T | C | C | C | G | C | C | T | C | T | A | A | C | C | C | G | T | T | T | C | G | T | C | C | T | C | T | G |  |
| AY.100 | T | T | T | C | G | T | C | C | T | T | T | T | T | T | C | T | G | G | G | T | T | T | T | G | C | C | T | A | G | T | T | C | G | C | T | #417645 |
| AY.100 | G | C | C | T | T | T | C | C | T | T | T | T | T | C | C | T | G | G | G | T | T | T | T | G | C | C | T | A | G | T | T | C | G | C | T |  |
| mut |  |  |  |  |  |  |  |  |  |  |  |  |  | C>T |  |  |  |  |  |  |  |  |  |  |  |  |  |  |  |  |  |  |  |  |  |  |

|  |  |  |  |  |  |  |  |  |  |  |  |  |  |
| --- | --- | --- | --- | --- | --- | --- | --- | --- | --- | --- | --- | --- | --- |
| pos | 6638 | 6884 | 7702 | 14559 | 15842 | 17041 | 27656 | 28084 | 28106 | 28868 | 29254 | 29734 |  |
| ref | C | G | A | G | C | T | T | A | T | C | G | G |  |
| AY.4 | T | A | G | T | C | T | C | A | T | T | T | C |  |
| AY.4 | T | A | G | T | C | T | T | C | G | C | G | G | #419161 |
| AY.4 | C | G | A | G | T | C | T | C | G | C | G | G |  |
| mut |  |  |  |  |  |  |  |  |  |  |  |  |  |

|  |  |  |  |  |  |  |  |  |  |  |  |  |  |  |  |  |  |  |  |  |  |  |  |  |  |  |  |  |
| --- | --- | --- | --- | --- | --- | --- | --- | --- | --- | --- | --- | --- | --- | --- | --- | --- | --- | --- | --- | --- | --- | --- | --- | --- | --- | --- | --- | --- |
| pos | 4181 | 5184 | 5526 | 5584 | 6402 | 7124 | 8829 | 8986 | 9053 | 9891 | 10029 | 11201 | 11332 | 11418 | 11514 | 13019 | 13667 | 18804 | 19006 | 19220 | 21846 | 21987 | 22227 | 25352 | 27345 | 27874 | 28916 |  |
| ref | G | C | C | A | C | C | C | C | G | C | C | A | A | T | C | C | C | C | G | C | C | G | C | G | A | C | G |  |
| AY.119 | T | C | C | A | T | T | T | T | T | C | T | G | G | T | C | C | C | C | T | T | T | G | C | G | A | T | T |  |
| B.1.617.2 | T | C | C | A | T | T | T | T | T | C | T | A | A | C | T | T | T | T | G | C | C | A | T | T | T | C | G | #421696 |
| AY.26 | G | T | T | G | C | C | C | C | G | T | C | A | A | C | T | T | C | T | G | C | C | A | T | T | T | C | G |  |
| mut |  |  |  |  |  |  |  |  |  |  |  |  |  |  |  |  | C>T |  |  |  |  |  |  |  |  |  |  |  |

|  |  |  |  |  |  |  |  |  |  |  |  |  |  |  |  |  |  |  |  |
| --- | --- | --- | --- | --- | --- | --- | --- | --- | --- | --- | --- | --- | --- | --- | --- | --- | --- | --- | --- |
| pos | 2564 | 3948 | 8112 | 11456 | 11562 | 20296 | 22783 | 25710 | 26107 | 27507 | 29050 | 29509 | 29700 | 29762 | 29769 | 29774 | 29779 | 29781 |  |
| ref | G | A | C | A | G | A | C | C | G | A | G | C | A | C | C | C | G | G |  |
| AY.25.1 | A | G | C | A | T | A | C | C | C | C | G | C | A | C | C | C | G | G |  |
| AY.3 | A | G | C | A | T | G | T | T | G | A | A | T | A | C | C | C | G | G | #421731 |
| AY.3 | G | A | T | G | G | A | T | T | G | A | A | T | - | - | - | - | - | - |  |
| mut |  |  |  |  |  | A>G |  |  |  |  |  |  | ->A | ->C | ->C | ->C | ->G | ->G |  |

|  |  |  |  |  |  |  |  |  |  |  |  |  |  |  |  |  |  |  |
| --- | --- | --- | --- | --- | --- | --- | --- | --- | --- | --- | --- | --- | --- | --- | --- | --- | --- | --- |
| pos | 884 | 2156 | 3261 | 4720 | 7851 | 16914 | 17040 | 17339 | 19542 | 19677 | 22027 | 22029 | 22030 | 22031 | 22032 | 22033 | 22034 |  |
| ref | C | C | C | G | C | G | T | C | G | G | T | A | G | T | T | C | A |  |
| AY.4 | C | T | T | G | T | G | C | T | T | G | T | - | - | - | - | - | - |  |
| AY.4 | C | T | T | G | T | G | C | T | T | G | G | A | A | A | G | T | G | #423969 |
| AY.114 | T | C | C | T | C | T | T | C | G | T | G | A | A | A | G | T | G |  |
| mut |  |  |  |  |  |  |  |  |  |  |  |  |  |  |  |  |  |  |

|  |  |  |  |  |  |  |  |  |  |  |  |  |  |
| --- | --- | --- | --- | --- | --- | --- | --- | --- | --- | --- | --- | --- | --- |
| pos | 1059 | 4321 | 10271 | 16800 | 20505 | 22027 | 22029 | 22030 | 22031 | 22032 | 22033 | 22034 |  |
| ref | C | C | G | T | T | T | A | G | T | T | C | A |  |
| AY.118 | T | C | G | C | C | T | - | - | - | - | - | - |  |
| AY.118 | T | C | G | C | C | G | A | A | A | G | T | G | #424743 |
| AY.118 | C | T | A | T | T | G | A | A | A | G | T | G |  |
| mut |  |  |  |  |  |  |  |  |  |  |  |  |  |

|  |  |  |  |  |  |  |  |  |  |  |  |  |  |  |
| --- | --- | --- | --- | --- | --- | --- | --- | --- | --- | --- | --- | --- | --- | --- |
| pos | 799 | 2062 | 8318 | 8597 | 14318 | 22026 | 22027 | 22029 | 22030 | 22031 | 22032 | 22033 | 22034 |  |
| ref | C | C | C | C | C | G | T | A | G | T | T | C | A |  |
| AY.100 | T | T | A | A | T | G | T | - | - | - | - | - | - |  |
| AY.100 | T | T | A | A | T | G | G | A | A | A | G | T | G | #427780 |
| AY.100 | C | C | C | C | C | T | G | A | A | A | G | T | G |  |
| mut |  |  |  |  |  |  |  |  |  |  |  |  |  |  |

|  |  |  |  |  |  |  |  |  |  |  |  |  |  |
| --- | --- | --- | --- | --- | --- | --- | --- | --- | --- | --- | --- | --- | --- |
| pos | 3261 | 8752 | 17236 | 18486 | 20451 | 21897 | 24208 | 26107 | 26759 | 27384 | 27507 | 28378 |  |
| ref | C | C | A | C | C | C | C | G | A | T | A | G |  |
| AY.103 | T | C | G | T | C | C | T | G | A | T | A | G |  |
| AY.25 | T | C | G | T | T | T | T | C | G | C | C | T | #427863 |
| AY.25 | C | T | A | C | T | T | C | C | G | T | C | G |  |
| mut |  |  |  |  |  |  | C>T |  |  | T>C |  | G>T |  |

|  |  |  |  |  |  |  |  |  |  |  |  |  |  |  |  |  |  |  |  |  |  |  |  |  |  |  |  |  |  |  |  |  |  |  |  |
| --- | --- | --- | --- | --- | --- | --- | --- | --- | --- | --- | --- | --- | --- | --- | --- | --- | --- | --- | --- | --- | --- | --- | --- | --- | --- | --- | --- | --- | --- | --- | --- | --- | --- | --- | --- |
| pos | 686 | 687 | 688 | 689 | 690 | 691 | 692 | 693 | 694 | 4181 | 5184 | 5584 | 6402 | 7124 | 8986 | 9053 | 9891 | 10029 | 11201 | 11332 | 11418 | 11514 | 13019 | 15952 | 18744 | 19220 | 22227 | 22792 | 26262 | 27676 | 27874 | 28299 | 28916 | 29520 |  |
| ref | A | A | G | T | C | A | T | T | T | G | C | A | C | C | C | G | C | C | A | A | T | C | C | C | C | C | C | C | G | G | C | A | G | C |  |
| AY.43 | A | A | G | T | C | A | T | T | T | T | C | A | T | T | T | T | C | T | G | G | T | C | C | A | T | T | C | C | A | G | T | T | T | C |  |
| B.1.617.2 | - | - | - | - | - | - | - | - | - | T | C | A | T | T | T | T | C | T | G | G | T | C | C | A | C | T | T | T | G | T | C | A | G | T | #429553 |
| B.1.617.2 | A | A | G | T | C | A | T | T | T | G | T | G | C | C | C | G | T | C | A | A | C | T | T | C | C | C | T | C | G | T | C | A | G | C |  |
| mut | A>- | A>- | G>- | T>- | C>- | A>- | T>- | T>- | T>- |  |  |  |  |  |  |  |  |  |  |  |  |  |  |  | T>C |  |  | C>T |  |  |  |  |  | C>T |  |

|  |  |  |  |  |  |  |  |  |  |  |  |  |  |
| --- | --- | --- | --- | --- | --- | --- | --- | --- | --- | --- | --- | --- | --- |
| pos | 219 | 15372 | 19162 | 19960 | 22027 | 22029 | 22030 | 22031 | 22032 | 22033 | 22034 | 24000 |  |
| ref | G | G | G | A | T | A | G | T | T | C | A | G |  |
| AY.4 | G | A | T | A | T | - | - | - | - | - | - | G |  |
| AY.4 | G | A | T | A | G | A | A | A | G | T | G | T | #429581 |
| AY.4 | A | G | G | G | G | A | A | A | G | T | G | G |  |
| mut |  |  |  |  |  |  |  |  |  |  |  | G>T |  |

|  |  |  |  |  |  |  |  |  |  |  |  |  |  |  |  |  |  |  |  |  |  |  |  |  |  |  |  |  |  |  |  |  |
| --- | --- | --- | --- | --- | --- | --- | --- | --- | --- | --- | --- | --- | --- | --- | --- | --- | --- | --- | --- | --- | --- | --- | --- | --- | --- | --- | --- | --- | --- | --- | --- | --- |
| pos | 4181 | 5184 | 5584 | 6402 | 7124 | 7393 | 8986 | 9053 | 9891 | 10029 | 11201 | 11332 | 11418 | 11514 | 13019 | 17236 | 18360 | 19220 | 21622 | 22227 | 22801 | 24208 | 24367 | 25702 | 26062 | 27112 | 27516 | 27874 | 28895 | 28916 | 29738 |  |
| ref | G | C | A | C | C | G | C | G | C | C | A | A | T | C | C | A | A | C | C | C | G | C | A | C | G | G | G | C | G | G | C |  |
| AY.75 | G | T | G | C | C | T | C | G | T | C | A | A | C | T | T | A | A | C | T | T | G | C | A | C | T | A | A | C | G | G | A |  |
| AY.75 | G | T | G | C | C | T | C | G | T | C | A | A | C | T | T | G | G | C | T | T | G | T | G | T | G | G | G | T | T | T | A | #429995 |
| AY.103 | T | C | A | T | T | G | T | T | C | T | G | G | T | C | C | G | G | T | C | C | T | T | G | T | G | G | G | T | T | T | C |  |
| mut |  |  |  |  |  |  |  |  |  |  |  |  |  |  |  | A>G | A>G |  |  |  |  |  |  |  |  |  |  |  |  |  | C>A |  |

|  |  |  |  |  |  |  |  |  |  |  |  |  |  |  |  |  |  |  |  |
| --- | --- | --- | --- | --- | --- | --- | --- | --- | --- | --- | --- | --- | --- | --- | --- | --- | --- | --- | --- |
| pos | 884 | 4720 | 4789 | 5544 | 7851 | 9112 | 16914 | 19677 | 19806 | 20726 | 22027 | 22029 | 22030 | 22031 | 22032 | 22033 | 22034 | 29288 |  |
| ref | C | G | T | C | C | C | G | G | C | A | T | A | G | T | T | C | A | T |  |
| AY.4 | C | G | C | T | T | T | G | G | T | G | T | - | - | - | - | - | - | C |  |
| AY.4 | C | G | C | T | T | T | G | G | T | G | G | A | A | A | G | T | G | C | #431516 |
| AY.114 | T | T | T | C | C | C | T | T | C | A | G | A | A | A | G | T | G | T |  |
| mut |  |  |  |  |  |  |  |  |  |  |  |  |  |  |  |  |  | T>C |  |

|  |  |  |  |  |  |  |  |  |  |  |  |  |  |  |  |  |  |  |  |  |  |  |  |  |  |  |  |  |  |  |  |  |  |  |  |  |  |  |  |  |
| --- | --- | --- | --- | --- | --- | --- | --- | --- | --- | --- | --- | --- | --- | --- | --- | --- | --- | --- | --- | --- | --- | --- | --- | --- | --- | --- | --- | --- | --- | --- | --- | --- | --- | --- | --- | --- | --- | --- | --- | --- |
| pos | 881 | 1191 | 1267 | 4181 | 5184 | 6027 | 6040 | 6402 | 7124 | 8986 | 9053 | 9203 | 9678 | 10029 | 11005 | 11201 | 11332 | 14014 | 14688 | 15451 | 16466 | 16915 | 17496 | 18086 | 19220 | 20131 | 20396 | 21792 | 26439 | 27874 | 28248 | 28249 | 28250 | 28251 | 28252 | 28253 | 28916 | 29554 | 29700 |  |
| ref | G | C | C | G | C | C | C | C | C | C | G | G | T | C | C | A | A | T | C | G | C | C | A | C | C | G | A | A | G | C | G | A | T | T | T | C | G | G | A |  |
| B.1.617.2 | G | T | T | G | T | A | C | C | C | C | G | A | C | C | A | A | A | T | T | G | C | T | G | T | C | G | G | C | G | C | G | A | T | T | T | T | G | G | A |  |
| B.1.617.2 | A | T | T | G | T | A | C | C | C | C | G | A | C | C | A | A | A | T | T | G | C | T | G | T | T | T | A | A | T | T | - | - | - | - | - | - | T | T | G | #432690 |
| AY.44 | G | C | C | T | C | C | T | T | T | T | T | G | T | T | C | G | G | G | C | A | T | C | A | C | T | T | A | A | T | T | - | - | - | - | - | - | T | T | G |  |
| mut | G>A |  |  |  |  |  |  |  |  |  |  |  |  |  |  |  |  |  |  |  |  |  |  |  |  |  |  |  |  |  |  |  |  |  |  |  |  |  |  |  |

|  |  |  |  |  |  |  |  |  |  |  |  |  |  |  |  |  |  |
| --- | --- | --- | --- | --- | --- | --- | --- | --- | --- | --- | --- | --- | --- | --- | --- | --- | --- |
| pos | 583 | 3768 | 22026 | 22027 | 22029 | 22030 | 22031 | 22032 | 22033 | 22034 | 22383 | 22537 | 24770 | 25904 | 25983 | 29648 |  |
| ref | C | C | G | T | A | G | T | T | C | A | C | T | G | C | A | G |  |
| AY.100 | C | C | T | G | A | A | A | G | T | G | C | T | G | C | A | G |  |
| AY.100 | T | T | T | G | A | A | A | G | T | G | T | C | T | T | T | T | #432788 |
| AY.100 | T | T | G | T | - | - | - | - | - | - | T | C | T | T | T | T |  |
| mut | C>T | C>T |  |  |  |  |  |  |  |  |  |  |  |  |  |  |  |

|  |  |  |  |  |  |  |  |  |  |  |  |  |
| --- | --- | --- | --- | --- | --- | --- | --- | --- | --- | --- | --- | --- |
| pos | 4084 | 11332 | 15495 | 19019 | 22027 | 22029 | 22030 | 22031 | 22032 | 22033 | 22034 |  |
| ref | C | A | C | C | T | A | G | T | T | C | A |  |
| AY.4 | T | T | T | C | T | - | - | - | - | - | - |  |
| AY.4 | T | T | T | C | T | A | A | A | G | T | G | #434965 |
| AY.4 | C | G | C | T | G | A | A | A | G | T | G |  |
| mut |  |  |  |  |  |  |  |  |  |  |  |  |

|  |  |  |  |  |  |  |  |  |  |  |  |  |  |  |  |  |  |  |  |  |  |  |  |  |  |  |  |  |  |  |  |  |  |  |  |  |
| --- | --- | --- | --- | --- | --- | --- | --- | --- | --- | --- | --- | --- | --- | --- | --- | --- | --- | --- | --- | --- | --- | --- | --- | --- | --- | --- | --- | --- | --- | --- | --- | --- | --- | --- | --- | --- |
| pos | 219 | 593 | 4181 | 5184 | 5584 | 5812 | 6402 | 7124 | 8651 | 8986 | 9053 | 9526 | 9559 | 9891 | 10029 | 11201 | 11332 | 11418 | 11479 | 11514 | 12756 | 13019 | 13665 | 15240 | 17122 | 19220 | 21642 | 21846 | 22227 | 23734 | 27021 | 27368 | 27874 | 28007 | 28916 |  |
| ref | G | C | G | C | A | C | C | C | A | C | G | G | C | C | C | A | A | T | C | C | C | C | C | C | G | C | C | C | C | A | G | A | C | T | G |  |
| AY.100 | G | C | T | C | A | C | T | T | A | T | T | G | T | C | T | G | G | T | C | C | C | C | C | T | T | T | C | T | C | A | G | A | T | C | T |  |
| B.1.617.2 | G | C | T | C | A | C | T | T | A | T | T | G | T | C | T | G | G | T | C | C | C | C | C | T | T | T | A | T | C | A | C | G | C | T | G | #435268 |
| AY.10 | T | T | G | T | G | T | C | C | G | C | G | T | C | T | C | A | A | C | T | T | T | T | T | C | G | C | C | C | T | G | C | A | C | T | G |  |
| mut |  |  |  |  |  |  |  |  |  |  |  |  |  |  |  |  |  |  |  |  |  |  |  |  |  |  | C>A |  |  |  |  | A>G |  |  |  |  |

|  |  |  |  |  |  |  |  |  |  |  |  |  |  |  |  |  |  |  |  |  |  |  |  |
| --- | --- | --- | --- | --- | --- | --- | --- | --- | --- | --- | --- | --- | --- | --- | --- | --- | --- | --- | --- | --- | --- | --- | --- |
| pos | 526 | 1387 | 1392 | 2094 | 4710 | 7711 | 7814 | 7851 | 11417 | 14841 | 16466 | 17040 | 19524 | 21846 | 22072 | 24816 | 27875 | 27881 | 27882 | 27883 | 28178 | 28724 |  |
| ref | G | C | C | C | C | T | T | C | G | A | C | T | C | C | T | G | T | C | G | C | G | C |  |
| AY.4 | G | T | T | T | T | C | C | T | T | C | C | C | C | T | C | C | T | C | G | C | G | C |  |
| AY.4 | G | T | T | T | T | C | C | T | T | C | C | C | C | T | C | C | C | T | C | T | T | T | #436589 |
| AY.98 | T | C | C | C | C | T | T | C | G | A | T | T | T | C | T | G | C | T | C | T | G | C |  |
| mut |  |  |  |  |  |  |  |  |  |  |  |  |  |  |  |  |  |  |  |  | G>T | C>T |  |

|  |  |  |  |  |  |  |  |  |  |  |  |  |  |  |  |  |
| --- | --- | --- | --- | --- | --- | --- | --- | --- | --- | --- | --- | --- | --- | --- | --- | --- |
| pos | 370 | 5392 | 8299 | 10450 | 11281 | 15346 | 17040 | 17721 | 19374 | 19887 | 22713 | 27643 | 27706 | 28191 | 29253 |  |
| ref | G | C | C | C | T | C | T | G | C | A | C | C | G | G | C |  |
| AY.4 | G | C | C | T | C | C | C | T | T | A | C | C | G | T | C |  |
| AY.4 | G | C | C | T | C | C | C | T | T | G | T | T | T | G | T | #438742 |
| AY.4 | T | T | T | C | T | T | T | G | C | A | T | T | T | G | T |  |
| mut |  |  |  |  |  |  |  |  |  | A>G |  |  |  |  |  |  |

|  |  |  |  |  |  |  |  |  |  |  |  |  |  |  |  |
| --- | --- | --- | --- | --- | --- | --- | --- | --- | --- | --- | --- | --- | --- | --- | --- |
| pos | 1418 | 2971 | 6040 | 7840 | 7926 | 12191 | 14014 | 21897 | 21987 | 26107 | 27507 | 29249 | 29670 | 29700 |  |
| ref | G | G | C | C | C | G | T | C | G | G | A | C | C | A |  |
| AY.44 | G | T | T | T | T | T | G | C | G | G | A | T | C | G |  |
| AY.44 | A | T | T | T | T | T | G | T | A | C | C | C | T | A | #439722 |
| AY.25 | G | G | C | C | C | G | T | T | A | C | C | C | C | A |  |
| mut | G>A |  |  |  |  |  |  |  |  |  |  |  | C>T |  |  |

|  |  |  |  |  |  |  |  |  |  |  |  |  |  |  |  |  |  |  |  |  |  |  |  |  |  |  |  |  |  |  |  |  |  |  |  |  |  |  |  |  |  |  |
| --- | --- | --- | --- | --- | --- | --- | --- | --- | --- | --- | --- | --- | --- | --- | --- | --- | --- | --- | --- | --- | --- | --- | --- | --- | --- | --- | --- | --- | --- | --- | --- | --- | --- | --- | --- | --- | --- | --- | --- | --- | --- | --- |
| pos | 507 | 508 | 509 | 518 | 1968 | 2165 | 2919 | 3971 | 4162 | 4181 | 5184 | 5512 | 5584 | 6040 | 6402 | 6638 | 7124 | 7926 | 8634 | 8986 | 9053 | 9891 | 10029 | 11201 | 11332 | 11418 | 11514 | 12056 | 13019 | 13515 | 13701 | 14014 | 16726 | 17122 | 19220 | 22227 | 26681 | 27874 | 28073 | 28916 | 29700 |  |
| ref | A | T | G | A | C | A | C | G | G | G | C | C | A | C | C | C | C | C | T | C | G | C | C | A | A | T | C | T | C | C | T | T | C | G | C | C | C | C | G | G | A |  |
| AY.44 | T | C | A | G | C | A | C | T | G | T | C | C | A | T | T | T | T | T | T | T | T | C | T | G | G | T | C | T | C | C | T | G | T | G | T | C | C | T | T | T | G |  |
| AY.60 | T | C | A | G | T | C | T | G | T | G | T | T | G | C | C | C | C | C | C | C | G | T | C | A | A | C | T | A | T | T | C | T | C | T | C | T | T | C | G | G | A | #440320 |
| AY.60 | A | T | G | A | T | C | T | G | T | G | T | T | G | C | C | C | C | C | C | C | G | T | C | A | A | C | T | A | T | T | C | T | C | T | C | T | T | C | G | G | A |  |
| mut |  |  |  |  |  |  |  |  |  |  |  |  |  |  |  |  |  |  |  |  |  |  |  |  |  |  |  |  |  |  |  |  |  |  |  |  |  |  |  |  |  |  |

|  |  |  |  |  |  |  |  |  |  |  |  |  |  |  |
| --- | --- | --- | --- | --- | --- | --- | --- | --- | --- | --- | --- | --- | --- | --- |
| pos | 373 | 9127 | 18108 | 19118 | 21987 | 22026 | 22027 | 22029 | 22030 | 22031 | 22032 | 22033 | 22034 |  |
| ref | G | T | A | C | G | G | T | A | G | T | T | C | A |  |
| AY.100 | A | C | T | T | G | G | T | - | - | - | - | - | - |  |
| AY.100 | A | C | T | T | G | T | G | A | A | A | G | T | G | #440808 |
| AY.100 | G | T | A | C | A | T | G | A | A | A | G | T | G |  |
| mut |  |  |  |  |  |  |  |  |  |  |  |  |  |  |

|  |  |  |  |  |  |  |  |  |  |  |  |  |  |  |  |  |  |  |  |  |
| --- | --- | --- | --- | --- | --- | --- | --- | --- | --- | --- | --- | --- | --- | --- | --- | --- | --- | --- | --- | --- |
| pos | 22027 | 22029 | 22030 | 22031 | 22032 | 22033 | 22034 | 23039 | 26497 | 26505 | 27059 | 27406 | 29253 | 29700 | 29762 | 29769 | 29774 | 29779 | 29781 |  |
| ref | T | A | G | T | T | C | A | C | T | G | C | C | C | A | C | C | C | G | G |  |
| AY.4 | G | A | A | A | G | T | G | C | T | G | C | C | C | A | C | C | C | G | G |  |
| AY.4 | G | A | A | A | G | T | G | G | C | C | T | A | T | A | C | C | C | G | G | #443410 |
| AY.4 | T | - | - | - | - | - | - | G | C | C | T | A | T | - | - | - | - | - | - |  |
| mut |  |  |  |  |  |  |  |  |  |  |  |  |  | ->A | ->C | ->C | ->C | ->G | ->G |  |

|  |  |  |  |  |  |  |  |  |  |  |  |  |  |
| --- | --- | --- | --- | --- | --- | --- | --- | --- | --- | --- | --- | --- | --- |
| pos | 1321 | 22027 | 22029 | 22030 | 22031 | 22032 | 22033 | 22034 | 27739 | 27828 | 29540 | 29645 |  |
| ref | A | T | A | G | T | T | C | A | C | C | G | G |  |
| AY.4 | A | G | A | A | A | G | T | G | C | C | G | G |  |
| AY.4 | C | G | A | A | A | G | T | G | A | T | A | T | #443499 |
| AY.4 | C | T | - | - | - | - | - | - | A | T | A | T |  |
| mut | A>C |  |  |  |  |  |  |  |  |  |  |  |  |

|  |  |  |  |  |  |  |  |  |  |  |  |  |  |  |  |  |  |  |
| --- | --- | --- | --- | --- | --- | --- | --- | --- | --- | --- | --- | --- | --- | --- | --- | --- | --- | --- |
| pos | 3689 | 5241 | 6492 | 11674 | 22026 | 22027 | 22029 | 22030 | 22031 | 22032 | 22033 | 22034 | 25297 | 25678 | 26681 | 28209 | 29660 |  |
| ref | G | C | T | C | G | T | A | G | T | T | C | A | G | C | C | G | T |  |
| AY.39 | G | C | T | C | T | G | A | A | A | G | T | G | G | C | C | G | T |  |
| AY.39 | G | C | T | T | T | G | A | A | A | G | T | G | A | T | T | C | A | #448284 |
| AY.39 | A | T | C | T | G | T | - | - | - | - | - | - | A | T | T | C | A |  |
| mut |  |  |  | C>T |  |  |  |  |  |  |  |  |  |  |  |  |  |  |

|  |  |  |  |  |  |  |  |  |  |  |  |  |  |  |  |
| --- | --- | --- | --- | --- | --- | --- | --- | --- | --- | --- | --- | --- | --- | --- | --- |
| pos | 3768 | 5526 | 7042 | 10271 | 11782 | 13694 | 13712 | 22027 | 22029 | 22030 | 22031 | 22032 | 22033 | 22034 |  |
| ref | C | C | G | G | A | C | A | T | A | G | T | T | C | A |  |
| AY.118 | T | T | T | G | G | T | G | T | - | - | - | - | - | - |  |
| AY.118 | T | T | T | G | G | T | G | G | A | A | A | G | T | G | #448295 |
| AY.118 | C | C | G | A | A | C | A | G | A | A | A | G | T | G |  |
| mut |  |  |  |  |  |  |  |  |  |  |  |  |  |  |  |

|  |  |  |  |  |  |  |  |  |  |  |  |  |  |  |  |  |  |  |  |  |  |
| --- | --- | --- | --- | --- | --- | --- | --- | --- | --- | --- | --- | --- | --- | --- | --- | --- | --- | --- | --- | --- | --- |
| pos | 929 | 7851 | 10691 | 10833 | 11344 | 12071 | 13424 | 14919 | 18252 | 19160 | 20968 | 22026 | 22027 | 22029 | 22030 | 22031 | 22032 | 22033 | 22034 | 28606 |  |
| ref | A | C | A | C | C | G | C | C | C | C | G | G | T | A | G | T | T | C | A | C |  |
| AY.116.1 | G | C | G | T | T | A | C | C | T | T | A | G | T | - | - | - | - | - | - | T |  |
| AY.116.1 | G | C | G | T | T | A | C | C | T | T | A | T | G | A | A | A | G | T | G | C | #448483 |
| AY.4 | A | T | A | C | C | G | T | T | C | C | G | T | G | A | A | A | G | T | G | C |  |
| mut |  |  |  |  |  |  |  |  |  |  |  |  |  |  |  |  |  |  |  |  |  |

|  |  |  |  |  |  |  |  |  |  |  |  |  |  |  |  |  |
| --- | --- | --- | --- | --- | --- | --- | --- | --- | --- | --- | --- | --- | --- | --- | --- | --- |
| pos | 7528 | 16878 | 22027 | 22029 | 22030 | 22031 | 22032 | 22033 | 22034 | 23043 | 25688 | 26763 | 27967 | 28838 | 29367 |  |
| ref | C | A | T | A | G | T | T | C | A | C | C | G | G | C | C |  |
| AY.4 | C | A | G | A | A | A | G | T | G | C | C | G | G | C | C |  |
| AY.4 | T | T | G | A | A | A | G | T | G | T | T | T | T | T | T | #449556 |
| AY.4 | T | T | T | - | - | - | - | - | - | T | T | T | T | T | T |  |
| mut | C>T | A>T |  |  |  |  |  |  |  |  |  |  |  |  |  |  |

|  |  |  |  |  |  |  |  |  |  |  |  |  |  |  |  |  |  |  |
| --- | --- | --- | --- | --- | --- | --- | --- | --- | --- | --- | --- | --- | --- | --- | --- | --- | --- | --- |
| pos | 884 | 3122 | 4720 | 4927 | 7851 | 16914 | 17040 | 17763 | 19677 | 19989 | 22027 | 22029 | 22030 | 22031 | 22032 | 22033 | 22034 |  |
| ref | C | G | G | C | C | G | T | T | G | T | T | A | G | T | T | C | A |  |
| AY.4 | C | T | G | T | T | G | C | C | G | C | T | - | - | - | - | - | - |  |
| AY.4 | C | T | G | T | T | G | C | C | G | C | G | A | A | A | G | T | G | #451123 |
| AY.114 | T | G | T | C | C | T | T | T | T | T | G | A | A | A | G | T | G |  |
| mut |  |  |  |  |  |  |  |  |  |  |  |  |  |  |  |  |  |  |

|  |  |  |  |  |  |  |  |  |  |  |  |  |  |  |  |  |  |  |  |
| --- | --- | --- | --- | --- | --- | --- | --- | --- | --- | --- | --- | --- | --- | --- | --- | --- | --- | --- | --- |
| pos | 884 | 4720 | 5566 | 7851 | 16914 | 18508 | 19480 | 19677 | 19862 | 22027 | 22029 | 22030 | 22031 | 22032 | 22033 | 22034 | 24821 | 26455 |  |
| ref | C | G | T | C | G | C | G | G | C | T | A | G | T | T | C | A | G | C |  |
| AY.4 | C | G | C | T | G | T | T | G | T | T | - | - | - | - | - | - | T | T |  |
| AY.4 | C | G | C | T | G | T | T | G | T | G | A | A | A | G | T | G | T | T | #451160 |
| AY.114 | T | T | T | C | T | C | G | T | C | G | A | A | A | G | T | G | G | C |  |
| mut |  |  |  |  |  |  |  |  |  |  |  |  |  |  |  |  | G>T | C>T |  |

|  |  |  |  |  |  |  |  |  |  |  |  |  |  |  |  |  |  |  |  |  |  |  |  |  |  |  |  |  |  |  |  |  |  |  |  |  |  |  |
| --- | --- | --- | --- | --- | --- | --- | --- | --- | --- | --- | --- | --- | --- | --- | --- | --- | --- | --- | --- | --- | --- | --- | --- | --- | --- | --- | --- | --- | --- | --- | --- | --- | --- | --- | --- | --- | --- | --- |
| pos | 2911 | 4181 | 5184 | 5584 | 5784 | 6402 | 7124 | 8986 | 9053 | 9891 | 10029 | 10075 | 11201 | 11332 | 11418 | 11456 | 11514 | 12097 | 13019 | 15114 | 19220 | 21318 | 22227 | 22741 | 25352 | 27345 | 27874 | 28916 | 29050 | 29509 | 29700 | 29750 | 29762 | 29769 | 29774 | 29779 | 29781 |  |
| ref | T | G | C | A | C | C | C | C | G | C | C | A | A | A | T | A | C | A | C | T | C | T | C | T | G | A | C | G | G | C | A | C | C | C | C | G | G |  |
| AY.26 | G | G | T | G | C | C | C | C | G | T | C | A | A | A | C | A | T | G | T | T | C | T | T | T | T | T | C | G | G | C | A | C | C | C | C | G | G |  |
| B.1.617.2 | G | G | T | G | C | C | C | C | G | T | C | A | A | A | C | A | T | G | T | T | C | T | C | C | G | A | T | T | A | T | - | T | - | - | - | - | - | #452874 |
| AY.3 | T | T | C | A | T | T | T | T | T | C | T | G | G | G | T | G | C | A | C | C | T | C | C | C | G | A | T | T | A | T | - | T | - | - | - | - | - |  |
| mut |  |  |  |  |  |  |  |  |  |  |  |  |  |  |  |  |  |  |  |  |  |  |  |  |  |  |  |  |  |  |  |  |  |  |  |  |  |  |

|  |  |  |  |  |  |  |  |  |  |  |  |  |  |  |  |  |  |  |  |  |  |  |  |  |  |  |  |
| --- | --- | --- | --- | --- | --- | --- | --- | --- | --- | --- | --- | --- | --- | --- | --- | --- | --- | --- | --- | --- | --- | --- | --- | --- | --- | --- | --- |
| pos | 3267 | 3365 | 5241 | 5607 | 7643 | 7712 | 9653 | 9857 | 12202 | 13335 | 14745 | 15933 | 15952 | 16716 | 16935 | 17562 | 18348 | 18693 | 18744 | 19086 | 19875 | 20083 | 20280 | 24781 | 28299 | 29751 |  |
| ref | C | C | C | A | G | C | G | C | G | C | T | C | C | C | G | G | C | C | C | G | C | G | A | G | A | G |  |
| B.1.617.2 | T | T | T | G | G | T | T | T | T | T | C | T | C | T | G | G | T | C | C | G | T | A | C | G | A | C |  |
| AY.43 | T | T | T | G | G | T | G | C | G | C | T | C | A | C | T | T | C | A | T | T | C | G | A | T | T | G | #454594 |
| AY.43 | C | C | C | A | T | C | G | C | G | C | T | C | A | C | T | T | C | A | T | T | C | G | A | T | T | G |  |
| mut |  |  |  |  |  |  |  |  |  |  |  |  |  |  |  |  |  |  |  |  |  |  |  |  |  |  |  |

|  |  |  |  |  |  |  |  |  |  |  |  |  |  |  |
| --- | --- | --- | --- | --- | --- | --- | --- | --- | --- | --- | --- | --- | --- | --- |
| pos | 2102 | 13423 | 13424 | 13425 | 22027 | 22029 | 22030 | 22031 | 22032 | 22033 | 22034 | 22102 | 28834 |  |
| ref | C | C | C | G | T | A | G | T | T | C | A | A | A |  |
| AY.107 | T | A | A | A | T | - | - | - | - | - | - | T | T |  |
| AY.107 | T | A | A | A | G | A | A | A | G | T | G | T | T | #456023 |
| AY.107 | C | C | C | G | G | A | A | A | G | T | G | A | A |  |
| mut |  |  |  |  |  |  |  |  |  |  |  | A>T | A>T |  |

|  |  |  |  |  |  |  |  |  |  |  |  |  |  |  |  |  |  |  |  |  |  |  |  |  |  |  |  |  |  |  |  |  |  |  |  |  |
| --- | --- | --- | --- | --- | --- | --- | --- | --- | --- | --- | --- | --- | --- | --- | --- | --- | --- | --- | --- | --- | --- | --- | --- | --- | --- | --- | --- | --- | --- | --- | --- | --- | --- | --- | --- | --- |
| pos | 1191 | 1267 | 2523 | 4181 | 5184 | 6402 | 7124 | 7528 | 8334 | 8986 | 9053 | 9891 | 10029 | 10977 | 11201 | 11332 | 11418 | 12778 | 12946 | 13862 | 16593 | 17019 | 18176 | 18468 | 19220 | 20262 | 21846 | 21987 | 23202 | 26076 | 27739 | 27874 | 28386 | 28916 | 29744 |  |
| ref | C | C | C | G | C | C | C | C | C | C | G | C | C | C | A | A | T | C | T | C | T | G | C | A | C | A | C | G | C | T | C | C | A | G | G |  |
| AY.46 | C | C | T | T | C | T | T | C | C | T | T | C | T | T | G | G | T | C | T | T | C | G | C | A | T | A | T | A | C | T | C | T | G | T | G |  |
| AY.46 | C | C | T | T | C | T | T | T | T | T | T | C | T | T | G | G | T | T | C | C | T | T | T | G | C | G | C | G | T | A | T | C | A | G | A | #456614 |
| AY.16 | T | T | C | G | T | C | C | C | C | C | G | T | C | C | A | A | C | C | C | C | T | T | T | G | C | G | C | A | C | A | T | C | A | G | A |  |
| mut |  |  |  |  |  |  |  | C>T | C>T |  |  |  |  |  |  |  |  | C>T |  |  |  |  |  |  |  |  |  | A>G | C>T |  |  |  |  |  |  |  |

|  |  |  |  |  |  |  |  |  |  |  |  |  |  |  |  |  |  |  |  |  |  |
| --- | --- | --- | --- | --- | --- | --- | --- | --- | --- | --- | --- | --- | --- | --- | --- | --- | --- | --- | --- | --- | --- |
| pos | 884 | 1710 | 4720 | 7851 | 10440 | 11332 | 16914 | 16992 | 17040 | 17747 | 19677 | 21668 | 22027 | 22029 | 22030 | 22031 | 22032 | 22033 | 22034 | 23029 |  |
| ref | C | C | G | C | C | A | G | A | T | C | G | G | T | A | G | T | T | C | A | C |  |
| AY.4 | C | T | G | T | T | T | G | G | C | T | G | T | T | - | - | - | - | - | - | T |  |
| AY.4 | C | T | G | T | T | T | G | G | C | T | G | T | G | A | A | A | G | T | G | C | #460012 |
| AY.114 | T | C | T | C | C | G | T | A | T | C | T | G | G | A | A | A | G | T | G | C |  |
| mut |  |  |  |  |  |  |  |  |  |  |  |  |  |  |  |  |  |  |  |  |  |

|  |  |  |  |  |  |  |  |  |  |  |  |  |  |  |  |  |  |  |
| --- | --- | --- | --- | --- | --- | --- | --- | --- | --- | --- | --- | --- | --- | --- | --- | --- | --- | --- |
| pos | 884 | 2216 | 4720 | 7015 | 7851 | 8895 | 11332 | 16914 | 19677 | 20838 | 22027 | 22029 | 22030 | 22031 | 22032 | 22033 | 22034 |  |
| ref | C | T | G | A | C | C | A | G | G | T | T | A | G | T | T | C | A |  |
| AY.4 | C | C | G | C | T | T | T | G | G | C | T | - | - | - | - | - | - |  |
| AY.4 | C | C | G | C | T | T | T | G | G | C | G | A | A | A | G | T | G | #460081 |
| AY.114 | T | T | T | A | C | C | G | T | T | T | G | A | A | A | G | T | G |  |
| mut |  |  |  |  |  |  |  |  |  |  |  |  |  |  |  |  |  |  |

|  |  |  |  |  |  |  |  |  |  |  |  |  |  |  |  |
| --- | --- | --- | --- | --- | --- | --- | --- | --- | --- | --- | --- | --- | --- | --- | --- |
| pos | 5221 | 5284 | 9241 | 10318 | 12151 | 17243 | 22027 | 22029 | 22030 | 22031 | 22032 | 22033 | 22034 | 29764 |  |
| ref | C | C | A | G | A | C | T | A | G | T | T | C | A | G |  |
| AY.20 | T | T | T | A | G | T | T | - | - | - | - | - | - | T |  |
| AY.20 | T | T | T | A | G | T | G | A | A | A | G | T | G | T | #460554 |
| AY.20 | C | C | A | G | A | C | G | A | A | A | G | T | G | G |  |
| mut |  |  |  |  |  |  |  |  |  |  |  |  |  | G>T |  |

|  |  |  |  |  |  |  |  |  |  |  |  |  |  |  |
| --- | --- | --- | --- | --- | --- | --- | --- | --- | --- | --- | --- | --- | --- | --- |
| pos | 520 | 7644 | 12473 | 14033 | 15397 | 22027 | 22029 | 22030 | 22031 | 22032 | 22033 | 22034 | 29555 |  |
| ref | G | C | C | A | T | T | A | G | T | T | C | A | C |  |
| AY.20 | G | C | C | G | T | T | - | - | - | - | - | - | T |  |
| AY.20 | G | C | T | G | T | G | A | A | A | G | T | G | T | #462967 |
| AY.20 | T | T | C | A | C | T | A | A | A | G | T | G | C |  |
| mut |  |  | C>T |  |  | T>G |  |  |  |  |  |  | C>T |  |

|  |  |  |  |  |  |  |  |  |  |  |  |  |  |  |  |  |  |  |  |
| --- | --- | --- | --- | --- | --- | --- | --- | --- | --- | --- | --- | --- | --- | --- | --- | --- | --- | --- | --- |
| pos | 884 | 4720 | 7229 | 7851 | 9104 | 13821 | 13965 | 16914 | 17040 | 19677 | 22027 | 22029 | 22030 | 22031 | 22032 | 22033 | 22034 | 26110 |  |
| ref | C | G | G | C | A | C | C | G | T | G | T | A | G | T | T | C | A | C |  |
| AY.4 | C | G | T | T | G | T | T | G | C | G | T | - | - | - | - | - | - | T |  |
| AY.4 | C | G | T | T | G | T | T | G | C | G | G | A | A | A | G | T | G | T | #465275 |
| AY.114 | T | T | G | C | A | C | C | T | T | T | G | A | A | A | G | T | G | C |  |
| mut |  |  |  |  |  |  |  |  |  |  |  |  |  |  |  |  |  | C>T |  |

|  |  |  |  |  |  |  |  |  |  |  |  |  |  |  |  |  |  |  |
| --- | --- | --- | --- | --- | --- | --- | --- | --- | --- | --- | --- | --- | --- | --- | --- | --- | --- | --- |
| pos | 263 | 884 | 4720 | 5991 | 7851 | 16914 | 18604 | 19387 | 19677 | 21516 | 22027 | 22029 | 22030 | 22031 | 22032 | 22033 | 22034 |  |
| ref | A | C | G | A | C | G | G | A | G | C | T | A | G | T | T | C | A |  |
| AY.4 | G | C | G | C | T | G | G | C | G | T | T | - | - | - | - | - | - |  |
| AY.4 | G | C | G | C | T | G | T | C | G | T | G | A | A | A | G | T | G | #465295 |
| AY.114 | A | T | T | A | C | T | G | A | T | C | G | A | A | A | G | T | G |  |
| mut |  |  |  |  |  |  | G>T |  |  |  |  |  |  |  |  |  |  |  |

|  |  |  |  |  |  |  |  |  |  |  |  |  |  |  |  |  |  |  |  |  |  |  |  |  |  |  |  |  |  |  |  |  |  |  |  |  |  |  |  |  |  |  |  |  |  |  |  |  |  |  |  |  |  |  |  |  |  |  |  |  |  |  |  |  |  |  |  |  |  |  |  |  |  |  |  |  |  |  |  |  |  |  |  |
| --- | --- | --- | --- | --- | --- | --- | --- | --- | --- | --- | --- | --- | --- | --- | --- | --- | --- | --- | --- | --- | --- | --- | --- | --- | --- | --- | --- | --- | --- | --- | --- | --- | --- | --- | --- | --- | --- | --- | --- | --- | --- | --- | --- | --- | --- | --- | --- | --- | --- | --- | --- | --- | --- | --- | --- | --- | --- | --- | --- | --- | --- | --- | --- | --- | --- | --- | --- | --- | --- | --- | --- | --- | --- | --- | --- | --- | --- | --- | --- | --- | --- | --- | --- |
| pos | 210 | 733 | 2749 | 3828 | 4181 | 5648 | 6319 | 6402 | 6613 | 7124 | 8986 | 9053 | 10029 | 11201 | 11288 | 11289 | 11290 | 11291 | 11292 | 11293 | 11294 | 11295 | 11296 | 11332 | 11537 | 12778 | 13860 | 15451 | 15522 | 15952 | 16466 | 17259 | 18744 | 19220 | 21614 | 21618 | 21621 | 21638 | 21987 | 22029 | 22030 | 22031 | 22032 | 22033 | 22034 | 22132 | 22812 | 22917 | 22995 | 23012 | 23063 | 23525 | 23604 | 24130 | 24410 | 24642 | 25088 | 25469 | 26149 | 26767 | 27638 | 27752 | 27874 | 28167 | 28248 | 28249 | 28250 | 28251 | 28252 | 28253 | 28271 | 28299 | 28461 | 28512 | 28877 | 28878 | 28881 | 28882 | 28883 | 28916 | 29402 | 29742 |  |
| ref | G | T | C | C | G | A | A | C | A | C | C | G | C | A | T | C | T | G | G | T | T | T | T | A | A | C | C | G | T | C | C | G | C | C | C | C | C | C | G | A | G | T | T | C | A | G | A | T | C | G | A | C | C | C | G | C | G | C | T | T | T | C | C | G | G | A | T | T | T | C | A | A | A | C | A | G | G | G | G | G | G | G |  |
| P.1 | G | C | T | T | G | C | G | C | G | C | C | G | C | A | - | - | - | - | - | - | - | - | - | A | A | T | T | G | T | C | C | T | C | C | T | C | A | T | G | A | G | T | T | C | A | T | C | T | C | A | T | T | C | C | G | T | T | C | C | T | T | C | C | A | G | A | T | T | T | C | A | A | A | G | T | C | A | A | C | G | G | G |  |
| AY.43 | G | C | T | T | G | C | G | C | G | C | T | T | T | G | T | C | T | G | G | T | T | T | T | G | G | C | C | A | C | A | T | G | T | T | C | G | C | C | A | - | - | - | - | - | - | G | A | G | A | G | A | C | G | T | A | C | G | T | T | C | C | T | T | G | - | - | - | - | - | - | - | T | G | C | A | G | T | G | G | T | T | T | #465910 |
| AY.43 | T | T | C | C | T | A | A | T | A | T | T | T | T | G | T | C | T | G | G | T | T | T | T | G | A | C | C | A | T | A | T | G | T | T | C | G | C | C | A | - | - | - | - | - | - | G | A | G | A | G | A | C | G | T | A | C | G | T | T | C | C | T | T | G | - | - | - | - | - | - | - | T | G | C | A | G | T | G | G | T | T | T |  |
| mut |  |  |  |  |  |  |  |  |  |  |  |  |  |  |  |  |  |  |  |  |  |  |  |  | A>G |  |  |  | T>C |  |  |  |  |  |  |  |  |  |  |  |  |  |  |  |  |  |  |  |  |  |  |  |  |  |  |  |  |  |  |  |  |  |  |  |  |  |  |  |  |  |  |  |  |  |  |  |  |  |  |  |  |  |  |

|  |  |  |  |  |  |  |  |  |  |  |  |  |  |  |  |  |  |  |  |  |  |  |  |  |  |  |  |  |  |  |  |
| --- | --- | --- | --- | --- | --- | --- | --- | --- | --- | --- | --- | --- | --- | --- | --- | --- | --- | --- | --- | --- | --- | --- | --- | --- | --- | --- | --- | --- | --- | --- | --- |
| pos | 1191 | 1267 | 1877 | 4181 | 5184 | 6040 | 6402 | 7124 | 7135 | 7926 | 8986 | 9053 | 9891 | 10029 | 11201 | 11332 | 11418 | 12946 | 14014 | 18176 | 19160 | 19220 | 20014 | 20262 | 20718 | 27739 | 27853 | 27874 | 28916 | 29700 |  |
| ref | C | C | T | G | C | C | C | C | T | C | C | G | C | C | A | A | T | T | T | C | C | C | G | A | G | C | A | C | G | A |  |
| AY.44 | C | C | T | T | C | T | T | T | C | T | T | T | C | T | G | G | T | T | G | C | C | T | G | A | G | C | A | T | T | G |  |
| AY.44 | C | C | T | T | C | T | T | T | C | T | T | T | C | T | G | G | T | T | G | C | C | T | G | A | G | T | T | C | G | A | #467049 |
| AY.14 | T | T | G | G | T | C | C | C | T | C | C | G | T | C | A | A | C | C | T | T | T | C | T | G | T | T | T | C | G | A |  |
| mut |  |  |  |  |  |  |  |  |  |  |  |  |  |  |  |  |  |  |  |  |  |  |  |  |  |  |  |  |  |  |  |

|  |  |  |  |  |  |  |  |  |  |  |  |  |  |  |  |
| --- | --- | --- | --- | --- | --- | --- | --- | --- | --- | --- | --- | --- | --- | --- | --- |
| pos | 526 | 2094 | 6896 | 7814 | 7851 | 15950 | 17040 | 19524 | 19983 | 21846 | 27875 | 27881 | 27882 | 27883 |  |
| ref | G | C | C | T | C | T | T | C | C | C | T | C | G | C |  |
| AY.4 | G | T | C | C | T | T | C | C | C | T | T | C | G | C |  |
| AY.4 | G | T | T | C | T | C | C | C | T | T | C | T | C | T | #468967 |
| AY.98 | T | C | C | T | C | T | T | T | C | C | C | T | C | T |  |
| mut |  |  | C>T |  |  | T>C |  |  | C>T |  |  |  |  |  |  |

|  |  |  |  |  |  |  |  |  |  |  |  |  |  |  |  |  |  |  |  |  |  |  |  |
| --- | --- | --- | --- | --- | --- | --- | --- | --- | --- | --- | --- | --- | --- | --- | --- | --- | --- | --- | --- | --- | --- | --- | --- |
| pos | 556 | 3714 | 5526 | 9967 | 11456 | 14395 | 17236 | 22480 | 23222 | 23284 | 24208 | 25339 | 25593 | 27654 | 29050 | 29227 | 29509 | 29762 | 29769 | 29774 | 29779 | 29781 |  |
| ref | C | C | C | C | A | A | A | C | G | T | C | C | G | C | G | G | C | C | C | C | G | G |  |
| AY.103 | C | C | T | T | A | A | G | C | C | T | T | C | G | C | G | A | C | C | C | C | G | G |  |
| AY.3 | C | T | T | T | A | A | A | T | G | C | C | T | A | T | A | G | T | C | C | C | G | G | #472124 |
| AY.3 | T | C | C | C | G | G | A | T | G | C | C | T | A | T | A | G | T | - | - | - | - | - |  |
| mut |  | C>T |  |  |  |  |  |  |  |  |  |  |  |  |  |  |  | ->C | ->C | ->C | ->G | ->G |  |

|  |  |  |  |  |  |  |  |  |  |  |  |  |  |  |
| --- | --- | --- | --- | --- | --- | --- | --- | --- | --- | --- | --- | --- | --- | --- |
| pos | 526 | 1391 | 7851 | 17040 | 19524 | 21846 | 21995 | 22227 | 25614 | 27875 | 27881 | 27882 | 27883 |  |
| ref | G | T | C | T | C | C | T | C | C | T | C | G | C |  |
| AY.4.2 | G | C | T | C | C | T | C | T | T | T | C | G | C |  |
| AY.4.2 | G | C | T | C | C | T | C | T | T | C | T | C | T | #473494 |
| AY.98 | T | T | C | T | T | C | T | C | C | C | T | C | T |  |
| mut |  |  |  |  |  |  |  |  |  |  |  |  |  |  |

|  |  |  |  |  |  |  |  |  |  |  |  |  |  |  |  |  |  |  |
| --- | --- | --- | --- | --- | --- | --- | --- | --- | --- | --- | --- | --- | --- | --- | --- | --- | --- | --- |
| pos | 526 | 1593 | 2518 | 3152 | 4237 | 6349 | 7851 | 7978 | 17002 | 17040 | 19524 | 21846 | 25567 | 28312 | 28378 | 28725 | 29024 |  |
| ref | G | C | G | G | T | G | C | A | C | T | C | C | G | C | G | C | G |  |
| AY.98 | T | C | T | T | T | G | C | A | C | T | T | C | G | C | G | T | G |  |
| AY.98 | T | C | T | T | T | G | C | A | T | C | C | T | A | A | C | C | T | #478391 |
| AY.4 | G | T | G | G | C | T | T | G | C | C | C | T | A | A | C | C | T |  |
| mut |  |  |  |  |  |  |  |  | C>T |  |  |  |  |  |  |  |  |  |

|  |  |  |  |  |  |  |  |  |  |  |  |  |  |  |  |  |  |
| --- | --- | --- | --- | --- | --- | --- | --- | --- | --- | --- | --- | --- | --- | --- | --- | --- | --- |
| pos | 224 | 412 | 3243 | 7423 | 11238 | 19862 | 21077 | 22027 | 22029 | 22030 | 22031 | 22032 | 22033 | 22034 | 29023 | 29616 |  |
| ref | T | C | G | C | G | C | C | T | A | G | T | T | C | A | T | G |  |
| AY.36 | C | T | G | T | A | T | T | T | - | - | - | - | - | - | A | T |  |
| AY.36 | C | T | G | T | A | T | T | G | A | A | A | G | T | G | A | T | #478611 |
| AY.36 | T | C | A | C | G | C | C | G | A | A | A | G | T | G | T | G |  |
| mut |  |  |  |  |  |  |  |  |  |  |  |  |  |  | T>A | G>T |  |

|  |  |  |  |  |  |  |  |  |  |  |  |  |  |  |  |  |  |  |  |  |  |  |  |  |
| --- | --- | --- | --- | --- | --- | --- | --- | --- | --- | --- | --- | --- | --- | --- | --- | --- | --- | --- | --- | --- | --- | --- | --- | --- |
| pos | 884 | 2003 | 4057 | 4582 | 4720 | 6790 | 7851 | 16914 | 18292 | 19677 | 21045 | 22027 | 22028 | 22029 | 22030 | 22031 | 22032 | 22033 | 22034 | 23709 | 24023 | 25587 | 28093 |  |
| ref | C | C | T | C | G | A | C | G | G | G | T | T | G | A | G | T | T | C | A | C | C | C | C |  |
| AY.4 | C | A | C | T | G | G | T | G | T | G | C | T | G | - | - | - | - | - | - | T | T | A | T |  |
| AY.4 | C | A | C | T | G | G | T | G | T | G | C | G | A | A | A | A | G | T | G | T | T | A | T | #478620 |
| AY.114 | T | C | T | C | T | A | C | T | G | T | T | G | G | A | A | A | G | T | G | C | C | C | C |  |
| mut |  |  |  |  |  |  |  |  |  |  |  |  | G>A |  |  |  |  |  |  | C>T | C>T | C>A | C>T |  |

|  |  |  |  |  |  |  |  |  |  |  |  |  |  |  |  |  |  |  |  |  |  |  |  |
| --- | --- | --- | --- | --- | --- | --- | --- | --- | --- | --- | --- | --- | --- | --- | --- | --- | --- | --- | --- | --- | --- | --- | --- |
| pos | 1588 | 2231 | 2632 | 2946 | 3304 | 3433 | 3476 | 6726 | 7669 | 8655 | 8841 | 13929 | 15654 | 19735 | 22027 | 22029 | 22030 | 22031 | 22032 | 22033 | 22034 | 27281 |  |
| ref | A | G | G | G | G | G | G | C | A | C | C | T | C | G | T | A | G | T | T | C | A | G |  |
| AY.39 | A | T | G | A | T | T | G | C | G | T | T | C | C | T | T | - | - | - | - | - | - | T |  |
| AY.39 | A | T | G | A | T | T | G | C | G | T | T | C | C | T | G | A | A | A | G | T | G | G | #480359 |
| AY.39 | G | G | T | G | G | G | A | T | A | C | C | T | T | G | G | A | A | A | G | T | G | G |  |
| mut |  |  |  |  |  |  |  |  |  |  |  |  |  |  |  |  |  |  |  |  |  |  |  |

|  |  |  |  |  |  |  |  |  |  |  |  |  |  |  |  |  |  |  |  |  |
| --- | --- | --- | --- | --- | --- | --- | --- | --- | --- | --- | --- | --- | --- | --- | --- | --- | --- | --- | --- | --- |
| pos | 884 | 3896 | 4720 | 4891 | 7851 | 13459 | 13517 | 16914 | 19677 | 20006 | 22027 | 22029 | 22030 | 22031 | 22032 | 22033 | 22034 | 28367 | 28690 |  |
| ref | C | G | G | C | C | G | C | G | G | G | T | A | G | T | T | C | A | C | G |  |
| AY.4 | C | T | G | T | T | T | T | G | G | T | T | - | - | - | - | - | - | T | G |  |
| AY.4 | C | T | G | T | T | T | T | G | G | T | G | A | A | A | G | T | G | T | T | #482849 |
| AY.114 | T | G | T | C | C | G | C | T | T | G | G | A | A | A | G | T | G | C | G |  |
| mut |  |  |  |  |  |  |  |  |  |  |  |  |  |  |  |  |  | C>T | G>T |  |

|  |  |  |  |  |  |  |  |  |  |  |  |  |  |  |  |  |  |  |  |  |  |  |  |  |
| --- | --- | --- | --- | --- | --- | --- | --- | --- | --- | --- | --- | --- | --- | --- | --- | --- | --- | --- | --- | --- | --- | --- | --- | --- |
| pos | 466 | 884 | 2156 | 3828 | 4551 | 4720 | 7851 | 8456 | 9118 | 10681 | 11824 | 14271 | 16914 | 17040 | 19677 | 20568 | 22027 | 22029 | 22030 | 22031 | 22032 | 22033 | 22034 |  |
| ref | C | C | C | C | C | G | C | A | C | C | C | A | G | T | G | G | T | A | G | T | T | C | A |  |
| AY.4 | T | C | T | T | T | G | T | G | T | T | T | G | G | C | G | T | T | - | - | - | - | - | - |  |
| AY.4 | T | C | T | T | T | G | T | G | T | T | T | G | G | C | G | T | G | A | A | A | G | T | G | #483039 |
| AY.114 | C | T | C | C | C | T | C | A | C | C | C | A | T | T | T | G | G | A | A | A | G | T | G |  |
| mut |  |  |  |  |  |  |  |  |  |  |  |  |  |  |  |  |  |  |  |  |  |  |  |  |

|  |  |  |  |  |  |  |  |  |  |  |  |  |  |  |  |  |  |  |  |  |  |  |  |  |  |  |  |  |  |  |  |  |  |  |  |  |  |  |  |
| --- | --- | --- | --- | --- | --- | --- | --- | --- | --- | --- | --- | --- | --- | --- | --- | --- | --- | --- | --- | --- | --- | --- | --- | --- | --- | --- | --- | --- | --- | --- | --- | --- | --- | --- | --- | --- | --- | --- | --- |
| pos | 2395 | 2973 | 4181 | 5184 | 5584 | 6402 | 7124 | 7933 | 8140 | 8986 | 9053 | 9891 | 10029 | 10702 | 11201 | 11332 | 11418 | 11514 | 13019 | 14202 | 16221 | 18377 | 19220 | 21897 | 22093 | 22227 | 22482 | 23127 | 25352 | 26107 | 26111 | 27262 | 27345 | 27507 | 27874 | 28916 | 29586 | 29762 |  |
| ref | C | C | G | C | A | C | C | A | C | C | G | C | C | C | A | A | T | C | C | G | G | C | C | C | G | C | C | C | G | G | C | A | A | A | C | G | C | C |  |
| AY.25 | T | C | T | C | A | T | T | G | T | T | T | C | T | C | G | G | T | C | C | G | G | C | T | T | G | C | C | T | G | C | C | G | A | C | T | T | T | C |  |
| B.1.617.2 | T | C | T | C | A | T | T | G | T | T | T | C | T | C | G | G | T | C | C | G | A | T | C | C | T | T | T | C | T | G | T | A | T | A | C | G | C | - | #483758 |
| AY.26 | C | T | G | T | G | C | C | A | C | C | G | T | C | T | A | A | C | T | T | A | A | C | C | C | T | T | T | C | T | G | T | A | T | A | C | G | C | - |  |
| mut |  |  |  |  |  |  |  |  |  |  |  |  |  |  |  |  |  |  |  |  |  | C>T |  |  |  |  |  |  |  |  |  |  |  |  |  |  |  |  |  |

|  |  |  |  |  |  |  |  |  |  |  |  |  |  |  |  |  |  |  |
| --- | --- | --- | --- | --- | --- | --- | --- | --- | --- | --- | --- | --- | --- | --- | --- | --- | --- | --- |
| pos | 884 | 1419 | 4331 | 4720 | 7851 | 15324 | 16914 | 19677 | 21123 | 22027 | 22029 | 22030 | 22031 | 22032 | 22033 | 22034 | 29431 |  |
| ref | C | C | C | G | C | C | G | G | G | T | A | G | T | T | C | A | G |  |
| AY.4 | C | T | T | G | T | T | G | G | T | T | - | - | - | - | - | - | T |  |
| AY.4 | C | T | T | G | T | T | G | G | T | G | A | A | A | G | T | G | T | #486061 |
| AY.114 | T | C | C | T | C | C | T | T | G | G | A | A | A | G | T | G | G |  |
| mut |  |  |  |  |  |  |  |  |  |  |  |  |  |  |  |  | G>T |  |

|  |  |  |  |  |  |  |  |  |  |  |  |  |  |  |  |  |  |  |  |  |  |  |  |  |
| --- | --- | --- | --- | --- | --- | --- | --- | --- | --- | --- | --- | --- | --- | --- | --- | --- | --- | --- | --- | --- | --- | --- | --- | --- |
| pos | 884 | 1931 | 4720 | 7360 | 7851 | 12191 | 15380 | 16395 | 16914 | 17040 | 17738 | 19677 | 22027 | 22029 | 22030 | 22031 | 22032 | 22033 | 22034 | 23651 | 25916 | 27750 | 28077 |  |
| ref | C | C | G | G | C | G | G | A | G | T | C | G | T | A | G | T | T | C | A | A | C | G | G |  |
| AY.4 | C | A | G | T | T | T | T | G | G | C | T | G | T | - | - | - | - | - | - | T | T | T | T |  |
| AY.4 | C | A | G | T | T | T | T | G | G | C | T | G | G | A | A | A | G | T | G | T | T | T | T | #486099 |
| AY.114 | T | C | T | G | C | G | G | A | T | T | C | T | G | A | A | A | G | T | G | A | C | G | G |  |
| mut |  |  |  |  |  |  |  |  |  |  |  |  |  |  |  |  |  |  |  | A>T | C>T | G>T | G>T |  |

|  |  |  |  |  |  |  |  |  |  |  |  |  |  |  |
| --- | --- | --- | --- | --- | --- | --- | --- | --- | --- | --- | --- | --- | --- | --- |
| pos | 6042 | 11621 | 16726 | 17010 | 21194 | 22027 | 22029 | 22030 | 22031 | 22032 | 22033 | 22034 | 28156 |  |
| ref | A | T | C | C | A | T | A | G | T | T | C | A | T |  |
| AY.129 | G | C | T | T | G | T | - | - | - | - | - | - | C |  |
| AY.129 | G | C | T | T | G | G | A | A | A | G | T | G | C | #488184 |
| B.1.617.2 | A | T | C | C | A | G | A | A | A | G | T | G | T |  |
| mut |  |  |  |  |  |  |  |  |  |  |  |  | T>C |  |

|  |  |  |  |  |  |  |  |  |  |  |  |  |  |  |  |  |  |  |  |
| --- | --- | --- | --- | --- | --- | --- | --- | --- | --- | --- | --- | --- | --- | --- | --- | --- | --- | --- | --- |
| pos | 246 | 884 | 2272 | 4720 | 7851 | 14185 | 16914 | 19677 | 21742 | 22027 | 22029 | 22030 | 22031 | 22032 | 22033 | 22034 | 25207 | 28817 |  |
| ref | G | C | G | G | C | A | G | G | C | T | A | G | T | T | C | A | C | G |  |
| AY.4 | A | C | A | G | T | G | G | G | T | T | - | - | - | - | - | - | T | T |  |
| AY.4 | A | C | A | G | T | G | G | G | T | G | A | A | A | G | T | G | T | T | #488345 |
| AY.114 | G | T | G | T | C | A | T | T | C | G | A | A | A | G | T | G | C | G |  |
| mut |  |  |  |  |  |  |  |  |  |  |  |  |  |  |  |  | C>T | G>T |  |

|  |  |  |  |  |  |  |  |  |  |  |  |  |  |  |  |  |
| --- | --- | --- | --- | --- | --- | --- | --- | --- | --- | --- | --- | --- | --- | --- | --- | --- |
| pos | 400 | 2058 | 2091 | 4158 | 10537 | 11923 | 15596 | 17040 | 17916 | 20274 | 22101 | 26763 | 27914 | 29367 | 29564 |  |
| ref | T | T | C | C | C | A | A | T | A | G | A | G | A | C | T |  |
| AY.4 | T | T | T | T | C | G | G | C | A | G | A | G | A | C | T |  |
| AY.4 | G | T | T | T | C | G | G | T | G | T | T | T | G | T | C | #492039 |
| AY.4 | T | C | C | C | T | A | A | T | G | T | T | T | G | T | C |  |
| mut | T>G |  |  |  |  |  |  |  |  |  |  |  |  |  |  |  |

|  |  |  |  |  |  |  |  |  |  |  |  |  |  |  |  |  |  |  |  |  |  |  |  |  |  |  |  |  |  |  |  |  |  |  |
| --- | --- | --- | --- | --- | --- | --- | --- | --- | --- | --- | --- | --- | --- | --- | --- | --- | --- | --- | --- | --- | --- | --- | --- | --- | --- | --- | --- | --- | --- | --- | --- | --- | --- | --- |
| pos | 59 | 4181 | 5184 | 5365 | 5584 | 6335 | 6402 | 6408 | 7124 | 7813 | 8986 | 9053 | 9891 | 10029 | 11201 | 11332 | 11418 | 11514 | 13019 | 15861 | 18457 | 19009 | 19220 | 20451 | 21846 | 22227 | 22792 | 24757 | 25003 | 27874 | 27881 | 28916 | 29449 |  |
| ref | C | G | C | C | A | T | C | C | C | T | C | G | C | C | A | A | T | C | C | A | C | G | C | C | C | C | C | G | A | C | C | G | G |  |
| AY.120 | C | T | C | T | A | T | T | T | T | C | T | T | C | T | G | G | T | C | C | A | C | G | T | C | T | C | C | G | G | T | C | T | G |  |
| AY.120 | T | T | C | T | A | T | T | T | T | C | T | T | C | T | G | G | T | C | C | A | C | G | T | C | C | T | T | T | A | C | T | G | T | #493884 |
| AY.9 | C | G | T | C | G | G | C | C | C | T | C | G | T | C | A | A | C | T | T | C | A | T | C | T | C | T | T | T | A | C | C | G | T |  |
| mut | C>T |  |  |  |  |  |  |  |  |  |  |  |  |  |  |  |  |  |  |  |  |  |  |  |  |  |  |  |  |  | C>T |  |  |  |

|  |  |  |  |  |  |  |  |  |  |  |  |  |  |  |  |  |  |  |
| --- | --- | --- | --- | --- | --- | --- | --- | --- | --- | --- | --- | --- | --- | --- | --- | --- | --- | --- |
| pos | 5581 | 7851 | 11365 | 13424 | 14919 | 19170 | 21635 | 22026 | 22027 | 22029 | 22030 | 22031 | 22032 | 22033 | 22034 | 28628 | 29553 |  |
| ref | C | C | G | C | C | C | C | G | T | A | G | T | T | C | A | G | G |  |
| B.1.617.2 | T | C | T | C | C | T | C | G | T | - | - | - | - | - | - | G | G |  |
| B.1.617.2 | T | C | T | C | C | T | T | T | G | A | A | A | G | T | G | T | T | #496367 |
| AY.4 | C | T | G | T | T | C | C | T | G | A | A | A | G | T | G | G | G |  |
| mut |  |  |  |  |  |  | C>T |  |  |  |  |  |  |  |  | G>T | G>T |  |

|  |  |  |  |  |  |  |  |  |  |  |  |  |  |  |  |  |  |  |  |  |  |  |  |  |  |  |  |  |  |  |  |  |  |  |  |  |  |  |  |
| --- | --- | --- | --- | --- | --- | --- | --- | --- | --- | --- | --- | --- | --- | --- | --- | --- | --- | --- | --- | --- | --- | --- | --- | --- | --- | --- | --- | --- | --- | --- | --- | --- | --- | --- | --- | --- | --- | --- | --- |
| pos | 1091 | 1498 | 4181 | 5184 | 5584 | 6337 | 6402 | 7124 | 8349 | 8956 | 8986 | 9053 | 9693 | 9891 | 10029 | 10323 | 11201 | 11332 | 11418 | 11514 | 13019 | 16403 | 17236 | 18177 | 18657 | 19220 | 22227 | 24106 | 24208 | 25352 | 27005 | 27345 | 27632 | 27874 | 27999 | 28915 | 28916 | 29666 |  |
| ref | C | C | G | C | A | G | C | C | G | C | C | G | C | C | C | A | A | A | T | C | C | A | A | T | C | C | C | C | C | G | C | A | G | C | C | C | G | C |  |
| AY.26 | C | T | G | T | G | G | C | C | G | C | C | G | T | T | C | A | A | A | C | T | T | A | A | T | C | C | T | C | C | T | C | T | T | C | C | T | G | C |  |
| AY.103 | C | T | G | T | A | A | T | T | A | T | T | T | C | C | T | G | G | G | T | C | C | G | G | C | T | T | C | T | T | G | T | A | G | T | T | C | T | T | #496620 |
| AY.103 | T | C | T | C | A | A | T | T | A | T | T | T | C | C | T | A | G | G | T | C | C | A | G | C | T | T | C | C | T | G | T | A | G | T | T | C | T | T |  |
| mut |  |  |  |  |  |  |  |  |  |  |  |  |  |  |  | A>G |  |  |  |  |  | A>G |  |  |  |  |  | C>T |  |  |  |  |  |  |  |  |  |  |  |

|  |  |  |  |  |  |  |  |  |  |  |  |  |  |  |  |  |  |  |  |  |  |  |  |  |  |  |  |  |  |  |  |  |  |  |  |
| --- | --- | --- | --- | --- | --- | --- | --- | --- | --- | --- | --- | --- | --- | --- | --- | --- | --- | --- | --- | --- | --- | --- | --- | --- | --- | --- | --- | --- | --- | --- | --- | --- | --- | --- | --- |
| pos | 454 | 710 | 1729 | 4181 | 5184 | 5584 | 6402 | 7124 | 7851 | 8986 | 9053 | 9088 | 9891 | 10029 | 11201 | 11332 | 11418 | 11514 | 13019 | 14030 | 15714 | 18086 | 18143 | 19220 | 20679 | 21846 | 22227 | 22792 | 26456 | 26885 | 27874 | 28098 | 28528 | 28916 |  |
| ref | A | C | G | G | C | A | C | C | C | C | G | T | C | C | A | A | T | C | C | G | C | C | C | C | G | C | C | C | C | C | C | T | C | G |  |
| AY.4 | A | T | G | T | C | A | T | T | T | T | T | T | C | T | G | G | T | C | C | G | T | C | A | T | A | T | C | C | T | T | T | T | C | T |  |
| AY.9 | A | T | G | T | C | A | T | T | T | T | T | C | C | T | A | A | C | T | T | A | C | T | C | C | G | C | T | T | C | C | C | C | T | G | #499847 |
| AY.9.2 | G | C | A | G | T | G | C | C | C | C | G | T | T | C | A | A | C | T | T | A | C | T | C | C | G | C | T | T | C | C | C | T | C | G |  |
| mut |  |  |  |  |  |  |  |  |  |  |  | T>C |  |  |  |  |  |  |  |  |  |  |  |  |  |  |  |  |  |  |  | T>C | C>T |  |  |

|  |  |  |  |  |  |  |  |  |  |  |  |  |  |  |  |  |  |  |  |  |  |  |  |  |  |  |  |  |  |  |  |  |  |
| --- | --- | --- | --- | --- | --- | --- | --- | --- | --- | --- | --- | --- | --- | --- | --- | --- | --- | --- | --- | --- | --- | --- | --- | --- | --- | --- | --- | --- | --- | --- | --- | --- | --- |
| pos | 203 | 1684 | 4181 | 5164 | 5184 | 5584 | 6402 | 7124 | 9251 | 9891 | 10029 | 10255 | 11201 | 11332 | 11418 | 11514 | 12049 | 13019 | 15026 | 19220 | 21405 | 22227 | 22427 | 22998 | 25931 | 26720 | 26801 | 27874 | 27990 | 28326 | 28735 | 28916 |  |
| ref | C | C | G | G | C | A | C | C | G | C | C | G | A | A | T | C | C | C | C | C | A | C | G | C | C | G | C | C | G | G | T | G |  |
| AY.90 | C | C | T | G | C | A | T | T | A | C | T | G | G | G | T | C | C | C | T | T | A | C | G | C | T | A | C | T | T | G | C | T |  |
| AY.47 | T | C | T | G | C | A | T | T | G | T | C | A | A | A | C | T | T | T | C | C | G | T | A | T | C | G | T | C | G | T | T | G | #499869 |
| AY.47 | C | T | G | T | T | G | C | C | G | T | C | G | A | A | C | T | T | T | C | C | G | T | A | T | C | G | T | C | G | T | T | G |  |
| mut | C>T |  |  |  |  |  |  |  |  |  |  | G>A |  |  |  |  |  |  |  |  |  |  |  |  |  |  |  |  |  |  |  |  |  |

|  |  |  |  |  |  |  |  |  |  |  |  |  |  |  |  |  |  |  |  |  |
| --- | --- | --- | --- | --- | --- | --- | --- | --- | --- | --- | --- | --- | --- | --- | --- | --- | --- | --- | --- | --- |
| pos | 421 | 884 | 2644 | 3251 | 4720 | 7851 | 12250 | 15933 | 16914 | 18713 | 19677 | 22027 | 22029 | 22030 | 22031 | 22032 | 22033 | 22034 | 29085 |  |
| ref | C | C | C | G | G | C | A | C | G | C | G | T | A | G | T | T | C | A | C |  |
| AY.4 | T | C | C | T | G | T | G | T | G | T | G | T | - | - | - | - | - | - | C |  |
| AY.4 | T | C | T | T | G | T | G | T | G | T | G | G | A | A | A | G | T | G | T | #501923 |
| AY.114 | C | T | C | G | T | C | A | C | T | C | T | G | A | A | A | G | T | G | C |  |
| mut |  |  | C>T |  |  |  |  |  |  |  |  |  |  |  |  |  |  |  | C>T |  |

|  |  |  |  |  |  |  |  |  |  |  |  |  |  |  |  |  |  |
| --- | --- | --- | --- | --- | --- | --- | --- | --- | --- | --- | --- | --- | --- | --- | --- | --- | --- |
| pos | 3243 | 10615 | 14407 | 15628 | 17268 | 18079 | 20532 | 20762 | 22027 | 22029 | 22030 | 22031 | 22032 | 22033 | 22034 | 22468 |  |
| ref | G | C | C | T | T | G | A | C | T | A | G | T | T | C | A | G |  |
| B.1.617.2 | G | T | T | T | C | G | A | T | T | - | - | - | - | - | - | T |  |
| B.1.617.2 | G | T | T | T | C | G | A | T | G | A | A | A | G | T | G | T | #501941 |
| AY.36 | A | C | C | C | T | T | G | C | G | A | A | A | G | T | G | G |  |
| mut |  |  |  |  |  |  |  |  |  |  |  |  |  |  |  | G>T |  |

|  |  |  |  |  |  |  |  |  |  |  |  |  |  |  |  |  |  |  |  |  |  |  |  |  |  |  |  |  |  |  |  |  |  |  |  |  |  |  |  |  |  |  |  |  |  |  |  |  |  |  |  |  |  |  |  |  |  |  |  |  |  |  |  |  |  |  |  |  |  |  |  |  |  |  |  |  |  |  |  |
| --- | --- | --- | --- | --- | --- | --- | --- | --- | --- | --- | --- | --- | --- | --- | --- | --- | --- | --- | --- | --- | --- | --- | --- | --- | --- | --- | --- | --- | --- | --- | --- | --- | --- | --- | --- | --- | --- | --- | --- | --- | --- | --- | --- | --- | --- | --- | --- | --- | --- | --- | --- | --- | --- | --- | --- | --- | --- | --- | --- | --- | --- | --- | --- | --- | --- | --- | --- | --- | --- | --- | --- | --- | --- | --- | --- | --- | --- | --- | --- |
| pos | 210 | 1048 | 3304 | 3852 | 3961 | 4181 | 4255 | 6070 | 6402 | 7124 | 7798 | 8986 | 9053 | 9072 | 9891 | 10029 | 10116 | 11201 | 11288 | 11289 | 11290 | 11291 | 11292 | 11293 | 11294 | 11295 | 11296 | 11332 | 15451 | 16466 | 18795 | 19170 | 19220 | 19299 | 20578 | 21618 | 21649 | 21846 | 21987 | 21991 | 21992 | 21993 | 22029 | 22030 | 22031 | 22032 | 22033 | 22034 | 22917 | 22995 | 23012 | 23287 | 23604 | 23948 | 24382 | 24410 | 25276 | 25360 | 25469 | 27638 | 27752 | 27874 | 28209 | 28248 | 28249 | 28250 | 28251 | 28252 | 28253 | 28271 | 28461 | 28881 | 28882 | 28883 | 28916 | 29402 | 29742 | 29769 |  |
| ref | G | G | G | A | C | G | G | C | C | C | G | C | G | C | C | C | C | A | T | C | T | G | G | T | T | T | T | A | G | C | C | C | C | T | G | C | T | C | G | T | T | A | A | G | T | T | C | A | T | C | G | T | C | G | C | G | C | A | C | T | C | C | G | G | A | T | T | T | C | A | A | G | G | G | G | G | G | C |  |
| B.1.1.318 | G | G | G | T | T | G | G | C | C | C | T | C | G | T | T | C | T | A | - | - | - | - | - | - | - | - | - | A | G | C | C | C | C | T | A | C | T | T | G | - | - | - | A | G | T | T | C | A | T | C | A | C | A | C | T | G | A | A | C | T | C | C | T | G | A | T | T | T | C | G | A | A | A | C | G | G | G | T |  |
| B.1.617.2 | G | G | G | T | T | G | A | T | C | C | T | T | T | C | C | T | C | G | T | C | T | G | G | T | T | T | T | G | A | T | T | T | T | C | G | G | C | C | A | T | T | A | - | - | - | - | - | - | G | A | G | T | G | G | C | A | C | G | T | C | T | T | G | - | - | - | - | - | - | - | G | T | G | G | T | T | T | C | #503891 |
| AY.122 | T | T | T | A | C | T | G | C | T | T | G | T | T | C | C | T | C | G | T | C | T | G | G | T | T | T | T | G | A | T | T | T | T | C | G | G | C | C | A | T | T | A | - | - | - | - | - | - | G | A | G | T | G | G | C | A | C | A | T | C | T | T | G | - | - | - | - | - | - | - | G | T | G | G | T | T | T | C |  |
| mut |  |  |  |  |  |  | G>A | C>T |  |  |  |  |  |  |  |  |  |  |  |  |  |  |  |  |  |  |  |  |  |  |  |  |  |  |  |  |  |  |  |  |  |  |  |  |  |  |  |  |  |  |  |  |  |  |  |  |  | A>G |  |  |  |  |  |  |  |  |  |  |  |  |  |  |  |  |  |  |  |  |  |

|  |  |  |  |  |  |  |  |  |  |  |  |  |  |  |
| --- | --- | --- | --- | --- | --- | --- | --- | --- | --- | --- | --- | --- | --- | --- |
| pos | 526 | 2564 | 3948 | 8634 | 11562 | 19524 | 22127 | 26107 | 27507 | 27875 | 27881 | 27882 | 27883 |  |
| ref | G | G | A | T | G | C | C | G | A | T | C | G | C |  |
| AY.25.1 | G | A | G | C | T | C | T | C | C | T | C | G | C |  |
| AY.25.1 | G | A | G | C | T | C | T | C | C | C | T | C | T | #505415 |
| AY.98 | T | G | A | T | G | T | C | G | A | C | T | C | T |  |
| mut |  |  |  |  |  |  |  |  |  |  |  |  |  |  |

|  |  |  |  |  |  |  |  |  |  |  |  |
| --- | --- | --- | --- | --- | --- | --- | --- | --- | --- | --- | --- |
| pos | 3004 | 3948 | 4890 | 11562 | 17236 | 24208 | 26107 | 27507 | 28035 | 29118 |  |
| ref | G | A | C | G | A | C | G | A | A | C |  |
| AY.103 | A | A | C | G | G | T | G | A | A | C |  |
| AY.25 | A | A | C | G | G | C | C | C | G | T | #505953 |
| AY.25.1 | G | G | T | T | A | C | C | C | G | T |  |
| mut |  |  |  |  |  |  |  |  |  |  |  |

|  |  |  |  |  |  |  |  |  |  |  |  |  |  |  |
| --- | --- | --- | --- | --- | --- | --- | --- | --- | --- | --- | --- | --- | --- | --- |
| pos | 7600 | 12274 | 18807 | 20578 | 25452 | 25734 | 27572 | 27573 | 27575 | 27576 | 29700 | 29779 | 29781 |  |
| ref | C | G | C | G | C | C | G | C | C | T | A | G | G |  |
| AY.120.1 | C | G | C | T | C | T | G | C | C | T | G | - | - |  |
| AY.120.1 | C | A | T | T | C | T | A | T | A | C | G | G | G | #510915 |
| AY.120.1 | T | G | C | G | T | C | A | T | A | C | A | G | G |  |
| mut |  | G>A | C>T |  |  |  |  |  |  |  | A>G |  |  |  |

|  |  |  |  |  |  |  |  |  |  |  |  |  |  |  |  |  |  |  |  |  |  |  |
| --- | --- | --- | --- | --- | --- | --- | --- | --- | --- | --- | --- | --- | --- | --- | --- | --- | --- | --- | --- | --- | --- | --- |
| pos | 186 | 357 | 884 | 4720 | 7851 | 15277 | 16914 | 17040 | 17403 | 17427 | 19677 | 21845 | 22027 | 22029 | 22030 | 22031 | 22032 | 22033 | 22034 | 27506 | 29445 |  |
| ref | C | T | C | G | C | C | G | T | C | G | G | A | T | A | G | T | T | C | A | G | C |  |
| AY.4 | T | T | C | G | T | T | G | C | T | T | G | G | T | - | - | - | - | - | - | G | C |  |
| AY.4 | T | C | C | G | T | T | G | C | T | T | G | G | G | A | A | A | G | T | G | T | T | #512471 |
| AY.114 | C | T | T | T | C | C | T | T | C | G | T | A | G | A | A | A | G | T | G | G | C |  |
| mut |  | T>C |  |  |  |  |  |  |  |  |  |  |  |  |  |  |  |  |  | G>T | C>T |  |

|  |  |  |  |  |  |  |  |  |  |  |  |  |  |  |  |  |  |  |  |  |  |  |  |  |  |  |  |  |  |  |  |  |  |  |  |  |  |
| --- | --- | --- | --- | --- | --- | --- | --- | --- | --- | --- | --- | --- | --- | --- | --- | --- | --- | --- | --- | --- | --- | --- | --- | --- | --- | --- | --- | --- | --- | --- | --- | --- | --- | --- | --- | --- | --- |
| pos | 1684 | 2445 | 4181 | 4891 | 5164 | 5184 | 5368 | 5584 | 6402 | 7124 | 8986 | 9053 | 9891 | 10029 | 11201 | 11332 | 11418 | 11514 | 12049 | 12466 | 12801 | 13019 | 14004 | 17550 | 19220 | 21240 | 22227 | 22427 | 22801 | 26107 | 27507 | 27874 | 28326 | 28807 | 28916 | 28980 |  |
| ref | C | C | G | C | G | C | A | A | C | C | C | G | C | C | A | A | T | C | C | A | G | C | A | C | C | T | C | G | G | G | A | C | G | C | G | G |  |
| AY.25 | C | C | T | C | G | C | A | A | T | T | T | T | C | T | G | G | T | C | C | C | A | C | G | T | T | T | C | G | T | C | C | T | G | C | T | G |  |
| AY.25 | C | C | T | C | G | C | A | A | T | T | T | T | C | T | G | G | T | C | C | C | A | C | G | T | T | T | C | G | T | G | A | C | T | T | G | T | #513122 |
| AY.47 | T | T | G | T | T | T | G | G | C | C | C | G | T | C | A | A | C | T | T | A | G | T | A | C | C | C | T | A | G | G | A | C | T | T | G | T |  |
| mut |  |  |  |  |  |  |  |  |  |  |  |  |  |  |  |  |  |  |  |  |  |  |  |  |  |  |  |  |  |  |  |  |  |  |  |  |  |

|  |  |  |  |  |  |  |  |  |  |  |  |  |  |  |  |  |  |  |  |
| --- | --- | --- | --- | --- | --- | --- | --- | --- | --- | --- | --- | --- | --- | --- | --- | --- | --- | --- | --- |
| pos | 884 | 4720 | 7851 | 15277 | 16471 | 16914 | 17040 | 17427 | 19677 | 21342 | 22027 | 22029 | 22030 | 22031 | 22032 | 22033 | 22034 | 24583 |  |
| ref | C | G | C | C | A | G | T | G | G | C | T | A | G | T | T | C | A | T |  |
| AY.4 | C | G | T | T | G | G | C | T | G | T | T | - | - | - | - | - | - | C |  |
| AY.4 | C | G | T | T | G | G | C | T | G | T | G | A | A | A | G | T | G | C | #515468 |
| AY.114 | T | T | C | C | A | T | T | G | T | C | G | A | A | A | G | T | G | T |  |
| mut |  |  |  |  |  |  |  |  |  |  |  |  |  |  |  |  |  | T>C |  |

|  |  |  |  |  |  |  |  |  |  |  |  |  |  |  |  |
| --- | --- | --- | --- | --- | --- | --- | --- | --- | --- | --- | --- | --- | --- | --- | --- |
| pos | 15063 | 18803 | 21987 | 22027 | 22029 | 22030 | 22031 | 22032 | 22033 | 22034 | 23683 | 25785 | 27654 | 29040 |  |
| ref | A | G | G | T | A | G | T | T | C | A | C | G | C | A |  |
| AY.4 | A | G | G | G | A | A | A | G | T | G | C | G | C | A |  |
| AY.4 | G | T | G | G | A | A | A | G | T | G | T | C | T | G | #515481 |
| AY.4 | G | T | A | T | - | - | - | - | - | - | T | C | T | G |  |
| mut | A>G | G>T |  |  |  |  |  |  |  |  |  |  |  |  |  |

|  |  |  |  |  |  |  |  |  |  |  |  |  |  |  |  |  |  |  |  |  |  |  |  |
| --- | --- | --- | --- | --- | --- | --- | --- | --- | --- | --- | --- | --- | --- | --- | --- | --- | --- | --- | --- | --- | --- | --- | --- |
| pos | 526 | 2518 | 2564 | 4816 | 5437 | 7768 | 9118 | 12451 | 14724 | 17236 | 18360 | 19117 | 19524 | 21525 | 22314 | 22329 | 24208 | 24236 | 25702 | 26153 | 28086 | 28895 |  |
| ref | G | G | G | T | G | C | C | A | C | A | A | G | C | T | C | C | C | G | C | G | G | G |  |
| AY.103 | G | G | A | T | G | C | C | A | C | G | G | G | C | T | C | C | T | G | T | T | G | T |  |
| B.1.617.2 | G | G | A | T | G | C | C | A | C | G | G | T | C | C | C | T | C | T | C | G | T | G | #516785 |
| AY.98.1 | T | T | G | C | T | T | T | T | T | A | A | G | T | T | T | C | C | T | C | G | T | G |  |
| mut |  |  |  |  |  |  |  |  |  |  |  | G>T |  | T>C |  | C>T |  |  |  |  |  |  |  |

|  |  |  |  |  |  |  |  |  |  |  |  |  |  |  |
| --- | --- | --- | --- | --- | --- | --- | --- | --- | --- | --- | --- | --- | --- | --- |
| pos | 503 | 829 | 4878 | 10834 | 22026 | 22027 | 22029 | 22030 | 22031 | 22032 | 22033 | 22034 | 28208 |  |
| ref | C | C | C | C | G | T | A | G | T | T | C | A | T |  |
| AY.39 | T | T | T | T | G | T | - | - | - | - | - | - | C |  |
| AY.39 | T | T | T | T | T | G | A | A | A | G | T | G | C | #517130 |
| AY.39 | C | C | C | C | T | G | A | A | A | G | T | G | T |  |
| mut |  |  |  |  |  |  |  |  |  |  |  |  | T>C |  |

|  |  |  |  |  |  |  |  |  |  |  |  |  |  |  |
| --- | --- | --- | --- | --- | --- | --- | --- | --- | --- | --- | --- | --- | --- | --- |
| pos | 823 | 2756 | 6408 | 8208 | 9593 | 10317 | 16075 | 18395 | 21637 | 22199 | 22224 | 24812 | 27610 |  |
| ref | C | A | C | C | T | A | G | C | C | G | C | G | C |  |
| AY.4 | C | A | T | T | T | A | G | C | C | G | C | G | C |  |
| AY.4 | C | A | T | T | G | A | A | T | T | T | T | T | T | #518807 |
| AY.4 | T | G | C | C | T | G | A | T | T | G | T | G | T |  |
| mut |  |  |  |  | T>G |  |  |  |  | G>T |  | G>T |  |  |

|  |  |  |  |  |  |  |  |  |  |  |  |  |  |
| --- | --- | --- | --- | --- | --- | --- | --- | --- | --- | --- | --- | --- | --- |
| pos | 22027 | 22029 | 22030 | 22031 | 22032 | 22033 | 22034 | 23140 | 24780 | 24958 | 24959 | 28032 |  |
| ref | T | A | G | T | T | C | A | A | A | T | G | A |  |
| AY.4 | G | A | A | A | G | T | G | A | A | T | G | A |  |
| AY.4 | G | A | A | A | G | T | G | G | T | C | A | G | #519605 |
| AY.4 | T | - | - | - | - | - | - | A | T | C | A | G |  |
| mut |  |  |  |  |  |  |  | A>G |  |  |  |  |  |

|  |  |  |  |  |  |  |  |  |  |  |  |  |  |  |  |  |  |  |  |  |
| --- | --- | --- | --- | --- | --- | --- | --- | --- | --- | --- | --- | --- | --- | --- | --- | --- | --- | --- | --- | --- |
| pos | 884 | 4504 | 4720 | 9165 | 10323 | 12772 | 12828 | 15240 | 16914 | 18444 | 19677 | 21707 | 22027 | 22029 | 22030 | 22031 | 22032 | 22033 | 22034 |  |
| ref | C | G | G | C | A | A | T | C | G | A | G | C | T | A | G | T | T | C | A |  |
| AY.113 | C | T | G | T | G | G | C | T | G | T | G | T | T | - | - | - | - | - | - |  |
| AY.113 | C | T | G | T | G | G | C | T | G | T | G | T | G | A | A | A | G | T | G | #521854 |
| AY.114 | T | G | T | C | A | A | T | C | T | A | T | C | G | A | A | A | G | T | G |  |
| mut |  |  |  |  |  |  |  |  |  |  |  |  |  |  |  |  |  |  |  |  |

|  |  |  |  |  |  |  |  |  |  |  |  |  |  |  |  |  |  |
| --- | --- | --- | --- | --- | --- | --- | --- | --- | --- | --- | --- | --- | --- | --- | --- | --- | --- |
| pos | 1588 | 2632 | 3476 | 5541 | 6230 | 6726 | 9496 | 15654 | 18258 | 22027 | 22029 | 22030 | 22031 | 22032 | 22033 | 22034 |  |
| ref | A | G | G | A | C | C | A | C | T | T | A | G | T | T | C | A |  |
| AY.39 | A | G | G | T | T | C | G | C | C | T | - | - | - | - | - | - |  |
| AY.39 | A | G | G | T | T | C | G | C | C | G | A | A | A | G | T | G | #521941 |
| AY.39 | G | T | A | A | C | T | A | T | T | G | A | A | A | G | T | G |  |
| mut |  |  |  |  |  |  |  |  |  |  |  |  |  |  |  |  |  |

|  |  |  |  |  |  |  |  |  |  |  |  |  |  |  |  |
| --- | --- | --- | --- | --- | --- | --- | --- | --- | --- | --- | --- | --- | --- | --- | --- |
| pos | 3476 | 6408 | 7459 | 7851 | 8473 | 13944 | 15277 | 17040 | 17427 | 25003 | 26488 | 26491 | 26492 | 26497 |  |
| ref | G | C | G | C | T | C | C | T | G | A | T | T | A | T |  |
| AY.4 | A | C | T | T | T | C | T | C | T | A | T | T | A | T |  |
| AY.4 | A | C | T | T | T | C | T | C | T | A | G | C | T | C | #524189 |
| AY.120 | G | T | G | C | C | T | C | T | G | G | G | C | T | C |  |
| mut |  |  |  |  |  |  |  |  |  |  |  |  |  |  |  |

|  |  |  |  |  |  |  |  |  |  |  |  |  |  |  |  |  |  |  |  |
| --- | --- | --- | --- | --- | --- | --- | --- | --- | --- | --- | --- | --- | --- | --- | --- | --- | --- | --- | --- |
| pos | 2396 | 3318 | 3967 | 4282 | 9857 | 10271 | 20569 | 22026 | 22027 | 22029 | 22030 | 22031 | 22032 | 22033 | 22034 | 24979 | 25983 | 28236 |  |
| ref | A | C | T | A | C | G | G | G | T | A | G | T | T | C | A | T | A | C |  |
| AY.118 | G | T | G | G | T | G | A | G | T | - | - | - | - | - | - | C | G | C |  |
| AY.118 | G | T | G | G | T | G | A | T | G | A | A | A | G | T | G | C | G | T | #526178 |
| AY.118 | A | C | T | A | C | A | G | T | G | A | A | A | G | T | G | T | A | C |  |
| mut |  |  |  |  |  |  |  |  |  |  |  |  |  |  |  | T>C | A>G | C>T |  |

|  |  |  |  |  |  |  |  |  |  |  |  |  |  |  |  |  |  |  |
| --- | --- | --- | --- | --- | --- | --- | --- | --- | --- | --- | --- | --- | --- | --- | --- | --- | --- | --- |
| pos | 2066 | 4197 | 4300 | 11596 | 16726 | 19816 | 19962 | 21137 | 22026 | 22027 | 22029 | 22030 | 22031 | 22032 | 22033 | 22034 | 24848 |  |
| ref | A | A | G | G | C | G | G | A | G | T | A | G | T | T | C | A | G |  |
| AY.100 | G | C | T | T | T | T | T | G | G | T | - | - | - | - | - | - | A |  |
| AY.100 | G | C | T | T | T | T | T | G | T | G | A | A | A | G | T | G | A | #526313 |
| AY.100 | A | A | G | G | C | G | G | A | T | G | A | A | A | G | T | G | G |  |
| mut |  |  |  |  |  |  |  |  |  |  |  |  |  |  |  |  | G>A |  |

|  |  |  |  |  |  |  |  |  |  |  |  |  |  |  |  |
| --- | --- | --- | --- | --- | --- | --- | --- | --- | --- | --- | --- | --- | --- | --- | --- |
| pos | 1514 | 3433 | 10423 | 11932 | 19338 | 22026 | 22027 | 22029 | 22030 | 22031 | 22032 | 22033 | 22034 | 24622 |  |
| ref | C | G | T | T | A | G | T | A | G | T | T | C | A | T |  |
| AY.39 | T | T | C | C | G | G | T | - | - | - | - | - | - | C |  |
| AY.39 | T | T | C | C | G | T | G | A | A | A | G | T | G | C | #526329 |
| AY.39 | C | G | T | T | A | T | G | A | A | A | G | T | G | T |  |
| mut |  |  |  |  |  |  |  |  |  |  |  |  |  | T>C |  |

|  |  |  |  |  |  |  |  |  |  |  |  |  |  |  |  |  |  |  |  |  |
| --- | --- | --- | --- | --- | --- | --- | --- | --- | --- | --- | --- | --- | --- | --- | --- | --- | --- | --- | --- | --- |
| pos | 221 | 3243 | 4229 | 5339 | 9049 | 14055 | 16734 | 20431 | 21077 | 22027 | 22029 | 22030 | 22031 | 22032 | 22033 | 22034 | 26184 | 27613 | 29781 |  |
| ref | T | G | A | C | G | G | A | A | C | T | A | G | T | T | C | A | T | G | G |  |
| AY.36 | T | G | G | C | T | T | G | G | T | T | - | - | - | - | - | - | C | A | T |  |
| AY.36 | C | G | G | T | T | T | G | A | C | G | A | A | A | G | T | G | T | A | T | #528720 |
| AY.36 | T | A | A | C | G | G | A | A | C | G | A | A | A | G | T | G | T | G | G |  |
| mut | T>C |  |  | C>T |  |  |  |  |  |  |  |  |  |  |  |  |  | G>A | G>T |  |

|  |  |  |  |  |  |  |  |  |  |  |  |  |  |  |  |  |  |  |  |
| --- | --- | --- | --- | --- | --- | --- | --- | --- | --- | --- | --- | --- | --- | --- | --- | --- | --- | --- | --- |
| pos | 440 | 884 | 3183 | 4237 | 4720 | 7851 | 10755 | 16914 | 17040 | 17460 | 19677 | 22027 | 22029 | 22030 | 22031 | 22032 | 22033 | 22034 |  |
| ref | G | C | A | T | G | C | C | G | T | A | G | T | A | G | T | T | C | A |  |
| AY.4 | T | C | G | C | G | T | T | G | C | G | G | T | - | - | - | - | - | - |  |
| AY.4 | T | C | G | C | G | T | T | G | C | G | G | G | A | A | A | G | T | G | #528794 |
| AY.114 | G | T | A | T | T | C | C | T | T | A | T | G | A | A | A | G | T | G |  |
| mut |  |  |  |  |  |  |  |  |  |  |  |  |  |  |  |  |  |  |  |

|  |  |  |  |  |  |  |  |  |  |  |  |  |  |
| --- | --- | --- | --- | --- | --- | --- | --- | --- | --- | --- | --- | --- | --- |
| pos | 6042 | 11621 | 16206 | 17010 | 22027 | 22029 | 22030 | 22031 | 22032 | 22033 | 22034 | 25681 |  |
| ref | A | T | G | C | T | A | G | T | T | C | A | G |  |
| AY.129 | A | T | T | C | T | - | - | - | - | - | - | A |  |
| AY.129 | A | T | T | C | G | A | A | A | G | T | G | A | #532936 |
| AY.129 | G | C | G | T | G | A | A | A | G | T | G | G |  |
| mut |  |  |  |  |  |  |  |  |  |  |  | G>A |  |

|  |  |  |  |  |  |  |  |  |  |  |  |  |  |
| --- | --- | --- | --- | --- | --- | --- | --- | --- | --- | --- | --- | --- | --- |
| pos | 1116 | 22027 | 22029 | 22030 | 22031 | 22032 | 22033 | 22034 | 22346 | 25513 | 25617 | 26972 |  |
| ref | T | T | A | G | T | T | C | A | G | C | G | T |  |
| AY.126 | T | G | A | A | A | G | T | G | G | C | G | T |  |
| AY.126 | G | G | A | A | A | G | T | G | T | T | T | C | #532940 |
| AY.126 | G | T | - | - | - | - | - | - | T | T | T | C |  |
| mut | T>G |  |  |  |  |  |  |  |  |  |  |  |  |

|  |  |  |  |  |  |  |  |  |  |  |  |  |  |  |  |  |  |  |  |  |  |  |  |
| --- | --- | --- | --- | --- | --- | --- | --- | --- | --- | --- | --- | --- | --- | --- | --- | --- | --- | --- | --- | --- | --- | --- | --- |
| pos | 1588 | 2231 | 2632 | 2946 | 3304 | 3433 | 3476 | 6726 | 8655 | 8849 | 13929 | 15654 | 18186 | 22027 | 22029 | 22030 | 22031 | 22032 | 22033 | 22034 | 25614 | 29521 |  |
| ref | A | G | G | G | G | G | G | C | C | G | T | C | G | T | A | G | T | T | C | A | C | A |  |
| AY.39 | A | T | G | A | T | T | G | C | T | A | C | C | G | T | - | - | - | - | - | - | C | A |  |
| AY.39 | A | T | G | A | T | T | G | C | T | A | C | C | T | G | A | A | A | G | T | G | T | T | #534002 |
| AY.39 | G | G | T | G | G | G | A | T | C | G | T | T | G | G | A | A | A | G | T | G | C | A |  |
| mut |  |  |  |  |  |  |  |  |  |  |  |  | G>T |  |  |  |  |  |  |  | C>T | A>T |  |

|  |  |  |  |  |  |  |  |  |  |  |  |  |  |  |  |
| --- | --- | --- | --- | --- | --- | --- | --- | --- | --- | --- | --- | --- | --- | --- | --- |
| pos | 2910 | 10844 | 10993 | 15629 | 17040 | 19596 | 20176 | 22000 | 22993 | 25316 | 25503 | 25720 | 29706 | 29734 |  |
| ref | C | A | C | G | T | C | G | C | C | T | A | G | G | G |  |
| AY.4 | T | G | C | A | T | T | T | T | C | C | G | G | A | G |  |
| AY.4 | T | G | C | A | T | T | T | T | T | T | A | A | G | T | #536330 |
| AY.4 | C | A | T | G | C | C | G | C | C | T | A | A | G | T |  |
| mut |  |  |  |  |  |  |  |  | C>T |  |  |  |  |  |  |

|  |  |  |  |  |  |  |  |  |  |  |  |  |  |  |  |
| --- | --- | --- | --- | --- | --- | --- | --- | --- | --- | --- | --- | --- | --- | --- | --- |
| pos | 6633 | 13792 | 17678 | 22027 | 22029 | 22030 | 22031 | 22032 | 22033 | 22034 | 23939 | 24844 | 27643 | 27681 |  |
| ref | C | C | C | T | A | G | T | T | C | A | C | C | C | T |  |
| AY.4 | C | C | C | G | A | A | A | G | T | G | C | C | C | T |  |
| AY.4 | T | T | T | G | A | A | A | G | T | G | A | A | T | C | #536856 |
| AY.4 | T | C | T | T | - | - | - | - | - | - | A | A | T | C |  |
| mut | C>T | C>T | C>T |  |  |  |  |  |  |  |  |  |  |  |  |

|  |  |  |  |  |  |  |  |  |  |  |  |  |  |
| --- | --- | --- | --- | --- | --- | --- | --- | --- | --- | --- | --- | --- | --- |
| pos | 4597 | 11165 | 12210 | 18462 | 22027 | 22029 | 22030 | 22031 | 22032 | 22033 | 22034 | 25726 |  |
| ref | T | G | A | G | T | A | G | T | T | C | A | G |  |
| AY.119.2 | C | T | G | A | T | - | - | - | - | - | - | G |  |
| AY.119.2 | C | T | G | A | G | A | A | A | G | T | G | T | #537763 |
| AY.119.2 | T | G | A | G | G | A | A | A | G | T | G | G |  |
| mut |  |  |  |  |  |  |  |  |  |  |  | G>T |  |

|  |  |  |  |  |  |  |  |  |  |  |  |  |  |  |  |  |  |  |
| --- | --- | --- | --- | --- | --- | --- | --- | --- | --- | --- | --- | --- | --- | --- | --- | --- | --- | --- |
| pos | 171 | 1628 | 3559 | 5298 | 6638 | 9979 | 15654 | 19527 | 22026 | 22027 | 22029 | 22030 | 22031 | 22032 | 22033 | 22034 | 29779 |  |
| ref | C | C | A | C | C | C | C | T | G | T | A | G | T | T | C | A | G |  |
| AY.39 | T | C | C | T | G | T | T | C | G | T | - | - | - | - | - | - | T |  |
| AY.39 | T | A | C | T | G | T | T | C | G | T | A | A | A | G | T | G | T | #537824 |
| AY.39 | C | C | A | C | C | C | C | T | T | G | A | A | A | G | T | G | G |  |
| mut |  | C>A |  |  |  |  |  |  |  |  |  |  |  |  |  |  | G>T |  |

|  |  |  |  |  |  |  |  |  |  |  |  |  |  |  |  |  |
| --- | --- | --- | --- | --- | --- | --- | --- | --- | --- | --- | --- | --- | --- | --- | --- | --- |
| pos | 7143 | 13459 | 13965 | 16428 | 20408 | 22026 | 22027 | 22029 | 22030 | 22031 | 22032 | 22033 | 22034 | 28646 | 29741 |  |
| ref | G | G | C | C | T | G | T | A | G | T | T | C | A | G | C |  |
| AY.100 | T | A | T | C | G | G | T | - | - | - | - | - | - | G | A |  |
| AY.100 | T | A | T | T | G | G | G | A | A | A | G | T | G | T | A | #537826 |
| AY.100 | G | G | C | C | T | T | G | A | A | A | G | T | G | G | C |  |
| mut |  |  |  | C>T |  |  |  |  |  |  |  |  |  | G>T | C>A |  |

|  |  |  |  |  |  |  |  |  |  |  |  |
| --- | --- | --- | --- | --- | --- | --- | --- | --- | --- | --- | --- |
| pos | 355 | 691 | 1043 | 3784 | 6473 | 14094 | 16929 | 18739 | 25386 | 26060 |  |
| ref | C | A | G | C | G | T | A | G | C | C |  |
| AY.4 | T | A | G | T | G | T | A | T | C | C |  |
| AY.4 | T | A | G | T | A | C | G | G | T | T | #540037 |
| AY.4 | C | G | A | C | A | T | G | G | T | T |  |
| mut |  |  |  |  |  | T>C |  |  |  |  |  |

|  |  |  |  |  |  |  |  |  |  |  |  |  |  |  |  |  |
| --- | --- | --- | --- | --- | --- | --- | --- | --- | --- | --- | --- | --- | --- | --- | --- | --- |
| pos | 4084 | 5668 | 6286 | 8985 | 9360 | 17913 | 22027 | 22029 | 22030 | 22031 | 22032 | 22033 | 22034 | 28245 | 28254 |  |
| ref | C | G | C | A | C | C | T | A | G | T | T | C | A | T | A |  |
| AY.117 | T | G | T | A | T | C | T | - | - | - | - | - | - | C | - |  |
| AY.117 | T | G | T | A | T | C | G | A | A | A | G | T | G | T | A | #541943 |
| AY.117 | C | A | C | G | C | T | G | A | A | A | G | T | G | T | A |  |
| mut |  |  |  |  |  |  |  |  |  |  |  |  |  |  |  |  |

|  |  |  |  |  |  |  |  |  |  |  |  |  |  |  |  |  |  |  |  |  |  |
| --- | --- | --- | --- | --- | --- | --- | --- | --- | --- | --- | --- | --- | --- | --- | --- | --- | --- | --- | --- | --- | --- |
| pos | 606 | 884 | 4720 | 4891 | 7851 | 8681 | 9180 | 10156 | 16733 | 16914 | 19677 | 22027 | 22029 | 22030 | 22031 | 22032 | 22033 | 22034 | 23608 | 28922 |  |
| ref | T | C | G | C | C | A | C | C | C | G | G | T | A | G | T | T | C | A | G | G |  |
| AY.4 | C | C | G | T | T | G | T | T | T | G | G | T | - | - | - | - | - | - | T | G |  |
| AY.4 | C | C | G | T | T | G | T | T | T | G | G | G | A | A | A | G | T | G | T | T | #544166 |
| AY.114 | T | T | T | C | C | A | C | C | C | T | T | G | A | A | A | G | T | G | G | G |  |
| mut |  |  |  |  |  |  |  |  |  |  |  |  |  |  |  |  |  |  | G>T | G>T |  |

|  |  |  |  |  |  |  |  |  |  |  |  |  |  |  |  |  |  |  |  |  |  |  |  |
| --- | --- | --- | --- | --- | --- | --- | --- | --- | --- | --- | --- | --- | --- | --- | --- | --- | --- | --- | --- | --- | --- | --- | --- |
| pos | 884 | 1177 | 1377 | 2944 | 4720 | 5512 | 7851 | 8604 | 16914 | 17010 | 19677 | 20663 | 21380 | 22027 | 22029 | 22030 | 22031 | 22032 | 22033 | 22034 | 25757 | 27463 |  |
| ref | C | T | C | G | G | C | C | T | G | C | G | G | C | T | A | G | T | T | C | A | G | G |  |
| AY.4 | C | C | T | A | G | T | T | C | G | T | G | A | T | T | - | - | - | - | - | - | T | T |  |
| AY.4 | C | C | T | A | G | T | T | C | G | T | G | A | T | G | A | A | A | G | T | G | T | G | #544199 |
| AY.114 | T | T | C | G | T | C | C | T | T | C | T | G | C | G | A | A | A | G | T | G | G | G |  |
| mut |  |  |  |  |  |  |  |  |  |  |  |  |  |  |  |  |  |  |  |  | G>T |  |  |

|  |  |  |  |  |  |  |  |  |  |  |  |  |  |  |
| --- | --- | --- | --- | --- | --- | --- | --- | --- | --- | --- | --- | --- | --- | --- |
| pos | 2024 | 12073 | 12189 | 15017 | 17658 | 21987 | 22027 | 22029 | 22030 | 22031 | 22032 | 22033 | 22034 |  |
| ref | A | C | T | C | G | G | T | A | G | T | T | C | A |  |
| AY.4.7 | G | T | G | A | T | G | T | - | - | - | - | - | - |  |
| AY.4.7 | G | T | G | A | T | G | G | A | A | A | G | T | G | #544215 |
| AY.4.7 | A | C | T | C | G | A | G | A | A | A | G | T | G |  |
| mut |  |  |  |  |  |  |  |  |  |  |  |  |  |  |

|  |  |  |  |  |  |  |  |  |  |  |  |  |  |  |  |  |  |  |  |  |  |
| --- | --- | --- | --- | --- | --- | --- | --- | --- | --- | --- | --- | --- | --- | --- | --- | --- | --- | --- | --- | --- | --- |
| pos | 884 | 2533 | 3004 | 3535 | 4720 | 7393 | 7851 | 9856 | 13585 | 16914 | 19677 | 22027 | 22029 | 22030 | 22031 | 22032 | 22033 | 22034 | 22302 | 22899 |  |
| ref | C | T | G | C | G | G | C | G | C | G | G | T | A | G | T | T | C | A | G | G |  |
| AY.4 | C | C | T | T | G | T | T | T | T | G | G | T | - | - | - | - | - | - | G | T |  |
| AY.4 | C | C | T | T | G | T | T | T | T | G | G | G | A | A | A | G | T | G | T | T | #547860 |
| AY.114 | T | T | G | C | T | G | C | G | C | T | T | G | A | A | A | G | T | G | G | G |  |
| mut |  |  |  |  |  |  |  |  |  |  |  |  |  |  |  |  |  |  | G>T | G>T |  |

|  |  |  |  |  |  |  |  |  |  |  |  |  |  |  |  |  |  |  |  |  |  |  |  |  |  |
| --- | --- | --- | --- | --- | --- | --- | --- | --- | --- | --- | --- | --- | --- | --- | --- | --- | --- | --- | --- | --- | --- | --- | --- | --- | --- |
| pos | 427 | 526 | 1820 | 2518 | 3152 | 4237 | 4504 | 6027 | 7851 | 8662 | 11198 | 11620 | 12202 | 15857 | 17040 | 19524 | 21846 | 23191 | 23587 | 26022 | 27389 | 28498 | 28725 | 29409 |  |
| ref | A | G | G | G | G | T | G | C | C | T | G | C | G | C | T | C | C | C | G | C | C | C | C | C |  |
| AY.98 | A | T | G | T | T | T | G | C | C | T | G | C | G | C | T | T | C | C | G | C | C | C | T | C |  |
| AY.4 | A | T | G | T | T | T | G | T | T | C | A | T | T | T | C | C | T | T | C | T | T | T | C | T | #550957 |
| AY.4 | G | G | A | G | G | C | T | T | T | C | A | T | G | C | C | C | T | C | C | T | T | T | C | T |  |
| mut |  |  |  |  |  |  |  |  |  |  |  |  | G>T | C>T |  |  |  | C>T |  |  |  |  |  |  |  |

|  |  |  |  |  |  |  |  |  |  |  |  |  |  |  |  |  |
| --- | --- | --- | --- | --- | --- | --- | --- | --- | --- | --- | --- | --- | --- | --- | --- | --- |
| pos | 44 | 583 | 1722 | 9127 | 10009 | 14120 | 18108 | 22026 | 22027 | 22029 | 22030 | 22031 | 22032 | 22033 | 22034 |  |
| ref | C | C | C | T | T | C | A | G | T | A | G | T | T | C | A |  |
| AY.100 | C | T | T | C | C | T | T | G | T | - | - | - | - | - | - |  |
| AY.100 | T | T | T | C | C | T | T | G | G | A | A | A | G | T | G | #555035 |
| AY.100 | C | C | C | T | T | C | A | T | G | A | A | A | G | T | G |  |
| mut | C>T |  |  |  |  |  |  |  |  |  |  |  |  |  |  |  |

|  |  |  |  |  |  |  |  |  |  |  |  |  |  |  |  |  |  |  |  |  |
| --- | --- | --- | --- | --- | --- | --- | --- | --- | --- | --- | --- | --- | --- | --- | --- | --- | --- | --- | --- | --- |
| pos | 2837 | 8829 | 10279 | 11270 | 11456 | 11962 | 19006 | 21846 | 23284 | 25339 | 26606 | 29050 | 29509 | 29700 | 29762 | 29769 | 29774 | 29779 | 29781 |  |
| ref | T | C | C | A | A | C | G | C | T | C | C | G | C | A | C | C | C | G | G |  |
| AY.119 | T | T | T | G | A | C | T | T | T | C | T | G | C | A | C | C | C | G | G |  |
| AY.3 | G | T | T | G | A | T | T | T | C | T | C | A | T | - | - | - | - | - | - | #558155 |
| AY.3 | T | C | C | A | G | C | G | C | C | T | C | A | T | - | - | - | - | - | - |  |
| mut | T>G |  |  |  |  | C>T |  |  |  |  |  |  |  |  |  |  |  |  |  |  |

|  |  |  |  |  |  |  |  |  |  |  |  |  |  |  |  |  |  |  |  |  |  |  |  |
| --- | --- | --- | --- | --- | --- | --- | --- | --- | --- | --- | --- | --- | --- | --- | --- | --- | --- | --- | --- | --- | --- | --- | --- |
| pos | 697 | 872 | 884 | 4720 | 7851 | 11266 | 11388 | 11401 | 16170 | 16914 | 18803 | 19677 | 21991 | 21992 | 21993 | 22027 | 22029 | 22030 | 22031 | 22032 | 22033 | 22034 |  |
| ref | C | G | C | G | C | G | T | G | T | G | G | G | T | T | A | T | A | G | T | T | C | A |  |
| AY.4 | A | T | C | G | T | T | C | T | C | G | A | G | - | - | - | T | - | - | - | - | - | - |  |
| AY.4 | A | T | C | G | T | T | C | T | C | G | A | G | - | - | - | G | A | A | A | G | T | G | #559125 |
| AY.114 | C | G | T | T | C | G | T | G | T | T | G | T | T | T | A | G | A | A | A | G | T | G |  |
| mut |  |  |  |  |  |  |  |  |  |  |  |  |  |  |  |  |  |  |  |  |  |  |  |

|  |  |  |  |  |  |  |  |  |  |  |  |  |  |  |  |  |  |
| --- | --- | --- | --- | --- | --- | --- | --- | --- | --- | --- | --- | --- | --- | --- | --- | --- | --- |
| pos | 2716 | 7643 | 7851 | 10721 | 15952 | 16935 | 17040 | 18693 | 18744 | 19086 | 19687 | 21846 | 24781 | 25425 | 28299 | 29635 |  |
| ref | C | G | C | T | C | G | T | C | C | G | C | C | G | A | A | C |  |
| AY.4 | C | G | T | T | C | G | C | C | C | G | T | T | G | G | A | T |  |
| AY.4 | T | G | T | C | C | G | C | A | T | T | C | C | T | A | T | C | #559339 |
| AY.43 | C | T | C | T | A | T | T | A | T | T | C | C | T | A | T | C |  |
| mut | C>T |  |  | T>C |  |  |  |  |  |  |  |  |  |  |  |  |  |

|  |  |  |  |  |  |  |  |  |  |  |  |  |  |  |
| --- | --- | --- | --- | --- | --- | --- | --- | --- | --- | --- | --- | --- | --- | --- |
| pos | 4668 | 21161 | 22027 | 22029 | 22030 | 22031 | 22032 | 22033 | 22034 | 25500 | 27484 | 27489 | 29274 |  |
| ref | C | C | T | A | G | T | T | C | A | G | T | A | C |  |
| AY.4 | C | C | G | A | A | A | G | T | G | G | T | A | C |  |
| AY.4 | T | T | G | A | A | A | G | T | G | T | C | C | T | #560801 |
| AY.4 | C | T | T | - | - | - | - | - | - | T | C | C | T |  |
| mut | C>T | C>T |  |  |  |  |  |  |  |  |  |  |  |  |

|  |  |  |  |  |  |  |  |  |  |  |  |  |  |  |  |  |  |  |
| --- | --- | --- | --- | --- | --- | --- | --- | --- | --- | --- | --- | --- | --- | --- | --- | --- | --- | --- |
| pos | 5015 | 5935 | 8299 | 10642 | 13078 | 14181 | 15963 | 20178 | 21851 | 22027 | 22029 | 22030 | 22031 | 22032 | 22033 | 22034 | 24415 |  |
| ref | G | A | C | G | T | G | C | C | A | T | A | G | T | T | C | A | G |  |
| AY.20 | A | A | T | T | G | A | T | T | G | T | - | - | - | - | - | - | G |  |
| AY.20 | A | G | T | T | G | A | T | T | G | G | A | A | A | G | T | G | T | #562218 |
| AY.20 | G | A | C | G | T | G | C | C | A | G | A | A | A | G | T | G | G |  |
| mut |  | A>G |  |  |  |  |  |  |  |  |  |  |  |  |  |  | G>T |  |

|  |  |  |  |  |  |  |  |  |  |  |  |  |  |  |  |
| --- | --- | --- | --- | --- | --- | --- | --- | --- | --- | --- | --- | --- | --- | --- | --- |
| pos | 2062 | 3433 | 5628 | 10423 | 17338 | 22026 | 22027 | 22029 | 22030 | 22031 | 22032 | 22033 | 22034 | 25599 |  |
| ref | C | G | C | T | G | G | T | A | G | T | T | C | A | G |  |
| AY.39 | T | T | T | C | T | G | T | - | - | - | - | - | - | T |  |
| AY.39 | T | T | T | C | T | T | G | A | A | A | G | T | G | T | #562656 |
| AY.39 | C | G | C | T | G | T | G | A | A | A | G | T | G | G |  |
| mut |  |  |  |  |  |  |  |  |  |  |  |  |  | G>T |  |

|  |  |  |  |  |  |  |  |  |  |  |  |  |  |  |  |  |  |  |  |  |  |
| --- | --- | --- | --- | --- | --- | --- | --- | --- | --- | --- | --- | --- | --- | --- | --- | --- | --- | --- | --- | --- | --- |
| pos | 526 | 7042 | 7851 | 9492 | 10317 | 16075 | 17977 | 19524 | 21219 | 21513 | 21846 | 24535 | 26093 | 26111 | 27610 | 27875 | 27881 | 27882 | 27883 | 29085 |  |
| ref | G | G | C | A | A | G | C | C | C | C | C | G | A | C | C | T | C | G | C | C |  |
| AY.4 | G | T | T | C | G | A | C | C | T | C | T | A | T | T | T | T | C | G | C | C |  |
| AY.4 | G | T | T | C | G | A | T | C | T | T | T | A | T | T | T | C | T | C | T | T | #568400 |
| AY.98 | T | G | C | A | A | G | C | T | C | C | C | G | A | C | C | C | T | C | T | C |  |
| mut |  |  |  |  |  |  | C>T |  |  | C>T |  |  |  |  |  |  |  |  |  | C>T |  |

|  |  |  |  |  |  |  |  |  |  |  |  |  |  |  |
| --- | --- | --- | --- | --- | --- | --- | --- | --- | --- | --- | --- | --- | --- | --- |
| pos | 1942 | 6425 | 10138 | 15928 | 16530 | 21987 | 22027 | 22029 | 22030 | 22031 | 22032 | 22033 | 22034 |  |
| ref | G | A | C | C | T | G | T | A | G | T | T | C | A |  |
| AY.4.2 | A | G | T | T | C | A | T | - | - | - | - | - | - |  |
| AY.4.2 | A | G | T | T | C | A | G | A | A | A | G | T | G | #571040 |
| AY.4.2 | G | A | C | C | T | G | G | A | A | A | G | T | G |  |
| mut |  |  |  |  |  |  |  |  |  |  |  |  |  |  |

|  |  |  |  |  |  |  |  |  |  |  |  |  |  |  |  |
| --- | --- | --- | --- | --- | --- | --- | --- | --- | --- | --- | --- | --- | --- | --- | --- |
| pos | 599 | 5192 | 5608 | 7764 | 9363 | 11767 | 17964 | 22027 | 22029 | 22030 | 22031 | 22032 | 22033 | 22034 |  |
| ref | G | C | A | C | C | C | G | T | A | G | T | T | C | A |  |
| AY.34.1 | A | C | A | T | C | C | T | T | - | - | - | - | - | - |  |
| AY.34.1 | A | C | A | T | C | C | T | G | A | A | A | G | T | G | #571273 |
| AY.34.1 | G | T | G | C | T | T | G | G | A | A | A | G | T | G |  |
| mut |  |  |  |  |  |  |  |  |  |  |  |  |  |  |  |

|  |  |  |  |  |  |  |  |  |  |  |  |  |  |  |  |  |  |
| --- | --- | --- | --- | --- | --- | --- | --- | --- | --- | --- | --- | --- | --- | --- | --- | --- | --- |
| pos | 255 | 1412 | 15192 | 15720 | 18687 | 19782 | 21304 | 21305 | 21987 | 22027 | 22029 | 22030 | 22031 | 22032 | 22033 | 22034 |  |
| ref | C | A | A | C | C | A | C | G | G | T | A | G | T | T | C | A |  |
| AY.4.2 | T | A | G | T | T | G | A | A | A | T | - | - | - | - | - | - |  |
| AY.4.2 | T | G | G | T | T | G | A | A | A | G | A | A | A | G | T | G | #574335 |
| AY.4.2 | C | A | A | C | C | A | C | G | G | G | A | A | A | G | T | G |  |
| mut |  | A>G |  |  |  |  |  |  |  |  |  |  |  |  |  |  |  |

|  |  |  |  |  |  |  |  |  |  |  |  |  |  |  |  |  |  |  |  |  |  |
| --- | --- | --- | --- | --- | --- | --- | --- | --- | --- | --- | --- | --- | --- | --- | --- | --- | --- | --- | --- | --- | --- |
| pos | 439 | 884 | 4720 | 7851 | 9520 | 13348 | 15277 | 16914 | 17040 | 17427 | 19677 | 22027 | 22029 | 22030 | 22031 | 22032 | 22033 | 22034 | 23542 | 29639 |  |
| ref | A | C | G | C | C | G | C | G | T | G | G | T | A | G | T | T | C | A | T | G |  |
| AY.4 | G | C | G | T | T | T | T | G | C | T | G | T | - | - | - | - | - | - | C | T |  |
| AY.4 | G | C | G | T | T | T | T | G | C | T | G | G | A | A | A | G | T | G | T | T | #574565 |
| AY.114 | A | T | T | C | C | G | C | T | T | G | T | G | A | A | A | G | T | G | T | G |  |
| mut |  |  |  |  |  |  |  |  |  |  |  |  |  |  |  |  |  |  |  | G>T |  |

|  |  |  |  |  |  |  |  |  |  |  |  |  |  |  |
| --- | --- | --- | --- | --- | --- | --- | --- | --- | --- | --- | --- | --- | --- | --- |
| pos | 3692 | 4021 | 5668 | 8985 | 9360 | 17913 | 22027 | 22029 | 22030 | 22031 | 22032 | 22033 | 22034 |  |
| ref | G | C | G | A | C | C | T | A | G | T | T | C | A |  |
| AY.117 | T | T | G | A | T | C | T | - | - | - | - | - | - |  |
| AY.117 | T | T | G | A | T | C | G | A | A | A | G | T | G | #575251 |
| AY.117 | G | C | A | G | C | T | G | A | A | A | G | T | G |  |
| mut |  |  |  |  |  |  |  |  |  |  |  |  |  |  |

|  |  |  |  |  |  |  |  |  |  |  |  |  |  |  |  |  |  |  |  |  |
| --- | --- | --- | --- | --- | --- | --- | --- | --- | --- | --- | --- | --- | --- | --- | --- | --- | --- | --- | --- | --- |
| pos | 1588 | 2632 | 2923 | 3476 | 6478 | 6726 | 8964 | 10747 | 12025 | 14273 | 15654 | 22027 | 22029 | 22030 | 22031 | 22032 | 22033 | 22034 | 22444 |  |
| ref | A | G | A | G | T | C | C | C | C | A | C | T | A | G | T | T | C | A | C |  |
| AY.39 | A | G | G | G | C | C | T | T | T | C | C | T | - | - | - | - | - | - | T |  |
| AY.39 | A | G | G | G | C | C | T | T | T | C | C | G | A | A | A | G | T | G | T | #575303 |
| AY.39 | G | T | A | A | T | T | C | C | C | A | T | G | A | A | A | G | T | G | C |  |
| mut |  |  |  |  |  |  |  |  |  |  |  |  |  |  |  |  |  |  | C>T |  |

|  |  |  |  |  |  |  |  |  |  |  |  |  |  |  |  |  |  |
| --- | --- | --- | --- | --- | --- | --- | --- | --- | --- | --- | --- | --- | --- | --- | --- | --- | --- |
| pos | 1405 | 2485 | 4237 | 4321 | 4897 | 10279 | 13458 | 21987 | 22027 | 22029 | 22030 | 22031 | 22032 | 22033 | 22034 | 25555 |  |
| ref | T | C | T | C | C | C | C | G | T | A | G | T | T | C | A | G |  |
| AY.4 | C | T | C | C | C | T | T | A | T | - | - | - | - | - | - | G |  |
| AY.4 | C | T | C | C | C | T | T | A | G | A | A | A | G | T | G | T | #578351 |
| AY.4 | T | C | T | T | T | C | C | G | G | A | A | A | G | T | G | T |  |
| mut |  |  |  |  |  |  |  |  |  |  |  |  |  |  |  |  |  |

|  |  |  |  |  |  |  |  |  |  |  |  |  |  |  |  |  |  |  |  |  |  |  |  |  |  |  |  |  |  |  |  |  |  |  |  |  |
| --- | --- | --- | --- | --- | --- | --- | --- | --- | --- | --- | --- | --- | --- | --- | --- | --- | --- | --- | --- | --- | --- | --- | --- | --- | --- | --- | --- | --- | --- | --- | --- | --- | --- | --- | --- | --- |
| pos | 3461 | 3948 | 4181 | 5184 | 5584 | 6402 | 7124 | 7393 | 8986 | 9053 | 9891 | 10029 | 10741 | 11201 | 11332 | 11418 | 11514 | 11562 | 13019 | 15543 | 17795 | 19220 | 21622 | 22205 | 22227 | 24319 | 25618 | 26062 | 26107 | 27112 | 27507 | 27516 | 27874 | 28916 | 29738 |  |
| ref | A | A | G | C | A | C | C | G | C | G | C | C | C | A | A | T | C | G | C | G | C | C | C | G | C | C | G | G | G | G | A | G | C | G | C |  |
| AY.75 | G | A | G | T | G | C | C | T | C | G | T | C | C | A | A | C | T | G | T | G | C | C | T | T | T | C | G | T | G | A | A | A | C | G | A |  |
| AY.75 | G | A | G | T | G | C | C | T | C | G | T | C | C | A | A | C | T | G | T | G | C | C | T | T | T | T | A | G | C | G | C | G | T | T | C | #578884 |
| AY.25.1 | A | G | T | C | A | T | T | G | T | T | C | T | T | G | G | T | C | T | C | T | G | T | C | G | C | T | G | G | C | G | C | G | T | T | C |  |
| mut |  |  |  |  |  |  |  |  |  |  |  |  |  |  |  |  |  |  |  |  |  |  |  |  |  |  | G>A |  |  |  |  |  |  |  |  |  |

|  |  |  |  |  |  |  |  |  |  |  |  |  |  |  |  |  |  |  |  |  |  |  |  |  |  |  |
| --- | --- | --- | --- | --- | --- | --- | --- | --- | --- | --- | --- | --- | --- | --- | --- | --- | --- | --- | --- | --- | --- | --- | --- | --- | --- | --- |
| pos | 2306 | 2564 | 3211 | 3948 | 8728 | 10870 | 11456 | 11562 | 19419 | 19992 | 24208 | 25489 | 25553 | 25976 | 26107 | 27507 | 29044 | 29050 | 29509 | 29700 | 29762 | 29769 | 29774 | 29779 | 29781 |  |
| ref | C | G | T | A | A | G | A | G | G | T | C | G | C | C | G | A | G | G | C | A | C | C | C | G | G |  |
| AY.25.1 | C | A | C | G | A | G | A | T | G | C | C | T | C | T | C | C | G | G | C | A | C | C | C | G | G |  |
| AY.3 | C | A | C | G | A | T | G | G | T | T | C | G | T | C | G | A | A | A | C | - | - | - | - | - | - | #579097 |
| AY.3 | T | G | T | A | G | T | G | G | T | T | T | G | T | C | G | A | A | A | T | - | - | - | - | - | - |  |
| mut |  |  |  |  |  |  |  |  |  |  | T>C |  |  |  |  |  |  |  | T>C |  |  |  |  |  |  |  |

|  |  |  |  |  |  |  |  |  |  |  |  |  |  |  |
| --- | --- | --- | --- | --- | --- | --- | --- | --- | --- | --- | --- | --- | --- | --- |
| pos | 526 | 6040 | 7926 | 9529 | 12693 | 14014 | 19524 | 21987 | 27875 | 27881 | 27882 | 27883 | 29700 |  |
| ref | G | C | C | A | A | T | C | G | T | C | G | C | A |  |
| AY.44 | G | T | T | A | T | G | C | G | T | C | G | C | G |  |
| AY.44 | G | T | T | G | T | G | C | G | C | T | C | T | G | #579202 |
| AY.98 | T | C | C | A | A | T | T | A | C | T | C | T | A |  |
| mut |  |  |  | A>G |  |  |  |  |  |  |  |  | A>G |  |

|  |  |  |  |  |  |  |  |  |  |  |  |  |  |  |  |  |  |  |
| --- | --- | --- | --- | --- | --- | --- | --- | --- | --- | --- | --- | --- | --- | --- | --- | --- | --- | --- |
| pos | 5203 | 7105 | 9451 | 12756 | 15567 | 15652 | 21534 | 22027 | 22029 | 22030 | 22031 | 22032 | 22033 | 22034 | 24620 | 28868 | 29700 |  |
| ref | C | C | C | C | T | G | T | T | A | G | T | T | C | A | G | C | A |  |
| AY.119 | T | T | T | T | C | T | C | T | - | - | - | - | - | - | T | A | A |  |
| AY.119 | T | T | T | T | C | T | C | G | A | A | A | G | T | G | T | A | A | #579572 |
| AY.119 | C | C | C | C | T | G | T | G | A | A | A | G | T | G | G | C | G |  |
| mut |  |  |  |  |  |  |  |  |  |  |  |  |  |  | G>T | C>A | G>A |  |

|  |  |  |  |  |  |  |  |  |  |  |  |  |  |  |  |  |  |
| --- | --- | --- | --- | --- | --- | --- | --- | --- | --- | --- | --- | --- | --- | --- | --- | --- | --- |
| pos | 805 | 1264 | 6525 | 16509 | 19017 | 21809 | 22026 | 22027 | 22029 | 22030 | 22031 | 22032 | 22033 | 22034 | 23282 | 25584 |  |
| ref | G | G | C | T | C | G | G | T | A | G | T | T | C | A | G | C |  |
| AY.100 | A | T | T | C | T | A | G | T | - | - | - | - | - | - | T | T |  |
| AY.100 | A | T | T | C | T | A | G | G | A | A | A | G | T | G | T | T | #579674 |
| AY.100 | G | G | C | T | C | G | T | G | A | A | A | G | T | G | G | C |  |
| mut |  |  |  |  |  |  |  |  |  |  |  |  |  |  | G>T | C>T |  |

|  |  |  |  |  |  |  |  |  |  |  |  |  |  |  |  |  |  |  |  |
| --- | --- | --- | --- | --- | --- | --- | --- | --- | --- | --- | --- | --- | --- | --- | --- | --- | --- | --- | --- |
| pos | 884 | 1639 | 4720 | 9803 | 10977 | 13901 | 16914 | 19677 | 21846 | 22027 | 22029 | 22030 | 22031 | 22032 | 22033 | 22034 | 25673 | 27754 |  |
| ref | C | A | G | C | C | A | G | G | C | T | A | G | T | T | C | A | T | G |  |
| AY.46.1 | C | C | G | T | T | G | G | G | C | T | - | - | - | - | - | - | C | G |  |
| AY.46.1 | C | C | G | T | T | G | G | G | C | G | A | A | A | G | T | G | C | C | #579956 |
| AY.114 | T | A | T | C | C | A | T | T | T | G | A | A | A | G | T | G | T | G |  |
| mut |  |  |  |  |  |  |  |  |  |  |  |  |  |  |  |  | T>C | G>C |  |

|  |  |  |  |  |  |  |  |  |  |  |  |  |  |  |  |  |  |  |  |  |  |  |  |  |  |  |  |  |  |  |  |  |  |  |  |  |  |  |  |  |
| --- | --- | --- | --- | --- | --- | --- | --- | --- | --- | --- | --- | --- | --- | --- | --- | --- | --- | --- | --- | --- | --- | --- | --- | --- | --- | --- | --- | --- | --- | --- | --- | --- | --- | --- | --- | --- | --- | --- | --- | --- |
| pos | 448 | 1684 | 4181 | 4893 | 5062 | 5164 | 5184 | 5209 | 5584 | 5612 | 6402 | 7124 | 7851 | 8986 | 9053 | 9891 | 10029 | 11124 | 11201 | 11332 | 11418 | 11446 | 11514 | 12049 | 12514 | 13019 | 17040 | 19220 | 19983 | 21846 | 22227 | 22427 | 23013 | 27874 | 28107 | 28326 | 28916 | 29700 | 29836 |  |
| ref | G | C | G | C | G | G | C | A | A | G | C | C | C | C | G | C | C | C | A | A | T | A | C | C | G | C | T | C | C | C | C | G | A | C | C | G | G | A | C |  |
| AY.47 | G | T | G | C | G | T | T | A | G | G | C | C | C | C | G | T | C | C | A | A | C | A | T | T | G | T | T | C | C | C | T | A | A | C | C | T | G | A | T |  |
| AY.47 | G | T | G | C | A | T | T | G | G | G | C | C | C | C | G | T | C | C | A | A | C | A | T | T | G | T | T | C | T | C | T | A | G | T | T | G | T | G | C | #583110 |
| AY.4 | T | C | T | T | G | G | C | A | A | T | T | T | T | T | T | C | T | T | G | G | T | G | C | C | A | C | C | T | C | T | C | G | G | T | C | G | T | A | C |  |
| mut |  |  |  |  | G>A |  |  | A>G |  |  |  |  |  |  |  |  |  |  |  |  |  |  |  |  |  |  |  |  | C>T |  |  |  |  |  | C>T |  |  | A>G |  |  |

|  |  |  |  |  |  |  |  |  |  |  |  |  |  |  |  |  |  |  |  |  |  |  |  |  |  |  |  |
| --- | --- | --- | --- | --- | --- | --- | --- | --- | --- | --- | --- | --- | --- | --- | --- | --- | --- | --- | --- | --- | --- | --- | --- | --- | --- | --- | --- |
| pos | 4012 | 7246 | 8829 | 10279 | 11456 | 12513 | 12756 | 13426 | 13701 | 15141 | 19006 | 21754 | 21846 | 23284 | 23987 | 24914 | 25339 | 29050 | 29509 | 29687 | 29700 | 29762 | 29769 | 29774 | 29779 | 29781 |  |
| ref | C | G | C | C | A | C | C | C | T | C | G | G | C | T | C | G | C | G | C | A | A | C | C | C | G | G |  |
| AY.119 | C | G | T | T | A | C | T | C | T | C | T | G | T | T | T | T | C | G | C | A | A | C | C | C | G | G |  |
| AY.3 | C | G | T | T | G | T | C | T | C | T | G | T | C | C | C | G | T | A | T | C | - | - | - | - | - | - | #583782 |
| AY.3 | T | T | C | C | G | T | C | C | C | T | G | G | C | C | C | G | T | A | T | C | - | - | - | - | - | - |  |
| mut |  |  |  |  |  |  |  | C>T |  |  |  | G>T |  |  |  |  |  |  |  |  |  |  |  |  |  |  |  |

|  |  |  |  |  |  |  |  |  |  |  |  |  |  |  |  |
| --- | --- | --- | --- | --- | --- | --- | --- | --- | --- | --- | --- | --- | --- | --- | --- |
| pos | 532 | 2508 | 5812 | 9133 | 17427 | 21806 | 22026 | 22027 | 22029 | 22030 | 22031 | 22032 | 22033 | 22034 |  |
| ref | A | C | C | C | G | C | G | T | A | G | T | T | C | A |  |
| AY.100 | T | T | T | A | T | A | G | T | - | - | - | - | - | - |  |
| AY.100 | T | T | T | A | T | A | G | G | A | A | A | G | T | G | #584501 |
| AY.100 | A | C | C | C | G | C | T | G | A | A | A | G | T | G |  |
| mut |  |  |  |  |  |  |  |  |  |  |  |  |  |  |  |

|  |  |  |  |  |  |  |  |  |  |  |  |  |  |  |  |  |  |  |  |
| --- | --- | --- | --- | --- | --- | --- | --- | --- | --- | --- | --- | --- | --- | --- | --- | --- | --- | --- | --- |
| pos | 526 | 2706 | 5206 | 7042 | 7851 | 12988 | 17040 | 18927 | 19524 | 20374 | 21846 | 22021 | 22899 | 25803 | 27875 | 27881 | 27882 | 27883 |  |
| ref | G | C | G | G | C | G | T | T | C | A | C | G | G | C | T | C | G | C |  |
| AY.4.1 | G | T | T | T | T | T | C | C | C | C | T | G | T | T | T | C | G | C |  |
| AY.4.1 | G | T | T | T | T | T | C | C | C | C | T | T | T | T | C | T | C | T | #586363 |
| AY.98 | T | C | G | G | C | G | T | T | T | A | C | G | G | C | C | T | C | T |  |
| mut |  |  |  |  |  |  |  |  |  |  |  | G>T |  |  |  |  |  |  |  |

|  |  |  |  |  |  |  |  |  |  |  |  |  |  |  |  |  |  |  |  |  |  |  |
| --- | --- | --- | --- | --- | --- | --- | --- | --- | --- | --- | --- | --- | --- | --- | --- | --- | --- | --- | --- | --- | --- | --- |
| pos | 61 | 884 | 1281 | 4720 | 5008 | 7851 | 16914 | 17040 | 18131 | 18814 | 19677 | 20006 | 21660 | 21987 | 22027 | 22029 | 22030 | 22031 | 22032 | 22033 | 22034 |  |
| ref | G | C | C | G | G | C | G | T | C | C | G | G | C | G | T | A | G | T | T | C | A |  |
| AY.4 | T | C | T | G | T | T | G | C | T | T | G | T | T | G | T | - | - | - | - | - | - |  |
| AY.4 | T | C | T | G | T | T | G | C | T | T | G | T | T | G | G | A | A | A | G | T | G | #586953 |
| AY.114 | G | T | C | T | G | C | T | T | C | C | T | G | C | A | G | A | A | A | G | T | G |  |
| mut |  |  |  |  |  |  |  |  |  |  |  |  |  |  |  |  |  |  |  |  |  |  |

|  |  |  |  |  |  |  |  |  |  |  |  |  |  |  |  |  |  |  |  |  |  |  |  |  |
| --- | --- | --- | --- | --- | --- | --- | --- | --- | --- | --- | --- | --- | --- | --- | --- | --- | --- | --- | --- | --- | --- | --- | --- | --- |
| pos | 1588 | 2632 | 3476 | 4237 | 4795 | 5281 | 6726 | 7851 | 8110 | 13210 | 14742 | 15654 | 17040 | 20451 | 21372 | 22027 | 22029 | 22030 | 22031 | 22032 | 22033 | 22034 | 27604 |  |
| ref | A | G | G | T | C | C | C | C | T | T | T | C | T | C | G | T | A | G | T | T | C | A | G |  |
| AY.4 | A | G | G | C | T | T | C | T | C | C | C | C | C | T | G | T | - | - | - | - | - | - | G |  |
| AY.4 | A | G | G | C | T | T | C | T | C | C | C | C | C | T | G | G | A | A | A | G | T | G | A | #586964 |
| AY.39 | G | T | A | T | C | C | T | C | T | T | T | T | T | C | T | G | A | A | A | G | T | G | A |  |
| mut |  |  |  |  |  |  |  |  |  |  |  |  |  |  |  |  |  |  |  |  |  |  |  |  |

|  |  |  |  |  |  |  |  |  |  |  |  |  |  |  |  |  |  |  |  |  |  |  |
| --- | --- | --- | --- | --- | --- | --- | --- | --- | --- | --- | --- | --- | --- | --- | --- | --- | --- | --- | --- | --- | --- | --- |
| pos | 335 | 721 | 884 | 1997 | 4300 | 4720 | 6070 | 7851 | 11173 | 16914 | 18981 | 19677 | 22027 | 22029 | 22030 | 22031 | 22032 | 22033 | 22034 | 28542 | 28713 |  |
| ref | C | T | C | C | G | G | C | C | C | G | C | G | T | A | G | T | T | C | A | C | G |  |
| AY.4 | T | C | C | T | T | G | T | T | T | G | T | G | T | - | - | - | - | - | - | T | A |  |
| AY.4 | T | C | C | T | T | G | T | T | T | G | T | G | G | A | A | A | G | T | G | T | A | #587062 |
| AY.114 | C | T | T | C | G | T | C | C | C | T | C | T | G | A | A | A | G | T | G | C | G |  |
| mut |  |  |  |  |  |  |  |  |  |  |  |  |  |  |  |  |  |  |  | C>T | G>A |  |

|  |  |  |  |  |  |  |  |  |  |  |  |  |  |  |  |
| --- | --- | --- | --- | --- | --- | --- | --- | --- | --- | --- | --- | --- | --- | --- | --- |
| pos | 526 | 1099 | 2336 | 10323 | 15277 | 17331 | 17805 | 19524 | 24905 | 25688 | 27875 | 27881 | 27882 | 27883 |  |
| ref | G | T | G | A | C | G | A | C | A | C | T | C | G | C |  |
| AY.6 | G | T | A | A | T | T | G | C | G | T | T | C | G | C |  |
| AY.6 | G | C | A | G | T | T | G | C | G | T | C | T | C | T | #589888 |
| AY.98 | T | T | G | A | C | G | A | T | A | C | C | T | C | T |  |
| mut |  | T>C |  | A>G |  |  |  |  |  |  |  |  |  |  |  |

|  |  |  |  |  |  |  |  |  |  |  |  |  |  |  |  |  |  |  |  |  |  |  |
| --- | --- | --- | --- | --- | --- | --- | --- | --- | --- | --- | --- | --- | --- | --- | --- | --- | --- | --- | --- | --- | --- | --- |
| pos | 203 | 222 | 811 | 1882 | 1958 | 4516 | 9532 | 10456 | 10870 | 17040 | 18348 | 18888 | 19723 | 20934 | 21767 | 22205 | 23039 | 23430 | 27406 | 27936 | 28139 |  |
| ref | C | C | C | G | G | T | C | C | G | T | C | C | G | T | C | G | C | C | C | G | C |  |
| AY.4 | C | C | T | T | A | C | T | C | T | C | T | C | G | C | T | T | C | C | C | A | T |  |
| AY.4 | - | - | T | T | A | C | C | T | G | T | C | T | T | T | C | G | G | T | C | A | T | #594260 |
| AY.4 | C | C | C | G | G | T | C | T | G | T | C | T | T | T | C | G | G | T | A | G | C |  |
| mut | C>- | C>- |  |  |  |  |  |  |  |  |  |  |  |  |  |  |  |  | A>C | G>A | C>T |  |

|  |  |  |  |  |  |  |  |  |  |  |  |  |  |  |  |  |  |  |  |  |  |
| --- | --- | --- | --- | --- | --- | --- | --- | --- | --- | --- | --- | --- | --- | --- | --- | --- | --- | --- | --- | --- | --- |
| pos | 4931 | 5040 | 7851 | 9165 | 10323 | 12772 | 15240 | 15285 | 15854 | 15952 | 18795 | 19273 | 22026 | 22027 | 22029 | 22030 | 22031 | 22032 | 22033 | 22034 |  |
| ref | C | A | C | C | A | A | C | G | T | C | C | C | G | T | A | G | T | T | C | A |  |
| AY.113 | A | A | C | T | G | G | T | T | T | T | T | T | G | T | - | - | - | - | - | - |  |
| AY.113 | A | G | C | T | G | G | T | T | A | T | T | T | T | G | A | A | A | G | T | G | #596108 |
| AY.4 | C | A | T | C | A | A | C | G | T | C | C | C | T | G | A | A | A | G | T | G |  |
| mut |  | A>G |  |  |  |  |  |  | T>A |  |  |  |  |  |  |  |  |  |  |  |  |

|  |  |  |  |  |  |  |  |  |  |  |  |  |  |  |
| --- | --- | --- | --- | --- | --- | --- | --- | --- | --- | --- | --- | --- | --- | --- |
| pos | 1651 | 14145 | 16075 | 18412 | 22027 | 22029 | 22030 | 22031 | 22032 | 22033 | 22034 | 22344 | 28083 |  |
| ref | T | T | G | G | T | A | G | T | T | C | A | G | G |  |
| AY.117 | C | C | T | T | T | - | - | - | - | - | - | T | G |  |
| AY.117 | C | C | T | T | G | A | A | A | G | T | G | T | T | #596127 |
| AY.117 | T | T | G | G | G | A | A | A | G | T | G | G | G |  |
| mut |  |  |  |  |  |  |  |  |  |  |  | G>T | G>T |  |

|  |  |  |  |  |  |  |  |  |  |  |  |  |  |  |  |  |  |  |  |  |
| --- | --- | --- | --- | --- | --- | --- | --- | --- | --- | --- | --- | --- | --- | --- | --- | --- | --- | --- | --- | --- |
| pos | 526 | 7851 | 10165 | 16425 | 17040 | 19524 | 21846 | 21995 | 22227 | 25614 | 25667 | 25855 | 26172 | 26727 | 27143 | 27875 | 27881 | 27882 | 27883 |  |
| ref | G | C | C | T | T | C | C | T | C | C | C | G | G | G | C | T | C | G | C |  |
| AY.4.2 | G | T | T | A | C | C | T | C | T | T | T | T | A | T | T | T | C | G | C |  |
| AY.4.2 | G | T | T | A | C | C | T | C | T | T | T | T | A | T | T | C | T | C | T | #598035 |
| AY.98 | T | C | C | T | T | T | C | T | C | C | C | G | G | G | C | C | T | C | T |  |
| mut |  |  |  |  |  |  |  |  |  |  |  |  |  |  |  |  |  |  |  |  |

|  |  |  |  |  |  |  |  |  |  |  |  |  |  |  |  |  |  |  |  |  |
| --- | --- | --- | --- | --- | --- | --- | --- | --- | --- | --- | --- | --- | --- | --- | --- | --- | --- | --- | --- | --- |
| pos | 878 | 1820 | 6336 | 9611 | 10988 | 11411 | 15352 | 18533 | 19955 | 22026 | 22027 | 22029 | 22030 | 22031 | 22032 | 22033 | 22034 | 22599 | 25785 |  |
| ref | C | G | C | C | A | A | C | A | C | G | T | A | G | T | T | C | A | G | G |  |
| AY.39 | T | A | C | T | G | G | T | G | T | G | T | - | - | - | - | - | - | T | T |  |
| AY.39 | T | A | T | T | G | G | T | G | C | T | G | A | A | A | G | T | G | T | T | #600043 |
| AY.39 | C | G | C | C | A | A | C | A | C | T | G | A | A | A | G | T | G | G | G |  |
| mut |  |  | C>T |  |  |  |  |  |  |  |  |  |  |  |  |  |  | G>T | G>T |  |

|  |  |  |  |  |  |  |  |  |  |  |  |  |  |  |
| --- | --- | --- | --- | --- | --- | --- | --- | --- | --- | --- | --- | --- | --- | --- |
| pos | 6408 | 6613 | 7420 | 7851 | 8809 | 17467 | 20758 | 21714 | 21974 | 25003 | 25919 | 29580 | 29762 |  |
| ref | C | A | C | C | C | T | G | C | G | A | C | C | C |  |
| AY.120 | T | A | T | C | T | T | G | T | G | G | C | C | C |  |
| AY.120 | T | G | T | C | T | T | G | C | C | A | T | T | - | #602056 |
| AY.4 | C | A | C | T | C | C | A | C | C | A | T | T | C |  |
| mut |  | A>G |  |  |  |  |  |  |  |  |  |  | C>- |  |

|  |  |  |  |  |  |  |  |  |  |  |  |  |  |  |  |  |  |  |  |  |  |  |  |
| --- | --- | --- | --- | --- | --- | --- | --- | --- | --- | --- | --- | --- | --- | --- | --- | --- | --- | --- | --- | --- | --- | --- | --- |
| pos | 884 | 1405 | 2485 | 3250 | 4237 | 4720 | 7851 | 10084 | 10279 | 13458 | 16914 | 17040 | 19677 | 20262 | 22027 | 22029 | 22030 | 22031 | 22032 | 22033 | 22034 | 22648 |  |
| ref | C | T | C | G | T | G | C | T | C | C | G | T | G | A | T | A | G | T | T | C | A | T |  |
| AY.4 | C | C | T | T | C | G | T | C | T | T | G | C | G | G | T | - | - | - | - | - | - | A |  |
| AY.4 | C | C | T | T | C | G | T | C | T | T | G | C | G | G | G | A | A | A | G | T | G | A | #602556 |
| AY.114 | T | T | C | G | T | T | C | T | C | C | T | T | T | A | G | A | A | A | G | T | G | T |  |
| mut |  |  |  |  |  |  |  |  |  |  |  |  |  |  |  |  |  |  |  |  |  | T>A |  |

|  |  |  |  |  |  |  |  |  |  |  |  |  |  |  |  |  |  |  |  |  |  |  |  |  |  |  |  |  |  |  |  |  |  |  |  |  |  |  |
| --- | --- | --- | --- | --- | --- | --- | --- | --- | --- | --- | --- | --- | --- | --- | --- | --- | --- | --- | --- | --- | --- | --- | --- | --- | --- | --- | --- | --- | --- | --- | --- | --- | --- | --- | --- | --- | --- | --- |
| pos | 1274 | 1820 | 4181 | 4276 | 5184 | 5584 | 6402 | 7124 | 7393 | 8986 | 9053 | 9891 | 10029 | 11201 | 11332 | 11418 | 11514 | 13019 | 15012 | 15982 | 18471 | 19220 | 21622 | 21846 | 22184 | 22227 | 24499 | 24863 | 26062 | 27112 | 27516 | 27874 | 28392 | 28916 | 28961 | 29358 | 29738 |  |
| ref | G | G | G | C | C | A | C | C | G | C | G | C | C | A | A | T | C | C | A | G | T | C | C | C | A | C | T | C | G | G | G | C | G | G | C | C | C |  |
| AY.107 | A | G | T | C | C | A | T | T | G | T | T | C | T | G | G | T | C | C | G | G | C | T | C | T | A | C | T | T | G | G | G | T | G | T | T | T | C |  |
| B.1.617.2 | A | A | T | C | C | A | T | T | G | T | T | C | T | G | G | T | C | C | G | G | C | T | C | T | G | C | C | C | T | A | A | C | T | G | C | C | A | #602842 |
| AY.75 | G | G | G | T | T | G | C | C | T | C | G | T | C | A | A | C | T | T | A | A | T | C | T | C | A | T | T | C | T | A | A | C | T | G | C | C | A |  |
| mut |  | G>A |  |  |  |  |  |  |  |  |  |  |  |  |  |  |  |  |  |  |  |  |  |  | A>G |  | T>C |  |  |  |  |  |  |  |  |  |  |  |

|  |  |  |  |  |  |  |  |  |  |  |  |  |  |
| --- | --- | --- | --- | --- | --- | --- | --- | --- | --- | --- | --- | --- | --- |
| pos | 17034 | 22027 | 22029 | 22030 | 22031 | 22032 | 22033 | 22034 | 22217 | 25396 | 26256 | 28000 |  |
| ref | T | T | A | G | T | T | C | A | G | G | C | C |  |
| AY.119 | T | G | A | A | A | G | T | G | G | G | C | C |  |
| AY.119 | A | G | A | A | A | G | T | G | A | T | T | T | #603828 |
| AY.119 | A | T | - | - | - | - | - | - | A | T | T | T |  |
| mut | T>A |  |  |  |  |  |  |  |  |  |  |  |  |

|  |  |  |  |  |  |  |  |  |  |  |  |  |  |  |  |  |  |  |  |  |
| --- | --- | --- | --- | --- | --- | --- | --- | --- | --- | --- | --- | --- | --- | --- | --- | --- | --- | --- | --- | --- |
| pos | 884 | 2902 | 4720 | 5878 | 5974 | 7851 | 16914 | 19677 | 20234 | 21144 | 22027 | 22029 | 22030 | 22031 | 22032 | 22033 | 22034 | 25096 | 29266 |  |
| ref | C | C | G | C | C | C | G | G | C | T | T | A | G | T | T | C | A | C | G |  |
| AY.4 | C | T | G | T | C | T | G | G | T | C | T | - | - | - | - | - | - | C | G |  |
| AY.4 | C | T | G | T | T | T | G | G | T | C | G | A | A | A | G | T | G | T | T | #606286 |
| AY.114 | T | C | T | C | C | C | T | T | C | T | G | A | A | A | G | T | G | C | G |  |
| mut |  |  |  |  | C>T |  |  |  |  |  |  |  |  |  |  |  |  | C>T | G>T |  |

|  |  |  |  |  |  |  |  |  |  |  |  |  |  |  |  |  |  |  |  |
| --- | --- | --- | --- | --- | --- | --- | --- | --- | --- | --- | --- | --- | --- | --- | --- | --- | --- | --- | --- |
| pos | 884 | 4720 | 5192 | 7851 | 12793 | 13006 | 16914 | 17040 | 19677 | 21468 | 22027 | 22029 | 22030 | 22031 | 22032 | 22033 | 22034 | 24926 |  |
| ref | C | G | C | C | G | T | G | T | G | G | T | A | G | T | T | C | A | G |  |
| AY.4 | C | G | T | T | T | C | G | C | G | T | T | - | - | - | - | - | - | A |  |
| AY.4 | C | G | T | T | T | C | G | C | G | T | G | A | A | A | G | T | G | A | #607252 |
| AY.114 | T | T | C | C | G | T | T | T | T | G | G | A | A | A | G | T | G | G |  |
| mut |  |  |  |  |  |  |  |  |  |  |  |  |  |  |  |  |  | G>A |  |

|  |  |  |  |  |  |  |  |  |  |  |  |  |  |  |  |  |  |
| --- | --- | --- | --- | --- | --- | --- | --- | --- | --- | --- | --- | --- | --- | --- | --- | --- | --- |
| pos | 5812 | 9491 | 10802 | 11417 | 19684 | 22026 | 22027 | 22029 | 22030 | 22031 | 22032 | 22033 | 22034 | 23191 | 25904 | 28917 |  |
| ref | C | C | C | G | G | G | T | A | G | T | T | C | A | C | C | G |  |
| AY.100 | T | T | T | T | A | G | T | - | - | - | - | - | - | T | T | C |  |
| AY.100 | T | T | T | T | A | T | G | A | A | A | G | T | G | T | T | C | #607303 |
| AY.100 | C | C | C | G | G | T | G | A | A | A | G | T | G | C | C | G |  |
| mut |  |  |  |  |  |  |  |  |  |  |  |  |  | C>T | C>T | G>C |  |

|  |  |  |  |  |  |  |  |  |  |  |  |  |  |  |  |  |  |
| --- | --- | --- | --- | --- | --- | --- | --- | --- | --- | --- | --- | --- | --- | --- | --- | --- | --- |
| pos | 2706 | 4158 | 5365 | 11365 | 17040 | 18927 | 19810 | 20374 | 22027 | 22029 | 22030 | 22031 | 22032 | 22033 | 22034 | 23770 |  |
| ref | C | C | C | G | T | T | A | A | T | A | G | T | T | C | A | A |  |
| AY.4.1 | T | C | T | G | C | C | A | C | T | - | - | - | - | - | - | A |  |
| AY.4.1 | T | C | T | G | C | C | A | C | G | A | A | A | G | T | G | A | #610737 |
| AY.4 | C | T | C | T | T | T | C | A | G | A | A | A | G | T | G | G |  |
| mut |  |  |  |  |  |  |  |  |  |  |  |  |  |  |  | G>A |  |

|  |  |  |  |  |  |  |  |  |  |  |  |  |  |  |  |  |  |  |  |
| --- | --- | --- | --- | --- | --- | --- | --- | --- | --- | --- | --- | --- | --- | --- | --- | --- | --- | --- | --- |
| pos | 3924 | 6730 | 10317 | 17320 | 18831 | 22027 | 22029 | 22030 | 22031 | 22032 | 22033 | 22034 | 24919 | 29762 | 29769 | 29774 | 29779 | 29781 |  |
| ref | C | C | A | G | T | T | A | G | T | T | C | A | C | C | C | C | G | G |  |
| AY.4 | T | T | G | T | C | T | - | - | - | - | - | - | T | - | - | - | G | G |  |
| AY.4 | T | T | G | T | C | G | A | A | A | G | T | G | T | - | - | - | - | - | #610905 |
| AY.4 | C | C | A | G | T | G | A | A | A | G | T | G | C | C | C | C | G | G |  |
| mut |  |  |  |  |  |  |  |  |  |  |  |  | C>T | C>- | C>- | C>- | G>- | G>- |  |

|  |  |  |  |  |  |  |  |  |  |  |  |  |  |  |  |  |  |  |  |  |  |
| --- | --- | --- | --- | --- | --- | --- | --- | --- | --- | --- | --- | --- | --- | --- | --- | --- | --- | --- | --- | --- | --- |
| pos | 1348 | 7851 | 8016 | 9165 | 10323 | 12241 | 12772 | 15240 | 21811 | 21952 | 22026 | 22027 | 22029 | 22030 | 22031 | 22032 | 22033 | 22034 | 24926 | 26151 |  |
| ref | C | C | C | C | A | C | A | C | C | C | G | T | A | G | T | T | C | A | G | C |  |
| AY.113 | C | C | T | T | G | T | G | T | A | T | G | T | - | - | - | - | - | - | T | C |  |
| AY.113 | A | C | T | T | G | T | G | T | A | T | T | G | A | A | A | G | T | G | T | T | #611930 |
| AY.4 | C | T | C | C | A | C | A | C | C | C | T | G | A | A | A | G | T | G | G | C |  |
| mut | C>A |  |  |  |  |  |  |  |  |  |  |  |  |  |  |  |  |  | G>T | C>T |  |

|  |  |  |  |  |  |  |  |  |  |  |  |  |  |  |  |  |  |
| --- | --- | --- | --- | --- | --- | --- | --- | --- | --- | --- | --- | --- | --- | --- | --- | --- | --- |
| pos | 364 | 3304 | 3433 | 3877 | 3979 | 4370 | 16644 | 22026 | 22027 | 22029 | 22030 | 22031 | 22032 | 22033 | 22034 | 27424 |  |
| ref | C | G | G | T | A | G | A | G | T | A | G | T | T | C | A | A |  |
| AY.39 | T | T | T | C | G | G | G | G | T | - | - | - | - | - | - | A |  |
| AY.39 | T | T | T | C | G | A | G | T | G | A | A | A | G | T | G | G | #612000 |
| AY.39 | C | G | G | T | A | G | A | T | G | A | A | A | G | T | G | A |  |
| mut |  |  |  |  |  | G>A |  |  |  |  |  |  |  |  |  | A>G |  |

|  |  |  |  |  |  |  |  |  |  |  |  |  |  |  |  |  |
| --- | --- | --- | --- | --- | --- | --- | --- | --- | --- | --- | --- | --- | --- | --- | --- | --- |
| pos | 2324 | 4276 | 4917 | 11001 | 12756 | 19586 | 22027 | 22029 | 22030 | 22031 | 22032 | 22033 | 22034 | 27590 | 29700 |  |
| ref | G | C | T | C | C | C | T | A | G | T | T | C | A | C | A |  |
| AY.119 | A | T | C | T | T | T | T | - | - | - | - | - | - | T | A |  |
| AY.119 | A | T | C | T | T | T | G | A | A | A | G | T | G | T | A | #612966 |
| AY.119 | G | C | T | C | C | C | G | A | A | A | G | T | G | C | G |  |
| mut |  |  |  |  |  |  |  |  |  |  |  |  |  | C>T | G>A |  |

|  |  |  |  |  |  |  |  |  |  |  |  |  |  |  |  |
| --- | --- | --- | --- | --- | --- | --- | --- | --- | --- | --- | --- | --- | --- | --- | --- |
| pos | 4666 | 12786 | 15783 | 18324 | 19017 | 20238 | 22027 | 22029 | 22030 | 22031 | 22032 | 22033 | 22034 | 23638 |  |
| ref | A | C | C | T | C | G | T | A | G | T | T | C | A | C |  |
| AY.109 | G | T | T | C | T | T | T | - | - | - | - | - | - | T |  |
| AY.109 | G | T | T | C | T | T | G | A | A | A | G | T | G | T | #617001 |
| AY.109 | A | C | C | T | C | G | G | A | A | A | G | T | G | C |  |
| mut |  |  |  |  |  |  |  |  |  |  |  |  |  | C>T |  |

|  |  |  |  |  |  |  |  |  |  |  |  |  |  |  |  |  |  |  |  |  |  |  |
| --- | --- | --- | --- | --- | --- | --- | --- | --- | --- | --- | --- | --- | --- | --- | --- | --- | --- | --- | --- | --- | --- | --- |
| pos | 1085 | 1738 | 3304 | 3433 | 3559 | 8223 | 9979 | 15654 | 16644 | 17799 | 22026 | 22027 | 22029 | 22030 | 22031 | 22032 | 22033 | 22034 | 25726 | 27538 | 28310 |  |
| ref | G | G | G | G | A | A | C | C | A | A | G | T | A | G | T | T | C | A | G | C | C |  |
| AY.39 | T | T | T | T | A | G | C | C | G | G | G | T | - | - | - | - | - | - | G | T | T |  |
| AY.39 | T | T | T | T | A | G | C | C | G | G | G | G | A | A | A | G | T | G | T | C | T | #617042 |
| AY.39 | G | G | G | G | C | A | T | T | A | A | T | G | A | A | A | G | T | G | T | C | C |  |
| mut |  |  |  |  |  |  |  |  |  |  |  |  |  |  |  |  |  |  |  |  | C>T |  |

|  |  |  |  |  |  |  |  |  |  |  |  |  |  |  |  |  |  |  |  |
| --- | --- | --- | --- | --- | --- | --- | --- | --- | --- | --- | --- | --- | --- | --- | --- | --- | --- | --- | --- |
| pos | 1668 | 4320 | 4897 | 5192 | 5608 | 6681 | 7764 | 9363 | 9763 | 11767 | 22027 | 22029 | 22030 | 22031 | 22032 | 22033 | 22034 | 22729 |  |
| ref | A | C | C | C | A | C | C | C | C | C | T | A | G | T | T | C | A | T |  |
| AY.34.1 | G | T | T | C | A | T | T | C | T | C | T | - | - | - | - | - | - | C |  |
| AY.34.1 | G | T | T | C | A | T | T | C | T | C | G | A | A | A | G | T | G | C | #617064 |
| AY.34.1 | A | C | C | T | G | C | C | T | C | T | G | A | A | A | G | T | G | T |  |
| mut |  |  |  |  |  |  |  |  |  |  |  |  |  |  |  |  |  | T>C |  |

|  |  |  |  |  |  |  |  |  |  |  |  |  |  |  |
| --- | --- | --- | --- | --- | --- | --- | --- | --- | --- | --- | --- | --- | --- | --- |
| pos | 1074 | 16827 | 18985 | 20801 | 21987 | 22027 | 22029 | 22030 | 22031 | 22032 | 22033 | 22034 | 24138 |  |
| ref | G | G | G | C | G | T | A | G | T | T | C | A | C |  |
| AY.4.2 | A | A | T | T | A | T | - | - | - | - | - | - | C |  |
| AY.4.2 | A | A | T | T | A | G | A | A | A | G | T | G | T | #619112 |
| AY.4.2 | G | G | G | C | G | G | A | A | A | G | T | G | C |  |
| mut |  |  |  |  |  |  |  |  |  |  |  |  | C>T |  |

|  |  |  |  |  |  |  |  |  |  |  |  |  |  |  |  |  |  |  |
| --- | --- | --- | --- | --- | --- | --- | --- | --- | --- | --- | --- | --- | --- | --- | --- | --- | --- | --- |
| pos | 409 | 1659 | 3291 | 5661 | 16425 | 16733 | 21987 | 22027 | 22029 | 22030 | 22031 | 22032 | 22033 | 22034 | 25680 | 25855 | 27143 |  |
| ref | T | G | A | A | T | C | G | T | A | G | T | T | C | A | C | G | C |  |
| AY.4.2 | C | T | T | G | A | C | A | T | - | - | - | - | - | - | C | T | T |  |
| AY.4.2 | C | T | T | G | A | T | A | T | A | A | A | G | T | G | T | T | T | #619441 |
| AY.4.2 | T | G | A | A | T | C | G | G | A | A | A | G | T | G | C | G | C |  |
| mut |  |  |  |  |  | C>T |  |  |  |  |  |  |  |  | C>T | G>T | C>T |  |

|  |  |  |  |  |  |  |  |  |  |  |  |  |  |  |  |
| --- | --- | --- | --- | --- | --- | --- | --- | --- | --- | --- | --- | --- | --- | --- | --- |
| pos | 906 | 3994 | 7262 | 9611 | 10317 | 22027 | 22029 | 22030 | 22031 | 22032 | 22033 | 22034 | 25826 | 29026 |  |
| ref | C | G | C | C | A | T | A | G | T | T | C | A | A | T |  |
| AY.4 | T | T | T | T | G | T | - | - | - | - | - | - | A | C |  |
| AY.4 | T | T | T | T | G | G | A | A | A | G | T | G | T | C | #622234 |
| AY.4 | C | G | C | C | A | G | A | A | A | G | T | G | A | T |  |
| mut |  |  |  |  |  |  |  |  |  |  |  |  | A>T | T>C |  |

|  |  |  |  |  |  |  |  |  |  |  |  |  |  |  |  |  |  |  |
| --- | --- | --- | --- | --- | --- | --- | --- | --- | --- | --- | --- | --- | --- | --- | --- | --- | --- | --- |
| pos | 2755 | 3119 | 6701 | 15192 | 15720 | 18687 | 21987 | 22026 | 22027 | 22029 | 22030 | 22031 | 22032 | 22033 | 22034 | 23227 | 29266 |  |
| ref | G | G | C | A | C | C | G | G | T | A | G | T | T | C | A | T | G |  |
| AY.4.2 | T | G | T | G | T | T | A | G | T | - | - | - | - | - | - | T | G |  |
| AY.4.2 | T | G | T | G | T | T | A | T | G | A | A | A | G | T | G | C | T | #622572 |
| AY.4.2 | G | C | C | A | C | C | G | T | G | A | A | A | G | T | G | T | G |  |
| mut |  |  |  |  |  |  |  |  |  |  |  |  |  |  |  | T>C | G>T |  |

|  |  |  |  |  |  |  |  |  |  |  |  |  |  |  |  |  |  |  |  |
| --- | --- | --- | --- | --- | --- | --- | --- | --- | --- | --- | --- | --- | --- | --- | --- | --- | --- | --- | --- |
| pos | 556 | 1192 | 3361 | 5497 | 6501 | 10252 | 15720 | 18687 | 21668 | 21987 | 22027 | 22029 | 22030 | 22031 | 22032 | 22033 | 22034 | 28878 |  |
| ref | C | A | A | C | C | C | C | C | G | G | T | A | G | T | T | C | A | G |  |
| AY.4.2.1 | T | G | A | T | T | T | T | T | T | A | T | - | - | - | - | - | - | T |  |
| AY.4.2.1 | T | G | G | T | T | T | T | T | T | G | G | A | A | A | G | T | G | T | #622600 |
| AY.4.2 | C | A | A | C | C | C | C | C | G | G | G | A | A | A | G | T | G | G |  |
| mut |  |  | A>G |  |  |  |  |  |  | A>G |  |  |  |  |  |  |  | G>T |  |

|  |  |  |  |  |  |  |  |  |  |  |  |  |  |  |  |
| --- | --- | --- | --- | --- | --- | --- | --- | --- | --- | --- | --- | --- | --- | --- | --- |
| pos | 3119 | 3244 | 15150 | 18630 | 21987 | 22026 | 22027 | 22029 | 22030 | 22031 | 22032 | 22033 | 22034 | 28677 |  |
| ref | G | C | G | T | G | G | T | A | G | T | T | C | A | C |  |
| AY.4.2 | G | T | T | C | A | G | T | - | - | - | - | - | - | T |  |
| AY.4.2 | G | T | T | C | A | T | G | A | A | A | G | T | G | T | #622625 |
| AY.4.2 | C | C | G | T | G | T | G | A | A | A | G | T | G | C |  |
| mut |  |  |  |  |  |  |  |  |  |  |  |  |  | C>T |  |

|  |  |  |  |  |  |  |  |  |  |  |  |  |  |  |  |  |  |  |  |  |  |  |  |  |  |  |  |  |  |  |  |  |  |
| --- | --- | --- | --- | --- | --- | --- | --- | --- | --- | --- | --- | --- | --- | --- | --- | --- | --- | --- | --- | --- | --- | --- | --- | --- | --- | --- | --- | --- | --- | --- | --- | --- | --- |
| pos | 826 | 3816 | 4181 | 5184 | 5584 | 6402 | 7124 | 8752 | 8986 | 9053 | 9891 | 10029 | 11201 | 11332 | 11418 | 11514 | 13019 | 14487 | 19220 | 19846 | 21846 | 22227 | 23728 | 25352 | 26079 | 26527 | 27338 | 27345 | 27874 | 28657 | 28916 | 29167 |  |
| ref | T | A | G | C | A | C | C | C | C | G | C | C | A | A | T | C | C | A | C | G | C | C | T | G | C | C | A | A | C | C | G | C |  |
| AY.118 | C | G | T | A | A | T | T | C | T | T | C | T | G | G | T | C | C | A | T | G | T | C | C | G | C | T | T | A | T | T | T | C |  |
| B.1.617.2 | C | G | T | A | A | T | T | T | T | T | C | T | G | G | T | C | C | A | C | A | C | T | T | T | T | C | A | T | C | C | G | T | #622954 |
| AY.26 | T | A | G | T | G | C | C | C | C | G | T | C | A | A | C | T | T | G | C | A | C | T | T | T | T | C | A | T | C | C | G | C |  |
| mut |  |  |  |  |  |  |  | C>T |  |  |  |  |  |  |  |  |  |  |  |  |  |  |  |  |  |  |  |  |  |  |  | C>T |  |

|  |  |  |  |  |  |  |  |  |  |  |  |  |  |  |
| --- | --- | --- | --- | --- | --- | --- | --- | --- | --- | --- | --- | --- | --- | --- |
| pos | 1191 | 2623 | 3832 | 7086 | 19018 | 22027 | 22029 | 22030 | 22031 | 22032 | 22033 | 22034 | 26655 |  |
| ref | C | C | C | C | C | T | A | G | T | T | C | A | T |  |
| AY.105 | C | T | A | T | T | T | - | - | - | - | - | - | C |  |
| AY.105 | C | T | A | T | T | G | A | A | A | G | T | G | C | #623743 |
| AY.105 | T | C | C | C | C | G | A | A | A | G | T | G | T |  |
| mut |  |  |  |  |  |  |  |  |  |  |  |  | T>C |  |

|  |  |  |  |  |  |  |  |  |  |  |  |  |  |  |  |  |  |  |  |  |  |  |  |  |  |
| --- | --- | --- | --- | --- | --- | --- | --- | --- | --- | --- | --- | --- | --- | --- | --- | --- | --- | --- | --- | --- | --- | --- | --- | --- | --- |
| pos | 1588 | 2632 | 3476 | 4237 | 6341 | 6726 | 7393 | 7851 | 11836 | 15654 | 16323 | 16848 | 17040 | 21123 | 21372 | 22027 | 22029 | 22030 | 22031 | 22032 | 22033 | 22034 | 22311 | 27604 |  |
| ref | A | G | G | T | G | C | G | C | T | C | C | C | T | G | G | T | A | G | T | T | C | A | C | G |  |
| AY.4 | A | G | G | C | A | C | T | T | A | C | T | T | C | T | G | T | - | - | - | - | - | - | A | G |  |
| AY.4 | A | G | G | C | A | C | T | T | A | C | T | T | C | T | G | G | A | A | A | G | T | G | A | A | #625490 |
| AY.39 | G | T | A | T | G | T | G | C | T | T | C | C | T | G | T | G | A | A | A | G | T | G | C | A |  |
| mut |  |  |  |  |  |  |  |  |  |  |  |  |  |  |  |  |  |  |  |  |  |  | C>A |  |  |

|  |  |  |  |  |  |  |  |  |  |  |  |  |  |  |  |
| --- | --- | --- | --- | --- | --- | --- | --- | --- | --- | --- | --- | --- | --- | --- | --- |
| pos | 486 | 5724 | 22027 | 22029 | 22030 | 22031 | 22032 | 22033 | 22034 | 22104 | 22381 | 23244 | 28727 | 29315 |  |
| ref | C | C | T | A | G | T | T | C | A | G | G | C | G | G |  |
| AY.4 | C | C | G | A | A | A | G | T | G | G | G | C | G | G |  |
| AY.4 | T | T | G | A | A | A | G | T | G | T | T | A | T | C | #625516 |
| AY.4 | T | T | T | - | - | - | - | - | - | T | T | A | T | C |  |
| mut | C>T | C>T |  |  |  |  |  |  |  |  |  |  |  |  |  |

|  |  |  |  |  |  |  |  |  |  |  |  |  |  |  |
| --- | --- | --- | --- | --- | --- | --- | --- | --- | --- | --- | --- | --- | --- | --- |
| pos | 4237 | 7757 | 11564 | 14118 | 21987 | 21995 | 22027 | 22029 | 22030 | 22031 | 22032 | 22033 | 22034 |  |
| ref | T | A | C | G | G | T | T | A | G | T | T | C | A |  |
| AY.4 | C | C | A | T | A | T | T | - | - | - | - | - | - |  |
| AY.4 | C | C | A | T | A | T | G | A | A | A | G | T | G | #625562 |
| AY.4 | T | A | C | G | G | C | G | A | A | A | G | T | G |  |
| mut |  |  |  |  |  |  |  |  |  |  |  |  |  |  |

|  |  |  |  |  |  |  |  |  |  |  |  |  |  |  |  |  |  |  |  |  |  |
| --- | --- | --- | --- | --- | --- | --- | --- | --- | --- | --- | --- | --- | --- | --- | --- | --- | --- | --- | --- | --- | --- |
| pos | 884 | 2246 | 4720 | 6091 | 7851 | 11539 | 12565 | 16914 | 17040 | 17694 | 19677 | 22027 | 22029 | 22030 | 22031 | 22032 | 22033 | 22034 | 27657 | 28483 |  |
| ref | C | G | G | C | C | T | G | G | T | T | G | T | A | G | T | T | C | A | C | A |  |
| AY.4 | C | A | G | T | T | C | A | G | C | G | G | T | - | - | - | - | - | - | T | T |  |
| AY.4 | C | A | G | T | T | C | A | G | C | G | G | G | A | A | A | G | T | G | T | T | #625582 |
| AY.114 | T | G | T | C | C | T | G | T | T | T | T | G | A | A | A | G | T | G | C | A |  |
| mut |  |  |  |  |  |  |  |  |  |  |  |  |  |  |  |  |  |  | C>T | A>T |  |

|  |  |  |  |  |  |  |  |  |  |  |  |  |  |  |  |  |  |  |  |
| --- | --- | --- | --- | --- | --- | --- | --- | --- | --- | --- | --- | --- | --- | --- | --- | --- | --- | --- | --- |
| pos | 6539 | 7393 | 8299 | 12817 | 15254 | 19983 | 20935 | 21987 | 22027 | 22029 | 22030 | 22031 | 22032 | 22033 | 22034 | 22458 | 27389 | 27440 |  |
| ref | C | G | C | G | T | C | A | G | T | A | G | T | T | C | A | C | C | A |  |
| AY.4.2.2 | C | T | C | T | G | C | G | A | T | - | - | - | - | - | - | T | C | G |  |
| AY.4.2.2 | C | T | C | T | G | C | G | A | G | A | A | A | G | T | G | T | C | G | #625795 |
| AY.4.2.2 | T | G | T | G | T | T | A | G | G | A | A | A | G | T | G | C | G | A |  |
| mut |  |  |  |  |  |  |  |  |  |  |  |  |  |  |  | C>T | G>C | A>G |  |

|  |  |  |  |  |  |  |  |  |  |  |  |  |  |  |  |  |  |  |  |  |
| --- | --- | --- | --- | --- | --- | --- | --- | --- | --- | --- | --- | --- | --- | --- | --- | --- | --- | --- | --- | --- |
| pos | 1900 | 2147 | 6022 | 6040 | 6638 | 7926 | 8956 | 14014 | 16726 | 17236 | 18657 | 24208 | 25452 | 27005 | 27999 | 28073 | 28460 | 29593 | 29700 |  |
| ref | A | C | A | C | C | C | C | T | C | A | C | C | C | C | C | G | G | G | A |  |
| AY.103 | G | T | G | C | C | C | T | T | C | G | T | T | T | T | T | G | G | G | A |  |
| AY.103 | G | T | G | C | C | C | T | T | C | G | T | C | C | C | C | T | A | T | G | #627114 |
| AY.44 | A | C | A | T | T | T | C | G | T | A | C | C | C | C | C | T | A | T | G |  |
| mut |  |  |  |  |  |  |  |  |  |  |  |  |  |  |  |  |  |  |  |  |

|  |  |  |  |  |  |  |  |  |  |  |  |  |  |  |  |  |  |  |  |  |  |  |  |  |
| --- | --- | --- | --- | --- | --- | --- | --- | --- | --- | --- | --- | --- | --- | --- | --- | --- | --- | --- | --- | --- | --- | --- | --- | --- |
| pos | 2062 | 15951 | 21987 | 22026 | 22027 | 22029 | 22030 | 22031 | 22032 | 22033 | 22034 | 22242 | 24412 | 25537 | 25599 | 26214 | 29700 | 29762 | 29769 | 29774 | 29779 | 29781 | 29838 |  |
| ref | C | C | G | G | T | A | G | T | T | C | A | T | T | G | G | T | A | C | C | C | G | G | C |  |
| AY.39 | T | C | A | T | G | A | A | A | G | T | G | T | T | G | T | T | A | C | C | C | G | G | C |  |
| AY.39 | C | C | A | T | G | A | A | A | G | T | G | C | C | T | G | C | A | C | C | C | G | G | T | #627628 |
| AY.39 | C | T | G | G | T | - | - | - | - | - | - | C | C | T | G | C | - | - | - | - | - | - | C |  |
| mut | T>C |  |  |  |  |  |  |  |  |  |  |  |  |  |  |  | ->A | ->C | ->C | ->C | ->G | ->G | C>T |  |

|  |  |  |  |  |  |  |  |  |  |  |  |  |  |  |  |
| --- | --- | --- | --- | --- | --- | --- | --- | --- | --- | --- | --- | --- | --- | --- | --- |
| pos | 3838 | 4393 | 19185 | 19275 | 19532 | 22027 | 22029 | 22030 | 22031 | 22032 | 22033 | 22034 | 23323 | 27674 |  |
| ref | G | A | C | T | A | T | A | G | T | T | C | A | T | A |  |
| AY.4 | T | C | T | A | G | T | - | - | - | - | - | - | T | T |  |
| AY.4 | T | C | T | A | G | G | A | A | A | G | T | G | C | T | #630236 |
| AY.4 | G | A | C | T | A | G | A | A | A | G | T | G | T | A |  |
| mut |  |  |  |  |  |  |  |  |  |  |  |  | T>C | A>T |  |

|  |  |  |  |  |  |  |  |  |  |  |  |  |  |  |  |  |  |  |
| --- | --- | --- | --- | --- | --- | --- | --- | --- | --- | --- | --- | --- | --- | --- | --- | --- | --- | --- |
| pos | 5512 | 5730 | 10795 | 11455 | 12129 | 13953 | 16896 | 18525 | 20752 | 22027 | 22029 | 22030 | 22031 | 22032 | 22033 | 22034 | 26415 |  |
| ref | C | C | T | C | C | A | T | G | G | T | A | G | T | T | C | A | C |  |
| AY.126 | T | T | T | T | A | G | C | T | G | T | - | - | - | - | - | - | T |  |
| AY.126 | T | T | A | T | A | G | C | T | T | G | A | A | A | G | T | G | C | #630390 |
| AY.126 | C | C | T | C | C | A | T | G | G | G | A | A | A | G | T | G | C |  |
| mut |  |  | T>A |  |  |  |  |  | G>T |  |  |  |  |  |  |  |  |  |

|  |  |  |  |  |  |  |  |  |  |  |  |  |  |  |  |
| --- | --- | --- | --- | --- | --- | --- | --- | --- | --- | --- | --- | --- | --- | --- | --- |
| pos | 177 | 691 | 2317 | 3937 | 4249 | 21310 | 21987 | 22027 | 22029 | 22030 | 22031 | 22032 | 22033 | 22034 |  |
| ref | T | A | A | G | T | C | G | T | A | G | T | T | C | A |  |
| AY.4.2 | C | T | G | G | C | C | A | T | - | - | - | - | - | - |  |
| AY.4.2 | C | T | G | C | C | A | G | G | A | A | A | G | T | G | #630568 |
| AY.4.2 | T | A | A | G | T | C | G | G | A | A | A | G | T | G |  |
| mut |  |  |  | G>C |  | C>A | A>G |  |  |  |  |  |  |  |  |

|  |  |  |  |  |  |  |  |  |  |  |  |  |  |  |  |  |  |
| --- | --- | --- | --- | --- | --- | --- | --- | --- | --- | --- | --- | --- | --- | --- | --- | --- | --- |
| pos | 221 | 6196 | 7945 | 8356 | 10642 | 17622 | 18264 | 21987 | 22027 | 22029 | 22030 | 22031 | 22032 | 22033 | 22034 | 29001 |  |
| ref | T | C | C | T | G | A | C | G | T | A | G | T | T | C | A | G |  |
| AY.4.2 | G | T | T | C | T | G | T | A | T | - | - | - | - | - | - | A |  |
| AY.4.2 | G | T | T | C | T | G | T | A | G | A | A | A | G | T | G | G | #630591 |
| AY.4.2 | T | C | C | T | G | A | C | G | G | A | A | A | G | T | G | G |  |
| mut |  |  |  |  |  |  |  |  |  |  |  |  |  |  |  |  |  |

|  |  |  |  |  |  |  |  |  |  |  |  |  |  |
| --- | --- | --- | --- | --- | --- | --- | --- | --- | --- | --- | --- | --- | --- |
| pos | 177 | 691 | 2317 | 21987 | 22026 | 22027 | 22029 | 22030 | 22031 | 22032 | 22033 | 22034 |  |
| ref | T | A | A | G | G | T | A | G | T | T | C | A |  |
| AY.4.2 | C | T | G | A | G | T | - | - | - | - | - | - |  |
| AY.4.2 | C | T | G | G | T | G | A | A | A | G | T | G | #630600 |
| AY.4.2 | T | A | A | G | T | G | A | A | A | G | T | G |  |
| mut |  |  |  | A>G |  |  |  |  |  |  |  |  |  |

|  |  |  |  |  |  |  |  |  |  |  |  |  |  |  |  |  |  |
| --- | --- | --- | --- | --- | --- | --- | --- | --- | --- | --- | --- | --- | --- | --- | --- | --- | --- |
| pos | 221 | 6196 | 7945 | 8356 | 10642 | 17622 | 18264 | 21987 | 22027 | 22029 | 22030 | 22031 | 22032 | 22033 | 22034 | 29001 |  |
| ref | T | C | C | T | G | A | C | G | T | A | G | T | T | C | A | G |  |
| AY.4.2 | G | T | T | C | T | G | T | A | T | - | - | - | - | - | - | A |  |
| AY.4.2 | G | T | T | C | T | G | T | A | T | A | A | A | G | T | G | G | #630661 |
| AY.4.2 | T | C | C | T | G | A | C | G | G | A | A | A | G | T | G | G |  |
| mut |  |  |  |  |  |  |  |  |  |  |  |  |  |  |  |  |  |

|  |  |  |  |  |  |  |  |  |  |  |  |  |  |  |  |
| --- | --- | --- | --- | --- | --- | --- | --- | --- | --- | --- | --- | --- | --- | --- | --- |
| pos | 247 | 1832 | 4510 | 5560 | 6317 | 22027 | 22029 | 22030 | 22031 | 22032 | 22033 | 22034 | 24865 | 27749 |  |
| ref | G | A | G | A | C | T | A | G | T | T | C | A | C | A |  |
| AY.39 | G | G | G | G | T | T | - | - | - | - | - | - | A | A |  |
| AY.39 | G | G | G | G | T | G | A | A | A | G | T | G | A | A | #631610 |
| AY.39 | A | A | T | A | C | G | A | A | A | G | T | G | C | C |  |
| mut |  |  |  |  |  |  |  |  |  |  |  |  | C>A | C>A |  |

|  |  |  |  |  |  |  |  |  |  |  |  |  |  |  |  |  |  |  |  |  |  |  |  |  |
| --- | --- | --- | --- | --- | --- | --- | --- | --- | --- | --- | --- | --- | --- | --- | --- | --- | --- | --- | --- | --- | --- | --- | --- | --- |
| pos | 341 | 884 | 4720 | 5473 | 9339 | 13188 | 16914 | 17974 | 18086 | 18687 | 19677 | 19862 | 20555 | 21786 | 22027 | 22029 | 22030 | 22031 | 22032 | 22033 | 22034 | 25273 | 29227 |  |
| ref | G | C | G | T | A | T | G | G | C | C | G | C | C | G | T | A | G | T | T | C | A | G | G |  |
| B.1.617.2 | A | C | G | C | G | C | G | G | T | T | G | T | T | T | T | - | - | - | - | - | - | T | T |  |
| B.1.617.2 | A | C | G | C | G | C | G | A | T | T | G | T | T | T | G | A | A | A | G | T | G | T | T | #631615 |
| AY.114 | G | T | T | T | A | T | T | G | C | C | T | C | C | G | G | A | A | A | G | T | G | G | G |  |
| mut |  |  |  |  |  |  |  | G>A |  |  |  |  |  |  |  |  |  |  |  |  |  | G>T | G>T |  |

|  |  |  |  |  |  |  |  |  |  |  |  |  |  |  |  |  |  |
| --- | --- | --- | --- | --- | --- | --- | --- | --- | --- | --- | --- | --- | --- | --- | --- | --- | --- |
| pos | 3660 | 6250 | 10271 | 11138 | 12513 | 12528 | 13446 | 15654 | 22026 | 22027 | 22029 | 22030 | 22031 | 22032 | 22033 | 22034 |  |
| ref | A | C | G | G | C | C | C | C | G | T | A | G | T | T | C | A |  |
| AY.118 | G | T | G | T | T | T | T | C | G | T | - | - | - | - | - | - |  |
| AY.118 | G | T | G | T | T | T | T | A | T | G | A | A | A | G | T | G | #631765 |
| AY.118 | A | C | A | G | C | C | C | C | T | G | A | A | A | G | T | G |  |
| mut |  |  |  |  |  |  |  | C>A |  |  |  |  |  |  |  |  |  |

|  |  |  |  |  |  |  |  |  |  |  |  |  |  |  |  |  |  |  |  |  |  |  |  |  |
| --- | --- | --- | --- | --- | --- | --- | --- | --- | --- | --- | --- | --- | --- | --- | --- | --- | --- | --- | --- | --- | --- | --- | --- | --- |
| pos | 1588 | 2231 | 2271 | 2632 | 2946 | 3304 | 3433 | 3476 | 6726 | 8642 | 8655 | 13929 | 15654 | 18167 | 18756 | 22027 | 22029 | 22030 | 22031 | 22032 | 22033 | 22034 | 27916 |  |
| ref | A | G | A | G | G | G | G | G | C | G | C | T | C | C | G | T | A | G | T | T | C | A | G |  |
| AY.39 | A | T | G | G | A | T | T | G | C | A | T | C | C | A | T | T | - | - | - | - | - | - | A |  |
| AY.39 | A | T | G | G | A | T | T | G | C | A | T | C | C | A | T | G | A | A | A | G | T | G | A | #631794 |
| AY.39 | G | G | A | T | G | G | G | A | T | G | C | T | T | C | G | G | A | A | A | G | T | G | G |  |
| mut |  |  |  |  |  |  |  |  |  |  |  |  |  |  |  |  |  |  |  |  |  |  | G>A |  |

|  |  |  |  |  |  |  |  |  |  |  |  |  |  |  |  |
| --- | --- | --- | --- | --- | --- | --- | --- | --- | --- | --- | --- | --- | --- | --- | --- |
| pos | 204 | 564 | 2447 | 2516 | 10507 | 18652 | 21987 | 22027 | 22029 | 22030 | 22031 | 22032 | 22033 | 22034 |  |
| ref | G | G | G | G | C | C | G | T | A | G | T | T | C | A |  |
| AY.4 | G | A | T | T | C | T | A | T | - | - | - | - | - | - |  |
| AY.4 | G | A | T | T | C | T | A | G | A | A | A | G | T | G | #633813 |
| AY.4 | T | G | G | G | T | C | G | G | A | A | A | G | T | G |  |
| mut |  |  |  |  |  |  |  |  |  |  |  |  |  |  |  |

|  |  |  |  |  |  |  |  |  |  |  |  |  |  |  |  |  |  |  |  |
| --- | --- | --- | --- | --- | --- | --- | --- | --- | --- | --- | --- | --- | --- | --- | --- | --- | --- | --- | --- |
| pos | 1599 | 5221 | 7125 | 10323 | 10624 | 12872 | 15720 | 18687 | 21668 | 21987 | 22027 | 22029 | 22030 | 22031 | 22032 | 22033 | 22034 | 28878 |  |
| ref | G | C | C | A | A | G | C | C | G | G | T | A | G | T | T | C | A | G |  |
| AY.4.2.1 | G | T | T | G | G | A | T | T | T | A | T | - | - | - | - | - | - | T |  |
| AY.4.2.1 | T | T | T | G | G | A | T | T | T | G | G | A | A | A | G | T | G | T | #633937 |
| AY.4.2 | G | C | C | A | A | G | C | C | G | G | G | A | A | A | G | T | G | G |  |
| mut | G>T |  |  |  |  |  |  |  |  | A>G |  |  |  |  |  |  |  | G>T |  |

|  |  |  |  |  |  |  |  |  |  |  |  |  |  |  |  |  |  |
| --- | --- | --- | --- | --- | --- | --- | --- | --- | --- | --- | --- | --- | --- | --- | --- | --- | --- |
| pos | 208 | 3304 | 3433 | 7299 | 10376 | 16644 | 22026 | 22027 | 22029 | 22030 | 22031 | 22032 | 22033 | 22034 | 25166 | 28744 |  |
| ref | G | G | G | C | C | A | G | T | A | G | T | T | C | A | G | C |  |
| AY.39 | T | T | T | T | T | G | G | T | - | - | - | - | - | - | A | T |  |
| AY.39 | T | T | T | T | T | G | T | G | A | A | A | G | T | G | G | T | #635291 |
| AY.39 | G | G | G | C | C | A | T | G | A | A | A | G | T | G | G | C |  |
| mut |  |  |  |  |  |  |  |  |  |  |  |  |  |  |  | C>T |  |

|  |  |  |  |  |  |  |  |  |  |  |  |  |  |  |  |  |
| --- | --- | --- | --- | --- | --- | --- | --- | --- | --- | --- | --- | --- | --- | --- | --- | --- |
| pos | 1264 | 2232 | 3811 | 6629 | 10615 | 22026 | 22027 | 22029 | 22030 | 22031 | 22032 | 22033 | 22034 | 25720 | 28382 |  |
| ref | G | C | C | C | C | G | T | A | G | T | T | C | A | G | T |  |
| AY.100 | T | T | T | T | T | G | T | - | - | - | - | - | - | T | C |  |
| AY.100 | T | T | T | T | T | T | G | A | A | A | G | T | G | T | C | #635304 |
| AY.100 | G | C | C | C | C | T | G | A | A | A | G | T | G | G | T |  |
| mut |  |  |  |  |  |  |  |  |  |  |  |  |  | G>T | T>C |  |

|  |  |  |  |  |  |  |  |  |  |  |  |  |  |  |  |  |
| --- | --- | --- | --- | --- | --- | --- | --- | --- | --- | --- | --- | --- | --- | --- | --- | --- |
| pos | 1204 | 1394 | 5784 | 12754 | 20178 | 20450 | 21974 | 22027 | 22029 | 22030 | 22031 | 22032 | 22033 | 22034 | 25599 |  |
| ref | C | G | C | C | C | A | G | T | A | G | T | T | C | A | G |  |
| AY.4 | T | G | C | T | C | G | C | T | - | - | - | - | - | - | G |  |
| AY.4 | T | G | C | T | T | G | C | G | A | A | A | G | T | G | T | #637023 |
| AY.4 | C | A | T | C | C | A | G | G | A | A | A | G | T | G | T |  |
| mut |  |  |  |  | C>T |  |  |  |  |  |  |  |  |  |  |  |

|  |  |  |  |  |  |  |  |  |  |  |  |  |  |  |  |
| --- | --- | --- | --- | --- | --- | --- | --- | --- | --- | --- | --- | --- | --- | --- | --- |
| pos | 5301 | 6896 | 10687 | 18366 | 21641 | 22026 | 22027 | 22029 | 22030 | 22031 | 22032 | 22033 | 22034 | 23282 |  |
| ref | C | C | T | A | G | G | T | A | G | T | T | C | A | G |  |
| AY.100 | T | T | C | G | T | G | T | - | - | - | - | - | - | T |  |
| AY.100 | T | T | C | G | T | G | G | A | A | A | G | T | G | G | #639061 |
| AY.100 | C | C | T | A | G | T | G | A | A | A | G | T | G | G |  |
| mut |  |  |  |  |  |  |  |  |  |  |  |  |  |  |  |

|  |  |  |  |  |  |  |  |  |  |  |  |  |  |  |  |  |  |  |  |  |  |
| --- | --- | --- | --- | --- | --- | --- | --- | --- | --- | --- | --- | --- | --- | --- | --- | --- | --- | --- | --- | --- | --- |
| pos | 884 | 3738 | 4068 | 4720 | 5071 | 7989 | 10369 | 15407 | 16914 | 19677 | 20104 | 21846 | 22027 | 22029 | 22030 | 22031 | 22032 | 22033 | 22034 | 25740 |  |
| ref | C | C | C | G | T | C | C | C | G | G | C | C | T | A | G | T | T | C | A | G |  |
| B.1.617.2 | C | T | C | G | C | C | T | T | G | G | T | C | T | - | - | - | - | - | - | T |  |
| B.1.617.2 | C | T | T | G | C | T | T | T | G | G | T | C | G | A | A | A | G | T | G | T | #640848 |
| AY.114 | T | C | C | T | T | C | C | C | T | T | C | T | G | A | A | A | G | T | G | G |  |
| mut |  |  | C>T |  |  | C>T |  |  |  |  |  |  |  |  |  |  |  |  |  | G>T |  |

|  |  |  |  |  |  |  |  |  |  |  |  |  |  |  |  |  |  |  |  |  |
| --- | --- | --- | --- | --- | --- | --- | --- | --- | --- | --- | --- | --- | --- | --- | --- | --- | --- | --- | --- | --- |
| pos | 186 | 6336 | 7121 | 7860 | 21987 | 22027 | 22029 | 22030 | 22031 | 22032 | 22033 | 22034 | 25387 | 25613 | 27478 | 27526 | 27689 | 28312 | 29108 |  |
| ref | C | C | A | C | G | T | A | G | T | T | C | A | G | C | G | C | C | C | C |  |
| AY.4 | C | C | A | C | G | G | A | A | A | G | T | G | G | C | G | C | C | C | C |  |
| AY.4 | T | C | G | C | G | G | A | A | A | G | T | G | C | T | T | T | T | T | T | #640989 |
| AY.4 | T | T | G | A | A | T | - | - | - | - | - | - | G | T | T | T | T | T | T |  |
| mut | C>T |  | A>G |  |  |  |  |  |  |  |  |  | G>C |  |  |  |  |  |  |  |

|  |  |  |  |  |  |  |  |  |  |  |  |  |  |  |  |  |
| --- | --- | --- | --- | --- | --- | --- | --- | --- | --- | --- | --- | --- | --- | --- | --- | --- |
| pos | 2110 | 6474 | 7195 | 14910 | 15857 | 17410 | 21987 | 22027 | 22029 | 22030 | 22031 | 22032 | 22033 | 22034 | 29763 |  |
| ref | C | A | C | T | C | C | G | T | A | G | T | T | C | A | A |  |
| AY.4.2 | T | C | T | C | T | T | A | T | - | - | - | - | - | - | G |  |
| AY.4.2 | T | C | T | C | T | T | A | T | A | A | A | G | T | G | A | #641092 |
| AY.4.2 | C | A | C | T | C | C | G | G | A | A | A | G | T | G | A |  |
| mut |  |  |  |  |  |  |  |  |  |  |  |  |  |  |  |  |

|  |  |  |  |  |  |  |  |  |  |  |  |  |  |  |  |  |
| --- | --- | --- | --- | --- | --- | --- | --- | --- | --- | --- | --- | --- | --- | --- | --- | --- |
| pos | 1391 | 5886 | 6040 | 7739 | 21987 | 22026 | 22027 | 22029 | 22030 | 22031 | 22032 | 22033 | 22034 | 23675 | 29221 |  |
| ref | T | C | C | G | G | G | T | A | G | T | T | C | A | G | A |  |
| AY.4.2 | C | T | T | A | A | G | T | - | - | - | - | - | - | G | T |  |
| AY.4.2 | C | T | T | A | G | T | G | A | A | A | G | T | G | T | T | #641140 |
| AY.4.2 | T | C | C | G | G | T | G | A | A | A | G | T | G | G | A |  |
| mut |  |  |  |  | A>G |  |  |  |  |  |  |  |  | G>T | A>T |  |

|  |  |  |  |  |  |  |  |  |  |  |  |  |  |  |  |  |
| --- | --- | --- | --- | --- | --- | --- | --- | --- | --- | --- | --- | --- | --- | --- | --- | --- |
| pos | 1268 | 1431 | 5079 | 6726 | 10156 | 10519 | 11322 | 22027 | 22029 | 22030 | 22031 | 22032 | 22033 | 22034 | 27525 |  |
| ref | G | A | C | C | C | C | C | T | A | G | T | T | C | A | A |  |
| AY.4 | A | T | T | T | T | T | T | T | - | - | - | - | - | - | G |  |
| AY.4 | A | T | T | T | T | T | T | G | A | A | A | G | T | G | G | #642397 |
| AY.4 | G | A | C | C | C | C | C | G | A | A | A | G | T | G | A |  |
| mut |  |  |  |  |  |  |  |  |  |  |  |  |  |  | A>G |  |

|  |  |  |  |  |  |  |  |  |  |  |  |  |  |  |  |  |
| --- | --- | --- | --- | --- | --- | --- | --- | --- | --- | --- | --- | --- | --- | --- | --- | --- |
| pos | 5422 | 11173 | 11280 | 11789 | 12076 | 16658 | 19118 | 22026 | 22027 | 22029 | 22030 | 22031 | 22032 | 22033 | 22034 |  |
| ref | T | C | C | A | C | C | C | G | T | A | G | T | T | C | A |  |
| AY.100 | C | T | A | G | T | A | T | G | T | - | - | - | - | - | - |  |
| AY.100 | C | T | A | G | T | A | T | G | G | A | A | A | G | T | G | #642462 |
| AY.100 | T | C | C | A | C | C | C | T | G | A | A | A | G | T | G |  |
| mut |  |  |  |  |  |  |  |  |  |  |  |  |  |  |  |  |

|  |  |  |  |  |  |  |  |  |  |  |  |  |  |  |  |
| --- | --- | --- | --- | --- | --- | --- | --- | --- | --- | --- | --- | --- | --- | --- | --- |
| pos | 247 | 4510 | 22027 | 22029 | 22030 | 22031 | 22032 | 22033 | 22034 | 23159 | 24675 | 26777 | 27749 | 29108 |  |
| ref | G | G | T | A | G | T | T | C | A | T | A | T | A | C |  |
| AY.39 | A | T | G | A | A | A | G | T | G | T | A | T | C | C |  |
| AY.39 | G | G | G | A | A | A | G | T | G | G | C | C | A | A | #642537 |
| AY.39 | G | G | T | - | - | - | - | - | - | G | C | C | A | A |  |
| mut | A>G | T>G |  |  |  |  |  |  |  |  |  |  |  |  |  |

|  |  |  |  |  |  |  |  |  |  |  |  |  |
| --- | --- | --- | --- | --- | --- | --- | --- | --- | --- | --- | --- | --- |
| pos | 1407 | 6408 | 6709 | 7049 | 8208 | 15199 | 25549 | 25614 | 27654 | 27900 | 28146 |  |
| ref | A | C | A | T | C | G | C | C | C | T | C |  |
| AY.4.2 | A | C | C | C | C | G | C | T | C | T | T |  |
| AY.4.2 | A | C | C | C | C | T | T | C | T | C | C | #643701 |
| AY.4.2 | G | T | A | T | T | T | T | C | T | C | C |  |
| mut |  |  |  |  |  |  |  |  |  |  |  |  |

|  |  |  |  |  |  |  |  |  |  |  |  |  |  |  |
| --- | --- | --- | --- | --- | --- | --- | --- | --- | --- | --- | --- | --- | --- | --- |
| pos | 945 | 1264 | 2453 | 10986 | 21987 | 22027 | 22029 | 22030 | 22031 | 22032 | 22033 | 22034 | 28077 |  |
| ref | G | G | C | G | G | T | A | G | T | T | C | A | G |  |
| AY.4 | A | T | T | T | A | T | - | - | - | - | - | - | T |  |
| AY.4 | A | T | T | T | A | G | A | A | A | G | T | G | T | #644206 |
| AY.4 | G | G | C | G | G | G | A | A | A | G | T | G | G |  |
| mut |  |  |  |  |  |  |  |  |  |  |  |  | G>T |  |

|  |  |  |  |  |  |  |  |  |  |  |  |  |  |  |  |  |  |  |  |  |  |  |
| --- | --- | --- | --- | --- | --- | --- | --- | --- | --- | --- | --- | --- | --- | --- | --- | --- | --- | --- | --- | --- | --- | --- |
| pos | 884 | 3950 | 4720 | 5164 | 7851 | 12085 | 15921 | 16075 | 16914 | 19677 | 21746 | 22027 | 22029 | 22030 | 22031 | 22032 | 22033 | 22034 | 25855 | 27907 | 28340 |  |
| ref | C | G | G | G | C | C | G | G | G | G | G | T | A | G | T | T | C | A | G | T | T |  |
| AY.4 | C | A | G | G | T | T | T | T | G | G | A | T | - | - | - | - | - | - | T | C | C |  |
| AY.4 | C | A | G | C | T | T | T | T | G | G | A | G | A | A | A | G | T | G | T | C | C | #644244 |
| AY.114 | T | G | T | G | C | C | G | G | T | T | G | G | A | A | A | G | T | G | G | T | T |  |
| mut |  |  |  | G>C |  |  |  |  |  |  |  |  |  |  |  |  |  |  | G>T | T>C | T>C |  |

|  |  |  |  |  |  |  |  |  |  |  |  |  |  |  |  |  |  |
| --- | --- | --- | --- | --- | --- | --- | --- | --- | --- | --- | --- | --- | --- | --- | --- | --- | --- |
| pos | 270 | 3119 | 7735 | 7958 | 9139 | 16231 | 21987 | 22027 | 22029 | 22030 | 22031 | 22032 | 22033 | 22034 | 22918 | 27493 |  |
| ref | A | G | C | A | T | T | G | T | A | G | T | T | C | A | G | C |  |
| AY.4.2 | G | C | T | G | C | T | A | T | - | - | - | - | - | - | A | C |  |
| AY.4.2 | G | C | T | G | C | C | A | G | A | A | A | G | T | G | G | T | #644512 |
| AY.4.2 | A | G | C | A | T | T | G | G | A | A | A | G | T | G | G | C |  |
| mut |  |  |  |  |  | T>C |  |  |  |  |  |  |  |  |  | C>T |  |

|  |  |  |  |  |  |  |  |  |  |  |  |  |  |  |  |  |  |  |  |  |  |  |
| --- | --- | --- | --- | --- | --- | --- | --- | --- | --- | --- | --- | --- | --- | --- | --- | --- | --- | --- | --- | --- | --- | --- |
| pos | 835 | 884 | 4720 | 6990 | 7851 | 8025 | 8821 | 10870 | 14407 | 16914 | 17249 | 17797 | 19677 | 22027 | 22029 | 22030 | 22031 | 22032 | 22033 | 22034 | 27659 |  |
| ref | C | C | G | C | C | C | A | G | C | G | C | G | G | T | A | G | T | T | C | A | G |  |
| AY.4 | T | C | G | C | T | T | G | T | C | G | T | A | G | T | - | - | - | - | - | - | T |  |
| AY.4 | T | C | G | T | T | T | G | T | T | G | T | A | G | G | A | A | A | G | T | G | G | #644568 |
| AY.114 | C | T | T | C | C | C | A | G | C | T | C | G | T | G | A | A | A | G | T | G | G |  |
| mut |  |  |  | C>T |  |  |  |  | C>T |  |  |  |  |  |  |  |  |  |  |  |  |  |

|  |  |  |  |  |  |  |  |  |  |  |  |  |  |  |  |  |  |  |  |  |  |  |  |  |  |  |  |  |  |  |  |  |  |
| --- | --- | --- | --- | --- | --- | --- | --- | --- | --- | --- | --- | --- | --- | --- | --- | --- | --- | --- | --- | --- | --- | --- | --- | --- | --- | --- | --- | --- | --- | --- | --- | --- | --- |
| pos | 529 | 2061 | 2277 | 2974 | 8112 | 9319 | 10647 | 11456 | 18090 | 19684 | 21137 | 21302 | 21304 | 21305 | 21846 | 21987 | 23758 | 24872 | 25710 | 27297 | 27322 | 28692 | 29050 | 29179 | 29509 | 29628 | 29700 | 29762 | 29769 | 29774 | 29779 | 29781 |  |
| ref | G | C | T | T | C | C | C | A | G | G | A | C | C | G | C | G | C | G | C | C | T | A | G | G | C | G | A | C | C | C | G | G |  |
| AY.3 | G | C | T | T | T | C | C | G | G | A | G | C | C | G | C | A | C | G | T | C | C | C | A | G | T | T | - | - | - | - | - | - |  |
| B.1.617.2 | G | C | T | T | T | C | C | G | G | A | G | C | C | G | C | A | C | T | C | T | T | A | G | T | C | G | A | C | C | C | G | G | #644638 |
| AY.20 | T | T | C | C | C | T | T | A | T | G | A | T | A | A | T | G | T | T | C | T | T | A | G | T | C | G | A | C | C | C | G | - |  |
| mut |  |  |  |  |  |  |  |  |  |  |  |  |  |  |  |  |  |  |  |  |  |  |  |  |  |  |  |  |  |  |  | ->G |  |

|  |  |  |  |  |  |  |  |  |  |  |  |  |  |  |
| --- | --- | --- | --- | --- | --- | --- | --- | --- | --- | --- | --- | --- | --- | --- |
| pos | 7869 | 10969 | 13812 | 15315 | 19972 | 22026 | 22027 | 22029 | 22030 | 22031 | 22032 | 22033 | 22034 |  |
| ref | C | C | G | C | C | G | T | A | G | T | T | C | A |  |
| AY.34.1 | T | C | T | C | T | G | T | - | - | - | - | - | - |  |
| AY.34.1 | T | C | T | C | T | T | G | A | A | A | G | T | G | #645408 |
| AY.34.1 | C | T | G | T | C | T | G | A | A | A | G | T | G |  |
| mut |  |  |  |  |  |  |  |  |  |  |  |  |  |  |

|  |  |  |  |  |  |  |  |  |  |  |  |  |  |  |  |
| --- | --- | --- | --- | --- | --- | --- | --- | --- | --- | --- | --- | --- | --- | --- | --- |
| pos | 10156 | 11596 | 14755 | 19170 | 21638 | 22026 | 22027 | 22029 | 22030 | 22031 | 22032 | 22033 | 22034 | 27328 |  |
| ref | C | G | C | C | C | G | T | A | G | T | T | C | A | T |  |
| AY.100 | T | T | T | T | T | G | T | - | - | - | - | - | - | C |  |
| AY.100 | T | T | T | T | T | G | G | A | A | A | G | T | G | C | #645455 |
| AY.100 | C | G | C | C | C | T | G | A | A | A | G | T | G | T |  |
| mut |  |  |  |  |  |  |  |  |  |  |  |  |  | T>C |  |

|  |  |  |  |  |  |  |  |  |  |  |  |  |  |  |  |  |  |  |
| --- | --- | --- | --- | --- | --- | --- | --- | --- | --- | --- | --- | --- | --- | --- | --- | --- | --- | --- |
| pos | 6408 | 7851 | 8179 | 8829 | 14999 | 18032 | 19006 | 19648 | 22027 | 22029 | 22030 | 22031 | 22032 | 22033 | 22034 | 25593 | 26217 |  |
| ref | C | C | G | C | G | C | G | G | T | A | G | T | T | C | A | G | G |  |
| AY.4 | T | T | T | C | A | T | G | T | T | - | - | - | - | - | - | T | G |  |
| AY.4 | T | T | T | C | A | T | G | T | G | A | A | A | G | T | G | T | T | #647013 |
| AY.119 | C | C | G | T | G | C | T | G | G | A | A | A | G | T | G | G | G |  |
| mut |  |  |  |  |  |  |  |  |  |  |  |  |  |  |  | G>T | G>T |  |

|  |  |  |  |  |  |  |  |  |  |  |  |  |  |  |  |  |  |  |  |  |  |  |
| --- | --- | --- | --- | --- | --- | --- | --- | --- | --- | --- | --- | --- | --- | --- | --- | --- | --- | --- | --- | --- | --- | --- |
| pos | 203 | 222 | 489 | 1635 | 1655 | 2227 | 6706 | 8664 | 8950 | 9163 | 9802 | 10870 | 13105 | 14418 | 15741 | 17040 | 18828 | 25775 | 26543 | 26720 | 27389 |  |
| ref | C | C | A | A | G | C | C | C | C | C | G | G | C | T | T | T | C | G | T | G | C |  |
| AY.4 | - | - | G | C | A | C | C | T | C | A | G | G | C | T | T | T | T | G | T | G | C |  |
| AY.4 | C | C | G | C | A | T | C | T | C | A | G | T | T | C | C | C | C | T | C | A | T | #649306 |
| AY.4 | T | C | A | A | G | C | T | C | T | C | T | T | T | C | C | C | C | T | C | A | T |  |
| mut | ->C | ->C |  |  |  | C>T |  |  |  |  |  |  |  |  |  |  |  |  |  |  |  |  |

|  |  |  |  |  |  |  |  |  |  |  |  |  |  |  |  |  |  |  |  |
| --- | --- | --- | --- | --- | --- | --- | --- | --- | --- | --- | --- | --- | --- | --- | --- | --- | --- | --- | --- |
| pos | 2536 | 2755 | 5031 | 11335 | 14585 | 15222 | 15852 | 22027 | 22029 | 22030 | 22031 | 22032 | 22033 | 22034 | 25339 | 25647 | 27503 | 29422 |  |
| ref | C | G | C | G | C | C | C | T | A | G | T | T | C | A | C | G | C | G |  |
| B.1.617.2 | T | G | C | A | T | T | T | T | - | - | - | - | - | - | T | G | C | T |  |
| AY.112 | T | T | T | A | T | T | T | G | A | A | A | G | T | G | C | T | T | G | #650313 |
| AY.112 | C | G | C | G | C | C | C | G | A | A | A | G | T | G | C | T | C | G |  |
| mut |  | G>T | C>T |  |  |  |  |  |  |  |  |  |  |  |  |  | C>T |  |  |

|  |  |  |  |  |  |  |  |  |  |  |  |  |  |  |  |
| --- | --- | --- | --- | --- | --- | --- | --- | --- | --- | --- | --- | --- | --- | --- | --- |
| pos | 337 | 920 | 10870 | 12513 | 16428 | 21987 | 22027 | 22029 | 22030 | 22031 | 22032 | 22033 | 22034 | 25275 |  |
| ref | C | C | G | C | C | G | T | A | G | T | T | C | A | C |  |
| AY.4.9 | T | T | G | T | T | A | T | - | - | - | - | - | - | T |  |
| AY.4.9 | T | T | G | T | T | A | G | A | A | A | G | T | G | T | #650320 |
| AY.4 | C | C | T | C | C | G | G | A | A | A | G | T | G | C |  |
| mut |  |  |  |  |  |  |  |  |  |  |  |  |  | C>T |  |

|  |  |  |  |  |  |  |  |  |  |  |  |  |  |  |  |  |  |  |  |  |  |  |  |  |
| --- | --- | --- | --- | --- | --- | --- | --- | --- | --- | --- | --- | --- | --- | --- | --- | --- | --- | --- | --- | --- | --- | --- | --- | --- |
| pos | 1186 | 3079 | 3182 | 4150 | 8829 | 9746 | 11803 | 13446 | 15237 | 17336 | 18885 | 19006 | 19476 | 20544 | 21137 | 22027 | 22029 | 22030 | 22031 | 22032 | 22033 | 22034 | 28727 |  |
| ref | G | A | G | T | C | C | T | C | C | C | C | G | A | C | A | T | A | G | T | T | C | A | G |  |
| AY.126 | G | A | G | C | C | T | A | T | T | T | T | G | G | T | G | T | - | - | - | - | - | - | T |  |
| AY.126 | G | A | G | C | C | T | A | T | T | T | T | G | G | T | G | G | A | A | A | G | T | G | T | #650448 |
| AY.119 | A | G | C | T | T | C | T | C | C | C | C | T | A | C | A | G | A | A | A | G | T | G | G |  |
| mut |  |  |  |  |  |  |  |  |  |  |  |  |  |  |  |  |  |  |  |  |  |  | G>T |  |

|  |  |  |  |  |  |  |  |  |  |  |  |  |  |  |  |  |  |  |  |  |  |
| --- | --- | --- | --- | --- | --- | --- | --- | --- | --- | --- | --- | --- | --- | --- | --- | --- | --- | --- | --- | --- | --- |
| pos | 7000 | 7334 | 7851 | 8829 | 15212 | 15435 | 19006 | 21853 | 22000 | 22027 | 22029 | 22030 | 22031 | 22032 | 22033 | 22034 | 23587 | 26464 | 28744 | 29543 |  |
| ref | C | C | C | C | C | A | G | G | C | T | A | G | T | T | C | A | G | C | C | G |  |
| AY.4 | T | T | T | C | T | A | G | T | T | T | - | - | - | - | - | - | G | T | C | C |  |
| AY.4 | T | T | T | C | T | A | G | T | T | G | A | A | A | G | T | G | G | T | T | C | #650553 |
| AY.119.2 | C | C | C | T | C | G | T | G | C | G | A | A | A | G | T | G | C | C | C | G |  |
| mut |  |  |  |  |  |  |  |  |  |  |  |  |  |  |  |  | C>G | C>T | C>T | G>C |  |

|  |  |  |  |  |  |  |  |  |  |  |  |  |  |  |  |  |
| --- | --- | --- | --- | --- | --- | --- | --- | --- | --- | --- | --- | --- | --- | --- | --- | --- |
| pos | 1225 | 2246 | 2941 | 3736 | 8296 | 17040 | 22026 | 22027 | 22029 | 22030 | 22031 | 22032 | 22033 | 22034 | 28483 |  |
| ref | C | G | A | C | T | T | G | T | A | G | T | T | C | A | A |  |
| AY.4 | T | A | G | T | C | C | G | T | - | - | - | - | - | - | T |  |
| AY.4 | T | A | G | T | C | C | T | G | A | A | A | G | T | G | T | #650570 |
| AY.4 | C | G | A | C | T | T | T | G | A | A | A | G | T | G | A |  |
| mut |  |  |  |  |  |  |  |  |  |  |  |  |  |  | A>T |  |

|  |  |  |  |  |  |  |  |  |  |  |  |  |  |  |  |  |  |  |  |  |  |  |  |  |
| --- | --- | --- | --- | --- | --- | --- | --- | --- | --- | --- | --- | --- | --- | --- | --- | --- | --- | --- | --- | --- | --- | --- | --- | --- |
| pos | 815 | 823 | 1588 | 1780 | 2632 | 3476 | 6726 | 12787 | 13115 | 13482 | 14513 | 15654 | 17555 | 21261 | 22027 | 22029 | 22030 | 22031 | 22032 | 22033 | 22034 | 24942 | 27632 |  |
| ref | C | C | A | C | G | G | C | A | C | A | A | C | C | A | T | A | G | T | T | C | A | A | G |  |
| AY.39.1 | T | T | A | T | G | G | C | G | T | G | C | C | T | G | T | - | - | - | - | - | - | G | A |  |
| AY.39.1 | T | T | A | T | G | G | C | G | T | G | C | C | T | G | G | A | A | A | G | T | G | G | A | #652407 |
| AY.39 | C | C | G | C | T | A | T | A | C | A | A | T | C | A | G | A | A | A | G | T | G | A | G |  |
| mut |  |  |  |  |  |  |  |  |  |  |  |  |  |  |  |  |  |  |  |  |  | A>G | G>A |  |

|  |  |  |  |  |  |  |  |  |  |  |  |  |  |  |  |  |  |  |
| --- | --- | --- | --- | --- | --- | --- | --- | --- | --- | --- | --- | --- | --- | --- | --- | --- | --- | --- |
| pos | 3264 | 5812 | 8612 | 8832 | 14293 | 14619 | 15012 | 17535 | 22026 | 22027 | 22029 | 22030 | 22031 | 22032 | 22033 | 22034 | 24415 |  |
| ref | C | C | G | C | C | T | A | T | G | T | A | G | T | T | C | A | G |  |
| AY.100 | T | T | T | T | C | C | G | T | G | T | - | - | - | - | - | - | A |  |
| AY.100 | T | T | T | T | T | C | G | C | G | G | A | A | A | G | T | G | A | #652451 |
| AY.100 | C | C | G | C | C | T | A | T | T | G | A | A | A | G | T | G | G |  |
| mut |  |  |  |  | C>T |  |  | T>C |  |  |  |  |  |  |  |  | G>A |  |

|  |  |  |  |  |  |  |  |  |  |  |  |  |  |  |  |  |  |  |  |  |
| --- | --- | --- | --- | --- | --- | --- | --- | --- | --- | --- | --- | --- | --- | --- | --- | --- | --- | --- | --- | --- |
| pos | 2509 | 8076 | 9064 | 18015 | 18148 | 21724 | 21987 | 22027 | 22029 | 22030 | 22031 | 22032 | 22033 | 22034 | 27572 | 27573 | 27575 | 27576 | 28903 |  |
| ref | C | T | T | A | G | G | G | T | A | G | T | T | C | A | G | C | C | T | G |  |
| AY.120.1 | T | C | C | G | T | T | G | T | - | - | - | - | - | - | A | T | A | C | C |  |
| AY.120.1 | T | C | C | G | T | T | G | G | A | A | A | G | T | G | G | C | C | T | C | #652506 |
| AY.120.1 | C | T | T | A | G | G | A | G | A | A | A | G | T | G | G | C | C | T | G |  |
| mut |  |  |  |  |  |  |  |  |  |  |  |  |  |  |  |  |  |  | G>C |  |

|  |  |  |  |  |  |  |  |  |  |  |  |  |  |  |  |
| --- | --- | --- | --- | --- | --- | --- | --- | --- | --- | --- | --- | --- | --- | --- | --- |
| pos | 219 | 763 | 1932 | 5711 | 14407 | 17751 | 18501 | 22027 | 22029 | 22030 | 22031 | 22032 | 22033 | 22034 |  |
| ref | G | C | A | C | C | T | C | T | A | G | T | T | C | A |  |
| AY.4 | G | T | C | A | T | A | T | T | - | - | - | - | - | - |  |
| AY.4 | G | T | C | A | T | A | T | G | A | A | A | G | T | G | #655037 |
| AY.4 | A | C | A | C | C | T | C | G | A | A | A | G | T | G |  |
| mut |  |  |  |  |  |  |  |  |  |  |  |  |  |  |  |

|  |  |  |  |  |  |  |  |  |  |  |  |  |  |  |  |  |  |  |
| --- | --- | --- | --- | --- | --- | --- | --- | --- | --- | --- | --- | --- | --- | --- | --- | --- | --- | --- |
| pos | 884 | 4720 | 5064 | 5175 | 7851 | 16914 | 17034 | 17944 | 19677 | 22027 | 22029 | 22030 | 22031 | 22032 | 22033 | 22034 | 22224 |  |
| ref | C | G | A | C | C | G | T | G | G | T | A | G | T | T | C | A | C |  |
| AY.4 | C | G | G | T | T | G | A | T | G | T | - | - | - | - | - | - | C |  |
| AY.4 | C | G | G | T | T | G | A | T | G | G | A | A | A | G | T | G | T | #655044 |
| AY.114 | T | T | A | C | C | T | T | G | T | G | A | A | A | G | T | G | C |  |
| mut |  |  |  |  |  |  |  |  |  |  |  |  |  |  |  |  | C>T |  |

|  |  |  |  |  |  |  |  |  |  |  |  |  |  |
| --- | --- | --- | --- | --- | --- | --- | --- | --- | --- | --- | --- | --- | --- |
| pos | 21987 | 22027 | 22029 | 22030 | 22031 | 22032 | 22033 | 22034 | 25912 | 28750 | 28818 | 29304 |  |
| ref | G | T | A | G | T | T | C | A | G | A | C | C |  |
| AY.4 | A | G | A | A | A | G | T | G | G | A | C | C |  |
| AY.4 | A | G | A | A | A | G | T | G | T | C | T | T | #655055 |
| AY.4 | G | T | - | - | - | - | - | - | T | C | T | T |  |
| mut |  |  |  |  |  |  |  |  |  |  |  |  |  |

|  |  |  |  |  |  |  |  |  |  |  |  |  |  |  |  |  |  |  |
| --- | --- | --- | --- | --- | --- | --- | --- | --- | --- | --- | --- | --- | --- | --- | --- | --- | --- | --- |
| pos | 660 | 4885 | 20098 | 22027 | 22029 | 22030 | 22031 | 22032 | 22033 | 22034 | 23368 | 23628 | 25350 | 27451 | 28690 | 28925 | 29254 |  |
| ref | G | T | G | T | A | G | T | T | C | A | A | G | C | T | G | G | G |  |
| AY.4 | G | T | G | G | A | A | A | G | T | G | A | G | C | T | G | G | G |  |
| AY.4 | A | C | T | G | A | A | A | G | T | G | T | T | T | C | T | T | T | #655065 |
| AY.4 | A | T | T | T | - | - | - | - | - | - | T | T | T | C | T | T | T |  |
| mut | G>A | T>C | G>T |  |  |  |  |  |  |  |  |  |  |  |  |  |  |  |

|  |  |  |  |  |  |  |  |  |  |  |  |  |  |  |  |  |
| --- | --- | --- | --- | --- | --- | --- | --- | --- | --- | --- | --- | --- | --- | --- | --- | --- |
| pos | 1659 | 7675 | 13730 | 16184 | 16425 | 21987 | 22027 | 22029 | 22030 | 22031 | 22032 | 22033 | 22034 | 25855 | 27143 |  |
| ref | G | G | C | A | T | G | T | A | G | T | T | C | A | G | C |  |
| AY.4.2 | T | T | T | A | A | A | T | - | - | - | - | - | - | T | T |  |
| AY.4.2 | T | T | T | G | A | A | G | A | A | A | G | T | G | T | T | #655237 |
| AY.4.2 | G | G | C | A | T | G | G | A | A | A | G | T | G | G | C |  |
| mut |  |  |  | A>G |  |  |  |  |  |  |  |  |  | G>T | C>T |  |

|  |  |  |  |  |  |  |  |  |  |  |  |  |  |  |  |
| --- | --- | --- | --- | --- | --- | --- | --- | --- | --- | --- | --- | --- | --- | --- | --- |
| pos | 280 | 4423 | 10236 | 16647 | 21987 | 22026 | 22027 | 22029 | 22030 | 22031 | 22032 | 22033 | 22034 | 23797 |  |
| ref | C | C | A | G | G | G | T | A | G | T | T | C | A | T |  |
| AY.39 | T | T | G | T | A | G | T | - | - | - | - | - | - | C |  |
| AY.39 | T | T | G | T | A | G | G | A | A | A | G | T | G | C | #656834 |
| AY.39 | C | C | A | G | G | T | G | A | A | A | G | T | G | T |  |
| mut |  |  |  |  |  |  |  |  |  |  |  |  |  | T>C |  |

|  |  |  |  |  |  |  |  |  |  |  |  |  |
| --- | --- | --- | --- | --- | --- | --- | --- | --- | --- | --- | --- | --- |
| pos | 4789 | 5544 | 14585 | 20726 | 22027 | 22029 | 22030 | 22031 | 22032 | 22033 | 22034 |  |
| ref | T | C | C | A | T | A | G | T | T | C | A |  |
| AY.4 | T | C | T | A | T | - | - | - | - | - | - |  |
| AY.4 | T | C | T | A | G | A | A | A | G | T | G | #658585 |
| AY.4 | C | T | C | G | G | A | A | A | G | T | G |  |
| mut |  |  |  |  |  |  |  |  |  |  |  |  |

|  |  |  |  |  |  |  |  |  |  |  |  |  |  |  |  |  |  |  |  |  |
| --- | --- | --- | --- | --- | --- | --- | --- | --- | --- | --- | --- | --- | --- | --- | --- | --- | --- | --- | --- | --- |
| pos | 372 | 874 | 884 | 2306 | 3250 | 4720 | 5869 | 7851 | 16914 | 19677 | 22027 | 22029 | 22030 | 22031 | 22032 | 22033 | 22034 | 25087 | 26826 |  |
| ref | A | C | C | C | G | G | C | C | G | G | T | A | G | T | T | C | A | A | C |  |
| AY.4 | C | T | C | T | T | G | T | T | G | G | T | - | - | - | - | - | - | C | T |  |
| AY.4 | C | T | C | T | T | G | T | T | G | G | G | A | A | A | G | T | G | C | T | #658623 |
| AY.114 | A | C | T | C | G | T | C | C | T | T | G | A | A | A | G | T | G | A | C |  |
| mut |  |  |  |  |  |  |  |  |  |  |  |  |  |  |  |  |  | A>C | C>T |  |

|  |  |  |  |  |  |  |  |  |  |  |  |  |  |  |  |  |  |  |  |  |  |
| --- | --- | --- | --- | --- | --- | --- | --- | --- | --- | --- | --- | --- | --- | --- | --- | --- | --- | --- | --- | --- | --- |
| pos | 884 | 2348 | 2410 | 3493 | 4720 | 7851 | 8452 | 14085 | 16914 | 19677 | 21974 | 22027 | 22029 | 22030 | 22031 | 22032 | 22033 | 22034 | 25260 | 28395 |  |
| ref | C | A | A | T | G | C | A | C | G | G | G | T | A | G | T | T | C | A | T | G |  |
| AY.4 | C | G | G | C | G | T | G | T | G | G | C | T | - | - | - | - | - | - | C | A |  |
| AY.4 | C | G | G | C | G | T | G | T | G | G | C | G | A | A | A | G | T | G | C | A | #658633 |
| AY.114 | T | A | A | T | T | C | A | C | T | T | G | G | A | A | A | G | T | G | T | G |  |
| mut |  |  |  |  |  |  |  |  |  |  |  |  |  |  |  |  |  |  | T>C | G>A |  |

|  |  |  |  |  |  |  |  |  |  |  |  |  |  |  |
| --- | --- | --- | --- | --- | --- | --- | --- | --- | --- | --- | --- | --- | --- | --- |
| pos | 3692 | 12786 | 22027 | 22029 | 22030 | 22031 | 22032 | 22033 | 22034 | 23959 | 27656 | 28310 | 29630 |  |
| ref | G | C | T | A | G | T | T | C | A | T | T | C | A |  |
| AY.4 | G | C | G | A | A | A | G | T | G | T | T | C | A |  |
| AY.4 | T | T | G | A | A | A | G | T | G | G | C | T | G | #658650 |
| AY.4 | T | T | T | - | - | - | - | - | - | G | C | T | G |  |
| mut | G>T | C>T |  |  |  |  |  |  |  |  |  |  |  |  |

|  |  |  |  |  |  |  |  |  |  |  |  |  |  |  |  |  |  |  |
| --- | --- | --- | --- | --- | --- | --- | --- | --- | --- | --- | --- | --- | --- | --- | --- | --- | --- | --- |
| pos | 5431 | 10228 | 10440 | 10560 | 10636 | 10998 | 21987 | 22027 | 22029 | 22030 | 22031 | 22032 | 22033 | 22034 | 25413 | 25687 | 27379 |  |
| ref | A | C | C | C | T | G | G | T | A | G | T | T | C | A | C | G | A |  |
| AY.4 | A | T | T | C | G | A | A | T | - | - | - | - | - | - | C | T | A |  |
| AY.4 | G | T | T | C | G | A | A | G | A | A | A | G | T | G | T | T | G | #658678 |
| AY.4 | A | C | C | T | T | G | G | G | A | A | A | G | T | G | C | G | A |  |
| mut | A>G |  |  |  |  |  |  |  |  |  |  |  |  |  | C>T | G>T | A>G |  |

|  |  |  |  |  |  |  |  |  |  |  |  |  |  |
| --- | --- | --- | --- | --- | --- | --- | --- | --- | --- | --- | --- | --- | --- |
| pos | 1218 | 7675 | 7716 | 13724 | 16184 | 22027 | 22029 | 22030 | 22031 | 22032 | 22033 | 22034 |  |
| ref | C | G | C | C | A | T | A | G | T | T | C | A |  |
| AY.4.2 | T | G | T | C | A | T | - | - | - | - | - | - |  |
| AY.4.2 | T | G | T | T | A | T | A | A | A | G | T | G | #658973 |
| AY.4.2 | C | T | C | C | G | G | A | A | A | G | T | G |  |
| mut |  |  |  | C>T |  |  |  |  |  |  |  |  |  |

|  |  |  |  |  |  |  |  |  |  |  |  |  |  |  |  |  |  |  |  |  |  |
| --- | --- | --- | --- | --- | --- | --- | --- | --- | --- | --- | --- | --- | --- | --- | --- | --- | --- | --- | --- | --- | --- |
| pos | 884 | 2091 | 2210 | 4158 | 4720 | 7851 | 8782 | 11923 | 15596 | 16914 | 17040 | 19677 | 21761 | 22027 | 22029 | 22030 | 22031 | 22032 | 22033 | 22034 |  |
| ref | C | C | G | C | G | C | C | A | A | G | T | G | G | T | A | G | T | T | C | A |  |
| AY.4 | C | T | T | T | G | T | T | G | G | G | C | G | T | T | - | - | - | - | - | - |  |
| AY.4 | C | T | T | T | G | T | T | G | G | G | C | G | T | G | A | A | A | G | T | G | #659106 |
| AY.114 | T | C | G | C | T | C | C | A | A | T | T | T | G | G | A | A | A | G | T | G |  |
| mut |  |  |  |  |  |  |  |  |  |  |  |  |  |  |  |  |  |  |  |  |  |

|  |  |  |  |  |  |  |  |  |  |  |  |  |  |  |  |  |
| --- | --- | --- | --- | --- | --- | --- | --- | --- | --- | --- | --- | --- | --- | --- | --- | --- |
| pos | 2102 | 4331 | 7851 | 21846 | 22006 | 22026 | 22027 | 22029 | 22030 | 22031 | 22032 | 22033 | 22034 | 24928 | 29253 |  |
| ref | C | C | C | C | C | G | T | A | G | T | T | C | A | G | C |  |
| AY.5 | T | T | C | C | T | G | T | - | - | - | - | - | - | T | T |  |
| AY.5 | T | T | C | C | T | T | G | A | A | A | G | T | G | T | T | #659159 |
| AY.4 | C | C | T | T | C | T | G | A | A | A | G | T | G | G | C |  |
| mut |  |  |  |  |  |  |  |  |  |  |  |  |  | G>T | C>T |  |

|  |  |  |  |  |  |  |  |  |  |  |  |  |  |  |  |  |  |  |  |  |
| --- | --- | --- | --- | --- | --- | --- | --- | --- | --- | --- | --- | --- | --- | --- | --- | --- | --- | --- | --- | --- |
| pos | 884 | 3372 | 3574 | 4720 | 6573 | 7851 | 9428 | 16914 | 17278 | 17573 | 19677 | 22027 | 22029 | 22030 | 22031 | 22032 | 22033 | 22034 | 25552 |  |
| ref | C | A | T | G | C | C | A | G | G | C | G | T | A | G | T | T | C | A | G |  |
| AY.4 | C | G | C | G | T | T | G | G | T | T | G | T | - | - | - | - | - | - | G |  |
| AY.4 | C | G | C | G | T | T | G | G | T | T | G | G | A | A | A | G | T | G | T | #659168 |
| AY.114 | T | A | T | T | C | C | A | T | G | C | T | G | A | A | A | G | T | G | G |  |
| mut |  |  |  |  |  |  |  |  |  |  |  |  |  |  |  |  |  |  | G>T |  |

|  |  |  |  |  |  |  |  |  |  |  |  |  |  |  |  |
| --- | --- | --- | --- | --- | --- | --- | --- | --- | --- | --- | --- | --- | --- | --- | --- |
| pos | 1185 | 21987 | 22027 | 22029 | 22030 | 22031 | 22032 | 22033 | 22034 | 24757 | 25129 | 25505 | 26208 | 28905 |  |
| ref | C | G | T | A | G | T | T | C | A | G | T | A | C | C |  |
| AY.4 | C | G | G | A | A | A | G | T | G | G | T | A | C | C |  |
| AY.4 | T | G | G | A | A | A | G | T | G | T | C | T | T | T | #659191 |
| AY.4 | T | A | T | - | - | - | - | - | - | T | T | T | T | T |  |
| mut | C>T |  |  |  |  |  |  |  |  |  | T>C |  |  |  |  |

|  |  |  |  |  |  |  |  |  |  |  |  |  |  |  |  |  |  |
| --- | --- | --- | --- | --- | --- | --- | --- | --- | --- | --- | --- | --- | --- | --- | --- | --- | --- |
| pos | 1613 | 2564 | 3948 | 5612 | 6040 | 6638 | 7926 | 11562 | 14014 | 16726 | 23996 | 26034 | 26107 | 27507 | 28073 | 29700 |  |
| ref | C | G | A | G | C | C | C | G | T | C | C | G | G | A | G | A |  |
| AY.44 | C | G | A | G | T | T | T | G | G | T | T | G | G | A | T | G |  |
| AY.44 | A | G | A | G | T | T | T | G | G | T | T | A | C | C | G | A | #660281 |
| AY.25.1 | C | A | G | A | C | C | C | T | T | C | C | A | C | C | G | A |  |
| mut | C>A |  |  |  |  |  |  |  |  |  |  |  |  |  |  |  |  |

|  |  |  |  |  |  |  |  |  |  |  |  |  |  |  |  |  |  |  |  |  |
| --- | --- | --- | --- | --- | --- | --- | --- | --- | --- | --- | --- | --- | --- | --- | --- | --- | --- | --- | --- | --- |
| pos | 7764 | 8956 | 9118 | 11620 | 12073 | 13482 | 19086 | 19894 | 21304 | 21538 | 22026 | 22027 | 22029 | 22030 | 22031 | 22032 | 22033 | 22034 | 26058 |  |
| ref | C | C | C | C | C | A | G | G | C | G | G | T | A | G | T | T | C | A | C |  |
| AY.39 | T | T | C | T | T | G | T | A | T | A | G | T | - | - | - | - | - | - | T |  |
| AY.39 | T | T | T | T | T | G | T | A | T | A | G | G | A | A | A | G | T | G | C | #660445 |
| AY.39 | C | C | C | C | C | A | G | G | C | G | T | G | A | A | A | G | T | G | C |  |
| mut |  |  | C>T |  |  |  |  |  |  |  |  |  |  |  |  |  |  |  |  |  |

|  |  |  |  |  |  |  |  |  |  |  |  |  |  |  |  |  |  |
| --- | --- | --- | --- | --- | --- | --- | --- | --- | --- | --- | --- | --- | --- | --- | --- | --- | --- |
| pos | 3044 | 4197 | 6528 | 8762 | 11596 | 19816 | 19962 | 22026 | 22027 | 22029 | 22030 | 22031 | 22032 | 22033 | 22034 | 28849 |  |
| ref | C | A | A | G | G | G | G | G | T | A | G | T | T | C | A | C |  |
| AY.100 | T | C | G | A | T | T | T | G | T | - | - | - | - | - | - | C |  |
| AY.100 | T | C | G | A | T | T | T | G | G | A | A | A | G | T | G | T | #660471 |
| AY.100 | C | A | A | G | G | G | G | T | G | A | A | A | G | T | G | C |  |
| mut |  |  |  |  |  |  |  |  |  |  |  |  |  |  |  | C>T |  |

|  |  |  |  |  |  |  |  |  |  |  |  |  |  |  |  |  |  |  |
| --- | --- | --- | --- | --- | --- | --- | --- | --- | --- | --- | --- | --- | --- | --- | --- | --- | --- | --- |
| pos | 2536 | 2878 | 4668 | 4795 | 5704 | 6543 | 6573 | 20937 | 22027 | 22029 | 22030 | 22031 | 22032 | 22033 | 22034 | 25062 | 29825 |  |
| ref | C | C | C | C | G | C | C | G | T | A | G | T | T | C | A | G | G |  |
| AY.127 | C | T | T | T | A | T | T | A | T | - | - | - | - | - | - | T | T |  |
| AY.127 | T | T | T | T | A | T | T | A | G | A | A | A | G | T | G | G | T | #660483 |
| AY.127 | C | C | C | C | G | C | C | G | G | A | A | A | G | T | G | G | G |  |
| mut | C>T |  |  |  |  |  |  |  |  |  |  |  |  |  |  |  | G>T |  |

|  |  |  |  |  |  |  |  |  |  |  |  |  |  |  |  |  |  |  |  |  |  |  |
| --- | --- | --- | --- | --- | --- | --- | --- | --- | --- | --- | --- | --- | --- | --- | --- | --- | --- | --- | --- | --- | --- | --- |
| pos | 2110 | 3874 | 4904 | 6408 | 7851 | 8473 | 10870 | 13944 | 15315 | 17023 | 17040 | 21995 | 22227 | 23557 | 25003 | 25189 | 25614 | 26488 | 26491 | 26492 | 26497 |  |
| ref | C | C | G | C | C | T | G | C | C | T | T | T | C | C | A | T | C | T | T | A | T |  |
| AY.4.2 | T | C | G | C | T | T | T | C | T | G | C | C | T | T | A | T | T | T | T | A | T |  |
| AY.4.2 | T | T | T | C | T | T | T | C | T | G | C | C | T | T | A | C | T | G | C | T | C | #662100 |
| AY.120 | C | C | G | T | C | C | G | T | C | T | T | T | C | C | G | T | C | G | C | T | C |  |
| mut |  | C>T | G>T |  |  |  |  |  |  |  |  |  |  |  |  | T>C |  |  |  |  |  |  |

|  |  |  |  |  |  |  |  |  |  |  |  |  |  |  |  |  |  |
| --- | --- | --- | --- | --- | --- | --- | --- | --- | --- | --- | --- | --- | --- | --- | --- | --- | --- |
| pos | 1594 | 3117 | 9389 | 9519 | 17940 | 20262 | 21987 | 22026 | 22027 | 22029 | 22030 | 22031 | 22032 | 22033 | 22034 | 29666 |  |
| ref | C | C | G | T | A | A | G | G | T | A | G | T | T | C | A | C |  |
| AY.4.2 | T | T | A | G | G | G | A | G | T | - | - | - | - | - | - | T |  |
| AY.4.2 | T | T | A | G | G | G | A | T | G | A | A | A | G | T | G | T | #662281 |
| AY.4.2 | C | C | G | T | A | A | G | T | G | A | A | A | G | T | G | C |  |
| mut |  |  |  |  |  |  |  |  |  |  |  |  |  |  |  | C>T |  |

|  |  |  |  |  |  |  |  |  |  |  |  |  |  |  |  |
| --- | --- | --- | --- | --- | --- | --- | --- | --- | --- | --- | --- | --- | --- | --- | --- |
| pos | 8140 | 12651 | 22027 | 22029 | 22030 | 22031 | 22032 | 22033 | 22034 | 22465 | 24985 | 25435 | 27800 | 29363 |  |
| ref | C | C | T | A | G | T | T | C | A | T | G | T | C | C |  |
| AY.4 | C | C | G | A | A | A | G | T | G | T | G | T | C | C |  |
| AY.4 | T | C | G | A | A | A | G | T | G | C | T | C | G | T | #662323 |
| AY.4 | T | T | T | - | - | - | - | - | - | C | T | C | G | C |  |
| mut | C>T |  |  |  |  |  |  |  |  |  |  |  |  | C>T |  |

|  |  |  |  |  |  |  |  |  |  |  |  |  |  |  |  |  |  |  |  |  |  |
| --- | --- | --- | --- | --- | --- | --- | --- | --- | --- | --- | --- | --- | --- | --- | --- | --- | --- | --- | --- | --- | --- |
| pos | 884 | 1820 | 2453 | 4720 | 5108 | 7851 | 10790 | 12770 | 16914 | 19677 | 19872 | 21762 | 22027 | 22029 | 22030 | 22031 | 22032 | 22033 | 22034 | 25770 |  |
| ref | C | G | C | G | G | C | C | T | G | G | G | C | T | A | G | T | T | C | A | G |  |
| AY.4 | C | A | T | G | T | T | T | T | G | G | T | T | T | - | - | - | - | - | - | T |  |
| AY.4 | C | A | T | G | T | T | T | C | G | G | T | T | G | A | A | A | G | T | G | T | #662333 |
| AY.114 | T | G | C | T | G | C | C | T | T | T | G | C | G | A | A | A | G | T | G | G |  |
| mut |  |  |  |  |  |  |  | T>C |  |  |  |  |  |  |  |  |  |  |  | G>T |  |

|  |  |  |  |  |  |  |  |  |  |  |  |  |  |  |  |
| --- | --- | --- | --- | --- | --- | --- | --- | --- | --- | --- | --- | --- | --- | --- | --- |
| pos | 2525 | 9389 | 17940 | 18312 | 20262 | 21987 | 22027 | 22029 | 22030 | 22031 | 22032 | 22033 | 22034 | 24406 |  |
| ref | G | G | A | C | A | G | T | A | G | T | T | C | A | T |  |
| AY.4.2 | A | A | G | T | G | A | T | - | - | - | - | - | - | C |  |
| AY.4.2 | A | A | G | T | G | A | G | A | A | A | G | T | G | C | #662377 |
| AY.4.2 | G | G | A | C | A | G | G | A | A | A | G | T | G | T |  |
| mut |  |  |  |  |  |  |  |  |  |  |  |  |  | T>C |  |

|  |  |  |  |  |  |  |  |  |  |  |  |  |  |  |  |  |  |  |  |  |  |  |  |
| --- | --- | --- | --- | --- | --- | --- | --- | --- | --- | --- | --- | --- | --- | --- | --- | --- | --- | --- | --- | --- | --- | --- | --- |
| pos | 925 | 2047 | 4654 | 6449 | 7966 | 16443 | 17040 | 18412 | 19292 | 20283 | 20404 | 21048 | 21485 | 22027 | 22029 | 22030 | 22031 | 22032 | 22033 | 22034 | 26951 | 27630 |  |
| ref | C | T | T | C | A | C | T | G | G | C | C | T | G | T | A | G | T | T | C | A | G | C |  |
| AY.4 | T | T | C | T | G | T | C | T | G | T | T | T | G | T | - | - | - | - | - | - | T | C |  |
| AY.4 | T | T | C | T | G | T | C | T | G | T | T | T | G | G | A | A | A | G | T | G | T | G | #662393 |
| AY.4 | C | C | T | C | A | C | T | G | T | C | C | C | T | G | A | A | A | G | T | G | G | C |  |
| mut |  |  |  |  |  |  |  |  |  |  |  |  |  |  |  |  |  |  |  |  | G>T | C>G |  |

|  |  |  |  |  |  |  |  |  |  |  |  |  |  |  |  |  |  |  |  |  |  |  |
| --- | --- | --- | --- | --- | --- | --- | --- | --- | --- | --- | --- | --- | --- | --- | --- | --- | --- | --- | --- | --- | --- | --- |
| pos | 1170 | 6816 | 9684 | 17040 | 20996 | 21995 | 22227 | 25614 | 25906 | 26176 | 28077 | 28237 | 28241 | 28243 | 28245 | 28247 | 28251 | 28252 | 28253 | 28254 | 28845 |  |
| ref | C | G | T | T | G | T | C | C | G | C | G | G | T | T | T | A | T | T | C | A | G |  |
| AY.4 | T | A | C | T | G | T | C | C | G | T | T | G | T | T | T | A | - | - | - | A | G |  |
| AY.4 | T | A | C | T | G | T | C | C | G | T | T | T | A | A | C | G | T | T | C | C | G | #662434 |
| AY.4.2 | C | G | T | C | T | C | T | T | T | C | G | T | A | A | C | G | - | T | T | C | A |  |
| mut |  |  |  |  |  |  |  |  |  |  |  |  |  |  |  |  | ->T |  | T>C |  | A>G |  |

|  |  |  |  |  |  |  |  |  |  |  |  |  |  |  |  |
| --- | --- | --- | --- | --- | --- | --- | --- | --- | --- | --- | --- | --- | --- | --- | --- |
| pos | 2295 | 4448 | 8566 | 21600 | 21987 | 22026 | 22027 | 22029 | 22030 | 22031 | 22032 | 22033 | 22034 | 27632 |  |
| ref | C | A | T | G | G | G | T | A | G | T | T | C | A | G |  |
| AY.4.2.2 | T | G | C | C | A | G | T | - | - | - | - | - | - | T |  |
| AY.4.2.2 | T | G | C | C | A | T | G | A | A | A | G | T | G | G | #662437 |
| AY.4.2.2 | C | A | T | G | G | T | G | A | A | A | G | T | G | G |  |
| mut |  |  |  |  |  |  |  |  |  |  |  |  |  |  |  |

|  |  |  |  |  |  |  |  |  |  |  |  |  |  |  |  |  |  |  |  |  |  |
| --- | --- | --- | --- | --- | --- | --- | --- | --- | --- | --- | --- | --- | --- | --- | --- | --- | --- | --- | --- | --- | --- |
| pos | 884 | 4012 | 4720 | 7851 | 10323 | 10376 | 15277 | 16914 | 17040 | 17427 | 19677 | 22027 | 22029 | 22030 | 22031 | 22032 | 22033 | 22034 | 25785 | 28780 |  |
| ref | C | C | G | C | A | C | C | G | T | G | G | T | A | G | T | T | C | A | G | A |  |
| AY.4 | C | T | G | T | G | T | T | G | C | T | G | T | - | - | - | - | - | - | T | G |  |
| AY.4 | C | T | G | T | G | T | T | G | C | T | G | G | A | A | A | G | T | G | T | G | #662467 |
| AY.114 | T | C | T | C | A | C | C | T | T | G | T | G | A | A | A | G | T | G | G | A |  |
| mut |  |  |  |  |  |  |  |  |  |  |  |  |  |  |  |  |  |  | G>T | A>G |  |

|  |  |  |  |  |  |  |  |  |  |  |  |  |  |  |  |  |  |  |  |
| --- | --- | --- | --- | --- | --- | --- | --- | --- | --- | --- | --- | --- | --- | --- | --- | --- | --- | --- | --- |
| pos | 312 | 4543 | 4774 | 5672 | 10263 | 10323 | 12756 | 16995 | 22027 | 22029 | 22030 | 22031 | 22032 | 22033 | 22034 | 26894 | 28193 | 29700 |  |
| ref | T | C | A | C | C | A | C | C | T | A | G | T | T | C | A | C | G | A |  |
| AY.119 | C | T | G | T | T | G | T | T | T | - | - | - | - | - | - | T | A | A |  |
| AY.119 | C | T | G | T | T | G | T | T | G | A | A | A | G | T | G | T | A | A | #663450 |
| AY.119 | T | C | A | C | C | A | C | C | G | A | A | A | G | T | G | C | G | G |  |
| mut |  |  |  |  |  |  |  |  |  |  |  |  |  |  |  | C>T | G>A | G>A |  |

|  |  |  |  |  |  |  |  |  |  |  |  |  |  |  |  |  |  |  |  |  |  |  |
| --- | --- | --- | --- | --- | --- | --- | --- | --- | --- | --- | --- | --- | --- | --- | --- | --- | --- | --- | --- | --- | --- | --- |
| pos | 186 | 507 | 508 | 509 | 1222 | 2832 | 4158 | 4640 | 6040 | 6638 | 7926 | 9051 | 9491 | 14014 | 15952 | 16726 | 25352 | 25708 | 28073 | 28299 | 29700 |  |
| ref | C | A | T | G | T | A | C | T | C | C | C | C | C | T | C | C | G | C | G | A | A |  |
| AY.43 | T | T | C | A | T | A | C | T | C | C | C | C | C | T | A | C | G | C | G | T | A |  |
| AY.44 | T | T | C | A | A | G | T | C | T | T | T | T | T | G | C | T | T | T | T | A | G | #663484 |
| AY.44 | C | A | T | G | A | G | T | C | T | T | T | T | C | G | C | T | T | T | T | A | G |  |
| mut |  |  |  |  |  |  |  |  |  |  |  |  | C>T |  |  |  |  |  |  |  |  |  |

|  |  |  |  |  |  |  |  |  |  |  |  |  |  |  |  |  |  |
| --- | --- | --- | --- | --- | --- | --- | --- | --- | --- | --- | --- | --- | --- | --- | --- | --- | --- |
| pos | 679 | 1627 | 9936 | 17748 | 18090 | 20703 | 22026 | 22027 | 22029 | 22030 | 22031 | 22032 | 22033 | 22034 | 25708 | 26290 |  |
| ref | C | C | C | T | G | C | G | T | A | G | T | T | C | A | C | A |  |
| AY.39 | T | T | T | A | A | T | G | T | - | - | - | - | - | - | T | G |  |
| AY.39 | T | T | T | A | A | T | G | G | A | A | A | G | T | G | T | G | #663511 |
| AY.39 | C | C | C | T | G | C | T | G | A | A | A | G | T | G | C | A |  |
| mut |  |  |  |  |  |  |  |  |  |  |  |  |  |  | C>T | A>G |  |

|  |  |  |  |  |  |  |  |  |  |  |  |  |  |  |  |  |  |  |  |  |  |  |  |
| --- | --- | --- | --- | --- | --- | --- | --- | --- | --- | --- | --- | --- | --- | --- | --- | --- | --- | --- | --- | --- | --- | --- | --- |
| pos | 5541 | 5622 | 6230 | 6458 | 6894 | 6896 | 9496 | 10969 | 12462 | 13959 | 20578 | 21724 | 22026 | 22027 | 22029 | 22030 | 22031 | 22032 | 22033 | 22034 | 23897 | 25702 |  |
| ref | A | C | C | A | G | C | A | C | C | C | G | G | G | T | A | G | T | T | C | A | C | C |  |
| AY.39 | T | T | T | G | T | T | G | T | T | T | T | C | G | T | - | - | - | - | - | - | A | A |  |
| AY.39 | T | T | T | G | T | T | G | T | T | T | T | C | T | G | A | A | A | G | T | G | A | A | #663542 |
| AY.39 | A | C | C | A | G | C | A | C | C | C | G | G | T | G | A | A | A | G | T | G | C | C |  |
| mut |  |  |  |  |  |  |  |  |  |  |  |  |  |  |  |  |  |  |  |  | C>A | C>A |  |

|  |  |  |  |  |  |  |  |  |  |  |  |  |  |  |  |  |
| --- | --- | --- | --- | --- | --- | --- | --- | --- | --- | --- | --- | --- | --- | --- | --- | --- |
| pos | 431 | 5227 | 10835 | 11273 | 12067 | 13255 | 20511 | 22027 | 22029 | 22030 | 22031 | 22032 | 22033 | 22034 | 27788 |  |
| ref | G | A | G | G | G | C | A | T | A | G | T | T | C | A | G |  |
| AY.107 | A | T | A | T | G | T | G | T | - | - | - | - | - | - | G |  |
| AY.107 | A | T | A | T | T | T | G | G | A | A | A | G | T | G | T | #663550 |
| AY.107 | G | A | G | G | G | C | A | G | A | A | A | G | T | G | G |  |
| mut |  |  |  |  | G>T |  |  |  |  |  |  |  |  |  | G>T |  |

|  |  |  |  |  |  |  |  |  |  |  |  |  |  |  |
| --- | --- | --- | --- | --- | --- | --- | --- | --- | --- | --- | --- | --- | --- | --- |
| pos | 526 | 5392 | 7851 | 8299 | 15258 | 15346 | 19524 | 21846 | 24595 | 27875 | 27881 | 27882 | 27883 |  |
| ref | G | C | C | C | T | C | C | C | A | T | C | G | C |  |
| AY.4 | G | T | T | T | C | T | C | T | G | T | C | G | C |  |
| AY.4 | G | T | T | T | C | T | C | T | G | C | T | C | T | #665336 |
| AY.98 | T | C | C | C | T | C | T | C | A | C | T | C | T |  |
| mut |  |  |  |  |  |  |  |  |  |  |  |  |  |  |

|  |  |  |  |  |  |  |  |  |  |  |  |  |  |  |
| --- | --- | --- | --- | --- | --- | --- | --- | --- | --- | --- | --- | --- | --- | --- |
| pos | 21770 | 21987 | 22027 | 22029 | 22030 | 22031 | 22032 | 22033 | 22034 | 24766 | 24904 | 25489 | 29624 |  |
| ref | G | G | T | A | G | T | T | C | A | C | C | G | T |  |
| AY.4.2 | G | G | G | A | A | A | G | T | G | C | C | G | T |  |
| AY.4.2 | A | G | G | A | A | A | G | T | G | T | T | T | C | #665843 |
| AY.4.2 | A | A | T | - | - | - | - | - | - | T | T | T | C |  |
| mut | G>A |  |  |  |  |  |  |  |  |  |  |  |  |  |

|  |  |  |  |  |  |  |  |  |  |  |  |  |  |  |
| --- | --- | --- | --- | --- | --- | --- | --- | --- | --- | --- | --- | --- | --- | --- |
| pos | 1387 | 1594 | 7165 | 13711 | 21987 | 22026 | 22027 | 22029 | 22030 | 22031 | 22032 | 22033 | 22034 |  |
| ref | C | C | C | A | G | G | T | A | G | T | T | C | A |  |
| AY.4.2.1 | T | T | C | G | A | G | T | - | - | - | - | - | - |  |
| AY.4.2.1 | T | T | C | G | A | T | G | A | A | A | G | T | G | #669271 |
| AY.4.2.1 | C | C | A | A | G | T | G | A | A | A | G | T | G |  |
| mut |  |  |  |  |  |  |  |  |  |  |  |  |  |  |

|  |  |  |  |  |  |  |  |  |  |  |  |  |  |  |
| --- | --- | --- | --- | --- | --- | --- | --- | --- | --- | --- | --- | --- | --- | --- |
| pos | 6525 | 8299 | 14697 | 17339 | 21987 | 22026 | 22027 | 22029 | 22030 | 22031 | 22032 | 22033 | 22034 |  |
| ref | C | C | C | C | G | G | T | A | G | T | T | C | A |  |
| AY.4.2.2 | T | T | T | T | A | G | T | - | - | - | - | - | - |  |
| AY.4.2.2 | T | T | T | T | A | G | G | A | A | A | G | T | G | #669329 |
| AY.4.2.2 | C | C | C | C | G | T | G | A | A | A | G | T | G |  |
| mut |  |  |  |  |  |  |  |  |  |  |  |  |  |  |

|  |  |  |  |  |  |  |  |  |  |  |  |  |  |  |  |  |  |  |  |  |  |  |  |  |
| --- | --- | --- | --- | --- | --- | --- | --- | --- | --- | --- | --- | --- | --- | --- | --- | --- | --- | --- | --- | --- | --- | --- | --- | --- |
| pos | 222 | 1048 | 1218 | 2329 | 4173 | 5392 | 6896 | 7211 | 9073 | 11878 | 12751 | 20078 | 20262 | 22026 | 22027 | 22029 | 22030 | 22031 | 22032 | 22033 | 22034 | 24095 | 28045 |  |
| ref | C | G | C | G | C | C | C | T | C | C | T | C | A | G | T | A | G | T | T | C | A | G | C |  |
| AY.100 | C | G | T | A | C | C | C | C | C | C | C | C | G | G | T | - | - | - | - | - | - | T | T |  |
| AY.100 | C | G | T | A | C | C | C | C | C | C | C | C | G | G | G | A | A | A | G | T | G | T | T | #669991 |
| AY.100 | T | T | C | G | T | T | T | T | T | T | T | T | A | T | G | A | A | A | G | T | G | G | C |  |
| mut |  |  |  |  |  |  |  |  |  |  |  |  |  |  |  |  |  |  |  |  |  | G>T | C>T |  |

|  |  |  |  |  |  |  |  |  |  |  |  |  |  |  |  |  |  |  |  |  |  |  |  |  |  |  |  |  |  |  |  |  |  |  |  |  |  |  |  |  |  |  |
| --- | --- | --- | --- | --- | --- | --- | --- | --- | --- | --- | --- | --- | --- | --- | --- | --- | --- | --- | --- | --- | --- | --- | --- | --- | --- | --- | --- | --- | --- | --- | --- | --- | --- | --- | --- | --- | --- | --- | --- | --- | --- | --- |
| pos | 449 | 2246 | 4181 | 5184 | 5584 | 5661 | 5738 | 6402 | 7124 | 7851 | 8031 | 8838 | 8986 | 9053 | 9805 | 9891 | 10029 | 11201 | 11306 | 11332 | 11418 | 11514 | 12025 | 13019 | 14097 | 14220 | 17040 | 19220 | 19788 | 21846 | 22227 | 22717 | 23185 | 24631 | 25333 | 25352 | 25466 | 27688 | 27874 | 28916 | 29705 |  |
| ref | C | G | G | C | A | A | A | C | C | C | A | T | C | G | G | C | C | A | G | A | T | C | C | C | C | C | T | C | G | C | C | T | C | T | T | G | C | C | C | G | G |  |
| AY.4 | C | A | T | C | A | A | T | T | T | T | A | T | T | T | G | C | T | G | G | G | T | C | C | C | C | T | C | T | T | T | C | G | C | T | T | G | T | C | T | T | T |  |
| AY.4 | C | A | T | C | A | A | T | T | T | T | A | T | T | T | G | C | T | G | G | G | T | C | C | C | C | T | C | C | G | C | T | T | T | C | C | T | C | T | C | G | G | #672079 |
| B.1.617.2 | T | G | G | T | G | T | A | C | C | C | G | C | C | G | T | T | C | A | A | A | C | T | T | T | T | C | T | C | G | C | T | T | T | C | C | T | C | T | C | G | G |  |
| mut |  |  |  |  |  |  |  |  |  |  |  |  |  |  |  |  |  |  |  |  |  |  |  |  |  |  |  |  |  |  |  |  |  |  |  |  |  |  |  |  |  |  |

|  |  |  |  |  |  |  |  |  |  |  |  |  |  |  |  |  |  |  |
| --- | --- | --- | --- | --- | --- | --- | --- | --- | --- | --- | --- | --- | --- | --- | --- | --- | --- | --- |
| pos | 355 | 1342 | 4050 | 4720 | 5018 | 5991 | 8679 | 8707 | 12793 | 17040 | 17391 | 21516 | 25767 | 25906 | 25964 | 26527 | 27741 |  |
| ref | C | C | A | G | G | A | A | T | G | T | T | C | G | G | A | C | C |  |
| AY.4 | T | T | C | G | G | C | A | C | G | T | T | T | G | G | A | C | C |  |
| AY.4 | C | T | C | G | G | C | C | C | G | T | T | C | T | T | T | T | A | #674172 |
| AY.4 | C | C | A | A | T | A | A | T | T | C | C | C | T | T | T | T | C |  |
| mut | T>C |  |  |  |  |  | A>C |  |  |  |  |  |  |  |  |  | C>A |  |

|  |  |  |  |  |  |  |  |  |  |  |  |  |  |  |  |
| --- | --- | --- | --- | --- | --- | --- | --- | --- | --- | --- | --- | --- | --- | --- | --- |
| pos | 219 | 897 | 8950 | 11417 | 14790 | 20481 | 22027 | 22029 | 22030 | 22031 | 22032 | 22033 | 22034 | 24794 |  |
| ref | G | C | C | G | C | A | T | A | G | T | T | C | A | G |  |
| AY.4 | G | C | T | T | T | A | T | - | - | - | - | - | - | T |  |
| AY.4 | G | T | T | T | T | G | G | A | A | A | G | T | G | T | #676114 |
| AY.4 | A | C | C | G | C | A | G | A | A | A | G | T | G | G |  |
| mut |  | C>T |  |  |  | A>G |  |  |  |  |  |  |  | G>T |  |

|  |  |  |  |  |  |  |  |  |  |  |  |  |  |  |  |  |
| --- | --- | --- | --- | --- | --- | --- | --- | --- | --- | --- | --- | --- | --- | --- | --- | --- |
| pos | 6906 | 9996 | 14371 | 14961 | 18744 | 19881 | 22026 | 22027 | 22029 | 22030 | 22031 | 22032 | 22033 | 22034 | 25672 |  |
| ref | C | C | G | T | C | C | G | T | A | G | T | T | C | A | C |  |
| AY.100 | T | T | T | C | T | T | G | T | - | - | - | - | - | - | T |  |
| AY.100 | T | T | T | C | T | T | G | G | A | A | A | G | T | G | T | #677364 |
| AY.100 | C | C | G | T | C | C | T | G | A | A | A | G | T | G | C |  |
| mut |  |  |  |  |  |  |  |  |  |  |  |  |  |  | C>T |  |

|  |  |  |  |  |  |  |  |  |  |  |  |  |  |  |  |  |  |  |  |  |  |
| --- | --- | --- | --- | --- | --- | --- | --- | --- | --- | --- | --- | --- | --- | --- | --- | --- | --- | --- | --- | --- | --- |
| pos | 213 | 514 | 884 | 4720 | 6408 | 6586 | 6587 | 6589 | 6694 | 7851 | 15925 | 16914 | 19677 | 22027 | 22029 | 22030 | 22031 | 22032 | 22033 | 22034 |  |
| ref | G | T | C | G | C | T | A | G | G | C | C | G | G | T | A | G | T | T | C | A |  |
| AY.4 | A | A | C | G | T | A | C | T | A | T | T | G | G | T | - | - | - | - | - | - |  |
| AY.4 | A | A | C | G | T | A | C | T | A | T | T | G | G | G | A | A | A | G | T | G | #678495 |
| AY.114 | G | T | T | T | C | T | A | G | G | C | C | T | T | G | A | A | A | G | T | G |  |
| mut |  |  |  |  |  |  |  |  |  |  |  |  |  |  |  |  |  |  |  |  |  |

|  |  |  |  |  |  |  |  |  |  |  |  |  |  |  |  |  |  |  |  |
| --- | --- | --- | --- | --- | --- | --- | --- | --- | --- | --- | --- | --- | --- | --- | --- | --- | --- | --- | --- |
| pos | 601 | 884 | 4720 | 5681 | 7851 | 8595 | 16852 | 16914 | 17040 | 18021 | 19677 | 22027 | 22029 | 22030 | 22031 | 22032 | 22033 | 22034 |  |
| ref | C | C | G | A | C | C | G | G | T | G | G | T | A | G | T | T | C | A |  |
| AY.4 | T | C | G | C | T | T | T | G | C | T | G | T | - | - | - | - | - | - |  |
| AY.4 | T | C | G | C | T | T | T | G | C | T | G | G | A | A | A | G | T | G | #678515 |
| AY.114 | C | T | T | A | C | C | G | T | T | G | T | G | A | A | A | G | T | G |  |
| mut |  |  |  |  |  |  |  |  |  |  |  |  |  |  |  |  |  |  |  |

|  |  |  |  |  |  |  |  |  |  |  |  |  |  |  |  |  |  |  |  |  |
| --- | --- | --- | --- | --- | --- | --- | --- | --- | --- | --- | --- | --- | --- | --- | --- | --- | --- | --- | --- | --- |
| pos | 224 | 884 | 2445 | 4720 | 4960 | 6990 | 7851 | 12942 | 16914 | 19677 | 22027 | 22029 | 22030 | 22031 | 22032 | 22033 | 22034 | 22321 | 28603 |  |
| ref | T | C | C | G | G | C | C | A | G | G | T | A | G | T | T | C | A | T | C |  |
| AY.4 | G | C | T | G | T | T | T | T | G | G | T | - | - | - | - | - | - | C | T |  |
| AY.4 | G | C | T | G | T | T | T | T | G | G | G | A | A | A | G | T | G | C | T | #678517 |
| AY.114 | T | T | C | T | G | C | C | A | T | T | G | A | A | A | G | T | G | T | C |  |
| mut |  |  |  |  |  |  |  |  |  |  |  |  |  |  |  |  |  | T>C | C>T |  |

|  |  |  |  |  |  |  |  |  |  |  |  |  |  |  |  |  |  |  |  |  |  |  |  |  |  |  |  |
| --- | --- | --- | --- | --- | --- | --- | --- | --- | --- | --- | --- | --- | --- | --- | --- | --- | --- | --- | --- | --- | --- | --- | --- | --- | --- | --- | --- |
| pos | 1376 | 1753 | 2627 | 3964 | 4795 | 5812 | 6848 | 16289 | 17477 | 20991 | 21987 | 22027 | 22029 | 22030 | 22031 | 22032 | 22033 | 22034 | 25252 | 28409 | 29728 | 29762 | 29769 | 29774 | 29779 | 29781 |  |
| ref | C | T | C | A | C | C | C | C | A | G | G | T | A | G | T | T | C | A | G | C | T | C | C | C | G | G |  |
| AY.120 | T | C | T | G | T | T | T | T | G | T | G | T | - | - | - | - | - | - | T | C | C | - | - | - | - | - |  |
| AY.120 | T | C | T | G | T | T | T | T | G | T | G | G | A | A | A | G | T | G | T | T | T | C | C | C | G | G | #680514 |
| AY.120 | C | T | C | A | C | C | C | C | A | G | A | G | A | A | A | G | T | G | G | C | T | C | C | C | G | G |  |
| mut |  |  |  |  |  |  |  |  |  |  |  |  |  |  |  |  |  |  | G>T | C>T |  |  |  |  |  |  |  |

|  |  |  |  |  |  |  |  |  |  |  |  |  |  |  |  |  |  |  |  |
| --- | --- | --- | --- | --- | --- | --- | --- | --- | --- | --- | --- | --- | --- | --- | --- | --- | --- | --- | --- |
| pos | 1281 | 4183 | 5383 | 7162 | 9165 | 16726 | 21638 | 21987 | 21995 | 22027 | 22029 | 22030 | 22031 | 22032 | 22033 | 22034 | 25614 | 28899 |  |
| ref | C | T | A | C | C | C | C | G | T | T | A | G | T | T | C | A | C | G |  |
| AY.4 | T | C | T | T | T | T | T | A | T | T | - | - | - | - | - | - | C | A |  |
| AY.4 | T | C | T | T | T | T | T | A | T | G | A | A | A | G | T | G | C | A | #680516 |
| AY.4.2 | C | T | A | C | C | C | C | G | C | G | A | A | A | G | T | G | T | G |  |
| mut |  |  |  |  |  |  |  |  |  |  |  |  |  |  |  |  | T>C | G>A |  |

|  |  |  |  |  |  |  |  |  |  |  |  |  |  |  |  |  |  |  |  |  |  |  |  |  |  |  |  |  |  |  |  |  |  |  |  |  |  |  |  |  |  |  |  |  |  |  |  |
| --- | --- | --- | --- | --- | --- | --- | --- | --- | --- | --- | --- | --- | --- | --- | --- | --- | --- | --- | --- | --- | --- | --- | --- | --- | --- | --- | --- | --- | --- | --- | --- | --- | --- | --- | --- | --- | --- | --- | --- | --- | --- | --- | --- | --- | --- | --- | --- |
| pos | 392 | 3140 | 3634 | 4181 | 5184 | 5584 | 6040 | 6402 | 6638 | 7124 | 7926 | 8617 | 8986 | 9053 | 9725 | 9857 | 9891 | 10029 | 10520 | 11201 | 11332 | 11418 | 11514 | 13019 | 14014 | 16726 | 16997 | 17410 | 17505 | 19220 | 19684 | 20930 | 22227 | 23601 | 25624 | 26681 | 27870 | 27874 | 28073 | 28460 | 28553 | 28916 | 28968 | 29593 | 29700 | 29762 |  |
| ref | C | C | C | G | C | A | C | C | C | C | C | T | C | G | T | C | C | C | T | A | A | T | C | C | T | C | C | C | T | C | G | C | C | C | C | C | G | C | G | G | A | G | G | G | A | C |  |
| AY.44 | C | T | C | T | C | A | T | T | T | T | T | T | T | T | C | C | C | T | T | G | G | T | C | C | G | T | C | C | T | T | G | C | C | C | C | T | G | T | T | A | A | T | G | T | G | C |  |
| AY.44 | C | T | C | T | C | A | T | T | T | T | T | T | T | T | C | C | C | T | T | G | G | T | C | C | G | T | C | T | C | C | T | T | T | T | T | C | T | C | G | G | G | G | G | G | A | T | #681283 |
| AY.56 | T | C | T | G | T | G | C | C | C | C | C | C | C | G | T | T | T | C | C | A | A | C | T | T | T | C | T | T | C | C | T | T | T | T | T | C | T | C | G | G | G | G | A | G | A | T |  |
| mut |  |  |  |  |  |  |  |  |  |  |  |  |  |  |  |  |  |  |  |  |  |  |  |  |  |  |  |  |  |  |  |  |  |  |  |  |  |  |  |  |  |  | A>G |  |  |  |  |

|  |  |  |  |  |  |  |  |  |  |  |  |  |  |  |  |  |  |  |  |
| --- | --- | --- | --- | --- | --- | --- | --- | --- | --- | --- | --- | --- | --- | --- | --- | --- | --- | --- | --- |
| pos | 3304 | 3433 | 6478 | 14273 | 15297 | 16644 | 19743 | 20402 | 21821 | 22026 | 22027 | 22029 | 22030 | 22031 | 22032 | 22033 | 22034 | 28084 |  |
| ref | G | G | T | A | T | A | A | C | A | G | T | A | G | T | T | C | A | A |  |
| AY.39 | T | T | T | A | C | G | G | C | G | G | T | - | - | - | - | - | - | T |  |
| AY.39 | T | T | T | A | C | G | G | C | G | G | G | A | A | A | G | T | G | T | #681544 |
| AY.39 | G | G | C | C | T | A | A | T | A | T | G | A | A | A | G | T | G | A |  |
| mut |  |  |  |  |  |  |  |  |  |  |  |  |  |  |  |  |  | A>T |  |

|  |  |  |  |  |  |  |  |  |  |  |  |  |  |  |
| --- | --- | --- | --- | --- | --- | --- | --- | --- | --- | --- | --- | --- | --- | --- |
| pos | 3411 | 9967 | 20134 | 21141 | 21255 | 22027 | 22029 | 22030 | 22031 | 22032 | 22033 | 22034 | 29029 |  |
| ref | C | C | G | A | G | T | A | G | T | T | C | A | T |  |
| AY.4 | T | T | T | G | C | T | - | - | - | - | - | - | C |  |
| AY.4 | T | T | T | G | C | G | A | A | A | G | T | G | C | #684463 |
| AY.4 | C | C | G | A | G | G | A | A | A | G | T | G | T |  |
| mut |  |  |  |  |  |  |  |  |  |  |  |  | T>C |  |

|  |  |  |  |  |  |  |  |  |  |  |  |  |  |  |
| --- | --- | --- | --- | --- | --- | --- | --- | --- | --- | --- | --- | --- | --- | --- |
| pos | 219 | 7175 | 9491 | 20932 | 22027 | 22029 | 22030 | 22031 | 22032 | 22033 | 22034 | 29171 | 29272 |  |
| ref | G | T | C | G | T | A | G | T | T | C | A | C | C |  |
| AY.4 | G | C | T | T | T | - | - | - | - | - | - | C | T |  |
| AY.4 | G | C | T | T | G | A | A | A | G | T | G | T | T | #684482 |
| AY.4 | A | T | C | G | G | A | A | A | G | T | G | C | C |  |
| mut |  |  |  |  |  |  |  |  |  |  |  | C>T | C>T |  |

|  |  |  |  |  |  |  |  |  |  |  |  |  |  |  |  |  |  |  |  |
| --- | --- | --- | --- | --- | --- | --- | --- | --- | --- | --- | --- | --- | --- | --- | --- | --- | --- | --- | --- |
| pos | 884 | 3476 | 4720 | 7459 | 7851 | 8131 | 15277 | 16914 | 17040 | 17427 | 19677 | 22027 | 22029 | 22030 | 22031 | 22032 | 22033 | 22034 |  |
| ref | C | G | G | G | C | G | C | G | T | G | G | T | A | G | T | T | C | A |  |
| AY.4 | C | A | G | T | T | T | T | G | C | T | G | T | - | - | - | - | - | - |  |
| AY.4 | C | A | G | T | T | T | T | G | C | T | G | G | A | A | A | G | T | G | #684492 |
| AY.114 | T | G | T | G | C | G | C | T | T | G | T | G | A | A | A | G | T | G |  |
| mut |  |  |  |  |  |  |  |  |  |  |  |  |  |  |  |  |  |  |  |

|  |  |  |  |  |  |  |  |  |  |  |  |  |  |  |  |  |  |  |  |  |  |  |
| --- | --- | --- | --- | --- | --- | --- | --- | --- | --- | --- | --- | --- | --- | --- | --- | --- | --- | --- | --- | --- | --- | --- |
| pos | 3737 | 3987 | 4891 | 6568 | 6997 | 11014 | 16935 | 17010 | 21302 | 21304 | 21305 | 22027 | 22029 | 22030 | 22031 | 22032 | 22033 | 22034 | 25471 | 28666 | 29511 |  |
| ref | C | C | C | C | C | G | G | C | C | C | G | T | A | G | T | T | C | A | G | A | G |  |
| AY.4 | T | C | T | T | C | A | T | T | C | C | G | T | - | - | - | - | - | - | G | A | T |  |
| AY.4 | T | C | T | T | C | A | T | T | C | C | G | G | A | A | A | G | T | G | T | G | T | #684536 |
| AY.4 | C | T | C | C | T | G | G | C | T | A | A | G | A | A | A | G | T | G | G | A | G |  |
| mut |  |  |  |  |  |  |  |  |  |  |  |  |  |  |  |  |  |  | G>T | A>G | G>T |  |

|  |  |  |  |  |  |  |  |  |  |  |  |  |  |  |  |  |  |  |  |  |  |  |  |  |
| --- | --- | --- | --- | --- | --- | --- | --- | --- | --- | --- | --- | --- | --- | --- | --- | --- | --- | --- | --- | --- | --- | --- | --- | --- |
| pos | 1102 | 3542 | 5548 | 6616 | 7851 | 8658 | 8715 | 9802 | 13482 | 13638 | 21261 | 21372 | 21742 | 22026 | 22027 | 22029 | 22030 | 22031 | 22032 | 22033 | 22034 | 22567 | 27604 |  |
| ref | C | A | C | A | C | A | C | G | A | T | A | G | C | G | T | A | G | T | T | C | A | G | G |  |
| AY.4 | C | G | T | A | T | G | T | T | A | C | A | G | T | G | T | - | - | - | - | - | - | G | G |  |
| AY.4 | C | G | T | A | T | G | T | T | A | C | A | G | T | G | G | A | A | A | G | T | G | T | G | #684568 |
| AY.39.1 | T | A | C | G | C | A | C | G | G | T | G | T | C | T | G | A | A | A | G | T | G | G | A |  |
| mut |  |  |  |  |  |  |  |  |  |  |  |  |  |  |  |  |  |  |  |  |  | G>T | A>G |  |

|  |  |  |  |  |  |  |  |  |  |  |  |  |  |  |  |
| --- | --- | --- | --- | --- | --- | --- | --- | --- | --- | --- | --- | --- | --- | --- | --- |
| pos | 6250 | 6355 | 6944 | 7622 | 15854 | 22026 | 22027 | 22029 | 22030 | 22031 | 22032 | 22033 | 22034 | 28378 |  |
| ref | C | A | A | G | T | G | T | A | G | T | T | C | A | G |  |
| AY.119 | C | G | G | A | T | G | T | - | - | - | - | - | - | G |  |
| AY.119 | C | G | G | A | T | G | T | A | A | A | G | T | G | G | #685445 |
| AY.119 | T | A | A | G | A | T | G | A | A | A | G | T | G | A |  |
| mut |  |  |  |  |  |  |  |  |  |  |  |  |  | A>G |  |

|  |  |  |  |  |  |  |  |  |  |  |  |  |  |  |  |
| --- | --- | --- | --- | --- | --- | --- | --- | --- | --- | --- | --- | --- | --- | --- | --- |
| pos | 4551 | 9165 | 9684 | 10507 | 22026 | 22027 | 22029 | 22030 | 22031 | 22032 | 22033 | 22034 | 27904 | 29703 |  |
| ref | C | C | T | C | G | T | A | G | T | T | C | A | T | G |  |
| AY.39 | T | T | T | T | G | T | - | - | - | - | - | - | C | G |  |
| AY.39 | T | T | T | T | G | G | A | A | A | G | T | G | C | A | #685448 |
| AY.39 | C | C | C | C | T | G | A | A | A | G | T | G | T | G |  |
| mut |  |  |  |  |  |  |  |  |  |  |  |  | T>C | G>A |  |

|  |  |  |  |  |  |  |  |  |  |  |  |  |  |  |  |  |  |  |  |  |  |  |  |
| --- | --- | --- | --- | --- | --- | --- | --- | --- | --- | --- | --- | --- | --- | --- | --- | --- | --- | --- | --- | --- | --- | --- | --- |
| pos | 4543 | 4680 | 6463 | 7851 | 8146 | 8299 | 8473 | 9165 | 10323 | 12772 | 15240 | 22026 | 22027 | 22029 | 22030 | 22031 | 22032 | 22033 | 22034 | 25398 | 28603 | 29737 |  |
| ref | C | C | G | C | C | C | T | C | A | A | C | G | T | A | G | T | T | C | A | T | C | G |  |
| AY.113 | C | T | T | C | T | T | C | T | G | G | T | G | T | - | - | - | - | - | - | C | C | T |  |
| AY.113 | T | T | T | C | T | T | C | T | G | G | T | T | G | A | A | A | G | T | G | C | T | T | #685497 |
| AY.4 | C | C | G | T | C | C | T | C | A | A | C | T | G | A | A | A | G | T | G | T | C | G |  |
| mut | C>T |  |  |  |  |  |  |  |  |  |  |  |  |  |  |  |  |  |  | T>C | C>T | G>T |  |

|  |  |  |  |  |  |  |  |  |  |  |  |  |  |  |  |  |  |  |  |  |  |  |  |
| --- | --- | --- | --- | --- | --- | --- | --- | --- | --- | --- | --- | --- | --- | --- | --- | --- | --- | --- | --- | --- | --- | --- | --- |
| pos | 386 | 829 | 4021 | 7851 | 9165 | 9640 | 10323 | 12559 | 12772 | 15240 | 15629 | 18082 | 21008 | 21976 | 22027 | 22029 | 22030 | 22031 | 22032 | 22033 | 22034 | 28087 |  |
| ref | G | C | C | C | C | T | A | C | A | C | G | A | C | T | T | A | G | T | T | C | A | C |  |
| AY.113 | G | T | A | C | T | C | G | T | G | T | G | A | T | C | T | - | - | - | - | - | - | T |  |
| AY.113 | A | T | A | C | T | C | G | T | G | T | G | A | T | C | G | A | A | A | G | T | G | T | #685524 |
| AY.4 | G | C | C | T | C | T | A | C | A | C | T | C | C | T | G | A | A | A | G | T | G | C |  |
| mut | G>A |  |  |  |  |  |  |  |  |  |  |  |  |  |  |  |  |  |  |  |  | C>T |  |

|  |  |  |  |  |  |  |  |  |  |  |  |  |  |  |
| --- | --- | --- | --- | --- | --- | --- | --- | --- | --- | --- | --- | --- | --- | --- |
| pos | 622 | 4993 | 12241 | 17747 | 19983 | 22027 | 22029 | 22030 | 22031 | 22032 | 22033 | 22034 | 29167 |  |
| ref | C | C | C | C | C | T | A | G | T | T | C | A | C |  |
| AY.121 | T | T | T | T | T | T | - | - | - | - | - | - | T |  |
| AY.121 | T | T | T | T | T | G | A | A | A | G | T | G | T | #688398 |
| AY.121 | C | C | C | C | C | G | A | A | A | G | T | G | C |  |
| mut |  |  |  |  |  |  |  |  |  |  |  |  | C>T |  |

|  |  |  |  |  |  |  |  |  |  |  |  |  |  |  |  |  |  |  |  |
| --- | --- | --- | --- | --- | --- | --- | --- | --- | --- | --- | --- | --- | --- | --- | --- | --- | --- | --- | --- |
| pos | 556 | 1192 | 2259 | 5497 | 6582 | 15720 | 17562 | 18687 | 21668 | 21987 | 22027 | 22029 | 22030 | 22031 | 22032 | 22033 | 22034 | 28878 |  |
| ref | C | A | T | C | C | C | G | C | G | G | T | A | G | T | T | C | A | G |  |
| AY.4.2.1 | T | G | G | T | T | T | T | T | T | A | T | - | - | - | - | - | - | T |  |
| AY.4.2.1 | T | G | G | T | T | T | T | T | T | G | G | A | A | A | G | T | G | T | #688752 |
| AY.4.2 | C | A | T | C | C | C | G | C | G | G | G | A | A | A | G | T | G | G |  |
| mut |  |  |  |  |  |  |  |  |  | A>G |  |  |  |  |  |  |  | G>T |  |

|  |  |  |  |  |  |  |  |  |  |  |  |  |  |  |  |  |  |  |  |  |  |
| --- | --- | --- | --- | --- | --- | --- | --- | --- | --- | --- | --- | --- | --- | --- | --- | --- | --- | --- | --- | --- | --- |
| pos | 1588 | 2632 | 3433 | 3476 | 6726 | 9693 | 9805 | 10138 | 10423 | 15654 | 15714 | 17280 | 21987 | 22027 | 22029 | 22030 | 22031 | 22032 | 22033 | 22034 |  |
| ref | A | G | G | G | C | C | G | C | T | C | C | G | G | T | A | G | T | T | C | A |  |
| AY.39 | A | G | T | G | C | T | T | T | C | C | T | A | G | T | - | - | - | - | - | - |  |
| AY.39 | A | G | T | G | C | T | T | T | C | C | T | A | G | G | A | A | A | G | T | G | #689517 |
| AY.39 | G | T | G | A | T | C | G | C | T | T | C | G | A | G | A | A | A | G | T | G |  |
| mut |  |  |  |  |  |  |  |  |  |  |  |  |  |  |  |  |  |  |  |  |  |

|  |  |  |  |  |  |  |  |  |  |  |  |  |  |  |  |  |
| --- | --- | --- | --- | --- | --- | --- | --- | --- | --- | --- | --- | --- | --- | --- | --- | --- |
| pos | 2324 | 3140 | 4917 | 6883 | 11001 | 13484 | 19586 | 22027 | 22029 | 22030 | 22031 | 22032 | 22033 | 22034 | 25768 |  |
| ref | G | C | T | C | C | G | C | T | A | G | T | T | C | A | A |  |
| AY.119 | G | T | T | T | C | A | C | T | - | - | - | - | - | - | A |  |
| AY.119 | G | T | T | T | C | A | C | G | A | A | A | G | T | G | G | #689530 |
| AY.119 | A | C | C | C | T | G | T | G | A | A | A | G | T | G | A |  |
| mut |  |  |  |  |  |  |  |  |  |  |  |  |  |  | A>G |  |

|  |  |  |  |  |  |  |  |  |  |  |  |  |  |  |  |  |  |  |  |
| --- | --- | --- | --- | --- | --- | --- | --- | --- | --- | --- | --- | --- | --- | --- | --- | --- | --- | --- | --- |
| pos | 1722 | 1741 | 6408 | 7851 | 8473 | 13944 | 17004 | 21987 | 24985 | 25003 | 25435 | 26488 | 26491 | 26492 | 26497 | 27737 | 27739 | 27741 |  |
| ref | C | A | C | C | T | C | C | G | G | A | T | T | T | A | T | C | C | C |  |
| AY.4 | C | G | C | T | T | C | C | G | T | A | C | T | T | A | T | A | A | T |  |
| AY.4 | T | G | C | T | T | C | T | A | T | A | C | G | C | T | C | C | C | C | #691949 |
| AY.120 | C | A | T | C | C | T | C | A | G | G | T | G | C | T | C | C | C | C |  |
| mut | C>T |  |  |  |  |  | C>T | G>A |  |  |  |  |  |  |  |  |  |  |  |

|  |  |  |  |  |  |  |  |  |  |  |  |  |  |  |  |  |  |
| --- | --- | --- | --- | --- | --- | --- | --- | --- | --- | --- | --- | --- | --- | --- | --- | --- | --- |
| pos | 3903 | 13120 | 15192 | 15720 | 18687 | 21304 | 21305 | 21987 | 22027 | 22029 | 22030 | 22031 | 22032 | 22033 | 22034 | 23593 |  |
| ref | C | T | A | C | C | C | G | G | T | A | G | T | T | C | A | G |  |
| AY.4.2 | T | C | G | T | T | A | A | A | T | - | - | - | - | - | - | T |  |
| AY.4.2 | T | C | G | T | T | A | A | G | G | A | A | A | G | T | G | T | #692280 |
| AY.4.2 | C | T | A | C | C | C | G | G | G | A | A | A | G | T | G | G |  |
| mut |  |  |  |  |  |  |  | A>G |  |  |  |  |  |  |  | G>T |  |

|  |  |  |  |  |  |  |  |  |  |  |  |  |  |  |  |  |
| --- | --- | --- | --- | --- | --- | --- | --- | --- | --- | --- | --- | --- | --- | --- | --- | --- |
| pos | 439 | 6567 | 15277 | 15720 | 17427 | 18687 | 21668 | 21995 | 22227 | 23683 | 25614 | 28090 | 28819 | 28878 | 29140 |  |
| ref | A | A | C | C | G | C | G | T | C | C | C | G | C | G | G |  |
| AY.4 | G | G | T | C | T | C | G | T | C | T | C | G | C | G | A |  |
| AY.4.2 | G | G | T | C | T | C | G | C | T | C | T | A | T | T | G | #695155 |
| AY.4.2.1 | A | A | C | T | G | T | T | C | T | C | T | A | C | T | G |  |
| mut |  |  |  |  |  |  |  |  |  |  |  |  | C>T |  |  |  |

|  |  |  |  |  |  |  |  |  |  |  |  |  |  |  |  |  |  |  |  |
| --- | --- | --- | --- | --- | --- | --- | --- | --- | --- | --- | --- | --- | --- | --- | --- | --- | --- | --- | --- |
| pos | 7165 | 9195 | 15720 | 18687 | 19999 | 20976 | 21550 | 21551 | 21668 | 21987 | 22027 | 22029 | 22030 | 22031 | 22032 | 22033 | 22034 | 28878 |  |
| ref | C | C | C | C | G | T | A | A | G | G | T | A | G | T | T | C | A | G |  |
| AY.4.2.1 | A | C | T | T | T | C | C | T | T | A | T | - | - | - | - | - | - | T |  |
| AY.4.2.1 | A | T | T | T | T | C | C | T | T | A | G | A | A | A | G | T | G | T | #695349 |
| AY.4.2 | C | C | C | C | G | T | A | A | G | G | G | A | A | A | G | T | G | G |  |
| mut |  | C>T |  |  |  |  |  |  |  |  |  |  |  |  |  |  |  | G>T |  |

|  |  |  |  |  |  |  |  |  |  |  |  |  |  |  |  |
| --- | --- | --- | --- | --- | --- | --- | --- | --- | --- | --- | --- | --- | --- | --- | --- |
| pos | 526 | 3811 | 6362 | 6629 | 7851 | 10279 | 19524 | 19960 | 20886 | 21846 | 25500 | 26078 | 29179 | 29736 |  |
| ref | G | C | G | C | C | C | C | A | A | C | G | C | G | G |  |
| AY.4 | G | C | A | T | T | T | C | G | G | T | T | T | T | G |  |
| AY.98 | T | T | A | T | T | T | C | G | G | T | G | C | G | C | #696327 |
| AY.98 | T | T | G | C | C | C | T | A | A | C | G | C | G | C |  |
| mut | G>T | C>T |  |  |  |  |  |  |  |  |  |  |  |  |  |

|  |  |  |  |  |  |  |  |  |  |  |  |  |  |  |  |  |
| --- | --- | --- | --- | --- | --- | --- | --- | --- | --- | --- | --- | --- | --- | --- | --- | --- |
| pos | 3085 | 4782 | 5668 | 9360 | 17913 | 18225 | 22027 | 22029 | 22030 | 22031 | 22032 | 22033 | 22034 | 22264 | 29743 |  |
| ref | G | C | G | C | C | G | T | A | G | T | T | C | A | C | C |  |
| AY.117 | T | T | G | T | C | T | T | - | - | - | - | - | - | T | A |  |
| AY.117 | T | T | G | T | C | T | G | A | A | A | G | T | G | T | A | #696783 |
| AY.117 | G | C | A | C | T | G | G | A | A | A | G | T | G | C | C |  |
| mut |  |  |  |  |  |  |  |  |  |  |  |  |  | C>T | C>A |  |

|  |  |  |  |  |  |  |  |  |  |  |  |  |  |  |  |  |  |
| --- | --- | --- | --- | --- | --- | --- | --- | --- | --- | --- | --- | --- | --- | --- | --- | --- | --- |
| pos | 1190 | 2088 | 9166 | 10623 | 12961 | 16575 | 17034 | 22027 | 22029 | 22030 | 22031 | 22032 | 22033 | 22034 | 22217 | 28098 |  |
| ref | C | T | C | C | A | C | T | T | A | G | T | T | C | A | G | T |  |
| AY.119 | C | G | A | T | G | T | A | T | - | - | - | - | - | - | A | T |  |
| AY.119 | T | G | A | T | G | T | A | G | A | A | A | G | T | G | A | C | #696798 |
| AY.119 | C | T | C | C | A | C | T | G | A | A | A | G | T | G | G | T |  |
| mut | C>T |  |  |  |  |  |  |  |  |  |  |  |  |  | G>A | T>C |  |

|  |  |  |  |  |  |  |  |  |  |  |  |  |  |  |  |  |  |  |
| --- | --- | --- | --- | --- | --- | --- | --- | --- | --- | --- | --- | --- | --- | --- | --- | --- | --- | --- |
| pos | 1820 | 3775 | 5457 | 7787 | 11284 | 13285 | 14724 | 14902 | 21608 | 22027 | 22029 | 22030 | 22031 | 22032 | 22033 | 22034 | 25046 |  |
| ref | G | C | C | G | T | A | C | A | G | T | A | G | T | T | C | A | C |  |
| AY.4 | A | T | C | G | G | G | T | G | G | T | - | - | - | - | - | - | T |  |
| AY.4 | A | T | C | G | G | G | T | G | T | G | A | A | A | G | T | G | T | #698131 |
| AY.4 | G | C | T | A | T | A | C | A | G | G | A | A | A | G | T | G | C |  |
| mut |  |  |  |  |  |  |  |  | G>T |  |  |  |  |  |  |  | C>T |  |

|  |  |  |  |  |  |  |  |  |  |  |  |  |  |  |  |  |  |  |  |  |  |  |  |  |  |  |  |  |  |
| --- | --- | --- | --- | --- | --- | --- | --- | --- | --- | --- | --- | --- | --- | --- | --- | --- | --- | --- | --- | --- | --- | --- | --- | --- | --- | --- | --- | --- | --- |
| pos | 527 | 1588 | 2632 | 3466 | 3476 | 4237 | 6726 | 7851 | 8110 | 11036 | 11860 | 13210 | 14742 | 15654 | 17040 | 20451 | 21137 | 21372 | 22027 | 22029 | 22030 | 22031 | 22032 | 22033 | 22034 | 25889 | 27604 | 29095 |  |
| ref | C | A | G | T | G | T | C | C | T | C | A | T | T | C | T | C | A | G | T | A | G | T | T | C | A | C | G | C |  |
| AY.4 | T | A | G | C | G | C | C | T | C | T | G | C | C | C | C | T | G | G | T | - | - | - | - | - | - | T | G | T |  |
| AY.4 | T | A | G | C | G | C | C | T | C | T | G | C | C | C | C | T | G | G | G | A | A | A | G | T | G | T | A | T | #698139 |
| AY.39 | C | G | T | T | A | T | T | C | T | C | A | T | T | T | T | C | A | T | G | A | A | A | G | T | G | C | A | C |  |
| mut |  |  |  |  |  |  |  |  |  |  |  |  |  |  |  |  |  |  |  |  |  |  |  |  |  | C>T |  | C>T |  |

|  |  |  |  |  |  |  |  |  |  |  |  |  |  |  |  |  |  |  |  |  |  |  |  |
| --- | --- | --- | --- | --- | --- | --- | --- | --- | --- | --- | --- | --- | --- | --- | --- | --- | --- | --- | --- | --- | --- | --- | --- |
| pos | 487 | 683 | 884 | 4124 | 4720 | 7259 | 7851 | 13297 | 13385 | 16914 | 17040 | 19677 | 19862 | 22027 | 22029 | 22030 | 22031 | 22032 | 22033 | 22034 | 26124 | 28632 |  |
| ref | G | C | C | G | G | A | C | C | G | G | T | G | C | T | A | G | T | T | C | A | C | G |  |
| AY.4 | A | T | C | A | G | G | T | T | A | G | C | G | T | T | - | - | - | - | - | - | A | C |  |
| AY.4 | A | T | C | A | G | G | T | T | A | G | C | G | T | G | A | A | A | G | T | G | A | C | #698159 |
| AY.114 | G | C | T | G | T | A | C | C | G | T | T | T | C | G | A | A | A | G | T | G | C | G |  |
| mut |  |  |  |  |  |  |  |  |  |  |  |  |  |  |  |  |  |  |  |  | C>A | G>C |  |

|  |  |  |  |  |  |  |  |  |  |  |  |  |  |  |  |  |  |  |
| --- | --- | --- | --- | --- | --- | --- | --- | --- | --- | --- | --- | --- | --- | --- | --- | --- | --- | --- |
| pos | 809 | 884 | 1312 | 3632 | 4720 | 7693 | 7851 | 16914 | 17040 | 19677 | 22027 | 22029 | 22030 | 22031 | 22032 | 22033 | 22034 |  |
| ref | T | C | G | A | G | G | C | G | T | G | T | A | G | T | T | C | A |  |
| AY.4 | C | C | T | G | G | T | T | G | C | G | T | - | - | - | - | - | - |  |
| AY.4 | C | C | T | G | G | T | T | G | C | G | G | A | A | A | G | T | G | #698161 |
| AY.114 | T | T | G | A | T | G | C | T | T | T | G | A | A | A | G | T | G |  |
| mut |  |  |  |  |  |  |  |  |  |  |  |  |  |  |  |  |  |  |

|  |  |  |  |  |  |  |  |  |  |  |  |  |  |  |  |  |  |  |  |  |  |  |  |
| --- | --- | --- | --- | --- | --- | --- | --- | --- | --- | --- | --- | --- | --- | --- | --- | --- | --- | --- | --- | --- | --- | --- | --- |
| pos | 884 | 4720 | 5653 | 7851 | 8106 | 15755 | 16338 | 16622 | 16914 | 17826 | 19677 | 19893 | 21811 | 22027 | 22029 | 22030 | 22031 | 22032 | 22033 | 22034 | 26828 | 27722 |  |
| ref | C | G | T | C | C | C | C | G | G | T | G | T | C | T | A | G | T | T | C | A | G | T |  |
| AY.4 | C | G | C | T | T | T | T | A | G | C | G | C | T | T | - | - | - | - | - | - | T | C |  |
| AY.4 | C | G | C | T | T | T | T | A | G | C | G | C | T | G | A | A | A | G | T | G | T | C | #698164 |
| AY.114 | T | T | T | C | C | C | C | G | T | T | T | T | C | G | A | A | A | G | T | G | G | T |  |
| mut |  |  |  |  |  |  |  |  |  |  |  |  |  |  |  |  |  |  |  |  | G>T | T>C |  |

|  |  |  |  |  |  |  |  |  |  |  |  |  |  |  |  |  |  |  |  |  |  |  |  |  |
| --- | --- | --- | --- | --- | --- | --- | --- | --- | --- | --- | --- | --- | --- | --- | --- | --- | --- | --- | --- | --- | --- | --- | --- | --- |
| pos | 2083 | 3551 | 6040 | 6638 | 7712 | 7926 | 12970 | 14014 | 16726 | 17236 | 20251 | 23551 | 24208 | 25563 | 27291 | 28073 | 28909 | 29700 | 29762 | 29769 | 29774 | 29779 | 29781 |  |
| ref | T | C | C | C | C | C | C | T | C | A | A | C | C | G | T | G | C | A | C | C | C | G | G |  |
| AY.103 | T | T | C | C | C | C | C | T | C | G | A | C | T | G | C | G | T | A | C | C | C | G | G |  |
| B.1.617.2 | T | T | C | C | T | C | T | T | C | G | G | T | C | T | T | T | C | - | - | - | - | - | - | #701359 |
| AY.44 | C | C | T | T | C | T | C | G | T | A | A | T | C | T | T | T | C | G | C | C | C | G | G |  |
| mut |  |  |  |  | C>T |  | C>T |  |  |  | A>G |  |  |  |  |  |  | G>- | C>- | C>- | C>- | G>- | G>- |  |

|  |  |  |  |  |  |  |  |  |  |  |  |  |  |  |
| --- | --- | --- | --- | --- | --- | --- | --- | --- | --- | --- | --- | --- | --- | --- |
| pos | 3262 | 6312 | 7471 | 20679 | 22027 | 22029 | 22030 | 22031 | 22032 | 22033 | 22034 | 25249 | 28889 |  |
| ref | A | C | C | G | T | A | G | T | T | C | A | G | T |  |
| AY.4 | T | T | T | A | T | - | - | - | - | - | - | T | C |  |
| AY.4 | T | T | T | A | G | A | A | A | G | T | G | T | C | #707306 |
| AY.4 | A | C | C | G | G | A | A | A | G | T | G | G | T |  |
| mut |  |  |  |  |  |  |  |  |  |  |  | G>T | T>C |  |

|  |  |  |  |  |  |  |  |  |  |  |  |  |  |  |  |  |  |  |  |  |  |  |
| --- | --- | --- | --- | --- | --- | --- | --- | --- | --- | --- | --- | --- | --- | --- | --- | --- | --- | --- | --- | --- | --- | --- |
| pos | 1762 | 2923 | 6478 | 6883 | 8964 | 10747 | 12025 | 12529 | 14273 | 16192 | 21710 | 22027 | 22029 | 22030 | 22031 | 22032 | 22033 | 22034 | 22444 | 24781 | 29628 |  |
| ref | C | A | T | C | C | C | C | A | A | C | T | T | A | G | T | T | C | A | C | G | G |  |
| AY.39 | T | A | T | T | C | C | C | G | A | T | G | T | - | - | - | - | - | - | C | C | A |  |
| AY.39 | T | A | T | T | C | C | C | G | A | T | G | G | A | A | A | G | T | G | T | C | A | #710601 |
| AY.39 | C | G | C | C | T | T | T | A | C | C | T | G | A | A | A | G | T | G | T | G | G |  |
| mut |  |  |  |  |  |  |  |  |  |  |  |  |  |  |  |  |  |  |  | G>C | G>A |  |

|  |  |  |  |  |  |  |  |  |  |  |  |  |  |  |  |  |  |  |  |  |  |
| --- | --- | --- | --- | --- | --- | --- | --- | --- | --- | --- | --- | --- | --- | --- | --- | --- | --- | --- | --- | --- | --- |
| pos | 7851 | 9165 | 10323 | 11782 | 12772 | 15240 | 16733 | 19662 | 19735 | 21697 | 22026 | 22027 | 22029 | 22030 | 22031 | 22032 | 22033 | 22034 | 23524 | 28960 |  |
| ref | C | C | A | A | A | C | C | C | G | C | G | T | A | G | T | T | C | A | A | G |  |
| AY.113 | C | T | G | G | G | T | T | T | T | T | G | T | - | - | - | - | - | - | G | G |  |
| AY.113 | C | T | G | G | G | T | T | T | T | T | T | G | A | A | A | G | T | G | G | T | #710835 |
| AY.4 | T | C | A | A | A | C | C | C | G | C | T | G | A | A | A | G | T | G | A | G |  |
| mut |  |  |  |  |  |  |  |  |  |  |  |  |  |  |  |  |  |  | A>G | G>T |  |

|  |  |  |  |  |  |  |  |  |  |  |  |  |  |  |  |  |  |
| --- | --- | --- | --- | --- | --- | --- | --- | --- | --- | --- | --- | --- | --- | --- | --- | --- | --- |
| pos | 2716 | 7936 | 10202 | 11095 | 18252 | 22027 | 22029 | 22030 | 22031 | 22032 | 22033 | 22034 | 25897 | 27575 | 27576 | 29675 |  |
| ref | C | G | C | C | C | T | A | G | T | T | C | A | A | C | T | C |  |
| AY.120.1 | T | T | T | T | T | T | - | - | - | - | - | - | G | A | C | T |  |
| AY.120.1 | T | T | T | T | T | G | A | A | A | G | T | G | G | C | T | T | #716032 |
| AY.120 | C | G | C | C | C | G | A | A | A | G | T | G | A | C | T | C |  |
| mut |  |  |  |  |  |  |  |  |  |  |  |  | A>G |  |  | C>T |  |

|  |  |  |  |  |  |  |  |  |  |  |  |  |  |  |  |  |  |  |  |
| --- | --- | --- | --- | --- | --- | --- | --- | --- | --- | --- | --- | --- | --- | --- | --- | --- | --- | --- | --- |
| pos | 1372 | 1513 | 2037 | 2591 | 6330 | 7764 | 9611 | 21108 | 22026 | 22027 | 22029 | 22030 | 22031 | 22032 | 22033 | 22034 | 23309 | 28006 |  |
| ref | T | C | C | G | C | C | C | C | G | T | A | G | T | T | C | A | G | C |  |
| AY.100 | C | T | T | A | T | T | T | C | G | T | - | - | - | - | - | - | G | T |  |
| AY.100 | C | T | T | A | T | T | T | T | G | T | A | A | A | G | T | G | C | T | #716139 |
| AY.100 | T | C | C | G | C | C | C | C | T | G | A | A | A | G | T | G | G | C |  |
| mut |  |  |  |  |  |  |  | C>T |  |  |  |  |  |  |  |  | G>C | C>T |  |

|  |  |  |  |  |  |  |  |  |  |  |  |  |  |  |  |  |  |  |  |  |  |  |  |  |  |
| --- | --- | --- | --- | --- | --- | --- | --- | --- | --- | --- | --- | --- | --- | --- | --- | --- | --- | --- | --- | --- | --- | --- | --- | --- | --- |
| pos | 408 | 1820 | 3523 | 5668 | 5730 | 6419 | 6647 | 13716 | 17790 | 17913 | 22027 | 22029 | 22030 | 22031 | 22032 | 22033 | 22034 | 25069 | 29700 | 29762 | 29769 | 29774 | 29779 | 29781 |  |
| ref | A | G | A | G | C | G | G | T | G | C | T | A | G | T | T | C | A | C | A | C | C | C | G | G |  |
| AY.117 | T | A | A | G | T | G | G | C | T | C | T | - | - | - | - | - | - | T | - | - | - | - | - | - |  |
| AY.117 | T | A | A | G | T | G | A | C | T | C | G | A | A | A | G | T | G | T | A | C | C | C | G | G | #716338 |
| AY.117 | A | G | C | A | C | A | G | T | G | T | G | A | A | A | G | T | G | C | A | C | C | C | G | G |  |
| mut |  |  |  |  |  |  | G>A |  |  |  |  |  |  |  |  |  |  | C>T |  |  |  |  |  |  |  |

|  |  |  |  |  |  |  |  |  |  |  |  |  |  |  |  |  |  |  |  |  |  |  |
| --- | --- | --- | --- | --- | --- | --- | --- | --- | --- | --- | --- | --- | --- | --- | --- | --- | --- | --- | --- | --- | --- | --- |
| pos | 44 | 1068 | 1722 | 4572 | 7936 | 8131 | 8139 | 9127 | 10009 | 10030 | 18108 | 22026 | 22027 | 22029 | 22030 | 22031 | 22032 | 22033 | 22034 | 27384 | 29422 |  |
| ref | C | G | C | A | G | G | C | T | T | C | A | G | T | A | G | T | T | C | A | T | G |  |
| AY.100 | C | T | C | G | G | T | T | T | T | T | A | G | T | - | - | - | - | - | - | T | T |  |
| AY.100 | C | T | C | G | G | T | T | T | T | T | A | T | G | A | A | A | G | T | G | T | G | #720731 |
| AY.100 | T | G | T | A | T | G | C | C | C | C | T | T | G | A | A | A | G | T | G | - | G |  |
| mut |  |  |  |  |  |  |  |  |  |  |  |  |  |  |  |  |  |  |  | ->T |  |  |

|  |  |  |  |  |  |  |  |  |  |  |  |  |  |  |  |  |  |  |  |  |  |
| --- | --- | --- | --- | --- | --- | --- | --- | --- | --- | --- | --- | --- | --- | --- | --- | --- | --- | --- | --- | --- | --- |
| pos | 912 | 2706 | 5365 | 5623 | 9559 | 17040 | 18927 | 19359 | 19596 | 20374 | 22027 | 22029 | 22030 | 22031 | 22032 | 22033 | 22034 | 22899 | 27247 | 29700 |  |
| ref | C | C | C | T | C | T | T | A | C | A | T | A | G | T | T | C | A | G | C | A |  |
| AY.4 | T | C | C | C | T | T | T | A | T | A | T | - | - | - | - | - | - | G | T | A |  |
| AY.4 | T | C | C | C | T | T | T | G | T | A | G | A | A | A | G | T | G | T | T | G | #723113 |
| AY.4.1 | C | T | T | T | C | C | C | A | C | C | G | A | A | A | G | T | G | T | C | A |  |
| mut |  |  |  |  |  |  |  | A>G |  |  |  |  |  |  |  |  |  |  | C>T | A>G |  |

|  |  |  |  |  |  |  |  |  |  |  |  |  |  |  |
| --- | --- | --- | --- | --- | --- | --- | --- | --- | --- | --- | --- | --- | --- | --- |
| pos | 12832 | 13812 | 18032 | 19524 | 20390 | 22027 | 22029 | 22030 | 22031 | 22032 | 22033 | 22034 | 26753 |  |
| ref | G | G | C | C | G | T | A | G | T | T | C | A | C |  |
| AY.4 | A | T | T | T | T | T | - | - | - | - | - | - | T |  |
| AY.4 | A | T | T | T | T | G | A | A | A | G | T | G | T | #723154 |
| AY.4 | G | G | C | C | G | G | A | A | A | G | T | G | C |  |
| mut |  |  |  |  |  |  |  |  |  |  |  |  | C>T |  |

|  |  |  |  |  |  |  |  |  |  |  |  |  |  |  |  |  |  |  |  |  |  |  |  |  |
| --- | --- | --- | --- | --- | --- | --- | --- | --- | --- | --- | --- | --- | --- | --- | --- | --- | --- | --- | --- | --- | --- | --- | --- | --- |
| pos | 691 | 1043 | 2246 | 3559 | 6473 | 7851 | 9979 | 15654 | 16929 | 17040 | 18714 | 21372 | 22026 | 22027 | 22029 | 22030 | 22031 | 22032 | 22033 | 22034 | 23997 | 25386 | 27604 |  |
| ref | A | G | G | A | G | C | C | C | A | T | C | G | G | T | A | G | T | T | C | A | C | C | G |  |
| AY.4 | G | A | A | A | A | T | C | C | G | C | T | G | G | T | - | - | - | - | - | - | C | T | G |  |
| AY.4 | G | A | A | A | A | T | C | C | G | C | T | G | G | G | A | A | A | G | T | G | T | T | G | #723156 |
| AY.39 | A | G | G | C | G | C | T | T | A | T | C | T | T | G | A | A | A | G | T | G | C | C | A |  |
| mut |  |  |  |  |  |  |  |  |  |  |  |  |  |  |  |  |  |  |  |  | C>T | C>T | A>G |  |

|  |  |  |  |  |  |  |  |  |  |  |  |  |  |  |  |  |  |  |  |  |  |  |  |
| --- | --- | --- | --- | --- | --- | --- | --- | --- | --- | --- | --- | --- | --- | --- | --- | --- | --- | --- | --- | --- | --- | --- | --- |
| pos | 884 | 1405 | 1479 | 2485 | 4237 | 4720 | 7851 | 10279 | 13458 | 16864 | 16914 | 17040 | 19677 | 20262 | 22027 | 22029 | 22030 | 22031 | 22032 | 22033 | 22034 | 24040 |  |
| ref | C | T | C | C | T | G | C | C | C | G | G | T | G | A | T | A | G | T | T | C | A | G |  |
| AY.4 | C | C | T | T | C | G | T | T | T | A | G | C | G | G | T | - | - | - | - | - | - | T |  |
| AY.4 | C | C | T | T | C | G | T | T | T | A | G | C | G | G | G | A | A | A | G | T | G | T | #723218 |
| AY.114 | T | T | C | C | T | T | C | C | C | G | T | T | T | A | G | A | A | A | G | T | G | G |  |
| mut |  |  |  |  |  |  |  |  |  |  |  |  |  |  |  |  |  |  |  |  |  | G>T |  |

|  |  |  |  |  |  |  |  |  |  |  |  |  |  |  |  |  |  |  |  |  |  |  |  |
| --- | --- | --- | --- | --- | --- | --- | --- | --- | --- | --- | --- | --- | --- | --- | --- | --- | --- | --- | --- | --- | --- | --- | --- |
| pos | 884 | 4720 | 4792 | 7851 | 14475 | 15006 | 15843 | 16914 | 17019 | 17040 | 18255 | 19677 | 21987 | 22027 | 22029 | 22030 | 22031 | 22032 | 22033 | 22034 | 25595 | 27393 |  |
| ref | C | G | T | C | A | G | T | G | G | T | G | G | G | T | A | G | T | T | C | A | G | C |  |
| AY.4 | C | G | C | T | G | T | T | G | T | C | T | G | G | T | - | - | - | - | - | - | T | T |  |
| AY.4 | C | G | C | T | G | T | C | G | T | C | T | G | G | G | A | A | A | G | T | G | T | T | #723237 |
| AY.114 | T | T | T | C | A | G | T | T | G | T | G | T | A | G | A | A | A | G | T | G | G | C |  |
| mut |  |  |  |  |  |  | T>C |  |  |  |  |  |  |  |  |  |  |  |  |  | G>T | C>T |  |

|  |  |  |  |  |  |  |  |  |  |  |  |  |  |  |  |  |  |  |  |
| --- | --- | --- | --- | --- | --- | --- | --- | --- | --- | --- | --- | --- | --- | --- | --- | --- | --- | --- | --- |
| pos | 1474 | 6500 | 7851 | 8699 | 14471 | 15952 | 16054 | 17259 | 18744 | 19069 | 22027 | 22029 | 22030 | 22031 | 22032 | 22033 | 22034 | 28299 |  |
| ref | T | C | C | A | C | C | C | G | C | G | T | A | G | T | T | C | A | A |  |
| AY.4 | C | C | T | A | T | C | T | G | C | T | T | - | - | - | - | - | - | A |  |
| AY.4 | C | C | T | A | T | C | T | G | C | T | G | A | A | A | G | T | G | T | #726333 |
| AY.43.8 | T | T | C | G | C | A | C | T | T | G | G | A | A | A | G | T | G | T |  |
| mut |  |  |  |  |  |  |  |  |  |  |  |  |  |  |  |  |  |  |  |

|  |  |  |  |  |  |  |  |  |  |  |  |  |  |  |  |  |  |  |  |  |
| --- | --- | --- | --- | --- | --- | --- | --- | --- | --- | --- | --- | --- | --- | --- | --- | --- | --- | --- | --- | --- |
| pos | 884 | 2348 | 3493 | 4720 | 7851 | 8027 | 13857 | 16914 | 19677 | 21974 | 22027 | 22029 | 22030 | 22031 | 22032 | 22033 | 22034 | 23042 | 28077 |  |
| ref | C | A | T | G | C | G | T | G | G | G | T | A | G | T | T | C | A | T | G |  |
| AY.4 | C | G | C | G | T | T | C | G | G | C | T | - | - | - | - | - | - | C | C |  |
| AY.4 | C | G | C | G | T | T | C | G | G | C | G | A | A | A | G | T | G | C | C | #726366 |
| AY.114 | T | A | T | T | C | G | T | T | T | G | G | A | A | A | G | T | G | T | G |  |
| mut |  |  |  |  |  |  |  |  |  |  |  |  |  |  |  |  |  | T>C | G>C |  |

|  |  |  |  |  |  |  |  |  |  |  |  |  |  |  |  |  |  |  |  |  |  |
| --- | --- | --- | --- | --- | --- | --- | --- | --- | --- | --- | --- | --- | --- | --- | --- | --- | --- | --- | --- | --- | --- |
| pos | 526 | 920 | 1912 | 2402 | 4300 | 5184 | 6408 | 6885 | 8016 | 19524 | 21646 | 21846 | 25003 | 25792 | 26006 | 27005 | 27875 | 27881 | 27882 | 27883 |  |
| ref | G | C | C | T | G | C | C | G | C | C | C | C | A | C | G | C | T | C | G | C |  |
| AY.120 | G | T | T | A | G | T | T | C | T | C | C | T | G | C | G | C | T | C | G | C |  |
| AY.120 | G | T | T | A | G | T | T | C | T | C | C | T | G | C | G | T | C | T | C | T | #727176 |
| AY.98 | T | C | C | T | T | C | C | G | C | T | T | C | A | T | A | T | C | T | C | T |  |
| mut |  |  |  |  |  |  |  |  |  |  |  |  |  |  |  |  |  |  |  |  |  |

|  |  |  |  |  |  |  |  |  |  |  |  |  |  |  |  |  |  |  |  |  |  |
| --- | --- | --- | --- | --- | --- | --- | --- | --- | --- | --- | --- | --- | --- | --- | --- | --- | --- | --- | --- | --- | --- |
| pos | 805 | 3069 | 4399 | 4681 | 5869 | 6841 | 7851 | 13711 | 18744 | 21987 | 22027 | 22029 | 22030 | 22031 | 22032 | 22033 | 22034 | 25549 | 26915 | 28378 |  |
| ref | G | A | G | A | C | A | C | A | C | G | T | A | G | T | T | C | A | C | A | G |  |
| B.1.617.2 | T | A | T | G | C | G | C | G | C | G | T | - | - | - | - | - | - | T | G | T |  |
| B.1.617.2 | T | A | T | G | C | G | C | G | C | G | G | A | A | A | G | T | G | T | G | T | #729650 |
| AY.4 | G | G | G | A | T | A | T | A | T | A | G | A | A | A | G | T | G | C | A | G |  |
| mut |  |  |  |  |  |  |  |  |  |  |  |  |  |  |  |  |  | C>T | A>G | G>T |  |

|  |  |  |  |  |  |  |  |  |  |  |  |  |  |
| --- | --- | --- | --- | --- | --- | --- | --- | --- | --- | --- | --- | --- | --- |
| pos | 337 | 22027 | 22029 | 22030 | 22031 | 22032 | 22033 | 22034 | 24620 | 24789 | 25555 | 27967 |  |
| ref | C | T | A | G | T | T | C | A | G | C | G | G |  |
| AY.4 | C | G | A | A | A | G | T | G | G | C | T | G |  |
| AY.4 | T | G | A | A | A | G | T | G | T | T | G | T | #729654 |
| AY.4 | T | T | - | - | - | - | - | - | T | T | G | T |  |
| mut | C>T |  |  |  |  |  |  |  |  |  |  |  |  |

|  |  |  |  |  |  |  |  |  |  |  |  |  |
| --- | --- | --- | --- | --- | --- | --- | --- | --- | --- | --- | --- | --- |
| pos | 526 | 7851 | 19524 | 21846 | 23481 | 23530 | 27005 | 27875 | 27881 | 27882 | 27883 |  |
| ref | G | C | C | C | C | C | C | T | C | G | C |  |
| AY.4 | G | T | C | T | T | T | T | T | C | G | C |  |
| AY.4 | G | T | C | T | T | T | T | C | T | C | T | #731990 |
| AY.98 | T | C | T | C | C | C | C | C | T | C | T |  |
| mut |  |  |  |  |  |  |  |  |  |  |  |  |

|  |  |  |  |  |  |  |  |  |  |  |  |  |  |  |
| --- | --- | --- | --- | --- | --- | --- | --- | --- | --- | --- | --- | --- | --- | --- |
| pos | 100 | 526 | 3076 | 4510 | 19524 | 24928 | 26230 | 27390 | 27875 | 27881 | 27882 | 27883 | 29253 |  |
| ref | C | G | A | G | C | G | G | G | T | C | G | C | C |  |
| AY.5 | C | G | G | A | C | T | T | T | T | C | G | C | T |  |
| AY.5 | T | G | G | A | C | T | T | T | C | T | C | T | T | #736357 |
| AY.98 | C | T | A | G | T | G | G | G | C | T | C | T | C |  |
| mut | C>T |  |  |  |  |  |  |  |  |  |  |  | C>T |  |

|  |  |  |  |  |  |  |  |  |  |  |  |  |  |  |  |  |  |  |  |
| --- | --- | --- | --- | --- | --- | --- | --- | --- | --- | --- | --- | --- | --- | --- | --- | --- | --- | --- | --- |
| pos | 4402 | 6552 | 9191 | 12756 | 13774 | 15978 | 16700 | 20759 | 22026 | 22027 | 22029 | 22030 | 22031 | 22032 | 22033 | 22034 | 24197 | 27377 |  |
| ref | T | T | G | C | C | T | G | C | G | T | A | G | T | T | C | A | G | A |  |
| AY.119 | C | C | T | T | T | T | A | T | G | T | - | - | - | - | - | - | T | T |  |
| AY.119 | C | C | T | T | T | C | A | T | T | G | A | A | A | G | T | G | T | T | #743493 |
| AY.119 | T | T | G | C | C | T | G | C | T | G | A | A | A | G | T | G | G | A |  |
| mut |  |  |  |  |  | T>C |  |  |  |  |  |  |  |  |  |  | G>T | A>T |  |

|  |  |  |  |  |  |  |  |  |  |  |  |  |  |  |  |  |  |
| --- | --- | --- | --- | --- | --- | --- | --- | --- | --- | --- | --- | --- | --- | --- | --- | --- | --- |
| pos | 3495 | 12193 | 12961 | 16575 | 17863 | 19841 | 22027 | 22029 | 22030 | 22031 | 22032 | 22033 | 22034 | 24781 | 27981 | 28690 |  |
| ref | A | T | A | C | T | A | T | A | G | T | T | C | A | G | C | G |  |
| AY.119 | G | C | G | T | C | G | T | - | - | - | - | - | - | G | A | A |  |
| AY.119 | G | C | G | T | C | G | G | A | A | A | G | T | G | C | A | A | #743689 |
| AY.119 | A | T | A | C | T | A | G | A | A | A | G | T | G | G | C | G |  |
| mut |  |  |  |  |  |  |  |  |  |  |  |  |  | G>C | C>A | G>A |  |

|  |  |  |  |  |  |  |  |  |  |  |  |  |  |  |  |
| --- | --- | --- | --- | --- | --- | --- | --- | --- | --- | --- | --- | --- | --- | --- | --- |
| pos | 427 | 910 | 911 | 912 | 4237 | 5392 | 8299 | 11620 | 14055 | 15346 | 16725 | 17040 | 24780 | 26022 |  |
| ref | A | G | T | C | T | C | C | C | G | C | A | T | A | C |  |
| AY.4 | G | G | T | C | C | C | C | T | G | C | A | C | A | T |  |
| AY.4 | G | G | T | C | C | C | C | C | T | T | G | T | C | C | #746901 |
| AY.110 | A | A | A | A | T | T | T | C | T | T | G | T | C | C |  |
| mut |  |  |  |  |  |  |  |  |  |  |  |  |  |  |  |

|  |  |  |  |  |  |  |  |  |  |  |  |  |  |  |  |  |
| --- | --- | --- | --- | --- | --- | --- | --- | --- | --- | --- | --- | --- | --- | --- | --- | --- |
| pos | 1911 | 4540 | 5654 | 7800 | 14511 | 21412 | 22026 | 22027 | 22029 | 22030 | 22031 | 22032 | 22033 | 22034 | 28245 |  |
| ref | C | C | C | C | G | C | G | T | A | G | T | T | C | A | T |  |
| AY.4 | T | C | T | T | T | C | G | T | - | - | - | - | - | - | T |  |
| AY.4 | T | C | T | T | T | A | G | G | A | A | A | G | T | G | T | #747335 |
| AY.4 | C | T | C | C | G | C | T | G | A | A | A | G | T | G | G |  |
| mut |  |  |  |  |  | C>A |  |  |  |  |  |  |  |  | G>T |  |

|  |  |  |  |  |  |  |  |  |  |  |  |  |  |  |  |  |  |  |  |  |
| --- | --- | --- | --- | --- | --- | --- | --- | --- | --- | --- | --- | --- | --- | --- | --- | --- | --- | --- | --- | --- |
| pos | 280 | 2202 | 3543 | 4901 | 5812 | 10432 | 10626 | 11235 | 16662 | 21987 | 22026 | 22027 | 22029 | 22030 | 22031 | 22032 | 22033 | 22034 | 25273 |  |
| ref | C | G | C | C | C | C | C | C | G | G | G | T | A | G | T | T | C | A | G |  |
| AY.100 | T | T | A | T | T | T | T | C | A | G | G | T | - | - | - | - | - | - | G |  |
| AY.100 | T | T | A | T | T | T | T | T | G | A | T | G | A | A | A | G | T | G | T | #748738 |
| AY.100 | C | G | C | C | C | C | C | C | G | A | T | G | A | A | A | G | T | G | G |  |
| mut |  |  |  |  |  |  |  | C>T |  |  |  |  |  |  |  |  |  |  | G>T |  |

|  |  |  |  |  |  |  |  |  |  |  |  |  |  |  |  |  |
| --- | --- | --- | --- | --- | --- | --- | --- | --- | --- | --- | --- | --- | --- | --- | --- | --- |
| pos | 5817 | 7081 | 17019 | 19788 | 22026 | 22027 | 22029 | 22030 | 22031 | 22032 | 22033 | 22034 | 25904 | 25906 | 28507 |  |
| ref | C | C | G | G | G | T | A | G | T | T | C | A | C | G | C |  |
| AY.100 | T | T | T | A | G | T | - | - | - | - | - | - | T | T | C |  |
| AY.100 | T | T | T | A | G | G | A | A | A | G | T | G | T | T | T | #748794 |
| AY.100 | C | C | G | G | T | G | A | A | A | G | T | G | C | G | C |  |
| mut |  |  |  |  |  |  |  |  |  |  |  |  | C>T | G>T | C>T |  |

|  |  |  |  |  |  |  |  |  |  |  |  |  |  |  |  |  |  |
| --- | --- | --- | --- | --- | --- | --- | --- | --- | --- | --- | --- | --- | --- | --- | --- | --- | --- |
| pos | 367 | 1945 | 2365 | 5986 | 7926 | 12817 | 14259 | 22026 | 22027 | 22029 | 22030 | 22031 | 22032 | 22033 | 22034 | 29555 |  |
| ref | C | T | T | C | C | G | T | G | T | A | G | T | T | C | A | C |  |
| AY.39 | C | T | C | T | T | G | T | G | T | - | - | - | - | - | - | T |  |
| AY.39 | C | G | C | T | T | G | T | G | G | A | A | A | G | T | G | T | #748874 |
| AY.39 | T | T | T | C | C | T | C | T | G | A | A | A | G | T | G | C |  |
| mut |  | T>G |  |  |  |  |  |  |  |  |  |  |  |  |  | C>T |  |

|  |  |  |  |  |  |  |  |  |  |  |  |  |  |  |  |
| --- | --- | --- | --- | --- | --- | --- | --- | --- | --- | --- | --- | --- | --- | --- | --- |
| pos | 1950 | 9085 | 14708 | 15779 | 15960 | 21987 | 22026 | 22027 | 22029 | 22030 | 22031 | 22032 | 22033 | 22034 |  |
| ref | T | A | C | A | C | G | G | T | A | G | T | T | C | A |  |
| AY.39 | C | G | T | G | T | G | G | T | - | - | - | - | - | - |  |
| AY.39 | C | G | T | G | T | G | G | G | A | A | A | G | T | G | #748969 |
| AY.39 | T | A | C | A | C | A | T | G | A | A | A | G | T | G |  |
| mut |  |  |  |  |  |  |  |  |  |  |  |  |  |  |  |

|  |  |  |  |  |  |  |  |  |  |  |  |  |  |  |  |  |  |  |
| --- | --- | --- | --- | --- | --- | --- | --- | --- | --- | --- | --- | --- | --- | --- | --- | --- | --- | --- |
| pos | 5192 | 5884 | 8215 | 8344 | 12809 | 14184 | 19164 | 21646 | 22026 | 22027 | 22029 | 22030 | 22031 | 22032 | 22033 | 22034 | 28048 |  |
| ref | C | C | T | C | C | C | C | C | G | T | A | G | T | T | C | A | G |  |
| AY.100 | T | T | C | T | T | T | T | T | G | T | - | - | - | - | - | - | T |  |
| AY.100 | T | T | C | T | T | T | T | T | G | G | A | A | A | G | T | G | T | #748989 |
| AY.100 | C | C | T | C | C | C | C | C | T | G | A | A | A | G | T | G | G |  |
| mut |  |  |  |  |  |  |  |  |  |  |  |  |  |  |  |  | G>T |  |

|  |  |  |  |  |  |  |  |  |  |  |  |  |  |  |  |  |  |  |  |  |  |  |  |  |  |
| --- | --- | --- | --- | --- | --- | --- | --- | --- | --- | --- | --- | --- | --- | --- | --- | --- | --- | --- | --- | --- | --- | --- | --- | --- | --- |
| pos | 44 | 1722 | 5192 | 5884 | 7936 | 8215 | 8344 | 9127 | 10009 | 12809 | 14184 | 18108 | 19164 | 21646 | 22026 | 22027 | 22029 | 22030 | 22031 | 22032 | 22033 | 22034 | 27384 | 28048 |  |
| ref | C | C | C | C | G | T | C | T | T | C | C | A | C | C | G | T | A | G | T | T | C | A | T | G |  |
| AY.100 | C | C | T | T | G | C | T | T | T | T | T | A | T | T | G | T | - | - | - | - | - | - | T | T |  |
| AY.100 | C | C | T | T | G | C | T | T | T | T | T | A | T | T | T | G | A | A | A | G | T | G | T | T | #748991 |
| AY.100 | T | T | C | C | T | T | C | C | C | C | C | T | C | C | T | G | A | A | A | G | T | G | - | G |  |
| mut |  |  |  |  |  |  |  |  |  |  |  |  |  |  |  |  |  |  |  |  |  |  | ->T | G>T |  |

|  |  |  |  |  |  |  |  |  |  |  |  |  |  |  |
| --- | --- | --- | --- | --- | --- | --- | --- | --- | --- | --- | --- | --- | --- | --- |
| pos | 7165 | 10605 | 11674 | 15279 | 20259 | 22026 | 22027 | 22029 | 22030 | 22031 | 22032 | 22033 | 22034 |  |
| ref | C | C | C | C | C | G | T | A | G | T | T | C | A |  |
| AY.39 | T | T | T | T | T | G | T | - | - | - | - | - | - |  |
| AY.39 | T | T | T | T | T | G | G | A | A | A | G | T | G | #748993 |
| AY.39 | C | C | C | C | C | T | G | A | A | A | G | T | G |  |
| mut |  |  |  |  |  |  |  |  |  |  |  |  |  |  |

|  |  |  |  |  |  |  |  |  |  |  |  |  |  |  |  |  |
| --- | --- | --- | --- | --- | --- | --- | --- | --- | --- | --- | --- | --- | --- | --- | --- | --- |
| pos | 526 | 599 | 7203 | 13482 | 19524 | 21261 | 21372 | 21846 | 22241 | 25855 | 27604 | 27875 | 27881 | 27882 | 27883 |  |
| ref | G | G | C | A | C | A | G | C | G | G | G | T | C | G | C |  |
| AY.39.1 | G | T | T | G | C | G | T | T | A | T | A | T | C | G | C |  |
| AY.39.1 | G | T | T | G | C | G | T | T | A | T | A | C | T | C | T | #749380 |
| AY.98 | T | G | C | A | T | A | G | C | G | G | G | C | T | C | T |  |
| mut |  |  |  |  |  |  |  |  |  |  |  |  |  |  |  |  |

|  |  |  |  |  |  |  |  |  |  |  |  |  |  |  |  |  |  |  |  |  |  |  |  |
| --- | --- | --- | --- | --- | --- | --- | --- | --- | --- | --- | --- | --- | --- | --- | --- | --- | --- | --- | --- | --- | --- | --- | --- |
| pos | 1594 | 2197 | 2336 | 2825 | 3401 | 4100 | 4237 | 6286 | 6547 | 6720 | 7851 | 8561 | 14748 | 17040 | 17331 | 18029 | 19964 | 21627 | 21762 | 21846 | 22023 | 29535 |  |
| ref | C | C | G | A | G | T | T | C | T | C | C | G | A | T | G | C | T | C | C | C | A | C |  |
| AY.6 | T | T | A | G | A | T | T | T | T | C | C | G | A | T | T | C | C | T | T | C | A | T |  |
| AY.4 | T | T | A | G | A | C | C | C | C | T | T | T | G | C | G | T | T | C | C | T | C | C | #752005 |
| AY.4 | C | C | G | A | G | C | C | C | C | C | T | T | G | C | G | T | T | C | C | T | A | C |  |
| mut |  |  |  |  |  |  |  |  |  | C>T |  |  |  |  |  |  |  |  |  |  | A>C |  |  |

|  |  |  |  |  |  |  |  |  |  |  |  |  |  |  |  |
| --- | --- | --- | --- | --- | --- | --- | --- | --- | --- | --- | --- | --- | --- | --- | --- |
| pos | 1885 | 10723 | 12789 | 14653 | 16185 | 22026 | 22027 | 22029 | 22030 | 22031 | 22032 | 22033 | 22034 | 22687 |  |
| ref | T | T | C | G | T | G | T | A | G | T | T | C | A | C |  |
| AY.100 | G | A | T | T | C | G | T | - | - | - | - | - | - | T |  |
| AY.100 | G | A | T | T | C | G | G | A | A | A | G | T | G | T | #753831 |
| AY.100 | T | T | C | G | T | T | G | A | A | A | G | T | G | C |  |
| mut |  |  |  |  |  |  |  |  |  |  |  |  |  | C>T |  |

|  |  |  |  |  |  |  |  |  |  |  |  |  |  |  |  |  |  |  |  |  |  |  |
| --- | --- | --- | --- | --- | --- | --- | --- | --- | --- | --- | --- | --- | --- | --- | --- | --- | --- | --- | --- | --- | --- | --- |
| pos | 222 | 3304 | 3433 | 7945 | 8223 | 10870 | 16644 | 16993 | 17799 | 22026 | 22027 | 22029 | 22030 | 22031 | 22032 | 22033 | 22034 | 23415 | 27014 | 28310 | 29543 |  |
| ref | C | G | G | C | A | G | A | T | A | G | T | A | G | T | T | C | A | C | G | C | G |  |
| AY.39 | T | T | T | T | G | T | G | C | G | G | T | - | - | - | - | - | - | C | T | T | T |  |
| AY.39 | T | T | T | T | G | T | G | C | G | T | G | A | A | A | G | T | G | T | T | T | T | #753910 |
| AY.39 | C | G | G | C | A | G | A | T | A | T | G | A | A | A | G | T | G | C | G | C | G |  |
| mut |  |  |  |  |  |  |  |  |  |  |  |  |  |  |  |  |  | C>T | G>T | C>T | G>T |  |

|  |  |  |  |  |  |  |  |  |  |  |  |  |  |  |  |  |  |  |  |  |  |  |  |
| --- | --- | --- | --- | --- | --- | --- | --- | --- | --- | --- | --- | --- | --- | --- | --- | --- | --- | --- | --- | --- | --- | --- | --- |
| pos | 222 | 949 | 3448 | 4041 | 5704 | 6354 | 6622 | 13721 | 14478 | 14511 | 15952 | 17528 | 18010 | 18255 | 18744 | 21057 | 21846 | 24130 | 27131 | 28299 | 29700 | 29781 |  |
| ref | C | A | C | T | G | C | G | C | C | G | C | C | A | G | C | C | C | C | C | A | A | G |  |
| AY.43 | C | A | T | T | G | T | G | T | C | G | A | T | G | G | T | C | C | T | T | T | A | T |  |
| B.1.617.2 | C | A | T | T | G | T | A | T | T | C | A | C | A | T | C | T | T | C | C | A | A | T | #757547 |
| B.1.617.2 | T | G | C | G | A | C | A | C | T | C | C | C | A | G | C | T | T | C | C | A | G | G |  |
| mut |  |  |  |  |  |  | G>A |  |  |  | C>A |  |  | G>T |  |  |  |  |  |  | G>A | G>T |  |

|  |  |  |  |  |  |  |  |  |  |  |  |  |  |  |
| --- | --- | --- | --- | --- | --- | --- | --- | --- | --- | --- | --- | --- | --- | --- |
| pos | 2923 | 4024 | 5221 | 6424 | 12525 | 22027 | 22029 | 22030 | 22031 | 22032 | 22033 | 22034 | 25844 |  |
| ref | A | G | C | A | C | T | A | G | T | T | C | A | C |  |
| AY.36 | G | A | T | C | C | T | - | - | - | - | - | - | T |  |
| AY.36 | G | A | T | C | C | G | A | A | A | G | T | G | T | #757585 |
| AY.36 | A | G | C | A | T | G | A | A | A | G | T | G | C |  |
| mut |  |  |  |  |  |  |  |  |  |  |  |  | C>T |  |

|  |  |  |  |  |  |  |  |  |  |  |  |  |  |  |  |  |  |  |  |  |  |  |  |  |  |  |  |
| --- | --- | --- | --- | --- | --- | --- | --- | --- | --- | --- | --- | --- | --- | --- | --- | --- | --- | --- | --- | --- | --- | --- | --- | --- | --- | --- | --- |
| pos | 526 | 1420 | 3811 | 4333 | 6410 | 6622 | 7752 | 11417 | 14478 | 14511 | 15006 | 17733 | 18255 | 18744 | 19524 | 20483 | 20757 | 21057 | 21846 | 25518 | 25538 | 26054 | 26122 | 28378 | 29558 | 29736 |  |
| ref | G | C | C | A | G | G | T | G | C | G | G | C | G | C | C | C | T | C | C | T | G | C | G | G | A | G |  |
| AY.98 | T | C | T | G | A | G | C | T | C | G | T | T | G | C | T | C | C | C | C | T | G | C | T | G | A | C |  |
| AY.98 | T | C | T | G | A | G | C | T | C | G | T | T | G | C | T | C | C | C | C | C | T | A | G | T | T | G | #757612 |
| AY.127 | G | T | C | A | G | A | T | G | T | C | G | C | T | T | C | T | T | T | T | C | T | A | G | T | T | G |  |
| mut |  |  |  |  |  |  |  |  |  |  |  |  |  |  |  |  |  |  |  |  |  |  |  |  |  |  |  |

|  |  |  |  |  |  |  |  |  |  |  |  |  |  |  |
| --- | --- | --- | --- | --- | --- | --- | --- | --- | --- | --- | --- | --- | --- | --- |
| pos | 16357 | 18568 | 20250 | 21707 | 22026 | 22027 | 22029 | 22030 | 22031 | 22032 | 22033 | 22034 | 28462 |  |
| ref | T | C | A | C | G | T | A | G | T | T | C | A | C |  |
| AY.100 | C | T | G | T | G | T | - | - | - | - | - | - | T |  |
| AY.100 | C | T | G | T | G | G | A | A | A | G | T | G | T | #758699 |
| AY.100 | T | C | A | C | T | G | A | A | A | G | T | G | C |  |
| mut |  |  |  |  |  |  |  |  |  |  |  |  | C>T |  |

|  |  |  |  |  |  |  |  |  |  |  |  |  |  |  |
| --- | --- | --- | --- | --- | --- | --- | --- | --- | --- | --- | --- | --- | --- | --- |
| pos | 4719 | 6823 | 22027 | 22029 | 22030 | 22031 | 22032 | 22033 | 22034 | 23683 | 27297 | 27311 | 29332 |  |
| ref | C | A | T | A | G | T | T | C | A | C | C | T | G |  |
| AY.113 | C | A | G | A | A | A | G | T | G | C | C | T | G |  |
| AY.113 | T | G | G | A | A | A | G | T | G | T | T | C | T | #758814 |
| AY.113 | C | G | T | - | - | - | - | - | - | T | T | C | T |  |
| mut | C>T | A>G |  |  |  |  |  |  |  |  |  |  |  |  |

|  |  |  |  |  |  |  |  |  |  |  |  |  |  |  |  |  |  |  |  |  |  |  |  |
| --- | --- | --- | --- | --- | --- | --- | --- | --- | --- | --- | --- | --- | --- | --- | --- | --- | --- | --- | --- | --- | --- | --- | --- |
| pos | 44 | 653 | 1722 | 7936 | 9127 | 10009 | 16357 | 18108 | 18568 | 20250 | 21707 | 22026 | 22027 | 22029 | 22030 | 22031 | 22032 | 22033 | 22034 | 27384 | 28462 | 28774 |  |
| ref | C | G | C | G | T | T | T | A | C | A | C | G | T | A | G | T | T | C | A | T | C | G |  |
| AY.100 | C | G | C | G | T | T | C | A | T | G | T | G | T | - | - | - | - | - | - | T | T | G |  |
| AY.100 | C | A | C | G | T | T | C | A | T | G | T | T | G | A | A | A | G | T | G | T | T | T | #758852 |
| AY.100 | T | G | T | T | C | C | T | T | C | A | C | T | G | A | A | A | G | T | G | - | C | G |  |
| mut |  | G>A |  |  |  |  |  |  |  |  |  |  |  |  |  |  |  |  |  | ->T | C>T | G>T |  |

|  |  |  |  |  |  |  |  |  |  |  |  |  |  |  |  |  |  |  |  |  |  |  |
| --- | --- | --- | --- | --- | --- | --- | --- | --- | --- | --- | --- | --- | --- | --- | --- | --- | --- | --- | --- | --- | --- | --- |
| pos | 1239 | 2062 | 6288 | 8318 | 8597 | 8729 | 9907 | 10645 | 12049 | 15378 | 17048 | 20849 | 22026 | 22027 | 22029 | 22030 | 22031 | 22032 | 22033 | 22034 | 24077 |  |
| ref | A | C | G | C | C | T | C | C | C | T | A | A | G | T | A | G | T | T | C | A | G |  |
| AY.100 | A | T | T | A | A | C | T | T | T | C | G | C | G | T | - | - | - | - | - | - | G |  |
| AY.100 | G | T | T | A | A | C | T | T | T | C | G | C | G | G | A | A | A | G | T | G | T | #758982 |
| AY.100 | A | C | G | C | C | T | C | C | C | T | A | A | T | G | A | A | A | G | T | G | G |  |
| mut | A>G |  |  |  |  |  |  |  |  |  |  |  |  |  |  |  |  |  |  |  | G>T |  |

|  |  |  |  |  |  |  |  |  |  |  |  |  |  |  |  |  |  |  |  |  |  |  |
| --- | --- | --- | --- | --- | --- | --- | --- | --- | --- | --- | --- | --- | --- | --- | --- | --- | --- | --- | --- | --- | --- | --- |
| pos | 1170 | 1439 | 1912 | 6816 | 9684 | 9746 | 12793 | 21648 | 23120 | 26111 | 26176 | 28077 | 28237 | 28241 | 28243 | 28245 | 28247 | 28251 | 28252 | 28253 | 28254 |  |
| ref | C | G | C | G | T | C | G | C | G | C | C | G | G | T | T | T | A | T | T | C | A |  |
| AY.4 | C | A | C | G | T | T | G | T | T | T | C | G | G | T | T | T | A | - | - | - | A |  |
| AY.4 | C | A | T | G | T | T | T | T | T | T | C | G | T | A | A | C | G | - | T | T | C | #761421 |
| AY.4 | T | G | C | A | C | C | G | C | G | C | T | T | T | A | A | C | G | T | T | C | C |  |
| mut |  |  | C>T |  |  |  | G>T |  |  |  |  |  |  |  |  |  |  | T>- |  | C>T |  |  |

|  |  |  |  |  |  |  |  |  |  |  |  |  |  |  |  |  |
| --- | --- | --- | --- | --- | --- | --- | --- | --- | --- | --- | --- | --- | --- | --- | --- | --- |
| pos | 6726 | 11695 | 22026 | 22027 | 22029 | 22030 | 22031 | 22032 | 22033 | 22034 | 23664 | 25517 | 25519 | 28570 | 28748 |  |
| ref | C | T | G | T | A | G | T | T | C | A | C | C | T | T | C |  |
| AY.39 | C | T | T | G | A | A | A | G | T | G | C | C | T | T | C |  |
| AY.39 | T | C | T | G | A | A | A | G | T | G | T | T | C | C | T | #763292 |
| AY.39 | T | C | G | T | - | - | - | - | - | - | T | T | C | T | T |  |
| mut | C>T | T>C |  |  |  |  |  |  |  |  |  |  |  | T>C |  |  |

|  |  |  |  |  |  |  |  |  |  |  |  |  |  |  |  |
| --- | --- | --- | --- | --- | --- | --- | --- | --- | --- | --- | --- | --- | --- | --- | --- |
| pos | 14801 | 16795 | 22027 | 22029 | 22030 | 22031 | 22032 | 22033 | 22034 | 27107 | 27879 | 28392 | 28677 | 29521 |  |
| ref | A | G | T | A | G | T | T | C | A | A | C | G | C | A |  |
| AY.4 | A | G | G | A | A | A | G | T | G | A | C | G | C | A |  |
| AY.4 | C | T | G | A | A | A | G | T | G | G | T | T | C | G | #765830 |
| AY.4 | C | T | T | - | - | - | - | - | - | G | T | T | T | G |  |
| mut | A>C | G>T |  |  |  |  |  |  |  |  |  |  | T>C |  |  |

|  |  |  |  |  |  |  |  |  |  |  |  |  |  |  |  |
| --- | --- | --- | --- | --- | --- | --- | --- | --- | --- | --- | --- | --- | --- | --- | --- |
| pos | 3168 | 4051 | 9154 | 16275 | 19872 | 22026 | 22027 | 22029 | 22030 | 22031 | 22032 | 22033 | 22034 | 24064 |  |
| ref | C | T | A | A | G | G | T | A | G | T | T | C | A | C |  |
| AY.100 | T | C | G | G | T | G | T | - | - | - | - | - | - | T |  |
| AY.100 | T | C | G | G | T | G | G | A | A | A | G | T | G | T | #766920 |
| AY.100 | C | T | A | A | G | T | G | A | A | A | G | T | G | C |  |
| mut |  |  |  |  |  |  |  |  |  |  |  |  |  | C>T |  |

|  |  |  |  |  |  |  |  |  |  |  |  |  |  |  |  |  |  |  |
| --- | --- | --- | --- | --- | --- | --- | --- | --- | --- | --- | --- | --- | --- | --- | --- | --- | --- | --- |
| pos | 1892 | 2544 | 6445 | 15192 | 15720 | 18687 | 21304 | 21305 | 21987 | 22027 | 22029 | 22030 | 22031 | 22032 | 22033 | 22034 | 22073 |  |
| ref | G | C | C | A | C | C | C | G | G | T | A | G | T | T | C | A | G |  |
| AY.4.2 | A | A | T | G | T | T | A | A | A | T | - | - | - | - | - | - | A |  |
| AY.4.2 | A | A | T | G | T | T | A | A | A | T | A | A | A | G | T | G | A | #770237 |
| AY.4.2 | G | C | C | A | C | C | C | G | G | G | A | A | A | G | T | G | G |  |
| mut |  |  |  |  |  |  |  |  |  |  |  |  |  |  |  |  | G>A |  |

|  |  |  |  |  |  |  |  |  |  |  |  |  |  |  |  |  |  |  |
| --- | --- | --- | --- | --- | --- | --- | --- | --- | --- | --- | --- | --- | --- | --- | --- | --- | --- | --- |
| pos | 1892 | 2544 | 6445 | 15192 | 15720 | 18687 | 21304 | 21305 | 21987 | 22027 | 22029 | 22030 | 22031 | 22032 | 22033 | 22034 | 22073 |  |
| ref | G | C | C | A | C | C | C | G | G | T | A | G | T | T | C | A | G |  |
| AY.4.2 | A | A | T | G | T | T | A | A | A | T | - | - | - | - | - | - | A |  |
| AY.4.2 | A | A | T | G | T | T | A | A | A | G | A | A | A | G | T | G | A | #770328 |
| AY.4.2 | G | C | C | A | C | C | C | G | G | G | A | A | A | G | T | G | G |  |
| mut |  |  |  |  |  |  |  |  |  |  |  |  |  |  |  |  | G>A |  |

|  |  |  |  |  |  |  |  |  |  |  |  |  |  |  |  |  |  |  |
| --- | --- | --- | --- | --- | --- | --- | --- | --- | --- | --- | --- | --- | --- | --- | --- | --- | --- | --- |
| pos | 3523 | 5273 | 5668 | 6419 | 9700 | 13433 | 17442 | 17913 | 20404 | 22027 | 22029 | 22030 | 22031 | 22032 | 22033 | 22034 | 22264 |  |
| ref | A | G | G | G | C | A | T | C | C | T | A | G | T | T | C | A | C |  |
| AY.117 | A | A | G | G | T | G | C | C | C | T | - | - | - | - | - | - | C |  |
| AY.117 | A | A | G | G | T | G | C | C | T | G | A | A | A | G | T | G | T | #771717 |
| AY.117 | C | G | A | A | C | A | T | T | C | G | A | A | A | G | T | G | C |  |
| mut |  |  |  |  |  |  |  |  | C>T |  |  |  |  |  |  |  | C>T |  |

|  |  |  |  |  |  |  |  |  |  |  |  |  |  |  |  |  |
| --- | --- | --- | --- | --- | --- | --- | --- | --- | --- | --- | --- | --- | --- | --- | --- | --- |
| pos | 361 | 445 | 2306 | 3333 | 9653 | 18400 | 22027 | 22029 | 22030 | 22031 | 22032 | 22033 | 22034 | 27644 | 28756 |  |
| ref | A | T | C | T | G | C | T | A | G | T | T | C | A | C | T |  |
| AY.119 | A | C | T | C | G | C | T | - | - | - | - | - | - | T | T |  |
| AY.119 | A | C | T | C | G | C | G | A | A | A | G | T | G | C | A | #786042 |
| AY.119 | G | T | C | T | A | T | G | A | A | A | G | T | G | C | T |  |
| mut |  |  |  |  |  |  |  |  |  |  |  |  |  |  | T>A |  |

|  |  |  |  |  |  |  |  |  |  |  |  |  |  |  |  |  |  |  |  |  |
| --- | --- | --- | --- | --- | --- | --- | --- | --- | --- | --- | --- | --- | --- | --- | --- | --- | --- | --- | --- | --- |
| pos | 599 | 3092 | 3433 | 8976 | 10323 | 10423 | 17676 | 18162 | 20429 | 20736 | 22026 | 22027 | 22029 | 22030 | 22031 | 22032 | 22033 | 22034 | 26801 |  |
| ref | G | C | G | A | A | T | C | C | C | C | G | T | A | G | T | T | C | A | C |  |
| AY.39 | T | T | T | A | G | C | T | T | T | T | G | T | - | - | - | - | - | - | T |  |
| AY.39 | T | T | T | C | G | C | T | T | T | T | T | G | A | A | A | G | T | G | T | #786106 |
| AY.39 | G | C | G | A | A | T | C | C | C | C | T | G | A | A | A | G | T | G | C |  |
| mut |  |  |  | A>C |  |  |  |  |  |  |  |  |  |  |  |  |  |  | C>T |  |

|  |  |  |  |  |  |  |  |  |  |  |  |  |  |  |  |
| --- | --- | --- | --- | --- | --- | --- | --- | --- | --- | --- | --- | --- | --- | --- | --- |
| pos | 7858 | 22026 | 22027 | 22029 | 22030 | 22031 | 22032 | 22033 | 22034 | 22093 | 24784 | 28651 | 29254 | 29722 |  |
| ref | C | G | T | A | G | T | T | C | A | G | C | C | G | C |  |
| AY.113 | C | T | G | A | A | A | G | T | G | G | C | C | G | C |  |
| AY.113 | T | T | G | A | A | A | G | T | G | C | T | T | T | T | #786113 |
| AY.113 | T | G | T | - | - | - | - | - | - | C | T | T | T | T |  |
| mut | C>T |  |  |  |  |  |  |  |  |  |  |  |  |  |  |

|  |  |  |  |  |  |  |  |  |  |  |  |  |  |  |  |  |  |
| --- | --- | --- | --- | --- | --- | --- | --- | --- | --- | --- | --- | --- | --- | --- | --- | --- | --- |
| pos | 222 | 1519 | 4051 | 7104 | 8379 | 12801 | 17236 | 19164 | 22801 | 24208 | 25354 | 25934 | 26107 | 27507 | 29659 | 29825 |  |
| ref | C | C | T | C | A | G | A | C | G | C | G | A | G | A | C | G |  |
| AY.25 | T | T | C | C | G | A | A | C | T | C | T | T | C | C | C | G |  |
| AY.103 | T | T | C | C | G | G | G | T | G | T | G | A | G | A | T | T | #786190 |
| AY.103 | C | C | T | T | A | G | G | T | G | T | G | A | G | A | C | G |  |
| mut |  |  |  |  |  |  |  |  |  |  |  |  |  |  | C>T | G>T |  |

|  |  |  |  |  |  |  |  |  |  |  |  |  |  |  |  |  |  |
| --- | --- | --- | --- | --- | --- | --- | --- | --- | --- | --- | --- | --- | --- | --- | --- | --- | --- |
| pos | 526 | 3965 | 9165 | 10323 | 11750 | 12772 | 15240 | 19524 | 21846 | 22017 | 22335 | 27371 | 27875 | 27881 | 27882 | 27883 |  |
| ref | G | G | C | A | C | A | C | C | C | G | G | C | T | C | G | C |  |
| AY.113 | G | T | T | G | C | G | T | C | T | T | T | T | T | C | G | C |  |
| AY.113 | G | T | T | G | T | G | T | C | T | G | T | T | C | T | C | T | #790878 |
| AY.98 | T | G | C | A | C | A | C | T | C | G | G | C | C | T | C | T |  |
| mut |  |  |  |  | C>T |  |  |  |  | T>G |  |  |  |  |  |  |  |

|  |  |  |  |  |  |  |  |  |  |  |  |  |  |  |  |  |  |  |  |  |  |  |  |  |
| --- | --- | --- | --- | --- | --- | --- | --- | --- | --- | --- | --- | --- | --- | --- | --- | --- | --- | --- | --- | --- | --- | --- | --- | --- |
| pos | 2272 | 6040 | 6376 | 6638 | 7926 | 8062 | 8805 | 9985 | 14014 | 16726 | 20436 | 21846 | 22335 | 22813 | 25273 | 25549 | 26056 | 26607 | 27516 | 28073 | 29700 | 29750 | 29768 |  |
| ref | G | C | T | C | C | A | A | C | T | C | C | C | G | G | G | C | G | C | G | G | A | C | A |  |
| AY.44 | G | T | C | T | T | A | A | T | G | T | C | C | G | G | G | C | G | C | G | T | G | C | G |  |
| AY.44 | G | T | C | T | T | A | A | T | G | T | T | T | T | T | C | T | T | T | T | G | A | T | A | #794708 |
| AY.1 | C | C | T | C | C | G | G | C | T | C | T | T | T | T | C | T | T | T | T | G | A | T | A |  |
| mut |  |  |  |  |  |  |  |  |  |  |  |  |  |  |  |  |  |  |  |  |  |  |  |  |

|  |  |  |  |  |  |  |  |  |  |  |  |  |  |  |  |  |  |  |  |
| --- | --- | --- | --- | --- | --- | --- | --- | --- | --- | --- | --- | --- | --- | --- | --- | --- | --- | --- | --- |
| pos | 1191 | 5646 | 10369 | 15264 | 16092 | 17259 | 19816 | 20132 | 22027 | 22029 | 22030 | 22031 | 22032 | 22033 | 22034 | 25626 | 27143 | 29095 |  |
| ref | C | C | C | T | C | G | G | C | T | A | G | T | T | C | A | C | C | C |  |
| AY.4 | T | T | C | C | T | T | G | T | T | - | - | - | - | - | - | C | C | C |  |
| AY.4 | T | T | T | C | T | T | G | T | G | A | A | A | G | T | G | T | T | T | #797434 |
| AY.4 | C | C | C | T | C | G | T | C | G | A | A | A | G | T | G | C | T | C |  |
| mut |  |  | C>T |  |  |  |  |  |  |  |  |  |  |  |  | C>T |  | C>T |  |

|  |  |  |  |  |  |  |  |  |  |  |  |  |  |  |  |
| --- | --- | --- | --- | --- | --- | --- | --- | --- | --- | --- | --- | --- | --- | --- | --- |
| pos | 799 | 825 | 3210 | 7273 | 10741 | 21987 | 22027 | 22029 | 22030 | 22031 | 22032 | 22033 | 22034 | 29466 |  |
| ref | C | A | A | G | C | G | T | A | G | T | T | C | A | C |  |
| AY.4.2 | T | A | G | A | T | A | T | - | - | - | - | - | - | T |  |
| AY.4.2 | T | G | G | A | T | A | G | A | A | A | G | T | G | T | #797606 |
| AY.4.2 | C | A | A | G | C | G | G | A | A | A | G | T | G | C |  |
| mut |  | A>G |  |  |  |  |  |  |  |  |  |  |  | C>T |  |

|  |  |  |  |  |  |  |  |  |  |  |  |  |  |  |  |  |  |  |  |  |  |  |  |  |  |  |  |  |  |  |  |  |  |
| --- | --- | --- | --- | --- | --- | --- | --- | --- | --- | --- | --- | --- | --- | --- | --- | --- | --- | --- | --- | --- | --- | --- | --- | --- | --- | --- | --- | --- | --- | --- | --- | --- | --- |
| pos | 1616 | 1684 | 4181 | 5164 | 5184 | 6402 | 6616 | 7124 | 7851 | 8131 | 8986 | 9053 | 9891 | 10029 | 10448 | 11201 | 11332 | 11418 | 11514 | 12049 | 13019 | 14925 | 18647 | 19220 | 21846 | 22227 | 22427 | 23635 | 27071 | 27874 | 28326 | 28916 |  |
| ref | C | C | G | G | C | C | A | C | C | G | C | G | C | C | C | A | A | T | C | C | C | C | C | C | C | C | G | C | T | C | G | G |  |
| AY.4 | C | C | T | G | C | T | A | T | T | G | T | T | C | T | C | G | G | T | C | C | C | T | C | T | T | C | G | C | C | T | G | T |  |
| AY.47 | C | C | T | G | C | C | G | C | C | T | C | G | T | C | T | A | A | C | T | T | T | C | T | C | C | T | A | T | T | C | T | G | #803762 |
| AY.47 | A | T | G | T | T | C | G | C | C | T | C | G | T | C | T | A | A | C | T | T | T | C | T | C | C | T | A | T | T | C | T | G |  |
| mut |  |  |  |  |  |  |  |  |  |  |  |  |  |  |  |  |  |  |  |  |  |  |  |  |  |  |  |  |  |  |  |  |  |

|  |  |  |  |  |  |  |  |  |  |  |  |  |  |  |  |
| --- | --- | --- | --- | --- | --- | --- | --- | --- | --- | --- | --- | --- | --- | --- | --- |
| pos | 6990 | 21987 | 22026 | 22027 | 22029 | 22030 | 22031 | 22032 | 22033 | 22034 | 22054 | 27561 | 28808 | 28879 |  |
| ref | C | G | G | T | A | G | T | T | C | A | T | G | G | T |  |
| AY.39 | C | G | T | G | A | A | A | G | T | G | T | G | G | T |  |
| AY.39 | T | G | T | G | A | A | A | G | T | G | C | T | A | C | #808317 |
| AY.39 | T | A | G | T | - | - | - | - | - | - | C | T | A | C |  |
| mut | C>T |  |  |  |  |  |  |  |  |  |  |  |  |  |  |

|  |  |  |  |  |  |  |  |  |  |  |  |  |  |  |  |  |  |  |  |  |  |
| --- | --- | --- | --- | --- | --- | --- | --- | --- | --- | --- | --- | --- | --- | --- | --- | --- | --- | --- | --- | --- | --- |
| pos | 252 | 3433 | 4891 | 5826 | 7312 | 9805 | 10116 | 10423 | 10448 | 11461 | 19151 | 22026 | 22027 | 22029 | 22030 | 22031 | 22032 | 22033 | 22034 | 28000 |  |
| ref | G | G | C | C | G | G | C | T | C | C | C | G | T | A | G | T | T | C | A | C |  |
| AY.39 | T | T | T | T | T | T | C | C | T | C | T | G | T | - | - | - | - | - | - | C |  |
| AY.39 | T | T | T | T | T | T | T | C | T | T | T | T | G | A | A | A | G | T | G | T | #808674 |
| AY.39 | G | G | C | C | G | G | C | T | C | C | C | T | G | A | A | A | G | T | G | C |  |
| mut |  |  |  |  |  |  | C>T |  |  | C>T |  |  |  |  |  |  |  |  |  | C>T |  |

|  |  |  |  |  |  |  |  |  |  |  |  |  |  |  |  |  |  |  |  |  |  |  |  |  |  |  |  |  |  |  |  |
| --- | --- | --- | --- | --- | --- | --- | --- | --- | --- | --- | --- | --- | --- | --- | --- | --- | --- | --- | --- | --- | --- | --- | --- | --- | --- | --- | --- | --- | --- | --- | --- |
| pos | 1170 | 6816 | 7764 | 7851 | 8956 | 9684 | 11620 | 12073 | 13482 | 19086 | 19894 | 21304 | 21372 | 21538 | 25463 | 25676 | 26058 | 26176 | 26828 | 27604 | 28077 | 28237 | 28241 | 28243 | 28245 | 28247 | 28251 | 28252 | 28253 | 28254 |  |
| ref | C | G | C | C | C | T | C | C | A | G | G | C | G | G | C | T | C | C | G | G | G | G | T | T | T | A | T | T | C | A |  |
| AY.39 | C | G | T | C | T | T | T | T | G | T | A | T | T | A | C | C | T | C | T | A | G | G | T | T | T | A | - | - | - | A |  |
| AY.39 | C | G | T | C | T | T | T | T | G | T | A | T | T | A | T | C | T | C | T | A | G | G | A | A | C | G | - | T | T | C | #812396 |
| AY.4 | T | A | C | T | C | C | C | C | A | G | G | C | G | G | C | T | C | T | G | G | T | T | A | A | C | G | T | T | C | C |  |
| mut |  |  |  |  |  |  |  |  |  |  |  |  |  |  | C>T |  |  |  |  |  |  |  |  |  |  |  | T>- |  | C>T |  |  |

|  |  |  |  |  |  |  |  |  |  |  |  |  |  |  |  |  |  |  |
| --- | --- | --- | --- | --- | --- | --- | --- | --- | --- | --- | --- | --- | --- | --- | --- | --- | --- | --- |
| pos | 3243 | 9943 | 10030 | 12599 | 14925 | 17439 | 20384 | 20679 | 21364 | 22026 | 22027 | 22029 | 22030 | 22031 | 22032 | 22033 | 22034 |  |
| ref | G | C | C | A | C | C | C | G | C | G | T | A | G | T | T | C | A |  |
| AY.100 | T | T | T | T | T | T | T | T | T | G | T | - | - | - | - | - | - |  |
| AY.100 | T | T | T | T | T | T | T | T | T | G | G | A | A | A | G | T | G | #812783 |
| AY.100 | G | C | C | A | C | C | C | G | C | T | G | A | A | A | G | T | G |  |
| mut |  |  |  |  |  |  |  |  |  |  |  |  |  |  |  |  |  |  |

|  |  |  |  |  |  |  |  |  |  |  |  |  |  |  |  |  |  |  |
| --- | --- | --- | --- | --- | --- | --- | --- | --- | --- | --- | --- | --- | --- | --- | --- | --- | --- | --- |
| pos | 237 | 5995 | 7911 | 12756 | 14262 | 19476 | 21638 | 22027 | 22029 | 22030 | 22031 | 22032 | 22033 | 22034 | 28085 | 29254 | 29700 |  |
| ref | G | A | G | C | C | A | C | T | A | G | T | T | C | A | G | G | A |  |
| AY.119 | G | G | T | T | T | G | T | T | - | - | - | - | - | - | T | G | A |  |
| AY.119 | T | G | T | T | T | G | T | G | A | A | A | G | T | G | T | T | A | #812785 |
| AY.119 | G | A | G | C | C | A | C | G | A | A | A | G | T | G | G | G | G |  |
| mut | G>T |  |  |  |  |  |  |  |  |  |  |  |  |  | G>T | G>T | G>A |  |

|  |  |  |  |  |  |  |  |  |  |  |  |  |  |  |  |  |  |  |
| --- | --- | --- | --- | --- | --- | --- | --- | --- | --- | --- | --- | --- | --- | --- | --- | --- | --- | --- |
| pos | 173 | 526 | 8756 | 9895 | 10977 | 12073 | 15569 | 16242 | 17331 | 19524 | 20517 | 21071 | 25552 | 27875 | 27881 | 27882 | 27883 |  |
| ref | C | G | C | T | C | C | C | T | G | C | T | A | G | T | C | G | C |  |
| AY.46.5 | T | G | C | C | T | T | T | G | A | C | C | G | T | T | C | G | C |  |
| AY.46.5 | T | G | T | C | T | T | T | G | A | C | C | G | T | C | T | C | T | #813563 |
| AY.98 | C | T | C | T | C | C | C | T | G | T | T | A | G | C | T | C | T |  |
| mut |  |  | C>T |  |  |  |  |  |  |  |  |  |  |  |  |  |  |  |

|  |  |  |  |  |  |  |  |  |  |  |  |  |  |  |  |  |  |  |  |  |  |  |  |
| --- | --- | --- | --- | --- | --- | --- | --- | --- | --- | --- | --- | --- | --- | --- | --- | --- | --- | --- | --- | --- | --- | --- | --- |
| pos | 3478 | 6709 | 6763 | 7049 | 8245 | 8759 | 15192 | 15720 | 17775 | 17977 | 18687 | 20839 | 21304 | 21305 | 21995 | 22227 | 25614 | 26101 | 28146 | 28326 | 28638 | 29200 |  |
| ref | T | A | T | T | A | G | A | C | A | C | C | G | C | G | T | C | C | G | C | G | C | C |  |
| AY.4 | T | A | C | T | T | T | A | C | G | T | C | G | C | G | T | C | C | A | C | T | T | C |  |
| AY.4 | T | A | C | T | T | T | A | C | G | T | C | G | C | G | T | C | C | A | T | G | C | T | #814569 |
| AY.4.2 | C | C | T | C | A | G | G | T | A | C | T | A | A | A | C | T | T | G | T | G | C | T |  |
| mut |  |  |  |  |  |  |  |  |  |  |  |  |  |  |  |  |  |  |  |  |  |  |  |

|  |  |  |  |  |  |  |  |  |  |  |  |  |  |  |
| --- | --- | --- | --- | --- | --- | --- | --- | --- | --- | --- | --- | --- | --- | --- |
| pos | 186 | 3802 | 4034 | 7121 | 15656 | 17040 | 19524 | 22051 | 25473 | 25613 | 26014 | 27526 | 29108 |  |
| ref | C | T | A | A | C | T | C | G | T | C | A | C | C |  |
| AY.4 | C | A | G | A | C | T | C | C | T | C | A | C | C |  |
| AY.4 | C | A | G | A | T | C | T | G | C | T | G | T | T | #816313 |
| AY.4 | T | T | A | G | T | C | C | G | C | T | G | T | T |  |
| mut |  |  |  |  |  |  | C>T |  |  |  |  |  |  |  |

|  |  |  |  |  |  |  |  |  |  |  |  |  |  |  |  |  |  |  |  |
| --- | --- | --- | --- | --- | --- | --- | --- | --- | --- | --- | --- | --- | --- | --- | --- | --- | --- | --- | --- |
| pos | 526 | 3811 | 6113 | 7749 | 7851 | 10357 | 10969 | 15264 | 15912 | 16884 | 19524 | 21846 | 22225 | 25218 | 26732 | 27059 | 28199 | 29736 |  |
| ref | G | C | C | C | C | T | C | T | T | T | C | C | G | G | T | C | T | G |  |
| AY.98 | T | T | C | T | C | C | T | T | C | A | T | C | G | T | C | C | C | C |  |
| AY.98 | T | T | C | T | C | C | T | C | T | T | C | T | T | G | T | T | T | G | #816844 |
| AY.4 | G | C | T | C | T | T | C | C | T | T | C | T | T | G | T | T | T | G |  |
| mut |  |  |  |  |  |  |  |  |  |  |  |  |  |  |  |  |  |  |  |

|  |  |  |  |  |  |  |  |  |  |  |  |  |  |  |  |  |
| --- | --- | --- | --- | --- | --- | --- | --- | --- | --- | --- | --- | --- | --- | --- | --- | --- |
| pos | 449 | 526 | 535 | 1741 | 3155 | 7851 | 11595 | 15380 | 19524 | 21846 | 26270 | 27875 | 27881 | 27882 | 27883 |  |
| ref | C | G | A | A | G | C | A | G | C | C | C | T | C | G | C |  |
| AY.4 | T | G | G | G | T | T | G | G | C | T | C | T | C | G | C |  |
| AY.4 | T | G | G | G | T | T | G | T | C | T | T | C | T | C | T | #819037 |
| AY.98 | C | T | A | A | G | C | A | G | T | C | C | C | T | C | T |  |
| mut |  |  |  |  |  |  |  | G>T |  |  | C>T |  |  |  |  |  |

|  |  |  |  |  |  |  |  |  |  |  |  |  |  |  |  |  |  |  |  |  |  |  |
| --- | --- | --- | --- | --- | --- | --- | --- | --- | --- | --- | --- | --- | --- | --- | --- | --- | --- | --- | --- | --- | --- | --- |
| pos | 884 | 2147 | 4720 | 7851 | 15277 | 16914 | 17040 | 17403 | 17427 | 19677 | 20055 | 21839 | 22027 | 22029 | 22030 | 22031 | 22032 | 22033 | 22034 | 23785 | 25667 |  |
| ref | C | C | G | C | C | G | T | C | G | G | A | G | T | A | G | T | T | C | A | C | C |  |
| AY.4 | C | T | G | T | T | G | C | T | T | G | G | T | T | - | - | - | - | - | - | T | T |  |
| AY.4 | C | T | G | T | T | G | C | T | T | G | G | T | G | A | A | A | G | T | G | T | T | #819060 |
| AY.114 | T | C | T | C | C | T | T | C | G | T | A | G | G | A | A | A | G | T | G | C | C |  |
| mut |  |  |  |  |  |  |  |  |  |  |  |  |  |  |  |  |  |  |  | C>T | C>T |  |

|  |  |  |  |  |  |  |  |  |  |  |  |  |  |
| --- | --- | --- | --- | --- | --- | --- | --- | --- | --- | --- | --- | --- | --- |
| pos | 5218 | 7393 | 11563 | 21583 | 22027 | 22029 | 22030 | 22031 | 22032 | 22033 | 22034 | 24415 |  |
| ref | T | G | C | A | T | A | G | T | T | C | A | G |  |
| AY.117 | C | T | T | G | T | - | - | - | - | - | - | T |  |
| AY.117 | C | T | T | G | G | A | A | A | G | T | G | T | #820211 |
| AY.117 | T | G | C | A | G | A | A | A | G | T | G | G |  |
| mut |  |  |  |  |  |  |  |  |  |  |  | G>T |  |

|  |  |  |  |  |  |  |  |  |  |  |  |  |  |  |
| --- | --- | --- | --- | --- | --- | --- | --- | --- | --- | --- | --- | --- | --- | --- |
| pos | 203 | 526 | 934 | 3871 | 7851 | 19524 | 21600 | 21846 | 26455 | 27875 | 27881 | 27882 | 27883 |  |
| ref | C | G | C | G | C | C | G | C | C | T | C | G | C |  |
| AY.4 | T | G | T | T | T | C | C | T | T | T | C | G | C |  |
| AY.4 | T | G | T | T | T | C | C | T | T | C | T | C | T | #820694 |
| AY.98 | C | T | C | G | C | T | G | C | C | C | T | C | T |  |
| mut |  |  |  |  |  |  |  |  |  |  |  |  |  |  |

|  |  |  |  |  |  |  |  |  |  |  |  |  |  |  |  |
| --- | --- | --- | --- | --- | --- | --- | --- | --- | --- | --- | --- | --- | --- | --- | --- |
| pos | 526 | 4237 | 7851 | 9565 | 13210 | 17040 | 18348 | 19524 | 21846 | 23680 | 27875 | 27881 | 27882 | 27883 |  |
| ref | G | T | C | C | T | T | C | C | C | T | T | C | G | C |  |
| AY.4 | G | C | T | T | C | C | T | C | T | C | T | C | G | C |  |
| AY.4 | G | C | T | T | C | C | T | C | T | C | C | T | C | T | #822053 |
| AY.98 | T | T | C | C | T | T | C | T | C | T | C | T | C | T |  |
| mut |  |  |  |  |  |  |  |  |  |  |  |  |  |  |  |

|  |  |  |  |  |  |  |  |  |  |  |  |  |  |  |  |  |  |  |  |  |  |
| --- | --- | --- | --- | --- | --- | --- | --- | --- | --- | --- | --- | --- | --- | --- | --- | --- | --- | --- | --- | --- | --- |
| pos | 793 | 884 | 4720 | 7851 | 8867 | 15277 | 16750 | 16914 | 17040 | 17403 | 17427 | 19070 | 19677 | 22027 | 22029 | 22030 | 22031 | 22032 | 22033 | 22034 |  |
| ref | G | C | G | C | G | C | C | G | T | C | G | C | G | T | A | G | T | T | C | A |  |
| AY.4 | T | C | G | T | A | T | T | G | C | T | T | T | G | T | - | - | - | - | - | - |  |
| AY.4 | T | C | G | T | A | T | T | G | C | T | T | T | G | G | A | A | A | G | T | G | #822260 |
| AY.114 | G | T | T | C | G | C | C | T | T | C | G | C | T | G | A | A | A | G | T | G |  |
| mut |  |  |  |  |  |  |  |  |  |  |  |  |  |  |  |  |  |  |  |  |  |

|  |  |  |  |  |  |  |  |  |  |  |  |  |  |  |  |
| --- | --- | --- | --- | --- | --- | --- | --- | --- | --- | --- | --- | --- | --- | --- | --- |
| pos | 3329 | 5884 | 12459 | 13821 | 18264 | 21987 | 22027 | 22029 | 22030 | 22031 | 22032 | 22033 | 22034 | 23278 |  |
| ref | A | C | C | C | C | G | T | A | G | T | T | C | A | T |  |
| AY.4.2 | G | T | T | T | T | A | T | - | - | - | - | - | - | C |  |
| AY.4.2 | G | T | T | T | T | A | G | A | A | A | G | T | G | C | #822421 |
| AY.4.2 | A | C | C | C | C | G | G | A | A | A | G | T | G | T |  |
| mut |  |  |  |  |  |  |  |  |  |  |  |  |  | T>C |  |

|  |  |  |  |  |  |  |  |  |  |  |  |  |  |  |  |  |  |  |  |
| --- | --- | --- | --- | --- | --- | --- | --- | --- | --- | --- | --- | --- | --- | --- | --- | --- | --- | --- | --- |
| pos | 1588 | 2632 | 3476 | 6726 | 12357 | 13482 | 14637 | 15654 | 16224 | 19961 | 22027 | 22029 | 22030 | 22031 | 22032 | 22033 | 22034 | 25156 |  |
| ref | A | G | G | C | C | A | T | C | T | C | T | A | G | T | T | C | A | C |  |
| AY.39.1 | A | G | G | C | T | G | A | C | C | T | T | - | - | - | - | - | - | T |  |
| AY.39.1 | A | G | G | C | T | G | A | C | C | T | G | A | A | A | G | T | G | T | #823295 |
| AY.39 | G | T | A | T | C | A | T | T | T | C | G | A | A | A | G | T | G | C |  |
| mut |  |  |  |  |  |  |  |  |  |  |  |  |  |  |  |  |  | C>T |  |

|  |  |  |  |  |  |  |  |  |  |  |  |  |  |  |  |  |  |  |  |  |  |
| --- | --- | --- | --- | --- | --- | --- | --- | --- | --- | --- | --- | --- | --- | --- | --- | --- | --- | --- | --- | --- | --- |
| pos | 1059 | 4321 | 4331 | 10271 | 14359 | 15597 | 16221 | 16800 | 18583 | 20505 | 22026 | 22027 | 22029 | 22030 | 22031 | 22032 | 22033 | 22034 | 28582 | 29702 |  |
| ref | C | C | C | G | A | T | G | T | G | T | G | T | A | G | T | T | C | A | T | G |  |
| AY.118 | T | C | T | G | G | C | A | C | A | C | G | T | - | - | - | - | - | - | C | A |  |
| AY.118 | T | C | T | G | G | C | A | C | A | C | T | G | A | A | A | G | T | G | T | A | #823326 |
| AY.118 | C | T | C | A | A | T | G | T | G | T | T | G | A | A | A | G | T | G | T | G |  |
| mut |  |  |  |  |  |  |  |  |  |  |  |  |  |  |  |  |  |  |  | G>A |  |

|  |  |  |  |  |  |  |  |  |  |  |  |  |  |  |  |  |
| --- | --- | --- | --- | --- | --- | --- | --- | --- | --- | --- | --- | --- | --- | --- | --- | --- |
| pos | 507 | 508 | 509 | 5812 | 13850 | 16435 | 21595 | 22026 | 22027 | 22029 | 22030 | 22031 | 22032 | 22033 | 22034 |  |
| ref | A | T | G | C | G | G | C | G | T | A | G | T | T | C | A |  |
| AY.100 | T | C | A | T | T | T | T | G | T | - | - | - | - | - | - |  |
| AY.100 | T | C | A | T | T | T | T | G | G | A | A | A | G | T | G | #827288 |
| AY.100 | A | T | G | C | G | G | C | T | G | A | A | A | G | T | G |  |
| mut |  |  |  |  |  |  |  |  |  |  |  |  |  |  |  |  |

|  |  |  |  |  |  |  |  |  |  |  |  |  |  |  |  |  |  |  |  |  |  |  |  |  |  |  |  |  |  |
| --- | --- | --- | --- | --- | --- | --- | --- | --- | --- | --- | --- | --- | --- | --- | --- | --- | --- | --- | --- | --- | --- | --- | --- | --- | --- | --- | --- | --- | --- |
| pos | 3559 | 6040 | 6638 | 7926 | 11411 | 11740 | 12846 | 12946 | 13554 | 14014 | 15483 | 15654 | 16726 | 20003 | 20060 | 21372 | 21381 | 21396 | 21772 | 21786 | 21846 | 26227 | 26464 | 26645 | 27604 | 28073 | 29546 | 29700 |  |
| ref | A | C | C | C | A | A | C | T | C | T | A | C | C | A | G | G | C | C | C | G | C | G | C | C | G | G | C | A |  |
| AY.44 | A | T | T | T | G | A | C | C | T | G | G | C | T | G | G | G | C | C | T | T | C | G | T | C | G | T | T | G |  |
| AY.44 | A | T | T | T | G | A | T | C | T | G | G | C | T | G | G | G | C | T | T | T | C | T | C | T | A | G | C | A | #827500 |
| AY.39 | C | C | C | C | A | G | C | T | C | T | A | T | C | A | T | T | T | C | C | G | T | T | C | T | A | G | C | A |  |
| mut |  |  |  |  |  |  | C>T |  |  |  |  |  |  |  |  |  |  | C>T |  |  |  |  |  |  |  |  |  |  |  |

|  |  |  |  |  |  |  |  |  |  |  |  |  |  |  |  |
| --- | --- | --- | --- | --- | --- | --- | --- | --- | --- | --- | --- | --- | --- | --- | --- |
| pos | 2842 | 3264 | 4789 | 5544 | 10201 | 20726 | 22021 | 22027 | 22029 | 22030 | 22031 | 22032 | 22033 | 22034 |  |
| ref | C | C | T | C | G | A | G | T | A | G | T | T | C | A |  |
| AY.4 | T | T | T | C | T | A | T | T | - | - | - | - | - | - |  |
| AY.4 | T | T | T | C | T | A | T | G | A | A | A | G | T | G | #831889 |
| AY.4 | C | C | C | T | G | G | G | G | A | A | A | G | T | G |  |
| mut |  |  |  |  |  |  |  |  |  |  |  |  |  |  |  |

|  |  |  |  |  |  |  |  |  |  |  |  |  |  |  |
| --- | --- | --- | --- | --- | --- | --- | --- | --- | --- | --- | --- | --- | --- | --- |
| pos | 174 | 1106 | 5704 | 5944 | 12511 | 18395 | 22027 | 22029 | 22030 | 22031 | 22032 | 22033 | 22034 |  |
| ref | G | A | G | C | T | C | T | A | G | T | T | C | A |  |
| AY.20 | A | G | A | T | C | T | T | - | - | - | - | - | - |  |
| AY.20 | A | G | A | T | C | T | G | A | A | A | G | T | G | #833362 |
| AY.20 | G | A | G | C | T | C | G | A | A | A | G | T | G |  |
| mut |  |  |  |  |  |  |  |  |  |  |  |  |  |  |

|  |  |  |  |  |  |  |  |  |  |  |  |  |  |  |  |  |  |  |  |  |  |
| --- | --- | --- | --- | --- | --- | --- | --- | --- | --- | --- | --- | --- | --- | --- | --- | --- | --- | --- | --- | --- | --- |
| pos | 1274 | 3784 | 6421 | 6542 | 6661 | 8208 | 8994 | 11540 | 12473 | 14154 | 17058 | 19101 | 22027 | 22029 | 22030 | 22031 | 22032 | 22033 | 22034 | 25538 |  |
| ref | G | C | G | A | T | C | C | G | C | T | G | G | T | A | G | T | T | C | A | G |  |
| AY.103 | G | T | G | G | C | T | T | G | T | C | T | T | T | - | - | - | - | - | - | T |  |
| AY.103 | T | T | G | G | C | T | T | G | T | C | T | T | T | A | A | A | G | T | G | T | #834035 |
| AY.103 | G | C | T | A | T | C | C | T | C | T | G | G | G | A | A | A | G | T | G | G |  |
| mut | G>T |  |  |  |  |  |  |  |  |  |  |  |  |  |  |  |  |  |  | G>T |  |

|  |  |  |  |  |  |  |  |  |  |  |  |  |  |  |  |  |  |
| --- | --- | --- | --- | --- | --- | --- | --- | --- | --- | --- | --- | --- | --- | --- | --- | --- | --- |
| pos | 7851 | 10165 | 16425 | 17040 | 21846 | 21995 | 22227 | 24928 | 25614 | 25855 | 26172 | 26727 | 27143 | 27549 | 28689 | 29253 |  |
| ref | C | C | T | T | C | T | C | G | C | G | G | G | C | C | T | C |  |
| AY.4.2 | T | T | A | C | T | C | T | G | T | T | A | T | T | C | T | C |  |
| AY.4.2 | T | T | A | C | T | C | T | G | C | G | G | G | C | T | C | C | #835996 |
| AY.5 | C | C | T | T | C | T | C | T | C | G | G | G | C | T | T | T |  |
| mut |  |  |  |  |  |  |  |  |  |  |  |  |  |  | T>C | T>C |  |

|  |  |  |  |  |  |  |  |  |  |  |  |  |  |  |  |  |  |  |  |  |  |  |  |  |  |  |  |  |  |
| --- | --- | --- | --- | --- | --- | --- | --- | --- | --- | --- | --- | --- | --- | --- | --- | --- | --- | --- | --- | --- | --- | --- | --- | --- | --- | --- | --- | --- | --- |
| pos | 2273 | 5183 | 6031 | 7163 | 7851 | 9072 | 10448 | 11173 | 11743 | 12880 | 13627 | 14829 | 15952 | 16405 | 18525 | 18744 | 21707 | 21846 | 21898 | 23426 | 23673 | 24095 | 25452 | 27484 | 27641 | 27669 | 28188 | 28299 |  |
| ref | G | C | C | A | C | C | C | C | G | C | G | G | C | G | G | C | C | C | G | G | C | G | C | T | C | A | G | A |  |
| AY.43 | A | C | T | A | C | C | C | C | G | C | A | G | A | A | G | T | C | C | G | G | C | G | C | T | C | A | A | T |  |
| AY.43 | A | C | T | A | C | C | C | C | G | C | A | G | A | A | G | T | C | C | G | T | T | T | T | C | T | G | G | A | #838054 |
| AY.4 | G | T | C | G | T | T | T | T | T | T | G | T | C | G | T | C | T | T | A | T | T | T | T | C | T | G | G | A |  |
| mut |  |  |  |  |  |  |  |  |  |  |  |  |  |  |  |  |  |  |  |  |  |  |  |  |  |  |  |  |  |

|  |  |  |  |  |  |  |  |  |  |  |  |  |  |  |  |  |  |  |  |  |  |  |  |  |  |  |
| --- | --- | --- | --- | --- | --- | --- | --- | --- | --- | --- | --- | --- | --- | --- | --- | --- | --- | --- | --- | --- | --- | --- | --- | --- | --- | --- |
| pos | 1048 | 1168 | 2080 | 2229 | 4455 | 6040 | 7926 | 9750 | 11514 | 11866 | 12188 | 14014 | 19006 | 21123 | 25072 | 27014 | 27128 | 27526 | 27527 | 27689 | 28209 | 29050 | 29543 | 29579 | 29700 |  |
| ref | G | A | T | G | C | C | C | A | C | C | G | T | G | G | T | G | C | C | C | C | G | G | G | G | A |  |
| AY.44 | G | A | C | A | C | T | T | G | C | T | G | G | T | T | T | A | C | C | C | C | G | T | G | T | G |  |
| AY.44 | G | A | C | A | C | T | T | G | C | T | T | G | T | T | G | G | T | T | T | A | C | G | T | G | A | #839215 |
| AY.122 | T | G | T | G | T | C | C | A | T | C | G | T | G | G | G | G | T | T | T | A | C | G | T | G | A |  |
| mut |  |  |  |  |  |  |  |  |  |  | G>T |  |  |  |  |  |  |  |  |  |  |  |  |  |  |  |

|  |  |  |  |  |  |  |  |  |  |  |  |  |  |
| --- | --- | --- | --- | --- | --- | --- | --- | --- | --- | --- | --- | --- | --- |
| pos | 7732 | 8809 | 9339 | 18498 | 22026 | 22027 | 22029 | 22030 | 22031 | 22032 | 22033 | 22034 |  |
| ref | C | C | A | G | G | T | A | G | T | T | C | A |  |
| AY.100 | T | T | G | T | G | T | - | - | - | - | - | - |  |
| AY.100 | T | T | G | T | G | G | A | A | A | G | T | G | #843245 |
| AY.100 | C | C | A | G | T | G | A | A | A | G | T | G |  |
| mut |  |  |  |  |  |  |  |  |  |  |  |  |  |

|  |  |  |  |  |  |  |  |  |  |  |  |  |  |  |
| --- | --- | --- | --- | --- | --- | --- | --- | --- | --- | --- | --- | --- | --- | --- |
| pos | 374 | 2058 | 17541 | 18122 | 22026 | 22027 | 22029 | 22030 | 22031 | 22032 | 22033 | 22034 | 29543 |  |
| ref | G | T | C | G | G | T | A | G | T | T | C | A | G |  |
| AY.100 | A | G | T | T | G | T | - | - | - | - | - | - | T |  |
| AY.100 | A | G | T | T | G | G | A | A | A | G | T | G | G | #843289 |
| AY.100 | G | T | C | G | T | G | A | A | A | G | T | G | G |  |
| mut |  |  |  |  |  |  |  |  |  |  |  |  |  |  |

|  |  |  |  |  |  |  |  |  |  |  |  |  |  |  |  |  |  |  |  |  |  |
| --- | --- | --- | --- | --- | --- | --- | --- | --- | --- | --- | --- | --- | --- | --- | --- | --- | --- | --- | --- | --- | --- |
| pos | 1348 | 2202 | 3523 | 5668 | 6419 | 7787 | 7851 | 9622 | 14186 | 15810 | 17913 | 21974 | 22027 | 22029 | 22030 | 22031 | 22032 | 22033 | 22034 | 28817 |  |
| ref | C | G | A | G | G | G | C | T | G | C | C | G | T | A | G | T | T | C | A | G |  |
| AY.4 | T | G | A | G | G | A | T | C | G | C | C | C | T | - | - | - | - | - | - | G |  |
| AY.4 | T | G | A | G | G | A | T | C | G | C | C | C | G | A | A | A | G | T | G | T | #843295 |
| AY.117 | C | T | C | A | A | G | C | T | T | T | T | G | G | A | A | A | G | T | G | G |  |
| mut |  |  |  |  |  |  |  |  |  |  |  |  |  |  |  |  |  |  |  | G>T |  |

|  |  |  |  |  |  |  |  |  |  |  |  |  |  |  |  |  |  |
| --- | --- | --- | --- | --- | --- | --- | --- | --- | --- | --- | --- | --- | --- | --- | --- | --- | --- |
| pos | 1663 | 6042 | 11621 | 13011 | 16726 | 17010 | 17298 | 22027 | 22029 | 22030 | 22031 | 22032 | 22033 | 22034 | 22420 | 25677 |  |
| ref | C | A | T | C | C | C | G | T | A | G | T | T | C | A | A | G |  |
| AY.129 | T | A | T | T | C | C | T | T | - | - | - | - | - | - | G | T |  |
| AY.129 | T | A | T | T | C | C | T | G | A | A | A | G | T | G | G | T | #843367 |
| AY.129 | C | G | C | C | T | T | G | G | A | A | A | G | T | G | A | G |  |
| mut |  |  |  |  |  |  |  |  |  |  |  |  |  |  | A>G | G>T |  |

|  |  |  |  |  |  |  |  |  |  |  |  |  |  |  |  |  |  |  |  |  |  |  |  |  |  |  |  |
| --- | --- | --- | --- | --- | --- | --- | --- | --- | --- | --- | --- | --- | --- | --- | --- | --- | --- | --- | --- | --- | --- | --- | --- | --- | --- | --- | --- |
| pos | 5869 | 6040 | 6638 | 7926 | 8548 | 8752 | 12486 | 14014 | 14818 | 15540 | 16726 | 18337 | 18583 | 19684 | 20451 | 21817 | 21897 | 23929 | 26107 | 26759 | 27507 | 27691 | 28073 | 28810 | 29449 | 29700 |  |
| ref | C | C | C | C | G | C | T | T | C | C | C | G | G | G | C | A | C | C | G | A | A | A | G | C | G | A |  |
| AY.25 | C | C | C | C | G | T | T | T | T | T | C | T | A | T | T | A | T | C | C | G | C | G | G | A | T | A |  |
| B.1.617.2 | C | C | C | C | G | T | T | T | C | C | T | G | G | G | C | G | C | T | G | A | A | A | T | C | G | G | #843595 |
| AY.44 | T | T | T | T | T | C | C | G | C | C | T | G | G | G | C | G | C | T | G | A | A | A | T | C | G | G |  |
| mut |  |  |  |  |  |  |  |  |  |  |  |  |  |  |  |  |  |  |  |  |  |  |  |  |  |  |  |

|  |  |  |  |  |  |  |  |  |  |  |  |  |  |  |  |  |  |  |  |  |  |  |
| --- | --- | --- | --- | --- | --- | --- | --- | --- | --- | --- | --- | --- | --- | --- | --- | --- | --- | --- | --- | --- | --- | --- |
| pos | 400 | 5567 | 7851 | 10833 | 15952 | 16173 | 16377 | 17040 | 17122 | 18744 | 19645 | 21304 | 21595 | 21796 | 21846 | 22304 | 24130 | 27271 | 27506 | 28093 | 28299 |  |
| ref | T | G | C | C | C | C | G | T | G | C | G | C | C | G | C | T | C | G | G | C | A |  |
| AY.4 | C | A | T | T | C | T | G | C | G | C | T | C | T | G | T | C | C | G | G | C | A |  |
| AY.4 | C | A | T | T | A | T | G | C | T | C | G | C | C | T | C | T | C | T | T | T | T | #846450 |
| AY.43 | T | G | C | C | A | C | A | T | T | T | G | A | C | T | C | T | T | G | T | C | T |  |
| mut |  |  |  |  | C>A |  |  |  |  | T>C |  | A>C |  |  |  |  | T>C | G>T |  | C>T |  |  |

|  |  |  |  |  |  |  |  |  |  |  |  |  |  |  |
| --- | --- | --- | --- | --- | --- | --- | --- | --- | --- | --- | --- | --- | --- | --- |
| pos | 3737 | 7945 | 18488 | 19862 | 21727 | 22027 | 22029 | 22030 | 22031 | 22032 | 22033 | 22034 | 22484 |  |
| ref | C | C | T | C | C | T | A | G | T | T | C | A | G |  |
| AY.4 | T | T | C | C | T | T | - | - | - | - | - | - | A |  |
| AY.4 | T | T | C | C | T | G | A | A | A | G | T | G | A | #849427 |
| AY.4 | C | C | T | T | C | G | A | A | A | G | T | G | G |  |
| mut |  |  |  |  |  |  |  |  |  |  |  |  | G>A |  |

|  |  |  |  |  |  |  |  |  |  |  |  |  |  |
| --- | --- | --- | --- | --- | --- | --- | --- | --- | --- | --- | --- | --- | --- |
| pos | 6694 | 14817 | 18493 | 20629 | 22027 | 22029 | 22030 | 22031 | 22032 | 22033 | 22034 | 25916 |  |
| ref | G | T | C | C | T | A | G | T | T | C | A | C |  |
| AY.125 | T | C | T | T | T | - | - | - | - | - | - | T |  |
| AY.125 | T | C | T | T | G | A | A | A | G | T | G | T | #850984 |
| AY.125 | G | T | C | C | G | A | A | A | G | T | G | C |  |
| mut |  |  |  |  |  |  |  |  |  |  |  | C>T |  |

|  |  |  |  |  |  |  |  |  |  |  |  |  |  |  |  |  |  |  |  |  |  |  |  |  |  |  |  |  |  |  |  |
| --- | --- | --- | --- | --- | --- | --- | --- | --- | --- | --- | --- | --- | --- | --- | --- | --- | --- | --- | --- | --- | --- | --- | --- | --- | --- | --- | --- | --- | --- | --- | --- |
| pos | 443 | 1048 | 1170 | 2453 | 6816 | 7851 | 9684 | 10748 | 12075 | 14599 | 18816 | 20060 | 20594 | 21846 | 23401 | 26176 | 27214 | 27527 | 28054 | 28077 | 28237 | 28241 | 28243 | 28245 | 28247 | 28251 | 28252 | 28253 | 28254 | 29700 |  |
| ref | G | G | C | C | G | C | T | C | A | C | G | G | C | C | G | C | G | C | C | G | G | T | T | T | A | T | T | C | A | A |  |
| AY.122 | G | T | C | C | G | C | T | A | G | T | A | T | T | C | T | C | C | T | T | G | G | T | T | T | A | - | - | - | A | G |  |
| AY.122 | T | T | C | T | G | C | T | A | G | T | A | T | T | C | T | C | C | T | T | G | G | A | A | C | G | - | T | T | C | G | #852932 |
| AY.4 | G | G | T | C | A | T | C | C | A | C | G | G | C | T | G | T | G | C | C | T | T | A | A | C | G | T | T | C | C | A |  |
| mut | G>T |  |  | C>T |  |  |  |  |  |  |  |  |  |  |  |  |  |  |  |  |  |  |  |  |  | T>- |  | C>T |  | A>G |  |

|  |  |  |  |  |  |  |  |  |  |  |  |  |  |  |  |  |  |  |  |  |  |  |
| --- | --- | --- | --- | --- | --- | --- | --- | --- | --- | --- | --- | --- | --- | --- | --- | --- | --- | --- | --- | --- | --- | --- |
| pos | 490 | 884 | 4720 | 7851 | 8408 | 10523 | 15009 | 15960 | 16914 | 16917 | 19677 | 22027 | 22029 | 22030 | 22031 | 22032 | 22033 | 22034 | 23911 | 25793 | 29614 |  |
| ref | T | C | G | C | G | G | T | C | G | G | G | T | A | G | T | T | C | A | A | G | C |  |
| AY.4 | A | C | G | T | G | C | C | C | G | T | G | T | - | - | - | - | - | - | G | A | T |  |
| AY.4 | A | C | G | T | A | C | C | T | G | T | G | G | A | A | A | G | T | G | G | A | T | #853663 |
| AY.114 | T | T | T | C | G | G | T | C | T | G | T | G | A | A | A | G | T | G | A | G | C |  |
| mut |  |  |  |  | G>A |  |  | C>T |  |  |  |  |  |  |  |  |  |  | A>G | G>A | C>T |  |

|  |  |  |  |  |  |  |  |  |  |  |  |  |  |  |  |  |
| --- | --- | --- | --- | --- | --- | --- | --- | --- | --- | --- | --- | --- | --- | --- | --- | --- |
| pos | 4084 | 8950 | 9073 | 9246 | 11997 | 12220 | 14571 | 15792 | 22027 | 22029 | 22030 | 22031 | 22032 | 22033 | 22034 |  |
| ref | C | C | C | C | T | T | C | A | T | A | G | T | T | C | A |  |
| B.1.617.2 | T | T | T | T | C | T | C | T | T | - | - | - | - | - | - |  |
| B.1.617.2 | T | T | T | T | C | T | C | T | G | A | A | A | G | T | G | #854165 |
| B.1.617.2 | C | C | C | C | T | C | T | A | G | A | A | A | G | T | G |  |
| mut |  |  |  |  |  |  |  |  |  |  |  |  |  |  |  |  |

|  |  |  |  |  |  |  |  |  |  |  |  |  |  |  |  |  |  |  |
| --- | --- | --- | --- | --- | --- | --- | --- | --- | --- | --- | --- | --- | --- | --- | --- | --- | --- | --- |
| pos | 3646 | 5239 | 5812 | 8219 | 13045 | 15928 | 18828 | 22026 | 22027 | 22029 | 22030 | 22031 | 22032 | 22033 | 22034 | 27598 | 27722 |  |
| ref | C | C | C | G | G | C | C | G | T | A | G | T | T | C | A | G | T |  |
| AY.100 | T | T | T | A | A | T | C | G | T | - | - | - | - | - | - | G | T |  |
| AY.100 | T | T | T | A | A | T | T | G | G | A | A | A | G | T | G | T | C | #854189 |
| AY.100 | C | C | C | G | G | C | C | T | G | A | A | A | G | T | G | G | T |  |
| mut |  |  |  |  |  |  | C>T |  |  |  |  |  |  |  |  | G>T | T>C |  |

|  |  |  |  |  |  |  |  |  |  |  |  |  |  |  |  |  |  |  |  |  |  |  |  |  |  |  |  |  |  |  |  |  |  |  |  |  |  |  |  |  |  |  |  |  |  |  |  |  |  |
| --- | --- | --- | --- | --- | --- | --- | --- | --- | --- | --- | --- | --- | --- | --- | --- | --- | --- | --- | --- | --- | --- | --- | --- | --- | --- | --- | --- | --- | --- | --- | --- | --- | --- | --- | --- | --- | --- | --- | --- | --- | --- | --- | --- | --- | --- | --- | --- | --- | --- |
| pos | 193 | 222 | 1191 | 1267 | 1942 | 3122 | 4181 | 4510 | 5184 | 5339 | 6040 | 6402 | 6638 | 7124 | 7926 | 8986 | 9053 | 9535 | 9891 | 10029 | 11201 | 11332 | 11418 | 12946 | 14014 | 14187 | 16726 | 16741 | 17907 | 18176 | 18417 | 19160 | 19220 | 20262 | 21205 | 21910 | 25216 | 25571 | 25641 | 25806 | 26428 | 27739 | 27874 | 27910 | 28073 | 28916 | 29464 | 29700 |  |
| ref | C | C | C | C | G | G | G | G | C | C | C | C | C | C | C | C | G | T | C | C | A | A | T | T | T | G | C | G | T | C | T | C | C | A | C | C | A | C | G | A | G | C | C | T | G | G | T | A |  |
| AY.54 | T | C | T | T | T | T | G | G | T | T | C | C | C | C | C | C | G | C | T | C | A | A | C | C | T | A | C | T | C | T | T | T | C | G | C | C | A | C | G | G | G | T | C | A | G | G | T | A |  |
| AY.44 | T | C | T | T | T | T | T | T | C | C | T | T | T | T | T | T | T | T | C | T | G | G | T | T | G | G | T | G | T | C | C | C | T | A | T | T | G | T | T | A | T | C | T | T | T | T | C | G | #854204 |
| AY.44 | C | T | C | C | G | G | T | T | C | C | T | T | T | T | T | T | T | T | C | T | G | G | T | T | G | G | T | G | T | C | T | C | T | A | C | T | G | T | T | A | T | C | T | T | T | T | T | G |  |
| mut |  |  |  |  |  |  |  |  |  |  |  |  |  |  |  |  |  |  |  |  |  |  |  |  |  |  |  |  |  |  | T>C |  |  |  | C>T |  |  |  |  |  |  |  |  |  |  |  | T>C |  |  |

|  |  |  |  |  |  |  |  |  |  |  |  |  |  |  |  |  |  |  |  |  |
| --- | --- | --- | --- | --- | --- | --- | --- | --- | --- | --- | --- | --- | --- | --- | --- | --- | --- | --- | --- | --- |
| pos | 2277 | 2497 | 3395 | 10416 | 12115 | 15857 | 19011 | 19578 | 20304 | 22000 | 22027 | 22029 | 22030 | 22031 | 22032 | 22033 | 22034 | 25855 | 28849 |  |
| ref | T | A | G | C | C | C | C | A | A | C | T | A | G | T | T | C | A | G | C |  |
| AY.20 | C | G | T | T | T | T | T | G | G | T | T | - | - | - | - | - | - | T | T |  |
| AY.20 | C | G | T | T | T | T | T | G | G | T | G | A | A | A | G | T | G | T | T | #854278 |
| AY.20 | T | A | G | C | C | C | C | A | A | C | G | A | A | A | G | T | G | G | C |  |
| mut |  |  |  |  |  |  |  |  |  |  |  |  |  |  |  |  |  | G>T | C>T |  |

|  |  |  |  |  |  |  |  |  |  |  |  |  |
| --- | --- | --- | --- | --- | --- | --- | --- | --- | --- | --- | --- | --- |
| pos | 2639 | 4084 | 6449 | 8752 | 22027 | 22029 | 22030 | 22031 | 22032 | 22033 | 22034 |  |
| ref | G | C | C | C | T | A | G | T | T | C | A |  |
| AY.4 | A | T | T | T | T | - | - | - | - | - | - |  |
| AY.4 | A | T | T | T | G | A | A | A | G | T | G | #854285 |
| AY.4 | G | C | C | C | G | A | A | A | G | T | G |  |
| mut |  |  |  |  |  |  |  |  |  |  |  |  |

|  |  |  |  |  |  |  |  |  |  |  |  |  |  |  |  |  |  |  |  |  |  |  |  |
| --- | --- | --- | --- | --- | --- | --- | --- | --- | --- | --- | --- | --- | --- | --- | --- | --- | --- | --- | --- | --- | --- | --- | --- |
| pos | 174 | 2277 | 2497 | 3395 | 10270 | 10416 | 12115 | 15857 | 19011 | 19578 | 20304 | 22000 | 22026 | 22027 | 22029 | 22030 | 22031 | 22032 | 22033 | 22034 | 25855 | 28849 |  |
| ref | G | T | A | G | T | C | C | C | C | A | A | C | G | T | A | G | T | T | C | A | G | C |  |
| AY.20 | G | C | G | T | T | T | T | T | T | G | G | T | G | T | - | - | - | - | - | - | T | T |  |
| AY.20 | G | C | G | T | T | T | T | T | T | G | G | T | T | G | A | A | A | G | T | G | T | T | #854308 |
| AY.20 | A | T | A | G | C | C | C | C | C | A | A | C | T | G | A | A | A | G | T | G | G | C |  |
| mut |  |  |  |  |  |  |  |  |  |  |  |  |  |  |  |  |  |  |  |  | G>T | C>T |  |

|  |  |  |  |  |  |  |  |  |  |  |  |  |  |  |
| --- | --- | --- | --- | --- | --- | --- | --- | --- | --- | --- | --- | --- | --- | --- |
| pos | 6286 | 6539 | 11109 | 21642 | 22026 | 22027 | 22029 | 22030 | 22031 | 22032 | 22033 | 22034 | 23533 |  |
| ref | C | C | C | C | G | T | A | G | T | T | C | A | C |  |
| AY.39 | T | T | C | T | G | T | - | - | - | - | - | - | T |  |
| AY.39 | T | T | C | T | G | G | A | A | A | G | T | G | T | #857758 |
| AY.39 | C | C | T | C | T | G | A | A | A | G | T | G | C |  |
| mut |  |  |  |  |  |  |  |  |  |  |  |  | C>T |  |

|  |  |  |  |  |  |  |  |  |  |  |  |  |  |  |  |  |  |  |
| --- | --- | --- | --- | --- | --- | --- | --- | --- | --- | --- | --- | --- | --- | --- | --- | --- | --- | --- |
| pos | 934 | 7083 | 7452 | 13482 | 14592 | 16687 | 21261 | 22026 | 22027 | 22029 | 22030 | 22031 | 22032 | 22033 | 22034 | 28774 | 29321 |  |
| ref | C | C | G | A | T | A | A | G | T | A | G | T | T | C | A | G | G |  |
| AY.39.1 | T | T | G | G | A | G | G | G | T | - | - | - | - | - | - | G | G |  |
| AY.39.1 | T | T | T | G | A | G | G | T | G | A | A | A | G | T | G | T | T | #857833 |
| AY.39 | C | C | G | A | T | A | A | T | G | A | A | A | G | T | G | G | G |  |
| mut |  |  | G>T |  |  |  |  |  |  |  |  |  |  |  |  | G>T | G>T |  |

|  |  |  |  |  |  |  |  |  |  |  |  |  |  |  |  |
| --- | --- | --- | --- | --- | --- | --- | --- | --- | --- | --- | --- | --- | --- | --- | --- |
| pos | 491 | 823 | 2845 | 6421 | 11540 | 22027 | 22029 | 22030 | 22031 | 22032 | 22033 | 22034 | 22324 | 28909 |  |
| ref | G | C | T | G | G | T | A | G | T | T | C | A | T | C |  |
| AY.103 | T | T | C | G | G | T | - | - | - | - | - | - | C | T |  |
| AY.103 | T | T | C | G | G | G | A | A | A | G | T | G | C | T | #862824 |
| AY.103 | G | C | T | T | T | G | A | A | A | G | T | G | T | C |  |
| mut |  |  |  |  |  |  |  |  |  |  |  |  | T>C | C>T |  |

|  |  |  |  |  |  |  |  |  |  |  |  |  |  |  |  |  |  |  |  |
| --- | --- | --- | --- | --- | --- | --- | --- | --- | --- | --- | --- | --- | --- | --- | --- | --- | --- | --- | --- |
| pos | 884 | 4720 | 5746 | 7851 | 13713 | 16914 | 17310 | 17331 | 19677 | 22027 | 22029 | 22030 | 22031 | 22032 | 22033 | 22034 | 22880 | 29686 |  |
| ref | C | G | C | C | G | G | T | G | G | T | A | G | T | T | C | A | A | C |  |
| AY.4 | C | G | T | T | T | G | C | T | G | T | - | - | - | - | - | - | T | G |  |
| AY.4 | C | G | T | T | T | G | C | T | G | G | A | A | A | G | T | G | T | G | #863560 |
| AY.114 | T | T | C | C | G | T | T | G | T | G | A | A | A | G | T | G | A | C |  |
| mut |  |  |  |  |  |  |  |  |  |  |  |  |  |  |  |  | A>T | C>G |  |

|  |  |  |  |  |  |  |  |  |  |  |  |  |  |  |  |  |  |  |  |
| --- | --- | --- | --- | --- | --- | --- | --- | --- | --- | --- | --- | --- | --- | --- | --- | --- | --- | --- | --- |
| pos | 2527 | 3695 | 4534 | 6042 | 6554 | 11621 | 17010 | 17104 | 21987 | 22027 | 22029 | 22030 | 22031 | 22032 | 22033 | 22034 | 26091 | 27753 |  |
| ref | G | C | C | A | G | T | C | C | G | T | A | G | T | T | C | A | C | A |  |
| AY.129 | T | T | T | A | A | T | C | T | G | T | - | - | - | - | - | - | T | G |  |
| AY.129 | T | T | T | A | A | T | C | T | G | G | A | A | A | G | T | G | T | G | #863581 |
| AY.129 | G | C | C | G | G | C | T | C | A | G | A | A | A | G | T | G | C | A |  |
| mut |  |  |  |  |  |  |  |  |  |  |  |  |  |  |  |  | C>T | A>G |  |

|  |  |  |  |  |  |  |  |  |  |  |  |  |  |  |  |  |  |  |  |  |
| --- | --- | --- | --- | --- | --- | --- | --- | --- | --- | --- | --- | --- | --- | --- | --- | --- | --- | --- | --- | --- |
| pos | 6040 | 6408 | 7600 | 7926 | 9860 | 10159 | 11095 | 12885 | 14014 | 15181 | 15738 | 18501 | 21846 | 25003 | 25452 | 27595 | 28637 | 29627 | 29700 |  |
| ref | C | C | C | C | T | A | C | C | T | G | C | C | C | A | C | C | C | C | A |  |
| AY.44 | T | C | C | T | T | A | C | C | G | T | C | T | C | A | C | C | C | C | G |  |
| AY.44 | T | C | C | T | T | G | C | C | G | T | T | C | T | G | T | T | T | T | A | #867631 |
| AY.120.1 | C | T | T | C | C | A | T | T | T | G | C | C | T | G | T | T | C | T | A |  |
| mut |  |  |  |  |  | A>G |  |  |  |  | C>T |  |  |  |  |  | C>T |  |  |  |

|  |  |  |  |  |  |  |  |  |  |  |  |  |  |  |  |  |  |
| --- | --- | --- | --- | --- | --- | --- | --- | --- | --- | --- | --- | --- | --- | --- | --- | --- | --- |
| pos | 2623 | 10271 | 13170 | 15026 | 15720 | 17976 | 22027 | 22029 | 22030 | 22031 | 22032 | 22033 | 22034 | 27389 | 27988 | 29374 |  |
| ref | C | G | C | C | C | C | T | A | G | T | T | C | A | C | T | G |  |
| AY.118 | T | G | A | T | T | T | T | - | - | - | - | - | - | C | T | T |  |
| AY.118 | T | G | A | T | T | T | G | A | A | A | G | T | G | T | C | T | #867782 |
| AY.118 | C | A | C | C | C | C | G | A | A | A | G | T | G | C | T | G |  |
| mut |  |  |  |  |  |  |  |  |  |  |  |  |  | C>T | T>C | G>T |  |

|  |  |  |  |  |  |  |  |  |  |  |  |  |  |  |  |  |  |  |  |  |  |  |  |  |  |
| --- | --- | --- | --- | --- | --- | --- | --- | --- | --- | --- | --- | --- | --- | --- | --- | --- | --- | --- | --- | --- | --- | --- | --- | --- | --- |
| pos | 2706 | 3876 | 5203 | 5365 | 7851 | 8829 | 9451 | 10138 | 12756 | 15804 | 17040 | 17130 | 18927 | 19006 | 20374 | 22027 | 22029 | 22030 | 22031 | 22032 | 22033 | 22034 | 22899 | 23987 |  |
| ref | C | C | C | C | C | C | C | C | C | T | T | C | T | G | A | T | A | G | T | T | C | A | G | C |  |
| AY.119 | C | T | T | C | C | T | T | T | T | C | T | T | T | T | A | T | - | - | - | - | - | - | G | T |  |
| AY.119 | C | T | T | C | C | T | T | T | T | C | T | T | T | T | A | G | A | A | A | G | T | G | T | T | #867878 |
| AY.4.1 | T | C | C | T | T | C | C | C | C | T | C | C | C | G | C | G | A | A | A | G | T | G | T | C |  |
| mut |  |  |  |  |  |  |  |  |  |  |  |  |  |  |  |  |  |  |  |  |  |  |  | C>T |  |

|  |  |  |  |  |  |  |  |  |  |  |  |  |  |  |  |  |
| --- | --- | --- | --- | --- | --- | --- | --- | --- | --- | --- | --- | --- | --- | --- | --- | --- |
| pos | 1820 | 17822 | 18176 | 21974 | 22027 | 22029 | 22030 | 22031 | 22032 | 22033 | 22034 | 22502 | 24370 | 26464 | 26885 |  |
| ref | G | C | C | G | T | A | G | T | T | C | A | C | C | C | C |  |
| AY.4 | G | C | C | G | G | A | A | A | G | T | G | C | C | C | C |  |
| AY.4 | A | T | T | G | G | A | A | A | G | T | G | A | T | T | T | #871262 |
| AY.4 | A | T | C | C | T | - | - | - | - | - | - | A | T | T | T |  |
| mut | G>A | C>T | C>T |  |  |  |  |  |  |  |  |  |  |  |  |  |

|  |  |  |  |  |  |  |  |  |  |  |  |  |  |  |  |  |  |  |  |  |  |  |  |  |
| --- | --- | --- | --- | --- | --- | --- | --- | --- | --- | --- | --- | --- | --- | --- | --- | --- | --- | --- | --- | --- | --- | --- | --- | --- |
| pos | 5730 | 22029 | 22030 | 22031 | 22032 | 22033 | 22034 | 22194 | 22195 | 22196 | 22197 | 22198 | 22202 | 22204 | 22673 | 22674 | 22679 | 22686 | 22882 | 22898 | 23599 | 23854 | 26709 |  |
| ref | C | A | G | T | T | C | A | A | T | T | T | A | C | T | T | C | T | C | T | G | T | C | G |  |
| BA.1 | C | A | G | T | T | C | A | - | - | - | T | A | C | T | C | T | C | T | T | G | T | C | G |  |
| BA.1 | C | - | - | - | - | - | - | A | T | T | T | A | C | T | T | C | T | C | G | A | G | A | A | #871356 |
| BA.1.18 | T | - | - | - | - | - | - | A | G | T | G | C | A | C | T | C | T | C | G | A | G | A | A |  |
| mut |  | A>- | G>- | T>- | T>- | C>- | A>- | ->A | ->T | ->T |  |  |  |  |  |  |  |  |  |  |  |  |  |  |

|  |  |  |  |  |  |  |  |  |  |  |  |  |  |  |  |  |  |  |  |  |  |
| --- | --- | --- | --- | --- | --- | --- | --- | --- | --- | --- | --- | --- | --- | --- | --- | --- | --- | --- | --- | --- | --- |
| pos | 242 | 526 | 4827 | 7851 | 8756 | 11651 | 12310 | 12439 | 17502 | 17944 | 19524 | 21846 | 22329 | 23948 | 24062 | 27875 | 27881 | 27882 | 27883 | 29543 |  |
| ref | G | G | C | C | C | C | G | C | C | G | C | C | C | G | A | T | C | G | C | G |  |
| AY.4 | A | G | T | T | T | T | A | T | T | T | C | T | T | T | A | T | C | G | C | T |  |
| AY.4 | A | G | T | T | T | T | A | T | T | T | C | T | T | T | G | C | T | C | T | T | #874316 |
| AY.98 | G | T | C | C | C | C | G | C | C | G | T | C | C | G | A | C | T | C | T | G |  |
| mut |  |  |  |  |  |  |  |  |  |  |  |  |  |  | A>G |  |  |  |  | G>T |  |

|  |  |  |  |  |  |  |  |  |  |  |  |  |  |  |  |  |  |  |  |  |  |  |  |
| --- | --- | --- | --- | --- | --- | --- | --- | --- | --- | --- | --- | --- | --- | --- | --- | --- | --- | --- | --- | --- | --- | --- | --- |
| pos | 4255 | 6309 | 7851 | 8829 | 9485 | 10317 | 16075 | 17012 | 18040 | 19006 | 19185 | 21622 | 22027 | 22029 | 22030 | 22031 | 22032 | 22033 | 22034 | 22987 | 27610 | 29026 |  |
| ref | G | G | C | C | T | A | G | C | G | G | C | C | T | A | G | T | T | C | A | C | C | T |  |
| AY.4 | A | G | T | C | T | G | A | T | T | G | T | T | T | - | - | - | - | - | - | C | T | C |  |
| AY.4 | A | G | T | C | T | G | A | T | T | G | T | T | G | A | A | A | G | T | G | T | T | C | #874961 |
| AY.119 | G | A | C | T | C | A | G | C | G | T | C | C | G | A | A | A | G | T | G | T | C | T |  |
| mut |  |  |  |  |  |  |  |  |  |  |  |  |  |  |  |  |  |  |  |  | C>T | T>C |  |

|  |  |  |  |  |  |  |  |  |  |  |  |  |  |  |  |  |  |  |  |
| --- | --- | --- | --- | --- | --- | --- | --- | --- | --- | --- | --- | --- | --- | --- | --- | --- | --- | --- | --- |
| pos | 884 | 4720 | 5991 | 7851 | 10015 | 13621 | 16914 | 19677 | 19718 | 21264 | 22027 | 22029 | 22030 | 22031 | 22032 | 22033 | 22034 | 29100 |  |
| ref | C | G | A | C | C | G | G | G | C | T | T | A | G | T | T | C | A | G |  |
| AY.4 | C | G | C | T | C | A | G | G | T | G | T | - | - | - | - | - | - | A |  |
| AY.4 | C | G | C | T | T | A | G | G | T | G | G | A | A | A | G | T | G | A | #875059 |
| AY.114 | T | T | A | C | C | G | T | T | C | T | G | A | A | A | G | T | G | G |  |
| mut |  |  |  |  | C>T |  |  |  |  |  |  |  |  |  |  |  |  | G>A |  |

|  |  |  |  |  |  |  |  |  |  |  |  |  |  |  |  |  |  |  |  |  |  |  |  |  |  |  |  |  |  |  |  |  |  |  |  |  |  |  |  |  |  |  |  |  |
| --- | --- | --- | --- | --- | --- | --- | --- | --- | --- | --- | --- | --- | --- | --- | --- | --- | --- | --- | --- | --- | --- | --- | --- | --- | --- | --- | --- | --- | --- | --- | --- | --- | --- | --- | --- | --- | --- | --- | --- | --- | --- | --- | --- | --- |
| pos | 1520 | 3732 | 4181 | 5184 | 5584 | 6040 | 6402 | 6638 | 7124 | 7393 | 7926 | 8131 | 8653 | 8986 | 9053 | 9274 | 9891 | 10029 | 11201 | 11332 | 11418 | 11514 | 13019 | 14014 | 16726 | 19220 | 20115 | 21058 | 21156 | 21622 | 22227 | 23997 | 26062 | 26607 | 27112 | 27382 | 27516 | 27874 | 27879 | 28073 | 28916 | 29700 | 29738 |  |
| ref | A | C | G | C | A | C | C | C | C | G | C | G | G | C | G | A | C | C | A | A | T | C | C | T | C | C | C | C | C | C | C | C | G | C | G | G | G | C | C | G | G | A | C |  |
| AY.75 | A | A | G | T | G | C | C | C | C | T | C | G | G | C | G | A | T | C | A | A | C | T | T | T | C | C | T | C | T | T | T | C | T | C | A | G | A | C | T | G | G | A | A |  |
| B.1.617.2 | A | A | G | T | G | C | C | C | C | T | C | G | G | C | G | A | C | T | G | G | T | C | C | G | T | T | C | T | C | C | C | T | G | T | G | T | G | T | C | T | T | G | C | #875387 |
| AY.44 | G | C | T | C | A | T | T | T | T | G | T | T | T | T | T | G | C | T | G | G | T | C | C | G | T | T | C | T | C | C | C | T | G | T | G | T | G | T | C | T | T | G | C |  |
| mut |  |  |  |  |  |  |  |  |  |  |  |  |  |  |  |  |  |  |  |  |  |  |  |  |  |  |  |  |  |  |  |  |  |  |  |  |  |  |  |  |  |  |  |  |

|  |  |  |  |  |  |  |  |  |  |  |  |  |  |  |  |  |  |  |  |  |  |  |  |  |  |
| --- | --- | --- | --- | --- | --- | --- | --- | --- | --- | --- | --- | --- | --- | --- | --- | --- | --- | --- | --- | --- | --- | --- | --- | --- | --- |
| pos | 284 | 475 | 2806 | 3148 | 6040 | 7086 | 7926 | 8867 | 9180 | 11456 | 13712 | 14014 | 17866 | 18315 | 20389 | 21987 | 22447 | 23284 | 25183 | 25339 | 25690 | 29050 | 29509 | 29700 |  |
| ref | G | C | A | A | C | C | C | G | C | A | A | T | G | G | C | G | T | T | G | C | G | G | C | A |  |
| AY.44 | G | T | G | G | T | C | T | G | T | A | G | G | A | A | C | G | T | T | T | C | T | G | C | G |  |
| AY.3 | G | T | G | G | C | T | C | A | C | G | A | T | G | G | T | A | C | C | G | T | G | A | T | A | #876195 |
| AY.3 | A | C | A | A | C | T | C | G | C | G | A | T | G | G | T | A | C | C | G | T | G | A | T | A |  |
| mut |  |  |  |  |  |  |  | G>A |  |  |  |  |  |  |  |  |  |  |  |  |  |  |  |  |  |

|  |  |  |  |  |  |  |  |  |  |  |  |  |  |  |  |  |
| --- | --- | --- | --- | --- | --- | --- | --- | --- | --- | --- | --- | --- | --- | --- | --- | --- |
| pos | 526 | 3535 | 6040 | 7926 | 14014 | 14838 | 15002 | 16575 | 18804 | 19524 | 27875 | 27881 | 27882 | 27883 | 29700 |  |
| ref | G | C | C | C | T | C | A | C | C | C | T | C | G | C | A |  |
| AY.44 | G | C | T | T | G | C | G | T | T | C | T | C | G | C | G |  |
| AY.44 | G | T | T | T | G | T | G | T | T | C | C | T | C | T | G | #876220 |
| AY.98 | T | C | C | C | T | C | A | C | C | T | C | T | C | T | A |  |
| mut |  | C>T |  |  |  | C>T |  |  |  |  |  |  |  |  | A>G |  |

|  |  |  |  |  |  |  |  |  |  |  |  |  |  |  |  |  |  |  |  |  |  |  |  |  |
| --- | --- | --- | --- | --- | --- | --- | --- | --- | --- | --- | --- | --- | --- | --- | --- | --- | --- | --- | --- | --- | --- | --- | --- | --- |
| pos | 1170 | 6816 | 9684 | 13821 | 17040 | 18264 | 20208 | 21987 | 21995 | 22227 | 23278 | 25614 | 26176 | 28077 | 28237 | 28241 | 28243 | 28245 | 28247 | 28251 | 28252 | 28253 | 28254 |  |
| ref | C | G | T | C | T | C | G | G | T | C | T | C | C | G | G | T | T | T | A | T | T | C | A |  |
| AY.4.2 | C | G | T | T | C | T | T | G | C | T | C | T | C | G | G | T | T | T | A | - | - | - | A |  |
| AY.4.2 | C | G | T | T | C | T | T | A | C | T | C | T | C | G | G | A | A | C | G | - | T | T | C | #876792 |
| AY.4 | T | A | C | C | T | C | G | A | T | C | T | C | T | T | T | A | A | C | G | T | T | C | C |  |
| mut |  |  |  |  |  |  |  | G>A |  |  |  |  |  |  |  |  |  |  |  | T>- |  | C>T |  |  |

|  |  |  |  |  |  |  |  |  |  |  |  |  |  |  |  |  |
| --- | --- | --- | --- | --- | --- | --- | --- | --- | --- | --- | --- | --- | --- | --- | --- | --- |
| pos | 3429 | 5301 | 5988 | 9587 | 13965 | 14398 | 19870 | 22027 | 22029 | 22030 | 22031 | 22032 | 22033 | 22034 | 28141 |  |
| ref | C | C | C | A | C | G | G | T | A | G | T | T | C | A | G |  |
| AY.42 | T | T | C | G | T | T | C | T | - | - | - | - | - | - | T |  |
| AY.42 | T | T | T | G | T | T | C | G | A | A | A | G | T | G | T | #881025 |
| AY.42 | C | C | C | A | C | G | G | G | A | A | A | G | T | G | G |  |
| mut |  |  | C>T |  |  |  |  |  |  |  |  |  |  |  | G>T |  |

|  |  |  |  |  |  |  |  |  |  |  |  |  |  |  |  |  |  |  |  |  |  |  |  |
| --- | --- | --- | --- | --- | --- | --- | --- | --- | --- | --- | --- | --- | --- | --- | --- | --- | --- | --- | --- | --- | --- | --- | --- |
| pos | 601 | 1385 | 2568 | 6539 | 10186 | 13424 | 13854 | 14618 | 14919 | 17040 | 17193 | 20887 | 22026 | 22027 | 22029 | 22030 | 22031 | 22032 | 22033 | 22034 | 22595 | 29836 |  |
| ref | C | C | A | C | C | C | T | C | C | T | G | G | G | T | A | G | T | T | C | A | A | C |  |
| AY.4 | T | T | A | T | T | C | C | T | C | C | T | G | G | T | - | - | - | - | - | - | T | C |  |
| AY.4 | T | T | G | T | T | C | C | T | C | C | T | A | G | G | A | A | A | G | T | G | T | T | #881108 |
| AY.4 | C | C | A | C | C | T | T | C | T | T | G | G | T | G | A | A | A | G | T | G | A | T |  |
| mut |  |  | A>G |  |  |  |  |  |  |  |  | G>A |  |  |  |  |  |  |  |  | A>T |  |  |

|  |  |  |  |  |  |  |  |  |  |  |  |  |  |  |  |  |  |  |  |  |  |  |  |  |  |  |  |
| --- | --- | --- | --- | --- | --- | --- | --- | --- | --- | --- | --- | --- | --- | --- | --- | --- | --- | --- | --- | --- | --- | --- | --- | --- | --- | --- | --- |
| pos | 507 | 508 | 509 | 518 | 3559 | 3971 | 6040 | 6638 | 7756 | 7926 | 9979 | 12037 | 13723 | 14014 | 15654 | 16726 | 17907 | 19118 | 19300 | 20055 | 20397 | 21372 | 21846 | 27604 | 28073 | 29700 |  |
| ref | A | T | G | A | A | G | C | C | G | C | C | T | G | T | C | C | T | C | G | A | G | G | C | G | G | A |  |
| AY.44 | T | C | A | G | A | T | T | T | G | T | C | T | G | G | C | T | T | C | G | A | G | G | C | G | T | G |  |
| AY.39 | T | C | A | G | C | G | C | C | T | C | T | C | T | T | T | C | C | T | T | G | A | T | T | A | G | A | #881676 |
| AY.39 | A | T | G | A | C | G | C | C | T | C | T | C | T | T | T | C | C | T | T | G | A | T | T | A | G | A |  |
| mut |  |  |  |  |  |  |  |  |  |  |  |  |  |  |  |  |  |  |  |  |  |  |  |  |  |  |  |

|  |  |  |  |  |  |  |  |  |  |  |  |  |  |  |  |  |  |  |  |
| --- | --- | --- | --- | --- | --- | --- | --- | --- | --- | --- | --- | --- | --- | --- | --- | --- | --- | --- | --- |
| pos | 1358 | 1499 | 6408 | 7829 | 7851 | 8473 | 12410 | 13426 | 13944 | 17040 | 19021 | 20946 | 25003 | 26488 | 26491 | 26492 | 26497 | 29543 |  |
| ref | G | T | C | G | C | T | A | C | C | T | G | C | A | T | T | A | T | G |  |
| AY.4 | A | G | C | T | T | T | G | T | C | C | A | T | A | T | T | A | T | T |  |
| AY.4 | A | G | C | T | T | T | G | T | C | C | A | T | A | G | C | T | C | T | #882353 |
| AY.120 | G | T | T | G | C | C | A | C | T | T | G | C | G | G | C | T | C | G |  |
| mut |  |  |  |  |  |  |  |  |  |  |  |  |  |  |  |  |  | G>T |  |

|  |  |  |  |  |  |  |  |  |  |  |  |  |  |  |  |  |  |  |  |  |  |  |  |  |
| --- | --- | --- | --- | --- | --- | --- | --- | --- | --- | --- | --- | --- | --- | --- | --- | --- | --- | --- | --- | --- | --- | --- | --- | --- |
| pos | 3562 | 4802 | 6312 | 6566 | 8349 | 8829 | 8956 | 10181 | 16092 | 17236 | 18032 | 18657 | 19006 | 21658 | 21846 | 23635 | 23987 | 24208 | 27005 | 27999 | 29247 | 29422 | 29659 |  |
| ref | G | G | C | G | G | C | C | A | C | A | C | C | G | C | C | C | C | C | C | C | C | G | C |  |
| AY.119 | G | G | T | T | G | T | C | C | T | A | C | C | T | C | T | C | T | C | C | C | T | G | C |  |
| AY.119 | G | G | T | T | G | T | C | C | T | A | C | C | T | C | T | C | C | T | T | T | C | T | T | #891208 |
| AY.103 | A | A | C | G | A | C | T | A | C | G | T | T | G | T | C | T | C | T | T | T | C | T | C |  |
| mut |  |  |  |  |  |  |  |  |  |  |  |  |  |  |  |  |  |  |  |  |  |  | C>T |  |

|  |  |  |  |  |  |  |  |  |  |  |  |  |  |  |  |  |  |  |
| --- | --- | --- | --- | --- | --- | --- | --- | --- | --- | --- | --- | --- | --- | --- | --- | --- | --- | --- |
| pos | 2587 | 2755 | 3439 | 5746 | 12794 | 14775 | 16128 | 22000 | 22027 | 22029 | 22030 | 22031 | 22032 | 22033 | 22034 | 23428 | 25074 |  |
| ref | T | G | T | C | G | T | A | C | T | A | G | T | T | C | A | T | G |  |
| AY.107 | C | T | C | T | A | C | G | T | T | - | - | - | - | - | - | C | T |  |
| AY.107 | C | T | C | T | A | C | G | T | G | A | A | A | G | T | G | C | G | #891268 |
| AY.107 | T | G | T | C | G | T | A | C | G | A | A | A | G | T | G | T | G |  |
| mut |  |  |  |  |  |  |  |  |  |  |  |  |  |  |  | T>C |  |  |

|  |  |  |  |  |  |  |  |  |  |  |  |  |  |  |  |  |  |  |
| --- | --- | --- | --- | --- | --- | --- | --- | --- | --- | --- | --- | --- | --- | --- | --- | --- | --- | --- |
| pos | 625 | 761 | 823 | 5203 | 12756 | 15183 | 18981 | 22027 | 22029 | 22030 | 22031 | 22032 | 22033 | 22034 | 25681 | 27600 | 29700 |  |
| ref | G | A | C | C | C | C | C | T | A | G | T | T | C | A | G | C | A |  |
| AY.119 | T | G | T | T | T | T | T | T | - | - | - | - | - | - | G | C | A |  |
| AY.119 | T | G | T | T | T | T | T | G | A | A | A | G | T | G | A | T | A | #891276 |
| AY.119 | G | A | C | C | C | C | C | G | A | A | A | G | T | G | G | C | G |  |
| mut |  |  |  |  |  |  |  |  |  |  |  |  |  |  | G>A | C>T | G>A |  |

|  |  |  |  |  |  |  |  |  |  |  |  |  |  |  |  |  |  |
| --- | --- | --- | --- | --- | --- | --- | --- | --- | --- | --- | --- | --- | --- | --- | --- | --- | --- |
| pos | 403 | 7305 | 7600 | 8809 | 9339 | 20719 | 22026 | 22027 | 22029 | 22030 | 22031 | 22032 | 22033 | 22034 | 25463 | 26907 |  |
| ref | T | T | C | C | A | C | G | T | A | G | T | T | C | A | C | C |  |
| AY.100 | A | C | C | T | G | T | G | T | - | - | - | - | - | - | T | T |  |
| AY.100 | A | C | T | T | G | T | G | G | A | A | A | G | T | G | T | T | #891349 |
| AY.100 | T | T | C | C | A | C | T | G | A | A | A | G | T | G | C | C |  |
| mut |  |  | C>T |  |  |  |  |  |  |  |  |  |  |  | C>T | C>T |  |

|  |  |  |  |  |  |  |  |  |  |  |  |  |  |  |  |  |  |  |  |  |  |  |  |  |  |  |  |  |  |  |
| --- | --- | --- | --- | --- | --- | --- | --- | --- | --- | --- | --- | --- | --- | --- | --- | --- | --- | --- | --- | --- | --- | --- | --- | --- | --- | --- | --- | --- | --- | --- |
| pos | 144 | 3094 | 7039 | 9951 | 10573 | 10829 | 11173 | 13768 | 16056 | 19540 | 19806 | 20166 | 21054 | 21108 | 23126 | 28241 | 28243 | 28245 | 28247 | 28251 | 28252 | 28253 | 28254 | 29700 | 29762 | 29769 | 29774 | 29779 | 29781 |  |
| ref | T | A | C | C | T | A | C | A | T | A | C | T | C | C | G | T | T | T | A | T | T | C | A | A | C | C | C | G | G |  |
| AY.3 | C | A | T | C | A | A | T | A | C | G | C | T | C | T | C | T | T | T | A | - | - | - | A | - | - | - | - | - | - |  |
| AY.3 | C | A | T | T | A | A | T | G | C | G | C | T | C | T | C | A | A | C | G | - | T | T | C | A | C | C | C | G | G | #894321 |
| AY.3 | T | G | C | C | T | C | C | A | T | A | T | C | T | C | G | A | A | C | G | T | T | - | - | A | C | C | C | G | G |  |
| mut |  |  |  | C>T |  |  |  | A>G |  |  |  |  |  |  |  |  |  |  |  | T>- |  | ->T | ->C |  |  |  |  |  |  |  |

|  |  |  |  |  |  |  |  |  |  |  |  |  |  |  |  |  |  |  |  |  |  |  |  |  |  |  |  |  |
| --- | --- | --- | --- | --- | --- | --- | --- | --- | --- | --- | --- | --- | --- | --- | --- | --- | --- | --- | --- | --- | --- | --- | --- | --- | --- | --- | --- | --- |
| pos | 2231 | 2268 | 2946 | 3304 | 3433 | 3948 | 8655 | 8849 | 9988 | 11562 | 13929 | 16818 | 17304 | 18312 | 19269 | 21372 | 21846 | 24098 | 25614 | 25644 | 26107 | 26499 | 27507 | 27604 | 29167 | 29521 | 29555 |  |
| ref | G | C | G | G | G | A | C | G | C | G | T | A | C | C | C | G | C | G | C | G | G | C | A | G | C | A | C |  |
| AY.39 | T | T | A | T | T | A | T | A | T | G | C | A | C | T | C | T | T | G | T | G | G | C | A | A | C | T | C |  |
| AY.39 | T | T | A | T | T | A | T | A | T | G | C | A | C | T | C | T | T | G | T | G | C | T | C | G | T | A | T | #894500 |
| AY.25.1 | G | C | G | G | G | G | C | G | C | T | T | G | T | C | T | G | C | T | C | T | C | T | C | G | T | A | T |  |
| mut |  |  |  |  |  |  |  |  |  |  |  |  |  |  |  |  |  |  |  |  |  |  |  |  |  |  |  |  |

|  |  |  |  |  |  |  |  |  |  |  |  |  |
| --- | --- | --- | --- | --- | --- | --- | --- | --- | --- | --- | --- | --- |
| pos | 7736 | 8118 | 8725 | 13768 | 22027 | 22029 | 22030 | 22031 | 22032 | 22033 | 22034 |  |
| ref | G | C | A | A | T | A | G | T | T | C | A |  |
| AY.119 | G | T | G | G | T | - | - | - | - | - | - |  |
| AY.119 | G | T | G | G | G | A | A | A | G | T | G | #894694 |
| AY.119 | A | C | A | A | G | A | A | A | G | T | G |  |
| mut |  |  |  |  |  |  |  |  |  |  |  |  |

|  |  |  |  |  |  |  |  |  |  |  |  |  |  |  |  |  |  |  |  |  |  |  |  |  |  |
| --- | --- | --- | --- | --- | --- | --- | --- | --- | --- | --- | --- | --- | --- | --- | --- | --- | --- | --- | --- | --- | --- | --- | --- | --- | --- |
| pos | 2556 | 4456 | 4833 | 6463 | 10310 | 12032 | 13482 | 15930 | 16915 | 17615 | 18086 | 19969 | 20208 | 22026 | 22027 | 22029 | 22030 | 22031 | 22032 | 22033 | 22034 | 22120 | 25678 | 29349 |  |
| ref | A | C | T | G | G | G | A | T | C | A | C | G | G | G | T | A | G | T | T | C | A | C | C | C |  |
| AY.39.1 | A | T | C | T | A | T | G | C | T | G | T | G | G | G | T | - | - | - | - | - | - | C | A | G |  |
| AY.39.1 | A | T | C | T | A | T | G | C | T | G | T | G | A | G | G | A | A | A | G | T | G | T | A | G | #894761 |
| AY.39 | T | C | T | G | G | G | A | T | C | A | C | T | G | T | G | A | A | A | G | T | G | T | C | C |  |
| mut |  |  |  |  |  |  |  |  |  |  |  |  | G>A |  |  |  |  |  |  |  |  |  | C>A | C>G |  |

|  |  |  |  |  |  |  |  |  |  |  |  |  |  |  |  |  |  |
| --- | --- | --- | --- | --- | --- | --- | --- | --- | --- | --- | --- | --- | --- | --- | --- | --- | --- |
| pos | 2165 | 6444 | 7728 | 18105 | 19164 | 22026 | 22027 | 22029 | 22030 | 22031 | 22032 | 22033 | 22034 | 23280 | 25854 | 25916 |  |
| ref | A | A | C | G | C | G | T | A | G | T | T | C | A | C | C | C |  |
| AY.100 | G | G | T | G | T | G | T | - | - | - | - | - | - | C | T | T |  |
| AY.100 | G | G | T | T | T | G | G | A | A | A | G | T | G | T | T | T | #894831 |
| AY.100 | A | A | C | G | C | T | G | A | A | A | G | T | G | C | C | C |  |
| mut |  |  |  | G>T |  |  |  |  |  |  |  |  |  | C>T | C>T | C>T |  |

|  |  |  |  |  |  |  |  |  |  |  |  |  |  |  |  |  |  |
| --- | --- | --- | --- | --- | --- | --- | --- | --- | --- | --- | --- | --- | --- | --- | --- | --- | --- |
| pos | 2100 | 5717 | 5812 | 10642 | 17304 | 21892 | 21893 | 22027 | 22029 | 22030 | 22031 | 22032 | 22033 | 22034 | 25528 | 27261 |  |
| ref | G | C | C | G | C | A | G | T | A | G | T | T | C | A | C | G |  |
| AY.119 | T | T | T | T | T | G | A | T | - | - | - | - | - | - | T | G |  |
| AY.119 | T | T | T | T | T | G | A | G | A | A | A | G | T | G | T | T | #894853 |
| AY.119 | G | C | C | G | C | A | G | G | A | A | A | G | T | G | C | G |  |
| mut |  |  |  |  |  |  |  |  |  |  |  |  |  |  | C>T | G>T |  |

|  |  |  |  |  |  |  |  |  |  |  |  |  |  |  |  |  |  |  |  |
| --- | --- | --- | --- | --- | --- | --- | --- | --- | --- | --- | --- | --- | --- | --- | --- | --- | --- | --- | --- |
| pos | 2180 | 8605 | 12513 | 14183 | 17002 | 20384 | 20679 | 21157 | 22026 | 22027 | 22029 | 22030 | 22031 | 22032 | 22033 | 22034 | 24023 | 25019 |  |
| ref | G | C | C | C | C | C | G | G | G | T | A | G | T | T | C | A | C | G |  |
| AY.100 | A | C | T | T | T | T | T | A | G | T | - | - | - | - | - | - | T | T |  |
| AY.100 | A | T | T | T | T | T | T | A | G | G | A | A | A | G | T | G | T | T | #894880 |
| AY.100 | G | C | C | C | C | C | G | G | T | G | A | A | A | G | T | G | C | G |  |
| mut |  | C>T |  |  |  |  |  |  |  |  |  |  |  |  |  |  | C>T | G>T |  |

|  |  |  |  |  |  |  |  |  |  |  |  |  |  |  |  |  |
| --- | --- | --- | --- | --- | --- | --- | --- | --- | --- | --- | --- | --- | --- | --- | --- | --- |
| pos | 3241 | 3653 | 5515 | 5924 | 6698 | 22882 | 22917 | 23599 | 23604 | 23664 | 23854 | 26530 | 26577 | 26709 | 26767 |  |
| ref | C | C | G | G | T | T | T | T | C | C | C | A | C | G | T |  |
| BA.1.17.2 | T | T | G | A | C | G | T | G | A | T | A | G | G | A | T |  |
| BA.1.17.2 | T | T | G | A | C | T | G | T | G | C | C | A | C | G | C | #896340 |
| BA.1 | C | C | T | G | T | T | G | T | G | C | C | A | C | G | C |  |
| mut |  |  |  |  |  | G>T |  |  |  |  |  |  |  |  |  |  |

|  |  |  |  |  |  |  |  |  |  |  |  |  |  |  |  |  |  |  |  |  |  |  |  |  |  |  |  |  |  |  |  |  |  |  |  |  |  |  |
| --- | --- | --- | --- | --- | --- | --- | --- | --- | --- | --- | --- | --- | --- | --- | --- | --- | --- | --- | --- | --- | --- | --- | --- | --- | --- | --- | --- | --- | --- | --- | --- | --- | --- | --- | --- | --- | --- | --- |
| pos | 3971 | 5164 | 5184 | 5584 | 6040 | 6402 | 6601 | 6638 | 7124 | 7926 | 8986 | 9053 | 9361 | 9891 | 10029 | 11201 | 11332 | 11418 | 11514 | 12049 | 13019 | 14014 | 15267 | 16049 | 16726 | 19220 | 22227 | 22427 | 24378 | 25669 | 25731 | 25886 | 27874 | 28073 | 28326 | 28916 | 29700 |  |
| ref | G | G | C | A | C | C | A | C | C | C | C | G | A | C | C | A | A | T | C | C | C | T | A | C | C | C | C | G | C | C | C | C | C | G | G | G | A |  |
| AY.47 | G | T | T | G | C | C | A | C | C | C | C | G | A | T | C | A | A | C | T | T | T | T | A | C | C | C | T | A | C | T | T | C | C | G | T | G | A |  |
| B.1.617.2 | T | T | T | G | C | C | A | C | C | C | C | G | G | C | T | G | G | T | C | C | C | G | G | T | T | T | C | G | T | C | C | T | T | T | G | T | G | #897177 |
| AY.44 | T | G | C | A | T | T | C | T | T | T | T | T | A | C | T | G | G | T | C | C | C | G | G | T | T | T | C | G | T | C | C | T | T | T | G | T | G |  |
| mut | G>T |  |  |  |  |  |  |  |  |  |  |  | A>G |  |  |  |  |  |  |  |  |  |  |  |  |  |  |  |  |  |  |  |  |  |  |  |  |  |

|  |  |  |  |  |  |  |  |  |  |  |  |  |  |  |  |  |  |
| --- | --- | --- | --- | --- | --- | --- | --- | --- | --- | --- | --- | --- | --- | --- | --- | --- | --- |
| pos | 3241 | 5515 | 5924 | 6988 | 12309 | 22882 | 22917 | 23599 | 23604 | 23664 | 23854 | 26530 | 26577 | 26709 | 26767 | 26890 |  |
| ref | C | G | G | T | A | T | T | T | C | C | C | A | C | G | T | C |  |
| BA.1.17.2 | T | G | A | C | G | G | T | G | A | T | A | G | G | A | T | C |  |
| BA.1.17.2 | T | G | A | C | G | T | G | T | G | C | C | A | C | G | C | T | #898770 |
| BA.1 | C | T | G | T | A | T | G | T | G | C | C | A | C | G | C | C |  |
| mut |  |  |  |  |  | G>T |  |  |  |  |  |  |  |  |  | C>T |  |

|  |  |  |  |  |  |  |  |  |  |  |  |  |  |  |  |  |  |  |  |  |  |  |  |  |  |  |
| --- | --- | --- | --- | --- | --- | --- | --- | --- | --- | --- | --- | --- | --- | --- | --- | --- | --- | --- | --- | --- | --- | --- | --- | --- | --- | --- |
| pos | 1147 | 9179 | 10354 | 11335 | 11456 | 11572 | 17236 | 18463 | 19906 | 20703 | 21321 | 23284 | 24208 | 25339 | 27014 | 27291 | 29050 | 29509 | 29700 | 29762 | 29769 | 29774 | 29779 | 29781 | 29836 |  |
| ref | T | T | G | G | A | C | A | C | A | C | T | T | C | C | G | T | G | C | A | C | C | C | G | G | C |  |
| AY.3 | T | C | G | T | G | C | A | C | G | C | T | C | C | T | G | T | A | T | - | - | - | - | - | - | C |  |
| AY.3 | T | C | G | T | G | C | A | C | G | C | C | T | T | C | T | C | G | C | A | C | C | C | G | G | T | #899982 |
| AY.103 | C | T | T | G | A | T | G | A | A | T | T | T | T | C | T | C | G | C | A | C | C | C | G | G | T |  |
| mut |  |  |  |  |  |  |  |  |  |  | T>C |  |  |  |  |  |  |  |  |  |  |  |  |  |  |  |

|  |  |  |  |  |  |  |  |  |  |  |  |  |  |  |  |  |  |  |  |  |  |  |  |  |  |
| --- | --- | --- | --- | --- | --- | --- | --- | --- | --- | --- | --- | --- | --- | --- | --- | --- | --- | --- | --- | --- | --- | --- | --- | --- | --- |
| pos | 347 | 1078 | 1093 | 1593 | 6255 | 7851 | 10323 | 11365 | 11758 | 13971 | 14998 | 15120 | 15924 | 17192 | 19710 | 21301 | 22213 | 23120 | 23625 | 25720 | 27488 | 28028 | 28198 | 29782 |  |
| ref | G | A | C | C | C | C | A | G | C | C | A | C | C | A | T | C | T | G | C | G | A | G | G | A |  |
| AY.124 | G | G | C | T | T | C | G | T | T | T | G | T | T | G | C | A | G | G | T | T | G | T | T | A |  |
| AY.124 | A | G | C | T | T | C | G | T | T | T | G | T | T | G | C | A | G | T | C | G | A | G | G | G | #904404 |
| AY.4 | G | A | T | C | C | T | A | G | C | C | A | C | C | A | T | C | T | T | C | G | A | G | G | A |  |
| mut | G>A |  |  |  |  |  |  |  |  |  |  |  |  |  |  |  |  |  |  |  |  |  |  | A>G |  |

|  |  |  |  |  |  |  |  |  |  |  |  |  |  |  |  |  |  |  |  |  |
| --- | --- | --- | --- | --- | --- | --- | --- | --- | --- | --- | --- | --- | --- | --- | --- | --- | --- | --- | --- | --- |
| pos | 2470 | 7926 | 9166 | 16744 | 22194 | 22195 | 22196 | 22197 | 22198 | 22202 | 22204 | 22882 | 22992 | 23013 | 23040 | 23048 | 23055 | 23063 | 23075 |  |
| ref | C | C | C | G | A | T | T | T | A | C | T | T | G | A | A | G | A | A | T |  |
| BA.1.1.18 | T | T | C | G | - | - | - | T | A | C | T | G | A | C | G | A | G | T | C |  |
| BA.1.1.18 | T | T | C | G | - | - | - | T | A | C | T | T | A | C | G | G | A | A | T | #908926 |
| BA.1.1 | C | C | T | A | A | G | T | G | C | A | C | G | G | A | A | G | A | A | T |  |
| mut |  |  |  |  |  |  |  |  |  |  |  | G>T |  |  |  |  |  |  |  |  |

|  |  |  |  |  |  |  |  |  |  |  |  |  |  |  |  |  |  |  |  |  |  |  |  |  |  |  |  |  |  |  |  |  |  |  |
| --- | --- | --- | --- | --- | --- | --- | --- | --- | --- | --- | --- | --- | --- | --- | --- | --- | --- | --- | --- | --- | --- | --- | --- | --- | --- | --- | --- | --- | --- | --- | --- | --- | --- | --- |
| pos | 2470 | 9166 | 16744 | 21595 | 22029 | 22030 | 22031 | 22032 | 22033 | 22034 | 22194 | 22195 | 22196 | 22197 | 22198 | 22202 | 22204 | 22578 | 22599 | 22673 | 22674 | 22679 | 22686 | 22882 | 22992 | 23013 | 23040 | 23048 | 23055 | 23063 | 23075 | 25566 | 28269 |  |
| ref | C | C | G | C | A | G | T | T | C | A | A | T | T | T | A | C | T | G | G | T | C | T | C | T | G | A | A | G | A | A | T | C | C |  |
| BA.1.1 | T | C | G | T | A | G | T | T | C | A | - | - | - | T | A | C | T | G | G | T | C | T | C | G | A | C | G | A | G | T | C | T | T |  |
| BA.1 | T | C | G | T | - | - | - | - | - | - | - | - | - | T | A | C | T | G | G | T | C | T | C | T | A | A | A | G | A | A | T | C | C | #908970 |
| BA.1.1 | C | T | A | C | A | G | T | T | C | A | A | G | T | G | C | A | C | A | A | C | T | C | T | G | G | A | A | G | A | A | T | C | C |  |
| mut |  |  |  |  | A>- | G>- | T>- | T>- | C>- | A>- |  |  |  |  |  |  |  |  |  |  |  |  |  | G>T |  |  |  |  |  |  |  |  |  |  |

|  |  |  |  |  |  |  |  |  |  |  |  |  |  |  |  |  |  |  |  |  |  |  |  |
| --- | --- | --- | --- | --- | --- | --- | --- | --- | --- | --- | --- | --- | --- | --- | --- | --- | --- | --- | --- | --- | --- | --- | --- |
| pos | 6408 | 7303 | 7420 | 7600 | 8771 | 8829 | 9733 | 11095 | 12088 | 13275 | 18744 | 19006 | 20259 | 21987 | 25003 | 25452 | 26606 | 26894 | 27572 | 27573 | 27575 | 27576 |  |
| ref | C | C | C | C | A | C | C | C | A | C | C | G | C | G | A | C | C | C | G | C | C | T |  |
| AY.119 | C | T | T | C | G | T | T | C | G | T | T | T | T | G | A | C | T | T | G | C | C | T |  |
| AY.119 | C | T | T | C | G | T | T | C | G | T | T | T | T | G | A | C | T | T | A | T | A | C | #913020 |
| AY.120.1 | T | C | C | T | A | C | C | T | A | C | C | G | C | A | G | T | C | C | A | T | A | C |  |
| mut |  |  |  |  |  |  |  |  |  |  |  |  |  |  |  |  |  |  |  |  |  |  |  |

|  |  |  |  |  |  |  |  |  |  |  |  |  |  |  |  |
| --- | --- | --- | --- | --- | --- | --- | --- | --- | --- | --- | --- | --- | --- | --- | --- |
| pos | 2470 | 3241 | 5924 | 14940 | 17241 | 22599 | 23664 | 25047 | 27377 | 27378 | 27379 | 27382 | 27383 | 27384 |  |
| ref | C | C | G | A | T | G | C | C | A | G | A | G | A | T |  |
| BA.2.2.1 | T | C | G | G | T | A | C | C | A | G | A | G | A | T |  |
| BA.1.1 | T | C | G | G | T | A | C | C | C | T | G | C | T | C | #925211 |
| BA.1.17.2 | C | T | A | A | C | G | T | T | C | T | G | C | T | C |  |
| mut |  |  |  |  |  |  |  |  |  |  |  |  |  |  |  |

|  |  |  |  |  |  |  |  |  |  |  |  |  |  |  |
| --- | --- | --- | --- | --- | --- | --- | --- | --- | --- | --- | --- | --- | --- | --- |
| pos | 4551 | 6428 | 15543 | 21618 | 22026 | 22027 | 22029 | 22030 | 22031 | 22032 | 22033 | 22034 | 27510 |  |
| ref | C | C | G | C | G | T | A | G | T | T | C | A | A |  |
| AY.39 | T | T | T | G | G | T | - | - | - | - | - | - | G |  |
| AY.39 | T | T | T | G | G | G | A | A | A | G | T | G | G | #925497 |
| AY.39 | C | C | G | T | T | G | A | A | A | G | T | G | A |  |
| mut |  |  |  |  |  |  |  |  |  |  |  |  | A>G |  |

|  |  |  |  |  |  |  |  |  |  |  |  |  |  |  |  |  |  |  |
| --- | --- | --- | --- | --- | --- | --- | --- | --- | --- | --- | --- | --- | --- | --- | --- | --- | --- | --- |
| pos | 352 | 390 | 1251 | 1820 | 5339 | 7851 | 10977 | 12880 | 15569 | 16242 | 17040 | 21846 | 21995 | 25614 | 25906 | 26885 | 28845 |  |
| ref | T | C | C | G | C | C | C | C | C | T | T | C | T | C | G | C | G |  |
| AY.46.5 | T | C | C | G | C | C | T | T | T | G | T | C | T | C | G | C | G |  |
| AY.46.5 | C | C | C | G | T | C | T | T | T | G | T | C | T | T | T | T | A | #929067 |
| AY.4.2 | T | T | T | A | C | T | C | C | C | T | C | T | C | T | T | T | A |  |
| mut | T>C |  |  |  | C>T |  |  |  |  |  |  |  |  |  |  |  |  |  |

|  |  |  |  |  |  |  |  |  |  |  |  |  |  |  |
| --- | --- | --- | --- | --- | --- | --- | --- | --- | --- | --- | --- | --- | --- | --- |
| pos | 2470 | 5183 | 6677 | 8326 | 8647 | 16575 | 21595 | 21622 | 22599 | 22813 | 22882 | 22898 | 22917 |  |
| ref | C | C | A | C | T | C | C | C | G | G | T | G | T |  |
| BA.1 | C | T | G | C | C | C | C | T | G | G | T | G | G |  |
| BA.1 | C | T | G | C | C | T | C | T | G | T | G | A | T | #934558 |
| BA.1.1 | T | C | A | T | T | T | T | C | A | T | T | A | T |  |
| mut |  |  |  |  |  |  | T>C | C>T | A>G |  | T>G |  |  |  |

|  |  |  |  |  |  |  |  |  |  |  |  |  |  |  |  |  |  |
| --- | --- | --- | --- | --- | --- | --- | --- | --- | --- | --- | --- | --- | --- | --- | --- | --- | --- |
| pos | 2470 | 3241 | 5175 | 5924 | 6988 | 12309 | 21595 | 22599 | 22813 | 22898 | 23664 | 27874 | 27875 | 27881 | 27882 | 27883 |  |
| ref | C | C | C | G | T | A | C | G | G | G | C | C | T | C | G | C |  |
| BA.1.1 | T | C | T | G | T | A | T | A | T | A | C | C | T | C | G | C |  |
| BA.1.1 | T | C | T | G | T | A | T | A | T | A | C | T | C | T | C | T | #936638 |
| BA.1.17.2 | C | T | C | A | C | G | C | G | G | G | T | T | C | T | C | T |  |
| mut |  |  |  |  |  |  |  |  |  |  |  |  |  |  |  |  |  |

|  |  |  |  |  |  |  |  |  |  |  |  |  |  |  |  |  |  |  |  |  |  |  |  |  |  |  |  |  |  |  |  |  |  |  |  |  |  |  |  |  |  |  |  |  |  |  |  |  |  |  |  |  |  |  |  |  |  |  |  |  |  |  |  |  |  |  |  |  |  |  |  |  |  |  |  |  |  |  |  |  |  |  |  |  |
| --- | --- | --- | --- | --- | --- | --- | --- | --- | --- | --- | --- | --- | --- | --- | --- | --- | --- | --- | --- | --- | --- | --- | --- | --- | --- | --- | --- | --- | --- | --- | --- | --- | --- | --- | --- | --- | --- | --- | --- | --- | --- | --- | --- | --- | --- | --- | --- | --- | --- | --- | --- | --- | --- | --- | --- | --- | --- | --- | --- | --- | --- | --- | --- | --- | --- | --- | --- | --- | --- | --- | --- | --- | --- | --- | --- | --- | --- | --- | --- | --- | --- | --- | --- | --- |
| pos | 670 | 2470 | 2790 | 2832 | 4184 | 4321 | 5386 | 6513 | 6514 | 6515 | 8393 | 9344 | 9424 | 9534 | 10198 | 10447 | 11283 | 11284 | 11285 | 11286 | 11287 | 11292 | 11293 | 11294 | 11295 | 11296 | 11537 | 12880 | 13195 | 15240 | 15714 | 17410 | 21595 | 21618 | 21633 | 21634 | 21635 | 21636 | 21637 | 21638 | 21639 | 21640 | 21641 | 21762 | 21765 | 21766 | 21767 | 21768 | 21769 | 21770 | 21846 | 21987 | 21988 | 21989 | 21990 | 21991 | 21992 | 21993 | 21994 | 21995 | 22195 | 22197 | 22198 | 22200 | 22202 | 22204 | 22599 | 22673 | 22688 | 22775 | 22786 | 22898 | 23048 | 23202 | 24130 | 24503 | 26060 | 26530 | 26858 | 27382 | 27383 | 27384 | 29510 |  |
| ref | T | C | C | A | G | C | T | G | T | T | G | C | A | C | C | G | G | T | T | T | G | G | T | T | T | T | A | C | T | C | C | C | C | C | T | A | C | C | C | C | C | T | G | C | T | A | C | A | T | G | C | G | T | G | T | T | T | A | T | T | T | T | A | T | C | T | G | T | A | G | A | G | G | C | C | C | C | A | C | G | A | T | A |  |
| BA.1.1 | T | T | C | G | G | C | G | - | - | - | A | C | A | C | C | G | - | - | - | - | - | G | T | T | T | T | G | C | C | T | C | C | T | C | T | A | C | C | C | C | C | T | G | T | - | - | - | - | - | - | T | - | - | - | - | - | - | - | - | - | G | G | C | T | A | C | A | C | A | G | A | A | A | A | A | T | C | G | C | G | A | T | A |  |
| XM | T | T | C | G | G | C | G | - | - | - | A | C | A | C | C | G | - | - | - | - | - | G | T | T | T | T | G | C | C | T | C | C | C | C | T | A | C | C | C | C | C | T | G | T | - | - | - | - | - | - | T | - | - | - | - | - | - | - | - | - | G | G | C | T | A | C | G | T | G | A | C | G | G | C | C | C | T | A | T | C | T | C | C | #938900 |
| BA.2 | G | C | T | A | A | T | T | G | T | T | G | T | G | T | T | A | G | T | T | T | G | - | - | - | - | - | A | T | T | C | T | T | C | T | - | - | - | - | - | - | - | - | - | C | T | A | C | A | T | G | C | A | T | G | T | T | T | A | T | T | T | T | A | G | C | T | G | T | G | A | C | G | G | C | C | C | T | A | T | C | T | C | C |  |
| mut |  |  |  |  |  |  |  |  |  |  |  |  |  |  |  |  |  |  |  |  |  |  |  |  |  |  |  |  |  |  |  |  | T>C |  |  |  |  |  |  |  |  |  |  |  |  |  |  |  |  |  |  |  |  |  |  |  |  |  |  |  |  |  |  |  |  |  | A>G |  |  |  |  |  |  |  |  |  |  |  |  |  |  |  |  |  |

|  |  |  |  |  |  |  |  |  |  |  |
| --- | --- | --- | --- | --- | --- | --- | --- | --- | --- | --- |
| pos | 3393 | 5703 | 5704 | 5705 | 6988 | 22813 | 22898 | 26607 | 29545 |  |
| ref | C | A | G | T | T | G | G | C | C |  |
| BA.1.17.2 | C | T | A | A | C | G | G | T | C |  |
| BA.1.17.2 | C | T | A | A | T | T | A | C | A | #939917 |
| BA.1.17.2 | T | A | G | T | T | T | A | C | A |  |
| mut |  |  |  |  |  |  |  |  |  |  |

|  |  |  |  |  |  |  |  |  |  |  |  |  |  |  |  |  |  |  |  |
| --- | --- | --- | --- | --- | --- | --- | --- | --- | --- | --- | --- | --- | --- | --- | --- | --- | --- | --- | --- |
| pos | 2470 | 3535 | 4810 | 9042 | 9049 | 14107 | 22194 | 22195 | 22196 | 22197 | 22198 | 22203 | 22204 | 22207 | 22599 | 22882 | 25000 | 29781 |  |
| ref | C | C | C | C | G | A | A | T | T | T | A | G | T | T | G | T | C | G |  |
| BA.1.1.14 | T | T | A | T | A | G | - | - | - | T | A | G | T | T | A | T | C | G |  |
| BA.1.1.14 | T | T | A | T | A | G | A | G | T | G | C | C | A | A | A | G | T | G | #941092 |
| BA.1 | C | C | C | C | G | A | A | G | T | G | C | C | A | A | G | G | T | - |  |
| mut |  |  |  |  |  |  |  |  |  |  |  |  |  |  | G>A |  |  | ->G |  |

|  |  |  |  |  |  |  |  |  |  |  |  |  |  |  |  |  |  |  |  |  |  |  |  |  |  |  |  |  |  |  |  |  |  |  |  |  |  |  |  |  |  |  |  |  |  |  |  |  |  |  |  |  |  |  |  |  |  |  |  |  |  |  |  |  |  |  |  |  |  |  |  |  |  |  |  |  |  |  |  |  |  |  |  |  |  |  |  |  |  |  |  |  |  |  |  |  |  |  |  |  |  |  |  |  |  |  |  |  |  |  |  |  |  |  |  |  |  |  |  |  |  |  |  |  |  |  |  |  |  |
| --- | --- | --- | --- | --- | --- | --- | --- | --- | --- | --- | --- | --- | --- | --- | --- | --- | --- | --- | --- | --- | --- | --- | --- | --- | --- | --- | --- | --- | --- | --- | --- | --- | --- | --- | --- | --- | --- | --- | --- | --- | --- | --- | --- | --- | --- | --- | --- | --- | --- | --- | --- | --- | --- | --- | --- | --- | --- | --- | --- | --- | --- | --- | --- | --- | --- | --- | --- | --- | --- | --- | --- | --- | --- | --- | --- | --- | --- | --- | --- | --- | --- | --- | --- | --- | --- | --- | --- | --- | --- | --- | --- | --- | --- | --- | --- | --- | --- | --- | --- | --- | --- | --- | --- | --- | --- | --- | --- | --- | --- | --- | --- | --- | --- | --- | --- | --- | --- | --- | --- | --- | --- | --- | --- | --- | --- | --- | --- | --- | --- |
| pos | 210 | 1390 | 1426 | 2255 | 2832 | 4181 | 5386 | 6402 | 6513 | 6514 | 6515 | 7124 | 7851 | 8393 | 8986 | 9053 | 10449 | 11201 | 11283 | 11284 | 11285 | 11286 | 11287 | 11288 | 11289 | 11290 | 11291 | 11332 | 11537 | 13195 | 14801 | 15240 | 15451 | 16466 | 18163 | 19220 | 21618 | 21762 | 21765 | 21766 | 21767 | 21768 | 21769 | 21770 | 21987 | 21988 | 21989 | 21990 | 21991 | 21992 | 21993 | 21994 | 21995 | 22029 | 22030 | 22031 | 22032 | 22033 | 22034 | 22051 | 22194 | 22195 | 22196 | 22578 | 22673 | 22674 | 22679 | 22686 | 22813 | 22882 | 22898 | 22917 | 22992 | 23013 | 23040 | 23048 | 23055 | 23063 | 23075 | 23202 | 23525 | 23599 | 23604 | 23854 | 23948 | 24130 | 24410 | 24424 | 24469 | 24503 | 25000 | 25469 | 25584 | 26270 | 26530 | 26577 | 26709 | 26767 | 27259 | 27638 | 27752 | 27807 | 27874 | 28248 | 28249 | 28250 | 28251 | 28252 | 28253 | 28271 | 28311 | 28362 | 28363 | 28364 | 28365 | 28366 | 28367 | 28368 | 28369 | 28370 | 28461 | 28881 | 28882 | 28883 | 28916 | 29402 | 29632 | 29742 |  |
| ref | G | T | C | A | A | G | T | C | G | T | T | C | C | G | C | G | C | A | G | T | T | T | G | T | C | T | G | A | A | T | A | C | G | C | A | C | C | C | T | A | C | A | T | G | G | T | G | T | T | T | A | T | T | A | G | T | T | C | A | G | A | T | T | G | T | C | T | C | G | T | G | T | G | A | A | G | A | A | T | C | C | T | C | C | G | C | G | A | T | C | C | C | C | C | A | C | G | T | A | T | C | C | C | G | A | T | T | T | C | A | C | G | A | G | A | A | C | G | C | A | A | G | G | G | G | G | C | G |  |
| AY.4 | T | C | C | A | A | T | T | T | G | T | T | T | T | G | T | T | C | G | G | T | T | T | G | T | C | T | G | G | A | T | C | C | A | T | A | T | G | C | T | A | C | A | T | G | A | T | G | T | T | T | A | T | T | - | - | - | - | - | - | C | A | T | T | G | T | C | T | C | G | T | G | G | G | A | A | G | A | A | T | C | C | T | G | C | G | C | A | A | T | C | C | T | C | C | A | C | G | C | A | C | T | C | T | - | - | - | - | - | - | - | C | G | A | G | A | A | C | G | C | A | G | T | G | G | T | T | C | T |  |
| XF | T | C | C | G | A | T | T | C | - | - | - | C | C | A | C | G | A | A | - | - | - | - | - | - | - | - | - | A | G | C | A | T | G | C | G | C | C | T | - | - | - | - | - | - | - | - | - | - | - | - | - | - | - | A | G | T | T | C | A | G | - | - | - | A | C | T | C | T | T | G | A | T | A | C | G | A | G | T | C | A | T | G | A | A | T | A | G | T | A | T | T | C | T | T | G | G | A | T | C | T | C | T | C | G | A | T | T | T | C | T | T | - | - | - | - | - | - | - | - | - | A | A | A | C | G | G | T | G | #946761 |
| BA.1 | G | T | T | A | G | G | G | C | - | - | - | C | C | A | C | G | A | A | - | - | - | - | - | - | - | - | - | A | G | C | A | T | G | C | G | C | C | T | - | - | - | - | - | - | - | - | - | - | - | - | - | - | - | A | G | T | T | C | A | G | - | - | - | A | C | T | C | T | T | G | A | T | A | C | G | A | G | T | C | A | T | G | A | A | T | A | G | T | A | T | T | C | T | T | G | G | A | T | C | T | C | T | C | G | A | T | T | T | C | T | T | - | - | - | - | - | - | - | - | - | A | A | A | C | G | G | T | G |  |
| mut |  |  |  | A>G |  |  |  |  |  |  |  |  |  |  |  |  |  |  |  |  |  |  |  |  |  |  |  |  |  |  |  |  |  |  |  |  |  |  |  |  |  |  |  |  |  |  |  |  |  |  |  |  |  |  |  |  |  |  |  |  |  |  |  |  |  |  |  |  |  |  |  |  |  |  |  |  |  |  |  |  |  |  |  |  |  |  |  |  |  |  |  |  |  |  |  |  |  |  |  |  |  |  |  |  |  |  |  |  |  |  |  |  |  |  |  |  |  |  |  |  |  |  |  |  |  |  |  |  |  |

|  |  |  |  |  |  |  |  |  |  |  |  |  |  |  |  |  |  |  |  |  |  |  |  |  |  |  |  |  |  |  |  |  |  |  |  |  |  |  |  |  |  |  |  |  |  |  |  |  |  |  |  |  |  |  |  |  |  |  |  |  |  |  |  |  |  |  |  |  |  |  |  |  |  |  |  |  |  |  |  |  |  |  |  |  |  |  |
| --- | --- | --- | --- | --- | --- | --- | --- | --- | --- | --- | --- | --- | --- | --- | --- | --- | --- | --- | --- | --- | --- | --- | --- | --- | --- | --- | --- | --- | --- | --- | --- | --- | --- | --- | --- | --- | --- | --- | --- | --- | --- | --- | --- | --- | --- | --- | --- | --- | --- | --- | --- | --- | --- | --- | --- | --- | --- | --- | --- | --- | --- | --- | --- | --- | --- | --- | --- | --- | --- | --- | --- | --- | --- | --- | --- | --- | --- | --- | --- | --- | --- | --- | --- | --- | --- | --- |
| pos | 44 | 670 | 2470 | 2790 | 2832 | 4184 | 4321 | 5386 | 6513 | 6514 | 6515 | 8393 | 9344 | 9424 | 9534 | 9866 | 10198 | 10447 | 11283 | 11284 | 11285 | 11286 | 11287 | 11292 | 11293 | 11294 | 11295 | 11296 | 11537 | 12880 | 15240 | 15714 | 17410 | 19955 | 20055 | 21595 | 21618 | 21633 | 21634 | 21635 | 21636 | 21637 | 21638 | 21639 | 21640 | 21641 | 21762 | 21765 | 21766 | 21767 | 21768 | 21769 | 21770 | 21846 | 21987 | 21988 | 21989 | 21990 | 21991 | 21992 | 21993 | 21994 | 21995 | 22194 | 22195 | 22196 | 22200 | 22599 | 22673 | 22688 | 22775 | 22786 | 22898 | 23048 | 23202 | 24130 | 24503 | 25416 | 26060 | 26530 | 26858 | 27382 | 27383 | 27384 | 29510 |  |
| ref | C | T | C | C | A | G | C | T | G | T | T | G | C | A | C | C | C | G | G | T | T | T | G | G | T | T | T | T | A | C | C | C | C | C | A | C | C | T | A | C | C | C | C | C | T | G | C | T | A | C | A | T | G | C | G | T | G | T | T | T | A | T | T | A | T | T | T | G | T | A | G | A | G | G | C | C | C | C | C | A | C | G | A | T | A |  |
| BA.1.1 | C | T | T | C | G | G | C | G | - | - | - | A | C | A | C | C | C | G | - | - | - | - | - | G | T | T | T | T | G | C | T | C | C | C | A | T | C | T | A | C | C | C | C | C | T | G | T | - | - | - | - | - | - | T | - | - | - | - | - | - | - | - | - | - | - | - | T | A | C | A | G | A | A | A | A | A | T | C | C | G | C | G | A | T | A |  |
| BA.2 | C | T | T | C | G | G | C | G | - | - | - | A | C | A | C | C | C | G | - | - | - | - | - | G | T | T | T | T | G | C | C | T | T | T | G | C | T | - | - | - | - | - | - | - | - | - | C | T | A | C | A | T | G | C | A | T | G | T | T | T | A | T | T | A | T | T | G | G | T | G | A | C | G | G | C | C | C | T | T | A | T | C | T | C | C | #960400 |
| BA.2 | T | G | C | T | A | A | T | T | G | T | T | G | T | G | T | T | T | A | G | T | T | T | G | - | - | - | - | - | A | T | C | T | T | T | G | C | T | - | - | - | - | - | - | - | - | - | C | T | A | C | A | T | G | C | A | T | G | T | T | T | A | T | T | A | T | T | G | G | T | G | A | C | G | G | C | C | C | T | T | A | T | C | T | C | C |  |
| mut |  |  |  |  |  |  |  |  |  |  |  |  |  |  |  |  |  |  |  |  |  |  |  |  |  |  |  |  |  |  | T>C |  |  |  |  |  |  |  |  |  |  |  |  |  |  |  |  |  |  |  |  |  |  |  |  |  |  |  |  |  |  |  |  |  |  |  |  |  |  |  |  |  |  |  |  |  |  |  |  |  |  |  |  |  |  |  |

|  |  |  |  |  |  |  |  |  |  |  |  |
| --- | --- | --- | --- | --- | --- | --- | --- | --- | --- | --- | --- |
| pos | 3241 | 5924 | 6988 | 10955 | 12459 | 20762 | 23664 | 23876 | 26607 | 29632 |  |
| ref | C | G | T | T | C | C | C | G | C | C |  |
| BA.1 | C | G | T | T | T | C | C | G | C | T |  |
| BA.1 | C | G | T | G | T | T | T | A | T | C | #960706 |
| BA.1.17.2 | T | A | C | T | C | T | T | A | T | C |  |
| mut |  |  |  | T>G |  |  |  |  |  |  |  |

|  |  |  |  |  |  |  |  |  |  |  |  |  |  |  |  |  |  |  |  |  |  |  |  |  |  |  |  |  |  |  |  |  |  |  |  |  |  |  |  |  |  |  |  |  |  |  |  |  |  |  |  |  |  |  |  |  |  |  |  |  |  |  |  |  |  |  |  |  |  |  |  |  |  |  |  |  |  |  |  |  |  |  |  |  |  |  |  |  |  |  |  |  |  |  |  |  |  |  |  |  |  |  |  |  |  |  |  |  |  |  |  |  |  |  |  |  |  |  |  |  |  |  |  |  |  |  |  |  |  |  |  |  |  |  |  |
| --- | --- | --- | --- | --- | --- | --- | --- | --- | --- | --- | --- | --- | --- | --- | --- | --- | --- | --- | --- | --- | --- | --- | --- | --- | --- | --- | --- | --- | --- | --- | --- | --- | --- | --- | --- | --- | --- | --- | --- | --- | --- | --- | --- | --- | --- | --- | --- | --- | --- | --- | --- | --- | --- | --- | --- | --- | --- | --- | --- | --- | --- | --- | --- | --- | --- | --- | --- | --- | --- | --- | --- | --- | --- | --- | --- | --- | --- | --- | --- | --- | --- | --- | --- | --- | --- | --- | --- | --- | --- | --- | --- | --- | --- | --- | --- | --- | --- | --- | --- | --- | --- | --- | --- | --- | --- | --- | --- | --- | --- | --- | --- | --- | --- | --- | --- | --- | --- | --- | --- | --- | --- | --- | --- | --- | --- | --- | --- | --- | --- | --- | --- | --- | --- | --- | --- |
| pos | 210 | 2832 | 2842 | 4181 | 5386 | 6402 | 6513 | 6514 | 6515 | 6740 | 7124 | 7851 | 8393 | 8986 | 9053 | 10449 | 11201 | 11283 | 11284 | 11285 | 11286 | 11287 | 11288 | 11289 | 11290 | 11291 | 11332 | 11537 | 13195 | 14457 | 14844 | 15240 | 15451 | 16466 | 18163 | 19220 | 21292 | 21618 | 21762 | 21765 | 21766 | 21767 | 21768 | 21769 | 21770 | 21898 | 21987 | 21988 | 21989 | 21990 | 21991 | 21992 | 21993 | 21994 | 21995 | 21997 | 22029 | 22030 | 22031 | 22032 | 22033 | 22034 | 22194 | 22195 | 22196 | 22578 | 22673 | 22674 | 22679 | 22686 | 22813 | 22882 | 22898 | 22917 | 22992 | 23013 | 23040 | 23048 | 23055 | 23063 | 23075 | 23202 | 23525 | 23599 | 23604 | 23854 | 23948 | 24130 | 24410 | 24424 | 24469 | 24503 | 25000 | 25469 | 25584 | 25714 | 26270 | 26530 | 26534 | 26577 | 26709 | 26767 | 26834 | 26873 | 27259 | 27638 | 27752 | 27807 | 27874 | 28248 | 28249 | 28250 | 28251 | 28252 | 28253 | 28271 | 28311 | 28362 | 28363 | 28364 | 28365 | 28366 | 28367 | 28368 | 28369 | 28370 | 28461 | 28881 | 28882 | 28883 | 28916 | 29402 | 29666 | 29742 |  |
| ref | G | A | C | G | T | C | G | T | T | C | C | C | G | C | G | C | A | G | T | T | T | G | T | C | T | G | A | A | T | A | A | C | G | C | A | C | C | C | C | T | A | C | A | T | G | G | G | T | G | T | T | T | A | T | T | C | A | G | T | T | C | A | A | T | T | G | T | C | T | C | G | T | G | T | G | A | A | G | A | A | T | C | C | T | C | C | G | C | G | A | T | C | C | C | C | C | C | A | C | C | G | T | G | C | A | T | C | C | C | G | A | T | T | T | C | A | C | G | A | G | A | A | C | G | C | A | A | G | G | G | G | G | C | G |  |
| AY.4 | T | A | T | T | T | T | G | T | T | T | T | T | G | T | T | C | G | G | T | T | T | G | T | C | T | G | G | A | T | T | A | C | A | T | A | T | T | G | C | T | A | C | A | T | G | T | A | T | G | T | T | T | A | T | T | T | - | - | - | - | - | - | A | T | T | G | T | C | T | C | G | T | G | G | G | A | A | G | A | A | T | C | C | T | G | C | G | C | A | A | T | C | C | T | C | T | C | A | C | C | G | C | T | C | A | C | T | C | T | - | - | - | - | - | - | - | C | G | A | G | A | A | C | G | C | A | G | T | G | G | T | T | C | T |  |
| BA.1 | T | A | T | T | T | T | G | T | T | T | T | T | G | T | T | C | G | G | T | T | T | G | T | C | T | G | G | A | T | T | G | C | A | T | A | T | T | G | T | - | - | - | - | - | - | G | - | - | - | - | - | - | - | - | - | C | A | G | T | T | C | A | - | - | - | A | C | T | C | T | T | G | A | T | A | C | G | A | G | T | C | A | T | G | A | A | T | A | G | T | A | T | T | C | T | C | T | G | T | G | A | T | G | T | C | T | C | T | C | G | A | T | T | T | C | T | T | - | - | - | - | - | - | - | - | - | A | A | A | C | G | G | T | G | #961285 |
| BA.1 | G | G | C | G | G | C | - | - | - | C | C | C | A | C | G | A | A | - | - | - | - | - | - | - | - | - | A | G | C | A | A | T | G | C | G | C | C | C | T | - | - | - | - | - | - | G | - | - | - | - | - | - | - | - | - | C | A | G | T | T | C | A | - | - | - | A | C | T | C | T | T | G | A | T | A | C | G | A | G | T | C | A | T | G | A | A | T | A | G | T | A | T | T | C | T | C | T | G | T | G | A | T | G | T | C | T | C | T | C | G | A | T | T | T | C | T | T | - | - | - | - | - | - | - | - | - | A | A | A | C | G | G | C | G |  |
| mut |  |  |  |  |  |  |  |  |  |  |  |  |  |  |  |  |  |  |  |  |  |  |  |  |  |  |  |  |  |  | A>G |  |  |  |  |  |  |  |  |  |  |  |  |  |  |  |  |  |  |  |  |  |  |  |  |  |  |  |  |  |  |  |  |  |  |  |  |  |  |  |  |  |  |  |  |  |  |  |  |  |  |  |  |  |  |  |  |  |  |  |  |  |  |  |  |  |  |  |  |  |  |  |  |  |  |  |  |  |  |  |  |  |  |  |  |  |  |  |  |  |  |  |  |  |  |  |  |  |  |  |  |  | C>T |  |  |

|  |  |  |  |  |  |  |  |  |  |  |  |  |  |  |  |  |  |  |  |  |  |  |  |  |  |  |  |  |  |  |  |  |  |  |  |  |  |  |  |  |  |  |  |  |  |  |  |  |  |  |  |  |  |  |  |  |  |  |  |  |  |  |  |  |  |  |  |  |  |  |  |  |  |  |  |  |  |  |  |  |  |  |  |  |  |
| --- | --- | --- | --- | --- | --- | --- | --- | --- | --- | --- | --- | --- | --- | --- | --- | --- | --- | --- | --- | --- | --- | --- | --- | --- | --- | --- | --- | --- | --- | --- | --- | --- | --- | --- | --- | --- | --- | --- | --- | --- | --- | --- | --- | --- | --- | --- | --- | --- | --- | --- | --- | --- | --- | --- | --- | --- | --- | --- | --- | --- | --- | --- | --- | --- | --- | --- | --- | --- | --- | --- | --- | --- | --- | --- | --- | --- | --- | --- | --- | --- | --- | --- | --- | --- | --- |
| pos | 670 | 2453 | 2790 | 2832 | 3241 | 4184 | 4321 | 5386 | 5924 | 6513 | 6514 | 6515 | 8393 | 9344 | 9424 | 9534 | 10198 | 10447 | 11283 | 11284 | 11285 | 11286 | 11287 | 11292 | 11293 | 11294 | 11295 | 11296 | 11537 | 12880 | 13195 | 15240 | 15714 | 17410 | 19955 | 20055 | 21618 | 21633 | 21634 | 21635 | 21636 | 21637 | 21638 | 21639 | 21640 | 21641 | 21762 | 21765 | 21766 | 21767 | 21768 | 21769 | 21770 | 21846 | 21987 | 21988 | 21989 | 21990 | 21991 | 21992 | 21993 | 21994 | 21995 | 22194 | 22195 | 22196 | 22200 | 22673 | 22688 | 22775 | 22786 | 22898 | 23048 | 23202 | 23664 | 24130 | 24503 | 26060 | 26530 | 26858 | 27382 | 27383 | 27384 | 29510 |  |
| ref | T | C | C | A | C | G | C | T | G | G | T | T | G | C | A | C | C | G | G | T | T | T | G | G | T | T | T | T | A | C | T | C | C | C | C | A | C | T | A | C | C | C | C | C | T | G | C | T | A | C | A | T | G | C | G | T | G | T | T | T | A | T | T | A | T | T | T | T | A | G | A | G | G | C | C | C | C | C | A | C | G | A | T | A |  |
| BA.2 | G | C | T | A | C | A | T | T | G | G | T | T | G | T | G | T | T | A | G | T | T | T | G | - | - | - | - | - | A | T | T | C | T | T | T | G | T | - | - | - | - | - | - | - | - | - | C | T | A | C | A | T | G | C | A | T | G | T | T | T | A | T | T | A | T | T | G | T | G | A | C | G | G | C | C | C | C | T | A | T | C | T | C | C |  |
| BA.2 | G | C | T | A | C | A | T | T | G | G | T | T | G | T | G | T | T | A | G | T | T | T | G | - | - | - | - | - | A | T | T | C | T | T | T | G | T | - | - | - | - | - | - | - | - | - | C | T | A | C | A | T | G | C | A | T | G | T | T | T | A | T | T | A | T | T | G | T | G | A | C | G | G | C | C | C | C | C | G | C | G | A | T | A | #964555 |
| BA.1.17.2 | T | T | C | G | T | G | C | G | A | - | - | - | A | C | A | C | C | G | - | - | - | - | - | G | T | T | T | T | G | C | C | T | C | C | C | A | C | T | A | C | C | C | C | C | T | G | T | - | - | - | - | - | - | T | - | - | - | - | - | - | - | - | - | - | - | - | T | C | A | G | A | A | A | A | T | A | T | C | G | C | G | A | T | A |  |
| mut |  |  |  |  |  |  |  |  |  |  |  |  |  |  |  |  |  |  |  |  |  |  |  |  |  |  |  |  |  |  |  |  |  |  |  |  |  |  |  |  |  |  |  |  |  |  |  |  |  |  |  |  |  |  |  |  |  |  |  |  |  |  |  |  |  |  |  |  |  |  |  |  |  |  |  |  |  |  |  |  |  |  |  |  |  |

|  |  |  |  |  |  |  |  |  |  |  |  |  |  |  |  |  |  |  |  |  |  |  |  |  |  |  |  |  |  |  |  |  |  |  |  |  |  |  |  |  |  |  |  |  |  |  |  |  |  |  |  |  |  |  |  |  |  |  |  |  |  |  |  |  |  |  |  |  |  |  |  |  |  |  |  |  |  |  |  |  |  |  |  |  |  |  |
| --- | --- | --- | --- | --- | --- | --- | --- | --- | --- | --- | --- | --- | --- | --- | --- | --- | --- | --- | --- | --- | --- | --- | --- | --- | --- | --- | --- | --- | --- | --- | --- | --- | --- | --- | --- | --- | --- | --- | --- | --- | --- | --- | --- | --- | --- | --- | --- | --- | --- | --- | --- | --- | --- | --- | --- | --- | --- | --- | --- | --- | --- | --- | --- | --- | --- | --- | --- | --- | --- | --- | --- | --- | --- | --- | --- | --- | --- | --- | --- | --- | --- | --- | --- | --- | --- | --- |
| pos | 670 | 2790 | 2832 | 3241 | 4184 | 4321 | 5386 | 5924 | 6513 | 6514 | 6515 | 8393 | 9344 | 9424 | 9534 | 10198 | 10447 | 11283 | 11284 | 11285 | 11286 | 11287 | 11292 | 11293 | 11294 | 11295 | 11296 | 11537 | 12880 | 13195 | 15240 | 15714 | 17410 | 19955 | 20055 | 21618 | 21633 | 21634 | 21635 | 21636 | 21637 | 21638 | 21639 | 21640 | 21641 | 21762 | 21765 | 21766 | 21767 | 21768 | 21769 | 21770 | 21846 | 21987 | 21988 | 21989 | 21990 | 21991 | 21992 | 21993 | 21994 | 21995 | 22194 | 22195 | 22196 | 22200 | 22224 | 22673 | 22688 | 22775 | 22786 | 22792 | 22813 | 23048 | 23202 | 23664 | 24130 | 24503 | 26060 | 26530 | 26858 | 27382 | 27383 | 27384 | 29510 |  |
| ref | T | C | A | C | G | C | T | G | G | T | T | G | C | A | C | C | G | G | T | T | T | G | G | T | T | T | T | A | C | T | C | C | C | C | A | C | T | A | C | C | C | C | C | T | G | C | T | A | C | A | T | G | C | G | T | G | T | T | T | A | T | T | A | T | T | T | C | T | A | G | A | C | G | G | C | C | C | C | C | A | C | G | A | T | A |  |
| BA.1.17.2 | T | C | G | T | G | C | G | A | - | - | - | A | C | A | C | C | G | - | - | - | - | - | G | T | T | T | T | G | C | C | T | C | C | C | A | C | T | A | C | C | C | C | C | T | G | T | - | - | - | - | - | - | T | - | - | - | - | - | - | - | - | - | - | - | - | T | T | C | A | G | A | C | G | A | A | T | A | T | C | G | C | G | A | T | A |  |
| BA.2 | T | C | G | T | G | C | G | A | - | - | - | A | C | A | C | C | G | G | T | T | T | G | - | - | - | - | - | A | T | T | C | T | T | T | G | T | - | - | - | - | - | - | - | - | - | C | T | A | C | A | T | G | C | A | T | G | T | T | T | A | T | T | A | T | T | G | C | T | G | A | C | T | T | G | C | C | C | C | T | A | T | C | T | C | C | #965353 |
| BA.2 | G | T | A | C | A | T | T | G | G | T | T | G | T | G | T | T | A | G | T | T | T | G | - | - | - | - | - | A | T | T | C | T | T | T | G | T | - | - | - | - | - | - | - | - | - | C | T | A | C | A | T | G | C | A | T | G | T | T | T | A | T | T | A | T | T | G | C | T | G | A | C | C | T | G | C | C | C | C | T | A | T | C | T | C | C |  |
| mut |  |  |  |  |  |  |  |  |  |  |  |  |  |  |  |  |  |  |  |  |  |  |  |  |  |  |  |  |  |  |  |  |  |  |  |  |  |  |  |  |  |  |  |  |  |  |  |  |  |  |  |  |  |  |  |  |  |  |  |  |  |  |  |  |  |  |  |  |  |  |  | C>T |  |  |  |  |  |  |  |  |  |  |  |  |  |  |

|  |  |  |  |  |  |  |  |  |  |  |  |  |  |  |  |  |  |  |  |  |  |  |  |  |  |  |  |  |  |  |  |  |  |  |  |  |  |  |  |  |  |  |  |  |  |  |  |  |  |  |  |  |  |  |  |  |  |  |  |  |  |  |  |  |  |  |  |  |  |  |  |  |  |  |  |  |  |  |  |  |  |  |  |  |  |  |
| --- | --- | --- | --- | --- | --- | --- | --- | --- | --- | --- | --- | --- | --- | --- | --- | --- | --- | --- | --- | --- | --- | --- | --- | --- | --- | --- | --- | --- | --- | --- | --- | --- | --- | --- | --- | --- | --- | --- | --- | --- | --- | --- | --- | --- | --- | --- | --- | --- | --- | --- | --- | --- | --- | --- | --- | --- | --- | --- | --- | --- | --- | --- | --- | --- | --- | --- | --- | --- | --- | --- | --- | --- | --- | --- | --- | --- | --- | --- | --- | --- | --- | --- | --- | --- | --- | --- |
| pos | 670 | 2790 | 2832 | 3241 | 4184 | 4321 | 5386 | 5924 | 6513 | 6514 | 6515 | 8393 | 9344 | 9424 | 9534 | 10198 | 10447 | 11283 | 11284 | 11285 | 11286 | 11287 | 11292 | 11293 | 11294 | 11295 | 11296 | 11537 | 12880 | 13195 | 15240 | 17410 | 18747 | 19955 | 20055 | 21597 | 21618 | 21633 | 21634 | 21635 | 21636 | 21637 | 21638 | 21639 | 21640 | 21641 | 21762 | 21765 | 21766 | 21767 | 21768 | 21769 | 21770 | 21846 | 21987 | 21988 | 21989 | 21990 | 21991 | 21992 | 21993 | 21994 | 21995 | 22194 | 22195 | 22196 | 22200 | 22673 | 22688 | 22775 | 22786 | 22898 | 23048 | 23202 | 23664 | 24130 | 24503 | 26060 | 26530 | 26858 | 26978 | 27382 | 27383 | 27384 | 29510 |  |
| ref | T | C | A | C | G | C | T | G | G | T | T | G | C | A | C | C | G | G | T | T | T | G | G | T | T | T | T | A | C | T | C | C | C | C | A | C | C | T | A | C | C | C | C | C | T | G | C | T | A | C | A | T | G | C | G | T | G | T | T | T | A | T | T | A | T | T | T | T | A | G | A | G | G | C | C | C | C | C | A | C | T | G | A | T | A |  |
| BA.1.17.2 | T | C | G | T | G | C | G | A | - | - | - | A | C | A | C | C | G | - | - | - | - | - | G | T | T | T | T | G | C | C | T | C | T | C | A | T | C | T | A | C | C | C | C | C | T | G | T | - | - | - | - | - | - | T | - | - | - | - | - | - | - | - | - | - | - | - | T | C | A | G | A | A | A | A | T | A | T | C | G | C | C | G | A | T | A |  |
| BA.2 | T | C | G | T | G | C | G | A | - | - | - | A | C | A | C | C | G | - | - | - | - | - | G | T | T | T | T | G | C | C | C | T | C | T | G | C | T | - | - | - | - | - | - | - | - | - | C | T | A | C | A | T | G | C | A | T | G | T | T | T | A | T | T | A | T | T | G | T | G | A | C | G | G | C | C | C | C | T | A | T | T | C | T | C | C | #966905 |
| BA.2 | G | T | A | C | A | T | T | G | G | T | T | G | T | G | T | T | A | G | T | T | T | G | - | - | - | - | - | A | T | T | C | T | C | T | G | C | T | - | - | - | - | - | - | - | - | - | C | T | A | C | A | T | G | C | A | T | G | T | T | T | A | T | T | A | T | T | G | T | G | A | C | G | G | C | C | C | C | T | A | T | T | C | T | C | C |  |
| mut |  |  |  |  |  |  |  |  |  |  |  |  |  |  |  |  |  |  |  |  |  |  |  |  |  |  |  |  |  |  | T>C |  |  |  |  |  |  |  |  |  |  |  |  |  |  |  |  |  |  |  |  |  |  |  |  |  |  |  |  |  |  |  |  |  |  |  |  |  |  |  |  |  |  |  |  |  |  |  |  |  |  |  |  |  |  |  |

|  |  |  |  |  |  |  |  |  |  |  |  |  |  |  |  |  |  |  |  |  |  |  |  |  |  |  |  |  |  |  |  |  |  |  |  |  |  |  |  |  |  |  |  |  |  |  |  |  |  |  |  |  |  |  |  |  |  |  |  |  |  |  |  |  |  |  |  |  |  |  |  |  |  |  |  |  |  |  |  |  |  |  |  |  |  |  |  |  |  |  |  |  |  |  |  |  |  |  |  |  |  |  |  |  |  |  |  |  |  |  |  |  |  |  |  |  |  |  |  |  |  |  |  |  |  |  |  |  |  |  |  |  |  |  |  |  |  |  |  |  |  |  |  |  |  |  |  |  |  |
| --- | --- | --- | --- | --- | --- | --- | --- | --- | --- | --- | --- | --- | --- | --- | --- | --- | --- | --- | --- | --- | --- | --- | --- | --- | --- | --- | --- | --- | --- | --- | --- | --- | --- | --- | --- | --- | --- | --- | --- | --- | --- | --- | --- | --- | --- | --- | --- | --- | --- | --- | --- | --- | --- | --- | --- | --- | --- | --- | --- | --- | --- | --- | --- | --- | --- | --- | --- | --- | --- | --- | --- | --- | --- | --- | --- | --- | --- | --- | --- | --- | --- | --- | --- | --- | --- | --- | --- | --- | --- | --- | --- | --- | --- | --- | --- | --- | --- | --- | --- | --- | --- | --- | --- | --- | --- | --- | --- | --- | --- | --- | --- | --- | --- | --- | --- | --- | --- | --- | --- | --- | --- | --- | --- | --- | --- | --- | --- | --- | --- | --- | --- | --- | --- | --- | --- | --- | --- | --- | --- | --- | --- | --- | --- | --- | --- | --- | --- | --- | --- |
| pos | 210 | 313 | 507 | 508 | 509 | 1684 | 1962 | 2470 | 2725 | 2832 | 2883 | 3984 | 4582 | 5164 | 5184 | 5386 | 5584 | 6513 | 6514 | 6515 | 7393 | 8393 | 8591 | 9891 | 10029 | 10449 | 11283 | 11284 | 11285 | 11286 | 11287 | 11288 | 11289 | 11290 | 11291 | 11418 | 11514 | 11537 | 12049 | 13019 | 13195 | 13366 | 14857 | 15240 | 15451 | 16466 | 18163 | 19101 | 20464 | 21618 | 21762 | 21765 | 21766 | 21767 | 21768 | 21769 | 21770 | 21846 | 21987 | 21988 | 21989 | 21990 | 21991 | 21992 | 21993 | 21994 | 21995 | 22029 | 22030 | 22031 | 22032 | 22033 | 22034 | 22195 | 22197 | 22198 | 22202 | 22204 | 22227 | 22427 | 22578 | 22599 | 22673 | 22674 | 22679 | 22686 | 22813 | 22879 | 22882 | 22898 | 22917 | 22992 | 23013 | 23040 | 23048 | 23055 | 23063 | 23075 | 23202 | 23525 | 23599 | 23604 | 23707 | 23854 | 23948 | 24130 | 24410 | 24424 | 24469 | 24503 | 25000 | 25455 | 25469 | 25584 | 26270 | 26530 | 26577 | 26709 | 26767 | 27259 | 27505 | 27638 | 27752 | 27807 | 28248 | 28249 | 28250 | 28251 | 28252 | 28253 | 28271 | 28311 | 28326 | 28362 | 28363 | 28364 | 28365 | 28366 | 28367 | 28368 | 28369 | 28370 | 28461 | 28881 | 28882 | 28883 | 29402 | 29742 |  |
| ref | G | C | A | T | G | C | C | C | A | A | G | C | C | G | C | T | A | G | T | T | G | G | G | C | C | C | G | T | T | T | G | T | C | T | G | T | C | A | C | C | T | C | G | C | G | C | A | G | G | C | C | T | A | C | A | T | G | C | G | T | G | T | T | T | A | T | T | A | G | T | T | C | A | T | T | A | C | T | C | G | G | G | T | C | T | C | G | C | T | G | T | G | A | A | G | A | A | T | C | C | T | C | C | C | G | C | G | A | T | C | C | G | C | C | C | A | C | G | T | A | G | T | C | C | G | A | T | T | T | C | A | C | G | G | A | G | A | A | C | G | C | A | A | G | G | G | G | G |  |
| BA.1.1 | G | C | T | C | A | C | C | T | A | G | G | C | C | G | C | G | A | - | - | - | G | A | G | C | T | A | - | - | - | - | - | - | - | - | - | T | C | G | C | C | C | T | G | T | G | C | G | A | G | C | T | - | - | - | - | - | - | T | - | - | - | - | - | - | - | - | - | A | G | T | T | C | A | G | G | C | A | C | C | G | A | A | C | T | C | T | T | T | G | A | T | A | C | G | A | G | T | C | A | T | G | A | C | A | T | A | G | T | A | T | T | G | C | T | T | G | G | A | T | C | T | T | C | T | G | A | T | T | T | C | T | T | G | - | - | - | - | - | - | - | - | - | A | A | A | C | G | G |  |
| AY.47 | G | C | T | C | A | C | C | T | G | G | G | C | C | G | C | G | A | G | T | T | A | G | A | T | C | C | G | T | T | T | G | T | C | T | G | C | T | A | T | T | T | C | T | C | A | T | A | G | T | G | C | T | A | C | A | T | G | C | A | T | G | T | T | T | A | T | T | - | - | - | - | - | - | T | T | A | C | T | T | A | G | G | T | C | T | C | G | C | T | G | G | G | A | A | G | A | A | T | C | C | T | G | T | C | G | C | A | A | T | C | C | T | T | C | C | A | C | G | C | A | G | C | T | C | - | - | - | - | - | - | - | C | T | G | A | G | A | A | C | G | C | A | G | T | G | G | T | T | #967112 |
| AY.47 | T | T | A | T | G | T | T | C | A | A | A | T | T | T | T | T | G | G | T | T | A | G | G | T | C | C | G | T | T | T | G | T | C | T | G | C | T | A | T | T | T | C | T | C | A | T | A | G | G | G | C | T | A | C | A | T | G | C | A | T | G | T | T | T | A | T | T | - | - | - | - | - | - | T | T | A | C | T | T | A | G | G | T | C | T | C | G | C | T | G | G | G | A | A | G | A | A | T | C | C | T | G | T | C | G | C | A | A | T | C | C | T | T | C | C | A | C | G | C | A | G | C | T | C | - | - | - | - | - | - | - | C | T | G | A | G | A | A | C | G | C | A | G | T | G | G | T | T |  |
| mut |  |  |  |  |  |  |  |  | A>G |  |  |  |  |  |  |  |  |  |  |  |  |  | G>A |  |  |  |  |  |  |  |  |  |  |  |  |  |  |  |  |  |  |  |  |  |  |  |  |  | G>T |  |  |  |  |  |  |  |  |  |  |  |  |  |  |  |  |  |  |  |  |  |  |  |  |  |  |  |  |  |  |  |  |  |  |  |  |  |  |  |  |  |  |  |  |  |  |  |  |  |  |  |  |  |  |  |  |  |  |  |  |  |  |  |  |  |  |  |  |  |  |  |  |  |  |  |  |  |  |  |  |  |  |  |  |  |  |  |  |  |  |  |  |  |  |  |  |  |  |  |  |

|  |  |  |  |  |  |  |  |  |  |  |  |  |  |  |  |  |  |
| --- | --- | --- | --- | --- | --- | --- | --- | --- | --- | --- | --- | --- | --- | --- | --- | --- | --- |
| pos | 1185 | 2346 | 6198 | 9988 | 18183 | 18508 | 21304 | 21952 | 22194 | 22195 | 22196 | 22197 | 22198 | 22202 | 22204 | 23432 |  |
| ref | C | T | C | C | C | C | C | C | A | T | T | T | A | C | T | A |  |
| BA.1 | C | C | C | T | T | T | A | C | - | - | - | T | A | C | T | C |  |
| BA.1 | T | C | T | T | T | T | C | T | A | G | T | G | C | A | C | A | #967397 |
| BA.1 | C | T | C | C | C | C | C | C | A | G | T | G | C | A | C | A |  |
| mut | C>T |  | C>T |  |  |  |  | C>T |  |  |  |  |  |  |  |  |  |

|  |  |  |  |  |  |  |  |  |  |  |  |  |  |  |  |  |  |  |  |
| --- | --- | --- | --- | --- | --- | --- | --- | --- | --- | --- | --- | --- | --- | --- | --- | --- | --- | --- | --- |
| pos | 1612 | 2470 | 5515 | 7926 | 8730 | 16178 | 22194 | 22195 | 22196 | 22197 | 22198 | 22202 | 22204 | 22599 | 27382 | 27383 | 27384 | 29374 |  |
| ref | C | C | G | C | C | C | A | T | T | T | A | C | T | G | G | A | T | G |  |
| BA.1 | C | C | T | C | T | T | - | - | - | T | A | C | T | G | G | A | T | G |  |
| BA.1 | T | C | T | C | T | T | A | G | T | G | C | A | C | G | C | T | C | A | #967732 |
| BA.1.1 | C | T | G | T | C | C | A | G | T | G | C | A | C | A | C | T | C | G |  |
| mut | C>T |  |  |  |  |  |  |  |  |  |  |  |  | A>G |  |  |  | G>A |  |

|  |  |  |  |  |  |  |  |  |  |  |  |  |
| --- | --- | --- | --- | --- | --- | --- | --- | --- | --- | --- | --- | --- |
| pos | 2470 | 2902 | 3041 | 7561 | 22599 | 26139 | 26485 | 26488 | 26491 | 26492 | 26497 |  |
| ref | C | C | C | G | G | C | T | T | T | A | T |  |
| BA.1 | C | T | C | A | G | T | T | T | T | A | T |  |
| BA.1 | C | T | T | A | G | T | T | G | C | T | C | #974002 |
| BA.1.1 | T | C | C | G | A | C | G | G | C | T | C |  |
| mut |  |  | C>T |  |  |  |  |  |  |  |  |  |

|  |  |  |  |  |  |  |  |  |  |  |  |  |  |  |  |  |
| --- | --- | --- | --- | --- | --- | --- | --- | --- | --- | --- | --- | --- | --- | --- | --- | --- |
| pos | 1577 | 2470 | 3241 | 3936 | 5924 | 20480 | 22599 | 22813 | 22898 | 23664 | 26522 | 29571 | 29572 | 29573 | 29575 |  |
| ref | G | C | C | A | G | C | G | G | G | C | C | A | C | G | T |  |
| BA.1.1 | G | T | C | T | G | C | A | T | A | C | T | A | C | G | T |  |
| BA.1.1 | A | T | C | T | G | C | A | T | A | C | T | T | T | A | A | #979290 |
| BA.1.17.2 | G | C | T | A | A | T | G | G | G | T | C | T | T | A | A |  |
| mut | G>A |  |  |  |  |  |  |  |  |  |  |  |  |  |  |  |

|  |  |  |  |  |  |  |  |  |  |  |  |  |  |  |  |  |  |  |  |  |  |  |  |  |  |  |  |  |  |  |  |  |  |  |  |  |  |  |  |  |  |  |  |  |  |  |  |  |  |  |  |  |  |  |  |  |  |  |  |  |  |  |  |  |  |  |  |  |  |  |  |  |  |  |  |  |  |  |  |  |  |  |  |  |  |  |  |  |  |  |  |  |  |  |  |  |  |  |  |  |  |  |  |  |  |  |  |  |  |  |  |  |  |  |  |  |  |  |  |  |  |  |  |  |  |  |  |  |  |  |  |  |  |  |  |  |  |  |  |  |
| --- | --- | --- | --- | --- | --- | --- | --- | --- | --- | --- | --- | --- | --- | --- | --- | --- | --- | --- | --- | --- | --- | --- | --- | --- | --- | --- | --- | --- | --- | --- | --- | --- | --- | --- | --- | --- | --- | --- | --- | --- | --- | --- | --- | --- | --- | --- | --- | --- | --- | --- | --- | --- | --- | --- | --- | --- | --- | --- | --- | --- | --- | --- | --- | --- | --- | --- | --- | --- | --- | --- | --- | --- | --- | --- | --- | --- | --- | --- | --- | --- | --- | --- | --- | --- | --- | --- | --- | --- | --- | --- | --- | --- | --- | --- | --- | --- | --- | --- | --- | --- | --- | --- | --- | --- | --- | --- | --- | --- | --- | --- | --- | --- | --- | --- | --- | --- | --- | --- | --- | --- | --- | --- | --- | --- | --- | --- | --- | --- | --- | --- | --- | --- | --- | --- | --- | --- | --- | --- | --- | --- |
| pos | 210 | 686 | 687 | 689 | 691 | 692 | 694 | 2668 | 2832 | 4181 | 5386 | 5526 | 6402 | 6513 | 6514 | 6515 | 7124 | 8393 | 8986 | 9053 | 10449 | 11201 | 11283 | 11284 | 11285 | 11286 | 11287 | 11288 | 11289 | 11290 | 11291 | 11332 | 11537 | 13195 | 15240 | 15451 | 16466 | 17014 | 18163 | 19220 | 21618 | 21762 | 21765 | 21766 | 21767 | 21768 | 21769 | 21770 | 21846 | 21987 | 21988 | 21989 | 21990 | 21991 | 21992 | 21993 | 21994 | 21995 | 22029 | 22030 | 22031 | 22032 | 22033 | 22034 | 22194 | 22195 | 22196 | 22578 | 22673 | 22674 | 22679 | 22686 | 22813 | 22882 | 22898 | 22917 | 22992 | 23013 | 23040 | 23048 | 23055 | 23063 | 23075 | 23202 | 23525 | 23599 | 23604 | 23854 | 23948 | 24130 | 24410 | 24424 | 24469 | 24503 | 24928 | 25000 | 25273 | 25469 | 25584 | 26010 | 26270 | 26530 | 26577 | 26709 | 26767 | 27259 | 27408 | 27610 | 27635 | 27638 | 27752 | 27807 | 27874 | 28248 | 28249 | 28250 | 28251 | 28252 | 28253 | 28271 | 28311 | 28362 | 28363 | 28364 | 28365 | 28366 | 28367 | 28368 | 28369 | 28370 | 28461 | 28881 | 28882 | 28883 | 28916 | 29253 | 29402 | 29632 | 29742 |  |
| ref | G | A | A | T | A | T | T | C | A | G | T | C | C | G | T | T | C | G | C | G | C | A | G | T | T | T | G | T | C | T | G | A | A | T | C | G | C | G | A | C | C | C | T | A | C | A | T | G | C | G | T | G | T | T | T | A | T | T | A | G | T | T | C | A | A | T | T | G | T | C | T | C | G | T | G | T | G | A | A | G | A | A | T | C | C | T | C | C | G | C | G | A | T | C | G | C | G | C | C | C | C | A | C | G | T | A | T | C | C | T | C | C | C | G | A | T | T | T | C | A | C | G | A | G | A | A | C | G | C | A | A | G | G | G | G | C | G | C | G |  |
| BA.1 | G | G | C | C | G | C | A | C | G | G | G | C | C | - | - | - | C | A | C | G | A | A | - | - | - | - | - | - | - | - | - | A | G | C | T | G | C | G | G | C | C | T | - | - | - | - | - | - | T | - | - | - | - | - | - | - | - | - | A | G | T | T | C | A | - | - | - | A | C | T | C | T | T | G | A | T | A | C | G | A | G | T | C | A | T | G | A | A | T | A | G | T | A | T | G | T | G | C | T | C | T | G | G | A | T | C | T | C | C | T | C | T | C | G | A | T | T | T | C | T | T | - | - | - | - | - | - | - | - | - | A | A | A | C | G | C | G | T | G |  |
| AY.5 | T | G | C | C | G | C | A | T | A | T | T | A | T | G | T | T | T | G | T | T | C | G | G | T | T | T | G | T | C | T | G | G | A | T | C | A | T | T | A | T | G | C | T | A | C | A | T | G | C | A | T | G | T | T | T | A | T | T | - | - | - | - | - | - | A | T | T | G | T | C | T | C | G | T | G | G | G | A | A | G | A | A | T | C | C | T | G | C | G | C | A | A | T | C | T | C | T | T | C | T | C | A | C | G | C | A | A | T | T | C | T | C | T | - | - | - | - | - | - | - | C | G | A | G | A | A | C | G | C | A | G | T | G | G | T | T | T | C | T | #982641 |
| AY.5 | T | A | A | T | A | T | T | T | A | T | T | A | T | G | T | T | T | G | T | T | C | G | G | T | T | T | G | T | C | T | G | G | A | T | C | A | T | T | A | T | G | C | T | A | C | A | T | G | C | A | T | G | T | T | T | A | T | T | - | - | - | - | - | - | A | T | T | G | T | C | T | C | G | T | G | G | G | A | A | G | A | A | T | C | C | T | G | C | G | C | A | A | T | C | T | C | T | T | C | C | C | A | C | G | C | A | A | T | T | C | T | C | T | - | - | - | - | - | - | - | C | G | A | G | A | A | C | G | C | A | G | T | G | G | T | T | T | C | T |  |
| mut | G>T |  |  |  |  |  |  |  |  |  |  |  |  |  |  |  |  |  |  |  |  |  |  |  |  |  |  |  |  |  |  |  |  |  |  |  |  |  |  |  |  |  |  |  |  |  |  |  |  |  |  |  |  |  |  |  |  |  |  |  |  |  |  |  |  |  |  |  |  |  |  |  |  |  |  |  |  |  |  |  |  |  |  |  |  |  |  |  |  |  |  |  |  |  |  |  |  |  |  | C>T |  |  |  |  |  |  |  |  |  |  |  |  |  |  |  |  |  |  |  |  |  |  |  |  |  |  |  |  |  |  |  |  |  |  |  |  |  |  |  |  |

|  |  |  |  |  |  |  |  |  |  |  |  |  |  |  |  |  |  |  |  |
| --- | --- | --- | --- | --- | --- | --- | --- | --- | --- | --- | --- | --- | --- | --- | --- | --- | --- | --- | --- |
| pos | 1989 | 2470 | 3241 | 4579 | 5924 | 11782 | 13234 | 17241 | 22599 | 22879 | 23664 | 25047 | 27377 | 27378 | 27379 | 27382 | 27383 | 27384 |  |
| ref | A | C | C | T | G | A | T | T | G | C | C | C | A | G | A | G | A | T |  |
| BA.1.1 | G | T | C | A | G | G | C | T | A | T | C | C | A | G | A | G | A | T |  |
| BA.1.1 | G | T | C | A | G | G | C | T | A | T | C | C | C | T | G | C | T | C | #983066 |
| BA.1.17.2 | A | C | T | T | A | A | T | C | G | C | T | T | C | T | G | C | T | C |  |
| mut |  |  |  |  |  |  |  |  |  |  |  |  |  |  |  |  |  |  |  |

|  |  |  |  |  |  |  |  |  |  |  |  |  |  |  |  |  |  |  |  |  |  |  |  |  |  |  |  |  |  |
| --- | --- | --- | --- | --- | --- | --- | --- | --- | --- | --- | --- | --- | --- | --- | --- | --- | --- | --- | --- | --- | --- | --- | --- | --- | --- | --- | --- | --- | --- |
| pos | 7926 | 21595 | 22029 | 22030 | 22031 | 22032 | 22033 | 22034 | 22194 | 22195 | 22196 | 22578 | 22599 | 22673 | 22674 | 22679 | 22686 | 22898 | 22917 | 23013 | 23040 | 23048 | 23055 | 23063 | 23075 | 25311 | 27874 | 28073 |  |
| ref | C | C | A | G | T | T | C | A | A | T | T | G | G | T | C | T | C | G | T | A | A | G | A | A | T | G | C | G |  |
| BA.1.1.18 | T | C | A | G | T | T | C | A | - | - | - | A | A | C | T | C | T | G | G | C | G | A | G | T | C | G | C | G |  |
| BA.1.1.18 | T | C | A | G | T | T | C | A | - | - | - | A | A | C | T | C | T | G | G | C | A | G | A | A | T | T | T | T | #990337 |
| BA.1 | C | T | - | - | - | - | - | - | A | T | T | G | G | T | C | T | C | A | T | A | A | G | A | A | T | G | T | T |  |
| mut |  |  |  |  |  |  |  |  |  |  |  |  |  |  |  |  |  |  |  |  |  |  |  |  |  | G>T |  |  |  |

|  |  |  |  |  |  |  |  |  |  |  |  |
| --- | --- | --- | --- | --- | --- | --- | --- | --- | --- | --- | --- |
| pos | 710 | 4013 | 5924 | 7754 | 10416 | 11473 | 16809 | 17999 | 23782 | 29632 |  |
| ref | C | A | G | A | C | T | T | C | G | C |  |
| BA.1 | T | A | G | A | C | C | T | C | G | T |  |
| BA.1 | T | A | G | A | T | C | C | T | T | C | #995433 |
| BA.1.17 | C | G | A | G | C | T | C | T | T | C |  |
| mut |  |  |  |  | C>T |  |  |  |  |  |  |

|  |  |  |  |  |  |  |  |  |  |  |  |  |  |  |  |  |  |  |  |  |  |  |  |  |  |  |  |  |  |  |  |  |  |  |  |  |  |  |  |  |  |  |  |  |  |  |  |  |  |  |  |  |  |  |  |  |  |  |  |  |  |  |  |  |  |  |  |  |  |  |  |  |  |  |  |  |  |  |  |  |  |  |  |  |  |  |  |
| --- | --- | --- | --- | --- | --- | --- | --- | --- | --- | --- | --- | --- | --- | --- | --- | --- | --- | --- | --- | --- | --- | --- | --- | --- | --- | --- | --- | --- | --- | --- | --- | --- | --- | --- | --- | --- | --- | --- | --- | --- | --- | --- | --- | --- | --- | --- | --- | --- | --- | --- | --- | --- | --- | --- | --- | --- | --- | --- | --- | --- | --- | --- | --- | --- | --- | --- | --- | --- | --- | --- | --- | --- | --- | --- | --- | --- | --- | --- | --- | --- | --- | --- | --- | --- | --- | --- | --- |
| pos | 44 | 670 | 1072 | 2790 | 2832 | 4184 | 4321 | 5386 | 6513 | 6514 | 6515 | 8393 | 9344 | 9424 | 9534 | 9866 | 10198 | 10447 | 11152 | 11283 | 11284 | 11285 | 11286 | 11287 | 11292 | 11293 | 11294 | 11295 | 11296 | 11537 | 12880 | 13195 | 15240 | 15714 | 16342 | 17410 | 19955 | 20055 | 21618 | 21633 | 21634 | 21635 | 21636 | 21637 | 21638 | 21639 | 21640 | 21641 | 21762 | 21765 | 21766 | 21767 | 21768 | 21769 | 21770 | 21846 | 21987 | 21988 | 21989 | 21990 | 21991 | 21992 | 21993 | 21994 | 21995 | 22194 | 22195 | 22196 | 22200 | 22673 | 22688 | 22775 | 22786 | 22898 | 23048 | 23202 | 24130 | 24503 | 26060 | 26530 | 26858 | 27382 | 27383 | 27384 | 29287 | 29510 |  |
| ref | C | T | A | C | A | G | C | T | G | T | T | G | C | A | C | C | C | G | C | G | T | T | T | G | G | T | T | T | T | A | C | T | C | C | T | C | C | A | C | T | A | C | C | C | C | C | T | G | C | T | A | C | A | T | G | C | G | T | G | T | T | T | A | T | T | A | T | T | T | T | A | G | A | G | G | C | C | C | C | A | C | G | A | T | A | A |  |
| BA.1 | C | T | G | C | G | G | C | G | - | - | - | A | C | A | C | C | C | G | C | - | - | - | - | - | G | T | T | T | T | G | C | C | T | C | T | C | C | A | C | T | A | C | C | C | C | C | T | G | T | - | - | - | - | - | - | T | - | - | - | - | - | - | - | - | - | - | - | - | T | C | A | G | A | A | A | A | A | T | C | G | C | G | A | T | G | A |  |
| BA.2.10 | C | T | G | C | G | G | T | T | G | T | T | G | T | G | T | T | T | A | A | G | T | T | T | G | - | - | - | - | - | A | T | T | C | T | C | T | T | G | T | - | - | - | - | - | - | - | - | - | C | T | A | C | A | T | G | C | A | T | G | T | T | T | A | T | T | A | T | T | G | T | G | A | C | G | G | C | C | C | T | A | T | C | T | C | A | C | #995727 |
| BA.2.10 | T | G | A | T | A | A | T | T | G | T | T | G | T | G | T | T | T | A | A | G | T | T | T | G | - | - | - | - | - | A | T | T | C | T | C | T | T | G | T | - | - | - | - | - | - | - | - | - | C | T | A | C | A | T | G | C | A | T | G | T | T | T | A | T | T | A | T | T | G | T | G | A | C | G | G | C | C | C | T | A | T | C | T | C | A | C |  |
| mut |  |  |  |  |  |  |  |  |  |  |  |  |  |  |  |  |  |  |  |  |  |  |  |  |  |  |  |  |  |  |  |  |  |  |  |  |  |  |  |  |  |  |  |  |  |  |  |  |  |  |  |  |  |  |  |  |  |  |  |  |  |  |  |  |  |  |  |  |  |  |  |  |  |  |  |  |  |  |  |  |  |  |  |  |  |  |  |

|  |  |  |  |  |  |  |  |  |  |  |  |  |  |  |  |  |
| --- | --- | --- | --- | --- | --- | --- | --- | --- | --- | --- | --- | --- | --- | --- | --- | --- |
| pos | 9120 | 10396 | 13575 | 17746 | 19559 | 21595 | 21646 | 22194 | 22195 | 22196 | 22197 | 22198 | 22202 | 22204 | 22225 |  |
| ref | C | G | T | C | G | C | C | A | T | T | T | A | C | T | G |  |
| BA.1.1 | T | T | C | C | T | T | T | A | G | T | G | C | A | C | G |  |
| BA.1.1 | T | T | C | T | T | T | C | - | - | - | T | A | C | T | A | #997085 |
| BA.1.1 | C | G | T | C | G | C | C | - | - | - | T | A | C | T | A |  |
| mut |  |  |  | C>T |  |  |  |  |  |  |  |  |  |  |  |  |

|  |  |  |  |  |  |  |  |  |  |  |  |  |  |  |  |  |  |  |  |  |  |  |  |  |  |  |  |  |  |  |  |  |  |  |  |  |  |  |  |  |  |  |  |  |  |  |  |  |  |  |  |  |  |  |  |  |  |  |  |  |  |  |  |  |  |  |  |  |  |  |  |  |  |  |  |  |  |  |  |  |  |  |  |  |  |  |  |  |  |  |  |  |  |  |  |  |  |  |  |  |  |  |  |  |  |  |  |  |  |  |  |  |  |  |  |  |  |  |  |  |  |  |  |  |  |  |  |  |  |  |  |  |  |  |  |  |  |  |  |  |  |  |
| --- | --- | --- | --- | --- | --- | --- | --- | --- | --- | --- | --- | --- | --- | --- | --- | --- | --- | --- | --- | --- | --- | --- | --- | --- | --- | --- | --- | --- | --- | --- | --- | --- | --- | --- | --- | --- | --- | --- | --- | --- | --- | --- | --- | --- | --- | --- | --- | --- | --- | --- | --- | --- | --- | --- | --- | --- | --- | --- | --- | --- | --- | --- | --- | --- | --- | --- | --- | --- | --- | --- | --- | --- | --- | --- | --- | --- | --- | --- | --- | --- | --- | --- | --- | --- | --- | --- | --- | --- | --- | --- | --- | --- | --- | --- | --- | --- | --- | --- | --- | --- | --- | --- | --- | --- | --- | --- | --- | --- | --- | --- | --- | --- | --- | --- | --- | --- | --- | --- | --- | --- | --- | --- | --- | --- | --- | --- | --- | --- | --- | --- | --- | --- | --- | --- | --- | --- | --- | --- | --- | --- | --- | --- |
| pos | 210 | 1549 | 2003 | 2106 | 2470 | 2832 | 4181 | 5365 | 5386 | 6196 | 6402 | 6513 | 6514 | 6515 | 7124 | 8393 | 8986 | 9053 | 10449 | 11201 | 11283 | 11284 | 11285 | 11286 | 11287 | 11288 | 11289 | 11290 | 11291 | 11332 | 11537 | 13195 | 15240 | 15451 | 16466 | 16692 | 17236 | 18163 | 19220 | 21301 | 21595 | 21618 | 21762 | 21765 | 21766 | 21767 | 21768 | 21769 | 21770 | 21800 | 21846 | 21987 | 21988 | 21989 | 21990 | 21991 | 21992 | 21993 | 21994 | 21995 | 22029 | 22030 | 22031 | 22032 | 22033 | 22034 | 22195 | 22197 | 22198 | 22202 | 22204 | 22578 | 22599 | 22673 | 22674 | 22679 | 22686 | 22813 | 22882 | 22898 | 22917 | 22992 | 23013 | 23040 | 23048 | 23055 | 23063 | 23075 | 23202 | 23525 | 23599 | 23604 | 23854 | 23948 | 24130 | 24208 | 24410 | 24424 | 24469 | 24503 | 25000 | 25469 | 25584 | 26270 | 26530 | 26577 | 26709 | 26767 | 27259 | 27513 | 27638 | 27752 | 27807 | 27874 | 28248 | 28249 | 28250 | 28251 | 28252 | 28253 | 28271 | 28311 | 28362 | 28363 | 28364 | 28365 | 28366 | 28367 | 28368 | 28369 | 28370 | 28461 | 28541 | 28881 | 28882 | 28883 | 28916 | 29095 | 29402 | 29541 | 29742 |  |
| ref | G | C | C | C | C | A | G | C | T | C | C | G | T | T | C | G | C | G | C | A | G | T | T | T | G | T | C | T | G | A | A | T | C | G | C | T | A | A | C | C | C | C | C | T | A | C | A | T | G | G | C | G | T | G | T | T | T | A | T | T | A | G | T | T | C | A | T | T | A | C | T | G | G | T | C | T | C | G | T | G | T | G | A | A | G | A | A | T | C | C | T | C | C | G | C | C | G | A | T | C | C | C | C | C | A | C | G | T | A | C | T | C | C | C | G | A | T | T | T | C | A | C | G | A | G | A | A | C | G | C | A | A | G | G | G | G | G | C | G | C | G |  |
| AY.103 | T | C | C | C | C | A | T | T | T | T | T | G | T | T | T | G | T | T | C | G | G | T | T | T | G | T | C | T | G | G | A | T | C | A | T | C | G | A | T | T | C | G | C | T | A | C | A | T | G | T | C | A | T | G | T | T | T | A | T | T | - | - | - | - | - | - | T | T | A | C | T | G | G | T | C | T | C | G | T | G | G | G | A | A | G | A | A | T | C | C | T | G | C | G | C | T | A | A | T | C | C | T | C | C | A | C | G | C | A | T | C | T | C | T | - | - | - | - | - | - | - | C | G | A | G | A | A | C | G | C | A | G | T | T | G | G | T | T | T | T | T |  |
| XS | T | C | C | C | C | A | T | T | T | T | T | G | T | T | T | G | T | T | A | A | - | - | - | - | - | - | - | - | - | A | G | C | T | G | C | T | A | G | C | C | T | C | T | - | - | - | - | - | - | G | T | - | - | - | - | - | - | - | - | - | A | G | T | T | C | A | G | G | C | A | C | A | A | C | T | C | T | T | G | A | T | A | C | G | A | G | T | C | A | T | G | A | A | T | A | C | G | T | A | T | T | C | T | T | G | G | A | T | C | C | T | C | T | C | G | A | T | T | T | C | T | T | - | - | - | - | - | - | - | - | - | A | G | A | A | C | G | C | G | C | G | #1000242 |
| BA.1.1 | G | T | T | T | T | G | G | C | G | C | C | - | - | - | C | A | C | G | A | A | - | - | - | - | - | - | - | - | - | A | G | C | T | G | C | T | A | G | C | C | C | C | T | - | - | - | - | - | - | G | T | - | - | - | - | - | - | - | - | - | A | G | T | T | C | A | G | G | C | A | C | A | A | C | T | C | T | T | G | A | T | A | C | G | A | G | T | C | A | T | G | A | A | T | A | C | G | T | A | T | T | C | T | T | G | G | A | T | C | C | T | C | T | C | G | A | T | T | T | C | T | T | - | - | - | - | - | - | - | - | - | A | G | A | A | C | G | C | G | C | G |  |
| mut |  |  |  |  |  |  |  |  |  |  |  |  |  |  |  |  |  |  |  |  |  |  |  |  |  |  |  |  |  |  |  |  |  |  |  |  |  |  |  |  | C>T |  |  |  |  |  |  |  |  |  |  |  |  |  |  |  |  |  |  |  |  |  |  |  |  |  |  |  |  |  |  |  |  |  |  |  |  |  |  |  |  |  |  |  |  |  |  |  |  |  |  |  |  |  |  |  |  |  |  |  |  |  |  |  |  |  |  |  |  |  |  |  |  |  |  |  |  |  |  |  |  |  |  |  |  |  |  |  |  |  |  |  |  |  |  |  |  |  |  |  |  |  |

|  |  |  |  |  |  |  |  |  |  |  |  |  |  |  |  |  |  |  |  |  |  |  |  |  |  |  |  |  |  |  |  |  |  |  |  |  |  |  |  |  |  |  |  |  |  |  |  |  |  |  |  |  |  |  |  |  |  |  |  |  |  |  |  |  |  |  |  |  |  |  |  |  |  |  |  |  |  |  |  |  |  |  |  |  |  |  |  |  |  |  |  |  |  |
| --- | --- | --- | --- | --- | --- | --- | --- | --- | --- | --- | --- | --- | --- | --- | --- | --- | --- | --- | --- | --- | --- | --- | --- | --- | --- | --- | --- | --- | --- | --- | --- | --- | --- | --- | --- | --- | --- | --- | --- | --- | --- | --- | --- | --- | --- | --- | --- | --- | --- | --- | --- | --- | --- | --- | --- | --- | --- | --- | --- | --- | --- | --- | --- | --- | --- | --- | --- | --- | --- | --- | --- | --- | --- | --- | --- | --- | --- | --- | --- | --- | --- | --- | --- | --- | --- | --- | --- | --- | --- | --- | --- | --- | --- |
| pos | 670 | 2470 | 2790 | 2832 | 3291 | 4184 | 4321 | 5386 | 6513 | 6514 | 6515 | 8393 | 9344 | 9424 | 9534 | 9866 | 10198 | 10323 | 10447 | 11283 | 11284 | 11285 | 11286 | 11287 | 11292 | 11293 | 11294 | 11295 | 11296 | 11537 | 12880 | 13195 | 13366 | 14940 | 15240 | 15714 | 17410 | 19955 | 20055 | 21618 | 21633 | 21634 | 21635 | 21636 | 21637 | 21638 | 21639 | 21640 | 21641 | 21762 | 21765 | 21766 | 21767 | 21768 | 21769 | 21770 | 21846 | 21987 | 21988 | 21989 | 21990 | 21991 | 21992 | 21993 | 21994 | 21995 | 22194 | 22195 | 22196 | 22200 | 22599 | 22673 | 22688 | 22775 | 22786 | 22792 | 22898 | 23048 | 23202 | 23610 | 24130 | 24503 | 25714 | 26060 | 26530 | 26858 | 27382 | 27383 | 27384 | 28426 | 29000 | 29510 |  |
| ref | T | C | C | A | A | G | C | T | G | T | T | G | C | A | C | C | C | A | G | G | T | T | T | G | G | T | T | T | T | A | C | T | C | A | C | C | C | C | A | C | T | A | C | C | C | C | C | T | G | C | T | A | C | A | T | G | C | G | T | G | T | T | T | A | T | T | A | T | T | T | G | T | A | G | A | C | G | G | C | G | C | C | C | C | A | C | G | A | T | T | G | A |  |
| BA.2.34 | G | C | T | A | A | A | T | T | G | T | T | G | T | G | T | T | T | A | A | G | T | T | T | G | - | - | - | - | - | A | T | T | C | A | C | T | T | T | G | T | - | - | - | - | - | - | - | - | - | C | T | A | C | A | T | G | C | A | T | G | T | T | T | A | T | T | A | T | T | G | G | T | G | A | C | T | G | G | C | G | C | C | C | T | A | T | C | T | C | T | A | C |  |
| BA.2 | G | C | T | A | A | A | T | T | G | T | T | G | T | G | T | T | T | A | A | G | T | T | T | G | - | - | - | - | - | A | T | T | C | A | C | T | T | C | A | C | T | A | C | C | C | C | C | T | G | T | - | - | - | - | - | - | T | - | - | - | - | - | - | - | - | - | - | - | - | T | A | C | A | G | A | C | A | A | A | T | A | T | G | C | G | C | G | A | T | C | G | A | #1002687 |
| BA.1.1 | T | T | C | G | G | G | C | G | - | - | - | A | C | A | C | C | C | G | G | - | - | - | - | - | G | T | T | T | T | G | C | C | T | G | T | C | C | C | A | C | T | A | C | C | C | C | C | T | G | T | - | - | - | - | - | - | T | - | - | - | - | - | - | - | - | - | - | - | - | T | A | C | A | G | A | C | A | A | A | G | A | T | C | C | G | C | G | A | T | C | G | A |  |
| mut |  |  |  |  |  |  |  |  |  |  |  |  |  |  |  |  |  |  |  |  |  |  |  |  |  |  |  |  |  |  |  |  |  |  |  |  |  |  |  |  |  |  |  |  |  |  |  |  |  |  |  |  |  |  |  |  |  |  |  |  |  |  |  |  |  |  |  |  |  |  |  |  |  |  |  |  |  |  |  | G>T |  |  | C>G |  |  |  |  |  |  |  |  |  |  |

|  |  |  |  |  |  |  |  |  |  |  |  |  |  |  |  |  |  |  |  |  |  |  |  |  |  |  |  |  |  |  |  |  |  |  |  |  |  |  |  |  |  |  |  |  |  |  |  |  |  |  |  |  |  |  |  |  |  |  |  |  |  |  |  |  |  |  |  |  |  |  |  |  |  |  |  |  |  |  |  |  |  |
| --- | --- | --- | --- | --- | --- | --- | --- | --- | --- | --- | --- | --- | --- | --- | --- | --- | --- | --- | --- | --- | --- | --- | --- | --- | --- | --- | --- | --- | --- | --- | --- | --- | --- | --- | --- | --- | --- | --- | --- | --- | --- | --- | --- | --- | --- | --- | --- | --- | --- | --- | --- | --- | --- | --- | --- | --- | --- | --- | --- | --- | --- | --- | --- | --- | --- | --- | --- | --- | --- | --- | --- | --- | --- | --- | --- | --- | --- | --- | --- | --- | --- |
| pos | 670 | 2470 | 2790 | 2832 | 4184 | 4321 | 5386 | 6513 | 6514 | 6515 | 8393 | 9344 | 9424 | 9534 | 10198 | 10447 | 11283 | 11284 | 11285 | 11286 | 11287 | 11292 | 11293 | 11294 | 11295 | 11296 | 11537 | 12880 | 13195 | 15240 | 15714 | 17410 | 21595 | 21633 | 21634 | 21635 | 21636 | 21637 | 21638 | 21639 | 21640 | 21641 | 21762 | 21765 | 21766 | 21767 | 21768 | 21769 | 21770 | 21846 | 21987 | 21988 | 21989 | 21990 | 21991 | 21992 | 21993 | 21994 | 21995 | 22194 | 22195 | 22196 | 22200 | 22599 | 22673 | 22688 | 22775 | 22786 | 22898 | 23048 | 23202 | 24130 | 24503 | 26060 | 26530 | 26858 | 27382 | 27383 | 27384 | 29510 |  |
| ref | T | C | C | A | G | C | T | G | T | T | G | C | A | C | C | G | G | T | T | T | G | G | T | T | T | T | A | C | T | C | C | C | C | T | A | C | C | C | C | C | T | G | C | T | A | C | A | T | G | C | G | T | G | T | T | T | A | T | T | A | T | T | T | G | T | A | G | A | G | G | C | C | C | C | A | C | G | A | T | A |  |
| BA.1.1 | T | T | C | G | G | C | G | - | - | - | A | C | A | C | C | G | - | - | - | - | - | G | T | T | T | T | G | C | C | T | C | C | T | T | A | C | C | C | C | C | T | G | T | - | - | - | - | - | - | T | - | - | - | - | - | - | - | - | - | - | - | - | T | A | C | A | G | A | A | A | A | A | T | C | G | C | G | A | T | A |  |
| XM | T | T | C | G | G | C | G | - | - | - | A | C | A | C | C | G | - | - | - | - | - | G | T | T | T | T | G | C | C | T | C | C | C | - | - | - | - | - | - | - | - | - | C | T | A | C | A | T | G | C | A | T | G | T | T | T | A | T | T | A | T | T | G | G | T | G | A | C | G | G | C | C | C | T | A | T | C | T | C | C | #1003220 |
| BA.2 | G | C | T | A | A | T | T | G | T | T | G | T | G | T | T | A | G | T | T | T | G | - | - | - | - | - | A | T | T | C | T | T | C | - | - | - | - | - | - | - | - | - | C | T | A | C | A | T | G | C | A | T | G | T | T | T | A | T | T | A | T | T | G | G | T | G | A | C | G | G | C | C | C | T | A | T | C | T | C | C |  |
| mut |  |  |  |  |  |  |  |  |  |  |  |  |  |  |  |  |  |  |  |  |  |  |  |  |  |  |  |  |  |  |  |  |  |  |  |  |  |  |  |  |  |  |  |  |  |  |  |  |  |  |  |  |  |  |  |  |  |  |  |  |  |  |  |  |  |  |  |  |  |  |  |  |  |  |  |  |  |  |  |  |  |

|  |  |  |  |  |  |  |  |  |  |  |  |  |  |  |  |  |
| --- | --- | --- | --- | --- | --- | --- | --- | --- | --- | --- | --- | --- | --- | --- | --- | --- |
| pos | 1523 | 2398 | 3003 | 5515 | 6040 | 6337 | 6504 | 13193 | 21587 | 22813 | 22882 | 22898 | 22917 | 23191 | 26198 |  |
| ref | T | G | A | G | C | G | C | G | C | G | T | G | T | C | C |  |
| BA.1 | A | G | G | G | T | T | T | G | C | G | T | G | G | C | C |  |
| BA.1 | A | G | G | G | T | T | T | G | T | T | G | A | T | C | T | #1008581 |
| BA.1 | T | A | A | T | C | G | C | A | T | T | G | A | T | T | C |  |
| mut |  |  |  |  |  |  |  |  |  |  |  |  |  | T>C | C>T |  |

|  |  |  |  |  |  |  |  |  |  |  |  |  |  |  |  |  |  |  |  |  |  |  |  |  |  |  |
| --- | --- | --- | --- | --- | --- | --- | --- | --- | --- | --- | --- | --- | --- | --- | --- | --- | --- | --- | --- | --- | --- | --- | --- | --- | --- | --- |
| pos | 210 | 2832 | 4181 | 5365 | 5386 | 5515 | 6196 | 6402 | 6513 | 6514 | 6515 | 7124 | 8393 | 8986 | 9053 | 21595 | 22194 | 22195 | 22196 | 22197 | 22198 | 22202 | 22204 | 22599 | 22813 |  |
| ref | G | A | G | C | T | G | C | C | G | T | T | C | G | C | G | C | A | T | T | T | A | C | T | G | G |  |
| XS | T | A | T | T | T | G | T | T | G | T | T | T | G | T | T | T | A | G | T | G | C | A | C | A | T |  |
| XS | T | A | T | T | T | G | T | T | G | T | T | T | G | T | T | T | - | - | - | T | A | C | T | A | G | #1014313 |
| BA.1 | G | G | G | C | G | T | C | C | - | - | - | C | A | C | G | C | - | - | - | T | A | C | T | G | G |  |
| mut |  |  |  |  |  |  |  |  |  |  |  |  |  |  |  |  |  |  |  |  |  |  |  | G>A |  |  |

|  |  |  |  |  |  |  |  |  |  |  |  |  |  |  |
| --- | --- | --- | --- | --- | --- | --- | --- | --- | --- | --- | --- | --- | --- | --- |
| pos | 1613 | 4305 | 17608 | 20946 | 22194 | 22195 | 22196 | 22197 | 22198 | 22202 | 22204 | 22882 | 23855 |  |
| ref | C | A | G | C | A | T | T | T | A | C | T | T | C |  |
| BA.1.15 | T | G | A | C | A | G | T | G | C | A | C | G | C |  |
| BA.1.15 | T | G | A | C | - | - | - | T | A | C | T | T | A | #1023548 |
| BA.1.15 | C | A | G | T | - | - | - | T | A | C | T | T | A |  |
| mut |  |  |  |  |  |  |  |  |  |  |  |  |  |  |

|  |  |  |  |  |  |  |  |  |  |  |  |  |  |  |  |  |
| --- | --- | --- | --- | --- | --- | --- | --- | --- | --- | --- | --- | --- | --- | --- | --- | --- |
| pos | 507 | 508 | 509 | 518 | 2470 | 6279 | 6541 | 6583 | 7926 | 10135 | 14268 | 22599 | 25708 | 27513 | 29301 |  |
| ref | A | T | G | A | C | C | C | T | C | T | G | G | C | C | A |  |
| BA.1.1.18 | T | C | A | G | T | C | C | T | T | T | G | A | C | C | A |  |
| BA.1.15 | T | C | A | G | C | A | T | C | C | C | A | G | T | A | G | #1028159 |
| BA.1.15 | A | T | G | A | C | A | T | C | C | C | A | G | T | A | G |  |
| mut |  |  |  |  |  |  |  |  |  |  |  |  |  |  |  |  |

|  |  |  |  |  |  |  |  |  |  |  |  |  |  |  |  |  |  |  |  |  |  |  |  |  |  |  |  |  |  |  |  |  |  |  |  |  |  |  |  |  |  |  |  |  |  |  |  |  |  |  |  |  |  |  |  |  |  |  |  |  |  |  |  |  |  |  |  |  |  |  |  |  |  |  |  |  |  |  |  |  |  |  |  |  |  |  |  |  |  |  |  |
| --- | --- | --- | --- | --- | --- | --- | --- | --- | --- | --- | --- | --- | --- | --- | --- | --- | --- | --- | --- | --- | --- | --- | --- | --- | --- | --- | --- | --- | --- | --- | --- | --- | --- | --- | --- | --- | --- | --- | --- | --- | --- | --- | --- | --- | --- | --- | --- | --- | --- | --- | --- | --- | --- | --- | --- | --- | --- | --- | --- | --- | --- | --- | --- | --- | --- | --- | --- | --- | --- | --- | --- | --- | --- | --- | --- | --- | --- | --- | --- | --- | --- | --- | --- | --- | --- | --- | --- | --- | --- | --- | --- |
| pos | 44 | 670 | 2470 | 2790 | 2832 | 4184 | 4321 | 5386 | 6513 | 6514 | 6515 | 7926 | 8393 | 9344 | 9424 | 9534 | 9866 | 10198 | 10447 | 11283 | 11284 | 11285 | 11286 | 11287 | 11292 | 11293 | 11294 | 11295 | 11296 | 11537 | 12880 | 13195 | 15240 | 15714 | 16342 | 17410 | 19955 | 20055 | 21618 | 21633 | 21634 | 21635 | 21636 | 21637 | 21638 | 21639 | 21640 | 21641 | 21762 | 21765 | 21766 | 21767 | 21768 | 21769 | 21770 | 21846 | 21987 | 21988 | 21989 | 21990 | 21991 | 21992 | 21993 | 21994 | 21995 | 22195 | 22197 | 22198 | 22200 | 22202 | 22204 | 22599 | 22673 | 22688 | 22775 | 22786 | 22898 | 23048 | 23202 | 24130 | 24503 | 25855 | 26060 | 26530 | 26858 | 27382 | 27383 | 27384 | 29510 | 29754 |  |
| ref | C | T | C | C | A | G | C | T | G | T | T | C | G | C | A | C | C | C | G | G | T | T | T | G | G | T | T | T | T | A | C | T | C | C | T | C | C | A | C | T | A | C | C | C | C | C | T | G | C | T | A | C | A | T | G | C | G | T | G | T | T | T | A | T | T | T | T | A | T | C | T | G | T | A | G | A | G | G | C | C | C | G | C | A | C | G | A | T | A | C |  |
| BA.2.10 | T | G | C | T | A | A | T | T | G | T | T | C | G | T | G | T | T | T | A | G | T | T | T | G | - | - | - | - | - | A | T | T | C | T | C | T | T | G | T | - | - | - | - | - | - | - | - | - | C | T | A | C | A | T | G | C | A | T | G | T | T | T | A | T | T | T | T | A | G | C | T | G | T | G | A | C | G | G | C | C | C | G | T | A | T | C | T | C | C | C |  |
| BA.2 | T | G | C | T | A | A | T | T | G | T | T | C | G | T | G | T | T | T | A | G | T | T | T | G | - | - | - | - | - | A | T | T | C | T | C | T | T | G | T | - | - | - | - | - | - | - | - | - | C | T | A | C | A | T | G | C | A | T | G | T | T | T | A | T | T | T | T | A | G | C | T | G | T | G | A | C | G | G | C | C | C | G | T | A | T | G | A | T | A | T | #1030562 |
| BA.1.1.18 | C | T | T | C | G | G | C | G | - | - | - | T | A | C | A | C | C | C | G | - | - | - | - | - | G | T | T | T | T | G | C | C | T | C | T | C | C | A | C | T | A | C | C | C | C | C | T | G | T | - | - | - | - | - | - | T | - | - | - | - | - | - | - | - | - | G | G | C | T | A | C | A | C | A | G | A | A | A | A | A | T | T | C | G | C | G | A | T | A | T |  |
| mut |  |  |  |  |  |  |  |  |  |  |  |  |  |  |  |  |  |  |  |  |  |  |  |  |  |  |  |  |  |  |  |  |  |  |  |  |  |  |  |  |  |  |  |  |  |  |  |  |  |  |  |  |  |  |  |  |  |  |  |  |  |  |  |  |  |  |  |  |  |  |  |  |  |  |  |  |  |  |  |  |  |  |  |  |  |  |  |  |  |  |  |

|  |  |  |  |  |  |  |  |  |  |  |  |  |  |  |  |  |  |  |  |  |  |  |  |  |  |  |  |  |  |  |  |  |  |  |  |  |  |  |  |  |  |  |  |  |  |  |  |  |  |  |  |  |  |  |  |  |  |  |  |  |  |  |  |  |  |  |  |  |  |  |  |  |  |  |  |  |  |  |  |  |  |  |  |  |  |  |  |  |
| --- | --- | --- | --- | --- | --- | --- | --- | --- | --- | --- | --- | --- | --- | --- | --- | --- | --- | --- | --- | --- | --- | --- | --- | --- | --- | --- | --- | --- | --- | --- | --- | --- | --- | --- | --- | --- | --- | --- | --- | --- | --- | --- | --- | --- | --- | --- | --- | --- | --- | --- | --- | --- | --- | --- | --- | --- | --- | --- | --- | --- | --- | --- | --- | --- | --- | --- | --- | --- | --- | --- | --- | --- | --- | --- | --- | --- | --- | --- | --- | --- | --- | --- | --- | --- | --- | --- | --- | --- |
| pos | 670 | 875 | 2790 | 2832 | 3241 | 4184 | 4321 | 5386 | 5924 | 6513 | 6514 | 6515 | 8393 | 9344 | 9424 | 9534 | 9866 | 10198 | 10447 | 11283 | 11284 | 11285 | 11286 | 11287 | 11292 | 11293 | 11294 | 11295 | 11296 | 11537 | 12880 | 13195 | 14229 | 15240 | 15714 | 17410 | 19955 | 20055 | 21618 | 21633 | 21634 | 21635 | 21636 | 21637 | 21638 | 21639 | 21640 | 21641 | 21762 | 21765 | 21766 | 21767 | 21768 | 21769 | 21770 | 21846 | 21987 | 21988 | 21989 | 21990 | 21991 | 21992 | 21993 | 21994 | 21995 | 22194 | 22195 | 22196 | 22200 | 22673 | 22688 | 22775 | 22786 | 22813 | 23048 | 23202 | 23664 | 24130 | 24503 | 26060 | 26530 | 26858 | 27382 | 27383 | 27384 | 29420 | 29510 |  |
| ref | T | C | C | A | C | G | C | T | G | G | T | T | G | C | A | C | C | C | G | G | T | T | T | G | G | T | T | T | T | A | C | T | G | C | C | C | C | A | C | T | A | C | C | C | C | C | T | G | C | T | A | C | A | T | G | C | G | T | G | T | T | T | A | T | T | A | T | T | T | T | A | G | A | G | G | C | C | C | C | C | A | C | G | A | T | C | A |  |
| BA.1.17.2 | T | T | C | G | T | G | C | G | A | - | - | - | A | C | A | C | C | C | G | - | - | - | - | - | G | T | T | T | T | G | C | C | G | T | C | C | C | A | C | T | A | C | C | C | C | C | T | G | T | - | - | - | - | - | - | T | - | - | - | - | - | - | - | - | - | - | - | - | T | C | A | G | A | G | A | A | T | A | T | C | G | C | G | A | T | A | A |  |
| XL | T | T | C | G | T | G | C | G | A | - | - | - | G | T | G | T | T | T | A | G | T | T | T | G | - | - | - | - | - | A | T | T | A | C | T | T | T | G | T | - | - | - | - | - | - | - | - | - | C | T | A | C | A | T | G | C | A | T | G | T | T | T | A | T | T | A | T | T | G | T | G | A | C | T | G | C | C | C | C | T | A | T | C | T | C | C | C | #1034619 |
| BA.2.5 | G | C | T | A | C | A | T | T | G | G | T | T | G | T | G | T | T | T | A | G | T | T | T | G | - | - | - | - | - | A | T | T | G | C | T | T | T | G | T | - | - | - | - | - | - | - | - | - | C | T | A | C | A | T | G | C | A | T | G | T | T | T | A | T | T | A | T | T | G | T | G | A | C | T | G | C | C | C | C | T | A | T | C | T | C | C | C |  |
| mut |  |  |  |  |  |  |  |  |  |  |  |  |  |  |  |  |  |  |  |  |  |  |  |  |  |  |  |  |  |  |  |  | G>A |  |  |  |  |  |  |  |  |  |  |  |  |  |  |  |  |  |  |  |  |  |  |  |  |  |  |  |  |  |  |  |  |  |  |  |  |  |  |  |  |  |  |  |  |  |  |  |  |  |  |  |  |  |  |  |

|  |  |  |  |  |  |  |  |  |  |  |  |  |  |
| --- | --- | --- | --- | --- | --- | --- | --- | --- | --- | --- | --- | --- | --- |
| pos | 2470 | 5730 | 21595 | 22599 | 24815 | 25922 | 29734 | 29735 | 29736 | 29760 | 29762 | 29766 |  |
| ref | C | C | C | G | G | G | G | A | G | T | C | G |  |
| BA.1.18 | C | T | C | G | G | A | G | A | G | T | - | G |  |
| BA.1.18 | C | T | C | G | G | A | A | G | A | C | C | T | #1044332 |
| BA.1.1 | T | C | T | A | A | G | A | G | A | C | C | T |  |
| mut |  |  |  |  |  |  |  |  |  |  |  |  |  |

|  |  |  |  |  |  |  |  |  |  |  |  |  |  |  |  |  |  |  |  |  |  |  |  |  |  |  |  |  |  |  |  |  |  |  |  |  |  |  |  |  |  |  |  |  |  |  |  |  |  |  |  |  |  |  |  |  |  |  |  |  |  |  |  |  |  |  |  |  |  |  |  |  |  |  |  |  |  |  |  |  |  |  |  |  |  |  |  |  |  |  |  |
| --- | --- | --- | --- | --- | --- | --- | --- | --- | --- | --- | --- | --- | --- | --- | --- | --- | --- | --- | --- | --- | --- | --- | --- | --- | --- | --- | --- | --- | --- | --- | --- | --- | --- | --- | --- | --- | --- | --- | --- | --- | --- | --- | --- | --- | --- | --- | --- | --- | --- | --- | --- | --- | --- | --- | --- | --- | --- | --- | --- | --- | --- | --- | --- | --- | --- | --- | --- | --- | --- | --- | --- | --- | --- | --- | --- | --- | --- | --- | --- | --- | --- | --- | --- | --- | --- | --- | --- | --- | --- | --- | --- |
| pos | 670 | 2335 | 2470 | 2790 | 2832 | 4184 | 4321 | 5386 | 6513 | 6514 | 6515 | 8393 | 9344 | 9424 | 9534 | 9866 | 10198 | 10447 | 11283 | 11284 | 11285 | 11286 | 11287 | 11292 | 11293 | 11294 | 11295 | 11296 | 11537 | 11674 | 12880 | 13195 | 15240 | 15714 | 17242 | 17410 | 19955 | 20055 | 21618 | 21633 | 21634 | 21635 | 21636 | 21637 | 21638 | 21639 | 21640 | 21641 | 21762 | 21765 | 21766 | 21767 | 21768 | 21769 | 21770 | 21846 | 21987 | 21988 | 21989 | 21990 | 21991 | 21992 | 21993 | 21994 | 21995 | 22194 | 22195 | 22196 | 22200 | 22599 | 22673 | 22688 | 22775 | 22786 | 22898 | 23048 | 23202 | 24130 | 24503 | 26060 | 26522 | 26530 | 26858 | 27382 | 27383 | 27384 | 27639 | 27688 | 29020 | 29510 |  |
| ref | T | T | C | C | A | G | C | T | G | T | T | G | C | A | C | C | C | G | G | T | T | T | G | G | T | T | T | T | A | C | C | T | C | C | G | C | C | A | C | T | A | C | C | C | C | C | T | G | C | T | A | C | A | T | G | C | G | T | G | T | T | T | A | T | T | A | T | T | T | G | T | A | G | A | G | G | C | C | C | C | C | A | C | G | A | T | T | C | A | A |  |
| BA.2 | G | C | C | T | A | A | T | T | G | T | T | G | T | G | T | T | T | A | G | T | T | T | G | - | - | - | - | - | A | C | T | T | C | T | T | T | T | G | T | - | - | - | - | - | - | - | - | - | C | T | A | C | A | T | G | C | A | T | G | T | T | T | A | T | T | A | T | T | G | G | T | G | A | C | G | G | C | C | C | T | C | A | T | C | T | C | A | C | G | C |  |
| BA.1 | G | C | C | T | A | A | T | T | G | T | T | G | T | G | T | T | T | A | - | - | - | - | - | G | T | T | T | T | G | T | C | C | T | C | G | C | C | A | C | T | A | C | C | C | C | C | T | G | T | - | - | - | - | - | - | T | - | - | - | - | - | - | - | - | - | - | - | - | T | A | C | A | G | A | A | A | A | A | T | C | T | G | C | G | A | T | T | T | A | A | #1048126 |
| BA.1.1 | T | T | T | C | G | G | C | G | - | - | - | A | C | A | C | C | C | G | - | - | - | - | - | G | T | T | T | T | G | T | C | C | T | C | G | C | C | A | C | T | A | C | C | C | C | C | T | G | T | - | - | - | - | - | - | T | - | - | - | - | - | - | - | - | - | - | - | - | T | A | C | A | G | A | A | A | A | A | T | C | T | G | C | G | A | T | T | T | A | A |  |
| mut |  |  |  |  |  |  |  |  |  |  |  |  |  |  |  |  |  |  |  |  |  |  |  |  |  |  |  |  |  |  |  |  |  |  |  |  |  |  |  |  |  |  |  |  |  |  |  |  |  |  |  |  |  |  |  |  |  |  |  |  |  |  |  |  |  |  |  |  |  |  |  |  |  |  |  |  |  |  |  |  |  |  |  |  |  |  |  |  |  |  |  |

|  |  |  |  |  |  |  |  |  |  |  |  |  |  |  |  |  |  |  |  |  |  |  |  |  |
| --- | --- | --- | --- | --- | --- | --- | --- | --- | --- | --- | --- | --- | --- | --- | --- | --- | --- | --- | --- | --- | --- | --- | --- | --- |
| pos | 2470 | 3241 | 5310 | 5924 | 16884 | 22194 | 22195 | 22196 | 22197 | 22198 | 22202 | 22204 | 22599 | 22933 | 23664 | 25906 | 26750 | 27384 | 27879 | 27880 | 27881 | 27882 | 27883 |  |
| ref | C | C | C | G | T | A | T | T | T | A | C | T | G | G | C | G | C | T | C | A | C | G | C |  |
| BA.1.1 | T | C | T | G | T | A | G | T | G | C | A | C | A | G | C | G | C | T | C | A | C | G | C |  |
| BA.1.1 | T | C | T | G | T | A | G | T | G | C | A | C | A | G | C | G | C | T | A | G | T | C | T | #1048896 |
| BA.1.17.2 | C | T | C | A | C | - | - | - | T | A | C | T | G | A | T | T | A | C | A | G | T | C | T |  |
| mut |  |  |  |  |  |  |  |  |  |  |  |  |  |  |  |  |  |  |  |  |  |  |  |  |

|  |  |  |  |  |  |  |  |  |  |  |  |  |  |  |  |  |  |
| --- | --- | --- | --- | --- | --- | --- | --- | --- | --- | --- | --- | --- | --- | --- | --- | --- | --- |
| pos | 2037 | 3864 | 7926 | 11250 | 11537 | 14940 | 17145 | 21742 | 22194 | 22195 | 22196 | 22197 | 22198 | 22202 | 22204 | 22227 |  |
| ref | C | A | C | G | A | A | C | C | A | T | T | T | A | C | T | C |  |
| BA.1.1.18 | T | C | T | G | G | A | T | C | A | G | T | G | C | A | C | C |  |
| BA.1.1.18 | T | C | T | G | G | A | T | C | - | - | - | T | A | C | T | T | #1052140 |
| BA.1.1 | C | A | C | A | A | G | C | A | - | - | - | T | A | C | T | T |  |
| mut |  |  |  |  |  |  |  |  |  |  |  |  |  |  |  |  |  |

|  |  |  |  |  |  |  |  |  |  |  |  |  |  |  |  |  |  |  |  |  |  |  |  |  |  |  |  |  |  |  |  |  |  |  |  |  |  |  |  |  |  |  |  |  |  |  |  |  |  |  |  |  |  |  |  |  |  |  |  |  |  |  |  |  |  |  |  |  |  |  |  |  |  |  |  |  |  |  |  |  |  |  |  |  |  |  |  |  |  |  |  |  |  |  |  |  |  |  |  |  |  |  |  |  |  |  |  |  |  |  |  |  |  |  |  |  |  |  |  |  |  |  |  |  |  |  |  |  |  |  |  |  |  |  |  |  |
| --- | --- | --- | --- | --- | --- | --- | --- | --- | --- | --- | --- | --- | --- | --- | --- | --- | --- | --- | --- | --- | --- | --- | --- | --- | --- | --- | --- | --- | --- | --- | --- | --- | --- | --- | --- | --- | --- | --- | --- | --- | --- | --- | --- | --- | --- | --- | --- | --- | --- | --- | --- | --- | --- | --- | --- | --- | --- | --- | --- | --- | --- | --- | --- | --- | --- | --- | --- | --- | --- | --- | --- | --- | --- | --- | --- | --- | --- | --- | --- | --- | --- | --- | --- | --- | --- | --- | --- | --- | --- | --- | --- | --- | --- | --- | --- | --- | --- | --- | --- | --- | --- | --- | --- | --- | --- | --- | --- | --- | --- | --- | --- | --- | --- | --- | --- | --- | --- | --- | --- | --- | --- | --- | --- | --- | --- | --- | --- | --- | --- | --- | --- | --- | --- | --- | --- | --- |
| pos | 210 | 829 | 949 | 1274 | 2470 | 2832 | 4041 | 4181 | 5386 | 6402 | 6513 | 6514 | 6515 | 7124 | 8302 | 8393 | 8986 | 9053 | 10449 | 11201 | 11283 | 11284 | 11285 | 11286 | 11287 | 11288 | 11289 | 11290 | 11291 | 11332 | 11537 | 13195 | 15240 | 15451 | 15960 | 16466 | 17236 | 18163 | 19220 | 21595 | 21618 | 21762 | 21765 | 21766 | 21767 | 21768 | 21769 | 21770 | 21846 | 21987 | 21988 | 21989 | 21990 | 21991 | 21992 | 21993 | 21994 | 21995 | 22029 | 22030 | 22031 | 22032 | 22033 | 22034 | 22194 | 22195 | 22196 | 22578 | 22599 | 22673 | 22674 | 22679 | 22686 | 22813 | 22882 | 22898 | 22917 | 22992 | 23013 | 23040 | 23048 | 23055 | 23063 | 23075 | 23202 | 23525 | 23599 | 23604 | 23854 | 23948 | 24130 | 24208 | 24410 | 24424 | 24469 | 24503 | 25000 | 25469 | 25584 | 26270 | 26530 | 26577 | 26709 | 26767 | 27259 | 27294 | 27638 | 27752 | 27807 | 27874 | 28248 | 28249 | 28250 | 28251 | 28252 | 28253 | 28271 | 28311 | 28362 | 28363 | 28364 | 28365 | 28366 | 28367 | 28368 | 28369 | 28370 | 28461 | 28808 | 28881 | 28882 | 28883 | 28916 | 29402 | 29742 |  |
| ref | G | C | A | G | C | A | T | G | T | C | G | T | T | C | A | G | C | G | C | A | G | T | T | T | G | T | C | T | G | A | A | T | C | G | C | C | A | A | C | C | C | C | T | A | C | A | T | G | C | G | T | G | T | T | T | A | T | T | A | G | T | T | C | A | A | T | T | G | G | T | C | T | C | G | T | G | T | G | A | A | G | A | A | T | C | C | T | C | C | G | C | C | G | A | T | C | C | C | C | C | A | C | G | T | A | C | T | C | C | C | G | A | T | T | T | C | A | C | G | A | G | A | A | C | G | C | A | A | G | G | G | G | G | G | G |  |
| AY.103 | T | T | G | T | C | A | T | T | T | T | G | T | T | T | A | G | T | T | C | G | G | T | T | T | G | T | C | T | G | G | A | T | C | A | C | T | G | A | T | C | G | C | T | A | C | A | T | G | C | A | T | G | T | T | T | A | T | T | - | - | - | - | - | - | A | T | T | G | G | T | C | T | C | G | T | G | G | G | A | A | G | A | A | T | C | C | T | G | C | G | C | T | A | A | T | C | C | T | C | C | A | C | G | C | A | T | C | T | C | T | - | - | - | - | - | - | - | C | G | A | G | A | A | C | G | C | A | G | A | T | G | G | T | T | T |  |
| BA.1 | T | T | G | T | C | A | C | T | T | C | - | - | - | C | G | A | C | G | A | A | - | - | - | - | - | - | - | - | - | A | G | C | T | G | T | C | A | G | C | T | C | T | - | - | - | - | - | - | T | - | - | - | - | - | - | - | - | - | A | G | T | T | C | A | - | - | - | A | A | C | T | C | T | T | G | A | T | A | C | G | A | G | T | C | A | T | G | A | A | T | A | C | G | T | A | T | T | C | T | T | G | G | A | T | C | C | T | C | T | C | G | A | T | T | T | C | T | T | - | - | - | - | - | - | - | - | - | A | G | A | A | C | G | G | G | #1053479 |
| BA.1.1 | G | C | A | G | T | G | T | G | G | C | - | - | - | C | G | A | C | G | A | A | - | - | - | - | - | - | - | - | - | A | G | C | T | G | C | C | A | G | C | T | C | T | - | - | - | - | - | - | T | - | - | - | - | - | - | - | - | - | A | G | T | T | C | A | - | - | - | A | A | C | T | C | T | T | G | A | T | A | C | G | A | G | T | C | A | T | G | A | A | T | A | C | G | T | A | T | T | C | T | T | G | G | A | T | C | C | T | C | T | C | G | A | T | T | T | C | T | T | - | - | - | - | - | - | - | - | - | A | G | A | A | C | G | G | G |  |
| mut |  |  |  |  |  |  | T>C |  |  |  |  |  |  |  |  |  |  |  |  |  |  |  |  |  |  |  |  |  |  |  |  |  |  |  | C>T |  |  |  |  |  |  |  |  |  |  |  |  |  |  |  |  |  |  |  |  |  |  |  |  |  |  |  |  |  |  |  |  |  |  |  |  |  |  |  |  |  |  |  |  |  |  |  |  |  |  |  |  |  |  |  |  |  |  |  |  |  |  |  |  |  |  |  |  |  |  |  |  |  |  |  |  |  |  |  |  |  |  |  |  |  |  |  |  |  |  |  |  |  |  |  |  |  |  |  |  |  |

|  |  |  |  |  |  |  |  |  |  |  |  |  |  |  |  |  |  |  |  |  |  |  |  |  |  |  |  |  |  |  |  |  |  |  |  |  |  |  |  |  |  |  |  |  |  |  |  |  |  |  |  |  |  |  |  |  |  |  |  |  |  |  |  |  |  |  |  |  |  |  |  |  |  |  |  |  |  |  |  |  |  |  |  |  |  |  |
| --- | --- | --- | --- | --- | --- | --- | --- | --- | --- | --- | --- | --- | --- | --- | --- | --- | --- | --- | --- | --- | --- | --- | --- | --- | --- | --- | --- | --- | --- | --- | --- | --- | --- | --- | --- | --- | --- | --- | --- | --- | --- | --- | --- | --- | --- | --- | --- | --- | --- | --- | --- | --- | --- | --- | --- | --- | --- | --- | --- | --- | --- | --- | --- | --- | --- | --- | --- | --- | --- | --- | --- | --- | --- | --- | --- | --- | --- | --- | --- | --- | --- | --- | --- | --- | --- | --- |
| pos | 670 | 2470 | 2790 | 2832 | 4184 | 4321 | 5386 | 6513 | 6514 | 6515 | 8393 | 9344 | 9424 | 9534 | 9866 | 10198 | 10447 | 11283 | 11284 | 11285 | 11286 | 11287 | 11292 | 11293 | 11294 | 11295 | 11296 | 11537 | 12880 | 13195 | 15240 | 15714 | 17410 | 18537 | 19955 | 20055 | 21618 | 21633 | 21634 | 21635 | 21636 | 21637 | 21638 | 21639 | 21640 | 21641 | 21762 | 21765 | 21766 | 21767 | 21768 | 21769 | 21770 | 21846 | 21987 | 21988 | 21989 | 21990 | 21991 | 21992 | 21993 | 21994 | 21995 | 22194 | 22195 | 22196 | 22200 | 22599 | 22673 | 22688 | 22775 | 22786 | 22898 | 22987 | 23048 | 23202 | 24130 | 24503 | 26060 | 26530 | 26858 | 27382 | 27383 | 27384 | 29510 |  |
| ref | T | C | C | A | G | C | T | G | T | T | G | C | A | C | C | C | G | G | T | T | T | G | G | T | T | T | T | A | C | T | C | C | C | T | C | A | C | T | A | C | C | C | C | C | T | G | C | T | A | C | A | T | G | C | G | T | G | T | T | T | A | T | T | A | T | T | T | G | T | A | G | A | G | C | G | C | C | C | C | A | C | G | A | T | A |  |
| BA.2 | G | C | T | A | A | T | T | G | T | T | G | T | G | T | T | T | A | G | T | T | T | G | - | - | - | - | - | A | T | T | C | T | T | T | T | G | T | - | - | - | - | - | - | - | - | - | C | T | A | C | A | T | G | C | A | T | G | T | T | T | A | T | T | A | T | T | G | G | T | G | A | C | G | T | G | C | C | C | T | A | T | C | T | C | C |  |
| BA.2 | G | C | T | A | A | T | T | G | T | T | G | T | G | T | T | T | A | G | T | T | T | G | - | - | - | - | - | A | T | T | T | C | C | G | C | A | C | T | A | C | C | C | C | C | T | G | T | - | - | - | - | - | - | T | - | - | - | - | - | - | - | - | - | - | - | - | T | A | C | A | G | A | A | C | A | A | A | T | C | G | C | G | A | T | A | #1055297 |
| BA.1.1 | T | T | C | G | G | C | G | - | - | - | A | C | A | C | C | C | G | - | - | - | - | - | G | T | T | T | T | G | C | C | T | C | C | G | C | A | C | T | A | C | C | C | C | C | T | G | T | - | - | - | - | - | - | T | - | - | - | - | - | - | - | - | - | - | - | - | T | A | C | A | G | A | A | C | A | A | A | T | C | G | C | G | A | T | A |  |
| mut |  |  |  |  |  |  |  |  |  |  |  |  |  |  |  |  |  |  |  |  |  |  |  |  |  |  |  |  |  |  | C>T |  |  |  |  |  |  |  |  |  |  |  |  |  |  |  |  |  |  |  |  |  |  |  |  |  |  |  |  |  |  |  |  |  |  |  |  |  |  |  |  |  |  |  |  |  |  |  |  |  |  |  |  |  |  |  |

|  |  |  |  |  |  |  |  |  |  |  |  |  |  |  |  |  |  |  |  |
| --- | --- | --- | --- | --- | --- | --- | --- | --- | --- | --- | --- | --- | --- | --- | --- | --- | --- | --- | --- |
| pos | 118 | 203 | 2470 | 4600 | 10576 | 11950 | 14184 | 15999 | 21595 | 22194 | 22195 | 22196 | 22197 | 22198 | 22202 | 22204 | 22599 | 26111 |  |
| ref | A | C | C | T | C | C | C | A | C | A | T | T | T | A | C | T | G | C |  |
| BA.1.1 | G | T | T | C | T | C | T | T | T | - | - | - | T | A | C | T | A | C |  |
| BA.1.1 | G | T | T | C | T | C | T | T | T | A | G | T | G | C | A | C | A | T | #1056796 |
| BA.1 | A | C | C | T | C | T | C | A | C | A | G | T | G | C | A | C | G | T |  |
| mut |  |  |  |  |  |  |  |  |  |  |  |  |  |  |  |  | G>A |  |  |

|  |  |  |  |  |  |  |  |  |  |  |  |  |  |  |  |  |  |  |  |  |  |  |  |  |  |  |  |  |  |  |  |  |  |  |  |  |  |  |  |  |  |  |  |  |  |  |  |  |  |  |  |  |  |  |  |  |  |  |  |  |  |  |  |  |  |  |  |  |  |  |  |  |  |  |  |  |  |  |  |  |  |  |  |  |  |  |  |  |  |  |  |  |  |  |  |  |  |  |  |  |  |  |  |  |  |  |  |  |  |  |  |  |  |  |  |  |  |  |  |  |  |  |  |  |  |  |  |  |  |  |  |  |  |  |  |  |  |  |  |  |  |  |  |  |  |  |  |  |  |  |
| --- | --- | --- | --- | --- | --- | --- | --- | --- | --- | --- | --- | --- | --- | --- | --- | --- | --- | --- | --- | --- | --- | --- | --- | --- | --- | --- | --- | --- | --- | --- | --- | --- | --- | --- | --- | --- | --- | --- | --- | --- | --- | --- | --- | --- | --- | --- | --- | --- | --- | --- | --- | --- | --- | --- | --- | --- | --- | --- | --- | --- | --- | --- | --- | --- | --- | --- | --- | --- | --- | --- | --- | --- | --- | --- | --- | --- | --- | --- | --- | --- | --- | --- | --- | --- | --- | --- | --- | --- | --- | --- | --- | --- | --- | --- | --- | --- | --- | --- | --- | --- | --- | --- | --- | --- | --- | --- | --- | --- | --- | --- | --- | --- | --- | --- | --- | --- | --- | --- | --- | --- | --- | --- | --- | --- | --- | --- | --- | --- | --- | --- | --- | --- | --- | --- | --- | --- | --- | --- | --- | --- | --- | --- | --- | --- | --- | --- | --- | --- | --- | --- |
| pos | 210 | 1820 | 2832 | 2992 | 4181 | 5386 | 6402 | 6513 | 6514 | 6515 | 7124 | 8063 | 8393 | 8752 | 8986 | 9053 | 10135 | 10449 | 11201 | 11283 | 11284 | 11285 | 11286 | 11287 | 11288 | 11289 | 11290 | 11291 | 11332 | 11537 | 11750 | 12513 | 13195 | 15240 | 15451 | 16466 | 17634 | 18163 | 19220 | 20451 | 21137 | 21618 | 21762 | 21765 | 21766 | 21767 | 21768 | 21769 | 21770 | 21846 | 21897 | 21987 | 21988 | 21989 | 21990 | 21991 | 21992 | 21993 | 21994 | 21995 | 22029 | 22030 | 22031 | 22032 | 22033 | 22034 | 22194 | 22195 | 22196 | 22578 | 22673 | 22674 | 22679 | 22686 | 22813 | 22882 | 22898 | 22917 | 22992 | 23013 | 23040 | 23048 | 23055 | 23063 | 23075 | 23202 | 23525 | 23599 | 23604 | 23854 | 23948 | 24130 | 24410 | 24424 | 24469 | 24503 | 25000 | 25469 | 25552 | 25584 | 25708 | 25886 | 26107 | 26270 | 26530 | 26577 | 26709 | 26759 | 26767 | 27259 | 27507 | 27638 | 27752 | 27807 | 27874 | 28077 | 28248 | 28249 | 28250 | 28251 | 28252 | 28253 | 28271 | 28311 | 28362 | 28363 | 28364 | 28365 | 28366 | 28367 | 28368 | 28369 | 28370 | 28461 | 28881 | 28882 | 28883 | 28916 | 29215 | 29301 | 29402 | 29686 | 29700 | 29742 | 29762 | 29769 | 29774 | 29779 | 29781 |  |
| ref | G | G | A | T | G | T | C | G | T | T | C | T | G | C | C | G | T | C | A | G | T | T | T | G | T | C | T | G | A | A | C | C | T | C | G | C | C | A | C | C | A | C | C | T | A | C | A | T | G | C | C | G | T | G | T | T | T | A | T | T | A | G | T | T | C | A | A | T | T | G | T | C | T | C | G | T | G | T | G | A | A | G | A | A | T | C | C | T | C | C | G | C | G | A | T | C | C | C | G | C | C | C | G | C | A | C | G | A | T | A | A | T | C | C | C | G | G | A | T | T | T | C | A | C | G | A | G | A | A | C | G | C | A | A | G | G | G | G | C | A | G | C | A | G | C | C | C | G | G |  |
| AY.25 | T | A | A | T | T | T | T | G | T | T | T | G | G | T | T | T | T | C | G | G | T | T | T | G | T | C | T | G | G | A | T | T | T | C | A | T | C | A | T | T | G | G | C | T | A | C | A | T | G | C | T | A | T | G | T | T | T | A | T | T | - | - | - | - | - | - | A | T | T | G | T | C | T | C | G | T | G | G | G | A | A | G | A | A | T | C | C | T | G | C | G | C | A | A | T | C | C | T | T | C | C | C | C | C | A | C | G | G | C | A | C | C | T | C | T | T | - | - | - | - | - | - | - | C | G | A | G | A | A | C | G | C | A | G | T | G | G | T | T | A | T | C | - | T | - | - | - | - | - |  |
| BA.1.15 | T | A | A | C | T | T | T | G | T | T | T | G | G | T | T | T | C | A | A | - | - | - | - | - | - | - | - | - | A | G | C | C | C | T | G | C | T | G | C | C | A | C | T | - | - | - | - | - | - | T | C | - | - | - | - | - | - | - | - | - | A | G | T | T | C | A | - | - | - | A | C | T | C | T | T | G | A | T | A | C | G | A | G | T | C | A | T | G | A | A | T | A | G | T | A | T | T | C | G | T | T | T | G | T | G | G | A | A | T | C | A | T | C | T | C | G | G | A | T | T | T | C | T | T | - | - | - | - | - | - | - | - | - | A | A | A | C | G | C | G | G | T | A | G | C | C | C | G | G | #1057084 |
| BA.1.15 | G | G | G | T | G | G | C | - | - | - | C | T | A | C | C | G | C | A | A | - | - | - | - | - | - | - | - | - | A | G | C | C | C | T | G | C | T | G | C | C | A | C | T | - | - | - | - | - | - | T | C | - | - | - | - | - | - | - | - | - | A | G | T | T | C | A | - | - | - | A | C | T | C | T | T | G | A | T | A | C | G | A | G | T | C | A | T | G | A | A | T | A | G | T | A | T | T | C | G | T | T | C | G | T | G | G | A | A | T | C | A | T | C | T | C | G | G | A | T | T | T | C | T | T | - | - | - | - | - | - | - | - | - | A | A | A | C | G | C | G | G | C | A | G | C | C | C | G | G |  |
| mut |  |  |  | T>C |  |  |  |  |  |  |  |  |  |  |  |  |  |  |  |  |  |  |  |  |  |  |  |  |  |  |  |  |  |  |  |  |  |  |  |  |  |  |  |  |  |  |  |  |  |  |  |  |  |  |  |  |  |  |  |  |  |  |  |  |  |  |  |  |  |  |  |  |  |  |  |  |  |  |  |  |  |  |  |  |  |  |  |  |  |  |  |  |  |  |  |  |  |  |  |  |  | C>T |  |  |  |  |  |  |  |  |  |  |  |  |  |  |  |  |  |  |  |  |  |  |  |  |  |  |  |  |  |  |  |  |  |  |  |  |  |  |  | C>T |  |  |  |  |  |  |  |  |

|  |  |  |  |  |  |  |  |  |  |  |  |  |  |  |  |  |  |  |  |  |  |  |  |  |  |  |  |  |  |  |  |  |  |  |  |  |  |  |  |  |  |  |  |  |  |  |  |  |  |  |  |  |  |  |  |  |  |  |  |  |  |  |  |  |  |  |  |  |  |  |  |  |  |  |  |  |  |  |  |  |  |  |  |  |  |  |  |  |  |  |  |  |  |  |  |  |  |  |  |  |  |  |  |  |  |  |  |  |  |  |  |  |  |  |  |  |  |  |  |  |  |  |  |  |  |  |  |  |  |  |  |  |  |  |  |  |  |  |  |  |
| --- | --- | --- | --- | --- | --- | --- | --- | --- | --- | --- | --- | --- | --- | --- | --- | --- | --- | --- | --- | --- | --- | --- | --- | --- | --- | --- | --- | --- | --- | --- | --- | --- | --- | --- | --- | --- | --- | --- | --- | --- | --- | --- | --- | --- | --- | --- | --- | --- | --- | --- | --- | --- | --- | --- | --- | --- | --- | --- | --- | --- | --- | --- | --- | --- | --- | --- | --- | --- | --- | --- | --- | --- | --- | --- | --- | --- | --- | --- | --- | --- | --- | --- | --- | --- | --- | --- | --- | --- | --- | --- | --- | --- | --- | --- | --- | --- | --- | --- | --- | --- | --- | --- | --- | --- | --- | --- | --- | --- | --- | --- | --- | --- | --- | --- | --- | --- | --- | --- | --- | --- | --- | --- | --- | --- | --- | --- | --- | --- | --- | --- | --- | --- | --- | --- | --- | --- | --- | --- | --- | --- |
| pos | 210 | 2470 | 2832 | 3505 | 4015 | 4181 | 5386 | 5584 | 6040 | 6402 | 6513 | 6514 | 6515 | 6638 | 7124 | 7926 | 8393 | 8986 | 9053 | 10449 | 11201 | 11283 | 11284 | 11285 | 11286 | 11287 | 11288 | 11289 | 11290 | 11291 | 11332 | 11537 | 13195 | 14014 | 15240 | 15451 | 16466 | 16726 | 18163 | 19220 | 21005 | 21595 | 21618 | 21762 | 21765 | 21766 | 21767 | 21768 | 21769 | 21770 | 21846 | 21987 | 21988 | 21989 | 21990 | 21991 | 21992 | 21993 | 21994 | 21995 | 22029 | 22030 | 22031 | 22032 | 22033 | 22034 | 22194 | 22195 | 22196 | 22578 | 22599 | 22673 | 22674 | 22679 | 22686 | 22813 | 22882 | 22898 | 22917 | 22992 | 23013 | 23040 | 23048 | 23055 | 23063 | 23075 | 23202 | 23525 | 23599 | 23604 | 23854 | 23948 | 24130 | 24410 | 24424 | 24469 | 24503 | 24826 | 25000 | 25469 | 25584 | 26270 | 26530 | 26577 | 26709 | 26767 | 27259 | 27638 | 27752 | 27807 | 27874 | 28073 | 28248 | 28249 | 28250 | 28251 | 28252 | 28253 | 28271 | 28311 | 28362 | 28363 | 28364 | 28365 | 28366 | 28367 | 28368 | 28369 | 28370 | 28461 | 28881 | 28882 | 28883 | 28916 | 29402 | 29575 | 29576 | 29700 | 29742 |  |
| ref | G | C | A | C | A | G | T | A | C | C | G | T | T | C | C | C | G | C | G | C | A | G | T | T | T | G | T | C | T | G | A | A | T | T | C | G | C | C | A | C | C | C | C | C | T | A | C | A | T | G | C | G | T | G | T | T | T | A | T | T | A | G | T | T | C | A | A | T | T | G | G | T | C | T | C | G | T | G | T | G | A | A | G | A | A | T | C | C | T | C | C | G | C | G | A | T | C | C | C | C | C | C | A | C | G | T | A | T | C | C | C | G | G | A | T | T | T | C | A | C | G | A | G | A | A | C | G | C | A | A | G | G | G | G | G | T | T | A | G |  |
| AY.44 | T | C | A | T | G | T | T | G | T | T | G | T | T | T | T | T | G | T | T | C | G | G | T | T | T | G | T | C | T | G | G | A | T | G | C | A | T | T | A | T | C | C | G | C | T | A | C | A | T | G | C | A | T | G | T | T | T | A | T | T | - | - | - | - | - | - | A | T | T | G | G | T | C | T | C | G | T | G | G | G | A | A | G | A | A | T | C | C | T | G | C | G | C | A | A | T | C | T | C | T | C | C | A | C | G | C | A | C | T | C | T | T | - | - | - | - | - | - | - | C | G | A | G | A | A | C | G | C | A | G | T | G | G | T | T | T | T | G | T |  |
| BA.1.1 | T | C | A | T | G | T | T | G | C | C | - | - | - | C | C | C | A | C | G | A | A | - | - | - | - | - | - | - | - | - | A | G | C | T | T | G | C | C | G | C | T | T | C | T | - | - | - | - | - | - | T | - | - | - | - | - | - | - | - | - | A | G | T | T | C | A | - | - | - | A | A | C | T | C | T | T | G | A | T | A | C | G | A | G | T | C | A | T | G | A | A | T | A | G | T | A | T | C | T | C | T | T | G | G | A | T | C | T | C | T | C | G | G | A | T | T | T | C | T | T | - | - | - | - | - | - | - | - | - | A | A | A | C | G | G | G | C | A | G | #1057278 |
| BA.1.1 | G | T | G | C | A | G | G | A | C | C | - | - | - | C | C | C | A | C | G | A | A | - | - | - | - | - | - | - | - | - | A | G | C | T | T | G | C | C | G | C | T | T | C | T | - | - | - | - | - | - | T | - | - | - | - | - | - | - | - | - | A | G | T | T | C | A | - | - | - | A | A | C | T | C | T | T | G | A | T | A | C | G | A | G | T | C | A | T | G | A | A | T | A | G | T | A | T | C | T | C | T | T | G | G | A | T | C | T | C | T | C | G | G | A | T | T | T | C | T | T | - | - | - | - | - | - | - | - | - | A | A | A | C | G | G | T | T | A | G |  |
| mut |  |  |  |  |  |  |  |  |  |  |  |  |  |  |  |  |  |  |  |  |  |  |  |  |  |  |  |  |  |  |  |  |  |  |  |  |  |  |  |  |  |  |  |  |  |  |  |  |  |  |  |  |  |  |  |  |  |  |  |  |  |  |  |  |  |  |  |  |  |  |  |  |  |  |  |  |  |  |  |  |  |  |  |  |  |  |  |  |  |  |  |  |  |  |  |  |  |  |  |  |  |  |  |  |  |  |  |  |  |  |  |  |  |  |  |  |  |  |  |  |  |  |  |  |  |  |  |  |  |  |  |  |  |  |  | T>G | T>C |  |  |  |

|  |  |  |  |  |  |  |  |  |  |  |  |  |  |  |  |  |  |  |  |  |  |  |  |  |  |  |  |  |  |  |  |  |  |  |  |  |  |  |  |  |  |  |  |  |  |  |  |  |  |  |  |  |  |  |  |  |  |  |  |  |  |  |  |  |  |  |  |  |  |  |  |  |  |  |  |  |  |  |  |  |  |  |  |  |  |  |  |  |  |  |  |  |  |  |
| --- | --- | --- | --- | --- | --- | --- | --- | --- | --- | --- | --- | --- | --- | --- | --- | --- | --- | --- | --- | --- | --- | --- | --- | --- | --- | --- | --- | --- | --- | --- | --- | --- | --- | --- | --- | --- | --- | --- | --- | --- | --- | --- | --- | --- | --- | --- | --- | --- | --- | --- | --- | --- | --- | --- | --- | --- | --- | --- | --- | --- | --- | --- | --- | --- | --- | --- | --- | --- | --- | --- | --- | --- | --- | --- | --- | --- | --- | --- | --- | --- | --- | --- | --- | --- | --- | --- | --- | --- | --- | --- | --- | --- | --- | --- |
| pos | 670 | 1314 | 2061 | 2470 | 2790 | 2832 | 4184 | 4321 | 5386 | 5463 | 6513 | 6514 | 6515 | 8393 | 9344 | 9424 | 9534 | 9866 | 10198 | 10447 | 11283 | 11284 | 11285 | 11286 | 11287 | 11292 | 11293 | 11294 | 11295 | 11296 | 11537 | 12880 | 13195 | 13570 | 14117 | 15240 | 15714 | 17410 | 17615 | 19955 | 20055 | 21618 | 21633 | 21634 | 21635 | 21636 | 21637 | 21638 | 21639 | 21640 | 21641 | 21762 | 21765 | 21766 | 21767 | 21768 | 21769 | 21770 | 21846 | 21987 | 21988 | 21989 | 21990 | 21991 | 21992 | 21993 | 21994 | 21995 | 22194 | 22195 | 22196 | 22200 | 22599 | 22673 | 22688 | 22775 | 22786 | 22792 | 22898 | 23048 | 23202 | 24023 | 24130 | 24503 | 25624 | 26060 | 26530 | 26858 | 27382 | 27383 | 27384 | 29510 | 29755 |  |
| ref | T | C | C | C | C | A | G | C | T | G | G | T | T | G | C | A | C | C | C | G | G | T | T | T | G | G | T | T | T | T | A | C | T | G | C | C | C | C | A | C | A | C | T | A | C | C | C | C | C | T | G | C | T | A | C | A | T | G | C | G | T | G | T | T | T | A | T | T | A | T | T | T | G | T | A | G | A | C | G | G | C | C | C | C | C | C | A | C | G | A | T | A | G |  |
| BA.1.1.15 | T | C | T | T | C | G | G | C | G | T | - | - | - | A | C | A | C | C | C | G | - | - | - | - | - | G | T | T | T | T | G | C | C | A | T | T | C | C | A | C | A | C | T | A | C | C | C | C | C | T | G | T | - | - | - | - | - | - | T | - | - | - | - | - | - | - | - | - | - | - | - | T | A | C | A | G | A | C | A | A | A | C | A | T | C | C | G | C | G | A | T | A | C |  |
| XQ | T | C | T | T | C | G | G | C | T | G | G | T | T | G | T | G | T | T | T | A | G | T | T | T | G | - | - | - | - | - | A | T | T | G | C | C | T | T | G | T | G | T | - | - | - | - | - | - | - | - | - | C | T | A | C | A | T | G | C | A | T | G | T | T | T | A | T | T | A | T | T | G | G | T | G | A | C | C | G | G | C | C | C | C | C | T | A | T | C | T | C | C | G | #1058654 |
| BA.2.9 | G | T | C | C | T | A | A | T | T | G | G | T | T | G | T | G | T | T | T | A | G | T | T | T | G | - | - | - | - | - | A | T | T | G | C | C | T | T | G | T | G | T | - | - | - | - | - | - | - | - | - | C | T | A | C | A | T | G | C | A | T | G | T | T | T | A | T | T | A | T | T | G | G | T | G | A | C | T | G | G | C | T | C | C | T | T | A | T | C | T | C | C | G |  |
| mut |  |  |  |  |  |  |  |  |  |  |  |  |  |  |  |  |  |  |  |  |  |  |  |  |  |  |  |  |  |  |  |  |  |  |  |  |  |  |  |  |  |  |  |  |  |  |  |  |  |  |  |  |  |  |  |  |  |  |  |  |  |  |  |  |  |  |  |  |  |  |  |  |  |  |  |  |  | T>C |  |  |  | T>C |  |  | T>C |  |  |  |  |  |  |  |  |  |

|  |  |  |  |  |  |  |  |  |  |  |  |  |  |  |  |  |  |  |  |  |  |  |  |  |  |  |  |  |  |  |  |  |  |  |  |  |  |  |  |  |  |  |  |  |  |  |  |  |  |  |  |  |  |  |  |  |  |  |  |  |  |  |  |  |  |  |  |  |  |  |  |  |  |  |  |  |  |  |  |  |  |  |  |  |  |  |  |  |  |
| --- | --- | --- | --- | --- | --- | --- | --- | --- | --- | --- | --- | --- | --- | --- | --- | --- | --- | --- | --- | --- | --- | --- | --- | --- | --- | --- | --- | --- | --- | --- | --- | --- | --- | --- | --- | --- | --- | --- | --- | --- | --- | --- | --- | --- | --- | --- | --- | --- | --- | --- | --- | --- | --- | --- | --- | --- | --- | --- | --- | --- | --- | --- | --- | --- | --- | --- | --- | --- | --- | --- | --- | --- | --- | --- | --- | --- | --- | --- | --- | --- | --- | --- | --- | --- | --- | --- | --- | --- | --- |
| pos | 44 | 670 | 2470 | 2790 | 2832 | 3264 | 4184 | 4321 | 5386 | 6513 | 6514 | 6515 | 8393 | 9344 | 9424 | 9534 | 9866 | 10198 | 10447 | 11283 | 11284 | 11285 | 11286 | 11287 | 11292 | 11293 | 11294 | 11295 | 11296 | 11537 | 12880 | 13195 | 15240 | 15714 | 16342 | 17410 | 19518 | 19955 | 20055 | 21595 | 21618 | 21633 | 21634 | 21635 | 21636 | 21637 | 21638 | 21639 | 21640 | 21641 | 21762 | 21765 | 21766 | 21767 | 21768 | 21769 | 21770 | 21846 | 21987 | 21988 | 21989 | 21990 | 21991 | 21992 | 21993 | 21994 | 21995 | 22194 | 22195 | 22196 | 22200 | 22599 | 22673 | 22688 | 22775 | 22786 | 22898 | 23048 | 23202 | 24130 | 24503 | 26060 | 26530 | 26858 | 27382 | 27383 | 27384 | 29510 |  |
| ref | C | T | C | C | A | C | G | C | T | G | T | T | G | C | A | C | C | C | G | G | T | T | T | G | G | T | T | T | T | A | C | T | C | C | T | C | G | C | A | C | C | T | A | C | C | C | C | C | T | G | C | T | A | C | A | T | G | C | G | T | G | T | T | T | A | T | T | A | T | T | T | G | T | A | G | A | G | G | C | C | C | C | A | C | G | A | T | A |  |
| BA.1.1 | C | T | T | C | G | T | G | C | G | - | - | - | A | C | A | C | C | C | G | - | - | - | - | - | G | T | T | T | T | G | C | C | T | C | T | C | G | C | A | T | C | T | A | C | C | C | C | C | T | G | T | - | - | - | - | - | - | T | - | - | - | - | - | - | - | - | - | - | - | - | T | A | C | A | G | A | A | A | A | A | T | C | G | C | G | A | T | A |  |
| BA.2 | C | T | T | C | G | T | G | C | G | - | - | - | A | T | G | T | T | T | A | G | T | T | T | G | - | - | - | - | - | A | T | T | C | T | C | T | T | T | G | C | T | - | - | - | - | - | - | - | - | - | C | T | A | C | A | T | G | C | A | T | G | T | T | T | A | T | T | A | T | T | G | G | T | G | A | C | G | G | C | C | C | T | A | T | C | T | C | C | #1059681 |
| BA.2.10 | T | G | C | T | A | C | A | T | T | G | T | T | G | T | G | T | T | T | A | G | T | T | T | G | - | - | - | - | - | A | T | T | C | T | C | T | G | T | G | C | T | - | - | - | - | - | - | - | - | - | C | T | A | C | A | T | G | C | A | T | G | T | T | T | A | T | T | A | T | T | G | G | T | G | A | C | G | G | C | C | C | T | A | T | C | T | C | C |  |
| mut |  |  |  |  |  |  |  |  |  |  |  |  |  |  |  |  |  |  |  |  |  |  |  |  |  |  |  |  |  |  |  |  |  |  |  |  | G>T |  |  |  |  |  |  |  |  |  |  |  |  |  |  |  |  |  |  |  |  |  |  |  |  |  |  |  |  |  |  |  |  |  |  |  |  |  |  |  |  |  |  |  |  |  |  |  |  |  |  |  |  |

|  |  |  |  |  |  |  |  |  |  |  |  |  |  |  |  |  |  |  |  |  |  |  |  |  |  |  |  |  |  |  |  |  |  |  |  |  |  |  |  |  |  |  |  |  |  |  |  |  |  |  |  |  |  |  |  |  |  |  |  |  |  |  |  |  |  |  |  |  |  |  |  |  |  |  |  |  |  |  |  |  |  |  |  |  |  |  |  |  |  |  |  |  |  |  |  |  |  |  |  |  |  |  |  |  |  |  |  |  |  |  |  |  |  |  |  |  |  |  |  |  |  |  |  |  |  |  |  |  |  |  |  |  |  |  |
| --- | --- | --- | --- | --- | --- | --- | --- | --- | --- | --- | --- | --- | --- | --- | --- | --- | --- | --- | --- | --- | --- | --- | --- | --- | --- | --- | --- | --- | --- | --- | --- | --- | --- | --- | --- | --- | --- | --- | --- | --- | --- | --- | --- | --- | --- | --- | --- | --- | --- | --- | --- | --- | --- | --- | --- | --- | --- | --- | --- | --- | --- | --- | --- | --- | --- | --- | --- | --- | --- | --- | --- | --- | --- | --- | --- | --- | --- | --- | --- | --- | --- | --- | --- | --- | --- | --- | --- | --- | --- | --- | --- | --- | --- | --- | --- | --- | --- | --- | --- | --- | --- | --- | --- | --- | --- | --- | --- | --- | --- | --- | --- | --- | --- | --- | --- | --- | --- | --- | --- | --- | --- | --- | --- | --- | --- | --- | --- | --- | --- | --- | --- | --- | --- | --- |
| pos | 210 | 300 | 2470 | 2832 | 4181 | 5386 | 6402 | 6513 | 6514 | 6515 | 7124 | 8393 | 8829 | 8986 | 9053 | 10449 | 11120 | 11201 | 11283 | 11284 | 11285 | 11286 | 11287 | 11288 | 11289 | 11290 | 11291 | 11332 | 11537 | 12793 | 13195 | 13428 | 15240 | 15451 | 16466 | 18163 | 19006 | 19220 | 21595 | 21618 | 21762 | 21765 | 21766 | 21767 | 21768 | 21769 | 21770 | 21987 | 21988 | 21989 | 21990 | 21991 | 21992 | 21993 | 21994 | 21995 | 22029 | 22030 | 22031 | 22032 | 22033 | 22034 | 22194 | 22195 | 22196 | 22578 | 22599 | 22673 | 22674 | 22679 | 22686 | 22813 | 22882 | 22898 | 22917 | 22992 | 23013 | 23040 | 23048 | 23055 | 23063 | 23075 | 23202 | 23525 | 23587 | 23599 | 23604 | 23854 | 23948 | 24130 | 24410 | 24424 | 24469 | 24503 | 25000 | 25469 | 25584 | 26270 | 26530 | 26577 | 26709 | 26767 | 27259 | 27638 | 27752 | 27768 | 27807 | 27874 | 28248 | 28249 | 28250 | 28251 | 28252 | 28253 | 28271 | 28311 | 28362 | 28363 | 28364 | 28365 | 28366 | 28367 | 28368 | 28369 | 28370 | 28461 | 28881 | 28882 | 28883 | 28916 | 29362 | 29402 | 29742 |  |
| ref | G | C | C | A | G | T | C | G | T | T | C | G | C | C | G | C | A | A | G | T | T | T | G | T | C | T | G | A | A | G | T | A | C | G | C | A | G | C | C | C | C | T | A | C | A | T | G | G | T | G | T | T | T | A | T | T | A | G | T | T | C | A | A | T | T | G | G | T | C | T | C | G | T | G | T | G | A | A | G | A | A | T | C | C | G | T | C | C | G | C | G | A | T | C | C | C | C | C | A | C | G | T | A | T | C | T | C | C | G | A | T | T | T | C | A | C | G | A | G | A | A | C | G | C | A | A | G | G | G | G | C | G | G |  |
| AY.119.2 | T | C | C | A | T | T | T | G | T | T | T | G | T | T | T | C | A | G | G | T | T | T | G | T | C | T | G | G | A | T | T | G | C | A | T | A | T | T | C | G | C | T | A | C | A | T | G | A | T | G | T | T | T | A | T | T | - | - | - | - | - | - | A | T | T | G | G | T | C | T | C | G | T | G | G | G | A | A | G | A | A | T | C | C | C | T | G | C | G | C | A | A | T | C | C | T | C | C | A | C | G | C | A | C | T | T | C | T | - | - | - | - | - | - | - | C | G | A | G | A | A | C | G | C | A | G | T | G | G | T | T | T | T |  |
| BA.1.1 | T | T | C | A | T | T | T | G | T | T | T | G | T | T | T | C | A | G | G | T | T | T | G | T | C | T | G | G | A | T | T | G | C | A | T | A | T | T | C | G | C | T | A | C | A | T | G | A | T | G | T | T | T | A | T | T | - | - | - | - | - | - | A | T | T | A | A | C | T | C | T | T | G | A | T | A | C | G | A | G | T | C | A | T | G | G | A | A | T | A | G | T | A | T | T | C | T | T | G | G | A | T | C | T | C | C | T | C | G | A | T | T | T | C | T | T | - | - | - | - | - | - | - | - | - | A | A | A | C | G | C | G | G | #1060983 |
| BA.1.1 | G | C | T | G | G | G | C | - | - | - | C | A | C | C | G | A | G | A | - | - | - | - | - | - | - | - | - | A | G | G | C | A | T | G | C | G | G | C | T | C | T | - | - | - | - | - | - | - | - | - | - | - | - | - | - | - | A | G | T | T | C | A | - | - | - | A | A | C | T | C | T | T | G | A | T | A | C | G | A | G | T | C | A | T | G | G | A | A | T | A | G | T | A | T | T | C | T | T | G | G | A | T | C | T | C | C | T | C | G | A | T | T | T | C | T | T | - | - | - | - | - | - | - | - | - | A | A | A | C | G | C | G | G |  |
| mut |  | C>T |  |  |  |  |  |  |  |  |  |  |  |  |  |  |  |  |  |  |  |  |  |  |  |  |  |  |  |  |  |  |  |  |  |  |  |  |  |  |  |  |  |  |  |  |  |  |  |  |  |  |  |  |  |  |  |  |  |  |  |  |  |  |  |  |  |  |  |  |  |  |  |  |  |  |  |  |  |  |  |  |  |  |  |  |  |  |  |  |  |  |  |  |  |  |  |  |  |  |  |  |  |  |  |  |  |  |  |  |  |  |  |  |  |  |  |  |  |  |  |  |  |  |  |  |  |  |  |  |  |  |  |  |

|  |  |  |  |  |  |  |  |  |  |  |  |  |  |  |  |  |  |  |  |  |  |  |  |  |  |  |  |  |  |  |  |  |  |  |  |  |  |  |  |  |  |  |  |  |  |  |  |  |  |  |  |  |  |  |  |  |  |  |  |  |  |  |  |  |  |  |  |  |  |  |  |  |  |  |  |  |  |  |  |  |  |  |  |  |  |  |  |  |  |  |  |
| --- | --- | --- | --- | --- | --- | --- | --- | --- | --- | --- | --- | --- | --- | --- | --- | --- | --- | --- | --- | --- | --- | --- | --- | --- | --- | --- | --- | --- | --- | --- | --- | --- | --- | --- | --- | --- | --- | --- | --- | --- | --- | --- | --- | --- | --- | --- | --- | --- | --- | --- | --- | --- | --- | --- | --- | --- | --- | --- | --- | --- | --- | --- | --- | --- | --- | --- | --- | --- | --- | --- | --- | --- | --- | --- | --- | --- | --- | --- | --- | --- | --- | --- | --- | --- | --- | --- | --- | --- | --- | --- | --- |
| pos | 670 | 2470 | 2790 | 2832 | 4184 | 4321 | 4893 | 5386 | 6026 | 6513 | 6514 | 6515 | 8393 | 9344 | 9424 | 9534 | 9866 | 10198 | 10447 | 11283 | 11284 | 11285 | 11286 | 11287 | 11292 | 11293 | 11294 | 11295 | 11296 | 11537 | 12880 | 13195 | 15240 | 15714 | 17410 | 19955 | 20055 | 21618 | 21633 | 21634 | 21635 | 21636 | 21637 | 21638 | 21639 | 21640 | 21641 | 21762 | 21765 | 21766 | 21767 | 21768 | 21769 | 21770 | 21846 | 21987 | 21988 | 21989 | 21990 | 21991 | 21992 | 21993 | 21994 | 21995 | 22194 | 22195 | 22196 | 22200 | 22599 | 22673 | 22688 | 22775 | 22786 | 22898 | 23048 | 23202 | 23734 | 24130 | 24503 | 25096 | 25224 | 26060 | 26424 | 26522 | 26530 | 26858 | 27382 | 27383 | 27384 | 29510 |  |
| ref | T | C | C | A | G | C | C | T | C | G | T | T | G | C | A | C | C | C | G | G | T | T | T | G | G | T | T | T | T | A | C | T | C | C | C | C | A | C | T | A | C | C | C | C | C | T | G | C | T | A | C | A | T | G | C | G | T | G | T | T | T | A | T | T | A | T | T | T | G | T | A | G | A | G | G | C | A | C | C | C | T | C | T | C | A | C | G | A | T | A |  |
| BA.2 | G | C | T | A | A | T | T | T | T | G | T | T | G | T | G | T | T | T | A | G | T | T | T | G | - | - | - | - | - | A | T | T | C | T | T | T | G | T | - | - | - | - | - | - | - | - | - | C | T | A | C | A | T | G | C | A | T | G | T | T | T | A | T | T | A | T | T | G | G | T | G | A | C | G | G | C | G | C | C | T | C | T | T | C | A | T | C | T | C | C |  |
| BA.2 | G | C | T | A | A | T | T | T | C | G | T | T | G | T | G | T | T | T | A | G | T | T | T | G | - | - | - | - | - | A | T | T | C | T | T | T | G | T | - | - | - | - | - | - | - | - | - | C | T | A | C | A | T | G | C | A | T | G | T | T | T | A | T | T | A | T | T | G | G | T | G | A | C | G | G | C | G | C | C | T | C | C | C | T | G | C | G | A | T | A | #1062830 |
| BA.1.1 | T | T | C | G | G | C | C | G | C | - | - | - | A | C | A | C | C | C | G | - | - | - | - | - | G | T | T | T | T | G | C | C | T | C | C | C | A | C | T | A | C | C | C | C | C | T | G | T | - | - | - | - | - | - | T | - | - | - | - | - | - | - | - | - | - | - | - | T | A | C | A | G | A | A | A | A | A | A | T | C | T | C | C | T | G | C | G | A | T | A |  |
| mut |  |  |  |  |  |  |  |  | T>C |  |  |  |  |  |  |  |  |  |  |  |  |  |  |  |  |  |  |  |  |  |  |  |  |  |  |  |  |  |  |  |  |  |  |  |  |  |  |  |  |  |  |  |  |  |  |  |  |  |  |  |  |  |  |  |  |  |  |  |  |  |  |  |  |  |  |  |  |  |  |  |  |  |  |  |  |  |  |  |  |  |  |

|  |  |  |  |  |  |  |  |  |  |  |  |  |  |  |  |  |  |  |  |  |  |  |  |  |  |  |  |  |
| --- | --- | --- | --- | --- | --- | --- | --- | --- | --- | --- | --- | --- | --- | --- | --- | --- | --- | --- | --- | --- | --- | --- | --- | --- | --- | --- | --- | --- |
| pos | 2470 | 3418 | 9868 | 11123 | 18555 | 20629 | 21595 | 22194 | 22195 | 22196 | 22197 | 22198 | 22202 | 22204 | 22599 | 28362 | 28363 | 28364 | 28365 | 28366 | 28367 | 28368 | 28369 | 28370 | 29732 | 29779 | 29781 |  |
| ref | C | G | T | G | C | C | C | A | T | T | T | A | C | T | G | G | A | G | A | A | C | G | C | A | C | G | G |  |
| BA.1.1 | T | T | T | A | C | T | T | - | - | - | T | A | C | T | A | - | - | - | - | - | - | - | - | - | C | G | G |  |
| BA.1.1 | T | T | T | A | C | T | T | A | G | T | G | C | A | C | A | - | - | - | - | - | - | - | - | - | T | G | G | #1063584 |
| BA.1 | C | G | C | G | T | C | C | A | G | T | G | C | A | C | G | G | A | G | A | A | C | G | C | A | T | - | - |  |
| mut |  |  |  |  |  |  |  |  |  |  |  |  |  |  | G>A | G>- | A>- | G>- | A>- | A>- | C>- | G>- | C>- | A>- |  | ->G | ->G |  |

|  |  |  |  |  |  |  |  |  |  |  |  |  |  |  |  |  |  |  |  |  |  |  |  |  |  |  |  |  |  |  |  |  |  |  |  |  |  |  |  |  |  |  |  |  |  |  |  |  |  |  |  |  |  |  |  |  |  |  |  |  |  |  |  |  |  |  |  |  |  |  |  |  |  |  |  |  |  |  |  |  |  |  |  |  |  |  |  |  |  |  |  |  |  |  |  |
| --- | --- | --- | --- | --- | --- | --- | --- | --- | --- | --- | --- | --- | --- | --- | --- | --- | --- | --- | --- | --- | --- | --- | --- | --- | --- | --- | --- | --- | --- | --- | --- | --- | --- | --- | --- | --- | --- | --- | --- | --- | --- | --- | --- | --- | --- | --- | --- | --- | --- | --- | --- | --- | --- | --- | --- | --- | --- | --- | --- | --- | --- | --- | --- | --- | --- | --- | --- | --- | --- | --- | --- | --- | --- | --- | --- | --- | --- | --- | --- | --- | --- | --- | --- | --- | --- | --- | --- | --- | --- | --- | --- | --- | --- | --- | --- |
| pos | 44 | 670 | 2790 | 2832 | 3241 | 4184 | 4197 | 4321 | 5386 | 5924 | 6513 | 6514 | 6515 | 6988 | 8393 | 9344 | 9424 | 9534 | 9866 | 10198 | 10447 | 11283 | 11284 | 11285 | 11286 | 11287 | 11292 | 11293 | 11294 | 11295 | 11296 | 11537 | 12309 | 12880 | 13195 | 15240 | 15714 | 16342 | 17410 | 19955 | 20055 | 21618 | 21633 | 21634 | 21635 | 21636 | 21637 | 21638 | 21639 | 21640 | 21641 | 21762 | 21765 | 21766 | 21767 | 21768 | 21769 | 21770 | 21846 | 21987 | 21988 | 21989 | 21990 | 21991 | 21992 | 21993 | 21994 | 21995 | 22194 | 22195 | 22196 | 22200 | 22673 | 22688 | 22775 | 22786 | 22813 | 23048 | 23202 | 23664 | 24130 | 24503 | 26060 | 26530 | 26858 | 27382 | 27383 | 27384 | 27874 | 27875 | 27881 | 27882 | 27883 | 29510 |  |
| ref | C | T | C | A | C | G | A | C | T | G | G | T | T | T | G | C | A | C | C | C | G | G | T | T | T | G | G | T | T | T | T | A | A | C | T | C | C | T | C | C | A | C | T | A | C | C | C | C | C | T | G | C | T | A | C | A | T | G | C | G | T | G | T | T | T | A | T | T | A | T | T | T | T | A | G | A | G | G | C | C | C | C | C | A | C | G | A | T | C | T | C | G | C | A |  |
| BA.2.10 | T | G | T | A | C | A | C | T | T | G | G | T | T | T | G | T | G | T | T | T | A | G | T | T | T | G | - | - | - | - | - | A | A | T | T | C | T | C | T | T | G | T | - | - | - | - | - | - | - | - | - | C | T | A | C | A | T | G | C | A | T | G | T | T | T | A | T | T | A | T | T | G | T | G | A | C | T | G | C | C | C | C | T | A | T | C | T | C | C | T | C | G | C | C |  |
| BA.2.10 | T | G | T | A | C | A | C | T | T | G | G | T | T | T | G | T | G | T | T | T | A | G | T | T | T | G | - | - | - | - | - | A | A | T | T | C | T | C | T | T | G | T | - | - | - | - | - | - | - | - | - | C | T | A | C | A | T | G | C | A | T | G | T | T | T | A | T | T | A | T | T | G | T | G | A | C | T | G | C | C | C | C | T | A | T | C | T | C | T | C | T | C | T | C | #1065871 |
| BA.1.17.2 | C | T | C | G | T | G | A | C | G | A | - | - | - | C | A | C | A | C | C | C | G | - | - | - | - | - | G | T | T | T | T | G | G | C | C | T | C | T | C | C | A | C | T | A | C | C | C | C | C | T | G | T | - | - | - | - | - | - | T | - | - | - | - | - | - | - | - | - | - | - | - | T | C | A | G | A | G | A | A | T | A | T | C | G | C | G | A | T | T | C | T | C | T | A |  |
| mut |  |  |  |  |  |  |  |  |  |  |  |  |  |  |  |  |  |  |  |  |  |  |  |  |  |  |  |  |  |  |  |  |  |  |  |  |  |  |  |  |  |  |  |  |  |  |  |  |  |  |  |  |  |  |  |  |  |  |  |  |  |  |  |  |  |  |  |  |  |  |  |  |  |  |  |  |  |  |  |  |  |  |  |  |  |  |  |  |  |  |  |  |  | A>C |  |

|  |  |  |  |  |  |  |  |  |  |  |  |  |  |  |  |  |  |
| --- | --- | --- | --- | --- | --- | --- | --- | --- | --- | --- | --- | --- | --- | --- | --- | --- | --- |
| pos | 906 | 2470 | 5515 | 10332 | 14599 | 21595 | 22194 | 22195 | 22196 | 22197 | 22198 | 22202 | 22204 | 22599 | 22813 | 23513 |  |
| ref | C | C | G | C | C | C | A | T | T | T | A | C | T | G | G | A |  |
| BA.1 | C | C | T | T | T | C | A | G | T | G | C | A | C | G | T | G |  |
| BA.1 | T | C | T | T | T | C | G | T | G | T | A | C | T | G | G | A | #1067541 |
| BA.1.1 | C | T | G | C | C | T | G | T | G | T | A | C | T | A | G | A |  |
| mut | C>T |  |  |  |  |  |  |  |  |  |  |  |  | A>G |  |  |  |

|  |  |  |  |  |  |  |  |  |  |  |  |  |  |  |  |  |  |
| --- | --- | --- | --- | --- | --- | --- | --- | --- | --- | --- | --- | --- | --- | --- | --- | --- | --- |
| pos | 1251 | 2007 | 10078 | 18457 | 18788 | 19073 | 22194 | 22195 | 22196 | 22197 | 22198 | 22202 | 22204 | 22813 | 29260 | 29700 |  |
| ref | C | T | C | C | C | A | A | T | T | T | A | C | T | G | G | A |  |
| BA.1.1 | T | C | C | T | C | G | A | G | T | G | C | A | C | G | G | G |  |
| BA.1.1 | T | C | C | T | T | G | - | - | - | T | A | C | T | T | T | G | #1067666 |
| BA.1.1 | C | T | T | C | C | A | - | - | - | T | A | C | T | T | T | A |  |
| mut |  |  |  |  | C>T |  |  |  |  |  |  |  |  |  |  | A>G |  |

|  |  |  |  |  |  |  |  |  |  |  |  |  |  |  |  |
| --- | --- | --- | --- | --- | --- | --- | --- | --- | --- | --- | --- | --- | --- | --- | --- |
| pos | 8692 | 9693 | 11956 | 22194 | 22195 | 22196 | 22197 | 22198 | 22202 | 22204 | 25626 | 26733 | 27561 | 28817 |  |
| ref | C | C | C | A | T | T | T | A | C | T | C | T | G | G |  |
| BA.1 | C | C | C | A | G | T | G | C | A | C | C | T | G | G |  |
| BA.1 | T | T | T | A | G | T | G | C | A | C | A | C | T | T | #1067976 |
| BA.1 | C | T | C | - | - | - | T | A | C | T | A | C | T | T |  |
| mut | C>T | C>T | C>T |  |  |  |  |  |  |  |  |  |  |  |  |

|  |  |  |  |  |  |  |  |  |  |  |  |  |  |  |  |  |
| --- | --- | --- | --- | --- | --- | --- | --- | --- | --- | --- | --- | --- | --- | --- | --- | --- |
| pos | 679 | 1344 | 1558 | 5576 | 6884 | 13019 | 22194 | 22195 | 22196 | 22197 | 22198 | 22202 | 22204 | 23938 | 23948 |  |
| ref | C | T | A | A | G | C | A | T | T | T | A | C | T | A | G |  |
| BA.1.1.18 | T | C | A | G | A | T | - | - | - | T | A | C | T | A | G |  |
| BA.1.1.18 | T | C | G | G | A | T | A | G | T | G | C | A | C | T | T | #1072020 |
| BA.1.1.18 | C | T | A | A | G | C | A | G | T | G | C | A | C | A | T |  |
| mut |  |  | A>G |  |  |  |  |  |  |  |  |  |  | A>T |  |  |

|  |  |  |  |  |  |  |  |  |  |  |  |  |  |  |  |  |
| --- | --- | --- | --- | --- | --- | --- | --- | --- | --- | --- | --- | --- | --- | --- | --- | --- |
| pos | 6906 | 11572 | 18471 | 21595 | 22194 | 22195 | 22196 | 22197 | 22198 | 22202 | 22204 | 24905 | 25452 | 26951 | 29721 |  |
| ref | C | C | T | C | A | T | T | T | A | C | T | A | C | G | C |  |
| BA.1.1 | C | C | C | T | A | G | T | G | C | A | C | A | C | G | C |  |
| BA.1.1 | T | T | C | C | A | G | T | G | C | A | C | G | T | T | T | #1072455 |
| BA.1.1 | T | T | T | C | - | - | - | T | A | C | T | G | T | T | T |  |
| mut | C>T | C>T |  | T>C |  |  |  |  |  |  |  |  |  |  |  |  |

|  |  |  |  |  |  |  |  |  |  |  |  |  |
| --- | --- | --- | --- | --- | --- | --- | --- | --- | --- | --- | --- | --- |
| pos | 5175 | 6226 | 8032 | 12525 | 22194 | 22195 | 22196 | 22197 | 22198 | 22202 | 22204 |  |
| ref | C | T | A | C | A | T | T | T | A | C | T |  |
| BA.1.1 | T | C | G | T | - | - | - | T | A | C | T |  |
| BA.1.1 | T | C | G | T | A | G | T | G | C | A | C | #1075207 |
| BA.1.1 | C | T | A | C | A | G | T | G | C | A | C |  |
| mut |  |  |  |  |  |  |  |  |  |  |  |  |

|  |  |  |  |  |  |  |  |  |  |  |  |  |  |  |  |  |  |
| --- | --- | --- | --- | --- | --- | --- | --- | --- | --- | --- | --- | --- | --- | --- | --- | --- | --- |
| pos | 1710 | 3579 | 6532 | 6659 | 7926 | 9724 | 15999 | 21595 | 22194 | 22195 | 22196 | 22197 | 22198 | 22202 | 22204 | 25720 |  |
| ref | C | T | G | A | C | C | A | C | A | T | T | T | A | C | T | G |  |
| BA.1.1 | T | C | T | A | C | T | C | T | - | - | - | T | A | C | T | T |  |
| BA.1.1 | T | C | T | A | C | T | A | T | A | G | T | G | C | A | C | T | #1078172 |
| BA.1.1.18 | C | T | G | G | T | C | A | C | A | G | T | G | C | A | C | G |  |
| mut |  |  |  |  |  |  |  | C>T |  |  |  |  |  |  |  | G>T |  |

|  |  |  |  |  |  |  |  |  |  |  |  |  |  |  |  |  |  |
| --- | --- | --- | --- | --- | --- | --- | --- | --- | --- | --- | --- | --- | --- | --- | --- | --- | --- |
| pos | 2662 | 5115 | 7926 | 14307 | 15451 | 18298 | 21156 | 22194 | 22195 | 22196 | 22197 | 22198 | 22202 | 22204 | 26111 | 26209 |  |
| ref | C | C | C | T | G | A | C | A | T | T | T | A | C | T | C | G |  |
| BA.1.1.18 | C | T | T | C | A | G | C | A | G | T | G | C | A | C | T | A |  |
| BA.1.1.18 | C | T | T | C | A | A | T | - | - | - | T | A | C | T | T | G | #1078501 |
| BA.1.1 | T | C | C | T | G | A | T | - | - | - | T | A | C | T | C | G |  |
| mut |  |  |  |  |  |  |  |  |  |  |  |  |  |  | C>T |  |  |

|  |  |  |  |  |  |  |  |  |  |  |  |  |  |  |  |
| --- | --- | --- | --- | --- | --- | --- | --- | --- | --- | --- | --- | --- | --- | --- | --- |
| pos | 991 | 2470 | 6759 | 13900 | 17280 | 21595 | 22194 | 22195 | 22196 | 22197 | 22198 | 22202 | 22204 | 22599 |  |
| ref | G | C | G | G | G | C | A | T | T | T | A | C | T | G |  |
| BA.1.1 | A | T | C | T | T | T | - | - | - | T | A | C | T | A |  |
| BA.1.1 | A | T | C | T | T | T | A | G | T | G | C | A | C | A | #1081228 |
| BA.1 | G | C | G | G | G | C | A | G | T | G | C | A | C | G |  |
| mut |  |  |  |  |  |  |  |  |  |  |  |  |  | G>A |  |

|  |  |  |  |  |  |  |  |  |  |  |  |  |  |  |
| --- | --- | --- | --- | --- | --- | --- | --- | --- | --- | --- | --- | --- | --- | --- |
| pos | 1342 | 5079 | 12624 | 18888 | 21036 | 22194 | 22195 | 22196 | 22197 | 22198 | 22202 | 22204 | 23010 |  |
| ref | C | C | C | C | C | A | T | T | T | A | C | T | T |  |
| BA.1.1 | T | T | T | G | T | - | - | - | T | A | C | T | C |  |
| BA.1.1 | T | T | T | G | T | A | G | T | G | C | A | C | C | #1081265 |
| BA.1.1 | C | C | C | C | C | A | G | T | G | C | A | C | T |  |
| mut |  |  |  |  |  |  |  |  |  |  |  |  | T>C |  |

|  |  |  |  |  |  |  |  |  |  |  |  |  |  |  |  |  |  |  |  |  |  |  |  |  |  |  |  |  |  |  |  |  |  |  |  |  |  |  |  |  |  |  |  |  |  |  |  |  |  |  |  |  |  |  |  |  |  |  |  |  |  |  |  |  |  |  |  |  |  |  |  |  |  |  |  |  |  |  |  |  |  |  |  |  |  |  |  |  |  |  |
| --- | --- | --- | --- | --- | --- | --- | --- | --- | --- | --- | --- | --- | --- | --- | --- | --- | --- | --- | --- | --- | --- | --- | --- | --- | --- | --- | --- | --- | --- | --- | --- | --- | --- | --- | --- | --- | --- | --- | --- | --- | --- | --- | --- | --- | --- | --- | --- | --- | --- | --- | --- | --- | --- | --- | --- | --- | --- | --- | --- | --- | --- | --- | --- | --- | --- | --- | --- | --- | --- | --- | --- | --- | --- | --- | --- | --- | --- | --- | --- | --- | --- | --- | --- | --- | --- | --- | --- | --- | --- | --- |
| pos | 670 | 2790 | 2832 | 2862 | 4184 | 4321 | 5386 | 5672 | 5924 | 6513 | 6514 | 6515 | 8393 | 9344 | 9424 | 9534 | 9866 | 10198 | 10447 | 11283 | 11284 | 11285 | 11286 | 11287 | 11292 | 11293 | 11294 | 11295 | 11296 | 11537 | 12880 | 13195 | 15240 | 15714 | 17410 | 19855 | 19955 | 20055 | 21618 | 21633 | 21634 | 21635 | 21636 | 21637 | 21638 | 21639 | 21640 | 21641 | 21762 | 21765 | 21766 | 21767 | 21768 | 21769 | 21770 | 21846 | 21987 | 21988 | 21989 | 21990 | 21991 | 21992 | 21993 | 21994 | 21995 | 22195 | 22197 | 22198 | 22200 | 22202 | 22204 | 22673 | 22688 | 22775 | 22786 | 22792 | 22898 | 23048 | 23202 | 24130 | 24503 | 25672 | 26060 | 26530 | 26858 | 27382 | 27383 | 27384 | 29510 |  |
| ref | T | C | A | C | G | C | T | C | G | G | T | T | G | C | A | C | C | C | G | G | T | T | T | G | G | T | T | T | T | A | C | T | C | C | C | A | C | A | C | T | A | C | C | C | C | C | T | G | C | T | A | C | A | T | G | C | G | T | G | T | T | T | A | T | T | T | T | A | T | C | T | T | A | G | A | C | G | G | C | C | C | C | C | A | C | G | A | T | A |  |
| BA.1.17 | T | C | G | C | G | C | G | T | A | - | - | - | A | C | A | C | C | C | G | - | - | - | - | - | G | T | T | T | T | G | C | C | T | C | C | A | C | A | C | T | A | C | C | C | C | C | T | G | T | - | - | - | - | - | - | T | - | - | - | - | - | - | - | - | - | G | G | C | T | A | C | C | A | G | A | C | A | A | A | A | T | C | C | G | C | G | A | T | A |  |
| XG | T | C | G | C | G | C | G | T | A | G | T | T | G | T | G | T | T | T | A | G | T | T | T | G | - | - | - | - | - | A | T | T | C | T | T | G | T | G | T | - | - | - | - | - | - | - | - | - | C | T | A | C | A | T | G | C | A | T | G | T | T | T | A | T | T | T | T | A | G | C | T | T | G | A | C | T | G | G | C | C | C | T | T | A | T | C | T | C | C | #1083412 |
| BA.2 | G | T | A | T | A | T | T | C | G | G | T | T | G | T | G | T | T | T | A | G | T | T | T | G | - | - | - | - | - | A | T | T | C | T | T | A | T | G | T | - | - | - | - | - | - | - | - | - | C | T | A | C | A | T | G | C | A | T | G | T | T | T | A | T | T | T | T | A | G | C | T | T | G | A | C | C | G | G | C | C | C | C | T | A | T | C | T | C | C |  |
| mut |  |  |  |  |  |  |  |  |  |  |  |  |  |  |  |  |  |  |  |  |  |  |  |  |  |  |  |  |  |  |  |  |  |  |  | A>G |  |  |  |  |  |  |  |  |  |  |  |  |  |  |  |  |  |  |  |  |  |  |  |  |  |  |  |  |  |  |  |  |  |  |  |  |  |  |  | C>T |  |  |  |  |  | C>T |  |  |  |  |  |  |  |  |

|  |  |  |  |  |  |  |  |  |  |  |  |  |  |  |  |  |  |  |  |  |  |  |  |
| --- | --- | --- | --- | --- | --- | --- | --- | --- | --- | --- | --- | --- | --- | --- | --- | --- | --- | --- | --- | --- | --- | --- | --- |
| pos | 686 | 687 | 688 | 689 | 690 | 691 | 692 | 693 | 694 | 4084 | 4399 | 7926 | 14940 | 21102 | 21771 | 22194 | 22195 | 22196 | 22197 | 22198 | 22202 | 22204 |  |
| ref | A | A | G | T | C | A | T | T | T | C | G | C | A | T | T | A | T | T | T | A | C | T |  |
| BA.1.1 | A | A | G | T | C | A | T | T | T | T | T | C | G | C | C | - | - | - | T | A | C | T |  |
| BA.1.1 | - | - | - | - | - | - | - | - | - | T | T | C | G | C | C | A | G | T | G | C | A | C | #1086738 |
| BA.1.1.18 | A | A | G | T | C | A | T | T | T | C | G | T | A | T | T | A | G | T | G | C | A | C |  |
| mut | A>- | A>- | G>- | T>- | C>- | A>- | T>- | T>- | T>- |  |  |  |  |  |  |  |  |  |  |  |  |  |  |

|  |  |  |  |  |  |  |  |  |  |  |  |  |  |  |
| --- | --- | --- | --- | --- | --- | --- | --- | --- | --- | --- | --- | --- | --- | --- |
| pos | 475 | 4543 | 11471 | 18329 | 20629 | 22194 | 22195 | 22196 | 22197 | 22198 | 22202 | 22204 | 22882 |  |
| ref | C | C | C | C | C | A | T | T | T | A | C | T | T |  |
| BA.1.1 | T | C | T | T | T | A | G | T | G | C | A | C | G |  |
| BA.1.1 | T | C | T | T | T | - | - | - | T | A | C | T | G | #1086794 |
| BA.1.1 | C | T | C | C | C | - | - | - | T | A | C | T | T |  |
| mut |  |  |  |  |  |  |  |  |  |  |  |  | T>G |  |

|  |  |  |  |  |  |  |  |  |  |  |  |  |  |  |  |  |  |  |  |  |  |
| --- | --- | --- | --- | --- | --- | --- | --- | --- | --- | --- | --- | --- | --- | --- | --- | --- | --- | --- | --- | --- | --- |
| pos | 493 | 809 | 2470 | 5515 | 7926 | 16596 | 19572 | 19866 | 20104 | 22194 | 22195 | 22196 | 22197 | 22198 | 22202 | 22204 | 22599 | 27038 | 27039 | 27040 |  |
| ref | T | T | C | G | C | C | C | T | C | A | T | T | T | A | C | T | G | A | T | C |  |
| BA.1 | T | C | C | T | C | T | T | C | T | A | G | T | G | C | A | C | G | A | T | C |  |
| BA.1 | C | C | C | T | C | T | T | C | T | - | - | - | T | A | C | T | G | T | A | A | #1086804 |
| BA.1.1.18 | T | T | T | G | T | C | C | T | C | - | - | - | T | A | C | T | A | T | A | A |  |
| mut | T>C |  |  |  |  |  |  |  |  |  |  |  |  |  |  |  | A>G |  |  |  |  |

|  |  |  |  |  |  |  |  |  |  |  |  |  |  |
| --- | --- | --- | --- | --- | --- | --- | --- | --- | --- | --- | --- | --- | --- |
| pos | 3929 | 3930 | 3931 | 7732 | 22194 | 22195 | 22196 | 22197 | 22198 | 22202 | 22204 | 22882 |  |
| ref | G | T | T | C | A | T | T | T | A | C | T | T |  |
| BA.1.1 | A | A | C | T | A | G | T | G | C | A | C | G |  |
| BA.1.1 | A | A | C | T | - | - | - | T | A | C | T | T | #1089986 |
| BA.1.1 | G | T | T | C | - | - | - | T | A | C | T | G |  |
| mut |  |  |  |  |  |  |  |  |  |  |  | G>T |  |

|  |  |  |  |  |  |  |  |  |  |  |  |  |  |  |  |  |
| --- | --- | --- | --- | --- | --- | --- | --- | --- | --- | --- | --- | --- | --- | --- | --- | --- |
| pos | 934 | 1438 | 1599 | 4975 | 6568 | 14033 | 21595 | 22194 | 22195 | 22196 | 22197 | 22198 | 22202 | 22204 | 25916 |  |
| ref | C | T | G | G | C | A | C | A | T | T | T | A | C | T | C |  |
| BA.1.1 | T | C | G | A | T | C | T | A | G | T | G | C | A | C | T |  |
| BA.1.1 | T | C | G | A | T | C | T | - | - | - | T | A | C | T | T | #1090284 |
| BA.1.1 | C | T | T | G | C | A | C | - | - | - | T | A | C | T | C |  |
| mut |  |  |  |  |  |  |  |  |  |  |  |  |  |  | C>T |  |

|  |  |  |  |  |  |  |  |  |  |  |  |  |  |  |  |  |  |  |  |  |
| --- | --- | --- | --- | --- | --- | --- | --- | --- | --- | --- | --- | --- | --- | --- | --- | --- | --- | --- | --- | --- |
| pos | 313 | 2470 | 6145 | 17010 | 17172 | 17336 | 17749 | 21302 | 21304 | 21305 | 21595 | 22194 | 22195 | 22196 | 22197 | 22198 | 22202 | 22204 | 22599 |  |
| ref | C | C | C | C | C | C | G | C | C | G | C | A | T | T | T | A | C | T | G |  |
| BA.1.1 | C | T | T | T | T | C | A | T | A | A | T | - | - | - | T | A | C | T | A |  |
| BA.1.1 | T | T | T | T | T | T | A | C | C | G | T | A | G | T | G | C | A | C | A | #1090460 |
| BA.1 | C | C | C | C | C | C | G | C | C | G | C | A | G | T | G | C | A | C | G |  |
| mut | C>T |  |  |  |  | C>T |  |  |  |  | C>T |  |  |  |  |  |  |  | G>A |  |

|  |  |  |  |  |  |  |  |  |  |  |  |  |  |  |  |  |  |  |  |  |
| --- | --- | --- | --- | --- | --- | --- | --- | --- | --- | --- | --- | --- | --- | --- | --- | --- | --- | --- | --- | --- |
| pos | 835 | 1346 | 2470 | 7840 | 8302 | 11950 | 12241 | 14805 | 20389 | 21595 | 22194 | 22195 | 22196 | 22197 | 22198 | 22202 | 22204 | 22599 | 25810 |  |
| ref | C | C | C | C | A | C | C | C | C | C | A | T | T | T | A | C | T | G | C |  |
| BA.1.1 | T | A | T | T | G | C | T | C | C | T | - | - | - | T | A | C | T | A | C |  |
| BA.1.1 | T | A | T | T | G | C | T | T | T | T | A | G | T | G | C | A | C | A | T | #1090473 |
| BA.1 | C | C | C | C | A | T | C | T | C | C | A | G | T | G | C | A | C | G | C |  |
| mut |  |  |  |  |  |  |  |  | C>T | C>T |  |  |  |  |  |  |  | G>A | C>T |  |

|  |  |  |  |  |  |  |  |  |  |  |  |  |  |  |  |  |  |  |  |  |  |  |  |  |  |  |  |  |  |
| --- | --- | --- | --- | --- | --- | --- | --- | --- | --- | --- | --- | --- | --- | --- | --- | --- | --- | --- | --- | --- | --- | --- | --- | --- | --- | --- | --- | --- | --- |
| pos | 203 | 897 | 2095 | 4010 | 7926 | 8832 | 13424 | 14940 | 15024 | 16543 | 22194 | 22195 | 22196 | 22197 | 22198 | 22202 | 22204 | 27143 | 29541 | 29543 | 29555 | 29614 | 29700 | 29762 | 29769 | 29774 | 29779 | 29781 |  |
| ref | C | C | G | C | C | C | C | A | C | G | A | T | T | T | A | C | T | C | C | G | C | C | A | C | C | C | G | G |  |
| BA.1.1 | T | T | A | C | C | C | C | G | T | A | - | - | - | T | A | C | T | C | - | - | - | - | - | - | - | - | - | - |  |
| BA.1.1 | T | T | A | T | C | T | C | G | T | G | A | G | T | G | C | A | C | T | C | G | C | C | A | C | C | C | G | G | #1090717 |
| BA.1.1.18 | C | C | G | C | T | C | T | A | C | G | A | G | T | G | C | A | C | T | C | G | C | C | A | C | C | C | G | G |  |
| mut |  |  |  | C>T |  | C>T |  |  |  |  |  |  |  |  |  |  |  |  |  |  |  |  |  |  |  |  |  |  |  |

|  |  |  |  |  |  |  |  |  |  |  |  |  |  |  |
| --- | --- | --- | --- | --- | --- | --- | --- | --- | --- | --- | --- | --- | --- | --- |
| pos | 1190 | 9559 | 15801 | 16471 | 21595 | 22194 | 22195 | 22196 | 22197 | 22198 | 22202 | 22204 | 22792 |  |
| ref | C | C | T | A | C | A | T | T | T | A | C | T | C |  |
| BA.1.1 | T | T | C | G | T | A | G | T | G | C | A | C | C |  |
| BA.1.1 | T | T | C | G | T | - | - | - | T | A | C | T | T | #1093413 |
| BA.1.1 | C | C | T | A | C | - | - | - | T | A | C | T | T |  |
| mut |  |  |  |  |  |  |  |  |  |  |  |  |  |  |

|  |  |  |  |  |  |  |  |  |  |  |  |  |  |  |  |  |  |  |  |  |  |  |  |  |  |  |  |  |  |  |  |  |  |  |  |  |  |  |  |  |  |  |  |  |  |  |  |  |  |  |  |  |  |  |  |  |  |  |  |  |  |  |  |  |  |  |  |  |  |  |  |  |  |  |  |  |  |  |  |  |  |  |  |  |  |  |  |
| --- | --- | --- | --- | --- | --- | --- | --- | --- | --- | --- | --- | --- | --- | --- | --- | --- | --- | --- | --- | --- | --- | --- | --- | --- | --- | --- | --- | --- | --- | --- | --- | --- | --- | --- | --- | --- | --- | --- | --- | --- | --- | --- | --- | --- | --- | --- | --- | --- | --- | --- | --- | --- | --- | --- | --- | --- | --- | --- | --- | --- | --- | --- | --- | --- | --- | --- | --- | --- | --- | --- | --- | --- | --- | --- | --- | --- | --- | --- | --- | --- | --- | --- | --- | --- | --- | --- | --- |
| pos | 44 | 670 | 2790 | 2832 | 4184 | 4321 | 5386 | 6513 | 6514 | 6515 | 8393 | 9344 | 9424 | 9534 | 9693 | 9866 | 10198 | 10447 | 11283 | 11284 | 11285 | 11286 | 11287 | 11292 | 11293 | 11294 | 11295 | 11296 | 11537 | 12880 | 13195 | 15002 | 15240 | 15714 | 17410 | 19955 | 20055 | 21417 | 21618 | 21633 | 21634 | 21635 | 21636 | 21637 | 21638 | 21639 | 21640 | 21641 | 21762 | 21765 | 21766 | 21767 | 21768 | 21769 | 21770 | 21846 | 21987 | 21988 | 21989 | 21990 | 21991 | 21992 | 21993 | 21994 | 21995 | 22194 | 22195 | 22196 | 22200 | 22673 | 22688 | 22775 | 22786 | 22898 | 23048 | 23202 | 24130 | 24503 | 26060 | 26530 | 26858 | 27382 | 27383 | 27384 | 29510 | 29632 |  |
| ref | C | T | C | A | G | C | T | G | T | T | G | C | A | C | C | C | C | G | G | T | T | T | G | G | T | T | T | T | A | C | T | A | C | C | C | C | A | A | C | T | A | C | C | C | C | C | T | G | C | T | A | C | A | T | G | C | G | T | G | T | T | T | A | T | T | A | T | T | T | T | A | G | A | G | G | C | C | C | C | A | C | G | A | T | A | C |  |
| BA.2 | T | G | T | A | A | T | T | G | T | T | G | T | G | T | C | T | T | A | G | T | T | T | G | - | - | - | - | - | A | T | T | G | C | T | T | T | G | A | T | - | - | - | - | - | - | - | - | - | C | T | A | C | A | T | G | C | A | T | G | T | T | T | A | T | T | A | T | T | G | T | G | A | C | G | G | C | C | C | T | A | T | C | T | C | C | C |  |
| BA.1 | T | G | T | A | A | T | T | G | T | T | A | C | A | C | T | C | C | G | - | - | - | - | - | G | T | T | T | T | G | C | C | A | T | C | C | C | A | G | C | T | A | C | C | C | C | C | T | G | T | - | - | - | - | - | - | T | - | - | - | - | - | - | - | - | - | - | - | - | T | C | A | G | A | A | A | A | A | T | C | G | C | G | A | T | A | T | #1095230 |
| BA.1 | C | T | C | G | G | C | G | - | - | - | A | C | A | C | T | C | C | G | - | - | - | - | - | G | T | T | T | T | G | C | C | A | T | C | C | C | A | A | C | T | A | C | C | C | C | C | T | G | T | - | - | - | - | - | - | T | - | - | - | - | - | - | - | - | - | - | - | - | T | C | A | G | A | A | A | A | A | T | C | G | C | G | A | T | A | T |  |
| mut |  |  |  |  |  |  |  |  |  |  |  |  |  |  |  |  |  |  |  |  |  |  |  |  |  |  |  |  |  |  |  |  |  |  |  |  |  | A>G |  |  |  |  |  |  |  |  |  |  |  |  |  |  |  |  |  |  |  |  |  |  |  |  |  |  |  |  |  |  |  |  |  |  |  |  |  |  |  |  |  |  |  |  |  |  |  |  |  |

|  |  |  |  |  |  |  |  |  |  |  |  |  |  |  |  |  |  |  |  |  |
| --- | --- | --- | --- | --- | --- | --- | --- | --- | --- | --- | --- | --- | --- | --- | --- | --- | --- | --- | --- | --- |
| pos | 1342 | 1415 | 2470 | 3802 | 5515 | 7926 | 9615 | 10198 | 16289 | 16596 | 22194 | 22195 | 22196 | 22197 | 22198 | 22202 | 22204 | 22599 | 29520 |  |
| ref | C | C | C | T | G | C | C | C | C | C | A | T | T | T | A | C | T | G | C |  |
| BA.1.1.18 | T | T | T | C | G | T | T | T | T | C | A | G | T | G | C | A | C | A | C |  |
| BA.1.1.18 | T | T | T | C | G | T | T | T | T | C | - | - | - | T | A | C | T | A | T | #1096800 |
| BA.1 | C | C | C | T | T | C | C | C | C | T | - | - | - | T | A | C | T | G | T |  |
| mut |  |  |  |  |  |  |  |  |  |  |  |  |  |  |  |  |  | G>A |  |  |

|  |  |  |  |  |  |  |  |  |  |  |  |  |  |  |  |  |
| --- | --- | --- | --- | --- | --- | --- | --- | --- | --- | --- | --- | --- | --- | --- | --- | --- |
| pos | 1077 | 2470 | 3041 | 8892 | 17304 | 21691 | 22194 | 22195 | 22196 | 22197 | 22198 | 22202 | 22204 | 22599 | 25819 |  |
| ref | C | C | C | C | C | C | A | T | T | T | A | C | T | G | G |  |
| BA.1.1 | T | T | G | T | T | T | - | - | - | T | A | C | T | A | A |  |
| BA.1.1 | T | T | G | T | T | T | A | G | T | G | C | A | C | A | A | #1099666 |
| BA.1 | C | C | C | C | C | C | A | G | T | G | C | A | C | G | G |  |
| mut |  |  |  |  |  |  |  |  |  |  |  |  |  | G>A | G>A |  |

|  |  |  |  |  |  |  |  |  |  |  |  |  |  |  |  |  |  |  |  |  |  |  |  |  |  |  |  |  |  |  |  |  |  |  |  |  |  |  |  |  |  |  |  |  |  |  |  |  |  |  |  |  |  |  |  |  |  |  |  |  |  |  |  |  |  |  |  |  |  |  |  |  |  |  |  |  |  |  |
| --- | --- | --- | --- | --- | --- | --- | --- | --- | --- | --- | --- | --- | --- | --- | --- | --- | --- | --- | --- | --- | --- | --- | --- | --- | --- | --- | --- | --- | --- | --- | --- | --- | --- | --- | --- | --- | --- | --- | --- | --- | --- | --- | --- | --- | --- | --- | --- | --- | --- | --- | --- | --- | --- | --- | --- | --- | --- | --- | --- | --- | --- | --- | --- | --- | --- | --- | --- | --- | --- | --- | --- | --- | --- | --- | --- | --- | --- | --- |
| pos | 670 | 2470 | 2790 | 2832 | 4184 | 4321 | 5386 | 6513 | 6514 | 6515 | 8393 | 9344 | 9424 | 9534 | 9866 | 10198 | 10447 | 11283 | 11284 | 11285 | 11286 | 11287 | 11292 | 11293 | 11294 | 11295 | 11296 | 11537 | 12880 | 13195 | 15240 | 15714 | 17410 | 19955 | 20055 | 21595 | 21618 | 21633 | 21634 | 21635 | 21636 | 21637 | 21638 | 21639 | 21640 | 21641 | 21762 | 21765 | 21766 | 21767 | 21768 | 21769 | 21770 | 21846 | 21987 | 21988 | 21989 | 21990 | 21991 | 21992 | 21993 | 21994 | 21995 | 22194 | 22195 | 22196 | 22200 | 22599 | 22673 | 22688 | 22775 | 22786 | 22898 | 23048 | 23202 | 24130 | 24503 |  |
| ref | T | C | C | A | G | C | T | G | T | T | G | C | A | C | C | C | G | G | T | T | T | G | G | T | T | T | T | A | C | T | C | C | C | C | A | C | C | T | A | C | C | C | C | C | T | G | C | T | A | C | A | T | G | C | G | T | G | T | T | T | A | T | T | A | T | T | T | G | T | A | G | A | G | G | C | C | C |  |
| BA.1.1 | T | T | C | G | G | C | G | - | - | - | A | C | A | C | T | C | G | - | - | - | - | - | G | T | T | T | T | G | C | C | T | C | C | C | A | T | C | T | A | C | C | C | C | C | T | G | T | - | - | - | - | - | - | T | - | - | - | - | - | - | - | - | - | - | - | - | T | A | C | A | G | A | A | A | A | A | T |  |
| BA.2 | T | T | C | G | G | C | G | - | - | - | A | C | A | C | T | T | A | G | T | T | T | G | - | - | - | - | - | A | T | T | C | T | T | T | G | C | T | - | - | - | - | - | - | - | - | - | C | T | A | C | A | T | G | C | A | T | G | T | T | T | A | T | T | A | T | T | G | G | T | G | A | C | G | G | C | C | C | #1099995 |
| BA.2 | G | C | T | A | A | T | T | G | T | T | G | T | G | T | C | T | A | G | T | T | T | G | - | - | - | - | - | A | T | T | C | T | T | T | G | C | T | - | - | - | - | - | - | - | - | - | C | T | A | C | A | T | G | C | A | T | G | T | T | T | A | T | T | A | T | T | G | G | T | G | A | C | G | G | C | C | C |  |
| mut |  |  |  |  |  |  |  |  |  |  |  |  |  |  |  |  |  |  |  |  |  |  |  |  |  |  |  |  |  |  |  |  |  |  |  |  |  |  |  |  |  |  |  |  |  |  |  |  |  |  |  |  |  |  |  |  |  |  |  |  |  |  |  |  |  |  |  |  |  |  |  |  |  |  |  |  |  |  |

|  |  |  |  |  |  |  |  |  |  |  |  |  |  |  |  |  |  |  |
| --- | --- | --- | --- | --- | --- | --- | --- | --- | --- | --- | --- | --- | --- | --- | --- | --- | --- | --- |
| pos | 2306 | 2470 | 3205 | 3784 | 5512 | 21987 | 21988 | 22004 | 22194 | 22195 | 22196 | 22197 | 22198 | 22202 | 22204 | 22599 | 29422 |  |
| ref | C | C | A | C | C | G | T | A | A | T | T | T | A | C | T | G | G |  |
| BA.1.1 | T | T | G | T | T | A | C | C | - | - | - | T | A | C | T | A | A |  |
| BA.1.1 | T | T | G | T | T | A | C | C | A | G | T | G | C | A | C | A | A | #1101289 |
| BA.1 | C | C | A | C | C | - | - | A | A | G | T | G | C | A | C | G | G |  |
| mut |  |  |  |  |  |  |  |  |  |  |  |  |  |  |  | G>A | G>A |  |

|  |  |  |  |  |  |  |  |  |  |  |  |  |  |  |  |  |  |
| --- | --- | --- | --- | --- | --- | --- | --- | --- | --- | --- | --- | --- | --- | --- | --- | --- | --- |
| pos | 1077 | 1851 | 2260 | 2470 | 5822 | 14940 | 18084 | 21102 | 22194 | 22195 | 22196 | 22197 | 22198 | 22202 | 22204 | 22599 |  |
| ref | C | T | C | C | C | A | C | T | A | T | T | T | A | C | T | G |  |
| BA.1.1 | C | C | T | T | T | G | T | C | - | - | - | T | A | C | T | A |  |
| BA.1.1 | T | C | T | T | T | G | T | C | A | G | T | G | C | A | C | A | #1101297 |
| BA.1 | C | T | C | C | C | A | C | T | A | G | T | G | C | A | C | G |  |
| mut | C>T |  |  |  |  |  |  |  |  |  |  |  |  |  |  | G>A |  |

|  |  |  |  |  |  |  |  |  |  |  |  |  |  |  |  |  |  |
| --- | --- | --- | --- | --- | --- | --- | --- | --- | --- | --- | --- | --- | --- | --- | --- | --- | --- |
| pos | 259 | 2470 | 13386 | 14760 | 16744 | 18756 | 18972 | 21595 | 22194 | 22195 | 22196 | 22197 | 22198 | 22202 | 22204 | 28475 |  |
| ref | A | C | G | C | G | G | G | C | A | T | T | T | A | C | T | C |  |
| BA.1.1 | A | C | T | T | T | A | T | C | - | - | - | T | A | C | T | G |  |
| BA.1.1 | A | C | T | T | T | A | T | C | A | G | T | G | C | A | C | G | #1101658 |
| BA.1.1 | G | T | G | C | G | G | G | T | A | G | T | G | C | A | C | C |  |
| mut |  |  |  |  |  |  |  |  |  |  |  |  |  |  |  | C>G |  |

|  |  |  |  |  |  |  |  |  |  |  |  |  |  |  |  |  |  |  |
| --- | --- | --- | --- | --- | --- | --- | --- | --- | --- | --- | --- | --- | --- | --- | --- | --- | --- | --- |
| pos | 259 | 2470 | 3729 | 5880 | 14760 | 16744 | 18756 | 18972 | 21595 | 22194 | 22195 | 22196 | 22197 | 22198 | 22202 | 22204 | 28475 |  |
| ref | A | C | G | G | C | G | G | G | C | A | T | T | T | A | C | T | C |  |
| BA.1.1 | A | C | G | G | T | T | A | T | C | - | - | - | T | A | C | T | G |  |
| BA.1.1 | A | C | A | A | T | T | A | T | C | A | G | T | G | C | A | C | G | #1101660 |
| BA.1.1 | G | T | G | G | C | G | G | G | T | A | G | T | G | C | A | C | C |  |
| mut |  |  | G>A | G>A |  |  |  |  |  |  |  |  |  |  |  |  | C>G |  |

|  |  |  |  |  |  |  |  |  |  |  |  |  |  |  |  |  |  |
| --- | --- | --- | --- | --- | --- | --- | --- | --- | --- | --- | --- | --- | --- | --- | --- | --- | --- |
| pos | 259 | 2470 | 14760 | 16744 | 17245 | 18756 | 18972 | 21595 | 22194 | 22195 | 22196 | 22197 | 22198 | 22202 | 22204 | 28475 |  |
| ref | A | C | C | G | C | G | G | C | A | T | T | T | A | C | T | C |  |
| BA.1.1 | A | C | T | T | T | A | T | C | - | - | - | T | A | C | T | G |  |
| BA.1.1 | A | C | T | T | T | A | T | C | A | G | T | G | C | A | C | G | #1101662 |
| BA.1.1 | G | T | C | G | C | G | G | T | A | G | T | G | C | A | C | C |  |
| mut |  |  |  |  |  |  |  |  |  |  |  |  |  |  |  | C>G |  |

|  |  |  |  |  |  |  |  |  |  |  |  |  |  |  |  |  |  |  |  |
| --- | --- | --- | --- | --- | --- | --- | --- | --- | --- | --- | --- | --- | --- | --- | --- | --- | --- | --- | --- |
| pos | 2470 | 2523 | 5780 | 8302 | 15981 | 18555 | 18695 | 21595 | 22194 | 22195 | 22196 | 22197 | 22198 | 22202 | 22204 | 22599 | 25544 | 29358 |  |
| ref | C | C | A | A | C | C | C | C | A | T | T | T | A | C | T | G | C | C |  |
| BA.1.1 | T | T | G | G | T | T | T | T | - | - | - | T | A | C | T | A | A | A |  |
| BA.1.1 | T | T | G | G | T | T | T | T | A | G | T | G | C | A | C | A | A | A | #1104755 |
| BA.1 | C | C | A | A | C | C | C | C | A | G | T | G | C | A | C | G | C | C |  |
| mut |  |  |  |  |  |  |  |  |  |  |  |  |  |  |  | G>A | C>A | C>A |  |

|  |  |  |  |  |  |  |  |  |  |  |  |  |  |  |  |  |  |
| --- | --- | --- | --- | --- | --- | --- | --- | --- | --- | --- | --- | --- | --- | --- | --- | --- | --- |
| pos | 5079 | 5465 | 7100 | 11442 | 18555 | 18768 | 19488 | 20437 | 22194 | 22195 | 22196 | 22197 | 22198 | 22202 | 22204 | 28254 |  |
| ref | C | T | G | C | C | T | C | A | A | T | T | T | A | C | T | A |  |
| BA.1.1 | T | C | A | T | T | C | T | T | A | G | T | G | C | A | C | A |  |
| BA.1.1 | T | C | A | T | T | C | T | T | - | - | - | T | A | C | T | - | #1104785 |
| BA.1.1 | C | T | G | C | C | T | C | A | - | - | - | T | A | C | T | A |  |
| mut |  |  |  |  |  |  |  |  |  |  |  |  |  |  |  | A>- |  |

|  |  |  |  |  |  |  |  |  |  |  |  |  |  |  |  |
| --- | --- | --- | --- | --- | --- | --- | --- | --- | --- | --- | --- | --- | --- | --- | --- |
| pos | 1188 | 12562 | 16178 | 17491 | 22194 | 22195 | 22196 | 22197 | 22198 | 22202 | 22204 | 23948 | 27384 | 28254 |  |
| ref | C | A | C | C | A | T | T | T | A | C | T | G | T | A |  |
| BA.1.1 | T | G | T | T | A | G | T | G | C | A | C | T | C | A |  |
| BA.1.1 | T | G | T | T | - | - | - | T | A | C | T | G | C | - | #1104942 |
| BA.1.1 | C | A | C | C | - | - | - | T | A | C | T | T | T | A |  |
| mut |  |  |  |  |  |  |  |  |  |  |  | T>G | T>C | A>- |  |

|  |  |  |  |  |  |  |  |  |  |  |  |  |  |  |  |  |  |  |  |  |  |  |  |  |  |  |  |  |  |  |  |  |  |  |  |  |  |  |  |  |  |  |  |  |  |  |  |  |  |  |  |  |  |  |  |  |  |  |  |  |  |  |  |  |  |  |  |  |  |  |  |  |  |  |  |  |  |  |  |  |  |  |  |  |  |  |  |  |  |  |
| --- | --- | --- | --- | --- | --- | --- | --- | --- | --- | --- | --- | --- | --- | --- | --- | --- | --- | --- | --- | --- | --- | --- | --- | --- | --- | --- | --- | --- | --- | --- | --- | --- | --- | --- | --- | --- | --- | --- | --- | --- | --- | --- | --- | --- | --- | --- | --- | --- | --- | --- | --- | --- | --- | --- | --- | --- | --- | --- | --- | --- | --- | --- | --- | --- | --- | --- | --- | --- | --- | --- | --- | --- | --- | --- | --- | --- | --- | --- | --- | --- | --- | --- | --- | --- | --- | --- | --- | --- | --- | --- |
| pos | 44 | 670 | 1436 | 2790 | 2832 | 3261 | 4184 | 4321 | 5386 | 6513 | 6514 | 6515 | 8260 | 8393 | 9344 | 9424 | 9534 | 9866 | 10198 | 10447 | 11283 | 11284 | 11285 | 11286 | 11287 | 11292 | 11293 | 11294 | 11295 | 11296 | 11537 | 12530 | 12880 | 13195 | 15002 | 15240 | 15714 | 17410 | 19955 | 20055 | 21618 | 21633 | 21634 | 21635 | 21636 | 21637 | 21638 | 21639 | 21640 | 21641 | 21762 | 21765 | 21766 | 21767 | 21768 | 21769 | 21770 | 21846 | 21987 | 21988 | 21989 | 21990 | 21991 | 21992 | 21993 | 21994 | 21995 | 22194 | 22195 | 22196 | 22200 | 22673 | 22688 | 22775 | 22786 | 22898 | 23048 | 23202 | 24130 | 24503 | 26060 | 26530 | 26858 | 27382 | 27383 | 27384 | 28087 | 29510 | 29739 |  |
| ref | C | T | T | C | A | C | G | C | T | G | T | T | T | G | C | A | C | C | C | G | G | T | T | T | G | G | T | T | T | T | A | T | C | T | A | C | C | C | C | A | C | T | A | C | C | C | C | C | T | G | C | T | A | C | A | T | G | C | G | T | G | T | T | T | A | T | T | A | T | T | T | T | A | G | A | G | G | C | C | C | C | A | C | G | A | T | C | A | C |  |
| BA.2 | T | G | T | T | A | C | A | T | T | G | T | T | T | G | T | G | T | T | T | A | G | T | T | T | G | - | - | - | - | - | A | T | T | T | G | C | T | T | T | G | T | - | - | - | - | - | - | - | - | - | C | T | A | C | A | T | G | C | A | T | G | T | T | T | A | T | T | A | T | T | G | T | G | A | C | G | G | C | C | C | T | A | T | C | T | C | C | C | C |  |
| BA.1 | T | G | T | T | A | C | A | T | T | - | - | - | C | A | C | A | C | C | C | G | - | - | - | - | - | G | T | T | T | T | G | C | C | C | A | T | C | C | C | A | C | T | A | C | C | C | C | C | T | G | T | - | - | - | - | - | - | T | - | - | - | - | - | - | - | - | - | - | - | - | T | C | A | G | A | A | A | A | A | T | C | G | C | G | A | T | T | A | T | #1106336 |
| BA.1 | C | T | G | C | G | T | G | C | G | - | - | - | C | A | C | A | C | C | C | G | - | - | - | - | - | G | T | T | T | T | G | C | C | C | A | T | C | C | C | A | C | T | A | C | C | C | C | C | T | G | T | - | - | - | - | - | - | T | - | - | - | - | - | - | - | - | - | - | - | - | T | C | A | G | A | A | A | A | A | T | C | G | C | G | A | T | T | A | T |  |
| mut |  |  |  |  |  |  |  |  |  |  |  |  |  |  |  |  |  |  |  |  |  |  |  |  |  |  |  |  |  |  |  |  |  |  |  |  |  |  |  |  |  |  |  |  |  |  |  |  |  |  |  |  |  |  |  |  |  |  |  |  |  |  |  |  |  |  |  |  |  |  |  |  |  |  |  |  |  |  |  |  |  |  |  |  |  |  |  |  |  |  |

|  |  |  |  |  |  |  |  |  |  |  |  |  |  |  |  |  |
| --- | --- | --- | --- | --- | --- | --- | --- | --- | --- | --- | --- | --- | --- | --- | --- | --- |
| pos | 2488 | 4455 | 6543 | 9693 | 14724 | 16393 | 21595 | 22194 | 22195 | 22196 | 22197 | 22198 | 22202 | 22204 | 26022 |  |
| ref | C | C | C | C | C | C | C | A | T | T | T | A | C | T | C |  |
| BA.1.1 | T | T | T | C | T | T | T | - | - | - | T | A | C | T | C |  |
| BA.1.1 | T | T | T | C | T | T | T | A | G | T | G | C | A | C | T | #1107147 |
| BA.1.1 | C | C | C | T | C | C | C | A | G | T | G | C | A | C | T |  |
| mut |  |  |  |  |  |  |  |  |  |  |  |  |  |  |  |  |

|  |  |  |  |  |  |  |  |  |  |  |  |  |  |  |
| --- | --- | --- | --- | --- | --- | --- | --- | --- | --- | --- | --- | --- | --- | --- |
| pos | 221 | 22194 | 22195 | 22196 | 22197 | 22198 | 22202 | 22204 | 26522 | 27365 | 29539 | 29625 | 29730 |  |
| ref | T | A | T | T | T | A | C | T | C | A | T | C | C |  |
| BA.1.1 | C | - | - | - | T | A | C | T | T | A | T | C | C |  |
| BA.1.1 | C | - | - | - | T | A | C | T | C | G | A | T | T | #1107217 |
| BA.1.1 | T | A | G | T | G | C | A | C | C | G | A | T | T |  |
| mut |  |  |  |  |  |  |  |  |  |  |  |  |  |  |

|  |  |  |  |  |  |  |  |  |  |  |  |  |  |  |  |  |  |  |  |  |  |  |
| --- | --- | --- | --- | --- | --- | --- | --- | --- | --- | --- | --- | --- | --- | --- | --- | --- | --- | --- | --- | --- | --- | --- |
| pos | 1572 | 1593 | 2695 | 3045 | 4206 | 8233 | 9852 | 14107 | 21595 | 22186 | 22194 | 22195 | 22196 | 22197 | 22198 | 22202 | 22204 | 25207 | 27384 | 27741 | 28998 |  |
| ref | C | C | C | C | C | A | A | A | C | G | A | T | T | T | A | C | T | C | T | C | A |  |
| BA.1.1.14 | T | C | T | C | T | G | C | G | C | A | - | - | - | T | A | C | T | C | C | T | A |  |
| BA.1.1.14 | T | C | T | C | T | G | C | G | C | A | A | G | T | G | C | A | C | T | C | T | G | #1107255 |
| BA.1.1 | C | T | C | T | C | A | A | A | T | G | A | G | T | G | C | A | C | T | T | C | A |  |
| mut |  |  |  |  |  |  |  |  |  |  |  |  |  |  |  |  |  |  | T>C | C>T | A>G |  |

|  |  |  |  |  |  |  |  |  |  |  |  |  |  |  |
| --- | --- | --- | --- | --- | --- | --- | --- | --- | --- | --- | --- | --- | --- | --- |
| pos | 1330 | 2470 | 7299 | 15194 | 16260 | 22194 | 22195 | 22196 | 22197 | 22198 | 22202 | 22204 | 22792 |  |
| ref | T | C | C | C | C | A | T | T | T | A | C | T | C |  |
| BA.1.1 | C | T | T | T | T | - | - | - | T | A | C | T | T |  |
| BA.1.1 | C | T | T | T | T | A | G | T | G | C | A | C | T | #1107261 |
| BA.1.1 | T | C | C | C | C | A | G | T | G | C | A | C | C |  |
| mut |  |  |  |  |  |  |  |  |  |  |  |  | C>T |  |

|  |  |  |  |  |  |  |  |  |  |  |  |  |  |  |  |  |
| --- | --- | --- | --- | --- | --- | --- | --- | --- | --- | --- | --- | --- | --- | --- | --- | --- |
| pos | 689 | 4545 | 9191 | 16476 | 20003 | 22194 | 22195 | 22196 | 22197 | 22198 | 22202 | 22204 | 27713 | 29041 | 29543 |  |
| ref | T | G | G | T | A | A | T | T | T | A | C | T | T | G | G |  |
| BA.1.1 | C | A | G | C | A | - | - | - | T | A | C | T | T | T | G |  |
| BA.1.1 | C | A | G | C | A | A | G | T | G | C | A | C | C | T | G | #1109239 |
| BA.1.1 | T | G | T | T | G | A | G | T | G | C | A | C | T | G | T |  |
| mut |  |  |  |  |  |  |  |  |  |  |  |  | T>C | G>T | T>G |  |

|  |  |  |  |  |  |  |  |  |  |  |  |  |  |  |  |  |  |
| --- | --- | --- | --- | --- | --- | --- | --- | --- | --- | --- | --- | --- | --- | --- | --- | --- | --- |
| pos | 2106 | 2470 | 5629 | 9913 | 14877 | 21249 | 22046 | 22194 | 22195 | 22196 | 22197 | 22198 | 22202 | 22204 | 22599 | 26007 |  |
| ref | C | C | G | T | C | G | A | A | T | T | T | A | C | T | G | T |  |
| BA.1.1 | T | T | T | C | T | T | C | - | - | - | T | A | C | T | A | C |  |
| BA.1.1 | T | T | T | C | T | T | C | A | G | T | G | C | A | C | A | C | #1109323 |
| BA.1 | C | C | G | T | C | G | A | A | G | T | G | C | A | C | G | T |  |
| mut |  |  |  |  |  |  |  |  |  |  |  |  |  |  | G>A | T>C |  |

|  |  |  |  |  |  |  |  |  |  |  |  |  |  |  |  |  |  |
| --- | --- | --- | --- | --- | --- | --- | --- | --- | --- | --- | --- | --- | --- | --- | --- | --- | --- |
| pos | 1371 | 4965 | 10261 | 13176 | 15240 | 17639 | 18294 | 20762 | 22194 | 22195 | 22196 | 22197 | 22198 | 22202 | 22204 | 27059 |  |
| ref | A | C | G | C | C | C | A | C | A | T | T | T | A | C | T | C |  |
| BA.1.15 | A | C | A | T | C | T | G | C | - | - | - | T | A | C | T | T |  |
| BA.1.15 | A | C | A | T | C | T | G | C | A | G | T | G | C | A | C | T | #1109353 |
| BA.1.15 | G | T | G | C | T | C | A | T | A | G | T | G | C | A | C | C |  |
| mut |  |  |  |  |  |  |  |  |  |  |  |  |  |  |  | C>T |  |

|  |  |  |  |  |  |  |  |  |  |  |  |  |  |  |  |  |
| --- | --- | --- | --- | --- | --- | --- | --- | --- | --- | --- | --- | --- | --- | --- | --- | --- |
| pos | 5730 | 6347 | 6474 | 12733 | 15024 | 20432 | 22078 | 22194 | 22195 | 22196 | 22197 | 22198 | 22202 | 22204 | 29762 |  |
| ref | C | C | A | C | C | T | T | A | T | T | T | A | C | T | C |  |
| BA.1.18 | T | T | G | C | C | C | G | - | - | - | T | A | C | T | - |  |
| BA.1.18 | T | T | G | T | T | C | G | A | G | T | G | C | A | C | - | #1109378 |
| BA.1 | C | C | A | C | C | T | T | A | G | T | G | C | A | C | C |  |
| mut |  |  |  | C>T | C>T |  |  |  |  |  |  |  |  |  | C>- |  |

|  |  |  |  |  |  |  |  |  |  |  |  |  |  |  |
| --- | --- | --- | --- | --- | --- | --- | --- | --- | --- | --- | --- | --- | --- | --- |
| pos | 1973 | 2470 | 3241 | 3653 | 3873 | 5924 | 13682 | 14206 | 18162 | 22599 | 23664 | 26082 | 27626 |  |
| ref | C | C | C | C | T | G | A | C | C | G | C | C | G |  |
| BA.1.1 | T | T | C | C | G | G | A | T | C | A | C | T | G |  |
| BA.1.17.2 | T | T | C | C | G | A | G | C | T | G | T | C | A | #1110637 |
| BA.1.17.2 | C | C | T | T | T | A | G | C | T | G | T | C | G |  |
| mut |  |  |  |  |  |  |  |  |  |  |  |  | G>A |  |

|  |  |  |  |  |  |  |  |  |  |  |  |  |  |  |  |  |  |  |  |  |  |
| --- | --- | --- | --- | --- | --- | --- | --- | --- | --- | --- | --- | --- | --- | --- | --- | --- | --- | --- | --- | --- | --- |
| pos | 670 | 2790 | 2832 | 3241 | 4184 | 4321 | 5055 | 5386 | 5924 | 6513 | 6514 | 6515 | 7969 | 8393 | 9344 | 9424 | 9534 | 9866 | 10198 | 10447 |  |
| ref | T | C | A | C | G | C | C | T | G | G | T | T | T | G | C | A | C | C | C | G |  |
| BA.2 | G | T | A | C | A | T | T | T | G | G | T | T | T | G | T | G | T | T | T | A |  |
| BA.2 | G | T | A | C | A | T | T | G | G | - | - | - | A | A | C | A | C | C | C | A | #1113544 |
| BA.2 | T | C | G | T | G | C | C | G | A | - | - | - | T | A | C | A | C | C | C | G |  |
| mut |  |  |  |  |  |  |  | T>G |  |  |  |  | T>A |  |  |  |  |  |  | G>A |  |

|  |  |  |  |  |  |  |  |  |  |  |  |  |  |  |  |  |  |  |  |  |  |  |  |  |  |  |
| --- | --- | --- | --- | --- | --- | --- | --- | --- | --- | --- | --- | --- | --- | --- | --- | --- | --- | --- | --- | --- | --- | --- | --- | --- | --- | --- |
| pos | 65 | 2470 | 4579 | 11986 | 14386 | 14940 | 20941 | 22194 | 22195 | 22196 | 22197 | 22198 | 22202 | 22204 | 22599 | 29541 | 29543 | 29555 | 29614 | 29700 | 29762 | 29769 | 29774 | 29779 | 29781 |  |
| ref | T | C | T | C | T | A | C | A | T | T | T | A | C | T | G | C | G | C | C | A | C | C | C | G | G |  |
| BA.1.1 | T | T | T | T | C | G | T | A | G | T | G | C | A | C | A | C | G | C | C | A | C | C | C | G | G |  |
| BA.1.1 | T | T | T | T | C | G | T | - | - | - | T | A | C | T | A | - | - | - | - | - | - | - | - | - | - | #1114933 |
| BA.1 | G | C | A | C | T | A | C | - | - | - | T | A | C | T | G | C | G | C | C | A | C | C | C | G | G |  |
| mut |  |  |  |  |  |  |  |  |  |  |  |  |  |  | G>A | C>- | G>- | C>- | C>- | A>- | C>- | C>- | C>- | G>- | G>- |  |

|  |  |  |  |  |  |  |  |  |  |  |  |  |  |  |  |
| --- | --- | --- | --- | --- | --- | --- | --- | --- | --- | --- | --- | --- | --- | --- | --- |
| pos | 1085 | 2470 | 3170 | 4993 | 14206 | 21595 | 22194 | 22195 | 22196 | 22197 | 22198 | 22202 | 22204 | 29300 |  |
| ref | G | C | C | C | C | C | A | T | T | T | A | C | T | G |  |
| BA.1.1 | T | C | T | T | T | C | - | - | - | T | A | C | T | C |  |
| BA.1.1 | T | C | T | T | T | C | A | G | T | G | C | A | C | C | #1118164 |
| BA.1.1 | G | T | C | C | C | T | A | G | T | G | C | A | C | G |  |
| mut |  |  |  |  |  |  |  |  |  |  |  |  |  | G>C |  |

|  |  |  |  |  |  |  |  |  |  |  |  |  |  |  |  |  |  |  |  |  |  |  |  |  |  |  |  |  |  |  |  |  |  |  |  |  |  |  |  |  |  |  |  |  |  |  |  |  |  |  |  |  |  |  |  |  |  |  |  |  |  |  |  |  |  |  |  |  |  |  |  |  |  |  |  |  |  |  |  |  |  |  |  |  |  |  |  |  |  |  |  |  |
| --- | --- | --- | --- | --- | --- | --- | --- | --- | --- | --- | --- | --- | --- | --- | --- | --- | --- | --- | --- | --- | --- | --- | --- | --- | --- | --- | --- | --- | --- | --- | --- | --- | --- | --- | --- | --- | --- | --- | --- | --- | --- | --- | --- | --- | --- | --- | --- | --- | --- | --- | --- | --- | --- | --- | --- | --- | --- | --- | --- | --- | --- | --- | --- | --- | --- | --- | --- | --- | --- | --- | --- | --- | --- | --- | --- | --- | --- | --- | --- | --- | --- | --- | --- | --- | --- | --- | --- | --- | --- | --- | --- | --- |
| pos | 670 | 962 | 2790 | 2832 | 4184 | 4321 | 5386 | 6513 | 6514 | 6515 | 8393 | 9344 | 9424 | 9534 | 9866 | 10135 | 10198 | 10447 | 11283 | 11284 | 11285 | 11286 | 11287 | 11292 | 11293 | 11294 | 11295 | 11296 | 11537 | 12565 | 12880 | 13195 | 15240 | 15714 | 16443 | 17410 | 18705 | 19955 | 20055 | 21618 | 21633 | 21634 | 21635 | 21636 | 21637 | 21638 | 21639 | 21640 | 21641 | 21762 | 21765 | 21766 | 21767 | 21768 | 21769 | 21770 | 21846 | 21987 | 21988 | 21989 | 21990 | 21991 | 21992 | 21993 | 21994 | 21995 | 22194 | 22195 | 22196 | 22200 | 22673 | 22688 | 22775 | 22786 | 22822 | 22898 | 23048 | 23202 | 24130 | 24325 | 24503 | 25708 | 26060 | 26530 | 26858 | 27382 | 27383 | 27384 | 27561 | 29301 | 29510 |  |
| ref | T | G | C | A | G | C | T | G | T | T | G | C | A | C | C | T | C | G | G | T | T | T | G | G | T | T | T | T | A | G | C | T | C | C | C | C | C | C | A | C | T | A | C | C | C | C | C | T | G | C | T | A | C | A | T | G | C | G | T | G | T | T | T | A | T | T | A | T | T | T | T | A | G | A | T | G | G | C | C | A | C | C | C | A | C | G | A | T | G | A | A |  |
| BA.2 | G | G | T | A | A | T | T | G | T | T | G | T | G | T | T | T | T | A | G | T | T | T | G | - | - | - | - | - | A | G | T | T | C | T | T | T | C | T | G | T | - | - | - | - | - | - | - | - | - | C | T | A | C | A | T | G | C | A | T | G | T | T | T | A | T | T | A | T | T | G | T | G | A | C | T | G | G | C | C | G | C | C | T | A | T | C | T | C | G | A | C |  |
| BA.2 | G | G | T | A | A | T | T | G | T | T | G | T | G | T | T | T | T | A | G | T | T | T | G | - | - | - | - | - | A | G | T | T | C | T | T | T | C | T | G | T | - | - | - | - | - | - | - | - | - | C | T | A | C | A | T | G | C | A | T | G | T | T | T | A | T | T | A | T | T | G | T | G | A | C | T | G | G | C | C | G | C | C | T | A | T | G | A | T | T | G | A | #1118591 |
| BA.1.15 | T | A | C | G | G | C | G | - | - | - | A | C | A | C | C | C | C | G | - | - | - | - | - | G | T | T | T | T | G | T | C | C | T | C | C | C | T | C | A | C | T | A | C | C | C | C | C | T | G | T | - | - | - | - | - | - | T | - | - | - | - | - | - | - | - | - | - | - | - | T | C | A | G | A | C | A | A | A | A | A | T | T | C | G | C | G | A | T | T | G | A |  |
| mut |  |  |  |  |  |  |  |  |  |  |  |  |  |  |  |  |  |  |  |  |  |  |  |  |  |  |  |  |  |  |  |  |  |  |  |  |  |  |  |  |  |  |  |  |  |  |  |  |  |  |  |  |  |  |  |  |  |  |  |  |  |  |  |  |  |  |  |  |  |  |  |  |  |  |  |  |  |  |  |  |  |  |  |  |  |  |  |  |  |  |  |  |

|  |  |  |  |  |  |  |  |  |  |  |  |  |  |  |  |  |  |  |  |  |  |  |  |  |  |  |  |  |  |  |  |  |  |  |  |  |  |  |  |  |  |  |  |  |  |  |  |  |  |  |  |  |  |  |  |  |  |  |  |  |  |  |  |  |  |  |  |  |  |  |  |  |  |  |  |  |  |  |  |  |  |  |  |  |  |  |  |  |  |  |  |
| --- | --- | --- | --- | --- | --- | --- | --- | --- | --- | --- | --- | --- | --- | --- | --- | --- | --- | --- | --- | --- | --- | --- | --- | --- | --- | --- | --- | --- | --- | --- | --- | --- | --- | --- | --- | --- | --- | --- | --- | --- | --- | --- | --- | --- | --- | --- | --- | --- | --- | --- | --- | --- | --- | --- | --- | --- | --- | --- | --- | --- | --- | --- | --- | --- | --- | --- | --- | --- | --- | --- | --- | --- | --- | --- | --- | --- | --- | --- | --- | --- | --- | --- | --- | --- | --- | --- | --- | --- | --- | --- | --- |
| pos | 670 | 2470 | 2790 | 2832 | 4184 | 4321 | 5386 | 6513 | 6514 | 6515 | 8393 | 9130 | 9344 | 9424 | 9534 | 9866 | 10198 | 10447 | 11283 | 11284 | 11285 | 11286 | 11287 | 11292 | 11293 | 11294 | 11295 | 11296 | 11537 | 12253 | 12880 | 13195 | 13860 | 14946 | 15240 | 15714 | 16317 | 17410 | 19955 | 20055 | 21595 | 21618 | 21633 | 21634 | 21635 | 21636 | 21637 | 21638 | 21639 | 21640 | 21641 | 21762 | 21765 | 21766 | 21767 | 21768 | 21769 | 21770 | 21846 | 21987 | 21988 | 21989 | 21990 | 21991 | 21992 | 21993 | 21994 | 21995 | 22194 | 22195 | 22196 | 22200 | 22599 | 22673 | 22688 | 22775 | 22786 | 22898 | 23048 | 23202 | 24130 | 24503 | 25606 | 26060 | 26530 | 26858 | 27382 | 27383 | 27384 | 29510 |  |
| ref | T | C | C | A | G | C | T | G | T | T | G | G | C | A | C | C | C | G | G | T | T | T | G | G | T | T | T | T | A | C | C | T | C | T | C | C | T | C | C | A | C | C | T | A | C | C | C | C | C | T | G | C | T | A | C | A | T | G | C | G | T | G | T | T | T | A | T | T | A | T | T | T | G | T | A | G | A | G | G | C | C | C | G | C | A | C | G | A | T | A |  |
| BA.1.1 | T | T | C | G | G | C | G | - | - | - | A | G | C | A | C | C | C | G | - | - | - | - | - | G | T | T | T | T | G | T | C | C | C | T | T | C | C | C | C | A | T | C | T | A | C | C | C | C | C | T | G | T | - | - | - | - | - | - | T | - | - | - | - | - | - | - | - | - | - | - | - | T | A | C | A | G | A | A | A | A | A | T | G | C | G | C | G | A | T | A |  |
| BA.2 | T | T | C | G | G | C | G | - | - | - | A | G | C | A | C | C | C | G | - | - | - | - | - | G | T | T | T | T | G | T | C | T | T | A | C | T | T | T | T | G | C | T | - | - | - | - | - | - | - | - | - | C | T | A | C | A | T | G | C | A | T | G | T | T | T | A | T | T | A | T | T | G | G | T | G | A | C | G | G | C | C | C | A | T | A | T | C | T | C | C | #1123300 |
| BA.2 | G | C | T | A | A | T | T | G | T | T | G | T | T | G | T | T | T | A | G | T | T | T | G | - | - | - | - | - | A | C | T | T | T | A | C | T | T | T | T | G | C | T | - | - | - | - | - | - | - | - | - | C | T | A | C | A | T | G | C | A | T | G | T | T | T | A | T | T | A | T | T | G | G | T | G | A | C | G | G | C | C | C | A | T | A | T | C | T | C | C |  |
| mut |  |  |  |  |  |  |  |  |  |  |  |  |  |  |  |  |  |  |  |  |  |  |  |  |  |  |  |  |  |  |  |  |  |  |  |  |  |  |  |  |  |  |  |  |  |  |  |  |  |  |  |  |  |  |  |  |  |  |  |  |  |  |  |  |  |  |  |  |  |  |  |  |  |  |  |  |  |  |  |  |  |  |  |  |  |  |  |  |  |  |  |

|  |  |  |  |  |  |  |  |  |  |  |  |  |  |
| --- | --- | --- | --- | --- | --- | --- | --- | --- | --- | --- | --- | --- | --- |
| pos | 44 | 6548 | 9300 | 16494 | 22600 | 22792 | 25416 | 25624 | 29729 | 29757 | 29758 | 29759 |  |
| ref | C | C | G | T | A | C | C | C | T | G | T | G |  |
| BA.2.9 | C | C | G | C | A | T | C | T | T | G | T | G |  |
| BA.2 | C | C | G | C | A | C | C | C | C | A | C | C | #1125189 |
| BA.2 | T | T | A | T | T | C | T | C | C | A | C | C |  |
| mut |  |  |  |  |  | T>C |  | T>C |  |  |  |  |  |

|  |  |  |  |  |  |  |  |  |  |  |  |  |  |  |  |  |
| --- | --- | --- | --- | --- | --- | --- | --- | --- | --- | --- | --- | --- | --- | --- | --- | --- |
| pos | 2470 | 4300 | 6616 | 8576 | 12781 | 16965 | 19983 | 22194 | 22195 | 22196 | 22197 | 22198 | 22202 | 22204 | 22599 |  |
| ref | C | G | A | A | C | A | C | A | T | T | T | A | C | T | G |  |
| BA.1.1 | T | T | G | C | T | G | T | - | - | - | T | A | C | T | A |  |
| BA.1.1 | T | T | G | C | T | G | T | A | G | T | G | C | A | C | A | #1126052 |
| BA.1 | C | G | A | A | C | A | C | A | G | T | G | C | A | C | G |  |
| mut |  |  |  |  |  |  |  |  |  |  |  |  |  |  | G>A |  |

|  |  |  |  |  |  |  |  |  |  |  |  |  |  |  |  |  |
| --- | --- | --- | --- | --- | --- | --- | --- | --- | --- | --- | --- | --- | --- | --- | --- | --- |
| pos | 1081 | 8327 | 13860 | 21034 | 21724 | 22194 | 22195 | 22196 | 22197 | 22198 | 22202 | 22204 | 24118 | 27579 | 27583 |  |
| ref | T | C | C | C | G | A | T | T | T | A | C | T | A | A | G |  |
| BA.1.1 | C | C | C | T | A | - | - | - | T | A | C | T | T | A | G |  |
| BA.1.1 | C | C | T | T | A | A | G | T | G | C | A | C | T | T | T | #1126362 |
| BA.1.1 | T | T | C | C | G | A | G | T | G | C | A | C | A | T | T |  |
| mut |  |  | C>T |  |  |  |  |  |  |  |  |  | A>T |  |  |  |

|  |  |  |  |  |  |  |  |  |  |  |  |  |  |  |  |  |  |  |  |  |  |  |  |  |  |  |  |  |
| --- | --- | --- | --- | --- | --- | --- | --- | --- | --- | --- | --- | --- | --- | --- | --- | --- | --- | --- | --- | --- | --- | --- | --- | --- | --- | --- | --- | --- |
| pos | 686 | 687 | 688 | 689 | 690 | 691 | 692 | 693 | 694 | 809 | 2470 | 5515 | 15380 | 16596 | 19572 | 19866 | 20104 | 21595 | 22194 | 22195 | 22196 | 22197 | 22198 | 22202 | 22204 | 22599 | 29072 |  |
| ref | A | A | G | T | C | A | T | T | T | T | C | G | G | C | C | T | C | C | A | T | T | T | A | C | T | G | G |  |
| BA.1 | A | A | G | T | C | A | T | T | T | C | C | T | G | T | T | C | T | C | A | G | T | G | C | A | C | G | G |  |
| BA.1 | - | - | - | - | - | - | - | - | - | C | C | T | T | T | T | C | T | C | - | - | - | T | A | C | T | G | T | #1126552 |
| BA.1.1 | A | A | G | T | C | A | T | T | T | T | T | G | G | C | C | T | C | T | - | - | - | T | A | C | T | A | T |  |
| mut | A>- | A>- | G>- | T>- | C>- | A>- | T>- | T>- | T>- |  |  |  | G>T |  |  |  |  |  |  |  |  |  |  |  |  | A>G |  |  |

|  |  |  |  |  |  |  |  |  |  |  |  |  |  |
| --- | --- | --- | --- | --- | --- | --- | --- | --- | --- | --- | --- | --- | --- |
| pos | 8290 | 12116 | 14940 | 21595 | 22045 | 22194 | 22195 | 22196 | 22197 | 22198 | 22202 | 22204 |  |
| ref | C | C | A | C | T | A | T | T | T | A | C | T |  |
| BA.1.1 | T | A | G | C | A | - | - | - | T | A | C | T |  |
| BA.1.1 | T | A | G | C | A | A | G | T | G | C | A | C | #1126575 |
| BA.1.1 | C | C | A | T | T | A | G | T | G | C | A | C |  |
| mut |  |  |  |  |  |  |  |  |  |  |  |  |  |

|  |  |  |  |  |  |  |  |  |  |  |  |  |  |  |  |  |  |  |  |  |  |  |  |  |  |  |  |  |  |  |  |  |  |  |  |  |  |  |  |  |  |  |  |  |  |  |  |  |  |  |  |  |  |  |  |  |  |  |  |  |  |  |  |  |  |  |  |  |  |  |  |  |  |  |  |  |  |  |  |  |  |  |  |  |  |  |  |  |  |  |  |
| --- | --- | --- | --- | --- | --- | --- | --- | --- | --- | --- | --- | --- | --- | --- | --- | --- | --- | --- | --- | --- | --- | --- | --- | --- | --- | --- | --- | --- | --- | --- | --- | --- | --- | --- | --- | --- | --- | --- | --- | --- | --- | --- | --- | --- | --- | --- | --- | --- | --- | --- | --- | --- | --- | --- | --- | --- | --- | --- | --- | --- | --- | --- | --- | --- | --- | --- | --- | --- | --- | --- | --- | --- | --- | --- | --- | --- | --- | --- | --- | --- | --- | --- | --- | --- | --- | --- | --- | --- | --- | --- | --- |
| pos | 670 | 2790 | 2832 | 4184 | 4321 | 5365 | 5386 | 5924 | 6513 | 6514 | 6515 | 8393 | 9209 | 9344 | 9424 | 9534 | 9866 | 10198 | 10447 | 11283 | 11284 | 11285 | 11286 | 11287 | 11292 | 11293 | 11294 | 11295 | 11296 | 11537 | 12880 | 13019 | 13195 | 15240 | 15714 | 17410 | 19955 | 20055 | 21618 | 21633 | 21634 | 21635 | 21636 | 21637 | 21638 | 21639 | 21640 | 21641 | 21762 | 21765 | 21766 | 21767 | 21768 | 21769 | 21770 | 21846 | 21987 | 21988 | 21989 | 21990 | 21991 | 21992 | 21993 | 21994 | 21995 | 22194 | 22195 | 22196 | 22200 | 22673 | 22688 | 22775 | 22786 | 22792 | 22898 | 23048 | 23202 | 24130 | 24503 | 25557 | 25624 | 26060 | 26530 | 26858 | 27382 | 27383 | 27384 | 27630 | 28903 | 29510 |  |
| ref | T | C | A | G | C | C | T | G | G | T | T | G | G | C | A | C | C | C | G | G | T | T | T | G | G | T | T | T | T | A | C | C | T | C | C | C | C | A | C | T | A | C | C | C | C | C | T | G | C | T | A | C | A | T | G | C | G | T | G | T | T | T | A | T | T | A | T | T | T | T | A | G | A | C | G | G | C | C | C | T | C | C | A | C | G | A | T | C | G | A |  |
| BA.1.17 | T | C | G | G | C | T | G | A | - | - | - | A | G | C | A | C | C | C | G | - | - | - | - | - | G | T | T | T | T | G | C | T | C | T | C | C | C | A | C | T | A | C | C | C | C | C | T | G | T | - | - | - | - | - | - | T | - | - | - | - | - | - | - | - | - | - | - | - | T | C | A | G | A | C | A | A | A | A | T | C | C | C | G | C | G | A | T | C | A | A |  |
| BA.2 | T | C | G | G | C | T | G | A | - | - | - | A | G | T | G | T | T | T | A | G | T | T | T | G | - | - | - | - | - | A | T | C | T | C | T | T | T | G | T | - | - | - | - | - | - | - | - | - | C | T | A | C | A | T | G | C | A | T | G | T | T | T | A | T | T | A | T | T | G | T | G | A | C | T | G | G | C | C | C | T | T | T | A | T | C | T | C | A | G | C | #1128440 |
| BA.2 | G | T | A | A | T | C | T | G | G | T | T | G | A | T | G | T | T | T | A | G | T | T | T | G | - | - | - | - | - | A | T | C | T | C | T | T | T | G | T | - | - | - | - | - | - | - | - | - | C | T | A | C | A | T | G | C | A | T | G | T | T | T | A | T | T | A | T | T | G | T | G | A | C | C | G | G | C | C | C | T | C | T | A | T | C | T | C | A | G | C |  |
| mut |  |  |  |  |  |  |  |  |  |  |  |  |  |  |  |  |  |  |  |  |  |  |  |  |  |  |  |  |  |  |  |  |  |  |  |  |  |  |  |  |  |  |  |  |  |  |  |  |  |  |  |  |  |  |  |  |  |  |  |  |  |  |  |  |  |  |  |  |  |  |  |  |  | C>T |  |  |  |  |  |  | C>T |  |  |  |  |  |  |  |  |  |  |

|  |  |  |  |  |  |  |  |  |  |  |  |  |  |  |  |  |  |
| --- | --- | --- | --- | --- | --- | --- | --- | --- | --- | --- | --- | --- | --- | --- | --- | --- | --- |
| pos | 2470 | 2865 | 3251 | 4917 | 11653 | 12644 | 13966 | 14940 | 22194 | 22195 | 22196 | 22197 | 22198 | 22202 | 22204 | 22599 |  |
| ref | C | A | G | T | C | A | G | A | A | T | T | T | A | C | T | G |  |
| BA.1.1 | T | C | A | C | T | G | A | G | - | - | - | T | A | C | T | A |  |
| BA.1.1 | T | C | A | C | T | G | A | G | A | G | T | G | C | A | C | A | #1129394 |
| BA.1 | C | A | G | T | C | A | G | A | A | G | T | G | C | A | C | G |  |
| mut |  |  |  |  |  |  |  |  |  |  |  |  |  |  |  | G>A |  |

|  |  |  |  |  |  |  |  |  |  |  |  |  |  |  |  |  |  |  |  |  |  |  |  |  |  |  |  |  |  |  |  |  |  |  |  |  |  |  |  |  |  |  |  |  |  |  |  |  |  |  |  |  |  |  |  |  |  |  |  |  |  |  |  |  |  |  |  |  |  |  |  |  |  |  |  |  |  |  |  |  |  |  |  |  |  |  |  |  |  |  |  |  |  |  |  |  |
| --- | --- | --- | --- | --- | --- | --- | --- | --- | --- | --- | --- | --- | --- | --- | --- | --- | --- | --- | --- | --- | --- | --- | --- | --- | --- | --- | --- | --- | --- | --- | --- | --- | --- | --- | --- | --- | --- | --- | --- | --- | --- | --- | --- | --- | --- | --- | --- | --- | --- | --- | --- | --- | --- | --- | --- | --- | --- | --- | --- | --- | --- | --- | --- | --- | --- | --- | --- | --- | --- | --- | --- | --- | --- | --- | --- | --- | --- | --- | --- | --- | --- | --- | --- | --- | --- | --- | --- | --- | --- | --- | --- | --- | --- | --- | --- | --- |
| pos | 44 | 670 | 2790 | 2832 | 2944 | 3104 | 4006 | 4184 | 4321 | 4527 | 5386 | 6513 | 6514 | 6515 | 8290 | 8393 | 9424 | 9534 | 9866 | 10198 | 10447 | 11283 | 11284 | 11285 | 11286 | 11287 | 11292 | 11293 | 11294 | 11295 | 11296 | 11537 | 12164 | 12880 | 13195 | 15240 | 15672 | 15714 | 16209 | 17410 | 19955 | 20055 | 20235 | 21618 | 21633 | 21634 | 21635 | 21636 | 21637 | 21638 | 21639 | 21640 | 21641 | 21762 | 21765 | 21766 | 21767 | 21768 | 21769 | 21770 | 21846 | 21987 | 21988 | 21989 | 21990 | 21991 | 21992 | 21993 | 21994 | 21995 | 22194 | 22195 | 22196 | 22200 | 22673 | 22688 | 22775 | 22786 | 22898 | 23048 | 23202 | 23673 | 24130 | 24503 | 25416 | 26060 | 26530 | 26858 | 27382 | 27383 | 27384 | 28399 | 29476 | 29510 | 29632 |  |
| ref | C | T | C | A | G | T | G | G | C | G | T | G | T | T | C | G | A | C | C | C | G | G | T | T | T | G | G | T | T | T | T | A | G | C | T | C | G | C | T | C | C | A | C | C | T | A | C | C | C | C | C | T | G | C | T | A | C | A | T | G | C | G | T | G | T | T | T | A | T | T | A | T | T | T | T | A | G | A | G | G | C | C | C | C | C | C | A | C | G | A | T | C | T | A | C |  |
| BA.1.16 | C | T | C | G | G | C | A | G | C | G | G | - | - | - | C | A | A | C | C | C | G | - | - | - | - | - | G | T | T | T | T | G | G | C | C | T | G | C | C | C | C | A | T | C | T | A | C | C | C | C | C | T | G | T | - | - | - | - | - | - | T | - | - | - | - | - | - | - | - | - | - | - | - | T | C | A | G | A | A | A | A | C | A | T | C | C | G | C | G | A | T | C | T | A | T |  |
| BA.2 | C | T | C | G | G | C | A | G | C | G | G | - | - | - | C | A | A | C | C | C | G | - | - | - | - | - | G | T | T | T | T | G | G | C | C | T | G | C | C | C | C | A | T | C | T | A | C | C | C | C | C | T | G | T | - | - | - | - | - | - | T | - | - | - | - | - | - | - | - | - | - | - | - | T | C | A | A | C | G | G | C | T | C | C | T | T | A | T | C | T | C | T | C | C | C | #1130789 |
| BA.2.37 | T | G | T | A | A | T | G | A | T | A | T | G | T | T | T | G | G | T | T | T | A | G | T | T | T | G | - | - | - | - | - | A | T | T | T | C | T | T | T | T | T | G | C | T | - | - | - | - | - | - | - | - | - | C | T | A | C | A | T | G | C | A | T | G | T | T | T | A | T | T | A | T | T | G | T | G | A | C | G | G | C | T | C | C | T | T | A | T | C | T | C | T | C | C | C |  |
| mut |  |  |  |  |  |  |  |  |  |  |  |  |  |  |  |  |  |  |  |  |  |  |  |  |  |  |  |  |  |  |  |  |  |  |  |  |  |  |  |  |  |  |  |  |  |  |  |  |  |  |  |  |  |  |  |  |  |  |  |  |  |  |  |  |  |  |  |  |  |  |  |  |  |  |  |  |  |  |  |  |  |  |  |  |  |  |  |  |  |  |  |  |  |  |  |  |

|  |  |  |  |  |  |  |  |  |  |  |  |  |  |  |  |  |  |  |
| --- | --- | --- | --- | --- | --- | --- | --- | --- | --- | --- | --- | --- | --- | --- | --- | --- | --- | --- |
| pos | 1116 | 1968 | 6541 | 6762 | 7926 | 9274 | 10676 | 18816 | 21595 | 22194 | 22195 | 22196 | 22197 | 22198 | 22202 | 22204 | 29837 |  |
| ref | T | C | C | C | C | A | T | G | C | A | T | T | T | A | C | T | C |  |
| BA.1.1 | C | T | T | C | C | G | C | T | T | - | - | - | T | A | C | T | C |  |
| BA.1.1 | C | T | T | C | C | G | C | G | T | A | G | T | G | C | A | C | A | #1132214 |
| BA.1.1.18 | T | C | C | T | T | A | T | G | C | A | G | T | G | C | A | C | A |  |
| mut |  |  |  |  |  |  |  |  | C>T |  |  |  |  |  |  |  |  |  |

|  |  |  |  |  |  |  |  |  |  |  |  |  |  |  |  |
| --- | --- | --- | --- | --- | --- | --- | --- | --- | --- | --- | --- | --- | --- | --- | --- |
| pos | 105 | 7420 | 12053 | 13273 | 22194 | 22195 | 22196 | 22197 | 22198 | 22202 | 22204 | 28877 | 28878 | 29738 |  |
| ref | G | C | C | T | A | T | T | T | A | C | T | A | G | C |  |
| BA.1.1 | T | T | T | C | A | G | T | G | C | A | C | A | G | C |  |
| BA.1.1 | T | T | T | C | - | - | - | T | A | C | T | T | C | T | #1134977 |
| BA.1.18 | G | C | C | T | - | - | - | T | A | C | T | A | G | T |  |
| mut |  |  |  |  |  |  |  |  |  |  |  | A>T | G>C |  |  |

|  |  |  |  |  |  |  |  |  |  |  |  |  |  |  |  |  |
| --- | --- | --- | --- | --- | --- | --- | --- | --- | --- | --- | --- | --- | --- | --- | --- | --- |
| pos | 1454 | 3425 | 8662 | 11115 | 14940 | 21102 | 21595 | 22194 | 22195 | 22196 | 22197 | 22198 | 22202 | 22204 | 29647 |  |
| ref | C | C | T | G | A | T | C | A | T | T | T | A | C | T | A |  |
| BA.1.1 | C | T | C | T | G | C | C | - | - | - | T | A | C | T | G |  |
| BA.1.1 | T | T | C | T | G | C | C | A | G | T | G | C | A | C | G | #1137446 |
| BA.1.1 | C | C | T | G | A | T | T | A | G | T | G | C | A | C | A |  |
| mut | C>T |  |  |  |  |  |  |  |  |  |  |  |  |  | A>G |  |

|  |  |  |  |  |  |  |  |  |  |  |  |  |
| --- | --- | --- | --- | --- | --- | --- | --- | --- | --- | --- | --- | --- |
| pos | 5497 | 5575 | 19440 | 20946 | 22194 | 22195 | 22196 | 22197 | 22198 | 22202 | 22204 |  |
| ref | C | C | A | C | A | T | T | T | A | C | T |  |
| BA.1.1 | T | T | G | T | - | - | - | T | A | C | T |  |
| BA.1.1 | T | T | G | T | A | G | T | G | C | A | C | #1137450 |
| BA.1.1 | C | C | A | C | A | G | T | G | C | A | C |  |
| mut |  |  |  |  |  |  |  |  |  |  |  |  |

|  |  |  |  |  |  |  |  |  |  |  |  |  |  |
| --- | --- | --- | --- | --- | --- | --- | --- | --- | --- | --- | --- | --- | --- |
| pos | 8302 | 8983 | 8984 | 20469 | 20629 | 22194 | 22195 | 22196 | 22197 | 22198 | 22202 | 22204 |  |
| ref | A | T | G | G | C | A | T | T | T | A | C | T |  |
| BA.1.1 | A | G | A | A | C | - | - | - | T | A | C | T |  |
| BA.1.1 | A | G | A | A | T | A | G | T | G | C | A | C | #1142146 |
| BA.1.1 | G | T | G | G | T | A | G | T | G | C | A | C |  |
| mut |  |  |  |  |  |  |  |  |  |  |  |  |  |

|  |  |  |  |  |  |  |  |  |  |  |  |  |  |  |  |  |  |  |  |  |  |  |  |  |  |  |  |  |  |  |  |  |  |  |  |  |  |  |  |  |  |  |  |  |  |  |  |  |  |  |  |  |  |  |  |  |  |  |  |  |  |  |  |  |  |  |  |  |  |  |  |  |  |  |  |  |  |  |  |  |  |  |  |  |  |  |  |  |  |  |
| --- | --- | --- | --- | --- | --- | --- | --- | --- | --- | --- | --- | --- | --- | --- | --- | --- | --- | --- | --- | --- | --- | --- | --- | --- | --- | --- | --- | --- | --- | --- | --- | --- | --- | --- | --- | --- | --- | --- | --- | --- | --- | --- | --- | --- | --- | --- | --- | --- | --- | --- | --- | --- | --- | --- | --- | --- | --- | --- | --- | --- | --- | --- | --- | --- | --- | --- | --- | --- | --- | --- | --- | --- | --- | --- | --- | --- | --- | --- | --- | --- | --- | --- | --- | --- | --- | --- | --- | --- | --- | --- |
| pos | 44 | 670 | 934 | 2470 | 2790 | 2832 | 3542 | 4184 | 4321 | 5386 | 6513 | 6514 | 6515 | 7834 | 8393 | 9344 | 9424 | 9534 | 9866 | 10198 | 10447 | 11283 | 11284 | 11285 | 11286 | 11287 | 11292 | 11293 | 11294 | 11295 | 11296 | 11537 | 12880 | 13195 | 15240 | 15714 | 16342 | 17410 | 19955 | 20055 | 21618 | 21633 | 21634 | 21635 | 21636 | 21637 | 21638 | 21639 | 21640 | 21641 | 21762 | 21765 | 21766 | 21767 | 21768 | 21769 | 21770 | 21846 | 21987 | 21988 | 21989 | 21990 | 21991 | 21992 | 21993 | 21994 | 21995 | 22194 | 22195 | 22196 | 22200 | 22599 | 22673 | 22688 | 22775 | 22786 | 22898 | 23048 | 23202 | 24130 | 24503 | 25919 | 26060 | 26530 | 26858 | 27382 | 27383 | 27384 | 29510 |  |
| ref | C | T | C | C | C | A | A | G | C | T | G | T | T | C | G | C | A | C | C | C | G | G | T | T | T | G | G | T | T | T | T | A | C | T | C | C | T | C | C | A | C | T | A | C | C | C | C | C | T | G | C | T | A | C | A | T | G | C | G | T | G | T | T | T | A | T | T | A | T | T | T | G | T | A | G | A | G | G | C | C | C | C | C | A | C | G | A | T | A |  |
| BA.2.10 | T | G | T | C | T | A | A | A | T | T | G | T | T | C | G | T | G | T | T | T | A | G | T | T | T | G | - | - | - | - | - | A | T | T | C | T | C | T | T | G | T | - | - | - | - | - | - | - | - | - | C | T | A | C | A | T | G | C | A | T | G | T | T | T | A | T | T | A | T | T | G | G | T | G | A | C | G | G | C | C | C | T | T | A | T | C | T | C | C |  |
| BA.1 | T | G | T | C | T | A | G | G | C | G | - | - | - | T | A | C | A | C | C | C | G | - | - | - | - | - | G | T | T | T | T | G | C | C | T | C | T | C | C | A | C | T | A | C | C | C | C | C | T | G | T | - | - | - | - | - | - | T | - | - | - | - | - | - | - | - | - | - | - | - | T | A | C | A | G | A | A | A | A | A | T | C | C | G | C | G | A | T | A | #1145912 |
| BA.1.1 | C | T | C | T | C | G | G | G | C | G | - | - | - | T | A | C | A | C | C | C | G | - | - | - | - | - | G | T | T | T | T | G | C | C | T | C | T | C | C | A | C | T | A | C | C | C | C | C | T | G | T | - | - | - | - | - | - | T | - | - | - | - | - | - | - | - | - | - | - | - | T | A | C | A | G | A | A | A | A | A | T | C | C | G | C | G | A | T | A |  |
| mut |  |  |  |  |  |  |  |  |  |  |  |  |  |  |  |  |  |  |  |  |  |  |  |  |  |  |  |  |  |  |  |  |  |  |  |  |  |  |  |  |  |  |  |  |  |  |  |  |  |  |  |  |  |  |  |  |  |  |  |  |  |  |  |  |  |  |  |  |  |  |  |  |  |  |  |  |  |  |  |  |  |  |  |  |  |  |  |  |  |  |

|  |  |  |  |  |  |  |  |  |  |  |  |  |  |  |  |  |  |  |  |  |  |  |  |  |  |  |  |  |  |  |  |  |  |  |  |  |  |  |  |  |  |  |  |  |  |  |  |  |  |  |  |  |  |  |  |  |  |  |  |  |  |  |  |  |  |  |  |  |  |  |  |  |  |  |  |  |  |  |  |  |  |  |  |  |  |  |  |  |  |  |  |  |  |  |  |  |  |
| --- | --- | --- | --- | --- | --- | --- | --- | --- | --- | --- | --- | --- | --- | --- | --- | --- | --- | --- | --- | --- | --- | --- | --- | --- | --- | --- | --- | --- | --- | --- | --- | --- | --- | --- | --- | --- | --- | --- | --- | --- | --- | --- | --- | --- | --- | --- | --- | --- | --- | --- | --- | --- | --- | --- | --- | --- | --- | --- | --- | --- | --- | --- | --- | --- | --- | --- | --- | --- | --- | --- | --- | --- | --- | --- | --- | --- | --- | --- | --- | --- | --- | --- | --- | --- | --- | --- | --- | --- | --- | --- | --- | --- | --- | --- | --- | --- | --- |
| pos | 44 | 670 | 2106 | 2470 | 2790 | 2832 | 4184 | 4321 | 5386 | 5810 | 6513 | 6514 | 6515 | 8393 | 8638 | 8733 | 9344 | 9424 | 9534 | 9866 | 10198 | 10447 | 11283 | 11284 | 11285 | 11286 | 11287 | 11292 | 11293 | 11294 | 11295 | 11296 | 11537 | 12880 | 13195 | 13570 | 14117 | 15240 | 15714 | 17410 | 19955 | 20055 | 20718 | 21618 | 21633 | 21634 | 21635 | 21636 | 21637 | 21638 | 21639 | 21640 | 21641 | 21762 | 21765 | 21766 | 21767 | 21768 | 21769 | 21770 | 21846 | 21987 | 21988 | 21989 | 21990 | 21991 | 21992 | 21993 | 21994 | 21995 | 22194 | 22195 | 22196 | 22200 | 22599 | 22673 | 22688 | 22775 | 22786 | 22898 | 23048 | 23202 | 24130 | 24503 | 25008 | 25416 | 25725 | 26060 | 26530 | 26858 | 27382 | 27383 | 27384 | 28079 | 29510 | 29755 |  |
| ref | C | T | C | C | C | A | G | C | T | G | G | T | T | G | A | C | C | A | C | C | C | G | G | T | T | T | G | G | T | T | T | T | A | C | T | G | C | C | C | C | C | A | G | C | T | A | C | C | C | C | C | T | G | C | T | A | C | A | T | G | C | G | T | G | T | T | T | A | T | T | A | T | T | T | G | T | A | G | A | G | G | C | C | C | A | C | A | C | A | C | G | A | T | G | A | G |  |
| BA.2 | T | G | C | C | T | A | A | T | T | G | G | T | T | G | A | C | T | G | T | T | T | A | G | T | T | T | G | - | - | - | - | - | A | T | T | G | C | C | T | T | T | G | G | T | - | - | - | - | - | - | - | - | - | C | T | A | C | A | T | G | C | A | T | G | T | T | T | A | T | T | A | T | T | G | G | T | G | A | C | G | G | C | C | C | G | T | G | T | A | T | C | T | C | T | C | G |  |
| BA.2 | T | G | C | C | T | A | A | T | T | G | G | T | T | G | G | C | T | G | T | T | T | A | G | T | T | T | G | - | - | - | - | - | A | T | T | G | C | C | T | T | T | G | G | C | T | A | C | C | C | C | C | T | G | T | - | - | - | - | - | - | T | - | - | - | - | - | - | - | - | - | - | - | - | T | A | C | A | G | A | A | A | A | A | T | A | C | A | C | G | C | G | A | T | G | A | C | #1145942 |
| BA.1.1.15 | C | T | T | T | C | G | G | C | G | A | - | - | - | A | A | T | C | A | C | C | C | G | - | - | - | - | - | G | T | T | T | T | G | C | C | A | T | T | C | C | C | A | T | C | T | A | C | C | C | C | C | T | G | T | - | - | - | - | - | - | T | - | - | - | - | - | - | - | - | - | - | - | - | T | A | C | A | G | A | A | A | A | A | T | A | C | A | C | G | C | G | A | T | G | A | C |  |
| mut |  |  |  |  |  |  |  |  |  |  |  |  |  |  | A>G |  |  |  |  |  |  |  |  |  |  |  |  |  |  |  |  |  |  |  |  |  |  |  |  |  |  |  |  |  |  |  |  |  |  |  |  |  |  |  |  |  |  |  |  |  |  |  |  |  |  |  |  |  |  |  |  |  |  |  |  |  |  |  |  |  |  |  |  |  |  |  |  |  |  |  |  |  |  |  |  |  |  |

|  |  |  |  |  |  |  |  |  |  |  |  |  |  |  |  |  |  |  |  |  |  |  |  |  |  |  |  |  |  |  |  |  |  |  |  |  |  |  |  |  |  |  |  |  |  |  |  |  |  |  |  |  |  |  |  |  |  |  |  |  |  |  |  |  |  |  |  |  |  |  |  |  |  |  |  |  |  |  |  |  |  |  |  |  |  |  |  |  |  |  |  |  |  |  |  |  |  |  |
| --- | --- | --- | --- | --- | --- | --- | --- | --- | --- | --- | --- | --- | --- | --- | --- | --- | --- | --- | --- | --- | --- | --- | --- | --- | --- | --- | --- | --- | --- | --- | --- | --- | --- | --- | --- | --- | --- | --- | --- | --- | --- | --- | --- | --- | --- | --- | --- | --- | --- | --- | --- | --- | --- | --- | --- | --- | --- | --- | --- | --- | --- | --- | --- | --- | --- | --- | --- | --- | --- | --- | --- | --- | --- | --- | --- | --- | --- | --- | --- | --- | --- | --- | --- | --- | --- | --- | --- | --- | --- | --- | --- | --- | --- | --- | --- | --- | --- | --- |
| pos | 670 | 1585 | 2470 | 2790 | 2832 | 4184 | 4321 | 5386 | 5619 | 6513 | 6514 | 6515 | 8393 | 9344 | 9424 | 9534 | 9866 | 10198 | 10447 | 11283 | 11284 | 11285 | 11286 | 11287 | 11292 | 11293 | 11294 | 11295 | 11296 | 11537 | 12312 | 12754 | 12880 | 13195 | 15240 | 15714 | 15738 | 16293 | 17410 | 19955 | 20055 | 21618 | 21633 | 21634 | 21635 | 21636 | 21637 | 21638 | 21639 | 21640 | 21641 | 21762 | 21765 | 21766 | 21767 | 21768 | 21769 | 21770 | 21846 | 21987 | 21988 | 21989 | 21990 | 21991 | 21992 | 21993 | 21994 | 21995 | 22194 | 22195 | 22196 | 22200 | 22599 | 22673 | 22688 | 22775 | 22786 | 22792 | 22898 | 23048 | 23202 | 24130 | 24503 | 25437 | 25624 | 26060 | 26530 | 26858 | 27382 | 27383 | 27384 | 27403 | 29510 | 29752 | 29762 | 29769 | 29774 |  |
| ref | T | A | C | C | A | G | C | T | T | G | T | T | G | C | A | C | C | C | G | G | T | T | T | G | G | T | T | T | T | A | C | C | C | T | C | C | C | C | C | C | A | C | T | A | C | C | C | C | C | T | G | C | T | A | C | A | T | G | C | G | T | G | T | T | T | A | T | T | A | T | T | T | G | T | A | G | A | C | G | G | C | C | C | G | C | C | A | C | G | A | T | A | A | A | C | C | C |  |
| BA.2.9 | G | C | C | T | A | A | T | T | C | G | T | T | G | T | G | T | T | T | A | G | T | T | T | G | - | - | - | - | - | A | C | T | T | T | C | T | T | T | T | T | G | T | - | - | - | - | - | - | - | - | - | C | T | A | C | A | T | G | C | A | T | G | T | T | T | A | T | T | A | T | T | G | G | T | G | A | C | T | G | G | C | C | C | T | T | T | A | T | C | T | C | A | C | A | - | - | - |  |
| BA.2 | G | C | C | T | A | A | T | T | C | G | T | T | G | T | G | T | T | T | A | G | T | T | T | G | - | - | - | - | - | A | C | T | T | T | C | T | T | T | T | T | G | T | - | - | - | - | - | - | - | - | - | C | T | A | C | A | T | G | C | A | T | G | T | T | T | A | T | T | A | T | T | G | G | T | G | A | C | T | G | G | C | C | C | T | T | T | A | C | G | A | T | G | A | T | C | C | C | #1147618 |
| BA.1.1 | T | A | T | C | G | G | C | G | T | - | - | - | A | C | A | C | C | C | G | - | - | - | - | - | G | T | T | T | T | G | T | C | C | C | T | C | C | C | C | C | A | C | T | A | C | C | C | C | C | T | G | T | - | - | - | - | - | - | T | - | - | - | - | - | - | - | - | - | - | - | - | T | A | C | A | G | A | C | A | A | A | A | T | G | C | C | G | C | G | A | T | G | A | T | C | C | C |  |
| mut |  |  |  |  |  |  |  |  |  |  |  |  |  |  |  |  |  |  |  |  |  |  |  |  |  |  |  |  |  |  |  |  |  |  |  |  |  |  |  |  |  |  |  |  |  |  |  |  |  |  |  |  |  |  |  |  |  |  |  |  |  |  |  |  |  |  |  |  |  |  |  |  |  |  |  |  |  |  |  |  |  |  |  |  |  |  |  |  |  |  |  |  |  |  |  |  |  |  |

|  |  |  |  |  |  |  |  |  |  |  |  |  |  |  |  |
| --- | --- | --- | --- | --- | --- | --- | --- | --- | --- | --- | --- | --- | --- | --- | --- |
| pos | 9439 | 13994 | 16221 | 17550 | 18253 | 19512 | 22194 | 22195 | 22196 | 22197 | 22198 | 22202 | 22204 | 29754 |  |
| ref | A | C | G | C | A | C | A | T | T | T | A | C | T | C |  |
| BA.1.1 | A | T | A | T | G | T | - | - | - | T | A | C | T | C |  |
| BA.1.1 | A | T | A | T | G | T | A | G | T | G | C | A | C | T | #1148133 |
| BA.1.1 | G | C | G | C | A | C | A | G | T | G | C | A | C | C |  |
| mut |  |  |  |  |  |  |  |  |  |  |  |  |  | C>T |  |

|  |  |  |  |  |  |  |  |  |  |  |  |  |  |  |  |  |
| --- | --- | --- | --- | --- | --- | --- | --- | --- | --- | --- | --- | --- | --- | --- | --- | --- |
| pos | 1599 | 1623 | 7119 | 13665 | 18958 | 20117 | 21589 | 21595 | 22194 | 22195 | 22196 | 22197 | 22198 | 22202 | 22204 |  |
| ref | G | T | C | C | G | C | A | C | A | T | T | T | A | C | T |  |
| BA.1.1 | G | C | T | T | A | T | C | T | A | G | T | G | C | A | C |  |
| BA.1.1 | G | C | T | T | A | T | C | T | - | - | - | T | A | C | T | #1150594 |
| BA.1.1 | T | T | C | C | G | C | A | C | - | - | - | T | A | C | T |  |
| mut |  |  |  |  |  |  |  |  |  |  |  |  |  |  |  |  |

|  |  |  |  |  |  |  |  |  |  |  |  |  |  |  |  |
| --- | --- | --- | --- | --- | --- | --- | --- | --- | --- | --- | --- | --- | --- | --- | --- |
| pos | 1599 | 2783 | 4832 | 9848 | 13288 | 21595 | 21772 | 22194 | 22195 | 22196 | 22197 | 22198 | 22202 | 22204 |  |
| ref | G | A | C | A | A | C | C | A | T | T | T | A | C | T |  |
| BA.1.1 | G | G | T | G | G | T | T | A | G | T | G | C | A | C |  |
| BA.1.1 | G | G | T | G | G | T | T | - | - | - | T | A | C | T | #1150753 |
| BA.1.1 | T | A | C | A | A | C | C | - | - | - | T | A | C | T |  |
| mut |  |  |  |  |  |  |  |  |  |  |  |  |  |  |  |

|  |  |  |  |  |  |  |  |  |  |  |  |  |  |  |  |  |  |  |  |  |  |  |  |  |  |  |  |  |  |  |  |  |  |  |  |  |  |  |  |  |  |  |  |  |  |  |  |  |  |  |  |  |  |  |  |  |  |  |  |  |  |  |  |  |  |  |  |  |  |  |  |  |  |  |  |  |  |  |  |  |  |  |  |  |  |  |  |  |  |  |  |  |  |
| --- | --- | --- | --- | --- | --- | --- | --- | --- | --- | --- | --- | --- | --- | --- | --- | --- | --- | --- | --- | --- | --- | --- | --- | --- | --- | --- | --- | --- | --- | --- | --- | --- | --- | --- | --- | --- | --- | --- | --- | --- | --- | --- | --- | --- | --- | --- | --- | --- | --- | --- | --- | --- | --- | --- | --- | --- | --- | --- | --- | --- | --- | --- | --- | --- | --- | --- | --- | --- | --- | --- | --- | --- | --- | --- | --- | --- | --- | --- | --- | --- | --- | --- | --- | --- | --- | --- | --- | --- | --- | --- | --- | --- | --- |
| pos | 670 | 2790 | 2832 | 4184 | 4321 | 5386 | 6079 | 6513 | 6514 | 6515 | 7528 | 8393 | 9344 | 9424 | 9534 | 9866 | 10135 | 10198 | 10447 | 10456 | 11283 | 11284 | 11285 | 11286 | 11287 | 11292 | 11293 | 11294 | 11295 | 11296 | 11537 | 12880 | 13195 | 15240 | 15714 | 17410 | 19374 | 19955 | 20055 | 21618 | 21633 | 21634 | 21635 | 21636 | 21637 | 21638 | 21639 | 21640 | 21641 | 21762 | 21765 | 21766 | 21767 | 21768 | 21769 | 21770 | 21846 | 21987 | 21988 | 21989 | 21990 | 21991 | 21992 | 21993 | 21994 | 21995 | 22194 | 22195 | 22196 | 22200 | 22673 | 22688 | 22775 | 22786 | 22898 | 23048 | 23202 | 23230 | 24023 | 24130 | 24503 | 25708 | 26060 | 26530 | 26858 | 27382 | 27383 | 27384 | 27509 | 29301 | 29510 | 29734 |  |
| ref | T | C | A | G | C | T | T | G | T | T | C | G | C | A | C | C | T | C | G | C | G | T | T | T | G | G | T | T | T | T | A | C | T | C | C | C | C | C | A | C | T | A | C | C | C | C | C | T | G | C | T | A | C | A | T | G | C | G | T | G | T | T | T | A | T | T | A | T | T | T | T | A | G | A | G | G | C | C | C | C | C | C | C | A | C | G | A | T | C | A | A | G |  |
| BA.1.15 | T | C | G | G | C | G | T | - | - | - | C | A | C | A | C | C | C | C | G | T | - | - | - | - | - | G | T | T | T | T | G | C | C | T | C | C | C | C | A | C | T | A | C | C | C | C | C | T | G | T | - | - | - | - | - | - | T | - | - | - | - | - | - | - | - | - | - | - | - | T | C | A | G | A | A | A | A | C | C | A | T | T | C | G | C | G | A | T | C | G | A | G |  |
| BA.2 | T | C | G | G | C | G | T | - | - | - | T | A | C | A | C | C | C | C | G | T | - | - | - | - | - | G | T | T | T | T | G | C | C | T | C | C | T | C | A | T | - | - | - | - | - | - | - | - | - | C | T | A | C | A | T | G | C | A | T | G | T | T | T | A | T | T | A | T | T | G | T | G | A | C | G | G | C | T | C | C | C | C | T | A | T | C | T | C | T | A | A | C | #1154548 |
| BA.2 | G | T | A | A | T | T | C | G | T | T | C | G | T | G | T | T | T | T | A | C | G | T | T | T | G | - | - | - | - | - | A | T | T | C | T | T | C | T | G | T | - | - | - | - | - | - | - | - | - | C | T | A | C | A | T | G | C | A | T | G | T | T | T | A | T | T | A | T | T | G | T | G | A | C | G | G | C | T | T | C | C | C | T | A | T | C | T | C | T | A | C | G |  |
| mut |  |  |  |  |  |  |  |  |  |  | C>T |  |  |  |  |  |  |  |  |  |  |  |  |  |  |  |  |  |  |  |  |  |  |  |  |  | C>T |  |  |  |  |  |  |  |  |  |  |  |  |  |  |  |  |  |  |  |  |  |  |  |  |  |  |  |  |  |  |  |  |  |  |  |  |  |  |  |  |  | T>C |  |  |  |  |  |  |  |  |  |  |  | C>A | G>C |  |

|  |  |  |  |  |  |  |  |  |  |  |  |  |  |  |  |  |  |  |  |  |  |  |  |  |  |  |  |  |  |  |  |  |  |  |  |  |  |  |  |  |  |  |  |  |  |  |  |  |  |  |  |  |  |  |  |  |  |  |  |  |  |  |  |  |  |  |  |  |  |  |  |  |  |  |  |  |  |  |  |  |  |  |  |
| --- | --- | --- | --- | --- | --- | --- | --- | --- | --- | --- | --- | --- | --- | --- | --- | --- | --- | --- | --- | --- | --- | --- | --- | --- | --- | --- | --- | --- | --- | --- | --- | --- | --- | --- | --- | --- | --- | --- | --- | --- | --- | --- | --- | --- | --- | --- | --- | --- | --- | --- | --- | --- | --- | --- | --- | --- | --- | --- | --- | --- | --- | --- | --- | --- | --- | --- | --- | --- | --- | --- | --- | --- | --- | --- | --- | --- | --- | --- | --- | --- | --- | --- | --- |
| pos | 670 | 2470 | 2790 | 2832 | 4184 | 4321 | 5386 | 6166 | 6513 | 6514 | 6515 | 8393 | 9344 | 9424 | 9534 | 10198 | 10447 | 11283 | 11284 | 11285 | 11286 | 11287 | 11292 | 11293 | 11294 | 11295 | 11296 | 11537 | 12880 | 13195 | 15240 | 15714 | 17410 | 21595 | 21618 | 21633 | 21634 | 21635 | 21636 | 21637 | 21638 | 21639 | 21640 | 21641 | 21762 | 21765 | 21766 | 21767 | 21768 | 21769 | 21770 | 21846 | 21987 | 21988 | 21989 | 21990 | 21991 | 21992 | 21993 | 21994 | 21995 | 22194 | 22195 | 22196 | 22200 | 22599 | 22673 | 22688 | 22775 | 22786 | 22898 | 23048 | 23202 | 24130 | 24503 | 26060 | 26530 | 26858 | 27382 | 27383 | 27384 | 29510 |  |
| ref | T | C | C | A | G | C | T | G | G | T | T | G | C | A | C | C | G | G | T | T | T | G | G | T | T | T | T | A | C | T | C | C | C | C | C | T | A | C | C | C | C | C | T | G | C | T | A | C | A | T | G | C | G | T | G | T | T | T | A | T | T | A | T | T | T | G | T | A | G | A | G | G | C | C | C | C | A | C | G | A | T | A |  |
| BA.1.1 | T | T | C | G | G | C | G | A | - | - | - | A | C | A | C | C | G | - | - | - | - | - | G | T | T | T | T | G | C | C | T | C | C | T | C | T | A | C | C | C | C | C | T | G | T | - | - | - | - | - | - | T | - | - | - | - | - | - | - | - | - | - | - | - | T | A | C | A | G | A | A | A | A | A | T | C | G | C | G | A | T | A |  |
| XM | T | T | C | G | G | C | G | A | - | - | - | A | C | A | C | C | G | - | - | - | - | - | G | T | T | T | T | G | C | C | T | C | C | C | C | T | A | C | C | C | C | C | T | G | T | - | - | - | - | - | - | T | - | - | - | - | - | - | - | - | - | A | T | T | G | G | T | G | A | C | G | G | C | C | C | T | A | T | C | T | C | C | #1158127 |
| BA.2 | G | C | T | A | A | T | T | G | G | T | T | G | T | G | T | T | A | G | T | T | T | G | - | - | - | - | - | A | T | T | C | T | T | C | T | - | - | - | - | - | - | - | - | - | C | T | A | C | A | T | G | C | A | T | G | T | T | T | A | T | T | A | T | T | G | G | T | G | A | C | G | G | C | C | C | T | A | T | C | T | C | C |  |
| mut |  |  |  |  |  |  |  |  |  |  |  |  |  |  |  |  |  |  |  |  |  |  |  |  |  |  |  |  |  |  |  |  |  | T>C |  |  |  |  |  |  |  |  |  |  |  |  |  |  |  |  |  |  |  |  |  |  |  |  |  |  |  |  |  |  |  |  |  |  |  |  |  |  |  |  |  |  |  |  |  |  |  |  |  |

|  |  |  |  |  |  |  |  |  |  |  |  |  |  |  |  |  |  |  |  |  |  |  |  |  |  |  |  |  |  |  |  |  |  |  |  |  |  |  |  |  |  |  |  |  |  |  |  |  |  |  |  |  |  |  |  |  |  |  |  |  |  |  |  |  |  |  |  |  |  |  |  |  |  |  |  |  |  |  |  |  |  |  |  |  |  |  |  |  |  |  |  |  |  |  |  |  |  |  |  |  |
| --- | --- | --- | --- | --- | --- | --- | --- | --- | --- | --- | --- | --- | --- | --- | --- | --- | --- | --- | --- | --- | --- | --- | --- | --- | --- | --- | --- | --- | --- | --- | --- | --- | --- | --- | --- | --- | --- | --- | --- | --- | --- | --- | --- | --- | --- | --- | --- | --- | --- | --- | --- | --- | --- | --- | --- | --- | --- | --- | --- | --- | --- | --- | --- | --- | --- | --- | --- | --- | --- | --- | --- | --- | --- | --- | --- | --- | --- | --- | --- | --- | --- | --- | --- | --- | --- | --- | --- | --- | --- | --- | --- | --- | --- | --- | --- | --- | --- | --- | --- | --- |
| pos | 670 | 2470 | 2790 | 2832 | 4184 | 4321 | 4475 | 5386 | 6513 | 6514 | 6515 | 7104 | 8393 | 9344 | 9424 | 9534 | 9866 | 10198 | 10447 | 10486 | 10697 | 11283 | 11284 | 11285 | 11286 | 11287 | 11292 | 11293 | 11294 | 11295 | 11296 | 11537 | 12880 | 13195 | 13945 | 14940 | 15240 | 15714 | 16323 | 17410 | 17746 | 18483 | 19955 | 20055 | 20178 | 21618 | 21633 | 21634 | 21635 | 21636 | 21637 | 21638 | 21639 | 21640 | 21641 | 21762 | 21765 | 21766 | 21767 | 21768 | 21769 | 21770 | 21846 | 21987 | 21988 | 21989 | 21990 | 21991 | 21992 | 21993 | 21994 | 21995 | 22194 | 22195 | 22196 | 22200 | 22599 | 22673 | 22688 | 22775 | 22786 | 22792 | 22898 | 23048 | 23202 | 24130 | 24503 | 25844 | 26060 | 26133 | 26408 | 26530 | 26858 | 27382 | 27383 | 27384 | 28877 | 28878 | 29510 |  |
| ref | T | C | C | A | G | C | C | T | G | T | T | C | G | C | A | C | C | C | G | A | G | G | T | T | T | G | G | T | T | T | T | A | C | T | C | A | C | C | C | C | C | C | C | A | C | C | T | A | C | C | C | C | C | T | G | C | T | A | C | A | T | G | C | G | T | G | T | T | T | A | T | T | A | T | T | T | G | T | A | G | A | C | G | G | C | C | C | C | C | C | C | A | C | G | A | T | A | G | A |  |
| BA.1.1 | T | T | C | G | G | C | C | G | - | - | - | T | A | C | A | C | C | C | G | G | G | - | - | - | - | - | G | T | T | T | T | G | C | C | C | G | T | C | T | C | T | T | C | A | C | C | T | A | C | C | C | C | C | T | G | T | - | - | - | - | - | - | T | - | - | - | - | - | - | - | - | - | - | - | - | T | A | C | A | G | A | C | A | A | A | A | T | T | C | C | C | G | C | G | A | T | A | G | A |  |
| XM | T | T | C | G | G | C | T | G | - | - | - | T | A | C | A | C | C | C | G | G | A | - | - | - | - | - | G | T | T | T | T | G | C | C | C | G | T | C | T | C | T | T | C | A | C | T | - | - | - | - | - | - | - | - | - | C | T | A | C | A | T | G | C | A | T | G | T | T | T | A | T | T | A | T | T | G | G | T | G | A | C | C | G | G | C | C | C | C | T | T | T | A | T | C | T | C | T | C | C | #1158182 |
| BA.2 | G | C | T | A | A | T | C | T | G | T | T | C | G | T | G | T | T | T | A | A | G | G | T | T | T | G | - | - | - | - | - | A | T | T | T | A | C | T | C | T | C | C | T | G | T | T | - | - | - | - | - | - | - | - | - | C | T | A | C | A | T | G | C | A | T | G | T | T | T | A | T | T | A | T | T | G | G | T | G | A | C | T | G | G | C | C | C | C | T | T | C | A | T | C | T | C | A | G | C |  |
| mut |  |  |  |  |  |  | C>T |  |  |  |  |  |  |  |  |  |  |  |  |  | G>A |  |  |  |  |  |  |  |  |  |  |  |  |  |  |  |  |  |  |  |  |  |  |  |  |  |  |  |  |  |  |  |  |  |  |  |  |  |  |  |  |  |  |  |  |  |  |  |  |  |  |  |  |  |  |  |  |  |  |  |  | T>C |  |  |  |  |  |  |  |  | C>T |  |  |  |  |  | A>T | G>C |  |  |

|  |  |  |  |  |  |  |  |  |  |  |  |  |  |  |  |  |  |  |  |  |
| --- | --- | --- | --- | --- | --- | --- | --- | --- | --- | --- | --- | --- | --- | --- | --- | --- | --- | --- | --- | --- |
| pos | 2470 | 3828 | 5230 | 10408 | 14318 | 17304 | 21595 | 22194 | 22195 | 22196 | 22197 | 22198 | 22202 | 22204 | 22599 | 26491 | 26492 | 26497 | 27393 |  |
| ref | C | C | G | C | C | C | C | A | T | T | T | A | C | T | G | T | A | T | C |  |
| BA.1.1 | T | T | G | T | A | T | T | A | G | T | G | C | A | C | A | T | A | T | T |  |
| BA.1.1 | T | T | G | T | A | T | T | - | - | - | T | A | C | T | A | C | T | C | T | #1158755 |
| BA.1 | C | C | A | C | C | C | C | - | - | - | T | A | C | T | G | C | T | C | C |  |
| mut |  |  |  |  |  |  |  |  |  |  |  |  |  |  | G>A |  |  |  | C>T |  |

|  |  |  |  |  |  |  |  |  |  |  |  |  |  |  |  |  |  |
| --- | --- | --- | --- | --- | --- | --- | --- | --- | --- | --- | --- | --- | --- | --- | --- | --- | --- |
| pos | 437 | 1371 | 1868 | 4965 | 5343 | 9592 | 12783 | 20762 | 22194 | 22195 | 22196 | 22197 | 22198 | 22202 | 22204 | 24226 |  |
| ref | A | A | G | C | C | C | A | C | A | T | T | T | A | C | T | T |  |
| BA.1.15 | G | A | A | C | T | C | G | C | - | - | - | T | A | C | T | T |  |
| BA.1.15 | G | A | A | C | T | T | G | C | A | G | T | G | C | A | C | C | #1158773 |
| BA.1.15 | A | G | G | T | C | C | A | T | A | G | T | G | C | A | C | T |  |
| mut |  |  |  |  |  | C>T |  |  |  |  |  |  |  |  |  | T>C |  |

|  |  |  |  |  |  |  |  |  |  |  |  |  |  |  |  |  |  |  |  |  |  |  |  |  |  |  |  |  |  |  |  |  |  |  |  |  |  |  |  |  |  |  |  |  |  |  |  |  |  |  |  |  |  |  |  |  |  |  |  |  |  |  |  |  |  |  |  |  |  |  |  |  |  |  |  |  |  |  |  |  |  |  |  |  |  |  |  |  |  |  |  |
| --- | --- | --- | --- | --- | --- | --- | --- | --- | --- | --- | --- | --- | --- | --- | --- | --- | --- | --- | --- | --- | --- | --- | --- | --- | --- | --- | --- | --- | --- | --- | --- | --- | --- | --- | --- | --- | --- | --- | --- | --- | --- | --- | --- | --- | --- | --- | --- | --- | --- | --- | --- | --- | --- | --- | --- | --- | --- | --- | --- | --- | --- | --- | --- | --- | --- | --- | --- | --- | --- | --- | --- | --- | --- | --- | --- | --- | --- | --- | --- | --- | --- | --- | --- | --- | --- | --- | --- | --- | --- | --- | --- |
| pos | 44 | 670 | 2470 | 2790 | 2832 | 4321 | 5386 | 6513 | 6514 | 6515 | 8393 | 9344 | 9424 | 9534 | 10198 | 10447 | 10507 | 11283 | 11284 | 11285 | 11286 | 11287 | 11292 | 11293 | 11294 | 11295 | 11296 | 11537 | 12756 | 12880 | 13195 | 13570 | 14117 | 15240 | 15714 | 16020 | 17410 | 19955 | 20055 | 21618 | 21633 | 21634 | 21635 | 21636 | 21637 | 21638 | 21639 | 21640 | 21641 | 21762 | 21765 | 21766 | 21767 | 21768 | 21769 | 21770 | 21846 | 21987 | 21988 | 21989 | 21990 | 21991 | 21992 | 21993 | 21994 | 21995 | 22194 | 22195 | 22196 | 22200 | 22599 | 22673 | 22688 | 22775 | 22786 | 22898 | 23048 | 23202 | 24130 | 24503 | 25002 | 26060 | 26530 | 26858 | 27382 | 27383 | 27384 | 29510 | 29744 | 29755 |  |
| ref | C | T | C | C | A | C | T | G | T | T | G | C | A | C | C | G | C | G | T | T | T | G | G | T | T | T | T | A | C | C | T | G | C | C | C | G | C | C | A | C | T | A | C | C | C | C | C | T | G | C | T | A | C | A | T | G | C | G | T | G | T | T | T | A | T | T | A | T | T | T | G | T | A | G | A | G | G | C | C | C | C | C | A | C | G | A | T | A | G | G |  |
| BA.1.1.15 | C | T | T | C | G | C | G | - | - | - | A | C | A | C | C | G | C | - | - | - | - | - | G | T | T | T | T | G | C | C | C | A | T | T | C | G | C | C | A | C | T | A | C | C | C | C | C | T | G | T | - | - | - | - | - | - | T | - | - | - | - | - | - | - | - | - | - | - | - | T | A | C | A | G | A | A | A | A | A | T | T | C | G | C | G | A | T | A | T | C |  |
| XW | C | T | T | C | G | T | T | G | T | T | G | T | G | T | T | A | T | G | T | T | T | G | - | - | - | - | - | A | T | T | T | G | C | C | T | T | T | T | G | T | - | - | - | - | - | - | - | - | - | C | T | A | C | A | T | G | C | A | T | G | T | T | T | A | T | T | A | T | T | G | G | T | G | A | C | G | G | C | C | C | C | T | A | T | C | T | C | C | G | G | #1159411 |
| BA.2 | T | G | C | T | A | T | T | G | T | T | G | T | G | T | T | A | T | G | T | T | T | G | - | - | - | - | - | A | C | T | T | G | C | C | T | T | T | T | G | T | - | - | - | - | - | - | - | - | - | C | T | A | C | A | T | G | C | A | T | G | T | T | T | A | T | T | A | T | T | G | G | T | G | A | C | G | G | C | C | C | C | T | A | T | C | T | C | C | G | G |  |
| mut |  |  |  |  |  |  |  |  |  |  |  |  |  |  |  |  |  |  |  |  |  |  |  |  |  |  |  |  | C>T |  |  |  |  |  |  |  |  |  |  |  |  |  |  |  |  |  |  |  |  |  |  |  |  |  |  |  |  |  |  |  |  |  |  |  |  |  |  |  |  |  |  |  |  |  |  |  |  |  |  |  |  |  |  |  |  |  |  |  |  |  |  |

|  |  |  |  |  |  |  |  |  |  |  |  |  |  |  |  |  |  |  |  |  |  |  |  |  |  |  |  |  |  |  |  |  |  |  |  |  |  |  |  |  |  |  |  |  |  |  |  |  |  |  |  |  |  |  |  |  |  |  |  |  |  |  |  |  |  |  |  |  |  |  |  |  |  |  |  |  |  |  |  |  |  |  |  |  |  |  |  |  |
| --- | --- | --- | --- | --- | --- | --- | --- | --- | --- | --- | --- | --- | --- | --- | --- | --- | --- | --- | --- | --- | --- | --- | --- | --- | --- | --- | --- | --- | --- | --- | --- | --- | --- | --- | --- | --- | --- | --- | --- | --- | --- | --- | --- | --- | --- | --- | --- | --- | --- | --- | --- | --- | --- | --- | --- | --- | --- | --- | --- | --- | --- | --- | --- | --- | --- | --- | --- | --- | --- | --- | --- | --- | --- | --- | --- | --- | --- | --- | --- | --- | --- | --- | --- | --- | --- | --- | --- | --- |
| pos | 670 | 2470 | 2790 | 2832 | 4184 | 4321 | 5386 | 6513 | 6514 | 6515 | 8393 | 8739 | 9344 | 9424 | 9534 | 9866 | 10198 | 10447 | 11283 | 11284 | 11285 | 11286 | 11287 | 11292 | 11293 | 11294 | 11295 | 11296 | 11537 | 12880 | 13195 | 14913 | 15240 | 15714 | 17410 | 19955 | 20055 | 21618 | 21633 | 21634 | 21635 | 21636 | 21637 | 21638 | 21639 | 21640 | 21641 | 21762 | 21765 | 21766 | 21767 | 21768 | 21769 | 21770 | 21846 | 21987 | 21988 | 21989 | 21990 | 21991 | 21992 | 21993 | 21994 | 21995 | 22194 | 22195 | 22196 | 22200 | 22599 | 22673 | 22688 | 22775 | 22786 | 22898 | 23048 | 23202 | 24130 | 24503 | 26060 | 26078 | 26530 | 26858 | 27382 | 27383 | 27384 | 28853 | 29510 |  |
| ref | T | C | C | A | G | C | T | G | T | T | G | C | C | A | C | C | C | G | G | T | T | T | G | G | T | T | T | T | A | C | T | C | C | C | C | C | A | C | T | A | C | C | C | C | C | T | G | C | T | A | C | A | T | G | C | G | T | G | T | T | T | A | T | T | A | T | T | T | G | T | A | G | A | G | G | C | C | C | C | C | A | C | G | A | T | T | A |  |
| BA.2 | G | C | T | A | A | T | T | G | T | T | G | T | T | G | T | T | T | A | G | T | T | T | G | - | - | - | - | - | A | T | T | C | C | T | T | T | G | T | - | - | - | - | - | - | - | - | - | C | T | A | C | A | T | G | C | A | T | G | T | T | T | A | T | T | A | T | T | G | G | T | G | A | C | G | G | C | C | C | T | C | A | T | C | T | C | T | C |  |
| BA.1 | G | C | T | A | A | T | T | G | T | T | G | T | T | G | T | T | T | A | - | - | - | - | - | G | T | T | T | T | G | C | C | T | T | C | C | C | A | C | T | A | C | C | C | C | C | T | G | T | - | - | - | - | - | - | T | - | - | - | - | - | - | - | - | - | - | - | - | T | A | C | A | G | A | A | A | A | A | T | C | T | G | C | G | A | T | C | A | #1162485 |
| BA.1.1.13 | T | T | C | G | G | C | G | - | - | - | A | C | C | A | C | C | C | G | - | - | - | - | - | G | T | T | T | T | G | C | C | T | T | C | C | C | A | C | T | A | C | C | C | C | C | T | G | T | - | - | - | - | - | - | T | - | - | - | - | - | - | - | - | - | - | - | - | T | A | C | A | G | A | A | A | A | A | T | C | T | G | C | G | A | T | C | A |  |
| mut |  |  |  |  |  |  |  |  |  |  |  |  |  |  |  |  |  |  |  |  |  |  |  |  |  |  |  |  |  |  |  |  |  |  |  |  |  |  |  |  |  |  |  |  |  |  |  |  |  |  |  |  |  |  |  |  |  |  |  |  |  |  |  |  |  |  |  |  |  |  |  |  |  |  |  |  |  |  |  |  |  |  |  |  |  |  |  |  |

|  |  |  |  |  |  |  |  |  |  |  |  |  |  |  |
| --- | --- | --- | --- | --- | --- | --- | --- | --- | --- | --- | --- | --- | --- | --- |
| pos | 319 | 3122 | 8302 | 11536 | 21644 | 22194 | 22195 | 22196 | 22197 | 22198 | 22202 | 22204 | 22882 |  |
| ref | G | G | A | T | T | A | T | T | T | A | C | T | T |  |
| BA.1.1 | A | A | A | C | C | A | G | T | G | C | A | C | G |  |
| BA.1.1 | A | A | A | C | C | - | - | - | T | A | C | T | T | #1162861 |
| BA.1.1 | G | G | G | T | T | - | - | - | T | A | C | T | G |  |
| mut |  |  |  |  |  |  |  |  |  |  |  |  | G>T |  |

|  |  |  |  |  |  |  |  |  |  |  |  |  |  |  |
| --- | --- | --- | --- | --- | --- | --- | --- | --- | --- | --- | --- | --- | --- | --- |
| pos | 3088 | 4039 | 5025 | 6968 | 7926 | 21595 | 22194 | 22195 | 22196 | 22197 | 22198 | 22202 | 22204 |  |
| ref | T | C | C | C | C | C | A | T | T | T | A | C | T |  |
| BA.1.1 | A | T | T | T | C | T | - | - | - | T | A | C | T |  |
| BA.1.1 | A | T | T | T | C | T | A | G | T | G | C | A | C | #1165931 |
| BA.1.1.18 | T | C | C | C | T | C | A | G | T | G | C | A | C |  |
| mut |  |  |  |  |  |  |  |  |  |  |  |  |  |  |

|  |  |  |  |  |  |  |  |  |  |  |  |  |  |  |  |  |
| --- | --- | --- | --- | --- | --- | --- | --- | --- | --- | --- | --- | --- | --- | --- | --- | --- |
| pos | 4197 | 5513 | 11674 | 15009 | 16342 | 21721 | 22917 | 23415 | 23673 | 25416 | 27874 | 27875 | 27881 | 27882 | 27883 |  |
| ref | A | G | C | T | T | C | T | C | C | C | C | T | C | G | C |  |
| BA.2.12.1 | A | G | T | C | T | T | A | C | T | T | C | T | C | G | C |  |
| BA.2.12.1 | A | A | T | C | T | T | A | T | T | T | T | C | T | C | T | #1166157 |
| BA.2.10 | C | G | C | T | C | C | T | C | C | C | T | C | T | C | T |  |
| mut |  | G>A |  |  |  |  |  | C>T |  |  |  |  |  |  |  |  |

|  |  |  |  |  |  |  |  |  |  |  |  |  |  |  |  |  |  |  |  |
| --- | --- | --- | --- | --- | --- | --- | --- | --- | --- | --- | --- | --- | --- | --- | --- | --- | --- | --- | --- |
| pos | 2470 | 3583 | 4399 | 11455 | 13069 | 15257 | 19152 | 19688 | 21595 | 22194 | 22195 | 22196 | 22197 | 22198 | 22202 | 22204 | 22599 | 28863 |  |
| ref | C | C | G | C | T | A | C | C | C | A | T | T | T | A | C | T | G | C |  |
| BA.1.1 | T | T | G | T | T | G | T | A | T | - | - | - | T | A | C | T | A | C |  |
| BA.1.1 | T | T | A | T | A | G | T | A | T | A | G | T | G | C | A | C | A | T | #1166222 |
| BA.1 | C | C | G | C | T | A | C | C | C | A | G | T | G | C | A | C | G | C |  |
| mut |  |  | G>A |  | T>A |  |  |  |  |  |  |  |  |  |  |  | G>A | C>T |  |

|  |  |  |  |  |  |  |  |  |  |  |  |  |  |  |  |  |  |  |  |  |  |  |  |  |  |  |  |  |  |  |  |  |  |  |  |  |  |  |  |  |  |  |  |  |  |  |  |  |  |  |  |  |  |  |  |  |  |  |  |  |  |  |  |  |  |  |  |  |  |  |  |  |  |  |  |  |  |  |  |  |  |  |  |  |  |  |  |  |
| --- | --- | --- | --- | --- | --- | --- | --- | --- | --- | --- | --- | --- | --- | --- | --- | --- | --- | --- | --- | --- | --- | --- | --- | --- | --- | --- | --- | --- | --- | --- | --- | --- | --- | --- | --- | --- | --- | --- | --- | --- | --- | --- | --- | --- | --- | --- | --- | --- | --- | --- | --- | --- | --- | --- | --- | --- | --- | --- | --- | --- | --- | --- | --- | --- | --- | --- | --- | --- | --- | --- | --- | --- | --- | --- | --- | --- | --- | --- | --- | --- | --- | --- | --- | --- | --- | --- | --- | --- |
| pos | 670 | 2081 | 2470 | 2790 | 2832 | 4184 | 4321 | 5386 | 5497 | 6513 | 6514 | 6515 | 8393 | 9344 | 9424 | 9534 | 9866 | 10198 | 10447 | 11283 | 11284 | 11285 | 11286 | 11287 | 11292 | 11293 | 11294 | 11295 | 11296 | 11537 | 12306 | 12880 | 13195 | 15240 | 15714 | 17410 | 19955 | 20055 | 21595 | 21618 | 21633 | 21634 | 21635 | 21636 | 21637 | 21638 | 21639 | 21640 | 21641 | 21762 | 21765 | 21766 | 21767 | 21768 | 21769 | 21770 | 21846 | 21987 | 21988 | 21989 | 21990 | 21991 | 21992 | 21993 | 21994 | 21995 | 22194 | 22195 | 22196 | 22200 | 22599 | 22673 | 22688 | 22775 | 22786 | 22898 | 23048 | 23202 | 24130 | 24503 | 26060 | 26530 | 26858 | 27382 | 27383 | 27384 | 29510 |  |
| ref | T | G | C | C | A | G | C | T | C | G | T | T | G | C | A | C | C | C | G | G | T | T | T | G | G | T | T | T | T | A | A | C | T | C | C | C | C | A | C | C | T | A | C | C | C | C | C | T | G | C | T | A | C | A | T | G | C | G | T | G | T | T | T | A | T | T | A | T | T | T | G | T | A | G | A | G | G | C | C | C | C | A | C | G | A | T | A |  |
| BA.2 | G | G | C | T | A | A | T | T | T | G | T | T | G | T | G | T | T | T | A | G | T | T | T | G | - | - | - | - | - | A | A | T | T | C | T | T | T | G | C | T | - | - | - | - | - | - | - | - | - | C | T | A | C | A | T | G | C | A | T | G | T | T | T | A | T | T | A | T | T | G | G | T | G | A | C | G | G | C | C | C | T | A | T | C | T | C | C |  |
| BA.1 | G | G | C | T | A | A | T | T | T | G | T | T | G | C | A | C | C | C | G | - | - | - | - | - | G | T | T | T | T | G | G | C | C | T | C | C | C | A | C | C | T | A | C | C | C | C | C | T | G | T | - | - | - | - | - | - | T | - | - | - | - | - | - | - | - | - | - | - | - | T | A | C | A | G | A | A | A | A | A | T | C | G | C | G | A | T | A | #1172823 |
| BA.1.1 | T | A | T | C | G | G | C | G | C | - | - | - | A | C | A | C | C | C | G | - | - | - | - | - | G | T | T | T | T | G | G | C | C | T | C | C | C | A | T | C | T | A | C | C | C | C | C | T | G | T | - | - | - | - | - | - | T | - | - | - | - | - | - | - | - | - | - | - | - | T | A | C | A | G | A | A | A | A | A | T | C | G | C | G | A | T | A |  |
| mut |  |  |  |  |  |  |  |  |  |  |  |  |  |  |  |  |  |  |  |  |  |  |  |  |  |  |  |  |  |  |  |  |  |  |  |  |  |  | T>C |  |  |  |  |  |  |  |  |  |  |  |  |  |  |  |  |  |  |  |  |  |  |  |  |  |  |  |  |  |  |  |  |  |  |  |  |  |  |  |  |  |  |  |  |  |  |  |  |  |

|  |  |  |  |  |  |  |  |  |  |  |  |  |  |  |  |  |
| --- | --- | --- | --- | --- | --- | --- | --- | --- | --- | --- | --- | --- | --- | --- | --- | --- |
| pos | 517 | 4459 | 6627 | 7926 | 8634 | 21588 | 21595 | 21969 | 22194 | 22195 | 22196 | 22197 | 22198 | 22202 | 22204 |  |
| ref | T | A | C | C | T | C | C | G | A | T | T | T | A | C | T |  |
| BA.1.1 | C | G | T | C | C | C | T | G | - | - | - | T | A | C | T |  |
| BA.1.1 | C | G | T | C | C | T | T | T | A | G | T | G | C | A | C | #1174700 |
| BA.1.1.18 | T | A | C | T | T | T | C | G | A | G | T | G | C | A | C |  |
| mut |  |  |  |  |  |  | C>T | G>T |  |  |  |  |  |  |  |  |

|  |  |  |  |  |  |  |  |  |  |  |  |  |  |  |  |  |  |  |  |  |  |  |  |  |  |  |  |  |  |  |  |  |  |  |  |  |  |  |  |  |  |  |  |  |  |  |  |  |  |  |  |  |  |  |  |  |  |  |  |  |  |  |  |  |  |  |  |  |  |  |  |  |  |  |  |  |  |  |  |  |
| --- | --- | --- | --- | --- | --- | --- | --- | --- | --- | --- | --- | --- | --- | --- | --- | --- | --- | --- | --- | --- | --- | --- | --- | --- | --- | --- | --- | --- | --- | --- | --- | --- | --- | --- | --- | --- | --- | --- | --- | --- | --- | --- | --- | --- | --- | --- | --- | --- | --- | --- | --- | --- | --- | --- | --- | --- | --- | --- | --- | --- | --- | --- | --- | --- | --- | --- | --- | --- | --- | --- | --- | --- | --- | --- | --- | --- | --- | --- | --- | --- |
| pos | 670 | 2470 | 2790 | 2832 | 4184 | 4321 | 5055 | 7969 | 10198 | 10447 | 11283 | 11284 | 11285 | 11286 | 11287 | 11292 | 11293 | 11294 | 11295 | 11296 | 11537 | 12880 | 13195 | 15240 | 15714 | 17410 | 17762 | 19955 | 20055 | 21595 | 21618 | 21633 | 21634 | 21635 | 21636 | 21637 | 21638 | 21639 | 21640 | 21641 | 21762 | 21765 | 21766 | 21767 | 21768 | 21769 | 21770 | 21846 | 21987 | 21988 | 21989 | 21990 | 21991 | 21992 | 21993 | 21994 | 21995 | 22194 | 22195 | 22196 | 22200 | 22599 | 22673 | 22688 | 22775 | 22786 | 22792 | 22898 | 23048 | 23202 | 24130 | 24503 | 26060 | 26530 | 26858 | 27382 | 27383 | 27384 | 29510 |  |
| ref | T | C | C | A | G | C | C | T | C | G | G | T | T | T | G | G | T | T | T | T | A | C | T | C | C | C | C | C | A | C | C | T | A | C | C | C | C | C | T | G | C | T | A | C | A | T | G | C | G | T | G | T | T | T | A | T | T | A | T | T | T | G | T | A | G | A | C | G | G | C | C | C | C | A | C | G | A | T | A |  |
| BA.1.1 | T | T | C | G | G | C | C | T | T | G | - | - | - | - | - | G | T | T | T | T | G | C | C | T | C | C | T | C | A | T | C | T | A | C | C | C | C | C | T | G | T | - | - | - | - | - | - | T | - | - | - | - | - | - | - | - | - | - | - | - | T | A | C | A | G | A | C | A | A | A | A | T | C | G | C | G | A | T | A |  |
| BA.2 | T | T | C | G | G | C | C | T | T | A | G | T | T | T | G | - | - | - | - | - | A | T | T | C | T | T | C | T | G | C | T | - | - | - | - | - | - | - | - | - | C | T | A | C | A | T | G | C | A | T | G | T | T | T | A | T | T | A | T | T | G | G | T | G | A | C | C | G | G | C | C | C | T | A | T | C | T | C | C | #1177107 |
| BA.2 | G | C | T | A | A | T | T | A | C | A | G | T | T | T | G | - | - | - | - | - | A | T | T | C | T | T | C | T | G | C | T | - | - | - | - | - | - | - | - | - | C | T | A | C | A | T | G | C | A | T | G | T | T | T | A | T | T | A | T | T | G | G | T | G | A | C | T | G | G | C | C | C | T | A | T | C | T | C | C |  |
| mut |  |  |  |  |  |  |  |  |  |  |  |  |  |  |  |  |  |  |  |  |  |  |  |  |  |  |  |  |  |  |  |  |  |  |  |  |  |  |  |  |  |  |  |  |  |  |  |  |  |  |  |  |  |  |  |  |  |  |  |  |  |  |  |  |  |  | T>C |  |  |  |  |  |  |  |  |  |  |  |  |  |

|  |  |  |  |  |  |  |  |  |  |  |  |  |  |  |  |  |  |
| --- | --- | --- | --- | --- | --- | --- | --- | --- | --- | --- | --- | --- | --- | --- | --- | --- | --- |
| pos | 1623 | 2158 | 2470 | 5806 | 7240 | 11563 | 21595 | 22186 | 22194 | 22195 | 22196 | 22197 | 22198 | 22202 | 22204 | 22599 |  |
| ref | T | T | C | C | A | C | C | G | A | T | T | T | A | C | T | G |  |
| BA.1.1 | C | C | T | T | T | T | T | T | - | - | - | T | A | C | T | A |  |
| BA.1.1 | C | C | T | T | T | T | T | T | A | G | T | G | C | A | C | A | #1177123 |
| BA.1 | T | T | C | C | A | C | C | G | A | G | T | G | C | A | C | G |  |
| mut |  |  |  |  |  |  |  |  |  |  |  |  |  |  |  | G>A |  |

|  |  |  |  |  |  |  |  |  |  |  |  |  |  |  |  |  |  |
| --- | --- | --- | --- | --- | --- | --- | --- | --- | --- | --- | --- | --- | --- | --- | --- | --- | --- |
| pos | 421 | 2470 | 4297 | 7835 | 8621 | 21595 | 22194 | 22195 | 22196 | 22197 | 22198 | 22202 | 22204 | 22599 | 27389 | 27511 |  |
| ref | C | C | A | T | A | C | A | T | T | T | A | C | T | G | C | T |  |
| BA.1.1 | T | T | T | C | G | T | - | - | - | T | A | C | T | A | C | C |  |
| BA.1.1 | T | T | T | C | G | T | A | G | T | G | C | A | C | A | T | T | #1179824 |
| BA.1 | C | C | A | T | A | C | A | G | T | G | C | A | C | G | C | T |  |
| mut |  |  |  |  |  |  |  |  |  |  |  |  |  | G>A | C>T |  |  |

|  |  |  |  |  |  |  |  |  |  |  |  |  |  |  |  |  |  |  |  |  |  |  |  |  |  |  |  |  |
| --- | --- | --- | --- | --- | --- | --- | --- | --- | --- | --- | --- | --- | --- | --- | --- | --- | --- | --- | --- | --- | --- | --- | --- | --- | --- | --- | --- | --- |
| pos | 1567 | 7720 | 8302 | 13247 | 18938 | 20134 | 21595 | 21736 | 22194 | 22195 | 22196 | 22197 | 22198 | 22202 | 22204 | 25317 | 27762 | 29541 | 29543 | 29555 | 29614 | 29700 | 29762 | 29769 | 29774 | 29779 | 29781 |  |
| ref | C | C | A | C | G | G | C | C | A | T | T | T | A | C | T | C | G | C | G | C | C | A | C | C | C | G | G |  |
| BA.1.1 | T | C | G | T | A | G | T | C | - | - | - | T | A | C | T | T | C | - | - | - | - | - | - | - | - | - | - |  |
| BA.1.1 | T | T | G | T | A | T | T | T | A | G | T | G | C | A | C | T | G | C | G | C | C | A | C | C | C | G | G | #1179833 |
| BA.1.1 | C | C | A | C | G | T | C | T | A | G | T | G | C | A | C | C | G | C | G | C | C | A | C | C | C | G | G |  |
| mut |  | C>T |  |  |  |  | C>T |  |  |  |  |  |  |  |  | C>T |  |  |  |  |  |  |  |  |  |  |  |  |

|  |  |  |  |  |  |  |  |  |  |  |  |  |  |  |  |  |  |  |  |  |  |  |  |
| --- | --- | --- | --- | --- | --- | --- | --- | --- | --- | --- | --- | --- | --- | --- | --- | --- | --- | --- | --- | --- | --- | --- | --- |
| pos | 7926 | 8302 | 10537 | 19896 | 22194 | 22195 | 22196 | 22197 | 22198 | 22202 | 22204 | 27321 | 29541 | 29543 | 29555 | 29614 | 29700 | 29762 | 29769 | 29774 | 29779 | 29781 |  |
| ref | C | A | C | T | A | T | T | T | A | C | T | A | C | G | C | C | A | C | C | C | G | G |  |
| BA.1.1 | T | G | T | C | - | - | - | T | A | C | T | T | - | - | - | - | - | - | - | - | - | - |  |
| BA.1.1 | T | G | T | C | A | G | T | G | C | A | C | T | C | G | C | C | A | C | C | C | G | G | #1179839 |
| BA.1.1 | C | A | C | T | A | G | T | G | C | A | C | A | C | G | C | C | A | C | C | C | G | G |  |
| mut |  |  |  |  |  |  |  |  |  |  |  | A>T |  |  |  |  |  |  |  |  |  |  |  |

|  |  |  |  |  |  |  |  |  |  |  |  |  |  |  |  |  |  |  |  |
| --- | --- | --- | --- | --- | --- | --- | --- | --- | --- | --- | --- | --- | --- | --- | --- | --- | --- | --- | --- |
| pos | 875 | 1191 | 5703 | 5704 | 5705 | 5869 | 10135 | 20016 | 22194 | 22195 | 22196 | 22197 | 22198 | 22202 | 22204 | 23595 | 25708 | 29301 |  |
| ref | C | C | A | G | T | C | T | C | A | T | T | T | A | C | T | C | C | A |  |
| BA.1.1 | C | C | T | A | A | C | T | C | - | - | - | T | A | C | T | C | C | A |  |
| BA.1.15 | C | T | T | A | A | T | C | T | A | G | T | G | C | A | C | T | T | G | #1183324 |
| BA.1.15 | T | C | A | G | T | C | C | C | A | G | T | G | C | A | C | T | T | G |  |
| mut |  | C>T |  |  |  | C>T |  | C>T |  |  |  |  |  |  |  |  |  |  |  |

|  |  |  |  |  |  |  |  |  |  |  |  |  |  |  |  |  |  |
| --- | --- | --- | --- | --- | --- | --- | --- | --- | --- | --- | --- | --- | --- | --- | --- | --- | --- |
| pos | 186 | 2189 | 3510 | 6422 | 6532 | 6719 | 14184 | 17019 | 22194 | 22195 | 22196 | 22197 | 22198 | 22202 | 22204 | 28291 |  |
| ref | C | C | C | G | G | A | C | G | A | T | T | T | A | C | T | C |  |
| BA.1.1 | C | C | T | A | T | A | T | T | - | - | - | T | A | C | T | C |  |
| BA.1.1 | T | T | T | A | T | G | T | T | A | G | T | G | C | A | C | T | #1183328 |
| BA.1.1 | C | C | C | G | G | A | C | G | A | G | T | G | C | A | C | T |  |
| mut | C>T | C>T |  |  |  | A>G |  |  |  |  |  |  |  |  |  |  |  |

|  |  |  |  |  |  |  |  |  |  |  |  |  |  |  |  |  |  |  |
| --- | --- | --- | --- | --- | --- | --- | --- | --- | --- | --- | --- | --- | --- | --- | --- | --- | --- | --- |
| pos | 277 | 6568 | 7926 | 9042 | 11919 | 15337 | 16395 | 20010 | 21595 | 22194 | 22195 | 22196 | 22197 | 22198 | 22202 | 22204 | 29614 |  |
| ref | T | C | C | C | C | A | A | A | C | A | T | T | T | A | C | T | C |  |
| BA.1.1 | T | T | C | T | T | G | T | G | T | - | - | - | T | A | C | T | T |  |
| BA.1.1 | A | T | C | T | T | G | T | G | T | A | G | T | G | C | A | C | T | #1183585 |
| BA.1.1.18 | T | C | T | C | C | A | A | A | C | A | G | T | G | C | A | C | C |  |
| mut | T>A |  |  |  |  |  |  |  |  |  |  |  |  |  |  |  | C>T |  |

|  |  |  |  |  |  |  |  |  |  |  |  |  |  |  |  |  |  |
| --- | --- | --- | --- | --- | --- | --- | --- | --- | --- | --- | --- | --- | --- | --- | --- | --- | --- |
| pos | 843 | 1392 | 1599 | 6525 | 18439 | 21137 | 21141 | 21595 | 22194 | 22195 | 22196 | 22197 | 22198 | 22202 | 22204 | 27384 |  |
| ref | C | C | G | C | T | A | A | C | A | T | T | T | A | C | T | T |  |
| BA.1.1 | T | T | G | T | A | G | G | T | A | G | T | G | C | A | C | C |  |
| BA.1.1 | T | T | G | T | A | G | G | T | - | - | - | T | A | C | T | C | #1186872 |
| BA.1.1 | C | C | T | C | T | A | A | C | - | - | - | T | A | C | T | T |  |
| mut |  |  |  |  |  |  |  |  |  |  |  |  |  |  |  | T>C |  |

|  |  |  |  |  |  |  |  |  |  |  |  |  |  |  |  |  |  |  |  |  |  |  |  |  |  |  |  |  |  |  |  |  |  |  |  |  |  |  |  |  |  |  |  |  |  |  |  |  |  |  |  |  |  |  |  |  |  |  |  |  |  |  |  |  |  |  |  |  |  |  |  |  |  |  |  |  |  |  |  |  |  |  |  |  |  |  |  |  |  |  |  |  |  |
| --- | --- | --- | --- | --- | --- | --- | --- | --- | --- | --- | --- | --- | --- | --- | --- | --- | --- | --- | --- | --- | --- | --- | --- | --- | --- | --- | --- | --- | --- | --- | --- | --- | --- | --- | --- | --- | --- | --- | --- | --- | --- | --- | --- | --- | --- | --- | --- | --- | --- | --- | --- | --- | --- | --- | --- | --- | --- | --- | --- | --- | --- | --- | --- | --- | --- | --- | --- | --- | --- | --- | --- | --- | --- | --- | --- | --- | --- | --- | --- | --- | --- | --- | --- | --- | --- | --- | --- | --- | --- | --- | --- | --- | --- |
| pos | 670 | 1585 | 2470 | 2790 | 2832 | 2910 | 4184 | 4321 | 5386 | 6513 | 6514 | 6515 | 8149 | 8393 | 9344 | 9424 | 9534 | 9866 | 10198 | 10447 | 11049 | 11075 | 11283 | 11284 | 11285 | 11286 | 11287 | 11292 | 11293 | 11294 | 11295 | 11296 | 11537 | 12085 | 12880 | 13195 | 15240 | 15714 | 17410 | 19955 | 20055 | 21618 | 21633 | 21634 | 21635 | 21636 | 21637 | 21638 | 21639 | 21640 | 21641 | 21762 | 21765 | 21766 | 21767 | 21768 | 21769 | 21770 | 21846 | 21987 | 21988 | 21989 | 21990 | 21991 | 21992 | 21993 | 21994 | 21995 | 22194 | 22195 | 22196 | 22200 | 22351 | 22599 | 22673 | 22688 | 22775 | 22786 | 22792 | 22898 | 23048 | 23202 | 24130 | 24503 | 26058 | 26060 | 26530 | 26858 | 27382 | 27383 | 27384 | 29510 |  |
| ref | T | A | C | C | A | C | G | C | T | G | T | T | T | G | C | A | C | C | C | G | T | T | G | T | T | T | G | G | T | T | T | T | A | C | C | T | C | C | C | C | A | C | T | A | C | C | C | C | C | T | G | C | T | A | C | A | T | G | C | G | T | G | T | T | T | A | T | T | A | T | T | T | A | G | T | A | G | A | C | G | G | C | C | C | C | C | A | C | G | A | T | A |  |
| BA.1.1 | T | A | T | C | G | C | G | C | G | - | - | - | T | A | C | A | C | C | C | G | C | T | - | - | - | - | - | G | T | T | T | T | G | C | C | C | T | C | C | C | A | C | T | A | C | C | C | C | C | T | G | T | - | - | - | - | - | - | T | - | - | - | - | - | - | - | - | - | - | - | - | T | A | A | C | A | G | A | C | A | A | A | A | T | C | C | G | C | G | A | T | A |  |
| XY | T | G | T | C | G | C | G | C | G | - | - | - | T | A | C | A | C | C | C | G | C | T | - | - | - | - | - | G | T | T | T | T | G | T | T | T | C | T | T | T | G | T | - | - | - | - | - | - | - | - | - | C | T | A | C | A | T | G | C | A | T | G | T | T | T | A | T | T | A | T | T | G | G | G | T | G | A | C | T | G | G | C | C | C | T | T | A | T | C | T | C | C | #1187989 |
| BA.2 | G | A | C | T | A | T | A | T | T | G | T | T | C | G | T | G | T | T | T | A | T | C | G | T | T | T | G | - | - | - | - | - | A | C | T | T | C | T | T | T | G | T | - | - | - | - | - | - | - | - | - | C | T | A | C | A | T | G | C | A | T | G | T | T | T | A | T | T | A | T | T | G | G | G | T | G | A | C | T | G | G | C | C | C | T | T | A | T | C | T | C | C |  |
| mut |  | A>G |  |  |  |  |  |  |  |  |  |  |  |  |  |  |  |  |  |  |  |  |  |  |  |  |  |  |  |  |  |  |  | C>T |  |  |  |  |  |  |  |  |  |  |  |  |  |  |  |  |  |  |  |  |  |  |  |  |  |  |  |  |  |  |  |  |  |  |  |  |  |  |  |  |  |  |  |  |  |  |  |  |  |  |  |  |  |  |  |  |  |  |  |

|  |  |  |  |  |  |  |  |  |  |  |  |  |  |  |  |  |  |  |  |  |  |  |  |  |  |  |  |  |  |  |  |  |  |
| --- | --- | --- | --- | --- | --- | --- | --- | --- | --- | --- | --- | --- | --- | --- | --- | --- | --- | --- | --- | --- | --- | --- | --- | --- | --- | --- | --- | --- | --- | --- | --- | --- | --- |
| pos | 686 | 687 | 688 | 689 | 690 | 691 | 692 | 693 | 694 | 9866 | 12160 | 14277 | 16342 | 21765 | 21766 | 21767 | 21768 | 21769 | 21770 | 22917 | 23018 | 23040 | 26529 | 26858 | 27259 | 27382 | 27383 | 27384 | 27788 | 27889 | 28724 | 29754 |  |
| ref | A | A | G | T | C | A | T | T | T | C | G | G | T | T | A | C | A | T | G | T | T | A | G | C | A | G | A | T | G | C | C | C |  |
| BA.4 | - | - | - | - | - | - | - | - | - | C | A | G | T | - | - | - | - | - | - | G | G | A | G | T | C | C | T | C | T | C | T | C |  |
| BA.5 | A | A | G | T | C | A | T | T | T | C | A | G | T | - | - | - | - | - | - | G | G | A | A | C | A | G | A | T | G | T | C | T | #1189192 |
| BA.2 | A | A | G | T | C | A | T | T | T | T | G | T | C | T | A | C | A | T | G | T | T | G | G | C | C | G | A | T | G | C | C | T |  |
| mut | ->A | ->A | ->G | ->T | ->C | ->A | ->T | ->T | ->T |  |  |  |  |  |  |  |  |  |  |  |  |  | G>A |  | C>A |  |  |  |  | C>T |  |  |  |

|  |  |  |  |  |  |  |  |  |  |  |  |  |  |  |  |  |  |  |  |  |  |  |  |  |  |  |  |  |  |  |  |  |  |  |  |  |  |  |  |  |  |  |  |  |  |  |  |  |  |  |  |  |  |  |  |  |  |  |  |  |  |  |  |  |  |  |  |  |  |  |  |  |  |  |  |  |  |  |  |  |  |  |  |  |  |  |  |  |  |  |  |  |  |  |
| --- | --- | --- | --- | --- | --- | --- | --- | --- | --- | --- | --- | --- | --- | --- | --- | --- | --- | --- | --- | --- | --- | --- | --- | --- | --- | --- | --- | --- | --- | --- | --- | --- | --- | --- | --- | --- | --- | --- | --- | --- | --- | --- | --- | --- | --- | --- | --- | --- | --- | --- | --- | --- | --- | --- | --- | --- | --- | --- | --- | --- | --- | --- | --- | --- | --- | --- | --- | --- | --- | --- | --- | --- | --- | --- | --- | --- | --- | --- | --- | --- | --- | --- | --- | --- | --- | --- | --- | --- | --- | --- | --- | --- | --- | --- |
| pos | 670 | 2470 | 2790 | 2832 | 4184 | 4321 | 5386 | 6513 | 6514 | 6515 | 6812 | 8393 | 8905 | 9344 | 9424 | 9534 | 9803 | 9866 | 10198 | 10447 | 11283 | 11284 | 11285 | 11286 | 11287 | 11292 | 11293 | 11294 | 11295 | 11296 | 11537 | 12880 | 13195 | 15240 | 15714 | 17410 | 17460 | 19955 | 20055 | 21618 | 21633 | 21634 | 21635 | 21636 | 21637 | 21638 | 21639 | 21640 | 21641 | 21762 | 21765 | 21766 | 21767 | 21768 | 21769 | 21770 | 21846 | 21987 | 21988 | 21989 | 21990 | 21991 | 21992 | 21993 | 21994 | 21995 | 22000 | 22194 | 22195 | 22196 | 22200 | 22599 | 22673 | 22688 | 22775 | 22786 | 22792 | 22898 | 23048 | 23202 | 24130 | 24503 | 26060 | 26522 | 26530 | 26622 | 26858 | 27382 | 27383 | 27384 | 27970 | 28015 | 29510 |  |
| ref | T | C | C | A | G | C | T | G | T | T | A | G | C | C | A | C | C | C | C | G | G | T | T | T | G | G | T | T | T | T | A | C | T | C | C | C | A | C | A | C | T | A | C | C | C | C | C | T | G | C | T | A | C | A | T | G | C | G | T | G | T | T | T | A | T | T | C | A | T | T | T | G | T | A | G | A | C | G | G | C | C | C | C | C | A | C | C | G | A | T | C | T | A |  |
| BA.1.1 | T | T | C | G | G | C | G | - | - | - | G | A | C | C | A | C | T | C | C | G | - | - | - | - | - | G | T | T | T | T | G | C | C | T | C | C | A | C | A | C | T | A | C | C | C | C | C | T | T | T | - | - | - | - | - | - | T | - | - | - | - | - | - | - | - | - | T | - | - | - | T | A | C | A | G | A | C | A | A | A | A | T | C | T | G | T | C | G | A | T | T | T | A |  |
| BA.2 | T | T | C | G | G | C | G | - | - | - | G | A | C | C | A | C | T | C | C | G | - | - | - | - | - | G | T | T | T | T | A | T | T | C | T | T | G | T | G | T | - | - | - | - | - | - | - | - | - | C | T | A | C | A | T | G | C | A | T | G | T | T | T | A | T | T | C | A | T | T | G | G | T | G | A | C | T | G | G | C | C | C | T | C | A | C | T | C | T | C | C | C | C | #1189559 |
| BA.2 | G | C | T | A | A | T | T | G | T | T | A | G | T | T | G | T | C | T | T | A | G | T | T | T | G | - | - | - | - | - | A | T | T | C | T | T | G | T | G | T | - | - | - | - | - | - | - | - | - | C | T | A | C | A | T | G | C | A | T | G | T | T | T | A | T | T | C | A | T | T | G | G | T | G | A | C | T | G | G | C | C | C | T | C | A | C | T | C | T | C | C | C | C |  |
| mut |  |  |  |  |  |  |  |  |  |  |  |  |  |  |  |  |  |  |  |  |  |  |  |  |  |  |  |  |  |  |  |  |  |  |  |  |  |  |  |  |  |  |  |  |  |  |  |  |  |  |  |  |  |  |  |  |  |  |  |  |  |  |  |  |  |  |  |  |  |  |  |  |  |  |  |  |  |  |  |  |  |  |  |  |  |  |  |  |  |  |  |  |  |  |

|  |  |  |  |  |  |  |  |  |  |  |  |  |  |  |  |  |  |
| --- | --- | --- | --- | --- | --- | --- | --- | --- | --- | --- | --- | --- | --- | --- | --- | --- | --- |
| pos | 379 | 2470 | 3183 | 6666 | 10317 | 21588 | 21595 | 22194 | 22195 | 22196 | 22197 | 22198 | 22202 | 22204 | 22599 | 29781 |  |
| ref | C | C | A | C | A | C | C | A | T | T | T | A | C | T | G | G |  |
| BA.1.1 | T | T | G | T | G | T | T | - | - | - | T | A | C | T | A | - |  |
| BA.1.1 | T | T | G | T | G | T | T | A | G | T | G | C | A | C | A | G | #1191507 |
| BA.1 | C | C | A | C | A | C | C | A | G | T | G | C | A | C | G | G |  |
| mut |  |  |  |  |  |  |  |  |  |  |  |  |  |  | G>A |  |  |

|  |  |  |  |  |  |  |  |  |  |  |  |  |  |  |  |  |  |  |  |
| --- | --- | --- | --- | --- | --- | --- | --- | --- | --- | --- | --- | --- | --- | --- | --- | --- | --- | --- | --- |
| pos | 2272 | 4301 | 7926 | 7987 | 9093 | 9935 | 10186 | 13033 | 15237 | 21595 | 22194 | 22195 | 22196 | 22197 | 22198 | 22202 | 22204 | 25521 |  |
| ref | G | C | C | G | T | A | C | T | C | C | A | T | T | T | A | C | T | C |  |
| BA.1.1 | T | C | C | G | C | A | T | C | C | T | - | - | - | T | A | C | T | T |  |
| BA.1.1 | T | T | C | A | C | A | T | C | T | T | A | G | T | G | C | A | C | C | #1191658 |
| BA.1.1.18 | G | C | T | G | T | G | C | T | T | C | A | G | T | G | C | A | C | C |  |
| mut |  | C>T |  | G>A |  |  |  |  |  | C>T |  |  |  |  |  |  |  |  |  |

|  |  |  |  |  |  |  |  |  |  |  |  |  |  |  |  |
| --- | --- | --- | --- | --- | --- | --- | --- | --- | --- | --- | --- | --- | --- | --- | --- |
| pos | 2470 | 3426 | 4296 | 6647 | 12786 | 16064 | 21595 | 22194 | 22195 | 22196 | 22197 | 22198 | 22202 | 22204 |  |
| ref | C | C | C | G | C | A | C | A | T | T | T | A | C | T |  |
| BA.1.1.1 | C | T | T | A | C | G | C | - | - | - | T | A | C | T |  |
| BA.1.1.1 | C | T | T | A | T | G | C | A | G | T | G | C | A | C | #1194353 |
| BA.1.1 | T | C | C | G | C | A | T | A | G | T | G | C | A | C |  |
| mut |  |  |  |  | C>T |  |  |  |  |  |  |  |  |  |  |

|  |  |  |  |  |  |  |  |  |  |  |  |  |  |  |  |  |  |  |  |  |  |  |  |  |  |  |  |  |  |  |  |  |  |  |  |  |  |  |  |  |  |  |  |  |  |  |  |  |  |  |  |  |  |  |  |  |  |  |  |  |  |  |  |  |  |  |  |  |  |  |  |  |  |  |  |  |  |  |  |  |  |  |  |  |  |  |  |
| --- | --- | --- | --- | --- | --- | --- | --- | --- | --- | --- | --- | --- | --- | --- | --- | --- | --- | --- | --- | --- | --- | --- | --- | --- | --- | --- | --- | --- | --- | --- | --- | --- | --- | --- | --- | --- | --- | --- | --- | --- | --- | --- | --- | --- | --- | --- | --- | --- | --- | --- | --- | --- | --- | --- | --- | --- | --- | --- | --- | --- | --- | --- | --- | --- | --- | --- | --- | --- | --- | --- | --- | --- | --- | --- | --- | --- | --- | --- | --- | --- | --- | --- | --- | --- | --- | --- | --- |
| pos | 670 | 2790 | 2832 | 4084 | 4184 | 4321 | 5386 | 5515 | 6513 | 6514 | 6515 | 8393 | 9344 | 9424 | 9534 | 9866 | 10198 | 10263 | 10447 | 11283 | 11284 | 11285 | 11286 | 11287 | 11292 | 11293 | 11294 | 11295 | 11296 | 11537 | 12880 | 13195 | 13653 | 15240 | 15714 | 17410 | 19955 | 20055 | 21618 | 21633 | 21634 | 21635 | 21636 | 21637 | 21638 | 21639 | 21640 | 21641 | 21762 | 21765 | 21766 | 21767 | 21768 | 21769 | 21770 | 21846 | 21987 | 21988 | 21989 | 21990 | 21991 | 21992 | 21993 | 21994 | 21995 | 22194 | 22195 | 22196 | 22200 | 22673 | 22688 | 22775 | 22786 | 22898 | 23048 | 23202 | 23485 | 24130 | 24503 | 26060 | 26530 | 26858 | 27382 | 27383 | 28498 | 29510 |  |
| ref | T | C | A | C | G | C | T | G | G | T | T | G | C | A | C | C | C | C | G | G | T | T | T | G | G | T | T | T | T | A | C | T | A | C | C | C | C | A | C | T | A | C | C | C | C | C | T | G | C | T | A | C | A | T | G | C | G | T | G | T | T | T | A | T | T | A | T | T | T | T | A | G | A | G | G | C | T | C | C | C | A | C | G | A | C | A |  |
| BA.1 | T | C | G | C | G | C | G | T | - | - | - | A | C | A | C | C | C | T | G | - | - | - | - | - | G | T | T | T | T | G | C | C | G | T | C | C | C | A | C | T | A | C | C | C | C | C | T | G | T | - | - | - | - | - | - | T | - | - | - | - | - | - | - | - | - | - | - | - | T | C | A | G | A | A | A | A | T | A | T | C | G | C | G | A | C | A |  |
| BA.2 | T | C | G | C | G | C | G | T | - | - | - | A | C | A | C | C | C | T | G | - | - | - | - | - | G | T | T | T | T | G | C | C | G | T | C | C | T | G | T | - | - | - | - | - | - | - | - | - | C | T | A | C | A | T | G | C | A | T | G | T | T | T | A | T | T | A | T | T | G | T | G | A | C | G | G | C | C | C | C | T | A | T | C | T | T | C | #1196233 |
| BA.2 | G | T | A | T | A | T | T | G | G | T | T | G | T | G | T | T | T | C | A | G | T | T | T | G | - | - | - | - | - | A | T | T | A | C | T | T | T | G | T | - | - | - | - | - | - | - | - | - | C | T | A | C | A | T | G | C | A | T | G | T | T | T | A | T | T | A | T | T | G | T | G | A | C | G | G | C | C | C | C | T | A | T | C | T | T | C |  |
| mut |  |  |  |  |  |  |  |  |  |  |  |  |  |  |  |  |  |  |  |  |  |  |  |  |  |  |  |  |  |  |  |  |  |  |  |  |  |  |  |  |  |  |  |  |  |  |  |  |  |  |  |  |  |  |  |  |  |  |  |  |  |  |  |  |  |  |  |  |  |  |  |  |  |  |  |  |  |  |  |  |  |  |  |  |  |  |  |

|  |  |  |  |  |  |  |  |  |  |  |  |  |  |  |  |  |  |  |  |  |  |  |  |  |  |  |  |  |  |  |  |  |  |  |  |  |  |  |  |  |  |  |  |  |  |  |  |  |  |  |  |  |  |  |  |  |  |  |  |  |  |  |  |  |  |  |  |  |  |  |  |  |  |  |  |  |  |  |  |  |  |  |  |  |  |  |  |  |
| --- | --- | --- | --- | --- | --- | --- | --- | --- | --- | --- | --- | --- | --- | --- | --- | --- | --- | --- | --- | --- | --- | --- | --- | --- | --- | --- | --- | --- | --- | --- | --- | --- | --- | --- | --- | --- | --- | --- | --- | --- | --- | --- | --- | --- | --- | --- | --- | --- | --- | --- | --- | --- | --- | --- | --- | --- | --- | --- | --- | --- | --- | --- | --- | --- | --- | --- | --- | --- | --- | --- | --- | --- | --- | --- | --- | --- | --- | --- | --- | --- | --- | --- | --- | --- | --- | --- | --- | --- |
| pos | 670 | 2094 | 2695 | 2790 | 2832 | 4184 | 4321 | 5386 | 6513 | 6514 | 6515 | 8393 | 9344 | 9424 | 9534 | 9866 | 10000 | 10198 | 10447 | 11283 | 11284 | 11285 | 11286 | 11287 | 11292 | 11293 | 11294 | 11295 | 11296 | 11537 | 12880 | 13195 | 15240 | 15714 | 17410 | 19955 | 20055 | 21618 | 21633 | 21634 | 21635 | 21636 | 21637 | 21638 | 21639 | 21640 | 21641 | 21762 | 21765 | 21766 | 21767 | 21768 | 21769 | 21770 | 21846 | 21987 | 21988 | 21989 | 21990 | 21991 | 21992 | 21993 | 21994 | 21995 | 22194 | 22195 | 22196 | 22200 | 22673 | 22688 | 22775 | 22786 | 22792 | 22898 | 23048 | 23202 | 24130 | 24503 | 26060 | 26530 | 26571 | 26858 | 27382 | 27383 | 27384 | 29510 | 29632 |  |
| ref | T | C | C | C | A | G | C | T | G | T | T | G | C | A | C | C | T | C | G | G | T | T | T | G | G | T | T | T | T | A | C | T | C | C | C | C | A | C | T | A | C | C | C | C | C | T | G | C | T | A | C | A | T | G | C | G | T | G | T | T | T | A | T | T | A | T | T | T | T | A | G | A | C | G | G | C | C | C | C | A | C | C | G | A | T | A | C |  |
| BA.1 | T | T | C | C | G | G | C | G | - | - | - | A | C | A | C | C | T | C | G | - | - | - | - | - | G | T | T | T | T | G | C | C | T | C | C | C | A | C | T | A | C | C | C | C | C | T | G | T | - | - | - | - | - | - | T | - | - | - | - | - | - | - | - | - | - | - | - | T | C | A | G | A | C | A | A | A | A | T | C | G | C | C | G | A | T | A | T |  |
| BA.2 | T | T | T | C | G | G | C | G | - | - | - | A | C | A | C | T | C | T | A | G | T | T | T | G | - | - | - | - | - | A | T | T | C | T | T | T | G | T | - | - | - | - | - | - | - | - | - | C | T | A | C | A | T | G | C | A | T | G | T | T | T | A | T | T | A | T | T | G | T | G | A | C | T | G | G | C | C | C | T | A | T | T | C | T | C | C | C | #1197273 |
| BA.2 | G | C | C | T | A | A | T | T | G | T | T | G | T | G | T | T | C | T | A | G | T | T | T | G | - | - | - | - | - | A | T | T | C | T | T | T | G | T | - | - | - | - | - | - | - | - | - | C | T | A | C | A | T | G | C | A | T | G | T | T | T | A | T | T | A | T | T | G | T | G | A | C | C | G | G | C | C | C | T | A | C | T | C | T | C | C | C |  |
| mut |  |  | C>T |  |  |  |  |  |  |  |  |  |  |  |  |  |  |  |  |  |  |  |  |  |  |  |  |  |  |  |  |  |  |  |  |  |  |  |  |  |  |  |  |  |  |  |  |  |  |  |  |  |  |  |  |  |  |  |  |  |  |  |  |  |  |  |  |  |  |  |  |  | C>T |  |  |  |  |  |  |  | C>T |  |  |  |  |  |  |  |

|  |  |  |  |  |  |  |  |  |  |  |  |  |  |  |  |  |  |  |
| --- | --- | --- | --- | --- | --- | --- | --- | --- | --- | --- | --- | --- | --- | --- | --- | --- | --- | --- |
| pos | 44 | 105 | 335 | 2647 | 6097 | 11674 | 15009 | 21721 | 22917 | 23673 | 23948 | 24892 | 25416 | 28254 | 28877 | 28878 | 29769 |  |
| ref | C | G | C | A | A | C | T | C | T | C | G | T | C | A | A | G | C |  |
| BA.2 | C | T | C | A | G | C | T | C | T | C | T | C | C | A | A | G | T |  |
| BA.2.12.1 | C | T | T | G | G | C | T | T | A | T | G | T | T | - | T | C | C | #1197908 |
| BA.2.12.1 | T | G | C | A | A | T | C | T | A | T | T | T | T | A | A | G | C |  |
| mut |  |  | C>T | A>G |  |  |  |  |  |  | T>G |  |  | A>- | A>T | G>C |  |  |

|  |  |  |  |  |  |  |  |  |  |  |  |  |  |  |  |  |  |  |  |  |  |  |  |  |
| --- | --- | --- | --- | --- | --- | --- | --- | --- | --- | --- | --- | --- | --- | --- | --- | --- | --- | --- | --- | --- | --- | --- | --- | --- |
| pos | 44 | 670 | 2790 | 2832 | 4184 | 4321 | 5386 | 6127 | 6513 | 6514 | 6515 | 9693 | 11407 | 16064 | 19723 | 21417 | 21622 | 22599 | 25677 | 25792 | 29541 | 29632 | 29736 |  |
| ref | C | T | C | A | G | C | T | G | G | T | T | C | C | A | G | A | C | G | G | C | C | C | G |  |
| BA.1 | T | G | T | A | A | T | T | G | G | T | T | T | C | A | G | G | C | G | G | C | C | T | G |  |
| BA.1 | T | G | T | A | A | T | T | A | - | - | - | C | T | G | A | A | C | A | T | T | T | C | C | #1200360 |
| BA.1.1.1 | C | T | C | G | G | C | G | G | - | - | - | C | T | G | A | A | T | A | T | C | T | C | C |  |
| mut |  |  |  |  |  |  |  | G>A |  |  |  |  |  |  |  |  | T>C |  |  | C>T |  |  |  |  |

|  |  |  |  |  |  |  |  |  |  |  |  |  |  |  |  |  |  |  |  |
| --- | --- | --- | --- | --- | --- | --- | --- | --- | --- | --- | --- | --- | --- | --- | --- | --- | --- | --- | --- |
| pos | 246 | 2470 | 5467 | 5907 | 6160 | 10642 | 13335 | 16323 | 21595 | 22194 | 22195 | 22196 | 22197 | 22198 | 22202 | 22204 | 28877 | 28878 |  |
| ref | G | C | C | C | T | G | C | C | C | A | T | T | T | A | C | T | A | G |  |
| BA.1.1 | A | C | T | T | C | T | T | T | C | - | - | - | T | A | C | T | T | C |  |
| BA.1.1 | A | C | T | T | C | T | T | T | C | A | G | T | G | C | A | C | T | C | #1202399 |
| BA.1.1 | G | T | C | C | T | G | C | C | T | A | G | T | G | C | A | C | A | G |  |
| mut |  |  |  |  |  |  |  |  |  |  |  |  |  |  |  |  | A>T | G>C |  |

|  |  |  |  |  |  |  |  |  |  |  |  |  |  |  |  |  |  |
| --- | --- | --- | --- | --- | --- | --- | --- | --- | --- | --- | --- | --- | --- | --- | --- | --- | --- |
| pos | 44 | 949 | 13945 | 14845 | 20578 | 22792 | 25416 | 25511 | 25783 | 27476 | 29192 | 29700 | 29728 | 29757 | 29758 | 29759 |  |
| ref | C | A | C | C | G | C | C | C | T | C | T | A | T | G | T | G |  |
| BA.2 | T | G | C | T | T | C | T | T | T | C | C | G | T | G | T | G |  |
| BA.2 | T | G | C | T | T | C | T | T | C | T | C | G | C | A | C | C | #1203710 |
| BA.2 | C | A | T | C | G | T | C | C | T | C | T | A | C | A | C | C |  |
| mut |  |  |  |  |  |  |  |  | T>C | C>T |  |  |  |  |  |  |  |

|  |  |  |  |  |  |  |  |  |  |  |  |  |  |  |  |
| --- | --- | --- | --- | --- | --- | --- | --- | --- | --- | --- | --- | --- | --- | --- | --- |
| pos | 615 | 835 | 5447 | 6337 | 8017 | 17213 | 22194 | 22195 | 22196 | 22197 | 22198 | 22202 | 22204 | 29541 |  |
| ref | C | C | G | G | G | C | A | T | T | T | A | C | T | C |  |
| BA.1.1 | T | C | G | A | T | T | A | G | T | G | C | A | C | T |  |
| BA.1.1 | T | C | G | A | T | T | - | - | - | T | A | C | T | T | #1207909 |
| BA.1.1 | C | T | T | G | G | C | - | - | - | T | A | C | T | C |  |
| mut |  |  |  |  |  |  |  |  |  |  |  |  |  | C>T |  |

|  |  |  |  |  |  |  |  |  |  |  |  |  |  |  |  |  |
| --- | --- | --- | --- | --- | --- | --- | --- | --- | --- | --- | --- | --- | --- | --- | --- | --- |
| pos | 521 | 1371 | 3867 | 4965 | 8320 | 12897 | 20762 | 22120 | 22194 | 22195 | 22196 | 22197 | 22198 | 22202 | 22204 |  |
| ref | G | A | A | C | C | C | C | C | A | T | T | T | A | C | T |  |
| BA.1.15 | G | A | G | C | T | T | C | T | - | - | - | T | A | C | T |  |
| BA.1.15 | T | A | G | C | T | T | C | T | A | G | T | G | C | A | C | #1208033 |
| BA.1.15 | G | G | A | T | C | C | T | C | A | G | T | G | C | A | C |  |
| mut | G>T |  |  |  |  |  |  |  |  |  |  |  |  |  |  |  |

|  |  |  |  |  |  |  |  |  |  |  |  |  |  |
| --- | --- | --- | --- | --- | --- | --- | --- | --- | --- | --- | --- | --- | --- |
| pos | 162 | 2997 | 8647 | 19859 | 22194 | 22195 | 22196 | 22197 | 22198 | 22202 | 22204 | 25626 |  |
| ref | C | C | T | C | A | T | T | T | A | C | T | C |  |
| BA.1.1 | T | T | C | T | - | - | - | T | A | C | T | C |  |
| BA.1.1 | T | T | C | T | A | G | T | G | C | A | C | T | #1208815 |
| BA.1.1 | C | C | T | C | A | G | T | G | C | A | C | T |  |
| mut |  |  |  |  |  |  |  |  |  |  |  |  |  |

|  |  |  |  |  |  |  |  |  |  |  |  |  |  |  |  |  |  |  |  |  |  |  |  |  |  |  |  |  |  |  |  |  |  |  |  |  |  |  |  |  |  |  |  |  |  |  |  |  |  |  |  |  |  |  |  |  |  |  |  |  |  |  |  |  |  |  |  |  |  |  |  |  |  |  |  |  |  |  |  |  |  |  |  |  |  |  |  |  |  |  |  |  |  |  |  |  |
| --- | --- | --- | --- | --- | --- | --- | --- | --- | --- | --- | --- | --- | --- | --- | --- | --- | --- | --- | --- | --- | --- | --- | --- | --- | --- | --- | --- | --- | --- | --- | --- | --- | --- | --- | --- | --- | --- | --- | --- | --- | --- | --- | --- | --- | --- | --- | --- | --- | --- | --- | --- | --- | --- | --- | --- | --- | --- | --- | --- | --- | --- | --- | --- | --- | --- | --- | --- | --- | --- | --- | --- | --- | --- | --- | --- | --- | --- | --- | --- | --- | --- | --- | --- | --- | --- | --- | --- | --- | --- | --- | --- | --- | --- | --- | --- | --- |
| pos | 44 | 670 | 2470 | 2790 | 2832 | 4184 | 4321 | 5386 | 6513 | 6514 | 6515 | 7564 | 8393 | 9344 | 9424 | 9534 | 9866 | 10198 | 10447 | 11283 | 11284 | 11285 | 11286 | 11287 | 11292 | 11293 | 11294 | 11295 | 11296 | 11537 | 11674 | 12880 | 13195 | 15009 | 15240 | 15714 | 17410 | 18673 | 18674 | 19955 | 20055 | 21618 | 21633 | 21634 | 21635 | 21636 | 21637 | 21638 | 21639 | 21640 | 21641 | 21721 | 21762 | 21765 | 21766 | 21767 | 21768 | 21769 | 21770 | 21846 | 21987 | 21988 | 21989 | 21990 | 21991 | 21992 | 21993 | 21994 | 21995 | 22194 | 22195 | 22196 | 22200 | 22599 | 22673 | 22688 | 22775 | 22786 | 22879 | 22898 | 22917 | 23048 | 23202 | 23440 | 23673 | 24130 | 24503 | 25416 | 26060 | 26530 | 26858 | 27382 | 27383 | 27384 | 29510 |  |
| ref | C | T | C | C | A | G | C | T | G | T | T | C | G | C | A | C | C | C | G | G | T | T | T | G | G | T | T | T | T | A | C | C | T | T | C | C | C | A | G | C | A | C | T | A | C | C | C | C | C | T | G | C | C | T | A | C | A | T | G | C | G | T | G | T | T | T | A | T | T | A | T | T | T | G | T | A | G | A | C | G | T | G | C | A | C | C | C | C | C | A | C | G | A | T | A |  |
| BA.1.1 | C | T | T | C | G | G | C | G | - | - | - | T | A | C | A | C | C | C | G | - | - | - | - | - | G | T | T | T | T | G | C | C | C | T | T | C | C | G | T | C | A | C | T | A | C | C | C | C | C | T | G | C | T | - | - | - | - | - | - | T | - | - | - | - | - | - | - | - | - | - | - | - | T | A | C | A | G | A | T | A | T | A | A | A | C | A | T | C | C | G | C | G | A | T | A |  |
| BA.2 | C | T | T | C | G | G | C | G | - | - | - | T | A | C | A | C | C | C | G | G | T | T | T | G | - | - | - | - | - | A | T | T | T | C | C | T | T | A | G | T | G | T | - | - | - | - | - | - | - | - | - | T | C | T | A | C | A | T | G | C | A | T | G | T | T | T | A | T | T | A | T | T | G | G | T | G | A | C | C | G | A | G | C | G | T | C | C | T | T | A | T | C | T | C | C | #1210797 |
| BA.2.12.1 | T | G | C | T | A | A | T | T | G | T | T | C | G | T | G | T | T | T | A | G | T | T | T | G | - | - | - | - | - | A | T | T | T | C | C | T | T | A | G | T | G | T | - | - | - | - | - | - | - | - | - | T | C | T | A | C | A | T | G | C | A | T | G | T | T | T | A | T | T | A | T | T | G | G | T | G | A | C | C | G | A | G | C | G | T | C | C | T | T | A | T | C | T | C | C |  |
| mut |  |  |  |  |  |  |  |  |  |  |  |  |  |  |  |  |  |  |  |  |  |  |  |  |  |  |  |  |  |  |  |  |  |  |  |  |  |  |  |  |  |  |  |  |  |  |  |  |  |  |  |  |  |  |  |  |  |  |  |  |  |  |  |  |  |  |  |  |  |  |  |  |  |  |  |  |  |  |  |  |  |  |  |  |  |  |  |  |  |  |  |  |  |  |  |  |

|  |  |  |  |  |  |  |  |  |  |  |
| --- | --- | --- | --- | --- | --- | --- | --- | --- | --- | --- |
| pos | 13945 | 22197 | 22792 | 24310 | 25091 | 29728 | 29757 | 29758 | 29759 |  |
| ref | C | T | C | C | G | T | G | T | G |  |
| BA.2 | C | C | C | T | A | T | G | T | G |  |
| BA.2 | C | C | C | T | A | C | A | C | C | #1212556 |
| BA.2 | T | T | T | C | G | C | A | C | C |  |
| mut |  |  |  |  |  |  |  |  |  |  |

|  |  |  |  |  |  |  |  |  |  |  |  |  |
| --- | --- | --- | --- | --- | --- | --- | --- | --- | --- | --- | --- | --- |
| pos | 1227 | 4893 | 8772 | 9344 | 22194 | 22195 | 22196 | 22197 | 22198 | 22202 | 22204 |  |
| ref | T | C | C | C | A | T | T | T | A | C | T |  |
| BA.1.1 | T | T | T | T | - | - | - | T | A | C | T |  |
| BA.1.1 | T | T | T | T | A | G | T | G | C | A | C | #1212752 |
| BA.1.1 | C | C | C | C | A | G | T | G | C | A | C |  |
| mut |  |  |  |  |  |  |  |  |  |  |  |  |

|  |  |  |  |  |  |  |  |  |  |  |  |  |  |  |  |  |  |  |  |
| --- | --- | --- | --- | --- | --- | --- | --- | --- | --- | --- | --- | --- | --- | --- | --- | --- | --- | --- | --- |
| pos | 1210 | 2470 | 2875 | 5431 | 7782 | 8655 | 12025 | 22120 | 22194 | 22195 | 22196 | 22197 | 22198 | 22202 | 22204 | 22599 | 27893 | 29781 |  |
| ref | G | C | G | A | A | C | C | C | A | T | T | T | A | C | T | G | C | G |  |
| BA.1.1 | T | T | A | A | A | T | T | T | - | - | - | T | A | C | T | A | C | - |  |
| BA.1.1 | T | T | A | G | G | T | T | T | A | G | T | G | C | A | C | A | A | G | #1214060 |
| BA.1 | G | C | G | A | A | C | C | C | A | G | T | G | C | A | C | G | C | G |  |
| mut |  |  |  | A>G | A>G |  |  |  |  |  |  |  |  |  |  | G>A | C>A |  |  |

|  |  |  |  |  |  |  |  |  |  |  |  |  |  |  |  |  |
| --- | --- | --- | --- | --- | --- | --- | --- | --- | --- | --- | --- | --- | --- | --- | --- | --- |
| pos | 7820 | 15439 | 21595 | 22194 | 22195 | 22196 | 22197 | 22198 | 22202 | 22204 | 25538 | 26882 | 27247 | 28816 | 29167 |  |
| ref | T | G | C | A | T | T | T | A | C | T | G | C | C | A | C |  |
| BA.1.1 | T | G | C | A | G | T | G | C | A | C | G | C | C | A | C |  |
| BA.1.1 | G | A | T | A | G | T | G | C | A | C | T | T | T | G | T | #1221949 |
| BA.1.1 | T | G | T | - | - | - | T | A | C | T | T | T | C | G | T |  |
| mut | T>G | G>A | C>T |  |  |  |  |  |  |  |  |  | C>T |  |  |  |

|  |  |  |  |  |  |  |  |  |  |  |  |  |  |  |  |
| --- | --- | --- | --- | --- | --- | --- | --- | --- | --- | --- | --- | --- | --- | --- | --- |
| pos | 44 | 507 | 508 | 509 | 2942 | 3637 | 5425 | 6196 | 8991 | 18479 | 23978 | 25810 | 28854 | 29223 |  |
| ref | C | A | T | G | C | A | G | C | C | A | T | C | C | T |  |
| BA.2 | T | T | C | A | C | A | T | T | C | A | C | C | C | T |  |
| BA.2.3 | T | T | C | A | T | C | G | C | T | G | T | T | T | C | #1221979 |
| BA.2.3 | C | A | T | G | T | C | G | C | T | G | T | T | T | T |  |
| mut |  |  |  |  |  |  |  |  |  |  |  |  |  | T>C |  |

|  |  |  |  |  |  |  |  |  |  |  |  |  |  |  |
| --- | --- | --- | --- | --- | --- | --- | --- | --- | --- | --- | --- | --- | --- | --- |
| pos | 6742 | 9693 | 13968 | 15774 | 22088 | 22194 | 22195 | 22196 | 22197 | 22198 | 22202 | 22204 | 28810 |  |
| ref | G | C | C | C | C | A | T | T | T | A | C | T | C |  |
| BA.1.1 | A | T | T | T | T | - | - | - | T | A | C | T | T |  |
| BA.1.1 | A | T | T | T | T | A | G | T | G | C | A | C | T | #1225055 |
| BA.1.1 | G | C | C | C | C | A | G | T | G | C | A | C | C |  |
| mut |  |  |  |  |  |  |  |  |  |  |  |  | C>T |  |

|  |  |  |  |  |  |  |  |  |  |  |  |  |  |
| --- | --- | --- | --- | --- | --- | --- | --- | --- | --- | --- | --- | --- | --- |
| pos | 6548 | 9300 | 11674 | 15009 | 21721 | 22600 | 22917 | 23673 | 29729 | 29757 | 29758 | 29759 |  |
| ref | C | G | C | T | C | A | T | C | T | G | T | G |  |
| BA.2.12.1 | C | G | T | C | T | A | A | T | T | G | T | G |  |
| BA.2.12.1 | C | G | T | C | T | A | A | T | C | A | C | C | #1226050 |
| BA.2 | T | A | C | T | C | T | T | C | C | A | C | C |  |
| mut |  |  |  |  |  |  |  |  |  |  |  |  |  |

|  |  |  |  |  |  |  |  |  |  |  |  |  |  |  |  |
| --- | --- | --- | --- | --- | --- | --- | --- | --- | --- | --- | --- | --- | --- | --- | --- |
| pos | 44 | 507 | 508 | 509 | 3787 | 9298 | 18060 | 20320 | 22792 | 28522 | 29179 | 29200 | 29614 | 29781 |  |
| ref | C | A | T | G | C | C | C | C | C | A | G | C | C | G |  |
| BA.2.3 | C | A | T | G | T | C | C | C | C | A | G | C | C | G |  |
| BA.2.3 | C | A | T | G | T | T | T | T | T | C | C | T | T | T | #1234809 |
| BA.2.3 | T | T | C | A | C | T | T | C | C | C | C | C | C | G |  |
| mut |  |  |  |  |  |  |  | C>T | C>T |  |  | C>T | C>T | G>T |  |

|  |  |  |  |  |  |  |  |  |  |  |  |  |
| --- | --- | --- | --- | --- | --- | --- | --- | --- | --- | --- | --- | --- |
| pos | 44 | 3961 | 8821 | 11266 | 11674 | 16293 | 21537 | 22917 | 23673 | 25416 | 28678 |  |
| ref | C | C | A | G | C | C | T | T | C | C | T |  |
| BA.2.64 | C | T | G | T | C | T | C | T | C | C | C |  |
| BA.2.64 | C | T | G | T | C | T | C | A | T | T | T | #1238543 |
| BA.2.12.1 | T | C | A | G | T | C | T | A | T | T | T |  |
| mut |  |  |  |  |  |  |  |  |  |  |  |  |

|  |  |  |  |  |  |  |  |  |  |  |  |  |  |  |  |  |  |  |  |  |  |  |  |  |  |  |  |  |  |  |
| --- | --- | --- | --- | --- | --- | --- | --- | --- | --- | --- | --- | --- | --- | --- | --- | --- | --- | --- | --- | --- | --- | --- | --- | --- | --- | --- | --- | --- | --- | --- |
| pos | 44 | 9866 | 10507 | 12160 | 19174 | 21765 | 21766 | 21767 | 21768 | 21769 | 21770 | 22792 | 22916 | 22917 | 23018 | 23040 | 23767 | 24368 | 26529 | 26858 | 26921 | 27259 | 27382 | 27383 | 27384 | 27438 | 27889 | 29666 | 29754 |  |
| ref | C | C | C | G | G | T | A | C | A | T | G | C | C | T | T | A | A | G | G | C | T | A | G | A | T | T | C | C | C |  |
| BA.5.1 | T | C | T | A | A | - | - | - | - | - | - | C | C | G | G | A | A | G | A | C | T | A | G | A | T | C | T | T | T |  |
| BA.2 | T | C | T | A | A | - | - | - | - | - | - | C | A | T | T | G | G | T | G | T | C | C | C | T | C | T | C | C | C | #1239723 |
| BA.2.13 | C | T | C | G | G | T | A | C | A | T | G | T | A | T | T | G | G | T | G | T | T | C | C | T | C | T | C | C | C |  |
| mut |  |  |  |  |  |  |  |  |  |  |  |  |  |  |  |  |  |  |  |  | T>C |  |  |  |  |  |  |  |  |  |

|  |  |  |  |  |  |  |  |  |  |  |  |  |  |  |  |  |  |  |  |  |  |  |  |  |  |  |
| --- | --- | --- | --- | --- | --- | --- | --- | --- | --- | --- | --- | --- | --- | --- | --- | --- | --- | --- | --- | --- | --- | --- | --- | --- | --- | --- |
| pos | 44 | 1914 | 3885 | 4080 | 8991 | 11522 | 11575 | 11674 | 11750 | 14857 | 15009 | 15552 | 16092 | 18814 | 21191 | 21721 | 22917 | 23673 | 25416 | 25521 | 25810 | 27765 | 27798 | 27945 | 28727 |  |
| ref | C | G | C | G | C | T | C | C | C | G | T | T | C | C | C | C | T | C | C | C | C | C | G | C | G |  |
| BA.2.3 | C | A | T | A | T | T | C | C | T | G | T | A | C | T | C | C | T | C | C | C | T | T | G | C | G |  |
| BA.2.12.1 | C | A | T | A | T | T | C | C | T | A | C | T | T | C | T | T | A | T | T | T | C | C | T | T | T | #1251638 |
| BA.2.12.1 | T | G | C | G | C | G | T | T | C | A | C | T | T | C | C | T | A | T | T | T | C | C | T | T | T |  |
| mut |  |  |  |  |  |  |  |  |  |  |  |  |  |  | C>T |  |  |  |  |  |  |  |  |  |  |  |

|  |  |  |  |  |  |  |  |  |  |  |  |  |  |  |  |  |  |  |  |  |  |  |  |  |  |  |  |  |  |  |  |  |  |  |  |  |  |  |  |  |  |  |  |  |  |  |  |  |  |  |  |  |  |  |  |  |  |  |  |  |  |  |  |  |  |  |  |  |  |  |  |  |  |  |  |  |  |  |  |  |  |  |  |  |  |  |  |  |  |  |  |  |  |  |  |
| --- | --- | --- | --- | --- | --- | --- | --- | --- | --- | --- | --- | --- | --- | --- | --- | --- | --- | --- | --- | --- | --- | --- | --- | --- | --- | --- | --- | --- | --- | --- | --- | --- | --- | --- | --- | --- | --- | --- | --- | --- | --- | --- | --- | --- | --- | --- | --- | --- | --- | --- | --- | --- | --- | --- | --- | --- | --- | --- | --- | --- | --- | --- | --- | --- | --- | --- | --- | --- | --- | --- | --- | --- | --- | --- | --- | --- | --- | --- | --- | --- | --- | --- | --- | --- | --- | --- | --- | --- | --- | --- | --- | --- | --- | --- | --- |
| pos | 44 | 670 | 686 | 687 | 688 | 689 | 690 | 691 | 692 | 693 | 694 | 1627 | 2790 | 2832 | 4184 | 4321 | 5386 | 5826 | 6513 | 6514 | 6515 | 6979 | 8393 | 9344 | 9424 | 9534 | 10198 | 10447 | 11283 | 11284 | 11285 | 11286 | 11287 | 11292 | 11293 | 11294 | 11295 | 11296 | 11537 | 12160 | 12880 | 13195 | 15240 | 15714 | 17410 | 19955 | 20055 | 21618 | 21633 | 21634 | 21635 | 21636 | 21637 | 21638 | 21639 | 21640 | 21641 | 21762 | 21846 | 21987 | 21988 | 21989 | 21990 | 21991 | 21992 | 21993 | 21994 | 21995 | 22110 | 22194 | 22195 | 22196 | 22200 | 22673 | 22688 | 22775 | 22786 | 22898 | 22917 | 23018 | 23040 | 23048 | 23202 | 24130 | 24503 | 26060 | 26529 | 26530 | 27038 | 27259 | 27889 | 28330 | 29510 | 29754 |  |
| ref | C | T | A | A | G | T | C | A | T | T | T | C | C | A | G | C | T | C | G | T | T | T | G | C | A | C | C | G | G | T | T | T | G | G | T | T | T | T | A | G | C | T | C | C | C | C | A | C | T | A | C | C | C | C | C | T | G | C | C | G | T | G | T | T | T | A | T | T | A | A | T | T | T | T | A | G | A | G | T | T | A | G | C | C | C | C | G | A | A | A | C | A | A | C |  |
| BA.1 | C | T | G | C | G | C | C | G | C | T | A | C | C | G | G | C | G | C | - | - | - | T | A | C | A | C | C | G | - | - | - | - | - | G | T | T | T | T | G | G | C | C | T | C | C | C | A | C | T | A | C | C | C | C | C | T | G | T | T | - | - | - | - | - | - | - | - | - | A | - | - | - | T | C | A | G | A | A | T | T | G | A | A | A | T | C | G | G | A | C | C | A | A | C |  |
| BA.5.2.1 | C | G | G | C | G | C | C | G | C | T | A | T | T | A | A | T | T | T | G | T | T | G | G | T | G | T | T | A | G | T | T | T | G | - | - | - | - | - | A | A | T | T | C | T | T | T | G | T | - | - | - | - | - | - | - | - | - | C | C | A | T | G | T | T | T | A | T | T | T | A | T | T | G | T | G | A | C | G | G | G | A | G | C | C | C | T | A | A | G | A | T | G | C | T | #1253364 |
| BA.5.2.1 | T | G | - | - | - | - | - | - | - | - | - | T | T | A | A | T | T | T | G | T | T | G | G | T | G | T | T | A | G | T | T | T | G | - | - | - | - | - | A | A | T | T | C | T | T | T | G | T | - | - | - | - | - | - | - | - | - | C | C | A | T | G | T | T | T | A | T | T | T | A | T | T | G | T | G | A | C | G | G | G | A | G | C | C | C | T | A | A | G | A | T | G | C | T |  |
| mut |  | T>G |  |  |  |  |  |  |  |  |  |  |  |  |  |  |  |  |  |  |  |  |  |  |  |  |  |  |  |  |  |  |  |  |  |  |  |  |  |  |  |  |  |  |  |  |  |  |  |  |  |  |  |  |  |  |  |  |  |  |  |  |  |  |  |  |  |  |  |  |  |  |  |  |  |  |  |  |  |  |  |  |  |  |  |  |  |  |  |  |  |  |  |  |  |

|  |  |  |  |  |  |  |  |  |  |  |  |  |  |  |  |  |  |  |
| --- | --- | --- | --- | --- | --- | --- | --- | --- | --- | --- | --- | --- | --- | --- | --- | --- | --- | --- |
| pos | 44 | 6351 | 8991 | 11674 | 11986 | 15009 | 18060 | 18582 | 21721 | 22020 | 22917 | 23673 | 25416 | 25810 | 26408 | 28708 | 29059 |  |
| ref | C | A | C | C | C | T | C | C | C | T | T | C | C | C | C | C | T |  |
| BA.2.3 | C | G | T | C | T | T | T | C | C | C | T | C | C | T | T | T | C |  |
| BA.2.12.1 | C | G | T | C | T | C | C | T | T | T | A | T | T | C | C | C | T | #1257069 |
| BA.2.12.1 | T | A | C | T | C | C | C | T | T | T | A | T | T | C | C | C | T |  |
| mut |  |  |  |  |  |  |  |  |  |  |  |  |  |  |  |  |  |  |

|  |  |  |  |  |  |  |  |  |  |  |  |  |
| --- | --- | --- | --- | --- | --- | --- | --- | --- | --- | --- | --- | --- |
| pos | 2897 | 4893 | 6821 | 11674 | 13665 | 14097 | 21721 | 22326 | 22917 | 23673 | 25416 |  |
| ref | G | C | A | C | C | C | C | C | T | C | C |  |
| BA.2 | T | T | G | C | T | T | C | T | T | C | C |  |
| BA.2.12.1 | T | T | G | C | T | T | T | C | A | T | T | #1257342 |
| BA.2.12.1 | G | C | A | T | C | C | T | C | A | T | T |  |
| mut |  |  |  |  |  |  |  |  |  |  |  |  |

|  |  |  |  |  |  |  |  |  |  |  |  |  |  |  |
| --- | --- | --- | --- | --- | --- | --- | --- | --- | --- | --- | --- | --- | --- | --- |
| pos | 44 | 1989 | 3901 | 11674 | 14793 | 15009 | 21721 | 22917 | 23673 | 24793 | 25416 | 26873 | 28817 |  |
| ref | C | A | G | C | C | T | C | T | C | T | C | C | G |  |
| BA.2.12.1 | T | A | G | T | T | C | T | A | T | T | T | C | G |  |
| BA.2 | T | G | G | T | T | C | C | T | C | C | C | T | T | #1258847 |
| BA.2 | C | A | A | C | C | T | C | T | C | C | C | C | T |  |
| mut |  | A>G |  |  |  |  |  |  |  |  |  | C>T |  |  |

|  |  |  |  |  |  |  |  |  |  |  |  |  |  |  |  |  |  |  |  |
| --- | --- | --- | --- | --- | --- | --- | --- | --- | --- | --- | --- | --- | --- | --- | --- | --- | --- | --- | --- |
| pos | 44 | 1376 | 1598 | 2061 | 2675 | 4993 | 9856 | 11674 | 14925 | 15009 | 18486 | 21721 | 22917 | 23673 | 23929 | 25416 | 28720 | 28934 |  |
| ref | C | C | G | C | C | C | G | C | C | T | C | C | T | C | C | C | C | T |  |
| BA.2 | C | T | G | T | T | T | T | C | T | T | C | C | T | C | C | C | C | C |  |
| BA.2.12.1 | C | T | G | T | T | T | G | T | C | C | T | T | A | T | T | T | T | T | #1259749 |
| BA.2.12.1 | T | C | A | C | C | C | G | T | C | C | T | T | A | T | T | T | T | T |  |
| mut |  |  |  |  |  |  |  |  |  |  |  |  |  |  |  |  |  |  |  |

|  |  |  |  |  |  |  |  |  |  |  |  |  |  |  |  |  |  |  |  |  |  |  |  |  |  |  |  |  |  |  |  |  |  |  |  |  |
| --- | --- | --- | --- | --- | --- | --- | --- | --- | --- | --- | --- | --- | --- | --- | --- | --- | --- | --- | --- | --- | --- | --- | --- | --- | --- | --- | --- | --- | --- | --- | --- | --- | --- | --- | --- | --- |
| pos | 44 | 1230 | 3610 | 4111 | 9866 | 12160 | 15342 | 16428 | 20178 | 21765 | 21766 | 21767 | 21768 | 21769 | 21770 | 22792 | 22917 | 23018 | 23040 | 24433 | 25350 | 25624 | 26529 | 26858 | 27259 | 27382 | 27383 | 27384 | 27438 | 27889 | 28472 | 29508 | 29666 | 29754 | 29762 |  |
| ref | C | A | T | T | C | G | C | C | C | T | A | C | A | T | G | C | T | T | A | A | C | C | G | C | A | G | A | T | T | C | C | G | C | C | C |  |
| BA.5.1.5 | T | A | T | T | C | A | C | T | C | - | - | - | - | - | - | C | G | G | A | A | A | C | A | C | A | G | A | T | C | T | C | G | T | T | C |  |
| BA.5.1.5 | T | A | T | T | C | A | C | T | C | - | - | - | - | - | - | C | G | G | A | A | A | C | A | C | A | G | A | T | C | C | T | T | C | C | T | #1263357 |
| BA.2.9 | C | G | C | C | T | G | T | C | T | T | A | C | A | T | G | T | T | T | G | G | C | T | G | T | C | C | T | C | T | C | T | T | C | C | C |  |
| mut |  |  |  |  |  |  |  |  |  |  |  |  |  |  |  |  |  |  |  |  |  |  |  |  |  |  |  |  |  |  |  |  |  |  | C>T |  |

|  |  |  |  |  |  |  |  |  |  |  |  |  |  |  |  |  |  |  |  |  |  |  |  |  |  |  |  |  |  |  |  |  |  |
| --- | --- | --- | --- | --- | --- | --- | --- | --- | --- | --- | --- | --- | --- | --- | --- | --- | --- | --- | --- | --- | --- | --- | --- | --- | --- | --- | --- | --- | --- | --- | --- | --- | --- |
| pos | 702 | 1422 | 1931 | 3875 | 9866 | 12068 | 12160 | 16935 | 21765 | 21766 | 21767 | 21768 | 21769 | 21770 | 22792 | 22917 | 23018 | 23040 | 23323 | 25416 | 25534 | 25537 | 26043 | 26529 | 26858 | 27259 | 27382 | 27383 | 27384 | 27889 | 28681 | 29754 |  |
| ref | G | A | C | G | C | C | G | G | T | A | C | A | T | G | C | T | T | A | T | C | G | G | T | G | C | A | G | A | T | C | G | C |  |
| BE.1 | G | C | A | A | C | T | A | A | - | - | - | - | - | - | T | G | G | A | C | C | T | T | T | A | C | A | G | A | T | T | T | T |  |
| BA.4 | A | C | A | A | C | T | A | A | - | - | - | - | - | - | T | G | G | A | C | T | G | G | C | G | T | C | C | T | C | C | G | C | #1264200 |
| BA.2 | G | A | C | G | T | C | G | G | T | A | C | A | T | G | C | T | T | G | T | T | G | G | T | G | T | C | C | T | C | C | G | C |  |
| mut | G>A |  |  |  |  |  |  |  |  |  |  |  |  |  |  |  |  |  |  |  |  |  | T>C |  |  |  |  |  |  |  |  |  |  |

|  |  |  |  |  |  |  |  |  |  |  |  |  |  |  |  |  |  |  |  |  |  |  |  |  |  |  |  |  |  |  |  |  |  |
| --- | --- | --- | --- | --- | --- | --- | --- | --- | --- | --- | --- | --- | --- | --- | --- | --- | --- | --- | --- | --- | --- | --- | --- | --- | --- | --- | --- | --- | --- | --- | --- | --- | --- |
| pos | 1931 | 8991 | 9866 | 11674 | 12160 | 15009 | 16935 | 18167 | 18788 | 21721 | 21765 | 21766 | 21767 | 21768 | 21769 | 21770 | 22917 | 23018 | 23040 | 23673 | 25416 | 26529 | 26858 | 27259 | 27382 | 27383 | 27384 | 27875 | 27889 | 28674 | 28681 | 29754 |  |
| ref | C | C | C | C | G | T | G | C | C | C | T | A | C | A | T | G | T | T | A | C | C | G | C | A | G | A | T | T | C | C | G | C |  |
| BE.1 | A | C | C | C | A | T | A | C | C | C | - | - | - | - | - | - | G | G | A | C | C | A | C | A | G | A | T | T | T | C | T | T |  |
| BA.2.12.1 | A | C | C | C | A | T | A | C | C | T | T | A | C | A | T | G | A | T | G | T | T | G | T | C | C | T | C | C | C | T | G | C | #1267873 |
| BA.2.12.1 | C | T | T | T | G | C | G | T | T | T | T | A | C | A | T | G | A | T | G | T | T | G | T | C | C | T | C | C | C | C | G | C |  |
| mut |  |  |  |  |  |  |  |  |  |  |  |  |  |  |  |  |  |  |  |  |  |  |  |  |  |  |  |  |  | C>T |  |  |  |

|  |  |  |  |  |  |  |  |  |  |  |  |  |  |  |  |  |  |  |  |  |  |  |  |  |  |  |  |  |  |  |  |
| --- | --- | --- | --- | --- | --- | --- | --- | --- | --- | --- | --- | --- | --- | --- | --- | --- | --- | --- | --- | --- | --- | --- | --- | --- | --- | --- | --- | --- | --- | --- | --- |
| pos | 362 | 686 | 687 | 688 | 689 | 690 | 691 | 692 | 693 | 694 | 1627 | 3165 | 14742 | 18084 | 22104 | 23535 | 24106 | 26529 | 26858 | 27038 | 27259 | 27382 | 27383 | 27384 | 27788 | 27889 | 28330 | 28724 | 28957 | 29754 |  |
| ref | G | A | A | G | T | C | A | T | T | T | C | C | T | C | G | A | C | G | C | A | A | G | A | T | G | C | A | C | C | C |  |
| BA.4 | G | - | - | - | - | - | - | - | - | - | C | T | C | C | G | G | T | G | T | A | C | C | T | C | T | C | A | T | C | C |  |
| BA.5.2.1 | A | - | - | - | - | - | - | - | - | - | C | T | C | C | G | G | T | A | C | G | A | G | A | T | G | T | G | C | T | T | #1270184 |
| BF.6 | G | A | A | G | T | C | A | T | T | T | T | C | T | T | C | A | C | A | C | G | A | G | A | T | G | T | G | C | T | T |  |
| mut | G>A |  |  |  |  |  |  |  |  |  |  |  |  |  |  |  |  |  |  |  |  |  |  |  |  |  |  |  |  |  |  |

|  |  |  |  |  |  |  |  |  |  |  |  |  |  |  |  |  |  |  |  |  |  |  |  |  |  |  |  |  |  |  |  |
| --- | --- | --- | --- | --- | --- | --- | --- | --- | --- | --- | --- | --- | --- | --- | --- | --- | --- | --- | --- | --- | --- | --- | --- | --- | --- | --- | --- | --- | --- | --- | --- |
| pos | 526 | 5259 | 6107 | 6979 | 7732 | 9866 | 11674 | 12160 | 12439 | 21765 | 21766 | 21767 | 21768 | 21769 | 21770 | 22917 | 23018 | 23040 | 23673 | 25416 | 26529 | 26858 | 27038 | 27259 | 27382 | 27383 | 27384 | 27889 | 28330 | 29754 |  |
| ref | G | C | A | T | C | C | C | G | C | T | A | C | A | T | G | T | T | A | C | C | G | C | A | A | G | A | T | C | A | C |  |
| BA.2.12.1 | G | C | G | T | C | T | T | G | C | T | A | C | A | T | G | A | T | G | T | T | G | T | A | C | C | T | C | C | A | C |  |
| BA.5.2.1 | T | T | G | T | T | T | T | G | T | T | A | C | A | T | G | A | T | G | T | C | A | C | G | A | G | A | T | T | G | T | #1271665 |
| BA.5.2.1 | G | C | A | G | C | C | C | A | C | - | - | - | - | - | - | G | G | A | C | C | A | C | G | A | G | A | T | T | G | T |  |
| mut | G>T | C>T |  |  | C>T |  |  |  | C>T |  |  |  |  |  |  |  |  |  |  |  |  |  |  |  |  |  |  |  |  |  |  |

|  |  |  |  |  |  |  |  |  |  |  |  |  |  |
| --- | --- | --- | --- | --- | --- | --- | --- | --- | --- | --- | --- | --- | --- |
| pos | 1627 | 3478 | 16376 | 18084 | 22104 | 27038 | 27438 | 27630 | 27656 | 28330 | 28957 | 29666 |  |
| ref | C | T | C | C | G | A | T | C | T | A | C | C |  |
| BA.5.1 | C | C | T | C | G | A | C | T | C | A | C | T |  |
| BA.5.2.1 | C | C | T | C | C | G | T | C | T | G | T | C | #1273374 |
| BF.6 | T | T | C | T | C | G | T | C | T | G | T | C |  |
| mut |  |  |  |  |  |  |  |  |  |  |  |  |  |

|  |  |  |  |  |  |  |  |  |  |  |  |  |
| --- | --- | --- | --- | --- | --- | --- | --- | --- | --- | --- | --- | --- |
| pos | 594 | 1627 | 4347 | 5284 | 27038 | 27438 | 28330 | 28370 | 28371 | 29614 | 29666 |  |
| ref | A | C | C | C | A | T | A | A | G | C | C |  |
| BA.5.1 | G | C | T | C | A | C | A | - | G | T | T |  |
| BA.5 | G | C | T | C | A | C | G | T | T | C | C | #1274474 |
| BA.5.2.1 | A | T | C | T | G | T | G | T | T | C | C |  |
| mut |  |  |  |  |  |  |  |  |  |  |  |  |

|  |  |  |  |  |  |  |  |  |  |  |  |  |  |  |  |  |  |  |  |  |  |  |  |  |  |  |
| --- | --- | --- | --- | --- | --- | --- | --- | --- | --- | --- | --- | --- | --- | --- | --- | --- | --- | --- | --- | --- | --- | --- | --- | --- | --- | --- |
| pos | 686 | 687 | 688 | 689 | 690 | 691 | 692 | 693 | 694 | 1931 | 21161 | 23535 | 25315 | 25644 | 26529 | 26621 | 26690 | 26858 | 27259 | 27382 | 27383 | 27384 | 27788 | 27889 | 29754 |  |
| ref | A | A | G | T | C | A | T | T | T | C | C | A | A | G | G | T | G | C | A | G | A | T | G | C | C |  |
| BA.4 | - | - | - | - | - | - | - | - | - | C | C | G | G | T | G | C | T | T | C | C | T | C | T | C | C |  |
| BA.5 | A | A | G | T | C | A | T | T | T | C | T | G | G | G | G | C | G | C | A | G | A | T | G | T | T | #1275419 |
| BA.5.3 | A | A | G | T | C | A | T | T | T | A | C | A | A | G | A | T | G | C | A | G | A | T | G | T | T |  |
| mut | ->A | ->A | ->G | ->T | ->C | ->A | ->T | ->T | ->T |  | C>T |  |  | T>G |  |  |  |  |  |  |  |  |  |  |  |  |

|  |  |  |  |  |  |  |  |  |  |  |  |  |  |  |  |  |  |  |  |  |  |  |  |  |  |  |  |  |  |  |  |  |
| --- | --- | --- | --- | --- | --- | --- | --- | --- | --- | --- | --- | --- | --- | --- | --- | --- | --- | --- | --- | --- | --- | --- | --- | --- | --- | --- | --- | --- | --- | --- | --- | --- |
| pos | 686 | 687 | 688 | 689 | 690 | 691 | 692 | 693 | 694 | 1627 | 1824 | 2701 | 3859 | 5173 | 6979 | 13078 | 13721 | 18657 | 20233 | 26529 | 26858 | 27038 | 27259 | 27382 | 27383 | 27384 | 27788 | 27889 | 28330 | 28724 | 29754 |  |
| ref | A | A | G | T | C | A | T | T | T | C | C | C | A | C | T | T | C | C | C | G | C | A | A | G | A | T | G | C | A | C | C |  |
| BA.4.2 | - | - | - | - | - | - | - | - | - | C | T | C | A | A | T | C | T | C | C | G | T | A | C | C | T | C | T | C | A | T | C |  |
| BA.5.2.1 | - | - | - | - | - | - | - | - | - | C | T | T | A | A | T | T | C | T | T | A | C | G | A | G | A | T | G | T | G | C | T | #1276533 |
| BA.5.2.1 | A | A | G | T | C | A | T | T | T | T | C | C | G | C | G | T | C | T | C | A | C | G | A | G | A | T | G | T | G | C | T |  |
| mut |  |  |  |  |  |  |  |  |  |  |  | C>T |  |  |  |  |  |  | C>T |  |  |  |  |  |  |  |  |  |  |  |  |  |

|  |  |  |  |  |  |  |  |  |  |  |  |  |  |  |  |  |  |  |  |  |  |  |  |  |  |  |  |  |  |  |  |
| --- | --- | --- | --- | --- | --- | --- | --- | --- | --- | --- | --- | --- | --- | --- | --- | --- | --- | --- | --- | --- | --- | --- | --- | --- | --- | --- | --- | --- | --- | --- | --- |
| pos | 7765 | 9866 | 12160 | 15594 | 16954 | 21765 | 21766 | 21767 | 21768 | 21769 | 21770 | 21789 | 22792 | 22916 | 22917 | 23018 | 23040 | 23120 | 23767 | 24383 | 26529 | 26858 | 27259 | 27382 | 27383 | 27384 | 27889 | 28507 | 29227 | 29754 |  |
| ref | C | C | G | G | C | T | A | C | A | T | G | C | C | C | T | T | A | G | A | A | G | C | A | G | A | T | C | C | G | C |  |
| BA.2.13.1 | C | T | G | G | T | T | A | C | A | T | G | C | T | A | T | T | G | G | G | T | G | T | C | C | T | C | C | T | G | C |  |
| BA.2 | T | T | G | T | T | T | A | C | A | T | G | C | T | A | T | T | G | T | G | T | A | C | A | G | A | T | T | C | A | T | #1276626 |
| BA.5.5 | C | C | A | G | C | - | - | - | - | - | - | T | C | C | G | G | A | G | A | A | A | C | A | G | A | T | T | C | A | T |  |
| mut | C>T |  |  | G>T |  |  |  |  |  |  |  |  |  |  |  |  |  | G>T |  |  |  |  |  |  |  |  |  |  |  |  |  |

|  |  |  |  |  |  |  |  |  |  |  |  |  |  |  |  |  |  |  |  |  |  |  |  |  |  |  |  |  |  |  |  |  |
| --- | --- | --- | --- | --- | --- | --- | --- | --- | --- | --- | --- | --- | --- | --- | --- | --- | --- | --- | --- | --- | --- | --- | --- | --- | --- | --- | --- | --- | --- | --- | --- | --- |
| pos | 44 | 120 | 4893 | 7393 | 7735 | 9866 | 12160 | 13665 | 18568 | 21765 | 21766 | 21767 | 21768 | 21769 | 21770 | 22917 | 23018 | 23040 | 26529 | 26858 | 27259 | 27382 | 27383 | 27384 | 27438 | 27889 | 28528 | 28975 | 29666 | 29669 | 29754 |  |
| ref | C | T | C | G | C | C | G | C | C | T | A | C | A | T | G | T | T | A | G | C | A | G | A | T | T | C | C | G | C | A | C |  |
| BA.2 | C | C | T | G | T | T | G | T | C | T | A | C | A | T | G | T | T | G | G | T | C | C | T | C | T | C | T | T | C | G | C |  |
| BA.5.1 | C | C | T | T | C | T | G | T | C | - | - | - | - | - | - | G | G | A | A | C | A | G | A | T | C | T | C | G | T | A | T | #1277345 |
| BA.5.1 | T | T | C | G | C | C | A | C | T | - | - | - | - | - | - | G | G | A | A | C | A | G | A | T | C | T | C | G | T | A | T |  |
| mut |  |  |  | G>T | T>C |  |  |  |  |  |  |  |  |  |  |  |  |  |  |  |  |  |  |  |  |  |  |  |  |  |  |  |

|  |  |  |  |  |  |  |  |  |  |  |  |  |  |
| --- | --- | --- | --- | --- | --- | --- | --- | --- | --- | --- | --- | --- | --- |
| pos | 625 | 1627 | 1931 | 2305 | 11750 | 12310 | 16616 | 16935 | 27012 | 27513 | 28330 | 28681 |  |
| ref | G | C | C | G | C | G | C | G | C | C | A | G |  |
| BA.5.2.3 | T | T | C | A | C | A | A | G | T | T | G | G |  |
| BE.1 | T | T | C | A | T | G | C | A | C | C | A | T | #1278883 |
| BE.1 | G | C | A | G | C | G | C | A | C | C | A | T |  |
| mut |  |  |  |  | C>T |  |  |  |  |  |  |  |  |

|  |  |  |  |  |  |  |  |  |  |  |  |  |  |  |  |  |
| --- | --- | --- | --- | --- | --- | --- | --- | --- | --- | --- | --- | --- | --- | --- | --- | --- |
| pos | 44 | 686 | 687 | 689 | 691 | 692 | 694 | 1627 | 5826 | 6979 | 22110 | 27038 | 27438 | 28330 | 29666 |  |
| ref | C | A | A | T | A | T | T | C | C | T | A | A | T | A | C |  |
| BA.5.2.1 | C | G | C | C | G | C | A | T | T | G | T | G | T | G | C |  |
| BA.5.1 | C | G | C | C | G | C | A | C | C | T | A | A | C | A | T | #1279026 |
| BA.5.1 | T | A | A | T | A | T | T | C | C | T | A | A | C | A | T |  |
| mut |  |  |  |  |  |  |  |  |  |  |  |  |  |  |  |  |

|  |  |  |  |  |  |  |  |  |  |  |  |  |  |  |  |  |  |  |  |  |  |  |  |  |  |  |  |  |  |  |  |  |
| --- | --- | --- | --- | --- | --- | --- | --- | --- | --- | --- | --- | --- | --- | --- | --- | --- | --- | --- | --- | --- | --- | --- | --- | --- | --- | --- | --- | --- | --- | --- | --- | --- |
| pos | 44 | 4343 | 4936 | 9866 | 10051 | 12160 | 20451 | 20937 | 21765 | 21766 | 21767 | 21768 | 21769 | 21770 | 22304 | 22792 | 22917 | 23018 | 23040 | 26529 | 26858 | 27259 | 27382 | 27383 | 27384 | 27438 | 27889 | 29167 | 29386 | 29666 | 29754 |  |
| ref | C | A | G | C | G | G | C | G | T | A | C | A | T | G | T | C | T | T | A | G | C | A | G | A | T | T | C | C | C | C | C |  |
| BA.2 | C | G | A | T | A | G | C | T | T | A | C | A | T | G | C | T | T | T | G | G | T | C | C | T | C | T | C | C | C | C | C |  |
| BA.5.1 | C | G | A | T | A | G | C | G | - | - | - | - | - | - | T | C | G | G | A | A | C | A | G | A | T | C | T | T | T | T | T | #1279042 |
| BA.5.1 | T | A | G | C | G | A | T | G | - | - | - | - | - | - | T | C | G | G | A | A | C | A | G | A | T | C | T | C | C | T | T |  |
| mut |  |  |  |  |  |  |  |  |  |  |  |  |  |  |  |  |  |  |  |  |  |  |  |  |  |  |  | C>T | C>T |  |  |  |

|  |  |  |  |  |  |  |  |  |  |  |  |  |  |  |  |  |  |  |  |  |  |  |  |  |  |  |  |  |
| --- | --- | --- | --- | --- | --- | --- | --- | --- | --- | --- | --- | --- | --- | --- | --- | --- | --- | --- | --- | --- | --- | --- | --- | --- | --- | --- | --- | --- |
| pos | 44 | 1931 | 2954 | 10376 | 11750 | 12160 | 16935 | 19955 | 20055 | 21765 | 21766 | 21767 | 21768 | 21769 | 21770 | 22917 | 23018 | 23040 | 26529 | 26858 | 27259 | 27382 | 27383 | 27384 | 27889 | 28681 | 29754 |  |
| ref | C | C | T | C | C | G | G | C | A | T | A | C | A | T | G | T | T | A | G | C | A | G | A | T | C | G | C |  |
| BE.1.1 | T | A | C | T | T | A | A | T | G | - | - | - | - | - | - | G | G | A | A | C | A | G | A | T | T | T | T |  |
| BA.4 | T | A | C | T | T | A | A | T | G | - | - | - | - | - | - | G | G | A | G | T | C | C | T | C | C | G | C | #1279995 |
| BA.2 | C | C | T | C | C | G | G | C | A | T | A | C | A | T | G | T | T | G | G | T | C | C | T | C | C | G | C |  |
| mut |  |  |  |  |  |  |  |  |  |  |  |  |  |  |  |  |  |  |  |  |  |  |  |  |  |  |  |  |

|  |  |  |  |  |  |  |  |  |  |  |  |  |  |  |  |  |  |  |  |  |  |  |  |  |  |  |  |  |  |  |
| --- | --- | --- | --- | --- | --- | --- | --- | --- | --- | --- | --- | --- | --- | --- | --- | --- | --- | --- | --- | --- | --- | --- | --- | --- | --- | --- | --- | --- | --- | --- |
| pos | 44 | 686 | 687 | 688 | 689 | 690 | 691 | 692 | 693 | 694 | 958 | 1627 | 3232 | 5826 | 6979 | 21570 | 22110 | 26529 | 26858 | 27038 | 27259 | 27382 | 27383 | 27384 | 27788 | 27889 | 28330 | 28724 | 29754 |  |
| ref | C | A | A | G | T | C | A | T | T | T | C | C | T | C | T | T | A | G | C | A | A | G | A | T | G | C | A | C | C |  |
| BA.5.2.1 | C | G | C | G | C | C | G | C | T | A | C | T | T | T | G | T | T | A | C | G | A | G | A | T | G | T | G | C | T |  |
| BA.4.1 | C | G | C | G | C | C | G | C | T | A | T | C | C | C | T | G | A | G | T | A | C | C | T | C | T | C | A | T | C | #1280342 |
| BA.4.1 | T | - | - | - | - | - | - | - | - | - | T | C | T | C | T | G | A | G | T | A | C | C | T | C | T | C | A | T | C |  |
| mut |  |  |  |  |  |  |  |  |  |  |  |  | T>C |  |  |  |  |  |  |  |  |  |  |  |  |  |  |  |  |  |

|  |  |  |  |  |  |  |  |  |  |  |  |  |  |  |  |  |  |
| --- | --- | --- | --- | --- | --- | --- | --- | --- | --- | --- | --- | --- | --- | --- | --- | --- | --- |
| pos | 1559 | 1931 | 3875 | 4763 | 12068 | 16935 | 17278 | 22792 | 23323 | 25534 | 25537 | 27438 | 28681 | 28877 | 28878 | 29666 |  |
| ref | G | C | G | C | C | G | G | C | T | G | G | T | G | A | G | C |  |
| BA.5.1.24 | T | C | G | T | C | G | T | C | T | G | G | C | G | T | C | T |  |
| BE.1 | T | C | G | T | C | G | T | T | C | T | T | T | T | A | G | C | #1281614 |
| BE.1 | G | A | A | C | T | A | G | T | C | T | T | T | T | A | G | C |  |
| mut |  |  |  |  |  |  |  | C>T |  |  |  |  |  |  |  |  |  |

|  |  |  |  |  |  |  |  |  |  |  |  |  |  |  |  |  |  |  |  |  |  |  |  |
| --- | --- | --- | --- | --- | --- | --- | --- | --- | --- | --- | --- | --- | --- | --- | --- | --- | --- | --- | --- | --- | --- | --- | --- |
| pos | 44 | 809 | 1906 | 7276 | 8991 | 10518 | 11674 | 12025 | 13860 | 13921 | 15009 | 21159 | 21721 | 22118 | 22917 | 23673 | 24079 | 25207 | 25416 | 25810 | 26408 | 29006 |  |
| ref | C | T | T | T | C | A | C | C | C | G | T | G | C | T | T | C | T | C | C | C | C | A |  |
| BA.2.3.2 | C | C | C | T | T | A | C | T | T | A | T | G | C | C | T | C | C | C | C | T | T | G |  |
| BA.2.12.1 | C | C | C | T | T | G | T | C | C | G | C | T | T | T | A | T | T | T | T | C | C | G | #1281893 |
| BA.2.12.1 | T | T | T | A | C | G | T | C | C | G | C | T | T | T | A | T | T | T | T | C | C | A |  |
| mut |  |  |  |  |  |  |  |  |  |  |  |  |  |  |  |  |  |  |  |  |  | A>G |  |

|  |  |  |  |  |  |  |  |  |  |  |  |  |  |  |  |  |  |  |  |  |  |  |  |  |  |  |  |  |  |  |  |  |
| --- | --- | --- | --- | --- | --- | --- | --- | --- | --- | --- | --- | --- | --- | --- | --- | --- | --- | --- | --- | --- | --- | --- | --- | --- | --- | --- | --- | --- | --- | --- | --- | --- |
| pos | 686 | 687 | 688 | 689 | 690 | 691 | 692 | 693 | 694 | 6312 | 7764 | 9866 | 11674 | 12160 | 15009 | 21570 | 21721 | 21765 | 21766 | 21767 | 21768 | 21769 | 21770 | 22917 | 23018 | 23040 | 23673 | 25416 | 26768 | 27788 | 28724 |  |
| ref | A | A | G | T | C | A | T | T | T | C | C | C | C | G | T | T | C | T | A | C | A | T | G | T | T | A | C | C | C | G | C |  |
| BA.2.12.1 | A | A | G | T | C | A | T | T | T | A | C | T | T | G | C | T | T | T | A | C | A | T | G | A | T | G | T | T | T | G | C |  |
| BA.4 | A | A | G | T | C | A | T | T | T | A | C | T | T | G | C | T | T | T | A | C | A | T | G | G | G | A | C | C | C | T | T | #1282899 |
| BA.4.1 | - | - | - | - | - | - | - | - | - | C | T | C | C | A | T | G | C | - | - | - | - | - | - | G | G | A | C | C | C | T | T |  |
| mut |  |  |  |  |  |  |  |  |  |  |  |  |  |  |  |  |  |  |  |  |  |  |  |  |  |  |  |  |  |  |  |  |

|  |  |  |  |  |  |  |  |  |  |  |  |  |  |  |  |  |
| --- | --- | --- | --- | --- | --- | --- | --- | --- | --- | --- | --- | --- | --- | --- | --- | --- |
| pos | 219 | 2245 | 3393 | 6070 | 8618 | 8917 | 10615 | 19536 | 22792 | 22812 | 23248 | 27881 | 28098 | 28392 | 29073 |  |
| ref | G | C | C | C | G | C | C | C | C | A | C | C | T | G | C |  |
| BA.2.18 | T | T | T | C | A | C | C | C | C | C | T | C | C | G | T |  |
| BA.2 | T | T | T | C | A | C | C | T | T | A | C | T | T | T | C | #1283164 |
| BA.2 | G | C | C | T | G | T | T | T | C | A | C | C | T | G | C |  |
| mut |  |  |  |  |  |  |  |  | C>T |  |  | C>T |  | G>T |  |  |

|  |  |  |  |  |  |  |  |  |  |  |  |  |  |
| --- | --- | --- | --- | --- | --- | --- | --- | --- | --- | --- | --- | --- | --- |
| pos | 105 | 823 | 1627 | 1630 | 6979 | 11671 | 17750 | 18759 | 27038 | 27438 | 28330 | 29666 |  |
| ref | G | C | C | A | T | C | C | T | A | T | A | C |  |
| BA.5.2.1 | T | T | T | A | G | C | T | T | G | T | G | C |  |
| BA.5.1 | T | T | T | T | G | C | C | C | A | C | A | T | #1287188 |
| BA.5.1 | G | C | C | A | T | A | C | C | A | C | A | T |  |
| mut |  |  |  | A>T |  |  |  |  |  |  |  |  |  |

|  |  |  |  |  |  |  |  |  |  |  |  |  |  |  |  |  |  |  |  |  |  |  |  |  |  |  |  |  |  |  |  |  |  |  |  |  |  |  |  |  |  |  |
| --- | --- | --- | --- | --- | --- | --- | --- | --- | --- | --- | --- | --- | --- | --- | --- | --- | --- | --- | --- | --- | --- | --- | --- | --- | --- | --- | --- | --- | --- | --- | --- | --- | --- | --- | --- | --- | --- | --- | --- | --- | --- | --- |
| pos | 1078 | 1686 | 1843 | 9866 | 11522 | 11575 | 11674 | 11747 | 12160 | 14619 | 15009 | 16092 | 21721 | 21765 | 21766 | 21767 | 21768 | 21769 | 21770 | 21991 | 21992 | 21993 | 22917 | 23018 | 23040 | 23673 | 25416 | 25521 | 26529 | 26858 | 27259 | 27382 | 27383 | 27384 | 27438 | 27798 | 27889 | 27945 | 28471 | 29666 | 29754 |  |
| ref | A | C | G | C | T | C | C | C | G | T | T | C | C | T | A | C | A | T | G | T | T | A | T | T | A | C | C | C | G | C | A | G | A | T | T | G | C | C | C | C | C |  |
| BA.2.12.1 | A | C | G | T | G | T | T | T | G | C | C | T | T | T | A | C | A | T | G | T | T | A | A | T | G | T | T | T | G | T | C | C | T | C | T | T | C | T | C | C | C |  |
| BA.5.1 | A | T | G | T | G | T | T | T | G | C | C | T | C | - | - | - | - | - | - | T | T | A | G | G | A | C | C | C | A | C | A | G | A | T | C | G | T | C | T | T | T | #1288801 |
| BA.5.1 | T | C | T | C | T | C | C | C | A | T | T | C | C | - | - | - | - | - | - | - | - | - | G | G | A | C | C | C | A | C | A | G | A | T | C | G | T | C | T | T | T |  |
| mut |  | C>T |  |  |  |  |  |  |  |  |  |  |  |  |  |  |  |  |  | ->T | ->T | ->A |  |  |  |  |  |  |  |  |  |  |  |  |  |  |  |  |  |  |  |  |

|  |  |  |  |  |  |  |  |  |  |  |  |  |  |  |  |  |  |  |  |
| --- | --- | --- | --- | --- | --- | --- | --- | --- | --- | --- | --- | --- | --- | --- | --- | --- | --- | --- | --- |
| pos | 140 | 501 | 1627 | 1931 | 3875 | 7687 | 12068 | 16935 | 18084 | 22104 | 22792 | 23323 | 25534 | 25537 | 27038 | 28330 | 28681 | 28957 |  |
| ref | C | C | C | C | G | A | C | G | C | G | C | T | G | G | A | A | G | C |  |
| BE.1 | T | C | C | A | A | A | T | A | C | G | T | C | T | T | A | A | T | C |  |
| BF.6 | T | T | C | A | A | C | T | A | C | C | C | T | G | G | G | G | G | T | #1288968 |
| BF.6 | C | C | T | C | G | A | C | G | T | C | C | T | G | G | G | G | G | T |  |
| mut |  | C>T |  |  |  | A>C |  |  |  |  |  |  |  |  |  |  |  |  |  |

|  |  |  |  |  |  |  |  |  |  |  |  |  |  |  |  |  |  |  |  |  |  |  |  |  |  |  |  |  |  |  |  |  |
| --- | --- | --- | --- | --- | --- | --- | --- | --- | --- | --- | --- | --- | --- | --- | --- | --- | --- | --- | --- | --- | --- | --- | --- | --- | --- | --- | --- | --- | --- | --- | --- | --- |
| pos | 44 | 7345 | 8604 | 9866 | 10450 | 12160 | 12473 | 12792 | 16616 | 21765 | 21766 | 21767 | 21768 | 21769 | 21770 | 22917 | 23018 | 23040 | 25574 | 26529 | 26858 | 27259 | 27382 | 27383 | 27384 | 27438 | 27889 | 28432 | 28567 | 29666 | 29754 |  |
| ref | C | T | T | C | C | G | C | A | C | T | A | C | A | T | G | T | T | A | A | G | C | A | G | A | T | T | C | C | C | C | C |  |
| BA.2 | C | C | T | T | T | G | C | G | C | T | A | C | A | T | G | T | T | G | G | G | T | C | C | T | C | T | C | T | C | C | C |  |
| BA.5.1 | C | C | C | T | T | G | C | G | T | - | - | - | - | - | - | G | G | A | A | A | C | A | G | A | T | C | T | C | T | T | T | #1289393 |
| BA.5.1 | T | T | T | C | C | A | T | A | T | - | - | - | - | - | - | G | G | A | A | A | C | A | G | A | T | C | T | C | C | T | T |  |
| mut |  |  | T>C |  |  |  |  |  |  |  |  |  |  |  |  |  |  |  |  |  |  |  |  |  |  |  |  |  | C>T |  |  |  |

|  |  |  |  |  |  |  |  |  |  |  |  |  |  |  |  |  |
| --- | --- | --- | --- | --- | --- | --- | --- | --- | --- | --- | --- | --- | --- | --- | --- | --- |
| pos | 1820 | 3043 | 6472 | 6979 | 8174 | 9967 | 12784 | 15372 | 19524 | 24620 | 27038 | 27389 | 27532 | 28330 | 28402 |  |
| ref | G | T | C | T | G | C | C | G | C | G | A | C | C | A | A |  |
| BA.5 | A | C | C | T | A | C | T | C | C | G | A | T | C | A | A |  |
| BF.5 | A | C | A | T | A | C | T | C | C | T | G | C | T | G | G | #1291130 |
| BF.5 | G | T | C | G | G | T | C | G | T | T | G | C | T | G | G |  |
| mut |  |  | C>A |  |  |  |  |  |  |  |  |  |  |  |  |  |

|  |  |  |  |  |  |  |  |  |  |  |  |  |  |  |  |  |  |  |  |  |  |  |  |  |  |  |  |  |  |  |  |  |  |  |  |  |
| --- | --- | --- | --- | --- | --- | --- | --- | --- | --- | --- | --- | --- | --- | --- | --- | --- | --- | --- | --- | --- | --- | --- | --- | --- | --- | --- | --- | --- | --- | --- | --- | --- | --- | --- | --- | --- |
| pos | 1627 | 9866 | 10116 | 11665 | 12160 | 12310 | 13653 | 16616 | 19812 | 21765 | 21766 | 21767 | 21768 | 21769 | 21770 | 22304 | 22599 | 22917 | 23018 | 23040 | 24676 | 25416 | 26180 | 26529 | 26858 | 27012 | 27259 | 27382 | 27383 | 27384 | 27513 | 27532 | 27889 | 28330 | 29754 |  |
| ref | C | C | C | C | G | G | A | C | A | T | A | C | A | T | G | T | G | T | T | A | A | C | T | G | C | C | A | G | A | T | C | C | C | A | C |  |
| BA.2.76 | C | T | T | C | G | G | A | C | G | T | A | C | A | T | G | A | C | T | T | G | A | T | T | G | T | C | C | C | T | C | C | T | C | A | C |  |
| XBG | C | T | T | T | G | G | A | C | G | T | A | C | A | T | G | A | C | G | G | A | G | C | C | A | C | T | A | G | A | T | T | C | T | G | T | #1291970 |
| BA.5.2 | T | C | C | C | A | A | G | A | A | - | - | - | - | - | - | T | G | G | G | A | G | C | T | A | C | T | A | G | A | T | T | C | T | G | T |  |
| mut |  |  |  | C>T |  |  |  |  |  |  |  |  |  |  |  |  |  |  |  |  |  |  | T>C |  |  |  |  |  |  |  |  |  |  |  |  |  |

|  |  |  |  |  |  |  |  |  |  |  |  |  |  |  |  |  |  |  |  |  |  |  |  |  |  |  |  |  |  |
| --- | --- | --- | --- | --- | --- | --- | --- | --- | --- | --- | --- | --- | --- | --- | --- | --- | --- | --- | --- | --- | --- | --- | --- | --- | --- | --- | --- | --- | --- |
| pos | 686 | 687 | 688 | 689 | 690 | 691 | 692 | 693 | 694 | 1931 | 2445 | 2954 | 11750 | 11881 | 16935 | 20883 | 26529 | 26858 | 27259 | 27382 | 27383 | 27384 | 27788 | 27889 | 28208 | 28681 | 28724 | 29754 |  |
| ref | A | A | G | T | C | A | T | T | T | C | C | T | C | A | G | T | G | C | A | G | A | T | G | C | T | G | C | C |  |
| BE.1.1 | A | A | G | T | C | A | T | T | T | A | T | C | T | A | A | C | A | C | A | G | A | T | G | T | T | T | C | T |  |
| BA.4 | A | A | G | T | C | A | T | T | T | A | T | C | T | G | A | T | G | T | C | C | T | C | T | C | C | G | T | C | #1292896 |
| BA.4 | - | - | - | - | - | - | - | - | - | C | C | T | C | A | G | T | G | T | C | C | T | C | T | C | T | G | T | C |  |
| mut |  |  |  |  |  |  |  |  |  |  |  |  |  | A>G |  |  |  |  |  |  |  |  |  |  | T>C |  |  |  |  |

|  |  |  |  |  |  |  |  |  |  |  |  |  |  |  |  |  |  |  |  |  |  |  |  |  |  |
| --- | --- | --- | --- | --- | --- | --- | --- | --- | --- | --- | --- | --- | --- | --- | --- | --- | --- | --- | --- | --- | --- | --- | --- | --- | --- |
| pos | 2902 | 4906 | 6107 | 6638 | 7573 | 9866 | 11674 | 12160 | 12756 | 17402 | 21077 | 21570 | 21765 | 21766 | 21767 | 21768 | 21769 | 21770 | 22917 | 23018 | 23040 | 23673 | 25416 | 26529 |  |
| ref | C | T | A | C | C | C | C | G | C | C | C | T | T | A | C | A | T | G | T | T | A | C | C | G |  |
| BA.2.12.1 | T | T | G | C | T | T | T | G | C | C | C | T | T | A | C | A | T | G | A | T | G | T | T | G |  |
| BA.2 | T | T | G | T | T | T | T | G | A | C | C | T | - | - | - | - | - | - | G | G | A | C | C | A | #1293317 |
| BA.4 | C | C | A | C | C | C | C | A | C | T | T | G | - | - | - | - | - | - | G | G | A | C | C | A |  |
| mut |  |  |  | C>T |  |  |  |  | C>A |  |  |  |  |  |  |  |  |  |  |  |  |  |  |  |  |

|  |  |  |  |  |  |  |  |  |  |  |  |  |  |  |  |  |  |  |  |  |  |  |  |  |  |  |  |  |  |  |  |
| --- | --- | --- | --- | --- | --- | --- | --- | --- | --- | --- | --- | --- | --- | --- | --- | --- | --- | --- | --- | --- | --- | --- | --- | --- | --- | --- | --- | --- | --- | --- | --- |
| pos | 44 | 9866 | 12160 | 13945 | 21765 | 21766 | 21767 | 21768 | 21769 | 21770 | 21789 | 22792 | 22917 | 23018 | 23040 | 23948 | 24423 | 26529 | 26858 | 27259 | 27382 | 27383 | 27384 | 27889 | 28417 | 29728 | 29754 | 29757 | 29758 | 29759 |  |
| ref | C | C | G | C | T | A | C | A | T | G | C | C | T | T | A | G | A | G | C | A | G | A | T | C | T | T | C | G | T | G |  |
| BA.5.5 | T | C | A | C | - | - | - | - | - | - | T | C | G | G | A | T | G | A | C | A | G | A | T | T | C | T | T | G | T | G |  |
| BA.5.5 | T | C | A | C | - | - | - | - | - | - | T | C | G | G | A | G | G | A | C | A | G | A | T | T | C | C | C | A | C | C | #1294408 |
| BA.2 | C | T | G | T | T | A | C | A | T | G | C | T | T | T | G | T | A | G | T | C | C | T | C | C | T | C | C | A | C | C |  |
| mut |  |  |  |  |  |  |  |  |  |  |  |  |  |  |  | T>G |  |  |  |  |  |  |  |  |  |  |  |  |  |  |  |

|  |  |  |  |  |  |  |  |  |  |  |  |  |  |  |  |  |  |  |  |  |  |  |  |  |  |  |  |  |  |  |  |  |  |  |  |
| --- | --- | --- | --- | --- | --- | --- | --- | --- | --- | --- | --- | --- | --- | --- | --- | --- | --- | --- | --- | --- | --- | --- | --- | --- | --- | --- | --- | --- | --- | --- | --- | --- | --- | --- | --- |
| pos | 44 | 4230 | 4576 | 4579 | 4906 | 9866 | 12160 | 13366 | 15972 | 16376 | 16779 | 17402 | 18957 | 18984 | 21077 | 21765 | 21766 | 21767 | 21768 | 21769 | 21770 | 22305 | 22917 | 23018 | 23040 | 25481 | 26529 | 26858 | 27259 | 27382 | 27383 | 27384 | 27889 | 29754 |  |
| ref | C | C | A | T | T | C | G | C | A | C | C | C | T | G | C | T | A | C | A | T | G | A | T | T | A | G | G | C | A | G | A | T | C | C |  |
| BA.5.6 | T | C | A | T | C | C | A | C | A | C | C | T | C | T | T | - | - | - | - | - | - | A | G | G | A | G | A | C | A | G | A | T | T | T |  |
| BA.2 | T | C | A | T | C | C | A | C | A | C | C | T | C | T | C | T | A | C | A | T | G | C | T | T | G | A | G | T | C | C | T | C | C | C | #1294443 |
| BA.2.48 | C | T | T | A | T | T | G | T | G | T | T | C | T | G | C | T | A | C | A | T | G | C | T | T | G | A | G | T | C | C | T | C | C | C |  |
| mut |  |  |  |  |  |  |  |  |  |  |  |  |  |  |  |  |  |  |  |  |  |  |  |  |  |  |  |  |  |  |  |  |  |  |  |

|  |  |  |  |  |  |  |  |  |  |  |  |  |  |  |  |  |  |
| --- | --- | --- | --- | --- | --- | --- | --- | --- | --- | --- | --- | --- | --- | --- | --- | --- | --- |
| pos | 105 | 823 | 1627 | 6979 | 8956 | 10183 | 13611 | 17750 | 21232 | 25855 | 27038 | 27438 | 28330 | 28531 | 29518 | 29666 |  |
| ref | G | C | C | T | C | C | A | C | G | G | A | T | A | C | C | C |  |
| BA.5.2.1 | T | T | T | G | T | A | A | T | G | G | G | T | G | C | C | C |  |
| BA.5.1 | T | T | T | G | T | A | A | C | A | T | A | C | A | T | T | T | #1295777 |
| BA.5.1 | G | C | C | T | C | C | G | C | A | T | A | C | A | T | T | T |  |
| mut |  |  |  |  |  |  |  |  |  |  |  |  |  |  |  |  |  |

|  |  |  |  |  |  |  |  |  |  |  |  |  |  |  |  |  |  |  |  |  |  |  |  |  |  |  |  |  |  |  |  |
| --- | --- | --- | --- | --- | --- | --- | --- | --- | --- | --- | --- | --- | --- | --- | --- | --- | --- | --- | --- | --- | --- | --- | --- | --- | --- | --- | --- | --- | --- | --- | --- |
| pos | 6255 | 6684 | 9866 | 11674 | 12160 | 13944 | 15009 | 17634 | 21721 | 21765 | 21766 | 21767 | 21768 | 21769 | 21770 | 22917 | 23018 | 23040 | 23673 | 25416 | 26529 | 26858 | 27259 | 27382 | 27383 | 27384 | 27438 | 27889 | 29666 | 29754 |  |
| ref | C | A | C | C | G | C | T | C | C | T | A | C | A | T | G | T | T | A | C | C | G | C | A | G | A | T | T | C | C | C |  |
| BA.2.12.1 | T | G | T | T | G | C | C | C | T | T | A | C | A | T | G | A | T | G | T | T | G | T | C | C | T | C | T | C | C | C |  |
| BA.5.1 | T | G | T | T | G | T | C | C | C | - | - | - | - | - | - | G | G | A | C | C | A | C | A | G | A | T | C | T | T | T | #1296795 |
| BA.5.1 | C | A | C | C | A | C | T | T | C | - | - | - | - | - | - | G | G | A | C | C | A | C | A | G | A | T | C | T | T | T |  |
| mut |  |  |  |  |  | C>T |  |  |  |  |  |  |  |  |  |  |  |  |  |  |  |  |  |  |  |  |  |  |  |  |  |

|  |  |  |  |  |  |  |  |  |  |  |  |  |  |  |  |  |  |  |  |  |  |  |  |  |  |  |  |  |  |  |  |  |
| --- | --- | --- | --- | --- | --- | --- | --- | --- | --- | --- | --- | --- | --- | --- | --- | --- | --- | --- | --- | --- | --- | --- | --- | --- | --- | --- | --- | --- | --- | --- | --- | --- |
| pos | 583 | 686 | 687 | 688 | 689 | 690 | 691 | 692 | 693 | 694 | 1513 | 8140 | 10969 | 13426 | 14599 | 18555 | 22599 | 23535 | 26529 | 26858 | 27259 | 27382 | 27383 | 27384 | 27438 | 27788 | 27889 | 28697 | 28724 | 29666 | 29754 |  |
| ref | C | A | A | G | T | C | A | T | T | T | C | C | C | C | C | C | G | A | G | C | A | G | A | T | T | G | C | C | C | C | C |  |
| BA.4.6 | C | - | - | - | - | - | - | - | - | - | C | T | T | T | T | C | C | G | G | T | C | C | T | C | T | T | C | C | T | C | C |  |
| BA.5.1 | C | - | - | - | - | - | - | - | - | - | C | T | T | T | T | T | C | G | A | C | A | G | A | T | C | G | T | T | C | T | T | #1297571 |
| BA.5.1 | T | A | A | G | T | C | A | T | T | T | T | C | C | C | C | C | G | A | A | C | A | G | A | T | C | G | T | T | C | T | T |  |
| mut |  |  |  |  |  |  |  |  |  |  |  |  |  |  |  | C>T |  |  |  |  |  |  |  |  |  |  |  |  |  |  |  |  |

|  |  |  |  |  |  |  |  |  |  |  |  |  |  |  |  |  |  |  |  |  |  |  |  |  |  |  |  |  |  |  |  |
| --- | --- | --- | --- | --- | --- | --- | --- | --- | --- | --- | --- | --- | --- | --- | --- | --- | --- | --- | --- | --- | --- | --- | --- | --- | --- | --- | --- | --- | --- | --- | --- |
| pos | 686 | 687 | 688 | 689 | 690 | 691 | 692 | 693 | 694 | 2110 | 3685 | 3686 | 4906 | 7069 | 14925 | 17402 | 18957 | 21077 | 21570 | 23948 | 26529 | 26858 | 27259 | 27382 | 27383 | 27384 | 27788 | 27889 | 28724 | 29754 |  |
| ref | A | A | G | T | C | A | T | T | T | C | G | C | T | A | C | C | T | C | T | G | G | C | A | G | A | T | G | C | C | C |  |
| BA.5.6 | A | A | G | T | C | A | T | T | T | C | A | A | C | A | T | T | C | T | T | T | A | C | A | G | A | T | G | T | C | T |  |
| BA.4 | A | A | G | T | C | A | T | T | T | C | A | A | C | A | T | T | C | T | T | T | G | T | C | C | T | C | T | C | T | C | #1298543 |
| BA.4.1 | - | - | - | - | - | - | - | - | - | T | G | C | T | G | C | C | T | C | G | G | G | T | C | C | T | C | T | C | T | C |  |
| mut |  |  |  |  |  |  |  |  |  |  |  |  |  |  |  |  |  |  |  |  |  |  |  |  |  |  |  |  |  |  |  |

|  |  |  |  |  |  |  |  |  |  |  |  |  |  |  |  |  |  |  |  |  |  |  |  |  |  |  |  |  |  |  |  |  |  |
| --- | --- | --- | --- | --- | --- | --- | --- | --- | --- | --- | --- | --- | --- | --- | --- | --- | --- | --- | --- | --- | --- | --- | --- | --- | --- | --- | --- | --- | --- | --- | --- | --- | --- |
| pos | 934 | 5371 | 5512 | 9424 | 9866 | 11674 | 12160 | 15009 | 16852 | 21721 | 21765 | 21766 | 21767 | 21768 | 21769 | 21770 | 21789 | 22917 | 23018 | 23040 | 23673 | 25416 | 26529 | 26858 | 27259 | 27382 | 27383 | 27384 | 27552 | 27889 | 28603 | 29754 |  |
| ref | C | A | C | A | C | C | G | T | G | C | T | A | C | A | T | G | C | T | T | A | C | C | G | C | A | G | A | T | A | C | C | C |  |
| BA.2.12.1 | C | G | C | T | T | T | G | C | G | T | T | A | C | A | T | G | C | A | T | G | T | T | G | T | C | C | T | C | A | C | T | C |  |
| BA.5.5 | T | G | C | T | T | T | A | T | T | C | - | - | - | - | - | - | T | G | G | A | C | C | A | C | A | G | A | T | G | T | C | T | #1301051 |
| BA.5.5 | C | A | T | G | C | C | A | T | T | C | - | - | - | - | - | - | T | G | G | A | C | C | A | C | A | G | A | T | G | T | C | T |  |
| mut | C>T |  |  |  |  |  |  |  |  |  |  |  |  |  |  |  |  |  |  |  |  |  |  |  |  |  |  |  |  |  |  |  |  |

|  |  |  |  |  |  |  |  |  |  |  |  |  |  |  |  |  |  |  |  |  |  |  |  |  |  |  |  |  |  |  |  |  |  |  |  |  |  |  |  |  |  |
| --- | --- | --- | --- | --- | --- | --- | --- | --- | --- | --- | --- | --- | --- | --- | --- | --- | --- | --- | --- | --- | --- | --- | --- | --- | --- | --- | --- | --- | --- | --- | --- | --- | --- | --- | --- | --- | --- | --- | --- | --- | --- |
| pos | 1627 | 8290 | 9866 | 11522 | 11575 | 11674 | 12160 | 12188 | 14652 | 15009 | 16092 | 21643 | 21721 | 21765 | 21766 | 21767 | 21768 | 21769 | 21770 | 22786 | 22917 | 23018 | 23040 | 23673 | 25416 | 25521 | 26529 | 26858 | 27038 | 27259 | 27382 | 27383 | 27384 | 27889 | 27945 | 28254 | 28330 | 28370 | 28371 | 29754 |  |
| ref | C | C | C | T | C | C | G | G | T | T | C | A | C | T | A | C | A | T | G | A | T | T | A | C | C | C | G | C | A | A | G | A | T | C | C | A | A | A | G | C |  |
| BA.2.12.1 | C | C | T | G | T | T | G | G | T | C | T | A | T | T | A | C | A | T | G | C | A | T | G | T | T | T | G | T | A | C | C | T | C | C | T | - | A | - | G | C |  |
| BA.2.12.1 | C | T | T | G | T | T | G | G | T | C | T | A | T | T | A | C | A | T | G | A | A | T | G | T | C | C | A | C | G | A | G | A | T | T | C | A | G | T | T | T | #1301257 |
| BA.5.2.1 | T | C | C | T | C | C | A | T | C | T | C | T | C | - | - | - | - | - | - | C | G | G | A | C | C | C | A | C | G | A | G | A | T | T | C | A | G | T | T | T |  |
| mut |  | C>T |  |  |  |  |  |  |  |  |  |  |  |  |  |  |  |  |  | C>A |  |  |  |  |  |  |  |  |  |  |  |  |  |  |  |  |  |  |  |  |  |

|  |  |  |  |  |  |  |  |  |  |  |  |  |  |  |  |  |  |  |  |  |  |  |  |  |  |  |  |  |  |
| --- | --- | --- | --- | --- | --- | --- | --- | --- | --- | --- | --- | --- | --- | --- | --- | --- | --- | --- | --- | --- | --- | --- | --- | --- | --- | --- | --- | --- | --- |
| pos | 44 | 823 | 1627 | 6979 | 9286 | 9866 | 12160 | 21765 | 21766 | 21767 | 21768 | 21769 | 21770 | 22812 | 22917 | 23018 | 23040 | 23248 | 26529 | 26858 | 27038 | 27259 | 27382 | 27383 | 27384 | 27889 | 28330 | 29754 |  |
| ref | C | C | C | T | C | C | G | T | A | C | A | T | G | A | T | T | A | C | G | C | A | A | G | A | T | C | A | C |  |
| BA.2.18 | C | C | C | T | C | T | G | T | A | C | A | T | G | C | T | T | G | T | G | T | A | C | C | T | C | C | A | C |  |
| BA.5 | C | T | C | T | T | T | G | T | A | C | A | T | G | C | T | T | G | T | A | C | G | A | G | A | T | T | G | T | #1301898 |
| BA.5.2.1 | T | C | T | G | C | C | A | - | - | - | - | - | - | A | G | G | A | C | A | C | G | A | G | A | T | T | G | T |  |
| mut |  | C>T |  |  | C>T |  |  |  |  |  |  |  |  |  |  |  |  |  |  |  |  |  |  |  |  |  |  |  |  |

|  |  |  |  |  |  |  |  |  |  |  |  |  |  |  |  |  |  |  |  |  |  |  |  |  |  |  |  |  |  |  |
| --- | --- | --- | --- | --- | --- | --- | --- | --- | --- | --- | --- | --- | --- | --- | --- | --- | --- | --- | --- | --- | --- | --- | --- | --- | --- | --- | --- | --- | --- | --- |
| pos | 155 | 686 | 687 | 688 | 689 | 690 | 691 | 692 | 693 | 694 | 1627 | 4780 | 5284 | 12310 | 16616 | 21570 | 22786 | 26529 | 26858 | 27259 | 27382 | 27383 | 27384 | 27788 | 27889 | 28297 | 28330 | 28724 | 29754 |  |
| ref | G | A | A | G | T | C | A | T | T | T | C | C | C | G | C | T | A | G | C | A | G | A | T | G | C | T | A | C | C |  |
| BA.5.2.1 | G | A | A | G | T | C | A | T | T | T | T | C | C | A | A | T | C | A | C | A | G | A | T | G | T | T | G | C | T |  |
| BA.5.2 | G | A | A | G | T | C | A | T | T | T | T | T | T | A | A | T | A | A | C | A | G | A | T | T | C | C | A | T | C | #1302531 |
| BA.4.1 | A | - | - | - | - | - | - | - | - | - | C | C | C | G | C | G | C | G | T | C | C | T | C | T | C | C | A | T | C |  |
| mut |  |  |  |  |  |  |  |  |  |  |  | C>T | C>T |  |  |  | C>A |  |  |  |  |  |  |  |  |  |  |  |  |  |

|  |  |  |  |  |  |  |  |  |  |  |  |  |  |  |  |  |  |  |  |  |  |  |  |  |  |  |  |  |  |  |  |  |  |  |  |  |  |  |  |
| --- | --- | --- | --- | --- | --- | --- | --- | --- | --- | --- | --- | --- | --- | --- | --- | --- | --- | --- | --- | --- | --- | --- | --- | --- | --- | --- | --- | --- | --- | --- | --- | --- | --- | --- | --- | --- | --- | --- | --- |
| pos | 140 | 1588 | 9866 | 11494 | 11573 | 11674 | 12160 | 12439 | 15009 | 15597 | 20016 | 21721 | 21765 | 21766 | 21767 | 21768 | 21769 | 21770 | 22054 | 22917 | 23018 | 23040 | 23673 | 25002 | 25416 | 26526 | 26529 | 26858 | 27218 | 27259 | 27382 | 27383 | 27384 | 27438 | 27879 | 27889 | 29666 | 29754 |  |
| ref | C | A | C | C | T | C | G | C | T | T | C | C | T | A | C | A | T | G | T | T | T | A | C | C | C | G | G | C | A | A | G | A | T | T | C | C | C | C |  |
| BA.2.12.1 | T | G | T | C | C | T | G | T | C | C | T | T | T | A | C | A | T | G | G | A | T | G | T | C | T | A | G | T | A | C | C | T | C | T | C | C | C | C |  |
| BA.5.1.30 | T | G | T | C | C | T | G | T | C | C | C | C | - | - | - | - | - | - | T | G | G | A | C | T | C | G | A | C | G | A | G | A | T | C | T | T | T | T | #1303463 |
| BA.5.1.25 | C | A | C | T | T | C | A | C | T | T | C | C | - | - | - | - | - | - | T | G | G | A | C | C | C | G | A | C | G | A | G | A | T | C | T | T | T | T |  |
| mut |  |  |  |  |  |  |  |  |  |  |  |  |  |  |  |  |  |  |  |  |  |  |  | C>T |  |  |  |  |  |  |  |  |  |  |  |  |  |  |  |

|  |  |  |  |  |  |  |  |  |  |  |  |  |  |  |  |  |  |  |  |  |  |  |  |  |  |  |  |  |  |  |
| --- | --- | --- | --- | --- | --- | --- | --- | --- | --- | --- | --- | --- | --- | --- | --- | --- | --- | --- | --- | --- | --- | --- | --- | --- | --- | --- | --- | --- | --- | --- |
| pos | 686 | 687 | 688 | 689 | 690 | 691 | 692 | 693 | 694 | 1627 | 6979 | 8140 | 12911 | 14599 | 18682 | 22599 | 23535 | 26529 | 26858 | 27038 | 27259 | 27382 | 27383 | 27384 | 27788 | 27889 | 28330 | 28724 | 29754 |  |
| ref | A | A | G | T | C | A | T | T | T | C | T | C | G | C | A | G | A | G | C | A | A | G | A | T | G | C | A | C | C |  |
| BA.4.6 | - | - | - | - | - | - | - | - | - | C | T | T | G | T | A | C | G | G | T | A | C | C | T | C | T | C | A | T | C |  |
| BA.5 | - | - | - | - | - | - | - | - | - | C | T | T | G | T | G | G | A | A | C | G | A | G | A | T | G | T | G | C | T | #1305793 |
| BA.5.2.1 | A | A | G | T | C | A | T | T | T | T | G | C | A | C | G | G | A | A | C | G | A | G | A | T | G | T | G | C | T |  |
| mut |  |  |  |  |  |  |  |  |  |  |  |  |  |  |  |  |  |  |  |  |  |  |  |  |  |  |  |  |  |  |

|  |  |  |  |  |  |  |  |  |  |  |  |  |  |  |  |
| --- | --- | --- | --- | --- | --- | --- | --- | --- | --- | --- | --- | --- | --- | --- | --- |
| pos | 1625 | 1627 | 1931 | 4644 | 16935 | 17750 | 19239 | 22021 | 25826 | 27038 | 27689 | 28330 | 28681 | 29358 |  |
| ref | C | C | C | A | G | C | A | G | A | A | C | A | G | C |  |
| BE.1 | T | C | A | C | A | T | G | T | A | A | C | A | T | C |  |
| BE.1 | T | C | A | C | A | T | G | T | A | G | T | G | G | C | #1307218 |
| BF.8 | C | T | C | A | G | C | A | G | T | G | T | G | G | A |  |
| mut |  |  |  |  |  |  |  |  |  |  |  |  |  | A>C |  |

|  |  |  |  |  |  |  |  |  |  |  |  |  |  |  |  |  |  |  |  |  |  |  |  |  |  |  |  |  |  |  |  |  |  |  |  |  |  |  |  |  |  |  |  |
| --- | --- | --- | --- | --- | --- | --- | --- | --- | --- | --- | --- | --- | --- | --- | --- | --- | --- | --- | --- | --- | --- | --- | --- | --- | --- | --- | --- | --- | --- | --- | --- | --- | --- | --- | --- | --- | --- | --- | --- | --- | --- | --- | --- |
| pos | 203 | 1627 | 4993 | 7299 | 8150 | 9866 | 9907 | 10156 | 11522 | 11575 | 11674 | 12160 | 15009 | 15181 | 16092 | 21721 | 21765 | 21766 | 21767 | 21768 | 21769 | 21770 | 22917 | 23018 | 23040 | 23673 | 25416 | 25521 | 26529 | 26810 | 26858 | 27259 | 27382 | 27383 | 27384 | 27798 | 27889 | 27945 | 28330 | 28657 | 28677 | 29754 |  |
| ref | C | C | C | C | G | C | C | C | T | C | C | G | T | G | C | C | T | A | C | A | T | G | T | T | A | C | C | C | G | C | C | A | G | A | T | G | C | C | A | C | C | C |  |
| BA.2.12.1 | T | C | T | C | A | T | T | T | G | T | T | G | C | G | T | T | T | A | C | A | T | G | A | T | G | T | T | T | G | C | T | C | C | T | C | T | C | T | A | A | T | C |  |
| BA.2.12.1 | T | C | T | T | A | T | T | T | G | T | T | G | C | A | T | T | T | A | C | A | T | G | A | T | G | T | T | T | G | T | T | A | G | A | T | G | T | C | G | C | C | T | #1309135 |
| BA.5.2 | C | T | C | C | G | C | C | C | T | C | C | A | T | G | C | C | - | - | - | - | - | - | G | G | A | C | C | C | A | C | C | A | G | A | T | G | T | C | G | C | C | T |  |
| mut |  |  |  | C>T |  |  |  |  |  |  |  |  |  | G>A |  |  |  |  |  |  |  |  |  |  |  |  |  |  |  | C>T |  |  |  |  |  |  |  |  |  |  |  |  |  |

|  |  |  |  |  |  |  |  |  |  |  |  |  |  |  |  |  |  |  |  |  |  |  |  |  |  |  |  |  |  |  |  |  |
| --- | --- | --- | --- | --- | --- | --- | --- | --- | --- | --- | --- | --- | --- | --- | --- | --- | --- | --- | --- | --- | --- | --- | --- | --- | --- | --- | --- | --- | --- | --- | --- | --- |
| pos | 44 | 3478 | 3539 | 6117 | 9679 | 9866 | 12160 | 16376 | 19647 | 21765 | 21766 | 21767 | 21768 | 21769 | 21770 | 22792 | 22917 | 23018 | 23040 | 26529 | 26858 | 27259 | 27382 | 27383 | 27384 | 27438 | 27630 | 27656 | 27889 | 29666 | 29754 |  |
| ref | C | T | G | C | C | C | G | C | T | T | A | C | A | T | G | C | T | T | A | G | C | A | G | A | T | T | C | T | C | C | C |  |
| BA.5.1 | T | C | T | C | T | C | A | T | C | - | - | - | - | - | - | C | G | G | A | A | C | A | G | A | T | C | T | C | T | T | T |  |
| BA.5 | T | C | T | C | T | C | A | T | C | - | - | - | - | - | - | C | G | G | A | A | C | A | G | A | T | T | C | T | C | C | C | #1312843 |
| BA.2 | C | T | G | T | C | T | G | C | T | T | A | C | A | T | G | T | T | T | G | G | T | C | C | T | C | T | C | T | C | C | C |  |
| mut |  |  |  |  |  |  |  |  |  |  |  |  |  |  |  |  |  |  |  |  |  |  |  |  |  |  |  |  |  |  |  |  |

|  |  |  |  |  |  |  |  |  |  |  |  |  |  |  |  |  |  |  |  |  |  |  |  |  |  |  |  |  |  |  |  |  |  |  |  |  |
| --- | --- | --- | --- | --- | --- | --- | --- | --- | --- | --- | --- | --- | --- | --- | --- | --- | --- | --- | --- | --- | --- | --- | --- | --- | --- | --- | --- | --- | --- | --- | --- | --- | --- | --- | --- | --- |
| pos | 686 | 687 | 688 | 689 | 690 | 691 | 692 | 693 | 694 | 2485 | 4109 | 5512 | 5777 | 9262 | 9430 | 9968 | 16692 | 18634 | 18981 | 21570 | 21789 | 25552 | 26529 | 26858 | 27259 | 27382 | 27383 | 27384 | 27497 | 27672 | 27788 | 27889 | 28077 | 28724 | 29754 |  |
| ref | A | A | G | T | C | A | T | T | T | C | G | C | C | T | C | G | T | G | C | T | C | G | G | C | A | G | A | T | G | T | G | C | G | C | C |  |
| BA.4.1 | - | - | - | - | - | - | - | - | - | C | G | C | T | T | T | G | T | G | T | G | C | G | G | T | C | C | T | C | G | T | T | C | G | T | C |  |
| BA.5 | - | - | - | - | - | - | - | - | - | T | G | C | T | T | T | A | T | G | T | G | C | A | A | C | A | G | A | T | C | C | G | T | T | C | T | #1313481 |
| BA.5.5 | A | A | G | T | C | A | T | T | T | C | T | T | C | C | C | G | C | T | C | T | T | A | A | C | A | G | A | T | C | C | G | T | T | C | T |  |
| mut |  |  |  |  |  |  |  |  |  | C>T |  |  |  |  |  | G>A |  |  |  |  |  |  |  |  |  |  |  |  |  |  |  |  |  |  |  |  |

|  |  |  |  |  |  |  |  |  |  |  |  |  |  |  |  |  |  |  |  |  |  |  |  |  |  |  |  |  |  |  |  |  |  |  |  |
| --- | --- | --- | --- | --- | --- | --- | --- | --- | --- | --- | --- | --- | --- | --- | --- | --- | --- | --- | --- | --- | --- | --- | --- | --- | --- | --- | --- | --- | --- | --- | --- | --- | --- | --- | --- |
| pos | 44 | 670 | 1627 | 2062 | 5241 | 9866 | 10060 | 12160 | 13945 | 21765 | 21766 | 21767 | 21768 | 21769 | 21770 | 22792 | 22917 | 23018 | 23040 | 26529 | 26858 | 27038 | 27259 | 27382 | 27383 | 27384 | 27889 | 27916 | 28330 | 29728 | 29754 | 29757 | 29758 | 29759 |  |
| ref | C | T | C | C | C | C | T | G | C | T | A | C | A | T | G | C | T | T | A | G | C | A | A | G | A | T | C | G | A | T | C | G | T | G |  |
| BA.5.2.1 | T | T | T | C | C | C | C | A | C | - | - | - | - | - | - | C | G | G | A | A | C | G | A | G | A | T | T | G | G | T | T | G | T | G |  |
| BA.5.2.1 | T | T | T | T | T | C | C | A | C | - | - | - | - | - | - | C | G | G | A | A | C | G | A | G | A | T | T | A | G | C | C | A | C | C | #1315204 |
| BA.2 | C | G | C | C | C | T | T | G | T | T | A | C | A | T | G | T | T | T | G | G | T | A | C | C | T | C | C | G | A | C | C | A | C | C |  |
| mut |  |  |  | C>T | C>T |  |  |  |  |  |  |  |  |  |  |  |  |  |  |  |  |  |  |  |  |  |  | G>A |  |  |  |  |  |  |  |

|  |  |  |  |  |  |  |  |  |  |  |  |  |  |  |  |  |  |  |  |  |  |  |  |  |  |  |  |  |  |  |  |  |  |  |
| --- | --- | --- | --- | --- | --- | --- | --- | --- | --- | --- | --- | --- | --- | --- | --- | --- | --- | --- | --- | --- | --- | --- | --- | --- | --- | --- | --- | --- | --- | --- | --- | --- | --- | --- |
| pos | 1942 | 6446 | 9866 | 11674 | 12160 | 15009 | 20935 | 21721 | 21765 | 21766 | 21767 | 21768 | 21769 | 21770 | 22323 | 22917 | 23018 | 23040 | 23673 | 25416 | 26529 | 26858 | 27259 | 27382 | 27383 | 27384 | 27438 | 27889 | 28254 | 28877 | 28878 | 29666 | 29754 |  |
| ref | G | G | C | C | G | T | A | C | T | A | C | A | T | G | C | T | T | A | C | C | G | C | A | G | A | T | T | C | A | A | G | C | C |  |
| BA.2.12.1 | G | A | T | T | G | C | A | T | T | A | C | A | T | G | C | A | T | G | T | T | G | T | C | C | T | C | T | C | A | T | C | C | C |  |
| BA.5.1 | T | A | T | T | G | C | G | T | T | A | C | A | T | G | C | A | T | G | T | C | A | C | A | G | A | T | C | T | - | A | G | T | T | #1316888 |
| BA.5.1 | G | G | C | C | A | T | A | C | - | - | - | - | - | - | T | G | G | A | C | C | A | C | A | G | A | T | C | T | A | A | G | T | T |  |
| mut | G>T |  |  |  |  |  | A>G |  |  |  |  |  |  |  |  |  |  |  |  |  |  |  |  |  |  |  |  |  | A>- |  |  |  |  |  |

|  |  |  |  |  |  |  |  |  |  |  |  |  |  |
| --- | --- | --- | --- | --- | --- | --- | --- | --- | --- | --- | --- | --- | --- |
| pos | 396 | 1627 | 1931 | 2954 | 4084 | 6979 | 11750 | 16935 | 19656 | 27038 | 28330 | 28681 |  |
| ref | A | C | C | T | C | T | C | G | G | A | A | G |  |
| BE.1.1 | A | C | A | C | T | T | T | A | G | A | A | T |  |
| BA.5 | A | C | A | C | T | T | C | G | A | G | G | G | #1317484 |
| BF.28 | T | T | C | T | C | G | C | G | A | G | G | G |  |
| mut |  |  |  |  |  |  | T>C |  |  |  |  |  |  |

|  |  |  |  |  |  |  |  |  |  |  |  |  |  |  |  |  |  |  |  |  |  |  |
| --- | --- | --- | --- | --- | --- | --- | --- | --- | --- | --- | --- | --- | --- | --- | --- | --- | --- | --- | --- | --- | --- | --- |
| pos | 686 | 687 | 688 | 689 | 690 | 691 | 692 | 693 | 694 | 1931 | 8695 | 21570 | 26529 | 26858 | 27259 | 27382 | 27383 | 27384 | 27788 | 27889 | 29754 |  |
| ref | A | A | G | T | C | A | T | T | T | C | G | T | G | C | A | G | A | T | G | C | C |  |
| BA.4.1 | - | - | - | - | - | - | - | - | - | C | A | G | G | T | C | C | T | C | T | C | C |  |
| BA.4 | - | - | - | - | - | - | - | - | - | C | A | G | G | T | A | G | A | T | G | T | T | #1317640 |
| BA.5.3 | A | A | G | T | C | A | T | T | T | A | G | T | A | C | A | G | A | T | G | T | T |  |
| mut |  |  |  |  |  |  |  |  |  |  |  |  |  |  |  |  |  |  |  |  |  |  |

|  |  |  |  |  |  |  |  |  |  |  |  |  |  |  |  |  |  |  |  |  |  |  |  |  |  |  |  |  |  |  |  |  |  |  |  |
| --- | --- | --- | --- | --- | --- | --- | --- | --- | --- | --- | --- | --- | --- | --- | --- | --- | --- | --- | --- | --- | --- | --- | --- | --- | --- | --- | --- | --- | --- | --- | --- | --- | --- | --- | --- |
| pos | 4906 | 9713 | 9866 | 11674 | 12160 | 15009 | 15987 | 17211 | 17402 | 18957 | 21077 | 21721 | 21765 | 21766 | 21767 | 21768 | 21769 | 21770 | 21859 | 22510 | 22917 | 23018 | 23040 | 23673 | 25416 | 26529 | 26858 | 27259 | 27382 | 27383 | 27384 | 27889 | 28807 | 29754 |  |
| ref | T | A | C | C | G | T | A | G | C | T | C | C | T | A | C | A | T | G | C | T | T | T | A | C | C | G | C | A | G | A | T | C | C | C |  |
| BA.2.12.1 | T | G | T | T | G | C | G | A | C | T | C | T | T | A | C | A | T | G | T | T | A | T | G | T | T | G | T | C | C | T | C | C | A | C |  |
| BA.5 | T | G | T | T | G | C | G | A | C | C | T | C | - | - | - | - | - | - | C | C | G | G | A | C | C | A | C | A | G | A | T | T | C | T | #1317771 |
| BA.5.6 | C | A | C | C | A | T | A | G | T | C | T | C | - | - | - | - | - | - | C | C | G | G | A | C | C | A | C | A | G | A | T | T | C | T |  |
| mut |  |  |  |  |  |  |  |  |  |  |  |  |  |  |  |  |  |  |  |  |  |  |  |  |  |  |  |  |  |  |  |  |  |  |  |

|  |  |  |  |  |  |  |  |  |  |  |  |  |  |  |  |  |  |  |  |  |  |  |  |  |  |  |  |  |  |  |  |  |  |
| --- | --- | --- | --- | --- | --- | --- | --- | --- | --- | --- | --- | --- | --- | --- | --- | --- | --- | --- | --- | --- | --- | --- | --- | --- | --- | --- | --- | --- | --- | --- | --- | --- | --- |
| pos | 5512 | 6633 | 9866 | 11674 | 12160 | 15009 | 20016 | 21721 | 21765 | 21766 | 21767 | 21768 | 21769 | 21770 | 21789 | 22786 | 22917 | 23018 | 23040 | 23673 | 24526 | 25416 | 25950 | 26529 | 26753 | 26858 | 27259 | 27382 | 27383 | 27384 | 27889 | 29754 |  |
| ref | C | C | C | C | G | T | C | C | T | A | C | A | T | G | C | A | T | T | A | C | G | C | T | G | C | C | A | G | A | T | C | C |  |
| BA.5.5 | T | T | C | C | A | T | C | C | - | - | - | - | - | - | T | A | G | G | A | C | G | C | T | A | C | C | A | G | A | T | T | T |  |
| BA.2.12.1 | C | T | C | C | A | T | C | T | T | A | C | A | T | G | C | C | A | T | G | T | A | T | C | G | T | T | C | C | T | C | C | C | #1318288 |
| BA.2.12.1 | C | C | T | T | G | C | T | T | T | A | C | A | T | G | C | C | A | T | G | T | A | T | T | G | T | T | C | C | T | C | C | C |  |
| mut | T>C |  |  |  |  |  |  |  |  |  |  |  |  |  |  |  |  |  |  |  |  |  | T>C |  |  |  |  |  |  |  |  |  |  |

|  |  |  |  |  |  |  |  |  |  |  |  |  |  |  |  |  |  |  |  |  |  |  |  |  |  |  |  |  |  |  |  |  |  |  |  |  |  |  |
| --- | --- | --- | --- | --- | --- | --- | --- | --- | --- | --- | --- | --- | --- | --- | --- | --- | --- | --- | --- | --- | --- | --- | --- | --- | --- | --- | --- | --- | --- | --- | --- | --- | --- | --- | --- | --- | --- | --- |
| pos | 1627 | 5284 | 6548 | 9300 | 9866 | 12160 | 16098 | 20629 | 21765 | 21766 | 21767 | 21768 | 21769 | 21770 | 22600 | 22917 | 23018 | 23040 | 25416 | 26529 | 26858 | 27038 | 27259 | 27382 | 27383 | 27384 | 27889 | 28057 | 28330 | 28370 | 28371 | 28909 | 29729 | 29754 | 29757 | 29758 | 29759 |  |
| ref | C | C | C | G | C | G | A | C | T | A | C | A | T | G | A | T | T | A | C | G | C | A | A | G | A | T | C | C | A | A | G | C | T | C | G | T | G |  |
| BA.5.2.1 | T | T | C | G | C | A | G | T | - | - | - | - | - | - | A | G | G | A | C | A | C | G | A | G | A | T | T | G | G | T | T | C | T | T | G | T | G |  |
| BA.5.2.1 | T | T | C | G | C | A | G | T | - | - | - | - | - | - | A | G | G | A | C | A | C | G | A | G | A | T | T | G | G | T | T | T | C | C | A | C | C | #1318366 |
| BA.2 | C | C | T | A | T | G | A | C | T | A | C | A | T | G | T | T | T | G | T | G | T | A | C | C | T | C | C | C | A | - | G | C | C | C | A | C | C |  |
| mut |  |  |  |  |  |  |  |  |  |  |  |  |  |  |  |  |  |  |  |  |  |  |  |  |  |  |  |  |  |  |  | C>T |  |  |  |  |  |  |

|  |  |  |  |  |  |  |  |  |  |  |  |  |  |  |  |
| --- | --- | --- | --- | --- | --- | --- | --- | --- | --- | --- | --- | --- | --- | --- | --- |
| pos | 1627 | 3859 | 5221 | 6913 | 6979 | 14827 | 18228 | 21204 | 27038 | 27218 | 27438 | 28330 | 29614 | 29666 |  |
| ref | C | A | C | T | T | A | T | G | A | A | T | A | C | C |  |
| BA.5.1.9 | C | A | T | T | T | G | T | G | A | G | C | A | T | T |  |
| BA.5 | C | A | T | C | T | G | T | T | G | A | T | G | C | C | #1318668 |
| BA.5.2.1 | T | G | C | T | G | A | C | T | G | A | T | G | C | C |  |
| mut |  |  |  | T>C |  |  |  |  |  |  |  |  |  |  |  |

|  |  |  |  |  |  |  |  |  |  |  |  |  |  |  |  |  |
| --- | --- | --- | --- | --- | --- | --- | --- | --- | --- | --- | --- | --- | --- | --- | --- | --- |
| pos | 1170 | 1858 | 4906 | 9826 | 9922 | 16647 | 17402 | 18957 | 20703 | 21077 | 22786 | 26067 | 27438 | 27696 | 29666 |  |
| ref | C | T | T | A | A | G | C | T | C | C | A | T | T | T | C |  |
| BA.5.1.23 | T | C | T | A | A | G | C | T | C | C | C | T | C | C | T |  |
| BA.5.6 | T | C | T | G | A | A | T | C | T | T | A | G | T | T | C | #1319113 |
| BA.5.6 | C | T | C | A | G | G | T | C | T | T | C | G | T | T | C |  |
| mut |  |  |  | A>G |  | G>A |  |  |  |  | C>A |  |  |  |  |  |

|  |  |  |  |  |  |  |  |  |  |  |  |  |  |  |  |  |  |  |  |  |  |  |  |  |  |  |
| --- | --- | --- | --- | --- | --- | --- | --- | --- | --- | --- | --- | --- | --- | --- | --- | --- | --- | --- | --- | --- | --- | --- | --- | --- | --- | --- |
| pos | 335 | 960 | 3796 | 3927 | 4586 | 5183 | 12444 | 12969 | 14259 | 15451 | 16342 | 18286 | 22001 | 22016 | 22033 | 22190 | 22331 | 22577 | 22898 | 22942 | 23040 | 23283 | 25416 | 26275 | 29254 |  |
| ref | C | G | C | C | C | C | A | A | T | G | T | G | A | T | C | A | G | G | G | T | A | A | C | A | G |  |
| BA.2.10 | T | A | C | C | C | C | A | A | T | G | C | A | A | T | C | A | G | G | G | T | G | A | C | A | T |  |
| BA.2.75 | C | A | C | C | C | C | G | G | C | A | T | G | G | C | A | G | A | C | A | G | A | T | T | G | G | #1320033 |
| BA.2.75.1 | C | G | T | T | T | T | G | A | T | A | T | G | G | C | A | G | A | C | A | G | A | T | T | G | G |  |
| mut | T>C |  |  |  |  |  |  | A>G | T>C |  |  |  |  |  |  |  |  |  |  |  |  |  |  |  |  |  |

|  |  |  |  |  |  |  |  |  |  |  |  |  |  |  |  |  |  |  |  |  |  |  |  |  |  |  |  |  |  |  |  |  |
| --- | --- | --- | --- | --- | --- | --- | --- | --- | --- | --- | --- | --- | --- | --- | --- | --- | --- | --- | --- | --- | --- | --- | --- | --- | --- | --- | --- | --- | --- | --- | --- | --- |
| pos | 44 | 1627 | 2177 | 6979 | 9866 | 9967 | 12160 | 18570 | 21765 | 21766 | 21767 | 21768 | 21769 | 21770 | 22917 | 23018 | 23040 | 24620 | 26529 | 26858 | 27038 | 27259 | 27382 | 27383 | 27384 | 27532 | 27739 | 27889 | 28330 | 28402 | 29754 |  |
| ref | C | C | G | T | C | C | G | C | T | A | C | A | T | G | T | T | A | G | G | C | A | A | G | A | T | C | C | C | A | A | C |  |
| BF.5 | T | T | T | G | C | T | A | C | - | - | - | - | - | - | G | G | A | T | A | C | G | A | G | A | T | T | C | T | G | G | T |  |
| BF.24 | T | T | T | G | C | T | A | T | - | - | - | - | - | - | G | G | A | T | A | C | G | A | G | A | T | C | T | C | A | A | C | #1321052 |
| BA.2 | C | C | G | T | T | C | G | C | T | A | C | A | T | G | T | T | G | G | G | T | A | C | C | T | C | C | T | C | A | A | C |  |
| mut |  |  |  |  |  |  |  | C>T |  |  |  |  |  |  |  |  |  |  |  |  |  |  |  |  |  | T>C |  |  |  |  |  |  |

|  |  |  |  |  |  |  |  |  |  |  |  |  |  |  |  |  |  |  |  |  |  |  |  |  |  |  |  |  |  |  |  |
| --- | --- | --- | --- | --- | --- | --- | --- | --- | --- | --- | --- | --- | --- | --- | --- | --- | --- | --- | --- | --- | --- | --- | --- | --- | --- | --- | --- | --- | --- | --- | --- |
| pos | 1627 | 6548 | 9300 | 9866 | 12160 | 21765 | 21766 | 21767 | 21768 | 21769 | 21770 | 22600 | 22917 | 23018 | 23040 | 25416 | 26529 | 26858 | 27038 | 27259 | 27382 | 27383 | 27384 | 27889 | 28330 | 29729 | 29754 | 29757 | 29758 | 29759 |  |
| ref | C | C | G | C | G | T | A | C | A | T | G | A | T | T | A | C | G | C | A | A | G | A | T | C | A | T | C | G | T | G |  |
| BA.5.2.1 | T | C | G | C | A | - | - | - | - | - | - | A | G | G | A | C | A | C | G | A | G | A | T | T | G | T | T | G | T | G |  |
| BA.5.2.1 | T | C | G | C | A | - | - | - | - | - | - | A | G | G | A | C | A | C | G | A | G | A | T | T | G | C | C | A | C | C | #1321396 |
| BA.2 | C | T | A | T | G | T | A | C | A | T | G | T | T | T | G | T | G | T | A | C | C | T | C | C | A | C | C | A | C | C |  |
| mut |  |  |  |  |  |  |  |  |  |  |  |  |  |  |  |  |  |  |  |  |  |  |  |  |  |  |  |  |  |  |  |

|  |  |  |  |  |  |  |  |  |  |  |  |  |  |  |  |  |
| --- | --- | --- | --- | --- | --- | --- | --- | --- | --- | --- | --- | --- | --- | --- | --- | --- |
| pos | 1627 | 4690 | 6979 | 18501 | 21520 | 22879 | 26029 | 26122 | 27038 | 27438 | 28330 | 28395 | 28677 | 29614 | 29666 |  |
| ref | C | T | T | C | G | C | C | G | A | T | A | G | C | C | C |  |
| BA.5.1 | C | T | T | T | G | C | A | G | A | C | A | A | T | T | T |  |
| BF.1 | C | T | T | T | T | T | C | A | G | T | G | G | C | C | C | #1321670 |
| BF.1 | T | C | G | C | T | T | C | A | G | T | G | G | C | C | C |  |
| mut |  |  |  |  |  |  |  |  |  |  |  |  |  |  |  |  |

|  |  |  |  |  |  |  |  |  |  |  |  |  |  |  |  |
| --- | --- | --- | --- | --- | --- | --- | --- | --- | --- | --- | --- | --- | --- | --- | --- |
| pos | 1820 | 1931 | 2890 | 12554 | 16935 | 21045 | 22377 | 25455 | 26111 | 27438 | 27548 | 28462 | 28681 | 29666 |  |
| ref | G | C | G | G | G | T | C | G | C | T | A | C | G | C |  |
| BA.5.1 | G | C | G | A | G | C | T | G | C | C | A | T | G | T |  |
| BE.1 | A | C | G | A | G | C | T | T | T | T | G | C | T | C | #1322026 |
| BE.1 | G | A | T | G | A | T | C | T | T | T | G | C | T | C |  |
| mut | G>A |  |  |  |  |  |  |  |  |  |  |  |  |  |  |

|  |  |  |  |  |  |  |  |  |  |  |  |  |  |  |  |
| --- | --- | --- | --- | --- | --- | --- | --- | --- | --- | --- | --- | --- | --- | --- | --- |
| pos | 1627 | 6979 | 9967 | 11138 | 19524 | 24620 | 25672 | 27038 | 27438 | 27532 | 28330 | 28402 | 29614 | 29666 |  |
| ref | C | T | C | G | C | G | C | A | T | C | A | A | C | C |  |
| BA.5.1.1 | C | T | C | A | C | G | A | A | C | C | A | A | T | T |  |
| BF.5 | C | T | C | A | C | T | C | G | T | T | G | G | C | C | #1324872 |
| BF.5 | T | G | T | G | T | T | C | G | T | T | G | G | C | C |  |
| mut |  |  |  |  |  |  |  |  |  |  |  |  |  |  |  |

|  |  |  |  |  |  |  |  |  |  |  |  |  |  |  |  |  |
| --- | --- | --- | --- | --- | --- | --- | --- | --- | --- | --- | --- | --- | --- | --- | --- | --- |
| pos | 1170 | 1627 | 5175 | 5950 | 12311 | 12651 | 16920 | 20429 | 25473 | 27038 | 27438 | 27696 | 28330 | 28374 | 29666 |  |
| ref | C | C | C | G | G | C | A | C | T | A | T | T | A | G | C |  |
| BA.5.2.1 | C | T | C | G | G | T | G | T | C | G | T | T | G | G | C |  |
| BA.5.1 | C | T | C | G | G | T | A | C | T | A | C | C | A | T | T | #1327685 |
| BA.5.1.23 | T | C | T | T | T | C | A | C | T | A | C | C | A | T | T |  |
| mut |  |  |  |  |  |  |  |  |  |  |  |  |  |  |  |  |

|  |  |  |  |  |  |  |  |  |  |  |  |  |  |  |  |  |  |  |  |  |
| --- | --- | --- | --- | --- | --- | --- | --- | --- | --- | --- | --- | --- | --- | --- | --- | --- | --- | --- | --- | --- |
| pos | 44 | 686 | 687 | 689 | 691 | 692 | 5826 | 6720 | 6979 | 12310 | 13136 | 15849 | 16616 | 22110 | 26625 | 27012 | 27038 | 27513 | 28690 |  |
| ref | C | A | A | T | A | T | C | C | T | G | A | T | C | A | C | C | A | C | G |  |
| BA.5.2.1 | C | G | C | C | G | C | T | C | G | G | A | T | C | T | C | C | G | C | G |  |
| BA.5.2 | C | G | C | C | G | C | C | T | T | A | G | A | A | A | T | T | A | T | T | #1330349 |
| BA.5.2 | T | A | A | T | A | T | C | T | T | A | G | A | A | A | T | T | A | T | T |  |
| mut |  |  |  |  |  |  |  |  |  |  |  |  |  |  |  |  |  |  |  |  |

|  |  |  |  |  |  |  |  |  |  |  |  |  |  |  |  |  |  |  |  |  |  |  |  |  |  |  |  |  |  |  |  |  |  |  |  |  |  |
| --- | --- | --- | --- | --- | --- | --- | --- | --- | --- | --- | --- | --- | --- | --- | --- | --- | --- | --- | --- | --- | --- | --- | --- | --- | --- | --- | --- | --- | --- | --- | --- | --- | --- | --- | --- | --- | --- |
| pos | 1931 | 3875 | 6548 | 9300 | 9866 | 12068 | 12160 | 16710 | 16935 | 21765 | 21766 | 21767 | 21768 | 21769 | 21770 | 22600 | 22792 | 22917 | 23018 | 23040 | 25416 | 25534 | 25537 | 26529 | 26858 | 27259 | 27382 | 27383 | 27384 | 27889 | 28681 | 29729 | 29754 | 29757 | 29758 | 29759 |  |
| ref | C | G | C | G | C | C | G | G | G | T | A | C | A | T | G | A | C | T | T | A | C | G | G | G | C | A | G | A | T | C | G | T | C | G | T | G |  |
| BE.1 | A | A | C | G | C | T | A | G | A | - | - | - | - | - | - | A | T | G | G | A | C | T | T | A | C | A | G | A | T | T | T | T | T | G | T | G |  |
| BE.1 | A | A | C | G | C | T | A | A | A | - | - | - | - | - | - | A | T | G | G | A | C | T | T | A | C | A | G | A | T | T | T | C | C | A | C | C | #1331484 |
| BA.2 | C | G | T | A | T | C | G | G | G | T | A | C | A | T | G | T | C | T | T | G | T | G | G | G | T | C | C | T | C | C | G | C | C | A | C | C |  |
| mut |  |  |  |  |  |  |  | G>A |  |  |  |  |  |  |  |  |  |  |  |  |  |  |  |  |  |  |  |  |  |  |  |  |  |  |  |  |  |

|  |  |  |  |  |  |  |  |  |  |  |  |  |  |  |
| --- | --- | --- | --- | --- | --- | --- | --- | --- | --- | --- | --- | --- | --- | --- |
| pos | 1627 | 6979 | 8882 | 9967 | 17285 | 19524 | 24620 | 27038 | 27438 | 27532 | 28330 | 28402 | 29666 |  |
| ref | C | T | A | C | C | C | G | A | T | C | A | A | C |  |
| BF.5 | T | G | G | T | C | T | T | G | T | T | G | G | C |  |
| BA.5.1.22 | T | G | G | T | T | C | G | A | C | C | A | A | T | #1333151 |
| BA.5.1.22 | C | T | A | C | T | C | G | A | C | C | A | A | T |  |
| mut |  |  |  |  |  |  |  |  |  |  |  |  |  |  |

|  |  |  |  |  |  |  |  |  |  |  |  |  |  |  |  |  |  |  |  |  |  |  |  |  |  |  |  |  |  |  |  |  |  |
| --- | --- | --- | --- | --- | --- | --- | --- | --- | --- | --- | --- | --- | --- | --- | --- | --- | --- | --- | --- | --- | --- | --- | --- | --- | --- | --- | --- | --- | --- | --- | --- | --- | --- |
| pos | 44 | 1627 | 2177 | 6979 | 9967 | 12160 | 13821 | 19955 | 20055 | 21765 | 21766 | 21767 | 21768 | 21769 | 21770 | 22917 | 23018 | 23040 | 24620 | 26529 | 26858 | 26873 | 27038 | 27259 | 27382 | 27383 | 27384 | 27532 | 27889 | 28330 | 28402 | 29754 |  |
| ref | C | C | G | T | C | G | C | C | A | T | A | C | A | T | G | T | T | A | G | G | C | C | A | A | G | A | T | C | C | A | A | C |  |
| BF.5 | T | T | T | G | T | A | T | T | G | - | - | - | - | - | - | G | G | A | T | A | C | C | G | A | G | A | T | T | T | G | G | T |  |
| BA.4 | T | T | T | G | T | A | T | T | G | - | - | - | - | - | - | G | G | A | T | G | T | T | A | C | C | T | C | C | C | A | A | C | #1333649 |
| BA.2 | C | C | G | T | C | G | C | C | A | T | A | C | A | T | G | T | T | G | G | G | T | C | A | C | C | T | C | C | C | A | A | C |  |
| mut |  |  |  |  |  |  |  |  |  |  |  |  |  |  |  |  |  |  |  |  |  | C>T |  |  |  |  |  |  |  |  |  |  |  |

|  |  |  |  |  |  |  |  |  |  |  |  |  |  |  |  |  |  |
| --- | --- | --- | --- | --- | --- | --- | --- | --- | --- | --- | --- | --- | --- | --- | --- | --- | --- |
| pos | 1627 | 5284 | 5929 | 9988 | 14599 | 15026 | 16293 | 20462 | 25553 | 27038 | 27438 | 28330 | 28370 | 28371 | 28651 | 29666 |  |
| ref | C | C | T | C | C | C | C | C | C | A | T | A | A | G | C | C |  |
| BA.5.1.7 | C | C | C | T | C | T | T | T | C | A | C | A | - | G | C | T |  |
| BA.5.1.7 | C | C | C | T | C | T | T | T | T | G | T | G | T | T | T | C | #1334856 |
| BA.5.2.1 | T | T | T | C | T | C | C | C | T | G | T | G | T | T | C | C |  |
| mut |  |  |  |  |  |  |  |  |  |  |  |  |  |  | C>T |  |  |

|  |  |  |  |  |  |  |  |  |  |  |  |  |  |  |  |  |
| --- | --- | --- | --- | --- | --- | --- | --- | --- | --- | --- | --- | --- | --- | --- | --- | --- |
| pos | 594 | 4347 | 7493 | 10507 | 11906 | 12081 | 15451 | 19174 | 20429 | 22841 | 28330 | 28370 | 28371 | 29284 | 29666 |  |
| ref | A | C | A | C | G | C | G | G | C | G | A | A | G | C | C |  |
| BA.5.1.21 | A | C | G | T | A | C | A | A | T | A | A | - | G | T | T |  |
| BA.5.1.21 | A | C | G | T | A | T | A | A | C | G | G | T | T | C | C | #1335050 |
| BA.5 | G | T | A | C | G | C | G | G | C | G | G | T | T | C | C |  |
| mut |  |  |  |  |  | C>T |  |  |  |  |  |  |  |  |  |  |

|  |  |  |  |  |  |  |  |  |  |  |  |  |  |  |  |  |  |
| --- | --- | --- | --- | --- | --- | --- | --- | --- | --- | --- | --- | --- | --- | --- | --- | --- | --- |
| pos | 1931 | 3875 | 12068 | 16935 | 17130 | 17746 | 18160 | 18501 | 22792 | 23323 | 25534 | 25537 | 27438 | 28681 | 29614 | 29666 |  |
| ref | C | G | C | G | C | C | G | C | C | T | G | G | T | G | C | C |  |
| BA.5.1 | C | G | C | G | T | T | A | C | C | T | G | G | C | G | T | T |  |
| BE.1 | C | G | C | G | T | T | A | T | T | C | T | T | T | T | C | C | #1335368 |
| BE.1 | A | A | T | A | C | C | G | C | T | C | T | T | T | T | C | C |  |
| mut |  |  |  |  |  |  |  | C>T | C>T |  |  |  |  |  |  |  |  |

|  |  |  |  |  |  |  |  |  |  |  |  |  |  |  |
| --- | --- | --- | --- | --- | --- | --- | --- | --- | --- | --- | --- | --- | --- | --- |
| pos | 1627 | 3096 | 4595 | 6894 | 11109 | 12310 | 15451 | 16616 | 27012 | 27438 | 27513 | 28330 | 29666 |  |
| ref | C | C | A | G | C | G | G | C | C | T | C | A | C |  |
| BA.5.1 | C | T | G | G | T | G | A | C | C | C | C | A | T |  |
| BA.5.2 | C | T | G | G | T | G | A | C | T | T | T | G | C | #1335483 |
| BA.5.2 | T | C | A | T | C | A | G | A | T | T | T | G | C |  |
| mut |  |  |  |  |  |  |  |  |  |  |  |  |  |  |

|  |  |  |  |  |  |  |  |  |  |  |  |  |  |  |  |  |  |  |  |  |  |  |  |  |  |  |  |  |  |  |  |  |  |  |  |  |
| --- | --- | --- | --- | --- | --- | --- | --- | --- | --- | --- | --- | --- | --- | --- | --- | --- | --- | --- | --- | --- | --- | --- | --- | --- | --- | --- | --- | --- | --- | --- | --- | --- | --- | --- | --- | --- |
| pos | 44 | 1593 | 1627 | 5012 | 5365 | 6573 | 9866 | 12160 | 12310 | 13436 | 16616 | 17232 | 18744 | 20462 | 21765 | 21766 | 21767 | 21768 | 21769 | 21770 | 22917 | 23018 | 23040 | 26529 | 26858 | 26942 | 27012 | 27259 | 27382 | 27383 | 27384 | 27513 | 27889 | 28330 | 29754 |  |
| ref | C | C | C | G | C | C | C | G | G | C | C | A | C | C | T | A | C | A | T | G | T | T | A | G | C | C | C | A | G | A | T | C | C | A | C |  |
| BA.2 | C | T | C | G | T | C | T | G | G | C | C | A | C | C | T | A | C | A | T | G | T | T | G | G | T | C | C | C | C | T | C | C | C | A | C |  |
| BA.5.2 | C | T | C | T | T | C | C | A | A | T | A | G | T | T | - | - | - | - | - | - | G | G | A | A | C | T | T | A | G | A | T | T | T | G | T | #1337115 |
| BA.5.2 | T | C | T | G | C | T | C | A | A | C | A | G | T | T | - | - | - | - | - | - | G | G | A | A | C | C | T | A | G | A | T | T | T | G | T |  |
| mut |  |  |  | G>T |  |  |  |  |  | C>T |  |  |  |  |  |  |  |  |  |  |  |  |  |  |  | C>T |  |  |  |  |  |  |  |  |  |  |

|  |  |  |  |  |  |  |  |  |  |  |  |  |  |  |  |  |  |  |  |  |  |  |  |  |  |  |  |  |  |  |  |  |
| --- | --- | --- | --- | --- | --- | --- | --- | --- | --- | --- | --- | --- | --- | --- | --- | --- | --- | --- | --- | --- | --- | --- | --- | --- | --- | --- | --- | --- | --- | --- | --- | --- |
| pos | 686 | 687 | 688 | 689 | 690 | 691 | 692 | 693 | 694 | 1627 | 4690 | 6979 | 10319 | 11837 | 13890 | 21520 | 21570 | 22879 | 25922 | 26529 | 26858 | 27038 | 27259 | 27382 | 27383 | 27384 | 27788 | 27889 | 28330 | 28724 | 29754 |  |
| ref | A | A | G | T | C | A | T | T | T | C | T | T | C | G | T | G | T | C | G | G | C | A | A | G | A | T | G | C | A | C | C |  |
| BF.1 | A | A | G | T | C | A | T | T | T | T | C | G | C | T | T | T | T | T | G | A | C | G | A | G | A | T | G | T | G | C | T |  |
| BA.4.1 | A | A | G | T | C | A | T | T | T | T | C | G | T | T | C | G | G | C | A | G | T | A | C | C | T | C | T | C | A | T | C | #1337674 |
| BA.4.1 | - | - | - | - | - | - | - | - | - | C | T | T | C | G | T | G | G | C | G | G | T | A | C | C | T | C | T | C | A | T | C |  |
| mut |  |  |  |  |  |  |  |  |  |  |  |  | C>T |  | T>C |  |  |  | G>A |  |  |  |  |  |  |  |  |  |  |  |  |  |

|  |  |  |  |  |  |  |  |  |  |  |  |  |  |  |  |  |  |  |  |
| --- | --- | --- | --- | --- | --- | --- | --- | --- | --- | --- | --- | --- | --- | --- | --- | --- | --- | --- | --- |
| pos | 219 | 1376 | 1627 | 1931 | 2121 | 2910 | 4586 | 5284 | 5869 | 10399 | 13342 | 16376 | 25546 | 27038 | 28330 | 28370 | 28371 | 28681 |  |
| ref | G | C | C | C | C | C | C | C | C | A | C | C | C | A | A | A | G | G |  |
| BE.2 | A | T | C | A | C | T | T | C | T | G | T | T | T | A | A | - | G | T |  |
| BE.2 | A | T | C | A | C | T | T | C | T | G | C | C | C | G | G | T | T | G | #1338277 |
| BA.5.2.1 | G | C | T | C | T | C | C | T | C | A | C | C | C | G | G | T | T | G |  |
| mut |  |  |  |  |  |  |  |  |  |  |  |  |  |  |  |  |  |  |  |

|  |  |  |  |  |  |  |  |  |  |  |  |  |  |  |  |  |  |  |  |  |  |  |  |  |  |  |  |  |  |  |  |  |  |  |  |
| --- | --- | --- | --- | --- | --- | --- | --- | --- | --- | --- | --- | --- | --- | --- | --- | --- | --- | --- | --- | --- | --- | --- | --- | --- | --- | --- | --- | --- | --- | --- | --- | --- | --- | --- | --- |
| pos | 44 | 396 | 421 | 1627 | 5549 | 6979 | 9319 | 9866 | 12160 | 18452 | 20703 | 21765 | 21766 | 21767 | 21768 | 21769 | 21770 | 22917 | 23018 | 23040 | 23557 | 23663 | 25091 | 25207 | 26529 | 26858 | 27038 | 27259 | 27382 | 27383 | 27384 | 27889 | 28330 | 29754 |  |
| ref | C | A | C | C | C | T | C | C | G | C | C | T | A | C | A | T | G | T | T | A | C | G | G | C | G | C | A | A | G | A | T | C | A | C |  |
| BA.2 | C | A | C | C | A | T | C | T | G | C | C | T | A | C | A | T | G | T | T | G | C | G | A | C | G | T | A | C | C | T | C | C | A | C |  |
| BF.28 | C | A | C | C | A | T | T | C | A | T | T | - | - | - | - | - | - | G | G | A | T | T | G | T | A | C | G | A | G | A | T | T | G | T | #1338501 |
| BF.28 | T | T | T | T | C | G | T | C | A | T | T | - | - | - | - | - | - | G | G | A | T | T | G | T | A | C | G | A | G | A | T | T | G | T |  |
| mut |  |  |  |  |  |  |  |  |  |  |  |  |  |  |  |  |  |  |  |  |  |  |  |  |  |  |  |  |  |  |  |  |  |  |  |

|  |  |  |  |  |  |  |  |  |  |  |  |  |  |  |  |  |  |  |  |  |  |  |  |  |  |  |  |  |
| --- | --- | --- | --- | --- | --- | --- | --- | --- | --- | --- | --- | --- | --- | --- | --- | --- | --- | --- | --- | --- | --- | --- | --- | --- | --- | --- | --- | --- |
| pos | 44 | 686 | 687 | 688 | 689 | 690 | 691 | 692 | 693 | 694 | 7042 | 8140 | 14599 | 22599 | 23535 | 26529 | 26858 | 27259 | 27382 | 27383 | 27384 | 27438 | 27788 | 27889 | 28724 | 29666 | 29754 |  |
| ref | C | A | A | G | T | C | A | T | T | T | G | C | C | G | A | G | C | A | G | A | T | T | G | C | C | C | C |  |
| BA.5.1 | C | G | C | G | C | C | G | C | T | A | G | C | C | G | A | A | C | A | G | A | T | C | G | T | C | T | T |  |
| BA.4.6 | C | G | C | G | C | C | G | C | T | A | T | T | T | C | G | G | T | C | C | T | C | T | T | C | T | C | C | #1338895 |
| BA.4.6 | T | - | - | - | - | - | - | - | - | - | T | T | T | C | G | G | T | C | C | T | C | T | T | C | T | C | C |  |
| mut |  |  |  |  |  |  |  |  |  |  |  |  |  |  |  |  |  |  |  |  |  |  |  |  |  |  |  |  |

|  |  |  |  |  |  |  |  |  |  |  |  |  |  |  |  |  |
| --- | --- | --- | --- | --- | --- | --- | --- | --- | --- | --- | --- | --- | --- | --- | --- | --- |
| pos | 1627 | 6979 | 7869 | 9967 | 11781 | 19524 | 24620 | 27038 | 27438 | 27532 | 28233 | 28330 | 28402 | 28462 | 29666 |  |
| ref | C | T | C | C | A | C | G | A | T | C | G | A | A | C | C |  |
| BF.5 | T | G | C | T | A | T | T | G | T | T | G | G | G | C | C |  |
| BA.5.1 | T | G | T | T | A | T | G | A | C | C | C | A | A | T | T | #1340062 |
| BA.5.1.10 | C | T | C | C | G | C | G | A | C | C | C | A | A | T | T |  |
| mut |  |  | C>T |  |  |  |  |  |  |  |  |  |  |  |  |  |

|  |  |  |  |  |  |  |  |  |  |  |  |  |  |  |  |  |  |  |  |  |  |  |  |  |  |  |  |  |  |  |  |  |  |  |  |
| --- | --- | --- | --- | --- | --- | --- | --- | --- | --- | --- | --- | --- | --- | --- | --- | --- | --- | --- | --- | --- | --- | --- | --- | --- | --- | --- | --- | --- | --- | --- | --- | --- | --- | --- | --- |
| pos | 686 | 687 | 688 | 689 | 690 | 691 | 692 | 693 | 694 | 1627 | 5284 | 8140 | 10486 | 11950 | 12310 | 14599 | 16616 | 18828 | 22599 | 23535 | 26529 | 26858 | 27012 | 27259 | 27382 | 27383 | 27384 | 27513 | 27788 | 27889 | 28330 | 28724 | 28792 | 29754 |  |
| ref | A | A | G | T | C | A | T | T | T | C | C | C | A | C | G | C | C | C | G | A | G | C | C | A | G | A | T | C | G | C | A | C | A | C |  |
| BA.4.6 | - | - | - | - | - | - | - | - | - | C | C | T | A | T | G | T | C | T | C | G | G | T | C | C | C | T | C | C | T | C | A | T | A | C |  |
| BA.5.2 | - | - | - | - | - | - | - | - | - | C | C | T | G | T | G | T | C | T | C | G | A | C | T | A | G | A | T | T | G | T | G | C | T | T | #1340173 |
| BA.5.2.21 | A | A | G | T | C | A | T | T | T | T | T | C | A | C | A | C | A | C | G | A | A | C | T | A | G | A | T | T | G | T | G | C | T | T |  |
| mut |  |  |  |  |  |  |  |  |  |  |  |  | A>G |  |  |  |  |  |  |  |  |  |  |  |  |  |  |  |  |  |  |  |  |  |  |

|  |  |  |  |  |  |  |  |  |  |  |  |  |  |  |
| --- | --- | --- | --- | --- | --- | --- | --- | --- | --- | --- | --- | --- | --- | --- |
| pos | 1627 | 6979 | 10507 | 20949 | 23202 | 24620 | 27038 | 27259 | 27382 | 27383 | 27384 | 27889 | 28330 |  |
| ref | C | T | C | T | C | G | A | A | G | A | T | C | A |  |
| BF.3 | T | G | C | T | T | T | G | A | G | A | T | T | G |  |
| BA.4 | T | G | T | C | T | T | G | C | C | T | C | C | A | #1341416 |
| BA.4 | C | T | C | T | C | G | A | C | C | T | C | C | A |  |
| mut |  |  | C>T | T>C |  |  |  |  |  |  |  |  |  |  |

|  |  |  |  |  |  |  |  |  |  |  |  |  |  |  |  |  |  |  |  |  |  |  |  |  |  |  |  |
| --- | --- | --- | --- | --- | --- | --- | --- | --- | --- | --- | --- | --- | --- | --- | --- | --- | --- | --- | --- | --- | --- | --- | --- | --- | --- | --- | --- |
| pos | 686 | 687 | 688 | 689 | 690 | 691 | 692 | 693 | 694 | 4906 | 17402 | 18957 | 18984 | 21077 | 22599 | 23535 | 26529 | 26858 | 27259 | 27382 | 27383 | 27384 | 27788 | 27889 | 28724 | 29754 |  |
| ref | A | A | G | T | C | A | T | T | T | T | C | T | G | C | G | A | G | C | A | G | A | T | G | C | C | C |  |
| BA.5.6 | A | A | G | T | C | A | T | T | T | C | T | C | T | T | G | A | A | C | A | G | A | T | G | T | C | T |  |
| BA.4 | A | A | G | T | C | A | T | T | T | C | T | C | T | T | C | G | G | T | C | C | T | C | T | C | T | C | #1341590 |
| BA.4 | - | - | - | - | - | - | - | - | - | T | C | T | G | C | G | G | G | T | C | C | T | C | T | C | T | C |  |
| mut |  |  |  |  |  |  |  |  |  |  |  |  |  |  | G>C |  |  |  |  |  |  |  |  |  |  |  |  |

|  |  |  |  |  |  |  |  |  |  |  |  |  |  |  |  |  |  |
| --- | --- | --- | --- | --- | --- | --- | --- | --- | --- | --- | --- | --- | --- | --- | --- | --- | --- |
| pos | 1627 | 5284 | 7303 | 8641 | 9430 | 11782 | 14184 | 17371 | 21789 | 25585 | 27038 | 27235 | 28330 | 28370 | 28371 | 29223 |  |
| ref | C | C | C | T | C | A | C | G | C | C | A | G | A | A | G | T |  |
| BA.5.5 | C | C | C | C | T | G | C | A | T | C | A | G | A | - | G | T |  |
| BA.5 | C | C | C | C | T | G | C | G | C | T | G | A | G | T | T | C | #1344192 |
| BA.5.2.1 | T | T | T | T | C | A | T | G | C | T | G | A | G | T | T | C |  |
| mut |  |  |  |  |  |  |  |  |  |  |  |  |  |  |  |  |  |

|  |  |  |  |  |  |  |  |  |  |  |  |  |  |  |  |  |  |  |  |  |  |  |  |  |  |  |  |  |  |  |
| --- | --- | --- | --- | --- | --- | --- | --- | --- | --- | --- | --- | --- | --- | --- | --- | --- | --- | --- | --- | --- | --- | --- | --- | --- | --- | --- | --- | --- | --- | --- |
| pos | 44 | 335 | 3411 | 8917 | 9866 | 12160 | 13423 | 13605 | 21676 | 21765 | 21766 | 21767 | 21768 | 21769 | 21770 | 21789 | 22917 | 23018 | 23040 | 25416 | 26529 | 26858 | 26895 | 27259 | 27382 | 27383 | 27384 | 27889 | 29754 |  |
| ref | C | C | C | C | C | G | C | C | C | T | A | C | A | T | G | C | T | T | A | C | G | C | C | A | G | A | T | C | C |  |
| BA.5.5 | T | T | T | T | C | A | T | T | C | - | - | - | - | - | - | T | G | G | A | C | A | C | C | A | G | A | T | T | T |  |
| BA.4 | T | T | T | T | C | A | T | T | T | - | - | - | - | - | - | T | G | G | A | T | G | T | T | C | C | T | C | C | C | #1344977 |
| BA.2 | C | C | C | C | T | G | C | C | C | T | A | C | A | T | G | C | T | T | G | T | G | T | T | C | C | T | C | C | C |  |
| mut |  |  |  |  |  |  |  |  | C>T |  |  |  |  |  |  |  |  |  |  |  |  |  |  |  |  |  |  |  |  |  |

|  |  |  |  |  |  |  |  |  |  |  |  |  |  |  |
| --- | --- | --- | --- | --- | --- | --- | --- | --- | --- | --- | --- | --- | --- | --- |
| pos | 1627 | 1931 | 6025 | 12310 | 16616 | 17207 | 17304 | 27012 | 27513 | 28330 | 28681 | 28724 | 29702 |  |
| ref | C | C | T | G | C | A | C | C | C | A | G | C | G |  |
| BE.3 | C | A | C | G | C | G | T | C | C | A | T | T | G |  |
| BA.5.2 | C | A | C | G | C | G | T | T | T | G | G | C | A | #1346141 |
| BA.5.2.1 | T | C | T | A | A | A | C | T | T | G | G | C | A |  |
| mut |  |  |  |  |  |  |  |  |  |  |  |  |  |  |

|  |  |  |  |  |  |  |  |  |  |  |  |  |  |  |  |  |  |  |  |  |  |  |  |  |  |  |  |  |  |  |  |  |  |  |  |  |  |  |  |  |  |
| --- | --- | --- | --- | --- | --- | --- | --- | --- | --- | --- | --- | --- | --- | --- | --- | --- | --- | --- | --- | --- | --- | --- | --- | --- | --- | --- | --- | --- | --- | --- | --- | --- | --- | --- | --- | --- | --- | --- | --- | --- | --- |
| pos | 832 | 2243 | 3096 | 4906 | 9866 | 11674 | 12076 | 12160 | 16647 | 16726 | 17402 | 18957 | 18984 | 21077 | 21765 | 21766 | 21767 | 21768 | 21769 | 21770 | 22917 | 23018 | 23040 | 23673 | 25352 | 25416 | 26529 | 26858 | 27259 | 27382 | 27383 | 27384 | 27654 | 27889 | 28166 | 28380 | 28657 | 29367 | 29754 | 29779 |  |
| ref | C | G | C | T | C | C | C | G | G | C | C | T | G | C | T | A | C | A | T | G | T | T | A | C | G | C | G | C | A | G | A | T | C | C | G | G | C | C | C | G |  |
| BA.2.12.1 | T | A | T | T | T | T | C | G | T | T | C | T | G | C | T | A | C | A | T | G | A | T | G | T | T | T | G | T | C | C | T | C | T | C | G | G | T | T | C | T |  |
| BA.5.6 | T | A | T | T | T | T | T | G | G | C | T | C | T | T | - | - | - | - | - | - | G | G | A | C | G | C | A | C | A | G | A | T | C | T | A | A | C | C | T | G | #1346198 |
| BA.5.6 | C | G | C | C | C | C | C | A | G | C | T | C | T | T | - | - | - | - | - | - | G | G | A | C | G | C | A | C | A | G | A | T | C | T | A | G | C | C | T | G |  |
| mut |  |  |  |  |  |  | C>T |  |  |  |  |  |  |  |  |  |  |  |  |  |  |  |  |  |  |  |  |  |  |  |  |  |  |  |  | G>A |  |  |  |  |  |

|  |  |  |  |  |  |  |  |  |  |  |  |  |  |  |  |  |  |  |  |  |  |  |  |
| --- | --- | --- | --- | --- | --- | --- | --- | --- | --- | --- | --- | --- | --- | --- | --- | --- | --- | --- | --- | --- | --- | --- | --- |
| pos | 1627 | 1729 | 1884 | 3787 | 7348 | 7732 | 9073 | 11494 | 11514 | 12310 | 13821 | 23013 | 24386 | 25049 | 25521 | 27438 | 27879 | 28079 | 28330 | 29254 | 29666 | 29670 |  |
| ref | C | G | C | C | T | C | C | C | C | G | C | A | G | G | C | T | C | G | A | G | C | C |  |
| BA.5.2 | T | G | C | C | C | T | C | C | T | A | C | T | G | G | T | T | C | T | G | G | C | C |  |
| BA.5.1.25 | T | G | C | C | C | T | C | C | T | A | T | T | G | G | T | C | T | G | A | T | T | T | #1348408 |
| BA.5.1.25 | C | A | T | T | T | C | T | T | C | G | C | C | A | C | C | C | T | G | A | T | T | T |  |
| mut |  |  |  |  |  |  |  |  |  |  | C>T |  |  |  |  |  |  |  |  |  |  |  |  |

|  |  |  |  |  |  |  |  |  |  |  |  |  |  |  |  |  |  |  |  |  |  |  |  |  |  |  |  |  |  |  |  |  |
| --- | --- | --- | --- | --- | --- | --- | --- | --- | --- | --- | --- | --- | --- | --- | --- | --- | --- | --- | --- | --- | --- | --- | --- | --- | --- | --- | --- | --- | --- | --- | --- | --- |
| pos | 1627 | 6979 | 9866 | 12160 | 19812 | 21255 | 21765 | 21766 | 21767 | 21768 | 21769 | 21770 | 22304 | 22599 | 22917 | 23018 | 23040 | 23202 | 24620 | 25416 | 26529 | 26858 | 27038 | 27259 | 27382 | 27383 | 27384 | 27549 | 27889 | 28330 | 29754 |  |
| ref | C | T | C | G | A | G | T | A | C | A | T | G | T | G | T | T | A | C | G | C | G | C | A | A | G | A | T | C | C | A | C |  |
| BA.2.76 | C | T | T | G | G | G | T | A | C | A | T | G | A | C | T | T | G | C | G | T | G | T | A | C | C | T | C | C | C | A | C |  |
| BF.3 | C | T | T | G | G | G | T | A | C | A | T | G | A | C | G | G | A | T | T | C | A | C | G | A | G | A | T | T | T | G | T | #1348822 |
| BF.3 | T | G | C | A | A | C | - | - | - | - | - | - | T | G | G | G | A | T | T | C | A | C | G | A | G | A | T | T | T | G | T |  |
| mut |  |  |  |  |  |  |  |  |  |  |  |  |  |  |  |  |  |  |  |  |  |  |  |  |  |  |  |  |  |  |  |  |

|  |  |  |  |  |  |  |  |  |  |  |  |  |  |  |  |  |  |  |  |  |  |  |  |  |  |  |  |  |  |  |  |  |  |
| --- | --- | --- | --- | --- | --- | --- | --- | --- | --- | --- | --- | --- | --- | --- | --- | --- | --- | --- | --- | --- | --- | --- | --- | --- | --- | --- | --- | --- | --- | --- | --- | --- | --- |
| pos | 686 | 687 | 688 | 689 | 690 | 691 | 692 | 693 | 694 | 1627 | 8629 | 10323 | 11782 | 17766 | 21570 | 22910 | 26529 | 26858 | 27038 | 27259 | 27382 | 27383 | 27384 | 27389 | 27625 | 27788 | 27889 | 28330 | 28435 | 28724 | 29358 | 29754 |  |
| ref | A | A | G | T | C | A | T | T | T | C | T | A | A | C | T | A | G | C | A | A | G | A | T | C | C | G | C | A | C | C | C | C |  |
| BA.4.1 | - | - | - | - | - | - | - | - | - | C | T | A | G | C | G | G | G | T | A | C | C | T | C | C | C | T | C | A | T | T | C | C |  |
| BA.5 | - | - | - | - | - | - | - | - | - | C | T | A | G | T | G | G | A | C | G | A | G | A | T | T | T | G | T | G | C | C | C | T | #1350045 |
| BF.8 | A | A | G | T | C | A | T | T | T | T | C | G | A | C | T | A | A | C | G | A | G | A | T | C | T | G | T | G | C | C | A | T |  |
| mut |  |  |  |  |  |  |  |  |  |  |  |  |  | C>T |  |  |  |  |  |  |  |  |  | C>T |  |  |  |  |  |  | A>C |  |  |

|  |  |  |  |  |  |  |  |  |  |  |  |  |  |  |  |  |  |  |  |  |  |  |  |  |  |  |  |  |  |  |  |  |
| --- | --- | --- | --- | --- | --- | --- | --- | --- | --- | --- | --- | --- | --- | --- | --- | --- | --- | --- | --- | --- | --- | --- | --- | --- | --- | --- | --- | --- | --- | --- | --- | --- |
| pos | 694 | 6033 | 6196 | 6594 | 9866 | 12160 | 12415 | 16329 | 19011 | 21765 | 21766 | 21767 | 21768 | 21769 | 21770 | 22917 | 23018 | 23040 | 23191 | 25314 | 26529 | 26858 | 27259 | 27382 | 27383 | 27384 | 27438 | 27889 | 29022 | 29666 | 29754 |  |
| ref | T | C | C | C | C | G | C | C | C | T | A | C | A | T | G | T | T | A | C | G | G | C | A | G | A | T | T | C | C | C | C |  |
| BA.2 | T | T | T | C | T | G | C | T | C | T | A | C | A | T | G | T | T | G | C | A | G | T | C | C | T | C | T | C | T | C | C |  |
| BA.5.1 | T | T | T | T | T | G | T | C | T | - | - | - | - | - | - | G | G | A | T | G | A | C | A | G | A | T | C | T | C | C | T | #1350338 |
| BA.5.1 | A | C | C | C | C | A | C | C | T | - | - | - | - | - | - | G | G | A | C | G | A | C | A | G | A | T | C | T | C | T | T |  |
| mut |  |  |  | C>T |  |  | C>T |  |  |  |  |  |  |  |  |  |  |  | C>T |  |  |  |  |  |  |  |  |  |  | T>C |  |  |

|  |  |  |  |  |  |  |  |  |  |  |  |  |
| --- | --- | --- | --- | --- | --- | --- | --- | --- | --- | --- | --- | --- |
| pos | 1627 | 4622 | 5653 | 6979 | 13458 | 17339 | 27038 | 27438 | 28310 | 28330 | 29666 |  |
| ref | C | G | T | T | C | C | A | T | C | A | C |  |
| BA.5.1.6 | C | A | C | T | C | C | A | C | C | A | T |  |
| BF.10 | C | A | C | T | T | T | G | T | T | G | C | #1351167 |
| BF.10 | T | G | T | G | C | T | G | T | T | G | C |  |
| mut |  |  |  |  | C>T |  |  |  |  |  |  |  |

|  |  |  |  |  |  |  |  |  |  |  |  |  |  |  |  |  |  |  |  |  |  |  |  |  |  |  |  |  |  |  |  |
| --- | --- | --- | --- | --- | --- | --- | --- | --- | --- | --- | --- | --- | --- | --- | --- | --- | --- | --- | --- | --- | --- | --- | --- | --- | --- | --- | --- | --- | --- | --- | --- |
| pos | 44 | 203 | 686 | 687 | 688 | 689 | 690 | 691 | 692 | 693 | 694 | 1453 | 1627 | 5826 | 6979 | 22110 | 23535 | 25460 | 26529 | 26858 | 27038 | 27259 | 27382 | 27383 | 27384 | 27788 | 27889 | 28330 | 28724 | 29754 |  |
| ref | C | C | A | A | G | T | C | A | T | T | T | T | C | C | T | A | A | C | G | C | A | A | G | A | T | G | C | A | C | C |  |
| BA.5.2.1 | C | C | G | C | G | C | C | G | C | T | A | T | T | T | G | T | A | C | A | C | G | A | G | A | T | G | T | G | C | T |  |
| BA.4 | C | T | G | C | G | C | C | G | C | T | A | C | C | C | T | A | G | T | G | T | A | C | C | T | C | T | C | A | T | C | #1356368 |
| BA.4 | T | T | - | - | - | - | - | - | - | - | - | C | C | C | T | A | G | C | G | T | A | C | C | T | C | T | C | A | T | C |  |
| mut |  | C>T |  |  |  |  |  |  |  |  |  |  |  |  |  |  |  | C>T |  |  |  |  |  |  |  |  |  |  |  |  |  |

|  |  |  |  |  |  |  |  |  |  |  |  |  |  |  |  |  |
| --- | --- | --- | --- | --- | --- | --- | --- | --- | --- | --- | --- | --- | --- | --- | --- | --- |
| pos | 219 | 1376 | 1627 | 1684 | 1931 | 2910 | 5869 | 10399 | 13019 | 13342 | 16726 | 25546 | 27038 | 28330 | 28681 |  |
| ref | G | C | C | C | C | C | C | A | C | C | C | C | A | A | G |  |
| BE.2 | A | T | C | C | A | T | T | G | C | T | C | T | A | A | T |  |
| BA.5.2.1 | A | T | C | T | A | T | C | A | T | C | T | C | G | G | G | #1356780 |
| BA.5.2.1 | G | C | T | C | C | C | C | A | T | C | T | C | G | G | G |  |
| mut |  |  |  | C>T |  |  | T>C |  |  |  |  |  |  |  |  |  |

|  |  |  |  |  |  |  |  |  |  |  |  |  |  |  |  |  |  |  |  |  |  |  |  |  |  |  |  |  |  |
| --- | --- | --- | --- | --- | --- | --- | --- | --- | --- | --- | --- | --- | --- | --- | --- | --- | --- | --- | --- | --- | --- | --- | --- | --- | --- | --- | --- | --- | --- |
| pos | 44 | 3620 | 3796 | 3927 | 4586 | 4816 | 5141 | 5183 | 9430 | 11399 | 12444 | 14171 | 15451 | 19999 | 22001 | 22016 | 22033 | 22190 | 22331 | 22577 | 22599 | 22792 | 22898 | 22942 | 23019 | 23040 | 25416 | 26275 |  |
| ref | C | G | C | C | C | T | A | C | C | A | A | T | G | G | A | T | C | A | G | G | G | C | G | T | T | A | C | A |  |
| BA.2 | C | A | C | C | C | T | A | C | C | A | A | T | G | T | A | T | C | A | G | G | G | T | G | T | T | G | C | A |  |
| BA.2 | C | A | C | C | C | A | A | C | T | T | G | C | A | G | G | C | A | G | A | C | C | C | A | G | C | A | T | G | #1357912 |
| BA.2.75.7 | T | G | T | T | T | T | G | T | C | A | G | C | A | G | G | C | A | G | A | C | C | C | A | G | C | A | T | G |  |
| mut |  |  |  |  |  | T>A |  |  | C>T | A>T |  |  |  |  |  |  |  |  |  |  |  |  |  |  |  |  |  |  |  |

|  |  |  |  |  |  |  |  |  |  |  |  |  |  |  |  |  |  |  |  |  |  |  |  |  |  |  |  |  |  |  |  |  |  |  |
| --- | --- | --- | --- | --- | --- | --- | --- | --- | --- | --- | --- | --- | --- | --- | --- | --- | --- | --- | --- | --- | --- | --- | --- | --- | --- | --- | --- | --- | --- | --- | --- | --- | --- | --- |
| pos | 686 | 687 | 688 | 689 | 690 | 691 | 692 | 693 | 694 | 943 | 1627 | 5284 | 8116 | 17449 | 21570 | 24007 | 26529 | 26858 | 27038 | 27259 | 27382 | 27383 | 27384 | 27494 | 27532 | 27788 | 27889 | 28330 | 28370 | 28371 | 28724 | 29580 | 29754 |  |
| ref | A | A | G | T | C | A | T | T | T | G | C | C | A | T | T | G | G | C | A | A | G | A | T | C | C | G | C | A | A | G | C | C | C |  |
| BA.5.2.1 | A | A | G | T | C | A | T | T | T | G | T | T | G | C | T | A | A | C | G | A | G | A | T | C | T | G | T | G | T | T | C | T | T |  |
| BA.5.2.1 | A | A | G | T | C | A | T | T | T | A | T | T | G | T | T | A | A | C | G | A | G | A | T | T | C | T | C | A | - | G | T | C | C | #1358271 |
| BA.4.1 | - | - | - | - | - | - | - | - | - | G | C | C | A | T | G | G | G | T | A | C | C | T | C | T | C | T | C | A | - | G | T | C | C |  |
| mut |  |  |  |  |  |  |  |  |  | G>A |  |  |  | C>T |  |  |  |  |  |  |  |  |  |  |  |  |  |  |  |  |  |  |  |  |

|  |  |  |  |  |  |  |  |  |  |  |  |  |  |  |  |  |  |  |  |  |  |  |  |  |  |  |  |  |  |  |  |  |  |  |
| --- | --- | --- | --- | --- | --- | --- | --- | --- | --- | --- | --- | --- | --- | --- | --- | --- | --- | --- | --- | --- | --- | --- | --- | --- | --- | --- | --- | --- | --- | --- | --- | --- | --- | --- |
| pos | 6548 | 9300 | 9866 | 12160 | 13515 | 17042 | 20425 | 21765 | 21766 | 21767 | 21768 | 21769 | 21770 | 22600 | 22917 | 23018 | 23040 | 25416 | 26529 | 26858 | 27214 | 27259 | 27382 | 27383 | 27384 | 27438 | 27889 | 29666 | 29729 | 29754 | 29757 | 29758 | 29759 |  |
| ref | C | G | C | G | C | A | A | T | A | C | A | T | G | A | T | T | A | C | G | C | G | A | G | A | T | T | C | C | T | C | G | T | G |  |
| BA.5.1.2 | C | G | C | A | C | A | T | - | - | - | - | - | - | A | G | G | A | C | A | C | A | A | G | A | T | C | T | T | T | T | G | T | G |  |
| BA.5.1.2 | C | G | C | A | T | G | T | - | - | - | - | - | - | A | G | G | A | C | A | C | A | A | G | A | T | C | T | T | C | C | A | C | C | #1358513 |
| BA.2 | T | A | T | G | C | A | A | T | A | C | A | T | G | T | T | T | G | T | G | T | G | C | C | T | C | T | C | C | C | C | A | C | C |  |
| mut |  |  |  |  | C>T | A>G |  |  |  |  |  |  |  |  |  |  |  |  |  |  |  |  |  |  |  |  |  |  |  |  |  |  |  |  |

|  |  |  |  |  |  |  |  |  |  |  |  |  |  |  |  |  |  |  |  |
| --- | --- | --- | --- | --- | --- | --- | --- | --- | --- | --- | --- | --- | --- | --- | --- | --- | --- | --- | --- |
| pos | 1627 | 1931 | 2710 | 10188 | 12310 | 13342 | 16616 | 18814 | 22894 | 24083 | 27012 | 27513 | 27800 | 28330 | 28681 | 29762 | 29769 | 29774 |  |
| ref | C | C | C | C | G | C | C | C | G | C | C | C | C | A | G | C | C | C |  |
| BE.4 | C | A | C | A | G | T | C | C | G | C | C | C | T | A | T | C | C | C |  |
| BA.5.2 | C | A | C | A | G | T | C | T | T | T | T | T | C | G | G | - | - | - | #1359331 |
| BA.5.2.24 | T | C | T | C | A | C | A | C | T | C | T | T | C | G | G | C | C | C |  |
| mut |  |  |  |  |  |  |  | C>T |  | C>T |  |  |  |  |  | C>- | C>- | C>- |  |

|  |  |  |  |  |  |  |  |  |  |  |  |  |  |  |  |  |
| --- | --- | --- | --- | --- | --- | --- | --- | --- | --- | --- | --- | --- | --- | --- | --- | --- |
| pos | 1567 | 1627 | 2232 | 5284 | 16323 | 18811 | 25614 | 27038 | 27218 | 27438 | 28330 | 28370 | 28371 | 28395 | 29666 |  |
| ref | C | C | C | C | C | G | C | A | A | T | A | A | G | G | C |  |
| BA.5.1.30 | A | C | C | C | T | A | T | A | G | C | A | - | G | G | T |  |
| BA.5.1.30 | A | C | C | C | T | A | T | A | G | C | G | T | T | A | C | #1360479 |
| BA.5.2.1 | C | T | T | T | C | G | C | G | A | T | G | T | T | A | C |  |
| mut |  |  |  |  |  |  |  |  |  |  |  |  |  |  |  |  |

|  |  |  |  |  |  |  |  |  |  |  |  |  |  |  |  |  |  |  |  |  |  |  |  |  |  |  |  |  |  |  |  |  |
| --- | --- | --- | --- | --- | --- | --- | --- | --- | --- | --- | --- | --- | --- | --- | --- | --- | --- | --- | --- | --- | --- | --- | --- | --- | --- | --- | --- | --- | --- | --- | --- | --- |
| pos | 686 | 687 | 688 | 689 | 690 | 691 | 692 | 693 | 694 | 1627 | 6067 | 8140 | 11454 | 14599 | 22599 | 23535 | 26529 | 26858 | 27038 | 27259 | 27382 | 27383 | 27384 | 27506 | 27788 | 27889 | 28330 | 28390 | 28724 | 29754 | 29762 |  |
| ref | A | A | G | T | C | A | T | T | T | C | T | C | C | C | G | A | G | C | A | A | G | A | T | G | G | C | A | A | C | C | C |  |
| BA.5.2.1 | A | A | G | T | C | A | T | T | T | T | C | C | C | C | G | A | A | C | G | A | G | A | T | C | G | T | G | A | C | T | C |  |
| BA.5.2.1 | A | A | G | T | C | A | T | T | T | T | C | C | T | C | G | A | A | C | G | A | G | A | T | C | G | C | A | G | T | C | T | #1360676 |
| BA.4.6.5 | - | - | - | - | - | - | - | - | - | C | T | T | C | T | C | G | G | T | A | C | C | T | C | G | T | C | A | G | T | C | C |  |
| mut |  |  |  |  |  |  |  |  |  |  |  |  | C>T |  |  |  |  |  |  |  |  |  |  |  |  |  |  |  |  |  | C>T |  |

|  |  |  |  |  |  |  |  |  |  |  |  |  |  |  |  |  |  |  |  |  |
| --- | --- | --- | --- | --- | --- | --- | --- | --- | --- | --- | --- | --- | --- | --- | --- | --- | --- | --- | --- | --- |
| pos | 1514 | 3685 | 3686 | 4067 | 4906 | 10279 | 10507 | 14396 | 14925 | 17402 | 17893 | 18957 | 19174 | 21077 | 22599 | 26753 | 27438 | 27482 | 29666 |  |
| ref | C | G | C | G | T | C | C | C | C | C | G | T | G | C | G | C | T | T | C |  |
| BA.5.1.18 | T | G | C | A | T | T | T | T | C | C | T | T | A | C | C | C | C | A | T |  |
| BA.5.1.18 | T | G | C | A | T | T | T | T | C | C | T | T | A | C | C | T | T | T | C | #1361403 |
| BA.5.6 | C | A | A | G | C | C | C | C | T | T | G | C | G | T | G | T | T | T | C |  |
| mut |  |  |  |  |  |  |  |  |  |  |  |  |  |  |  |  |  |  |  |  |

|  |  |  |  |  |  |  |  |  |  |  |  |  |  |  |  |  |  |  |  |  |  |  |  |  |  |  |  |  |  |  |  |  |  |  |  |  |  |  |  |  |  |  |  |  |  |  |  |  |  |
| --- | --- | --- | --- | --- | --- | --- | --- | --- | --- | --- | --- | --- | --- | --- | --- | --- | --- | --- | --- | --- | --- | --- | --- | --- | --- | --- | --- | --- | --- | --- | --- | --- | --- | --- | --- | --- | --- | --- | --- | --- | --- | --- | --- | --- | --- | --- | --- | --- | --- |
| pos | 1627 | 2326 | 3796 | 3859 | 3927 | 4586 | 5183 | 6532 | 6979 | 7764 | 9866 | 10030 | 12160 | 12444 | 13785 | 15451 | 15810 | 18583 | 18657 | 21765 | 21766 | 21767 | 21768 | 21769 | 21770 | 22001 | 22016 | 22033 | 22190 | 22331 | 22577 | 22898 | 22917 | 22942 | 23018 | 25416 | 26275 | 26529 | 26858 | 27038 | 27259 | 27382 | 27383 | 27384 | 27752 | 27889 | 28330 | 29754 |  |
| ref | C | T | C | A | C | C | C | G | T | C | C | C | G | A | A | G | C | G | C | T | A | C | A | T | G | A | T | C | A | G | G | G | T | T | T | C | A | G | C | A | A | G | A | T | C | C | A | C |  |
| BA.2.75.3 | C | T | T | A | T | T | T | G | T | C | T | C | G | G | A | A | C | A | C | T | A | C | A | T | G | G | C | A | G | A | C | A | T | G | T | T | G | G | T | A | C | C | T | C | C | C | A | C |  |
| BA.2.75 | C | C | T | A | T | T | T | G | T | T | T | C | G | G | G | A | C | A | C | T | A | C | A | T | G | G | C | A | G | A | C | A | T | G | T | T | G | A | C | G | A | G | A | T | T | T | G | T | #1361724 |
| BA.5.2.1 | T | T | C | G | C | C | C | A | G | C | C | T | A | A | A | G | T | G | T | - | - | - | - | - | - | A | T | C | A | G | G | G | G | T | G | C | A | A | C | G | A | G | A | T | T | T | G | T |  |
| mut |  | T>C |  |  |  |  |  |  |  | C>T |  |  |  |  | A>G |  |  |  |  |  |  |  |  |  |  |  |  |  |  |  |  |  |  |  |  |  |  |  |  |  |  |  |  |  |  |  |  |  |  |

|  |  |  |  |  |  |  |  |  |  |  |  |  |  |  |  |  |  |  |  |  |
| --- | --- | --- | --- | --- | --- | --- | --- | --- | --- | --- | --- | --- | --- | --- | --- | --- | --- | --- | --- | --- |
| pos | 1593 | 1627 | 5703 | 5704 | 5705 | 5707 | 6894 | 12310 | 16541 | 16616 | 19961 | 20892 | 21137 | 27012 | 27438 | 27513 | 28330 | 29554 | 29666 |  |
| ref | C | C | A | G | T | T | G | G | A | C | C | T | A | C | T | C | A | G | C |  |
| BA.5.1 | C | C | T | A | A | C | G | G | G | C | T | A | G | C | C | C | A | A | T |  |
| BA.5.2 | T | C | T | A | A | C | G | G | G | C | T | A | G | T | T | T | G | G | C | #1362089 |
| BA.5.2 | C | T | A | G | T | T | T | A | A | A | C | T | A | T | T | T | G | G | C |  |
| mut | C>T |  |  |  |  |  |  |  |  |  |  |  |  |  |  |  |  |  |  |  |

|  |  |  |  |  |  |  |  |  |  |  |  |  |  |  |  |  |  |  |  |  |  |  |  |  |  |  |  |  |  |  |  |  |  |  |  |  |  |  |  |  |
| --- | --- | --- | --- | --- | --- | --- | --- | --- | --- | --- | --- | --- | --- | --- | --- | --- | --- | --- | --- | --- | --- | --- | --- | --- | --- | --- | --- | --- | --- | --- | --- | --- | --- | --- | --- | --- | --- | --- | --- | --- |
| pos | 2378 | 2554 | 3601 | 4234 | 6548 | 8179 | 9300 | 9866 | 12160 | 18751 | 21765 | 21766 | 21767 | 21768 | 21769 | 21770 | 22600 | 22917 | 23018 | 23040 | 25416 | 25824 | 26529 | 26858 | 26882 | 27259 | 27382 | 27383 | 27384 | 27438 | 27889 | 28754 | 29614 | 29666 | 29729 | 29754 | 29757 | 29758 | 29759 |  |
| ref | T | A | A | C | C | G | G | C | G | A | T | A | C | A | T | G | A | T | T | A | C | C | G | C | C | A | G | A | T | T | C | C | C | C | T | C | G | T | G |  |
| BA.5.1 | A | A | G | T | C | G | G | C | A | G | - | - | - | - | - | - | A | G | G | A | C | C | A | C | T | A | G | A | T | C | T | T | T | T | T | T | G | T | G |  |
| BA.5.1 | A | G | G | T | C | T | G | C | A | G | - | - | - | - | - | - | A | G | G | A | C | T | A | C | T | A | G | A | T | C | T | T | T | T | C | C | A | C | C | #1362718 |
| BA.2 | T | A | A | C | T | G | A | T | G | A | T | A | C | A | T | G | T | T | T | G | T | C | G | T | C | C | C | T | C | T | C | C | C | C | C | C | A | C | C |  |
| mut |  | A>G |  |  |  | G>T |  |  |  |  |  |  |  |  |  |  |  |  |  |  |  | C>T |  |  |  |  |  |  |  |  |  |  |  |  |  |  |  |  |  |  |

|  |  |  |  |  |  |  |  |  |  |  |  |  |  |  |  |  |
| --- | --- | --- | --- | --- | --- | --- | --- | --- | --- | --- | --- | --- | --- | --- | --- | --- |
| pos | 4011 | 6551 | 9782 | 11962 | 17278 | 20594 | 21232 | 22033 | 22522 | 25047 | 25855 | 27438 | 28531 | 29032 | 29666 |  |
| ref | T | A | A | C | G | C | G | C | C | C | G | T | C | G | C |  |
| BA.5.1 | G | A | G | T | G | C | A | C | C | C | T | C | T | G | T |  |
| BA.5.8 | G | A | G | T | T | T | G | G | A | T | G | T | C | A | C | #1363811 |
| BA.5.8 | T | C | A | C | T | T | G | C | C | T | G | T | C | A | C |  |
| mut |  |  |  |  |  |  |  | C>G | C>A |  |  |  |  |  |  |  |

|  |  |  |  |  |  |  |  |  |  |  |  |  |  |  |  |  |  |
| --- | --- | --- | --- | --- | --- | --- | --- | --- | --- | --- | --- | --- | --- | --- | --- | --- | --- |
| pos | 222 | 4622 | 5512 | 5653 | 6195 | 8092 | 12008 | 17336 | 21789 | 22898 | 26252 | 27438 | 27552 | 28077 | 29666 | 29781 |  |
| ref | C | G | C | T | C | C | C | C | C | G | C | T | A | G | C | G |  |
| BA.5.5 | C | G | T | T | C | C | T | C | T | G | C | T | A | T | C | G |  |
| BA.5.1 | T | G | T | T | T | T | T | C | T | G | T | C | G | G | T | G | #1363896 |
| BA.5.1.6 | C | A | C | C | C | C | C | T | C | A | T | C | G | G | T | - |  |
| mut | C>T |  |  |  | C>T | C>T |  |  |  |  |  |  |  |  |  | ->G |  |

|  |  |  |  |  |  |  |  |  |  |  |  |  |  |  |
| --- | --- | --- | --- | --- | --- | --- | --- | --- | --- | --- | --- | --- | --- | --- |
| pos | 1627 | 5284 | 5512 | 12569 | 14547 | 15738 | 21789 | 23569 | 23638 | 27038 | 28330 | 28370 | 28371 |  |
| ref | C | C | C | G | G | C | C | T | C | A | A | A | G |  |
| BA.5.5 | C | C | T | A | G | T | T | C | T | A | A | - | G |  |
| BA.5.5 | C | C | T | A | G | T | T | C | T | G | G | T | T | #1365029 |
| BA.5.2.1 | T | T | C | G | A | C | C | T | C | G | G | T | T |  |
| mut |  |  |  |  |  |  |  |  |  |  |  |  |  |  |

|  |  |  |  |  |  |  |  |  |  |  |  |  |  |  |  |  |  |  |
| --- | --- | --- | --- | --- | --- | --- | --- | --- | --- | --- | --- | --- | --- | --- | --- | --- | --- | --- |
| pos | 1627 | 4690 | 4930 | 5665 | 6979 | 12439 | 19866 | 20468 | 21520 | 22427 | 22879 | 26122 | 26256 | 27038 | 27438 | 28330 | 29666 |  |
| ref | C | T | T | G | T | C | T | C | G | G | C | G | C | A | T | A | C |  |
| BF.1 | T | C | T | G | G | C | C | C | T | G | T | A | C | G | T | G | C |  |
| BA.5.1 | T | C | T | A | G | T | C | C | T | G | C | G | T | A | C | A | T | #1365526 |
| BA.5.1.3 | C | T | C | G | T | C | T | T | G | A | C | G | T | A | C | A | T |  |
| mut |  |  |  | G>A |  | C>T |  |  |  |  |  |  |  |  |  |  |  |  |

|  |  |  |  |  |  |  |  |  |  |  |  |  |  |  |  |  |  |
| --- | --- | --- | --- | --- | --- | --- | --- | --- | --- | --- | --- | --- | --- | --- | --- | --- | --- |
| pos | 1627 | 2973 | 4906 | 5284 | 5829 | 14925 | 17402 | 18957 | 20483 | 21077 | 26726 | 27038 | 28209 | 28330 | 28370 | 28371 |  |
| ref | C | C | T | C | A | C | C | T | C | C | T | A | G | A | A | G |  |
| BA.5.2.1 | T | T | T | T | G | C | C | T | C | C | T | G | T | G | T | T |  |
| BA.5.6 | T | T | T | T | G | T | T | C | T | T | C | A | G | A | - | G | #1366397 |
| BA.5.6 | C | C | C | C | A | T | T | C | C | T | C | A | G | A | - | G |  |
| mut |  |  |  |  |  |  |  |  | C>T |  |  |  |  |  |  |  |  |

|  |  |  |  |  |  |  |  |  |  |  |  |  |  |  |  |  |  |  |  |  |  |  |
| --- | --- | --- | --- | --- | --- | --- | --- | --- | --- | --- | --- | --- | --- | --- | --- | --- | --- | --- | --- | --- | --- | --- |
| pos | 1931 | 3411 | 3875 | 5672 | 6349 | 11866 | 12068 | 16700 | 16935 | 21724 | 22458 | 22792 | 23323 | 25534 | 25537 | 27438 | 27465 | 28681 | 29114 | 29614 | 29666 |  |
| ref | C | C | G | C | G | C | C | G | G | G | C | C | T | G | G | T | T | G | C | C | C |  |
| BE.1 | A | C | A | C | G | C | T | G | A | C | C | T | C | T | T | T | T | T | C | C | C |  |
| BA.5.1 | A | C | A | C | G | C | T | G | A | C | T | T | T | G | G | C | C | G | A | T | T | #1366682 |
| BA.5.1 | C | T | G | T | A | T | C | T | G | G | T | C | T | G | G | C | T | G | C | C | T |  |
| mut |  |  |  |  |  |  |  |  |  |  |  | C>T |  |  |  |  | T>C |  | C>A | C>T |  |  |

|  |  |  |  |  |  |  |  |  |  |  |  |  |  |  |  |  |  |
| --- | --- | --- | --- | --- | --- | --- | --- | --- | --- | --- | --- | --- | --- | --- | --- | --- | --- |
| pos | 4794 | 6979 | 8278 | 11036 | 12310 | 12325 | 16616 | 17339 | 19983 | 22786 | 23380 | 27012 | 27038 | 27506 | 27513 | 28310 |  |
| ref | C | T | T | C | G | C | C | C | C | A | C | C | A | G | C | C |  |
| BF.10 | T | G | T | C | G | T | C | T | C | A | C | C | G | G | C | T |  |
| BA.5.2 | T | G | T | T | G | T | A | C | C | C | T | T | A | T | T | C | #1370716 |
| BA.5.2 | C | T | C | C | A | C | A | C | T | C | T | T | A | G | T | C |  |
| mut |  |  |  | C>T |  |  |  |  | T>C |  |  |  |  | G>T |  |  |  |

|  |  |  |  |  |  |  |  |  |  |  |  |  |  |  |  |  |  |  |
| --- | --- | --- | --- | --- | --- | --- | --- | --- | --- | --- | --- | --- | --- | --- | --- | --- | --- | --- |
| pos | 492 | 2039 | 4561 | 5812 | 8755 | 14724 | 17964 | 19518 | 21789 | 22639 | 25903 | 26153 | 26840 | 27297 | 27438 | 28063 | 29666 |  |
| ref | C | A | G | C | A | C | G | G | C | C | T | G | G | C | T | T | C |  |
| BA.5.1 | C | G | G | T | G | C | T | T | C | T | G | G | G | C | C | T | T |  |
| BA.5 | C | G | G | T | G | T | T | T | C | T | T | T | A | T | T | C | C | #1370927 |
| BA.5.5 | T | A | A | C | A | C | G | G | T | C | T | G | A | C | T | C | C |  |
| mut |  |  |  |  |  | C>T |  |  |  |  |  | G>T |  | C>T |  |  |  |  |

|  |  |  |  |  |  |  |  |  |  |  |  |  |  |  |  |  |  |
| --- | --- | --- | --- | --- | --- | --- | --- | --- | --- | --- | --- | --- | --- | --- | --- | --- | --- |
| pos | 376 | 1627 | 1931 | 5284 | 6968 | 8251 | 15621 | 16915 | 16935 | 18826 | 27038 | 28330 | 28370 | 28371 | 28681 | 29218 |  |
| ref | G | C | C | C | C | C | T | C | G | G | A | A | A | G | G | C |  |
| BA.5.2.1 | G | T | C | T | T | C | T | C | G | G | G | G | T | T | G | C |  |
| BE.1.4 | G | T | C | T | T | T | C | T | A | A | A | A | - | G | T | T | #1370930 |
| BE.1.4 | A | C | A | C | C | T | C | T | A | A | A | A | - | G | T | T |  |
| mut |  |  |  |  |  |  |  |  |  |  |  |  |  |  |  |  |  |

|  |  |  |  |  |  |  |  |  |  |  |  |  |  |  |  |  |  |  |  |  |  |  |
| --- | --- | --- | --- | --- | --- | --- | --- | --- | --- | --- | --- | --- | --- | --- | --- | --- | --- | --- | --- | --- | --- | --- |
| pos | 1514 | 1627 | 5284 | 6316 | 7484 | 10279 | 10507 | 14396 | 17893 | 18163 | 19174 | 22498 | 22599 | 27038 | 27179 | 27438 | 27482 | 28330 | 28370 | 28371 | 29666 |  |
| ref | C | C | C | A | T | C | C | C | G | A | G | C | G | A | G | T | T | A | A | G | C |  |
| BA.5.2.1 | C | T | T | A | T | C | C | C | G | G | G | C | G | G | A | T | T | G | T | T | C |  |
| BA.5.1.18 | C | T | C | A | T | C | C | C | T | T | A | T | C | A | G | C | A | A | - | G | T | #1370994 |
| BA.5.1.18 | T | C | C | G | C | T | T | T | T | T | A | T | C | A | G | C | A | A | - | G | T |  |
| mut |  |  | T>C |  |  |  |  |  |  |  |  |  |  |  |  |  |  |  |  |  |  |  |

|  |  |  |  |  |  |  |  |  |  |  |  |  |  |  |  |  |  |  |  |  |  |  |  |
| --- | --- | --- | --- | --- | --- | --- | --- | --- | --- | --- | --- | --- | --- | --- | --- | --- | --- | --- | --- | --- | --- | --- | --- |
| pos | 44 | 686 | 687 | 688 | 689 | 690 | 691 | 692 | 693 | 694 | 1931 | 3875 | 5629 | 12068 | 16935 | 18712 | 22792 | 25534 | 25537 | 27438 | 28681 | 29666 |  |
| ref | C | A | A | G | T | C | A | T | T | T | C | G | G | C | G | G | C | G | G | T | G | C |  |
| BA.5.1 | C | G | C | G | C | C | G | C | T | A | C | G | G | C | G | G | C | G | G | C | G | T |  |
| BE.1 | C | G | C | G | C | C | G | C | T | A | A | A | A | T | A | A | T | T | T | T | T | C | #1372032 |
| BE.1 | T | - | - | - | - | - | - | - | - | - | A | A | G | T | A | A | T | T | T | T | T | C |  |
| mut |  |  |  |  |  |  |  |  |  |  |  |  | G>A |  |  |  |  |  |  |  |  |  |  |

|  |  |  |  |  |  |  |  |  |  |  |  |  |  |  |  |  |  |  |  |  |  |  |  |  |  |  |  |  |  |  |  |  |  |  |  |
| --- | --- | --- | --- | --- | --- | --- | --- | --- | --- | --- | --- | --- | --- | --- | --- | --- | --- | --- | --- | --- | --- | --- | --- | --- | --- | --- | --- | --- | --- | --- | --- | --- | --- | --- | --- |
| pos | 44 | 1263 | 1627 | 1966 | 2245 | 5284 | 9866 | 12160 | 13165 | 14593 | 21765 | 21766 | 21767 | 21768 | 21769 | 21770 | 22812 | 22917 | 23018 | 23040 | 23248 | 26529 | 26858 | 27038 | 27259 | 27382 | 27383 | 27384 | 27889 | 28330 | 28370 | 28371 | 29614 | 29754 |  |
| ref | C | C | C | A | C | C | C | G | A | G | T | A | C | A | T | G | A | T | T | A | C | G | C | A | A | G | A | T | C | A | A | G | C | C |  |
| BA.2.18 | C | T | C | A | T | C | T | G | A | G | T | A | C | A | T | G | C | T | T | G | T | G | T | A | C | C | T | C | C | A | - | G | C | C |  |
| BF.26 | C | T | C | G | T | C | T | G | G | A | - | - | - | - | - | - | A | G | G | A | C | A | C | G | A | G | A | T | T | G | T | T | T | T | #1373146 |
| BF.26 | T | C | T | A | C | T | C | A | G | A | - | - | - | - | - | - | A | G | G | A | C | A | C | G | A | G | A | T | T | G | T | T | C | T |  |
| mut |  |  |  | A>G |  |  |  |  |  |  |  |  |  |  |  |  |  |  |  |  |  |  |  |  |  |  |  |  |  |  |  |  | C>T |  |  |

|  |  |  |  |  |  |  |  |  |  |  |  |  |  |  |  |  |  |  |  |  |  |  |  |  |  |  |  |  |  |  |  |  |  |  |  |  |  |  |  |  |
| --- | --- | --- | --- | --- | --- | --- | --- | --- | --- | --- | --- | --- | --- | --- | --- | --- | --- | --- | --- | --- | --- | --- | --- | --- | --- | --- | --- | --- | --- | --- | --- | --- | --- | --- | --- | --- | --- | --- | --- | --- |
| pos | 1319 | 1627 | 4893 | 5806 | 6548 | 6979 | 8616 | 9300 | 9866 | 11986 | 12160 | 21114 | 21765 | 21766 | 21767 | 21768 | 21769 | 21770 | 22600 | 22917 | 23018 | 23040 | 25416 | 26529 | 26858 | 27038 | 27259 | 27382 | 27383 | 27384 | 27400 | 27889 | 28087 | 28330 | 29729 | 29754 | 29757 | 29758 | 29759 |  |
| ref | G | C | C | C | C | T | C | G | C | C | G | C | T | A | C | A | T | G | A | T | T | A | C | G | C | A | A | G | A | T | A | C | C | A | T | C | G | T | G |  |
| BF.27 | A | T | T | C | C | A | T | G | C | C | A | T | - | - | - | - | - | - | A | G | G | A | C | A | C | G | A | G | A | T | G | T | A | G | T | T | G | T | G |  |
| BF.27 | A | T | T | T | C | A | T | G | C | T | A | T | - | - | - | - | - | - | A | G | G | A | C | A | C | G | A | G | A | T | G | T | A | G | C | C | A | C | C | #1373412 |
| BA.2 | G | C | C | C | T | T | C | A | T | C | G | C | T | A | C | A | T | G | T | T | T | G | T | G | T | A | C | C | T | C | A | C | C | A | C | C | A | C | C |  |
| mut |  |  |  | C>T |  |  |  |  |  | C>T |  |  |  |  |  |  |  |  |  |  |  |  |  |  |  |  |  |  |  |  |  |  |  |  |  |  |  |  |  |  |

|  |  |  |  |  |  |  |  |  |  |  |  |  |  |  |  |  |
| --- | --- | --- | --- | --- | --- | --- | --- | --- | --- | --- | --- | --- | --- | --- | --- | --- |
| pos | 1627 | 5284 | 10798 | 12747 | 14547 | 16686 | 17285 | 19152 | 27038 | 27438 | 28330 | 28370 | 28371 | 28603 | 29666 |  |
| ref | C | C | C | C | G | T | C | C | A | T | A | A | G | C | C |  |
| BA.5.1.22 | C | C | T | T | G | C | T | A | A | C | A | - | G | T | T |  |
| BA.5 | C | C | T | T | G | C | T | A | G | T | G | T | T | C | C | #1374207 |
| BA.5.2.1 | T | T | C | C | A | T | C | C | G | T | G | T | T | C | C |  |
| mut |  |  |  |  |  |  |  |  |  |  |  |  |  |  |  |  |

|  |  |  |  |  |  |  |  |  |  |  |  |  |  |  |  |  |  |  |  |  |  |  |  |  |  |  |  |  |  |
| --- | --- | --- | --- | --- | --- | --- | --- | --- | --- | --- | --- | --- | --- | --- | --- | --- | --- | --- | --- | --- | --- | --- | --- | --- | --- | --- | --- | --- | --- |
| pos | 227 | 3796 | 3927 | 4586 | 5183 | 5192 | 6548 | 9300 | 12444 | 15451 | 22001 | 22016 | 22033 | 22190 | 22331 | 22577 | 22599 | 22600 | 22898 | 22942 | 23019 | 23040 | 25157 | 26275 | 29729 | 29757 | 29758 | 29759 |  |
| ref | G | C | C | C | C | C | C | G | A | G | A | T | C | A | G | G | G | A | G | T | T | A | G | A | T | G | T | G |  |
| BA.2.75.2 | G | T | T | T | T | T | C | G | G | A | G | C | A | G | A | C | C | A | A | G | C | A | A | G | T | G | T | G |  |
| BA.2.75.2 | A | T | T | T | T | T | C | G | G | A | G | C | A | G | A | C | C | A | A | G | C | A | A | G | C | A | C | C | #1375056 |
| BA.2 | G | C | C | C | C | C | T | A | A | G | A | T | C | A | G | G | G | T | G | T | T | G | G | A | C | A | C | C |  |
| mut | G>A |  |  |  |  |  |  |  |  |  |  |  |  |  |  |  |  |  |  |  |  |  |  |  |  |  |  |  |  |

|  |  |  |  |  |  |  |  |  |  |  |  |  |  |  |  |  |  |  |
| --- | --- | --- | --- | --- | --- | --- | --- | --- | --- | --- | --- | --- | --- | --- | --- | --- | --- | --- |
| pos | 219 | 1376 | 1627 | 1931 | 2910 | 3838 | 5284 | 5869 | 10399 | 13342 | 16376 | 25546 | 27038 | 28330 | 28370 | 28371 | 28681 |  |
| ref | G | C | C | C | C | G | C | C | A | C | C | C | A | A | A | G | G |  |
| BA.5.2.1 | G | C | T | C | C | A | T | C | A | C | C | C | G | G | T | T | G |  |
| BA.5.3.1 | G | C | T | C | C | A | T | C | A | T | T | T | A | A | - | G | T | #1375130 |
| BE.2 | A | T | C | A | T | G | C | T | G | T | T | T | A | A | - | G | T |  |
| mut |  |  |  |  |  |  |  |  |  |  |  |  |  |  |  |  |  |  |

|  |  |  |  |  |  |  |  |  |  |  |  |  |  |  |  |  |  |  |  |  |  |  |  |
| --- | --- | --- | --- | --- | --- | --- | --- | --- | --- | --- | --- | --- | --- | --- | --- | --- | --- | --- | --- | --- | --- | --- | --- |
| pos | 44 | 389 | 686 | 687 | 688 | 689 | 690 | 691 | 692 | 693 | 694 | 961 | 8140 | 10771 | 14353 | 14599 | 22599 | 25315 | 25644 | 26621 | 26690 | 28791 |  |
| ref | C | G | A | A | G | T | C | A | T | T | T | T | C | T | A | C | G | A | G | T | G | C |  |
| BA.4.6 | C | G | - | - | - | - | - | - | - | - | - | C | T | T | A | T | C | A | G | T | G | T |  |
| BA.4 | C | A | G | C | G | C | C | G | C | T | A | C | C | C | C | C | G | G | T | C | T | C | #1376015 |
| BA.4 | T | G | - | - | - | - | - | - | - | - | - | T | C | T | A | C | G | G | T | C | T | C |  |
| mut |  | G>A | ->G | ->C | ->G | ->C | ->C | ->G | ->C | ->T | ->A |  |  | T>C | A>C |  |  |  |  |  |  |  |  |

|  |  |  |  |  |  |  |  |  |  |  |  |  |  |  |
| --- | --- | --- | --- | --- | --- | --- | --- | --- | --- | --- | --- | --- | --- | --- |
| pos | 44 | 518 | 686 | 687 | 689 | 691 | 692 | 5194 | 5826 | 6979 | 12310 | 22110 | 27038 |  |
| ref | C | A | A | A | T | A | T | G | C | T | G | A | A |  |
| BA.5.2.1 | C | A | G | C | C | G | C | G | T | G | G | T | G |  |
| BA.5.2 | C | G | G | C | C | G | C | T | C | T | A | A | A | #1376031 |
| BA.5.2 | T | G | A | A | T | A | T | T | C | T | A | A | A |  |
| mut |  | A>G |  |  |  |  |  |  |  |  |  |  |  |  |

|  |  |  |  |  |  |  |  |  |  |  |  |  |  |  |  |  |  |  |  |  |  |  |  |  |  |  |  |  |  |  |  |  |  |  |  |  |  |  |
| --- | --- | --- | --- | --- | --- | --- | --- | --- | --- | --- | --- | --- | --- | --- | --- | --- | --- | --- | --- | --- | --- | --- | --- | --- | --- | --- | --- | --- | --- | --- | --- | --- | --- | --- | --- | --- | --- | --- |
| pos | 400 | 686 | 687 | 688 | 689 | 690 | 691 | 692 | 693 | 694 | 1627 | 1779 | 8140 | 10323 | 13620 | 14599 | 22599 | 23535 | 23587 | 25352 | 26529 | 26750 | 26858 | 27038 | 27259 | 27382 | 27383 | 27384 | 27788 | 27889 | 28087 | 28254 | 28330 | 28724 | 29227 | 29358 | 29754 |  |
| ref | T | A | A | G | T | C | A | T | T | T | C | C | C | A | C | C | G | A | G | G | G | C | C | A | A | G | A | T | G | C | C | A | A | C | G | C | C |  |
| BA.4.6 | T | - | - | - | - | - | - | - | - | - | C | T | T | A | C | T | C | G | G | T | G | C | T | A | C | C | T | C | T | C | C | A | A | T | G | C | C |  |
| BA.5.2.1 | G | - | - | - | - | - | - | - | - | - | C | T | T | A | C | T | C | G | C | G | A | A | C | G | A | G | A | T | G | T | A | - | G | C | A | C | T | #1376425 |
| BF.8 | T | A | A | G | T | C | A | T | T | T | T | C | C | G | T | C | G | A | C | G | A | C | C | G | A | G | A | T | G | T | C | A | G | C | G | A | T |  |
| mut | T>G |  |  |  |  |  |  |  |  |  |  |  |  |  |  |  |  |  |  |  |  | C>A |  |  |  |  |  |  |  |  | C>A | A>- |  |  | G>A | A>C |  |  |

|  |  |  |  |  |  |  |  |  |  |  |  |  |  |  |  |  |  |  |  |  |  |  |  |  |  |  |  |  |  |  |  |  |  |  |  |  |  |  |  |  |  |  |  |  |
| --- | --- | --- | --- | --- | --- | --- | --- | --- | --- | --- | --- | --- | --- | --- | --- | --- | --- | --- | --- | --- | --- | --- | --- | --- | --- | --- | --- | --- | --- | --- | --- | --- | --- | --- | --- | --- | --- | --- | --- | --- | --- | --- | --- | --- |
| pos | 1627 | 3796 | 3927 | 4586 | 5183 | 5192 | 6979 | 9866 | 12160 | 12444 | 15451 | 21765 | 21766 | 21767 | 21768 | 21769 | 21770 | 22001 | 22016 | 22033 | 22190 | 22331 | 22577 | 22599 | 22898 | 22917 | 22942 | 23018 | 23019 | 24620 | 25157 | 25416 | 26275 | 26529 | 26858 | 27038 | 27259 | 27382 | 27383 | 27384 | 27889 | 28330 | 29754 |  |
| ref | C | C | C | C | C | C | T | C | G | A | G | T | A | C | A | T | G | A | T | C | A | G | G | G | G | T | T | T | T | G | G | C | A | G | C | A | A | G | A | T | C | A | C |  |
| BA.2.75.2 | C | T | T | T | T | T | T | T | G | G | A | T | A | C | A | T | G | G | C | A | G | A | C | C | A | T | G | T | C | G | A | T | G | G | T | A | C | C | T | C | C | A | C |  |
| XBD | C | T | T | T | T | T | T | T | G | G | A | T | A | C | A | T | G | G | C | A | G | A | C | C | A | T | G | T | C | T | G | C | A | A | C | G | A | G | A | T | T | G | T | #1378208 |
| BA.5.2.1 | T | C | C | C | C | C | G | C | A | A | G | - | - | - | - | - | - | A | T | C | A | G | G | G | G | G | T | G | T | T | G | C | A | A | C | G | A | G | A | T | T | G | T |  |
| mut |  |  |  |  |  |  |  |  |  |  |  |  |  |  |  |  |  |  |  |  |  |  |  |  |  |  |  |  |  |  |  |  |  |  |  |  |  |  |  |  |  |  |  |  |

|  |  |  |  |  |  |  |  |  |  |  |  |  |  |  |  |  |  |  |  |  |  |  |  |  |  |  |  |
| --- | --- | --- | --- | --- | --- | --- | --- | --- | --- | --- | --- | --- | --- | --- | --- | --- | --- | --- | --- | --- | --- | --- | --- | --- | --- | --- | --- |
| pos | 44 | 222 | 3796 | 3927 | 4586 | 5183 | 7528 | 12444 | 12789 | 13730 | 15451 | 22001 | 22016 | 22033 | 22190 | 22331 | 22577 | 22599 | 22898 | 22942 | 23019 | 23040 | 25157 | 25416 | 26275 | 28079 |  |
| ref | C | C | C | C | C | C | C | A | C | C | G | A | T | C | A | G | G | G | G | T | T | A | G | C | A | G |  |
| BA.2.1 | C | T | C | C | C | C | C | A | T | T | G | A | T | C | A | G | G | G | G | T | T | G | G | C | A | G |  |
| XBH | C | C | C | C | C | C | T | A | C | T | G | G | C | A | G | A | C | C | A | G | C | A | A | T | G | T | #1379419 |
| BA.2.75.2 | T | C | T | T | T | T | C | G | C | C | A | G | C | A | G | A | C | C | A | G | C | A | A | T | G | G |  |
| mut |  | T>C |  |  |  |  | C>T |  | T>C |  |  |  |  |  |  |  |  |  |  |  |  |  |  |  |  | G>T |  |

|  |  |  |  |  |  |  |  |  |  |  |  |  |  |  |  |  |  |  |  |  |  |  |  |  |  |  |  |  |  |  |  |  |  |  |
| --- | --- | --- | --- | --- | --- | --- | --- | --- | --- | --- | --- | --- | --- | --- | --- | --- | --- | --- | --- | --- | --- | --- | --- | --- | --- | --- | --- | --- | --- | --- | --- | --- | --- | --- |
| pos | 686 | 687 | 688 | 689 | 690 | 691 | 692 | 693 | 694 | 1627 | 3838 | 5008 | 5284 | 8140 | 14599 | 15480 | 22599 | 23535 | 26529 | 26618 | 26858 | 27038 | 27259 | 27382 | 27383 | 27384 | 27788 | 27889 | 28330 | 28370 | 28371 | 28739 | 29754 |  |
| ref | A | A | G | T | C | A | T | T | T | C | G | G | C | C | C | C | G | A | G | T | C | A | A | G | A | T | G | C | A | A | G | G | C |  |
| BA.5.2.1 | A | A | G | T | C | A | T | T | T | T | A | T | T | C | C | C | G | A | A | T | C | G | A | G | A | T | G | T | G | T | T | G | T |  |
| BA.4 | A | A | G | T | C | A | T | T | T | T | A | T | T | C | C | T | C | G | G | C | T | A | C | C | T | C | T | C | A | - | G | T | C | #1379763 |
| BA.4.6 | - | - | - | - | - | - | - | - | - | C | G | G | C | T | T | C | C | G | G | C | T | A | C | C | T | C | T | C | A | - | G | T | C |  |
| mut |  |  |  |  |  |  |  |  |  |  |  |  |  |  |  | C>T | G>C |  |  |  |  |  |  |  |  |  |  |  |  |  |  |  |  |  |

|  |  |  |  |  |  |  |  |  |  |  |  |  |  |  |  |  |  |  |  |  |  |  |  |  |  |  |  |  |  |  |  |  |  |  |
| --- | --- | --- | --- | --- | --- | --- | --- | --- | --- | --- | --- | --- | --- | --- | --- | --- | --- | --- | --- | --- | --- | --- | --- | --- | --- | --- | --- | --- | --- | --- | --- | --- | --- | --- |
| pos | 44 | 1627 | 3859 | 4933 | 6979 | 7735 | 12160 | 18657 | 19955 | 20055 | 20160 | 21765 | 21766 | 21767 | 21768 | 21769 | 21770 | 21991 | 21992 | 21993 | 22917 | 23018 | 23040 | 26529 | 26858 | 27038 | 27259 | 27382 | 27383 | 27384 | 27889 | 28330 | 29754 |  |
| ref | C | C | A | T | T | C | G | C | C | A | G | T | A | C | A | T | G | T | T | A | T | T | A | G | C | A | A | G | A | T | C | A | C |  |
| BA.5.2.1 | T | T | G | T | G | T | A | C | T | G | G | - | - | - | - | - | - | T | T | A | G | G | A | A | C | G | A | G | A | T | T | G | T |  |
| BA.4 | T | T | G | C | G | T | A | T | T | G | A | - | - | - | - | - | - | - | - | - | G | G | A | A | C | G | A | C | T | C | C | A | C | #1381228 |
| BA.2 | C | C | A | T | T | C | G | C | C | A | G | T | A | C | A | T | G | T | T | A | T | T | G | G | T | A | C | C | T | C | C | A | C |  |
| mut |  |  |  | T>C |  |  |  | C>T |  |  | G>A |  |  |  |  |  |  | T>- | T>- | A>- |  |  |  |  |  |  |  |  |  |  |  |  |  |  |

|  |  |  |  |  |  |  |  |  |  |  |  |  |  |  |  |  |  |  |
| --- | --- | --- | --- | --- | --- | --- | --- | --- | --- | --- | --- | --- | --- | --- | --- | --- | --- | --- |
| pos | 2375 | 3259 | 3859 | 6979 | 7675 | 7909 | 8984 | 12310 | 15126 | 16616 | 19851 | 24199 | 25479 | 27012 | 27038 | 27513 | 27870 |  |
| ref | A | G | A | T | G | A | G | G | C | C | G | G | T | C | A | C | G |  |
| BA.5.2.1 | A | G | G | G | T | A | G | G | T | C | A | G | T | C | G | C | G |  |
| BA.5.2 | G | G | G | G | T | G | G | A | C | A | G | A | C | T | A | T | T | #1381343 |
| BA.5.2 | A | T | A | T | G | A | A | A | C | A | G | G | C | T | A | T | T |  |
| mut | A>G |  |  |  |  | A>G |  |  |  |  |  | G>A |  |  |  |  |  |  |

|  |  |  |  |  |  |  |  |  |  |  |  |  |  |  |  |  |  |  |  |  |  |  |  |  |  |  |  |  |  |  |  |  |  |  |
| --- | --- | --- | --- | --- | --- | --- | --- | --- | --- | --- | --- | --- | --- | --- | --- | --- | --- | --- | --- | --- | --- | --- | --- | --- | --- | --- | --- | --- | --- | --- | --- | --- | --- | --- |
| pos | 337 | 600 | 686 | 687 | 688 | 689 | 690 | 691 | 692 | 693 | 694 | 936 | 1627 | 2334 | 5974 | 8140 | 12310 | 14599 | 22599 | 23535 | 26529 | 26858 | 26934 | 27259 | 27382 | 27383 | 27384 | 27788 | 27889 | 28330 | 28724 | 29686 | 29754 |  |
| ref | C | G | A | A | G | T | C | A | T | T | T | C | C | C | C | C | G | C | G | A | G | C | C | A | G | A | T | G | C | A | C | C | C |  |
| BA.4.6 | T | A | - | - | - | - | - | - | - | - | - | C | C | T | T | T | G | T | C | G | G | T | C | C | C | T | C | T | C | A | T | C | C |  |
| BA.5 | T | A | - | - | - | - | - | - | - | - | - | C | C | T | T | T | G | T | C | G | A | C | G | A | G | A | T | G | T | G | C | T | T | #1381416 |
| BA.5.2 | C | G | A | A | G | T | C | A | T | T | T | T | T | C | C | C | A | C | G | A | A | C | C | A | G | A | T | G | T | G | C | T | T |  |
| mut |  |  |  |  |  |  |  |  |  |  |  |  |  |  |  |  |  |  |  |  |  |  | C>G |  |  |  |  |  |  |  |  |  |  |  |

|  |  |  |  |  |  |  |  |  |  |  |  |  |  |  |  |  |
| --- | --- | --- | --- | --- | --- | --- | --- | --- | --- | --- | --- | --- | --- | --- | --- | --- |
| pos | 1627 | 1931 | 7945 | 8606 | 9286 | 16935 | 20148 | 22337 | 24919 | 26840 | 27038 | 27297 | 28330 | 28681 | 29167 |  |
| ref | C | C | C | C | C | G | C | A | C | G | A | C | A | G | C |  |
| BE.1 | C | A | C | C | T | A | T | A | C | A | A | T | A | T | T |  |
| BA.5 | C | A | C | C | T | A | T | A | T | G | G | C | G | G | C | #1382090 |
| BF.4 | T | C | T | A | C | G | C | G | T | G | G | C | G | G | C |  |
| mut |  |  |  |  |  |  |  |  |  |  |  |  |  |  |  |  |

|  |  |  |  |  |  |  |  |  |  |  |  |  |  |  |
| --- | --- | --- | --- | --- | --- | --- | --- | --- | --- | --- | --- | --- | --- | --- |
| pos | 1627 | 6894 | 12310 | 14922 | 15895 | 16616 | 17285 | 18084 | 27012 | 27438 | 27513 | 28330 | 29666 |  |
| ref | C | G | G | C | C | C | C | C | C | T | C | A | C |  |
| BA.5.1.22 | C | G | G | A | A | C | T | T | C | C | C | A | T |  |
| BA.5.2 | C | G | G | A | A | C | T | T | T | T | T | G | C | #1382185 |
| BA.5.2 | T | T | A | C | C | A | C | C | T | T | T | G | C |  |
| mut |  |  |  |  |  |  |  |  |  |  |  |  |  |  |

|  |  |  |  |  |  |  |  |  |  |  |  |  |  |  |  |  |  |  |  |  |
| --- | --- | --- | --- | --- | --- | --- | --- | --- | --- | --- | --- | --- | --- | --- | --- | --- | --- | --- | --- | --- |
| pos | 919 | 1627 | 3153 | 4133 | 12310 | 13868 | 16616 | 16897 | 21040 | 22286 | 23683 | 24745 | 25587 | 27012 | 27438 | 27463 | 27513 | 28330 | 29666 |  |
| ref | A | C | G | G | G | A | C | G | A | C | C | C | C | C | T | G | C | A | C |  |
| BA.5.1 | G | C | G | G | G | G | C | T | G | T | T | C | T | C | C | T | C | A | T |  |
| BA.5.2 | G | C | G | G | G | G | C | T | G | T | T | T | C | T | T | G | T | G | T | #1382683 |
| BA.5.2 | A | T | A | A | A | A | A | G | A | C | C | T | C | T | T | G | T | G | C |  |
| mut |  |  |  |  |  |  |  |  |  |  |  |  |  |  |  |  |  |  | C>T |  |

|  |  |  |  |  |  |  |  |  |  |  |  |  |  |  |  |  |  |  |  |
| --- | --- | --- | --- | --- | --- | --- | --- | --- | --- | --- | --- | --- | --- | --- | --- | --- | --- | --- | --- |
| pos | 6979 | 12310 | 16616 | 17339 | 17745 | 18877 | 20759 | 22097 | 22910 | 24785 | 25433 | 26227 | 27012 | 27038 | 27513 | 27546 | 28310 | 29524 |  |
| ref | T | G | C | C | C | C | C | C | A | T | C | G | C | A | C | T | C | T |  |
| BA.5.2.19 | T | A | A | C | T | T | T | T | G | C | T | G | T | A | T | C | C | T |  |
| BA.5.2 | T | A | A | C | T | T | T | C | A | T | C | T | C | G | C | T | T | A | #1382884 |
| BF.10 | G | G | C | T | C | C | C | C | A | T | C | T | C | G | C | T | T | T |  |
| mut |  |  |  |  |  |  |  |  |  |  |  |  |  |  |  |  |  | T>A |  |

|  |  |  |  |  |  |  |  |  |  |  |  |  |  |  |  |  |  |  |  |  |  |  |  |  |  |  |  |  |  |  |  |  |  |  |  |  |  |
| --- | --- | --- | --- | --- | --- | --- | --- | --- | --- | --- | --- | --- | --- | --- | --- | --- | --- | --- | --- | --- | --- | --- | --- | --- | --- | --- | --- | --- | --- | --- | --- | --- | --- | --- | --- | --- | --- |
| pos | 2476 | 6529 | 9866 | 10977 | 11674 | 12160 | 12180 | 15009 | 15684 | 16323 | 16394 | 21721 | 21765 | 21766 | 21767 | 21768 | 21769 | 21770 | 22017 | 22917 | 23018 | 23040 | 23673 | 25416 | 26529 | 26858 | 27218 | 27259 | 27382 | 27383 | 27384 | 27438 | 27889 | 28297 | 29666 | 29754 |  |
| ref | A | A | C | C | C | G | A | T | T | C | C | C | T | A | C | A | T | G | G | T | T | A | C | C | G | C | A | A | G | A | T | T | C | T | C | C |  |
| BG.5 | A | A | T | C | T | G | G | C | C | C | C | T | T | A | C | A | T | G | G | A | T | G | T | T | G | T | A | C | C | T | C | T | C | C | C | C |  |
| BA.5.1.30 | A | G | T | C | T | G | G | C | C | C | C | C | - | - | - | - | - | - | T | G | G | A | C | C | A | C | G | A | G | A | T | C | T | T | T | T | #1383149 |
| BA.5.1.30 | G | A | C | T | C | A | A | T | T | T | T | C | - | - | - | - | - | - | G | G | G | A | C | C | A | C | G | A | G | A | T | C | T | T | T | T |  |
| mut |  | A>G |  |  |  |  |  |  |  |  |  |  |  |  |  |  |  |  | G>T |  |  |  |  |  |  |  |  |  |  |  |  |  |  |  |  |  |  |

|  |  |  |  |  |  |  |  |  |  |  |  |  |  |  |  |  |  |  |  |  |  |  |  |  |  |  |  |  |  |  |  |  |  |  |  |  |  |  |  |  |  |
| --- | --- | --- | --- | --- | --- | --- | --- | --- | --- | --- | --- | --- | --- | --- | --- | --- | --- | --- | --- | --- | --- | --- | --- | --- | --- | --- | --- | --- | --- | --- | --- | --- | --- | --- | --- | --- | --- | --- | --- | --- | --- |
| pos | 44 | 538 | 1471 | 2445 | 4057 | 4197 | 5406 | 6770 | 6786 | 8208 | 8991 | 10189 | 10323 | 12030 | 13904 | 16342 | 16474 | 17678 | 18252 | 22020 | 22054 | 22295 | 22332 | 22893 | 22910 | 22916 | 22942 | 23012 | 23013 | 23040 | 23779 | 23948 | 25810 | 26822 | 27874 | 27875 | 27881 | 27882 | 27883 | 28550 |  |
| ref | C | A | C | C | T | A | T | A | C | C | C | C | A | A | A | T | A | C | C | T | T | C | G | A | A | C | T | G | A | A | A | G | C | C | C | T | C | G | C | C |  |
| CM.4 | C | G | T | T | C | A | C | G | T | T | T | T | A | G | G | T | A | T | T | C | G | A | A | G | G | A | G | A | G | A | A | G | T | T | C | T | C | G | C | A |  |
| CM.4 | C | G | T | T | C | A | C | G | T | T | T | T | G | G | G | T | G | T | T | C | G | A | A | G | G | A | G | A | G | A | G | T | T | T | T | C | T | C | T | A | #1383981 |
| BA.2.10 | T | A | C | C | T | C | T | A | C | C | C | C | A | A | A | C | A | C | C | T | T | C | G | A | A | C | T | G | C | G | A | T | C | C | T | C | T | C | T | C |  |
| mut |  |  |  |  |  |  |  |  |  |  |  |  | A>G |  |  |  | A>G |  |  |  |  |  |  |  |  |  |  |  |  |  | A>G | G>T |  |  |  |  |  |  |  | C>A |  |

|  |  |  |  |  |  |  |  |  |  |  |  |  |  |  |  |  |  |  |
| --- | --- | --- | --- | --- | --- | --- | --- | --- | --- | --- | --- | --- | --- | --- | --- | --- | --- | --- |
| pos | 1627 | 1748 | 6979 | 8882 | 9967 | 17285 | 17373 | 19524 | 24620 | 27038 | 27438 | 27532 | 27998 | 28330 | 28402 | 29374 | 29666 |  |
| ref | C | G | T | A | C | C | C | C | G | A | T | C | C | A | A | G | C |  |
| BF.5 | T | G | G | G | T | C | C | T | T | G | T | T | C | G | G | G | C |  |
| BF.5 | T | G | G | G | T | C | C | T | T | G | T | T | T | A | A | A | T | #1385832 |
| BA.5.1.22 | C | A | T | A | C | T | T | C | G | A | C | C | T | A | A | A | T |  |
| mut |  |  |  |  |  |  |  |  |  |  |  |  |  |  |  |  |  |  |

|  |  |  |  |  |  |  |  |  |  |  |  |  |  |  |  |  |  |  |  |  |  |  |  |  |  |  |  |  |  |  |  |  |  |  |  |
| --- | --- | --- | --- | --- | --- | --- | --- | --- | --- | --- | --- | --- | --- | --- | --- | --- | --- | --- | --- | --- | --- | --- | --- | --- | --- | --- | --- | --- | --- | --- | --- | --- | --- | --- | --- |
| pos | 92 | 686 | 687 | 688 | 689 | 690 | 691 | 692 | 693 | 694 | 3429 | 5549 | 8140 | 11781 | 12628 | 14599 | 16596 | 21034 | 22599 | 23535 | 25463 | 26529 | 26858 | 27259 | 27382 | 27383 | 27384 | 27438 | 27788 | 27889 | 28462 | 28724 | 29666 | 29754 |  |
| ref | C | A | A | G | T | C | A | T | T | T | C | C | C | A | T | C | C | C | G | A | C | G | C | A | G | A | T | T | G | C | C | C | C | C |  |
| BA.4.6 | T | - | - | - | - | - | - | - | - | - | C | C | T | A | C | T | C | C | C | G | C | G | T | C | C | T | C | T | T | C | C | T | C | C |  |
| BA.5.1 | T | - | - | - | - | - | - | - | - | - | C | C | T | A | C | T | C | C | C | G | T | A | C | A | G | A | T | C | G | T | T | C | T | T | #1388466 |
| BA.5.1.10 | C | A | A | G | T | C | A | T | T | T | T | A | C | G | T | C | T | T | G | A | T | A | C | A | G | A | T | C | G | T | T | C | T | T |  |
| mut |  |  |  |  |  |  |  |  |  |  |  |  |  |  |  |  |  |  |  |  |  |  |  |  |  |  |  |  |  |  |  |  |  |  |  |

|  |  |  |  |  |  |  |  |  |  |  |  |  |  |  |  |  |  |  |  |  |  |  |  |  |  |  |  |  |  |  |  |  |  |
| --- | --- | --- | --- | --- | --- | --- | --- | --- | --- | --- | --- | --- | --- | --- | --- | --- | --- | --- | --- | --- | --- | --- | --- | --- | --- | --- | --- | --- | --- | --- | --- | --- | --- |
| pos | 686 | 687 | 688 | 689 | 690 | 691 | 692 | 693 | 694 | 1931 | 2902 | 8140 | 10188 | 12769 | 13342 | 14599 | 15960 | 19203 | 22599 | 23535 | 26529 | 26858 | 27259 | 27382 | 27383 | 27384 | 27788 | 27800 | 27889 | 28681 | 28724 | 29754 |  |
| ref | A | A | G | T | C | A | T | T | T | C | C | C | C | G | C | C | C | T | G | A | G | C | A | G | A | T | G | C | C | G | C | C |  |
| BA.4.6 | - | - | - | - | - | - | - | - | - | C | C | T | C | A | C | T | C | T | C | G | G | T | C | C | T | C | T | C | C | G | T | C |  |
| BA.5 | - | - | - | - | - | - | - | - | - | C | T | T | C | A | C | T | T | C | C | A | A | C | A | G | A | T | G | T | T | T | C | T | #1389175 |
| BE.4 | A | A | G | T | C | A | T | T | T | A | C | C | A | G | T | C | C | T | G | A | A | C | A | G | A | T | G | T | T | T | C | T |  |
| mut |  |  |  |  |  |  |  |  |  |  | C>T |  |  |  |  |  | C>T | T>C |  |  |  |  |  |  |  |  |  |  |  |  |  |  |  |

|  |  |  |  |  |  |  |  |  |  |  |  |  |  |  |  |  |  |  |  |  |  |  |  |  |  |  |  |  |  |  |  |  |  |  |
| --- | --- | --- | --- | --- | --- | --- | --- | --- | --- | --- | --- | --- | --- | --- | --- | --- | --- | --- | --- | --- | --- | --- | --- | --- | --- | --- | --- | --- | --- | --- | --- | --- | --- | --- |
| pos | 335 | 686 | 687 | 688 | 689 | 690 | 691 | 692 | 693 | 694 | 1170 | 5833 | 15852 | 15957 | 22326 | 23380 | 23535 | 26529 | 26858 | 27259 | 27382 | 27383 | 27384 | 27438 | 27484 | 27788 | 27889 | 28063 | 28254 | 28390 | 28724 | 29666 | 29754 |  |
| ref | C | A | A | G | T | C | A | T | T | T | C | C | C | G | C | C | A | G | C | A | G | A | T | T | T | G | C | T | A | A | C | C | C |  |
| BA.5.1.23 | C | A | A | G | T | C | A | T | T | T | T | C | C | A | T | T | A | A | C | A | G | A | T | C | T | G | T | T | - | A | C | T | T |  |
| BA.5 | C | A | A | G | T | C | A | T | T | T | T | C | C | A | T | T | A | A | C | A | G | A | T | T | C | T | C | C | A | G | T | C | C | #1389607 |
| BA.4 | T | - | - | - | - | - | - | - | - | - | C | T | T | G | C | C | G | G | T | C | C | T | C | T | C | T | C | T | A | A | T | C | C |  |
| mut |  |  |  |  |  |  |  |  |  |  |  |  |  |  |  |  |  |  |  |  |  |  |  |  |  |  |  | T>C |  | A>G |  |  |  |  |

|  |  |  |  |  |  |  |  |  |  |  |  |  |  |  |  |  |  |  |  |
| --- | --- | --- | --- | --- | --- | --- | --- | --- | --- | --- | --- | --- | --- | --- | --- | --- | --- | --- | --- |
| pos | 344 | 1627 | 1684 | 4906 | 5284 | 6735 | 6819 | 17402 | 18957 | 18984 | 19554 | 21077 | 22588 | 25710 | 27038 | 28330 | 28370 | 28371 |  |
| ref | C | C | C | T | C | T | G | C | T | G | C | C | T | C | A | A | A | G |  |
| BA.5.6 | T | C | C | C | C | C | A | T | C | T | C | T | C | C | A | A | - | G |  |
| BA.5.6 | T | C | C | C | C | T | A | T | C | T | T | T | C | T | G | G | T | T | #1390340 |
| BA.5.2.1 | C | T | T | T | T | T | G | C | T | G | C | C | T | T | G | G | T | T |  |
| mut |  |  |  |  |  | C>T |  |  |  |  | C>T |  |  |  |  |  |  |  |  |

|  |  |  |  |  |  |  |  |  |  |  |  |  |  |  |
| --- | --- | --- | --- | --- | --- | --- | --- | --- | --- | --- | --- | --- | --- | --- |
| pos | 1627 | 7235 | 9319 | 12310 | 16616 | 17615 | 27012 | 27438 | 27513 | 28330 | 29284 | 29614 | 29666 |  |
| ref | C | G | C | G | C | A | C | T | C | A | C | C | C |  |
| BA.5.1 | C | A | C | G | C | G | C | C | C | A | C | T | T |  |
| BA.5.2 | C | A | T | G | C | G | T | T | T | G | T | C | C | #1390557 |
| BA.5.2 | T | G | C | A | A | A | T | T | T | G | T | C | C |  |
| mut |  |  | C>T |  |  |  |  |  |  |  |  |  |  |  |

|  |  |  |  |  |  |  |  |  |  |  |  |  |  |  |  |  |  |  |  |  |  |  |  |  |
| --- | --- | --- | --- | --- | --- | --- | --- | --- | --- | --- | --- | --- | --- | --- | --- | --- | --- | --- | --- | --- | --- | --- | --- | --- |
| pos | 1427 | 1627 | 1931 | 3601 | 3743 | 5812 | 7528 | 10834 | 11750 | 16935 | 21462 | 21991 | 21992 | 21993 | 22599 | 22851 | 23603 | 27038 | 27889 | 28330 | 28370 | 28371 | 28681 |  |
| ref | C | C | C | A | C | C | C | C | C | G | T | T | T | A | G | C | C | A | C | A | A | G | G |  |
| BF.7.13.1 | T | T | C | A | T | C | T | C | C | G | C | T | T | A | C | T | T | G | T | G | T | T | G |  |
| BF.7.13 | T | T | C | G | T | T | T | C | T | A | T | T | T | A | G | C | C | A | C | A | - | G | T | #1391762 |
| BE.1 | C | C | A | A | C | C | C | T | C | A | T | - | - | - | G | C | C | A | C | A | - | G | T |  |
| mut |  |  |  | A>G |  | C>T |  |  | C>T |  |  | ->T | ->T | ->A |  |  |  |  |  |  |  |  |  |  |

|  |  |  |  |  |  |  |  |  |  |  |  |  |  |  |  |  |  |  |
| --- | --- | --- | --- | --- | --- | --- | --- | --- | --- | --- | --- | --- | --- | --- | --- | --- | --- | --- |
| pos | 396 | 1627 | 1932 | 2065 | 3483 | 6941 | 6979 | 7417 | 11751 | 14472 | 14747 | 16705 | 27038 | 27438 | 28330 | 29416 | 29666 |  |
| ref | A | C | A | C | G | C | T | C | T | T | A | G | A | T | A | C | C |  |
| BF.28 | T | T | A | C | A | T | G | T | T | C | A | T | G | T | G | C | C |  |
| BF.28 | T | T | G | C | A | T | G | T | T | C | A | T | A | C | A | T | T | #1391791 |
| BA.5.1 | A | C | A | T | G | C | T | C | A | T | G | G | A | C | A | C | T |  |
| mut |  |  | A>G |  |  |  |  |  |  |  |  |  |  |  |  | C>T |  |  |

|  |  |  |  |  |  |  |  |  |  |  |  |  |
| --- | --- | --- | --- | --- | --- | --- | --- | --- | --- | --- | --- | --- |
| pos | 1931 | 2857 | 2954 | 11750 | 16935 | 19662 | 22427 | 27438 | 28681 | 29666 | 29808 |  |
| ref | C | C | T | C | G | C | G | T | G | C | G |  |
| BE.1.1 | A | C | C | T | A | T | G | T | T | C | T |  |
| BE.1.1 | A | T | C | T | A | T | A | C | G | T | G | #1394979 |
| BA.5.1.3 | C | C | T | C | G | C | A | C | G | T | G |  |
| mut |  | C>T |  |  |  |  |  |  |  |  |  |  |

|  |  |  |  |  |  |  |  |  |  |  |  |  |  |  |  |  |  |  |  |  |
| --- | --- | --- | --- | --- | --- | --- | --- | --- | --- | --- | --- | --- | --- | --- | --- | --- | --- | --- | --- | --- |
| pos | 589 | 1627 | 2445 | 2841 | 6145 | 10369 | 12310 | 15002 | 16428 | 17750 | 18698 | 24381 | 25350 | 27059 | 27438 | 28330 | 28510 | 29140 | 29666 |  |
| ref | C | C | C | C | C | C | G | A | C | C | C | C | C | C | T | A | T | G | C |  |
| BA.5.2.9 | T | T | T | T | T | T | A | G | C | C | T | A | C | T | T | G | T | G | C |  |
| BA.5.1.5 | T | T | T | T | T | T | A | G | T | T | C | C | A | C | C | A | C | T | T | #1395297 |
| BA.5.1.5 | C | C | C | C | C | C | G | A | T | C | C | C | A | C | C | A | C | T | T |  |
| mut |  |  |  |  |  |  |  |  |  | C>T |  |  |  |  |  |  |  |  |  |  |

|  |  |  |  |  |  |  |  |  |  |  |  |  |  |  |  |  |  |  |  |  |  |  |  |  |  |  |  |  |  |  |  |  |  |  |  |  |  |  |  |
| --- | --- | --- | --- | --- | --- | --- | --- | --- | --- | --- | --- | --- | --- | --- | --- | --- | --- | --- | --- | --- | --- | --- | --- | --- | --- | --- | --- | --- | --- | --- | --- | --- | --- | --- | --- | --- | --- | --- | --- |
| pos | 44 | 261 | 405 | 3796 | 3927 | 4586 | 5183 | 12444 | 15738 | 15939 | 16342 | 17859 | 18583 | 21810 | 21991 | 21992 | 21993 | 22000 | 22001 | 22016 | 22033 | 22109 | 22190 | 22200 | 22331 | 22577 | 22599 | 22664 | 22895 | 22896 | 22898 | 22942 | 23019 | 23031 | 23040 | 25416 | 25700 | 26275 |  |
| ref | C | G | A | C | C | C | C | A | C | T | T | T | G | T | T | T | A | C | A | T | C | C | A | T | G | G | G | C | G | T | G | T | T | T | A | C | C | A |  |
| BA.2.10 | C | G | A | C | C | C | C | A | C | T | C | T | G | T | T | T | A | C | A | T | C | C | A | G | G | G | G | C | G | T | G | T | T | T | G | C | T | A |  |
| XBB | C | G | G | C | C | C | C | A | T | C | C | C | G | C | - | - | - | A | A | T | C | G | A | A | G | C | C | A | C | C | A | G | C | C | A | T | C | G | #1396207 |
| BM.1.1.1 | T | A | A | T | T | T | T | G | C | T | T | T | A | T | T | T | A | C | G | C | A | C | G | G | A | C | C | C | G | T | A | G | C | C | A | T | C | G |  |
| mut |  |  | A>G |  |  |  |  |  | C>T | T>C |  | T>C |  | T>C | T>- | T>- | A>- | C>A |  |  |  | C>G |  | G>A |  |  |  | C>A | G>C | T>C |  |  |  |  |  |  |  |  |  |

|  |  |  |  |  |  |  |  |  |  |  |  |  |  |  |  |  |  |  |  |  |  |  |  |  |  |  |  |  |  |  |  |  |  |  |  |  |
| --- | --- | --- | --- | --- | --- | --- | --- | --- | --- | --- | --- | --- | --- | --- | --- | --- | --- | --- | --- | --- | --- | --- | --- | --- | --- | --- | --- | --- | --- | --- | --- | --- | --- | --- | --- | --- |
| pos | 376 | 686 | 687 | 688 | 689 | 690 | 691 | 692 | 693 | 694 | 941 | 1931 | 8140 | 10287 | 12970 | 14599 | 16935 | 18826 | 19152 | 21137 | 23535 | 25614 | 26529 | 26858 | 27259 | 27382 | 27383 | 27384 | 27788 | 27889 | 28170 | 28312 | 28681 | 28724 | 29754 |  |
| ref | G | A | A | G | T | C | A | T | T | T | A | C | C | T | C | C | G | G | C | A | A | C | G | C | A | G | A | T | G | C | C | C | G | C | C |  |
| BA.4.6 | G | - | - | - | - | - | - | - | - | - | G | C | T | T | C | T | G | G | T | A | G | C | G | T | C | C | T | C | T | C | C | C | G | T | C |  |
| BE.1.4 | G | - | - | - | - | - | - | - | - | - | G | C | T | T | T | C | A | A | C | G | A | T | A | C | A | G | A | T | G | T | A | T | T | C | T | #1398473 |
| BE.1.4.2 | A | A | A | G | T | C | A | T | T | T | A | A | C | C | C | C | A | A | C | G | A | T | A | C | A | G | A | T | G | T | A | T | T | C | T |  |
| mut |  |  |  |  |  |  |  |  |  |  |  |  |  |  | C>T |  |  |  |  |  |  |  |  |  |  |  |  |  |  |  |  |  |  |  |  |  |

|  |  |  |  |  |  |  |  |  |  |  |  |  |  |  |  |  |  |  |  |  |  |  |  |  |  |  |  |  |  |  |  |  |  |  |  |  |  |  |  |  |  |  |  |  |  |  |  |  |  |  |
| --- | --- | --- | --- | --- | --- | --- | --- | --- | --- | --- | --- | --- | --- | --- | --- | --- | --- | --- | --- | --- | --- | --- | --- | --- | --- | --- | --- | --- | --- | --- | --- | --- | --- | --- | --- | --- | --- | --- | --- | --- | --- | --- | --- | --- | --- | --- | --- | --- | --- | --- |
| pos | 1424 | 1627 | 2306 | 2509 | 3796 | 3927 | 4586 | 5183 | 8090 | 9335 | 9424 | 9866 | 12160 | 12310 | 12444 | 15451 | 16616 | 21765 | 21766 | 21767 | 21768 | 21769 | 21770 | 22001 | 22016 | 22033 | 22104 | 22190 | 22331 | 22577 | 22629 | 22898 | 22917 | 22942 | 23018 | 25416 | 26275 | 26529 | 26858 | 27012 | 27259 | 27382 | 27383 | 27384 | 27513 | 27889 | 28330 | 28791 | 29754 |  |
| ref | T | C | C | C | C | C | C | C | C | G | A | C | G | G | A | G | C | T | A | C | A | T | G | A | T | C | G | A | G | G | A | G | T | T | T | C | A | G | C | C | A | G | A | T | C | C | A | C | C |  |
| BA.2.75.5 | T | C | C | C | T | T | T | T | C | G | G | T | G | G | G | A | C | T | A | C | A | T | G | G | C | A | G | G | A | C | C | A | T | G | T | T | G | G | T | C | C | C | T | C | C | C | A | C | C |  |
| BA.2.75 | T | C | C | C | T | T | T | T | T | T | G | T | G | G | G | A | C | T | A | C | A | T | G | G | C | A | G | G | A | C | C | A | T | G | T | T | G | G | C | T | A | G | A | T | T | T | G | T | T | #1400720 |
| BA.5.2 | C | T | T | A | C | C | C | C | C | G | A | C | A | A | A | G | A | - | - | - | - | - | - | A | T | C | T | A | G | G | A | G | G | T | G | C | A | A | C | T | A | G | A | T | T | T | G | T | T |  |
| mut |  |  |  |  |  |  |  |  | C>T | G>T |  |  |  |  |  |  |  |  |  |  |  |  |  |  |  |  |  |  |  |  |  |  |  |  |  |  |  |  |  |  |  |  |  |  |  |  |  |  |  |  |

|  |  |  |  |  |  |  |  |  |  |  |  |  |  |  |  |  |  |  |  |  |  |  |  |  |  |  |  |  |  |  |  |  |  |  |  |  |  |  |  |  |  |  |  |  |  |  |  |  |  |  |  |  |  |  |
| --- | --- | --- | --- | --- | --- | --- | --- | --- | --- | --- | --- | --- | --- | --- | --- | --- | --- | --- | --- | --- | --- | --- | --- | --- | --- | --- | --- | --- | --- | --- | --- | --- | --- | --- | --- | --- | --- | --- | --- | --- | --- | --- | --- | --- | --- | --- | --- | --- | --- | --- | --- | --- | --- | --- |
| pos | 1320 | 1931 | 2954 | 3040 | 3796 | 3927 | 4586 | 5183 | 9866 | 11750 | 12160 | 12444 | 14257 | 15451 | 16935 | 17969 | 18814 | 20235 | 21765 | 21766 | 21767 | 21768 | 21769 | 21770 | 22001 | 22016 | 22033 | 22190 | 22331 | 22577 | 22599 | 22629 | 22893 | 22898 | 22917 | 22942 | 23018 | 23031 | 24977 | 25416 | 26078 | 26275 | 26529 | 26858 | 27259 | 27382 | 27383 | 27384 | 27600 | 27889 | 28312 | 28681 | 29754 |  |
| ref | A | C | T | C | C | C | C | C | C | C | G | A | T | G | G | A | C | C | T | A | C | A | T | G | A | T | C | A | G | G | G | A | A | G | T | T | T | T | G | C | C | A | G | C | A | G | A | T | C | C | C | G | C |  |
| BQ.1.10 | C | A | C | T | C | C | C | C | C | T | A | A | C | G | A | A | C | C | - | - | - | - | - | - | A | T | C | A | G | G | G | A | C | G | G | A | G | T | G | C | C | A | A | C | A | G | A | T | T | T | T | T | T |  |
| BA.2 | C | A | C | T | C | C | C | C | C | T | A | A | C | G | A | C | T | C | T | A | C | A | T | G | G | C | A | G | A | C | C | C | A | A | T | G | T | C | C | T | T | G | G | T | C | C | T | C | C | C | C | G | C | #1401612 |
| BN.1.3 | A | C | T | C | T | T | T | T | T | C | G | G | T | A | G | A | C | T | T | A | C | A | T | G | G | C | A | G | A | C | C | C | A | A | T | G | T | C | G | T | T | G | G | T | C | C | T | C | C | C | C | G | C |  |
| mut |  |  |  |  |  |  |  |  |  |  |  |  |  |  |  | A>C | C>T |  |  |  |  |  |  |  |  |  |  |  |  |  |  |  |  |  |  |  |  |  | G>C |  |  |  |  |  |  |  |  |  |  |  |  |  |  |  |

|  |  |  |  |  |  |  |  |  |  |  |  |  |
| --- | --- | --- | --- | --- | --- | --- | --- | --- | --- | --- | --- | --- |
| pos | 1288 | 3859 | 5284 | 6317 | 6979 | 12315 | 14883 | 19602 | 28370 | 28371 | 29769 |  |
| ref | C | A | C | C | T | G | T | C | A | G | C |  |
| BA.5.2.1 | C | A | T | C | T | G | T | C | T | T | C |  |
| BA.5.2.1 | C | A | T | C | T | A | C | T | - | G | T | #1403515 |
| BA.5.2.1 | T | G | C | T | G | A | C | T | - | G | T |  |
| mut |  |  |  |  |  |  |  |  |  |  |  |  |

|  |  |  |  |  |  |  |  |  |  |  |  |  |  |  |  |  |  |  |  |  |  |  |  |  |  |  |  |  |  |  |  |  |  |  |  |  |  |  |  |  |  |  |  |  |  |  |  |  |  |  |  |  |  |  |  |
| --- | --- | --- | --- | --- | --- | --- | --- | --- | --- | --- | --- | --- | --- | --- | --- | --- | --- | --- | --- | --- | --- | --- | --- | --- | --- | --- | --- | --- | --- | --- | --- | --- | --- | --- | --- | --- | --- | --- | --- | --- | --- | --- | --- | --- | --- | --- | --- | --- | --- | --- | --- | --- | --- | --- | --- |
| pos | 593 | 1627 | 3796 | 3927 | 4551 | 4586 | 5183 | 7390 | 9866 | 11146 | 12160 | 12310 | 12444 | 15451 | 16616 | 20235 | 21765 | 21766 | 21767 | 21768 | 21769 | 21770 | 22001 | 22016 | 22033 | 22190 | 22331 | 22577 | 22599 | 22629 | 22898 | 22917 | 22942 | 23018 | 23031 | 23707 | 24378 | 25416 | 25626 | 26078 | 26275 | 26529 | 26858 | 27012 | 27259 | 27382 | 27383 | 27384 | 27513 | 27889 | 28330 | 28975 | 29134 | 29754 |  |
| ref | C | C | C | C | C | C | C | C | C | G | G | G | A | G | C | C | T | A | C | A | T | G | A | T | C | A | G | G | G | A | G | T | T | T | T | C | C | C | C | C | A | G | C | C | A | G | A | T | C | C | A | G | G | C |  |
| BA.5.2.20 | C | T | C | C | T | C | C | C | C | G | A | A | A | G | A | C | - | - | - | - | - | - | A | T | C | A | G | G | G | A | G | G | T | G | T | T | C | C | T | C | A | A | C | T | A | G | A | T | T | T | G | G | G | T |  |
| BN.1.3.7 | C | T | C | C | T | T | T | A | T | T | G | G | G | A | C | T | T | A | C | A | T | G | G | C | A | G | A | C | C | C | A | T | G | T | C | C | T | T | C | T | G | G | T | C | C | C | T | C | C | C | A | C | A | C | #1404808 |
| BN.1.3 | T | C | T | T | C | T | T | A | T | T | G | G | G | A | C | T | T | A | C | A | T | G | G | C | A | G | A | C | C | C | A | T | G | T | C | C | C | T | C | T | G | G | T | C | C | C | T | C | C | C | A | G | G | C |  |
| mut |  |  |  |  |  |  |  |  |  |  |  |  |  |  |  |  |  |  |  |  |  |  |  |  |  |  |  |  |  |  |  |  |  |  |  |  | C>T |  |  |  |  |  |  |  |  |  |  |  |  |  |  | G>C | G>A |  |  |

|  |  |  |  |  |  |  |  |  |  |  |  |  |  |  |  |  |  |  |  |  |  |  |  |  |  |  |  |  |  |  |  |  |  |  |  |  |  |  |  |  |  |  |  |  |  |  |  |  |  |
| --- | --- | --- | --- | --- | --- | --- | --- | --- | --- | --- | --- | --- | --- | --- | --- | --- | --- | --- | --- | --- | --- | --- | --- | --- | --- | --- | --- | --- | --- | --- | --- | --- | --- | --- | --- | --- | --- | --- | --- | --- | --- | --- | --- | --- | --- | --- | --- | --- | --- |
| pos | 1627 | 1878 | 3796 | 3927 | 4586 | 5183 | 5284 | 9866 | 11837 | 12160 | 12444 | 16448 | 20235 | 21765 | 21766 | 21767 | 21768 | 21769 | 21770 | 22001 | 22016 | 22033 | 22190 | 22331 | 22577 | 22599 | 22629 | 22898 | 22917 | 22942 | 23018 | 23031 | 23042 | 25416 | 26275 | 26529 | 26858 | 27038 | 27259 | 27382 | 27383 | 27384 | 27889 | 28330 | 28370 | 28371 | 28585 | 29754 |  |
| ref | C | C | C | C | C | C | C | C | G | G | A | A | C | T | A | C | A | T | G | A | T | C | A | G | G | G | A | G | T | T | T | T | T | C | A | G | C | A | A | G | A | T | C | A | A | G | C | C |  |
| BA.5.2.1 | T | T | C | C | C | C | T | C | G | A | A | A | C | - | - | - | - | - | - | A | T | C | A | G | G | G | A | G | G | T | G | T | T | C | A | A | C | G | A | G | A | T | T | G | T | T | C | T |  |
| BA.2 | T | T | C | C | C | C | T | C | A | A | A | T | T | T | A | C | A | T | G | G | C | A | G | A | C | C | C | A | T | G | T | C | C | T | G | G | T | A | C | C | T | C | C | A | - | G | T | C | #1404920 |
| BN.1.1 | C | C | T | T | T | T | C | T | G | G | G | A | T | T | A | C | A | T | G | G | C | A | G | A | C | C | C | A | T | G | T | C | C | T | G | G | T | A | C | C | T | C | C | A | - | G | T | C |  |
| mut |  |  |  |  |  |  |  |  | G>A |  |  | A>T |  |  |  |  |  |  |  |  |  |  |  |  |  |  |  |  |  |  |  |  |  |  |  |  |  |  |  |  |  |  |  |  |  |  |  |  |  |

|  |  |  |  |  |  |  |  |  |  |  |  |  |  |  |  |  |  |  |  |
| --- | --- | --- | --- | --- | --- | --- | --- | --- | --- | --- | --- | --- | --- | --- | --- | --- | --- | --- | --- |
| pos | 1931 | 2954 | 7083 | 11750 | 14257 | 16428 | 16935 | 17039 | 17528 | 22599 | 22893 | 22942 | 25350 | 27438 | 28312 | 28510 | 28681 | 29666 |  |
| ref | C | T | C | C | T | C | G | A | C | G | A | T | C | T | C | T | G | C |  |
| BQ.1.1 | A | C | C | T | C | C | A | G | C | C | C | A | C | T | T | T | T | C |  |
| BQ.1.1.17 | A | C | T | T | C | C | A | G | A | C | C | A | C | T | C | C | G | T | #1406252 |
| BA.5.1.5 | C | T | C | C | T | T | G | A | C | G | A | T | A | C | C | C | G | T |  |
| mut |  |  | C>T |  |  |  |  |  | C>A |  |  |  |  |  |  |  |  |  |  |

|  |  |  |  |  |  |  |  |  |  |  |  |  |  |  |  |  |  |  |  |  |  |  |  |  |  |  |  |  |  |  |  |  |  |  |  |  |  |  |  |  |  |  |  |
| --- | --- | --- | --- | --- | --- | --- | --- | --- | --- | --- | --- | --- | --- | --- | --- | --- | --- | --- | --- | --- | --- | --- | --- | --- | --- | --- | --- | --- | --- | --- | --- | --- | --- | --- | --- | --- | --- | --- | --- | --- | --- | --- | --- |
| pos | 203 | 222 | 686 | 687 | 688 | 689 | 690 | 691 | 692 | 693 | 694 | 1627 | 7528 | 9365 | 13096 | 13694 | 17746 | 18348 | 18366 | 21180 | 21483 | 21570 | 22599 | 25177 | 26529 | 26858 | 27038 | 27259 | 27294 | 27382 | 27383 | 27384 | 27788 | 27834 | 27889 | 27942 | 28330 | 28370 | 28371 | 28724 | 28770 | 29754 |  |
| ref | C | C | A | A | G | T | C | A | T | T | T | C | C | C | T | C | C | C | A | T | T | T | G | G | G | C | A | A | C | G | A | T | G | A | C | C | A | A | G | C | C | C |  |
| BF.7.4.1 | - | - | A | A | G | T | C | A | T | T | T | T | T | T | C | T | T | C | A | C | C | T | C | G | A | C | G | A | T | G | A | T | G | A | T | T | G | T | T | C | C | T |  |
| BF.7.4.1 | C | C | A | A | G | T | C | A | T | T | T | T | T | T | C | T | T | C | A | C | C | T | C | T | A | C | G | A | T | G | A | T | T | G | C | C | A | - | G | T | T | C | #1406297 |
| BA.4.1 | C | C | - | - | - | - | - | - | - | - | - | C | C | C | T | C | C | T | G | T | T | G | G | G | G | T | A | C | C | C | T | C | T | A | C | C | A | - | G | T | T | C |  |
| mut | ->C | ->C |  |  |  |  |  |  |  |  |  |  |  |  |  |  |  |  |  |  |  |  |  | G>T |  |  |  |  |  |  |  |  |  | A>G |  |  |  |  |  |  |  |  |  |

|  |  |  |  |  |  |  |  |  |  |  |  |  |  |  |  |  |  |  |  |  |  |  |  |  |
| --- | --- | --- | --- | --- | --- | --- | --- | --- | --- | --- | --- | --- | --- | --- | --- | --- | --- | --- | --- | --- | --- | --- | --- | --- |
| pos | 625 | 1931 | 2954 | 3482 | 4906 | 6982 | 10116 | 11750 | 14257 | 14932 | 16935 | 17039 | 17402 | 17979 | 18957 | 20117 | 21077 | 22599 | 22893 | 22942 | 28312 | 28681 | 29260 |  |
| ref | G | C | T | G | T | C | C | C | T | C | G | A | C | T | T | C | C | G | A | T | C | G | G |  |
| BA.5.6 | G | C | T | A | C | T | T | C | T | T | G | A | T | T | C | T | T | G | A | T | C | G | G |  |
| BQ.1.1 | T | C | T | A | C | C | C | T | C | C | A | G | C | A | T | C | C | C | C | A | T | T | A | #1407172 |
| BQ.1.1 | G | A | C | G | T | C | C | T | C | C | A | G | C | A | T | C | C | C | C | A | T | T | A |  |
| mut | G>T |  |  |  |  |  |  |  |  |  |  |  |  |  |  |  |  |  |  |  |  |  |  |  |

|  |  |  |  |  |  |  |  |  |  |  |  |  |  |  |  |  |  |  |  |  |  |  |  |  |  |  |  |  |  |  |  |  |  |  |  |  |  |  |  |  |  |  |  |  |  |  |  |  |  |  |  |  |  |  |
| --- | --- | --- | --- | --- | --- | --- | --- | --- | --- | --- | --- | --- | --- | --- | --- | --- | --- | --- | --- | --- | --- | --- | --- | --- | --- | --- | --- | --- | --- | --- | --- | --- | --- | --- | --- | --- | --- | --- | --- | --- | --- | --- | --- | --- | --- | --- | --- | --- | --- | --- | --- | --- | --- | --- |
| pos | 261 | 1931 | 2954 | 3796 | 3927 | 4540 | 4586 | 5006 | 5183 | 8090 | 9349 | 9529 | 9866 | 11750 | 12160 | 12444 | 14257 | 15451 | 16935 | 18583 | 19185 | 21765 | 21766 | 21767 | 21768 | 21769 | 21770 | 22001 | 22016 | 22033 | 22190 | 22331 | 22577 | 22599 | 22893 | 22898 | 22917 | 22942 | 23018 | 23019 | 23031 | 25416 | 26275 | 26529 | 26858 | 27259 | 27382 | 27383 | 27384 | 27889 | 28312 | 28681 | 29754 |  |
| ref | G | C | T | C | C | C | C | A | C | C | T | A | C | C | G | A | T | G | G | G | C | T | A | C | A | T | G | A | T | C | A | G | G | G | A | G | T | T | T | T | T | C | A | G | C | A | G | A | T | C | C | G | C |  |
| BQ.1 | G | A | C | C | C | T | C | A | C | C | T | A | C | T | A | A | C | G | A | G | C | - | - | - | - | - | - | A | T | C | A | G | G | G | C | G | G | A | G | T | T | C | A | A | C | A | G | A | T | T | T | T | T |  |
| BA.2 | G | A | C | C | C | T | C | A | C | T | C | G | T | C | G | G | T | A | G | A | T | T | A | C | A | T | G | G | C | A | G | A | C | C | A | A | T | G | C | C | C | T | G | G | T | C | C | T | C | C | C | G | C | #1407336 |
| CJ.1.2 | A | C | T | T | T | C | T | G | T | T | T | A | T | C | G | G | T | A | G | A | C | T | A | C | A | T | G | G | C | A | G | A | C | C | A | A | T | G | C | C | C | T | G | G | T | C | C | T | C | C | C | G | C |  |
| mut |  |  |  |  |  |  |  |  |  |  | T>C | A>G |  |  |  |  |  |  |  |  | C>T |  |  |  |  |  |  |  |  |  |  |  |  |  |  |  |  |  |  |  |  |  |  |  |  |  |  |  |  |  |  |  |  |  |

|  |  |  |  |  |  |  |  |  |  |  |  |  |  |  |  |  |  |  |  |  |  |  |  |  |  |  |  |  |  |  |  |  |  |  |  |
| --- | --- | --- | --- | --- | --- | --- | --- | --- | --- | --- | --- | --- | --- | --- | --- | --- | --- | --- | --- | --- | --- | --- | --- | --- | --- | --- | --- | --- | --- | --- | --- | --- | --- | --- | --- |
| pos | 44 | 1426 | 1931 | 2197 | 8179 | 9866 | 10323 | 10834 | 12160 | 18975 | 21765 | 21766 | 21767 | 21768 | 21769 | 21770 | 22599 | 22792 | 22917 | 23018 | 23040 | 25624 | 26529 | 26858 | 27259 | 27382 | 27383 | 27384 | 27889 | 28435 | 28681 | 28693 | 29359 | 29754 |  |
| ref | C | C | C | C | G | C | A | C | G | T | T | A | C | A | T | G | G | C | T | T | A | C | G | C | A | G | A | T | C | C | G | T | A | C |  |
| BA.2.9 | C | T | C | T | G | T | A | C | G | T | T | A | C | A | T | G | G | T | T | T | G | T | G | T | C | C | T | C | C | T | G | T | A | C |  |
| BE.12 | C | T | C | T | C | T | G | T | A | C | - | - | - | - | - | - | C | C | G | G | A | C | A | C | A | G | A | T | T | C | T | C | G | T | #1409074 |
| BE.12 | T | C | A | C | G | C | A | T | A | C | - | - | - | - | - | - | G | C | G | G | A | C | A | C | A | G | A | T | T | C | T | C | G | T |  |
| mut |  |  |  |  | G>C |  | A>G |  |  |  |  |  |  |  |  |  | G>C |  |  |  |  |  |  |  |  |  |  |  |  |  |  |  |  |  |  |

|  |  |  |  |  |  |  |  |  |  |  |  |  |  |  |  |  |  |  |  |  |  |  |  |  |
| --- | --- | --- | --- | --- | --- | --- | --- | --- | --- | --- | --- | --- | --- | --- | --- | --- | --- | --- | --- | --- | --- | --- | --- | --- |
| pos | 1685 | 1931 | 2939 | 2954 | 3787 | 6373 | 10789 | 11750 | 12890 | 14257 | 16428 | 16935 | 17039 | 19374 | 22599 | 22893 | 22942 | 27438 | 28254 | 28312 | 28681 | 29074 | 29666 |  |
| ref | G | C | C | T | C | G | C | C | C | T | C | G | A | C | G | A | T | T | A | C | G | A | C |  |
| BA.5.1 | A | C | T | T | T | T | T | C | T | T | T | G | A | T | G | A | T | C | - | C | G | T | T |  |
| BQ.1.1 | A | C | T | T | T | G | C | T | C | C | C | A | G | C | C | C | A | T | A | T | T | A | C | #1410844 |
| BQ.1.1 | G | A | C | C | C | G | C | T | C | C | C | A | G | C | C | C | A | T | A | T | T | A | C |  |
| mut |  |  |  |  |  |  |  |  |  |  |  |  |  |  |  |  |  |  |  |  |  |  |  |  |

|  |  |  |  |  |  |  |  |  |  |  |  |  |  |  |  |  |  |  |  |  |  |
| --- | --- | --- | --- | --- | --- | --- | --- | --- | --- | --- | --- | --- | --- | --- | --- | --- | --- | --- | --- | --- | --- |
| pos | 1931 | 2710 | 2954 | 11322 | 11750 | 14257 | 16935 | 17039 | 21991 | 21992 | 21993 | 22893 | 22942 | 23277 | 27438 | 28312 | 28393 | 28681 | 29317 | 29666 |  |
| ref | C | C | T | C | C | T | G | A | T | T | A | A | T | C | T | C | T | G | T | C |  |
| BQ.1 | A | C | C | C | T | C | A | G | T | T | A | C | A | T | T | T | G | T | T | C |  |
| BQ.1 | A | T | C | T | T | C | A | G | - | - | - | C | A | T | C | C | T | G | C | T | #1411996 |
| BA.5.1 | C | C | T | C | C | T | G | A | T | T | A | A | T | C | C | C | T | G | C | T |  |
| mut |  | C>T |  | C>T |  |  |  |  | T>- | T>- | A>- |  |  |  |  |  |  |  |  |  |  |

|  |  |  |  |  |  |  |  |  |  |  |  |  |  |  |  |  |  |  |  |  |  |  |  |  |  |  |  |  |
| --- | --- | --- | --- | --- | --- | --- | --- | --- | --- | --- | --- | --- | --- | --- | --- | --- | --- | --- | --- | --- | --- | --- | --- | --- | --- | --- | --- | --- |
| pos | 1426 | 1627 | 1779 | 1931 | 2954 | 6133 | 7735 | 8251 | 10123 | 10340 | 11750 | 14257 | 16935 | 17039 | 21991 | 21992 | 21993 | 22599 | 22893 | 22942 | 26151 | 27038 | 27484 | 28271 | 28312 | 28330 | 28681 |  |
| ref | C | C | C | C | T | A | C | C | T | C | C | T | G | A | T | T | A | G | A | T | C | A | T | A | C | A | G |  |
| BQ.1.8 | T | C | T | A | C | A | T | T | T | C | T | C | A | G | - | - | - | G | C | A | C | A | C | T | T | A | T |  |
| BQ.1.8 | T | C | T | A | C | G | T | T | C | T | T | C | A | G | - | - | - | C | A | T | T | G | T | C | C | G | G | #1412422 |
| BF.11 | C | T | C | C | T | A | C | C | T | C | C | T | G | A | T | T | A | C | A | T | T | G | T | C | C | G | G |  |
| mut |  |  |  |  |  | A>G |  |  | T>C | C>T |  |  |  |  |  |  |  | G>C |  |  |  |  |  |  |  |  |  |  |

|  |  |  |  |  |  |  |  |  |  |  |  |  |  |  |  |  |  |  |  |  |  |  |  |
| --- | --- | --- | --- | --- | --- | --- | --- | --- | --- | --- | --- | --- | --- | --- | --- | --- | --- | --- | --- | --- | --- | --- | --- |
| pos | 1627 | 1931 | 2954 | 5421 | 6399 | 11750 | 12310 | 12892 | 14257 | 16616 | 16935 | 17039 | 22599 | 22893 | 22942 | 23260 | 27012 | 27513 | 28312 | 28330 | 28681 | 28727 |  |
| ref | C | C | T | A | A | C | G | G | T | C | G | A | G | A | T | C | C | C | C | A | G | G |  |
| BA.5.2 | T | C | T | G | A | C | A | G | T | A | G | A | G | A | T | C | T | T | C | G | G | T |  |
| BA.5.2.1 | T | C | T | G | G | C | A | G | T | A | G | A | C | C | A | A | C | C | T | A | T | G | #1413005 |
| BQ.1.1 | C | A | C | A | A | T | G | A | C | C | A | G | C | C | A | A | C | C | T | A | T | G |  |
| mut |  |  |  |  | A>G |  |  |  |  |  |  |  | G>C |  |  |  |  |  |  |  |  |  |  |

|  |  |  |  |  |  |  |  |  |  |  |  |  |  |  |  |  |  |  |  |  |  |  |  |  |  |  |  |  |  |  |  |  |  |  |  |  |  |  |  |  |  |  |  |  |
| --- | --- | --- | --- | --- | --- | --- | --- | --- | --- | --- | --- | --- | --- | --- | --- | --- | --- | --- | --- | --- | --- | --- | --- | --- | --- | --- | --- | --- | --- | --- | --- | --- | --- | --- | --- | --- | --- | --- | --- | --- | --- | --- | --- | --- |
| pos | 1931 | 2954 | 6525 | 6548 | 9300 | 9866 | 11305 | 11750 | 12160 | 14257 | 16935 | 21765 | 21766 | 21767 | 21768 | 21769 | 21770 | 21991 | 21992 | 21993 | 22600 | 22893 | 22917 | 22942 | 23018 | 23040 | 24844 | 25416 | 25603 | 26529 | 26858 | 27259 | 27382 | 27383 | 27384 | 27889 | 28312 | 28681 | 29729 | 29754 | 29757 | 29758 | 29759 |  |
| ref | C | T | C | C | G | C | A | C | G | T | G | T | A | C | A | T | G | T | T | A | A | A | T | T | T | A | C | C | C | G | C | A | G | A | T | C | C | G | T | C | G | T | G |  |
| BQ.1 | A | C | C | C | G | C | A | T | A | C | A | - | - | - | - | - | - | T | T | A | A | C | G | A | G | A | C | C | C | A | C | A | G | A | T | T | T | T | T | T | G | T | G |  |
| BQ.1.21 | A | C | T | C | G | C | G | T | A | C | A | - | - | - | - | - | - | - | - | - | C | C | G | A | G | A | T | C | T | A | C | A | G | A | T | T | T | T | C | C | A | C | C | #1418709 |
| BA.2 | C | T | C | T | A | T | A | C | G | T | G | T | A | C | A | T | G | T | T | A | T | A | T | T | T | G | C | T | C | G | T | C | C | T | C | C | C | G | C | C | A | C | C |  |
| mut |  |  | C>T |  |  |  | A>G |  |  |  |  |  |  |  |  |  |  | T>- | T>- | A>- | A>C |  |  |  |  |  | C>T |  | C>T |  |  |  |  |  |  |  |  |  |  |  |  |  |  |  |

|  |  |  |  |  |  |  |  |  |  |  |  |  |  |  |  |  |  |  |  |  |  |  |  |  |  |  |  |  |  |  |  |  |  |  |  |  |  |  |  |  |  |
| --- | --- | --- | --- | --- | --- | --- | --- | --- | --- | --- | --- | --- | --- | --- | --- | --- | --- | --- | --- | --- | --- | --- | --- | --- | --- | --- | --- | --- | --- | --- | --- | --- | --- | --- | --- | --- | --- | --- | --- | --- | --- |
| pos | 686 | 687 | 688 | 689 | 690 | 691 | 692 | 693 | 694 | 1931 | 2954 | 6286 | 6335 | 8140 | 9893 | 11485 | 11750 | 14257 | 14599 | 16389 | 16935 | 17039 | 17721 | 22017 | 22893 | 22942 | 23535 | 26529 | 26601 | 26858 | 27259 | 27382 | 27383 | 27384 | 27788 | 27889 | 28312 | 28681 | 28724 | 29754 |  |
| ref | A | A | G | T | C | A | T | T | T | C | T | C | T | C | C | T | C | T | C | T | G | A | G | G | A | T | A | G | C | C | A | G | A | T | G | C | C | G | C | C |  |
| BA.4.6.1 | - | - | - | - | - | - | - | - | - | C | T | C | T | T | C | T | C | T | T | T | G | A | T | T | A | T | G | G | C | T | C | C | T | C | T | C | C | G | T | C |  |
| BA.5.2.1 | - | - | - | - | - | - | - | - | - | C | T | T | T | T | T | T | C | T | T | T | G | A | T | T | C | A | A | A | T | C | A | G | A | T | G | T | T | T | C | T | #1419248 |
| BQ.1.1 | A | A | G | T | C | A | T | T | T | A | C | C | A | C | C | C | T | C | C | C | A | G | G | G | C | A | A | A | T | C | A | G | A | T | G | T | T | T | C | T |  |
| mut |  |  |  |  |  |  |  |  |  |  |  | C>T |  |  | C>T |  |  |  |  |  |  |  |  |  |  |  |  |  |  |  |  |  |  |  |  |  |  |  |  |  |  |

|  |  |  |  |  |  |  |  |  |  |  |  |  |  |  |  |  |  |  |  |  |  |  |  |  |  |  |  |  |  |  |  |  |  |  |  |  |  |  |  |  |  |  |  |  |  |  |  |  |  |  |  |  |  |  |  |  |  |  |  |  |  |
| --- | --- | --- | --- | --- | --- | --- | --- | --- | --- | --- | --- | --- | --- | --- | --- | --- | --- | --- | --- | --- | --- | --- | --- | --- | --- | --- | --- | --- | --- | --- | --- | --- | --- | --- | --- | --- | --- | --- | --- | --- | --- | --- | --- | --- | --- | --- | --- | --- | --- | --- | --- | --- | --- | --- | --- | --- | --- | --- | --- | --- | --- |
| pos | 1093 | 1440 | 1931 | 1964 | 2954 | 3796 | 3927 | 4586 | 5183 | 5844 | 8038 | 8597 | 9866 | 11750 | 12160 | 12440 | 12444 | 14257 | 15451 | 16935 | 21077 | 21765 | 21766 | 21767 | 21768 | 21769 | 21770 | 21976 | 22001 | 22016 | 22033 | 22190 | 22331 | 22577 | 22599 | 22629 | 22855 | 22893 | 22898 | 22910 | 22917 | 22942 | 23018 | 23031 | 25416 | 25471 | 25553 | 25569 | 26031 | 26275 | 26529 | 26858 | 27259 | 27382 | 27383 | 27384 | 27889 | 28312 | 28681 | 29754 |  |
| ref | C | G | C | A | T | C | C | C | C | A | T | C | C | C | G | T | A | T | G | G | C | T | A | C | A | T | G | T | A | T | C | A | G | G | G | A | C | A | G | A | T | T | T | T | C | G | C | T | G | A | G | C | A | G | A | T | C | C | G | C |  |
| BN.3.1 | T | A | C | G | T | T | T | T | T | G | T | T | T | C | G | C | G | T | A | G | T | T | A | C | A | T | G | T | G | C | A | G | A | C | G | C | C | A | A | G | T | G | T | C | T | G | C | T | G | G | G | T | C | C | T | C | C | C | G | C |  |
| XBR | T | A | C | G | T | T | T | T | T | G | T | T | T | C | G | C | G | T | A | G | T | T | A | C | A | T | G | C | G | C | A | A | G | G | C | A | T | C | G | A | G | A | G | T | C | C | T | C | A | A | A | C | A | G | A | T | T | T | T | T | #1420166 |
| BQ.1.25 | C | G | A | A | C | C | C | C | C | A | C | C | C | T | A | T | A | C | G | A | C | - | - | - | - | - | - | T | A | T | C | A | G | G | C | A | T | C | G | A | G | A | G | T | C | G | T | C | A | A | A | C | A | G | A | T | T | T | T | T |  |
| mut |  |  |  |  |  |  |  |  |  |  |  |  |  |  |  |  |  |  |  |  |  |  |  |  |  |  |  | T>C |  |  |  |  |  |  |  |  |  |  |  |  |  |  |  |  |  | G>C |  |  |  |  |  |  |  |  |  |  |  |  |  |  |  |

|  |  |  |  |  |  |  |  |  |  |  |  |  |  |  |  |  |  |  |  |  |  |  |  |  |  |  |  |  |  |  |  |  |  |  |  |  |  |  |  |  |  |  |  |  |  |  |  |  |  |
| --- | --- | --- | --- | --- | --- | --- | --- | --- | --- | --- | --- | --- | --- | --- | --- | --- | --- | --- | --- | --- | --- | --- | --- | --- | --- | --- | --- | --- | --- | --- | --- | --- | --- | --- | --- | --- | --- | --- | --- | --- | --- | --- | --- | --- | --- | --- | --- | --- | --- |
| pos | 261 | 625 | 1627 | 3339 | 3796 | 3927 | 4586 | 5006 | 5183 | 6070 | 8692 | 9866 | 12160 | 12444 | 15451 | 18583 | 21765 | 21766 | 21767 | 21768 | 21769 | 21770 | 22001 | 22016 | 22033 | 22190 | 22331 | 22577 | 22599 | 22898 | 22917 | 22942 | 23018 | 23019 | 23031 | 25416 | 26275 | 26529 | 26813 | 26858 | 27038 | 27259 | 27382 | 27383 | 27384 | 27889 | 28330 | 29754 |  |
| ref | G | G | C | T | C | C | C | A | C | C | C | C | G | A | G | G | T | A | C | A | T | G | A | T | C | A | G | G | G | G | T | T | T | T | T | C | A | G | T | C | A | A | G | A | T | C | A | C |  |
| BA.5.2.1 | G | T | T | T | C | C | C | A | C | C | C | C | A | A | G | G | - | - | - | - | - | - | A | T | C | A | G | G | G | G | G | T | G | T | T | C | A | A | C | C | G | A | G | A | T | T | G | T |  |
| XBF | G | T | T | C | C | C | C | A | C | T | T | T | G | G | A | A | T | A | C | A | T | G | G | C | A | G | A | C | C | A | T | G | C | C | C | T | G | G | T | T | A | C | C | T | C | C | A | C | #1420385 |
| CJ.1 | A | G | C | T | T | T | T | G | T | C | C | T | G | G | A | A | T | A | C | A | T | G | G | C | A | G | A | C | C | A | T | G | C | C | C | T | G | G | T | T | A | C | C | T | C | C | A | C |  |
| mut |  |  |  | T>C |  |  |  |  |  | C>T | C>T |  |  |  |  |  |  |  |  |  |  |  |  |  |  |  |  |  |  |  |  |  |  |  |  |  |  |  |  |  |  |  |  |  |  |  |  |  |  |

|  |  |  |  |  |  |  |  |  |  |  |  |  |  |  |  |  |  |  |  |  |  |  |  |  |
| --- | --- | --- | --- | --- | --- | --- | --- | --- | --- | --- | --- | --- | --- | --- | --- | --- | --- | --- | --- | --- | --- | --- | --- | --- |
| pos | 882 | 1178 | 1931 | 2954 | 5549 | 6532 | 11750 | 11781 | 12534 | 14257 | 16935 | 19186 | 20332 | 22893 | 22942 | 25340 | 25855 | 27438 | 28312 | 28462 | 28681 | 28697 | 29666 |  |
| ref | C | C | C | T | C | G | C | A | C | T | G | C | G | A | T | G | G | T | C | C | G | C | C |  |
| BQ.1 | C | C | A | C | C | T | T | A | C | C | A | C | G | C | A | G | G | T | T | C | T | C | C |  |
| BQ.1 | C | C | A | C | C | T | T | A | C | C | A | T | G | C | A | T | G | C | C | T | G | T | C | #1421539 |
| BA.5.1.10 | T | T | C | T | A | G | C | G | T | T | G | C | A | A | T | G | C | C | C | T | G | T | T |  |
| mut |  |  |  |  |  |  |  |  |  |  |  | C>T |  |  |  | G>T |  |  |  |  |  |  | T>C |  |

|  |  |  |  |  |  |  |  |  |  |  |  |  |  |  |  |  |  |  |  |  |  |  |  |
| --- | --- | --- | --- | --- | --- | --- | --- | --- | --- | --- | --- | --- | --- | --- | --- | --- | --- | --- | --- | --- | --- | --- | --- |
| pos | 1216 | 1627 | 1931 | 2110 | 2954 | 8251 | 11750 | 12310 | 14257 | 15346 | 16935 | 17039 | 21991 | 21992 | 21993 | 22599 | 22893 | 22942 | 28312 | 28330 | 28376 | 28681 |  |
| ref | T | C | C | C | T | C | C | G | T | C | G | A | T | T | A | G | A | T | C | A | G | G |  |
| BA.5.2 | C | T | C | C | T | C | C | A | T | C | G | A | T | T | A | G | A | T | C | G | G | G |  |
| BA.5.2.1 | C | T | C | T | T | T | C | A | T | T | G | A | - | - | - | C | C | A | T | A | A | T | #1422524 |
| BQ.1.1 | T | C | A | C | C | C | T | G | C | C | A | G | T | T | A | C | C | A | T | A | A | T |  |
| mut |  |  |  | C>T |  | C>T |  |  |  | C>T |  |  | T>- | T>- | A>- | G>C |  |  |  |  |  |  |  |

|  |  |  |  |  |  |  |  |  |  |  |  |  |  |  |  |  |  |  |  |  |  |
| --- | --- | --- | --- | --- | --- | --- | --- | --- | --- | --- | --- | --- | --- | --- | --- | --- | --- | --- | --- | --- | --- |
| pos | 625 | 1627 | 1931 | 2954 | 6894 | 9118 | 11750 | 12310 | 14257 | 16616 | 16935 | 17039 | 22599 | 22893 | 22942 | 27012 | 27513 | 28312 | 28330 | 28681 |  |
| ref | G | C | C | T | G | C | C | G | T | C | G | A | G | A | T | C | C | C | A | G |  |
| BQ.1.1 | G | C | A | C | G | C | T | G | C | C | A | G | C | C | A | C | C | T | A | T |  |
| BQ.1.1 | A | C | A | C | G | T | T | G | C | C | A | G | C | C | A | T | T | C | G | G | #1423858 |
| BA.5.2 | G | T | C | T | T | C | C | A | T | A | G | A | G | A | T | T | T | C | G | G |  |
| mut | G>A |  |  |  |  | C>T |  |  |  |  |  |  |  |  |  |  |  |  |  |  |  |

|  |  |  |  |  |  |  |  |  |  |  |  |  |  |  |  |  |  |  |  |  |  |  |  |  |  |  |  |  |  |  |  |  |  |  |  |  |
| --- | --- | --- | --- | --- | --- | --- | --- | --- | --- | --- | --- | --- | --- | --- | --- | --- | --- | --- | --- | --- | --- | --- | --- | --- | --- | --- | --- | --- | --- | --- | --- | --- | --- | --- | --- | --- |
| pos | 920 | 3796 | 3857 | 3927 | 4586 | 5183 | 6196 | 7540 | 12444 | 13441 | 15451 | 18583 | 20741 | 21811 | 22001 | 22016 | 22033 | 22190 | 22331 | 22577 | 22599 | 22893 | 22898 | 22917 | 22942 | 23019 | 23040 | 25416 | 26275 | 26488 | 26491 | 26492 | 26497 | 28093 | 29358 |  |
| ref | C | C | C | C | C | C | C | T | A | G | G | G | A | C | A | T | C | A | G | G | G | A | G | T | T | T | A | C | A | T | T | A | T | C | C |  |
| CH.1.1.14 | C | T | A | T | T | T | C | T | G | A | A | A | G | T | G | C | A | G | A | C | C | C | A | G | G | C | A | T | G | T | T | A | T | T | T |  |
| CH.1.1.14 | T | T | A | T | T | T | C | C | G | A | A | A | G | T | G | C | A | G | A | C | C | C | A | G | G | C | A | T | G | G | C | T | C | T | T | #1425714 |
| BA.2 | C | C | C | C | C | C | T | T | A | G | G | G | A | C | A | T | C | A | G | G | G | A | G | T | T | T | G | C | A | G | C | T | C | C | C |  |
| mut | C>T |  |  |  |  |  |  | T>C |  |  |  |  |  |  |  |  |  |  |  |  |  |  |  |  |  |  |  |  |  |  |  |  |  | C>T | C>T |  |

|  |  |  |  |  |  |  |  |  |  |  |  |  |  |  |  |  |  |  |  |  |  |  |  |  |
| --- | --- | --- | --- | --- | --- | --- | --- | --- | --- | --- | --- | --- | --- | --- | --- | --- | --- | --- | --- | --- | --- | --- | --- | --- |
| pos | 1627 | 1931 | 2954 | 10335 | 11355 | 11750 | 12310 | 14257 | 16616 | 16935 | 19690 | 21991 | 21992 | 21993 | 22893 | 22942 | 23621 | 27012 | 27513 | 27600 | 28312 | 28330 | 28681 |  |
| ref | C | C | T | C | C | C | G | T | C | G | G | T | T | A | A | T | G | C | C | C | C | A | G |  |
| BU.1 | T | C | T | T | T | C | A | T | A | G | G | - | - | - | T | G | G | T | T | C | C | G | G |  |
| BA.5.2.1 | T | C | T | T | T | C | A | T | A | G | A | - | - | - | C | A | A | C | C | T | T | A | T | #1427576 |
| BQ.1.10 | C | A | C | C | C | T | G | C | C | A | G | T | T | A | C | A | G | C | C | T | T | A | T |  |
| mut |  |  |  |  |  |  |  |  |  |  | G>A |  |  |  |  |  | G>A |  |  |  |  |  |  |  |

|  |  |  |  |  |  |  |  |  |  |  |  |  |  |  |  |  |  |  |  |  |  |  |  |  |  |  |  |  |  |  |  |  |  |  |  |  |  |
| --- | --- | --- | --- | --- | --- | --- | --- | --- | --- | --- | --- | --- | --- | --- | --- | --- | --- | --- | --- | --- | --- | --- | --- | --- | --- | --- | --- | --- | --- | --- | --- | --- | --- | --- | --- | --- | --- |
| pos | 931 | 2788 | 3796 | 3857 | 3927 | 4586 | 5183 | 6548 | 6676 | 9300 | 12444 | 15451 | 16342 | 18583 | 20741 | 22001 | 22016 | 22033 | 22190 | 22331 | 22577 | 22599 | 22600 | 22893 | 22898 | 22917 | 22942 | 23019 | 23040 | 24304 | 26275 | 27576 | 29729 | 29757 | 29758 | 29759 |  |
| ref | T | C | C | C | C | C | C | C | T | G | A | G | T | G | A | A | T | C | A | G | G | G | A | A | G | T | T | T | A | T | A | T | T | G | T | G |  |
| CH.1.1.24 | C | C | T | A | T | T | T | C | C | G | G | A | T | A | G | G | C | A | G | A | C | C | A | C | A | G | G | C | A | T | G | C | T | G | T | G |  |
| CH.1.1.24 | C | T | T | A | T | T | T | C | C | G | G | A | C | A | G | G | C | A | G | A | C | C | A | C | A | G | G | C | A | C | G | C | C | A | C | C | #1427736 |
| BA.2 | T | C | C | C | C | C | C | T | T | A | A | G | T | G | A | A | T | C | A | G | G | G | T | A | G | T | T | T | G | T | A | T | C | A | C | C |  |
| mut |  | C>T |  |  |  |  |  |  |  |  |  |  | T>C |  |  |  |  |  |  |  |  |  |  |  |  |  |  |  |  | T>C |  |  |  |  |  |  |  |

|  |  |  |  |  |  |  |  |  |  |  |  |  |  |  |  |  |  |  |  |  |  |  |  |  |  |  |  |  |  |  |  |  |  |  |  |  |  |  |  |
| --- | --- | --- | --- | --- | --- | --- | --- | --- | --- | --- | --- | --- | --- | --- | --- | --- | --- | --- | --- | --- | --- | --- | --- | --- | --- | --- | --- | --- | --- | --- | --- | --- | --- | --- | --- | --- | --- | --- | --- |
| pos | 686 | 687 | 688 | 689 | 690 | 691 | 692 | 693 | 694 | 1627 | 1931 | 2653 | 2954 | 3838 | 5284 | 9136 | 10787 | 11080 | 11750 | 14257 | 16935 | 17039 | 21465 | 21991 | 21992 | 21993 | 22024 | 22893 | 22896 | 22942 | 26013 | 26179 | 27038 | 28312 | 28330 | 28370 | 28371 | 28681 |  |
| ref | A | A | G | T | C | A | T | T | T | C | C | A | T | G | C | G | G | T | C | T | G | A | T | T | T | A | A | A | T | T | C | A | A | C | A | A | G | G |  |
| BF.40 | A | A | G | T | C | A | T | T | T | T | C | A | T | A | T | A | G | T | C | T | G | A | T | T | T | A | A | A | C | T | T | A | G | C | G | T | T | G |  |
| BA.5.2.1 | - | - | - | - | - | - | - | - | - | T | C | A | T | A | T | A | G | C | C | C | A | G | C | - | - | - | G | C | T | A | C | C | A | T | A | - | G | T | #1428039 |
| BQ.1.23 | A | A | G | T | C | A | T | T | T | C | A | G | C | G | C | G | A | T | T | C | A | G | C | - | - | - | G | C | T | A | C | C | A | T | A | - | G | T |  |
| mut | A>- | A>- | G>- | T>- | C>- | A>- | T>- | T>- | T>- |  |  |  |  |  |  |  |  | T>C |  |  |  |  |  |  |  |  |  |  |  |  |  |  |  |  |  |  |  |  |  |

|  |  |  |  |  |  |  |  |  |  |  |  |  |  |  |  |  |  |  |  |  |  |  |  |  |  |  |  |  |  |  |  |  |  |  |  |  |  |  |  |  |  |  |  |  |  |  |  |  |  |  |  |  |  |  |  |  |  |  |  |  |  |  |  |
| --- | --- | --- | --- | --- | --- | --- | --- | --- | --- | --- | --- | --- | --- | --- | --- | --- | --- | --- | --- | --- | --- | --- | --- | --- | --- | --- | --- | --- | --- | --- | --- | --- | --- | --- | --- | --- | --- | --- | --- | --- | --- | --- | --- | --- | --- | --- | --- | --- | --- | --- | --- | --- | --- | --- | --- | --- | --- | --- | --- | --- | --- | --- | --- |
| pos | 44 | 378 | 405 | 1931 | 2954 | 6532 | 7063 | 7164 | 9866 | 10688 | 11750 | 12160 | 14120 | 14257 | 15451 | 15738 | 15939 | 16342 | 16935 | 17124 | 17859 | 19326 | 20428 | 21765 | 21766 | 21767 | 21768 | 21769 | 21770 | 21810 | 21991 | 21992 | 21993 | 22000 | 22109 | 22200 | 22317 | 22577 | 22599 | 22664 | 22893 | 22895 | 22896 | 22898 | 22917 | 22942 | 23018 | 23019 | 23031 | 25416 | 26275 | 26529 | 26858 | 27259 | 27382 | 27383 | 27384 | 27889 | 27915 | 28312 | 28681 | 29754 |  |
| ref | C | T | A | C | T | G | C | C | C | G | C | G | C | T | G | C | T | T | G | T | T | A | C | T | A | C | A | T | G | T | T | T | A | C | C | T | G | G | G | C | A | G | T | G | T | T | T | T | T | C | A | G | C | A | G | A | T | C | G | C | G | C |  |
| BQ.1 | T | T | A | A | C | T | C | C | C | G | T | A | C | C | G | C | T | T | A | T | T | A | C | - | - | - | - | - | - | T | T | T | A | C | C | G | G | G | G | C | C | G | T | G | G | A | G | T | T | C | A | A | C | A | G | A | T | T | G | T | T | T |  |
| XBB.1 | T | C | A | A | C | T | C | C | C | G | T | A | C | C | G | C | T | T | A | T | T | A | T | T | A | C | A | T | G | C | - | - | - | A | G | A | T | C | C | A | A | C | C | A | T | G | C | C | C | T | G | G | T | C | C | T | C | C | T | C | G | C | #1429711 |
| XBB.1.5.18 | C | T | G | C | T | G | T | T | T | A | C | G | T | T | A | T | C | C | G | C | C | G | T | T | A | C | A | T | G | C | - | - | - | A | G | A | T | C | C | A | A | C | C | A | T | G | C | C | C | T | G | G | T | C | C | T | C | C | T | C | G | C |  |
| mut |  | T>C |  |  |  |  |  |  |  |  |  |  |  |  |  |  |  |  |  |  |  |  |  |  |  |  |  |  |  |  |  |  |  |  |  |  |  |  |  |  |  |  |  |  |  |  |  |  |  |  |  |  |  |  |  |  |  |  |  |  |  |  |  |

|  |  |  |  |  |  |  |  |  |  |  |  |  |  |  |  |  |  |  |  |  |  |  |  |  |  |  |  |  |  |  |  |  |  |  |  |  |  |  |  |  |  |  |  |  |  |  |  |  |  |  |  |  |  |  |  |  |  |  |
| --- | --- | --- | --- | --- | --- | --- | --- | --- | --- | --- | --- | --- | --- | --- | --- | --- | --- | --- | --- | --- | --- | --- | --- | --- | --- | --- | --- | --- | --- | --- | --- | --- | --- | --- | --- | --- | --- | --- | --- | --- | --- | --- | --- | --- | --- | --- | --- | --- | --- | --- | --- | --- | --- | --- | --- | --- | --- | --- |
| pos | 44 | 405 | 686 | 687 | 688 | 689 | 690 | 691 | 692 | 693 | 694 | 2973 | 3796 | 3857 | 3927 | 4009 | 4586 | 5183 | 5802 | 7145 | 8182 | 12444 | 15738 | 15939 | 16207 | 16342 | 17859 | 18583 | 19326 | 20741 | 21005 | 21220 | 21810 | 21991 | 21992 | 21993 | 22000 | 22001 | 22016 | 22033 | 22109 | 22190 | 22200 | 22317 | 22331 | 22664 | 22893 | 22895 | 22896 | 22917 | 23015 | 23031 | 24210 | 25015 | 25463 | 27576 | 27915 |  |
| ref | C | A | A | A | G | T | C | A | T | T | T | C | C | C | C | C | C | C | A | G | A | A | C | T | G | T | T | G | A | A | C | G | T | T | T | A | C | A | T | C | C | A | T | G | G | C | A | G | T | T | G | T | C | G | C | T | G |  |
| XBB.1 | C | G | A | A | G | T | C | A | T | T | T | C | C | C | C | C | C | C | A | G | A | A | T | C | G | C | C | G | G | A | T | T | C | - | - | - | A | A | T | C | G | A | A | T | G | A | A | C | C | T | G | C | C | T | T | T | T |  |
| BA.2 | C | G | - | - | - | - | - | - | - | - | - | T | C | C | C | C | C | C | A | G | A | A | T | C | T | C | C | G | G | G | C | G | T | T | T | A | C | G | C | A | C | G | G | G | A | C | C | G | T | G | C | T | T | G | C | C | G | #1430261 |
| CH.1.1.2 | T | A | A | A | G | T | C | A | T | T | T | C | T | A | T | T | T | T | G | A | G | G | C | T | G | T | T | A | A | G | C | G | T | T | T | A | C | G | C | A | C | G | G | G | A | C | C | G | T | G | C | T | T | G | C | C | G |  |
| mut |  |  | A>- | A>- | G>- | T>- | C>- | A>- | T>- | T>- | T>- | C>T |  |  |  |  |  |  |  |  |  |  |  |  | G>T |  |  |  |  |  |  |  |  |  |  |  |  |  |  |  |  |  |  |  |  |  |  |  |  |  |  |  |  |  |  |  |  |  |

|  |  |  |  |  |  |  |  |  |  |  |  |  |  |  |  |  |  |  |  |  |  |  |  |  |  |  |  |  |  |  |  |  |  |  |  |  |  |  |  |  |  |  |  |  |  |  |  |  |  |  |  |  |  |  |
| --- | --- | --- | --- | --- | --- | --- | --- | --- | --- | --- | --- | --- | --- | --- | --- | --- | --- | --- | --- | --- | --- | --- | --- | --- | --- | --- | --- | --- | --- | --- | --- | --- | --- | --- | --- | --- | --- | --- | --- | --- | --- | --- | --- | --- | --- | --- | --- | --- | --- | --- | --- | --- | --- | --- |
| pos | 1063 | 1931 | 2700 | 2954 | 3796 | 3857 | 3927 | 4586 | 5183 | 5802 | 7145 | 8293 | 9866 | 10323 | 11750 | 12160 | 12444 | 14257 | 15451 | 16935 | 18583 | 20741 | 21765 | 21766 | 21767 | 21768 | 21769 | 21770 | 22001 | 22016 | 22033 | 22190 | 22331 | 22577 | 22599 | 22898 | 22942 | 23018 | 23019 | 24210 | 25416 | 26275 | 26529 | 26858 | 27259 | 27382 | 27383 | 27384 | 27576 | 27889 | 28312 | 28681 | 29754 |  |
| ref | C | C | C | T | C | C | C | C | C | A | G | C | C | A | C | G | A | T | G | G | G | A | T | A | C | A | T | G | A | T | C | A | G | G | G | G | T | T | T | C | C | A | G | C | A | G | A | T | T | C | C | G | C |  |
| BQ.1 | T | A | C | C | C | C | C | C | C | A | G | T | C | A | T | A | A | C | G | A | G | A | - | - | - | - | - | - | A | T | C | A | G | G | G | G | A | G | T | C | C | A | A | C | A | G | A | T | T | T | T | T | T |  |
| BA.2 | T | A | T | C | C | C | C | C | C | A | G | T | C | G | T | G | G | T | A | G | A | G | T | A | C | A | T | G | G | C | A | G | A | C | C | A | G | T | C | T | T | G | G | T | C | C | T | C | C | C | C | G | C | #1430452 |
| CH.1.1.2 | C | C | C | T | T | A | T | T | T | G | A | C | T | A | C | G | G | T | A | G | A | G | T | A | C | A | T | G | G | C | A | G | A | C | C | A | G | T | C | T | T | G | G | T | C | C | T | C | C | C | C | G | C |  |
| mut |  |  | C>T |  |  |  |  |  |  |  |  |  |  | A>G |  |  |  |  |  |  |  |  |  |  |  |  |  |  |  |  |  |  |  |  |  |  |  |  |  |  |  |  |  |  |  |  |  |  |  |  |  |  |  |  |

|  |  |  |  |  |  |  |  |  |  |  |  |  |  |
| --- | --- | --- | --- | --- | --- | --- | --- | --- | --- | --- | --- | --- | --- |
| pos | 510 | 1545 | 4402 | 5812 | 6198 | 10765 | 19886 | 21811 | 22115 | 24991 | 25721 | 25936 |  |
| ref | G | C | T | C | C | C | A | C | A | T | C | C |  |
| CH.1.1.1 | G | T | A | C | T | T | G | C | G | A | T | C |  |
| CH.1.1 | A | T | A | T | T | T | G | T | A | T | C | T | #1431988 |
| CH.1.1 | G | C | T | C | C | C | A | T | A | T | C | C |  |
| mut | G>A |  |  | C>T |  |  |  |  |  |  |  | C>T |  |

|  |  |  |  |  |  |  |  |  |  |  |  |  |  |  |  |  |  |  |  |  |  |  |  |  |  |  |  |  |  |  |  |  |  |  |  |  |  |  |  |  |  |  |  |  |  |  |  |  |  |  |  |  |  |  |  |  |  |  |  |  |  |  |  |  |  |  |  |  |
| --- | --- | --- | --- | --- | --- | --- | --- | --- | --- | --- | --- | --- | --- | --- | --- | --- | --- | --- | --- | --- | --- | --- | --- | --- | --- | --- | --- | --- | --- | --- | --- | --- | --- | --- | --- | --- | --- | --- | --- | --- | --- | --- | --- | --- | --- | --- | --- | --- | --- | --- | --- | --- | --- | --- | --- | --- | --- | --- | --- | --- | --- | --- | --- | --- | --- | --- | --- | --- |
| pos | 686 | 687 | 688 | 689 | 690 | 691 | 692 | 693 | 694 | 1545 | 1931 | 2954 | 3796 | 3857 | 3927 | 4276 | 4402 | 4586 | 5183 | 6388 | 7850 | 9866 | 11374 | 11750 | 12160 | 12444 | 14257 | 15451 | 16935 | 17039 | 18583 | 19886 | 20741 | 21765 | 21766 | 21767 | 21768 | 21769 | 21770 | 22001 | 22016 | 22033 | 22115 | 22190 | 22331 | 22577 | 22898 | 22942 | 23018 | 23019 | 23401 | 23587 | 24991 | 25416 | 25721 | 26275 | 26529 | 26858 | 27259 | 27382 | 27383 | 27384 | 27889 | 28312 | 28681 | 28770 | 29754 |  |
| ref | A | A | G | T | C | A | T | T | T | C | C | T | C | C | C | C | T | C | C | C | G | C | T | C | G | A | T | G | G | A | G | A | A | T | A | C | A | T | G | A | T | C | A | A | G | G | G | T | T | T | G | G | T | C | C | A | G | C | A | G | A | T | C | C | G | C | C |  |
| BQ.1.1.29 | - | - | - | - | - | - | - | - | - | C | A | C | C | C | C | C | T | C | C | T | A | C | C | T | A | A | C | G | A | G | G | A | A | - | - | - | - | - | - | A | T | C | A | A | G | G | G | A | G | T | T | G | T | C | C | A | A | C | A | G | A | T | T | T | T | C | T |  |
| BA.2 | A | A | G | T | C | A | T | T | T | C | A | C | C | C | C | T | T | C | C | T | A | C | C | T | A | A | C | G | A | G | G | A | A | T | A | C | A | T | G | G | C | A | G | G | A | C | A | G | T | C | G | C | A | T | T | G | G | T | C | C | T | C | C | C | G | T | C | #1432902 |
| CH.1.1.1 | A | A | G | T | C | A | T | T | T | T | C | T | T | A | T | C | A | T | T | C | G | T | T | C | G | G | T | A | G | A | A | G | G | T | A | C | A | T | G | G | C | A | G | G | A | C | A | G | T | C | G | G | A | T | T | G | G | T | C | C | T | C | C | C | G | C | C |  |
| mut | ->A | ->A | ->G | ->T | ->C | ->A | ->T | ->T | ->T |  |  |  |  |  |  | C>T |  |  |  |  |  |  |  |  |  |  |  |  |  |  |  |  |  |  |  |  |  |  |  |  |  |  |  |  |  |  |  |  |  |  |  | G>C |  |  |  |  |  |  |  |  |  |  |  |  |  | C>T |  |  |

|  |  |  |  |  |  |  |  |  |  |  |  |  |  |  |  |  |  |  |  |  |  |  |  |  |  |  |  |  |  |  |  |  |  |  |  |  |  |  |  |
| --- | --- | --- | --- | --- | --- | --- | --- | --- | --- | --- | --- | --- | --- | --- | --- | --- | --- | --- | --- | --- | --- | --- | --- | --- | --- | --- | --- | --- | --- | --- | --- | --- | --- | --- | --- | --- | --- | --- | --- |
| pos | 44 | 405 | 6548 | 7732 | 9300 | 15451 | 15738 | 15939 | 16342 | 17124 | 17859 | 19326 | 21810 | 21991 | 21992 | 21993 | 22000 | 22109 | 22200 | 22317 | 22577 | 22599 | 22600 | 22664 | 22895 | 22896 | 22898 | 22942 | 23019 | 23031 | 23040 | 24210 | 26275 | 27915 | 29729 | 29757 | 29758 | 29759 |  |
| ref | C | A | C | C | G | G | C | T | T | T | T | A | T | T | T | A | C | C | T | G | G | G | A | C | G | T | G | T | T | T | A | C | A | G | T | G | T | G |  |
| XBB.1.4 | C | G | C | C | G | A | T | C | C | T | C | G | C | - | - | - | A | G | A | T | C | C | A | A | C | C | A | G | C | C | A | T | G | T | T | G | T | G |  |
| XBB.1.5.4 | C | G | C | T | G | A | T | C | C | C | C | G | C | - | - | - | A | G | A | T | C | C | A | A | C | C | A | G | C | C | A | T | G | T | C | A | C | C | #1436032 |
| BA.2 | T | A | T | C | A | G | C | T | T | T | T | A | T | T | T | A | C | C | G | G | G | G | T | C | G | T | G | T | T | T | G | C | A | G | C | A | C | C |  |
| mut |  |  |  | C>T |  |  |  |  |  | T>C |  |  |  |  |  |  |  |  |  |  |  |  |  |  |  |  |  |  |  |  |  |  |  |  |  |  |  |  |  |
